# What explains the effect of education on cardiovascular disease? Applying Mendelian randomization to identify the consequences of education inequality

**DOI:** 10.1101/488254

**Authors:** Alice R Carter, Dipender Gill, Neil M Davies, Amy E Taylor, Taavi Tillmann, Julien Vaucher, Robyn E Wootton, Marcus R Munafò, Gibran Hemani, Rainer Malik, Sudha Seshadri, Daniel Woo, Stephen Burgess, George Davey Smith, Michael V Holmes, Ioanna Tzoulaki, Laura D Howe, Abbas Dehghan

## Abstract

**Question:** What is the role of body mass index, systolic blood pressure and smoking in mediating the effect of education on cardiovascular disease risk?

**Finding:** We find consistent evidence that body mass index, systolic blood pressure and smoking mediate the effect of education, explaining up to 18%, 27% and 33% respectively. Including all three risk factors in a model together explains around 40% of the effect of education.

**Meaning:** Intervening on body mass index, systolic blood pressure and smoking would lead to reductions in cases of CVD attributable to lower levels of education. Over half of the effect of education on risk of cardiovascular disease is not mediated through these risk factors.

**Importance:** Lower levels of education are causally related to higher cardiovascular risk, but the extent to which this is driven by modifiable risk factors also associated with education is unknown.

**Objective:** To investigate the role of body mass index, systolic blood pressure and smoking in explaining the effect of education on risk of cardiovascular disease outcomes.

**Design:** Multivariable regression analysis of observational data and Mendelian randomization (MR) analysis of genetic data.

**Setting:** UK Biobank and international genome-wide association study consortia.

**Participants:** Predominantly individuals of European ancestry.

**Main outcomes and measures:** The effects of education (per 1-standard deviation increase, equivalent to 3.6 years) on coronary heart disease, cardiovascular disease (all subtypes), myocardial infarction and stroke risk (all measured in odds ratio, OR), and the degree to which this is mediated through body mass index, systolic blood pressure and smoking.

**Results:** Each additional standard deviation of education associated with 13% lower risk of coronary heart disease (OR 0.87, 95% confidence interval [CI] 0.84 to 0.89) in observational analysis and 37% lower risk (OR 0.63, 95% CI 0.60 to 0.67) in Mendelian randomization analysis. As a proportion of the total risk reduction, body mass index mediated 15% (95% CI 13% to 17%) and 18% (95% CI 14% to 23%) in the observational and Mendelian randomization estimates, respectively. Corresponding estimates for systolic blood pressure were 11% (95% CI 9% to 13%) and 21% (95% CI 15% to 27%), and for smoking, 19% (15% to 22%) and 33% (95% CI 17% to 49%). All three risk factors combined mediated 42% (95% CI 36% to 48%) and 36% (95 % CI 16% to 63%) of the effect of education on coronary heart disease in observational and Mendelian randomization respectively. Similar results were obtained when investigating risk of stroke, myocardial infarction and all-cause cardiovascular disease.

**Conclusions and relevance:** BMI, SBP and smoking mediate a substantial proportion of the protective effect of education on risk of cardiovascular outcomes and intervening on these would lead to reductions in cases of CVD attributable to lower levels of education. However, more than half of the protective effect of education remains unexplained and requires further investigation.

## Introduction

Cardiovascular disease (CVD) is the leading cause of mortality worldwide, accounting for over 17 million deaths annually (1). Insights into aetiological mechanisms have improved prevention through the modification of risk factors (1, 2). Recent studies have suggested that socioeconomic risk factors such as education play a causal role in the aetiology of CVD (3-5). Education may prevent CVD, in part through its effects on modifiable risk factors (6-8), and understanding this relationship can be used to inform public health policy.

Existing studies suggest that body mass index (BMI), systolic blood pressure (SBP) and smoking behaviour at least partly explain differences in CVD risk related to educational attainment (6-8). However, these studies have relied on observational mediation analyses that may suffer from biases. Traditional methods use a single snapshot of any given risk factor, which may thus incompletely capture a person’s lifetime exposure (9). For example, blood pressure measured at a single timepoint will suffer from measurement error due to day-to-day fluctuations and will not capture changes across the life course. This measurement error can lead to an underestimation of mediation (9). Furthermore, other biases such as unmeasured confounding cannot be addressed using observational methodologies (10).

Mendelian randomization (MR) uses genetic variants as instruments to estimate the effect of an exposure on an outcome of interest (11), exploiting the random allocation of genetic variants to infer causal effects that are robust to non-differential measurement error and confounding from environmental factors (11). Two-step Mendelian randomization was introduced for mediation analysis; unlike traditional observational mediation analysis approaches, it is both sensitive to the causal effects of the mediator and corrects for measurement error in the mediator (12). Recent genome-wide association study (GWAS) meta-analyses have identified a number of genetic variants for educational attainment that may be used as instrumental variables (13, 14). In this study, we investigated the role of BMI, SBP and lifetime smoking in mediating the causal effect of educational attainment on CVD risk using three complementary approaches: multivariable regression of observational data, one-sample MR and two-sample MR. BMI, SBP and smoking were selected as intermediate risk factors based on previous literature implicating them as both being affected by education and as risk factors for CVD, with suitable data available across all three complementary methods used.

## Methods

### Overall study design

This study used complementary approaches to investigate whether lower BMI, SBP and lifetime smoking explain the protective effect of education on risk of cardiovascular disease. Multivariable regression of observational data, one-sample MR of individual level genetic data and two-sample MR of summary level genetic data investigating coronary heart disease (CHD) myocardial infarction (MI), stroke risk and CVD (all subtypes) were carried out.

### Data sources

#### UK Biobank

The UK Biobank recruited 503,317 UK adults between 2006 and 2010. Participants attended baseline assessment centres involving questionnaires, interviews, anthropometric, physical and genetic measurements (15, 16). In the observational analysis, we included 217,013 White British individuals, with complete data on genotypes, age, sex, educational attainment, cardiovascular outcomes, BMI, smoking status, blood pressure, socioeconomic status (as measured by Townsend Deprivation Index at birth [TDI]) and place of birth. Exclusion criteria and flowchart are available in the Supplementary Methods (Supplementary Figures 1 and 2). Individuals of White British descent were defined using both self-reported questionnaire data with similar genetic ancestry based on the genetic principal components (PC) (17).

Data from the baseline assessment centre on highest qualifications completed and smoking, clinic measurements of BMI and SBP, and all covariate measures were used for the analyses (see Supplementary Methods). A continuous lifetime measure of smoking, incorporating smoking initiation, duration, heaviness and cessation is used in this analysis. Details of the derivation of this and applications have been detailed elsewhere (18). Participants reported their highest qualification and age of leaving school if they did not have a degree at the baseline assessment centre. These were converted to the International Standard Classification for Education (ISCED) coding of educational attainment (Supplementary Table 1) (14). Available follow-up data were used where baseline data were missing. To account for the effects of anti-hypertensive treatment, participants who were taking antihypertensive medication had 10mmHg added to their measured SBP (19). CVD diagnoses (including diagnoses of stroke, MI and CHD) and events were ascertained through linkage mortality data and hospital episode statistics (HES), with cases defined according to ICD-9 and ICD-10 codes (Supplementary Table 2) (20). Individuals who had experienced a CVD event prior to the baseline assessment (prevalent cases) were excluded and only first event, incident cases following the assessment centre were considered. Date of diagnoses are provided by HES data, which was linked with the date of assessment centre provided by UK Biobank.

#### GWAS meta-analyses

In the two-sample MR analysis, we obtained summary genetic associations from GWAS data for each respective phenotype. For education, this was the Social Science Genetic Association Consortium (SSGAC) GWAS meta-analysis of years of schooling in 1,131,881 individuals of European ancestry (14), with summary data made available for 766,345 of these participants. Instruments were selected as the 1,271 independent (pairwise r^2^<0.1) genome-wide significant SNPs after analysing data from the full sample (14). Education was assessed at or above the age of 30 years, with comparability between studies heterogeneous in their educational systems maximised by mapping major educational qualifications on to one of seven categories of the ISCED (13, 14). We obtained genetic estimates for BMI from the Genetic Investigation of ANthropometric Traits (GIANT) consortium’s 2018 GWAS meta-analysis of 681,275 individuals of European decent (21). Genetic association estimates for SBP and smoking were estimated from a GWAS of 318,417 White British individuals in UK Biobank, protocol details for which are available in the Supplementary Methods. Instruments for BMI, SBP and smoking were identified as the lead SNPs in loci reaching genome-wide significance after clumping summary estimates from the largest available GWAS for linkage disequilibrium (LD) r^2^<0.001 using a 1000 genomes European reference panel through the TwoSampleMR package of the statistical software R (22). For risk of CHD, we used publicly available genetic association estimates from the CARDIoGRAMplusC4D 1000 Genomes-based GWAS metaanalysis of 60,801 cases and 123,504 controls (23). Participants were of European, east Asian, south Asian, Hispanic and African American ancestry, and adjustment was made for population stratification using the genomic control method (23). The definition for CHD was broad and inclusive, considering acute coronary syndrome, myocardial infarction, angina with one or angiographic stenoses of greater than 50%, and chronic stable angina. Details of the studies used to obtain genetic association estimates for MI and stroke are provided in the Supplementary Methods. All genetic association estimates used in each two-sample MR analyses are provided in Supplementary Tables 7-21.

### Statistical Analysis

#### Effect of education on cardiovascular disease

In the observational analysis of UK Biobank data, multivariable logistic regression was used to estimate the association of education with CVD and its subtypes. Analyses were adjusted for potential confounders; age, sex, place of birth, birth distance from London, and TDI. Full details of derived variables (birth distance from London and TDI at birth) are available in the supplementary methods.

In the two-sample MR analysis, the effects of education on CVD subtypes were investigated using ratio method MR with standard errors derived using the delta method (24). Fixed-effect inverse-variance weighted (IVW) meta-analysis was used to pool MR estimates across individual SNPs and calculate an overall estimate (25).

#### Mediation by body mass index, systolic blood pressure and smoking

When investigating the degree to which the effects of education on CVD and its subtypes are mediated through each risk factor (BMI, SBP and smoking) individually, the product of coefficients method was used to estimate the indirect effect (26). This involved first estimating the effect of education on each risk factor individually, then multiplying this with the effect of that risk factor on the outcome after adjusting for education, to estimate the mediated (indirect) effect (i.e. the effect of education on CVD that goes through the risk factor). We estimated the proportion of the overall effect of education on CVD subtypes that was mediated by each risk factor by dividing the indirect effect by the total effect. Standard errors were derived by bootstrapping in the observational and one-sample MR analysis and using the delta method in the two-sample MR analysis.

When investigating the role of all three risk factors together on the association between education and CVD, we used the difference method (27). This involved estimating the total effect of education on each CVD subtype, as described above. We estimated the direct effect of education on each CVD subtype controlling for all three risk factors together, using either multivariable regression or multivariable MR, in observational and MR analyses respectively. The direct effect was then taken away from the total effect to find the indirect effect, which was divided by the total effect to give an estimate of the amount of mediation.

In the observational analysis, multivariable linear regression was used to estimate the association of education with each risk factor after adjusting for confounders (as in the total effects models). The effect of each risk factor on the individual CVD subtypes was then estimated with additional adjustment for self-reported educational attainment (27). The two estimates were multiplied together to estimate the indirect effect (of education, through the risk factor). Considering all three risk factors together, a multivariable logistic model for the effect of education on CVD (and subtypes) adjusting for all three risk factors was used to estimate the direct effect of education independently of the risk factors. This was taken away from the total effect to estimate the indirect effect of education through the three risk factors collectively.

For the two-sample MR, the IVW MR approach was used to estimate the effect of education on each risk factor and regression-based multivariable MR was used to estimate the effect of each risk factor on risk of the considered CVD subtypes, adjusting for genetic effect of the instruments on education (28). The indirect effect of education on risk of each CVD subtype through the considered risk factor was estimated by multiplying results from these two MR analyses. To estimate the total effect of education mediated through all three risk factors collectively, the direct effect of education after adjusting for the three risk factors together was estimated using MVMR and taken away from the total effect of education previously estimated using IVW MR; this indirect effect was divided by the total effect to estimate the proportion mediated.

#### One-sample MR and sensitivity analyses

One sample MR analysis was carried out in UK Biobank. Analyses using UK Biobank were replicated using the risk difference scale. A range of sensitivity analyses were carried out, including to explore the assumption of no pleiotropy within MR analyses. Full details are provided in the Supplementary Methods.

### Statistical software and ethical approval

Analysis was performed using Stata version 14 (StataCorp LP) and R version 3.4.3 (The R Foundation for Statistical Computing). The mrrobust package for Stata and the TwoSample MR package for R were used to facilitate MR analyses (22, 29). The code used for all analyses is available on request and details of genetic variants used are provided in the Supplementary Tables. Ethical approval was not sought for publicly available data because all participating studies had already obtained relevant authorisation. Project approval was obtained from UK Biobank (study ID: 10953).

## Results

### UK Biobank Cohort Description

The UK Biobank sample used in the observational and one-sample MR analysis was comparable to the participants in UK Biobank as a whole. In the analysis sample 38% of individuals had over 19 years of education, equivalent to a vocational qualification or degree. Comparatively, only 17% of individuals left school with no formal qualifications after seven years. Full details are available in Supplementary Table 3. The standard deviations (SD) of educational attainment was 3.6 years, BMI was 4.69 kg/m2 and SBP was 18.68 mm Hg. For lifetime smoking, 1-SD increase is equivalent to, for example, an individual smoking 20 cigarettes a day for 15 years and stopping 17 years ago, or an individual smoking 60 cigarettes a day for 13 years and stopping 22 years ago. For further explanation of this, see Wootton and colleagues (18).

### Effect of education on risk of CHD and stroke

In observational analyses, 1-SD higher education was associated with a 13% lower risk of CHD with an odds ratio (OR) of 0.87 (95% CI 0.84 to 0.89). The two-sample MR analysis indicated a stronger protective effect, with OR 0.63 (95% CI 0.60-0.67) (Figure 1).

**Figure 1:**
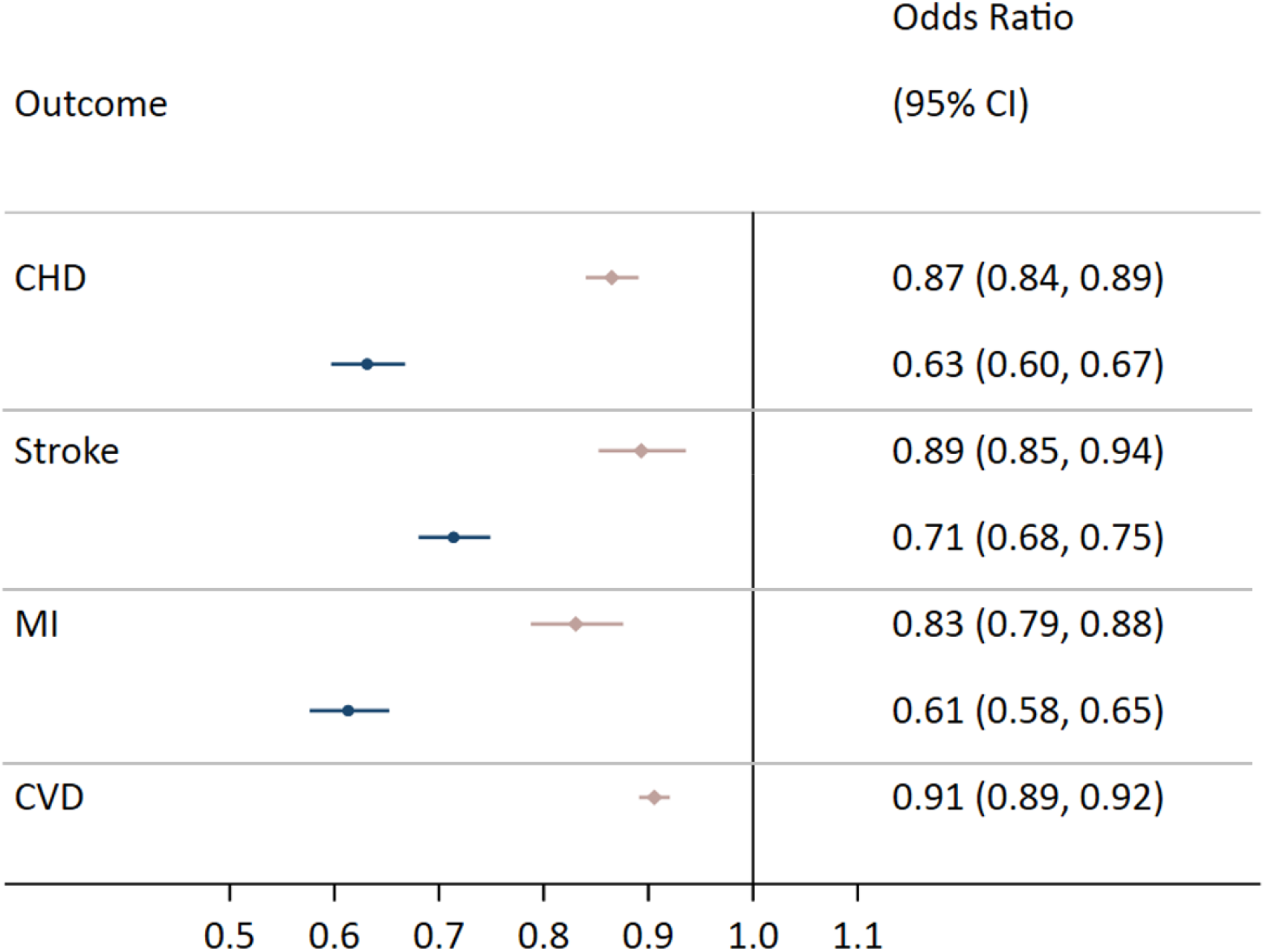
The effect of a 1-SD increase in education on the risk of cardiovascular disease and its subtypes. Observational multivariable estimates are plotted in pink and two-sample MR estimates plotted in navy. Multivariable analyses adjusted for: age, sex, place of birth and Townsend deprivation index at birth. BMI, SBP and smoking were measured in 1-SD units. CVD (All subtypes) was not available for analysis in two-sample MR analysis.

Similar protective associations were found for the effect of education on other CVD subtypes (Figure 1). In observational analyses, a 1-SD higher education was associated with an 11% lower risk of stroke, with an OR of 0.89 (95% CI 0.85, 0.94). In two-sample MR analyses the protective effect was stronger with OR 0.71 (0.68, 0.75).

One sample-MR analyses also provided consistent evidence for a protective effect of a 1-SD increase in education with CVD risk and its subtypes (Supplementary Figure 3).

### Effect of education on BMI, SBP and smoking

In all methods, a longer time in education was associated with lower ORs of BMI, SBP and smoking (Figure 2 and Supplementary Figure 4). Both multivariable observational analyses and two-sample MR found the largest association of education with smoking, where each additional 1-SD of education associated with 0.13 SD-units (95% CI 0.13 to 0.13) lower lifetime smoking in observational multivariable analyses, and 0.32 (95% CI 0.31 to 0.33) in two-sample MR. For each additional 1-SD of education, BMI was lower by 0.10 SD-units (95% CI 0.10 to 0.09) in observational analyses and 0.22 SD-units (95% CI 0.20 to 0.24) in two-sample MR. The smallest association was seen for SBP, where 1-SD higher education was associated with 0.06 SD-units (95% CI 0.06 to 0.07) lower SBP in multivariable observational analyses, and 0.15 SD-units (95% CI 0.14 to 0.16) in two-sample MR analyses (Figure 2).

**Figure 2:**
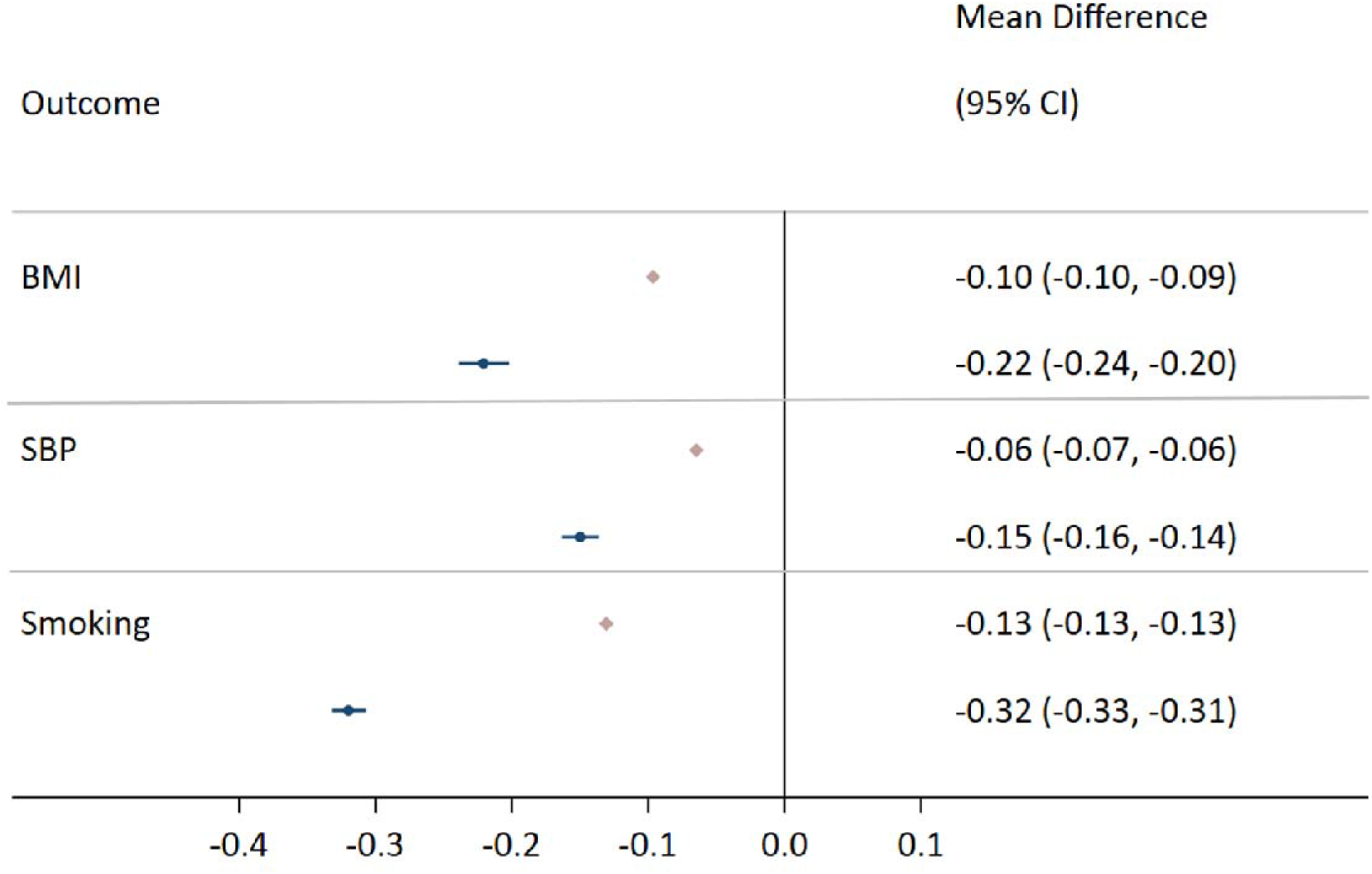
Observational and two-sample MR estimates for the association between 1-SD higher education and BMI, SBP and lifetime smoking respectively. All outcomes are in 1-SD units. Observational multivariable results are plotted in pink, with two sample MR estimates plotted in navy.

### Effect of BMI, SBP and smoking on risk of CVD subtypes

Both observational and two-sample MR analyses consistently found evidence to support an increased risk of CHD with higher BMI, SBP and smoking (Figure 3). The strongest risk factor for CHD was SBP, where a 1-SD increase in SBP increased the risk of CHD with an OR of 1.28 (95% CI 1.25 to 1.32) in multivariable observational analyses and 1.90 (95% CI 1.64 to 2.19) in two-sample MR analyses. A 1-SD higher BMI was associated with an OR of 1.26 (95% CI 1.22 to 1.30) for CHD in observational analyses and 1.47 (95% CI 1.35 to 1.59) in two-sample MR. A 1-SD increase in lifetime smoking was associated with a CHD OR of 1.23 (95% CI 1.20 to 1.26) in multivariable observational analyses and 1.61 (95% CI 1.28 to 2.02) in two-sample MR analyses.

**Figure 3:**
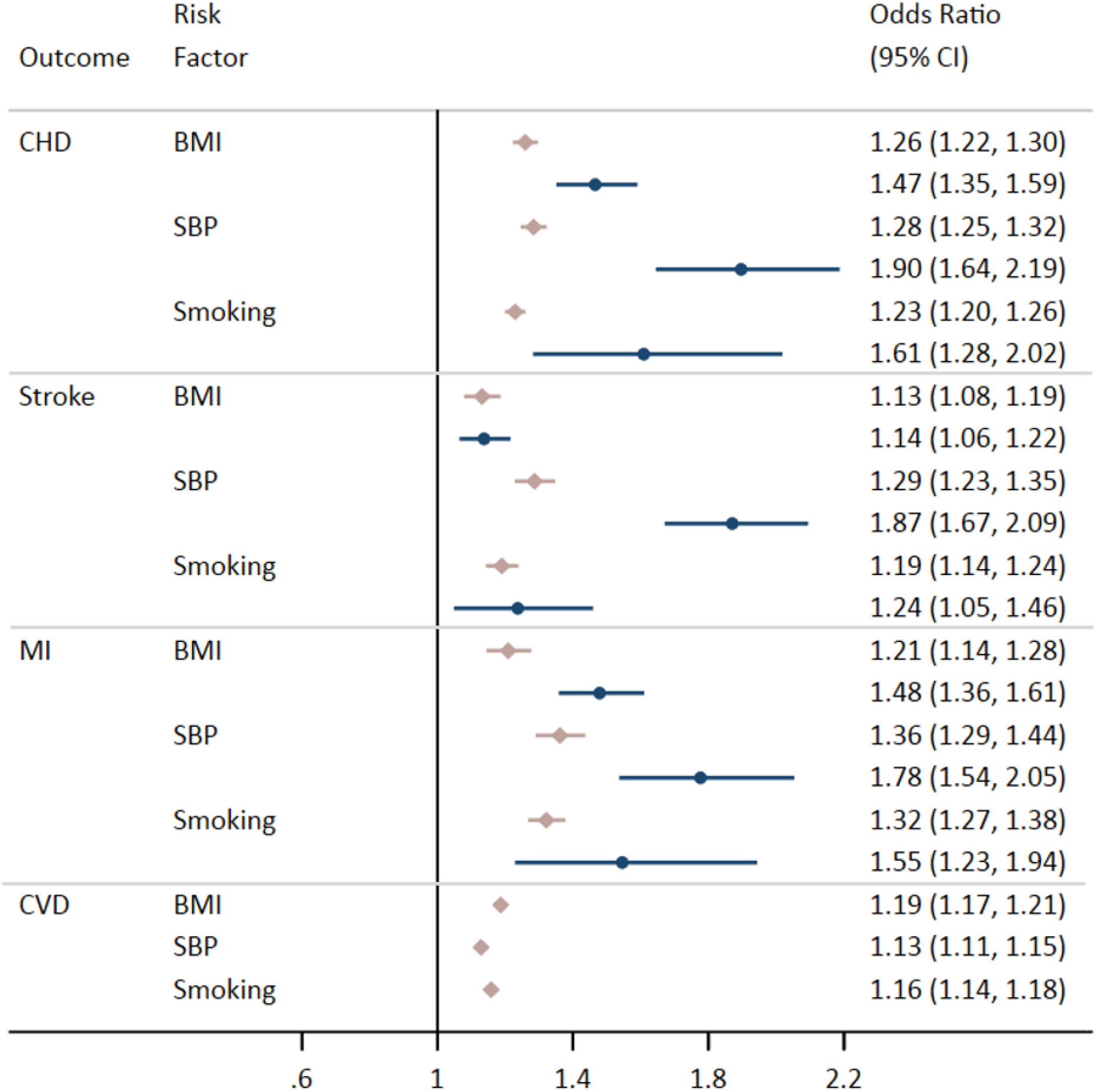
Observational and two-sample MR associations of 1-SD higher BMI, SBP and lifetime smoking on the risk of cardiovascular disease and its subtypes. Observational multivariable results are plotted in pink, with two sample MR estimates plotted in navy.

For all other outcomes, multivariable observational and two-sample MR analyses consistently found evidence to support an increased risk of CVD, stroke and MI with higher BMI, SBP and smoking (Supplementary Figure 6). The strongest risk factor for stroke was also SBP, with a 1-SD increase associated with OR 1.36 (95% CI 1.29, 1.44) in multivariable observational analyses and 1.87 (95% CI 1.67, 2.09) in two-sample MR analyses.

As expected, the effect of each risk factor was less consistent in the one-sample MR and estimates had wide confidence intervals (Supplementary Figure 5).

### Mediation by BMI, SBP and smoking

In the observational analysis, the proportion of the effect of education on CHD risk mediated by BMI was 15% (95% CI 13% to 17%), 11% for SBP (95% CI 9% to 13%) and 19% for smoking (95% CI 15% to 22%) (Figure 4). In the two-sample MR analysis, the percentage mediated by BMI was 18% (95% CI 14% to 23%), 21% by SBP (95% CI 15% to 27%) and 33% by smoking (95% CI 17% to 49%) (Figure 4, Supplementary Table 22). In observational analyses, combining all three risk factors together explained 42% (95% CI 36% to 48%) of the effect of education on risk of CHD (Figure 4). In two-sample MR the combined effect of all three risk factors on CHD as 36% (95% CI 5% to 68%).

**Figure 4:**
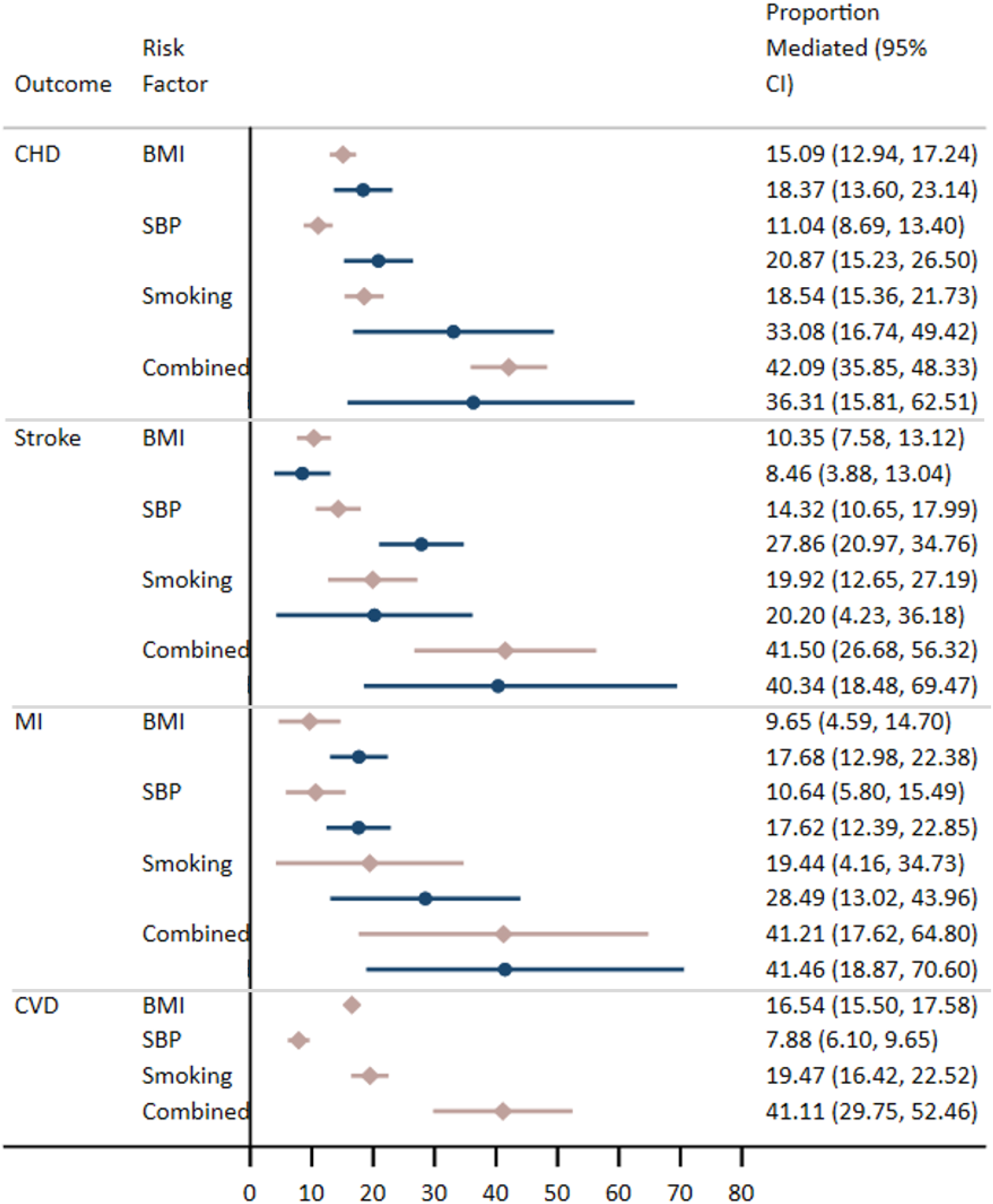
Estimates for the effect of education on CVD and its subtypes explained by BMI, SBP and smoking respectively. Results are provided for the multivariable observational analysis (plotted in pink) and two-sample MR (plotted in navy).

Similar results were found for other CVD subtypes in multivariable observational analyses. Smoking consistently mediated around 20% of the association. BMI explained between 10% and 17% of the association between education and CVD and its subtypes, whilst SBP explained between 8% and 18%. In two-sample MR analyses, smoking explained up to 28% of the association between education and CVD subtypes, whilst BMI estimated up to 18% and SBP up to 28% of the association.

One-sample MR analyses estimated similar amounts of the association explained by SBP and smoking but was less consistent for BMI (Supplementary figure 7).

### Sensitivity analyses

Results from sensitivity analyses were comparable but produced less precise estimates with wider confidence intervals (Supplementary Figure 2-8, Supplementary Tables 22 and 23). Unadjusted and age and sex adjusted models were also consistent with the main fully adjusted models for multivariable analyses (Supplementary Tables 24 and 25). The effects of each mediator individually, and combined, estimated on the risk difference scale and using the difference method in individual data were consistent with main analyses on the log OR scale (Supplementary Figure 8).

## Discussion

Our observational and genetic analyses provide complementary evidence that the effect of education on risk of CVD is mediated by up to one third of the effect through any of BMI, SBP or smoking. When investigating all three risk factors together, it was estimated that around 40% of the association between education and CVD is explained by the three risk factors, both in observational and MR analyses. However, it is important that over half of the effects of education remain unexplained in these analyses. This portion of the pathway may be due to other lifestyle behaviours associated with lower levels of education, such as access to healthcare, health-seeking behaviours and general lifestyle (30-36). Alternatively, biological factors, including lipid and glycaemic traits, may also play a part.

### Findings in context

A number of studies have used observational multivariable regression methods to support mediating roles of BMI, SBP and smoking in the pathway between education and CVD risk (7, 8, 37, 38), with consistent results obtained using various measures of education, including time spent in schooling and academic qualifications. In an analysis of Dutch individuals, Kershaw *et al*, attributed almost 27% of the association between education and CHD to be due to smoking, with 10% and 5% attributed to obesity and hypertension respectively (37). Similarly, Dégano *et al* found 7% and 14% of the association between education and CVD could be explained by BMI and hypertension respectively (8). However, they did not find any evidence of smoking mediating the association. Veronesi *et al* analysed their data stratified by sex, but consistently found mediating effects of SBP and smoking in both males and females (38). The findings in our study show that observational estimates underestimate the mediating role of smoking, BMI and SBP compared with MR, likely due to measurement error in the mediators that bias observational estimates towards the null, but do not affect MR estimates to the same degree (12). Given the importance of measurement error as a source of bias in mediation analysis (9), the MR approach offers favorable opportunities for understanding mediation.

### Strengths and limitations

The major strength of our work is that it allowed for us to assess the causal role of mediators using MR, an approach that is robust to non-differential measurement error in the mediator. We have used multiple data sources and approaches, each with different potential sources of biases, to thus improve the reliability of our findings through triangulation (39). Furthermore, the mediated effects estimated were consistent across the two approaches and in statistical sensitivity analyses. The imprecision in the one-sample MR analysis demonstrated the need for very large sample sizes to achieve sufficient statistical power when estimating mediation in an MR framework. The results were complemented by the two-sample MR approach, which had greater statistical power, but may be susceptible to alternative sources of bias, including those related to participant overlap in the samples used to obtain genetic association estimates for the exposures and outcomes (40). Existing SBP GWAS meta-analyses have adjusted for BMI as a covariate, which could introduce collider bias (41, 42), and for this reason we performed a GWAS of SBP in UK Biobank to select instruments, without adjusting for BMI. We also applied a ‘split sample’ SBP GWAS approach for use in individual-level data (one-sample) MR to avoid overlapping populations in the genetic association estimates for the exposure and outcome (43), and any associated bias (40, 44).

For all CVD subtypes and individual risk factors considered, the largest effects of education were consistently seen with the MR approaches, with smaller effects seen in the analysis of observational data. Furthermore, MR is not affected by measurement error in the same way as observational analysis, and particular discrepancies to the observational analysis for the proportion of the effect of educational attainment mediated by each the risk factors. For example, BMI is precisely measured and has little daily variation – and correspondingly the estimates of the proportion of effect mediated by BMI in the observational and MR analyses are similar (15% and 18% respectively). In contrast, SBP and lifetime smoking are difficult to measure precisely – and the estimated proportion mediated is smaller in the observational analysis than the MR (11% vs. 21% for SBP, and 19% vs. 33% for smoking). Non-differential measurement error in a mediator leads to an underestimation of the proportion mediated, so this discrepancy between our observational and MR analyses may be attributable to genetic variants instrumenting a lifetime exposure and therefore MR analyses suffering less from measurement error (9). The estimates for all three risk factors together were more similar between observational and MR estimates, although for all models, the confidence intervals were wide.

A limitation of the UK Biobank is that it is not representative of the UK population and is subject to healthy volunteer bias (16). Selection bias can induce biases in the results of instrumental variable analyses (45). However, this analysis shows that the estimates from UK Biobank are consistent with those from large-scale GWAS consortia that meta-analyse across multiple studies. This suggests that if selection bias is present, it is unlikely to invalidate our conclusions (46). Furthermore, data on circulating lipid levels and glycaemic traits were limited in this population, and we were therefore unable to investigate whether the effect of education on cardiovascular risk was also mediated through these pathways.

A limitation of the methods used for mediation analysis is that different methods have been used when considering the risk factors individually and jointly.

When estimating the indirect effects of a mediator on a binary outcome, the product of coefficients method results in the least amount of bias (27), and as such we used this approach for estimating the effects of education through each risk factor individually. However, this method cannot currently be used to consider multiple mediators simultaneously in an MR model. For this reason, we used the difference method for estimating the effect of education through the three considered risk factors collectively with MR. Although such an approach may introduce theoretical risk of bias when investigating a binary outcome, individual level data analyses in UK Biobank were also carried out on a linear risk difference scale to minimise any related issues. Estimates for the effect of education through the risk factors collectively were consistent between different scales in these analyses, and as such we would not expect any potential biases to alter the interpretation of our results.

### Clinical and public health implications

Intervening on education is difficult to achieve without social and political reform. Targeting BMI, SBP and smoking could therefore help to reduce inequalities in CVD risk related to educational attainment. However, these three risk factors together explain less than half of the overall effect of education. Further research identifying the other related factors and the interplay between them will be key to reducing social injustice.

### Conclusion

Using distinct analytical methodologies, including genetic approaches that are able draw causal inference, our results suggest that interventions aimed at reducing BMI, SBP and smoking in the population would lead to reductions in cases of CVD attributable to lower levels of education. Importantly, over half of the effect of education on risk of cardiovascular disease is not mediated through these risk factors and further work is required towards investigating this.

## Supporting information

## Acknowledgements

No funding body has influenced data collection, analysis or its interpretations. This publication is the work of the authors, who serve as the guarantors for the contents of this paper. This work was carried out using the computational facilities of the Advanced Computing Research Centre - http://www.bris.ac.uk/acrc/ and the Research Data Storage Facility of the University of Bristol - http://www.bris.ac.uk/acrc/storage/. This research was conducted using the UK Biobank Resource using application 10953.

## Author contributions

ARC and DG devised the project, analyzed and cleaned the data, interpreted results, wrote and revised the manuscript. AET, NMD, TT, JV, SB, GDS, MVH, JT, LDH and AD devised the project, interpreted the results and revised the manuscript. TT, JV, GH, SB and GDS contributed to the design of the project and critically revised the manuscript. RW, MM, RM, SS and DW provided data, and critically reviewed and revised the manuscript.

