## Supplementary material for "What explains the effect of education on cardiovascular disease? Applying Mendelian randomization to identify the consequences of education inequality"

Supplementary methods, figures and tables

### Supplementary methods

#### UK Biobank exclusion criteria

Individuals were excluded if their genetic sex differed to their reported gender from the assessment centre or for having aneuploidy of their sex chromosomes. Further individuals were excluded for being outliers for their heterozygosity and any missing genetic data. Finally, related individuals were also excluded from analyses, and the remaining subset was a maximal set of unrelated individuals. This exclusion list was derived in-house using an algorithm applied to the list of all the related pairs provided by UK Biobank (3rd degree or closer) (Supplementary Figure 1). It preferentially removes the individuals related to the greatest number of other individuals until no related pairs remain (1). Additionally, individuals were excluded if there were any missing data for education, body mass index (BMI), systolic blood pressure (SBP), smoking, cardiovascular disease (CVD) status or for any of the variables considered as confounders.

#### UK Biobank observational phenotypes

##### Body mass index

BMI was measured at the baseline assessment centre from baseline measures of height and weight used to calculate BMI (kg/m^2^).

In individual level MR analysis, we used 77 SNPs which had attained genome-wide significance for BMI in the GIANT consortium genome-wide association study (GWAS) analysis of individuals with European ancestry (2). Alleles were harmonised to all reflect BMI increasing SNPs and individual variants were recoded as 0, 1 or 2 according to the number of BMI increasing alleles. A genetic score for BMI was created by weighting each SNP by its relative effect size in the GWAS and summing all variants together in an additive model.

##### Blood pressure

Blood pressure was recorded both automatically and manually at the baseline assessment centre. All participants had an automatic reading, but manual readings were only taken for a subset. Each reading was taken twice, two minutes apart. This analysis uses the second reading of the automated blood pressure, where missing data were replaced with the first measure or any follow up assessment centre measures.

Participants were required to take all medication they are using to the assessment centre, details of which were recorded by nurses. We generated a variable for antihypertensive use based on the treatments recorded and 10mmHg was added to SBP measurements for these individuals, consistent with previous studies (3).

##### Smoking

A measure of lifetime smoking was constructed in the UK Biobank from self-reported age at initiation, age at cessation and cigarettes per day. From this information, smoking duration and time since cessation were calculated. The lifetime smoking measure further includes a simulated constant (half-life) which captures the exponentially decreasing effect of cigarettes on health over time. Aspects of smoking behaviour were combined into one score ranging from 0 (for non-smokers) to 4.17. The mean lifetime smoking score was 0.35 (standard deviation = 0.69). Full details of score construction can be found elsewhere (4).

##### Covariates

Variables considered as covariates were measured at the baseline assessment centres through interviews. Sex and ethnicity were confirmed according to genetic data. Place of Birth was adjusted for by the northing and easting birth location coordinates. Although the Townsend Deprivation Index (TDI) of historic birth locations are not recorded in UK Biobank, this has been estimated from the index of multiple deprivation indices using the current TDI of birth location as a proxy for historic birth place TDI. Mendelian Randomization models were also adjusted for the same confounders. Although a core assumption of MR is that the genetic variants are unrelated to confounders, there is some evidence of small associations with place of birth for the educational attainment variants in UK Biobank (5). These were only considered in observational and one sample MR analyses, where individual level data were available.

#### One-sample MR instrument selection (including GWAS methods)

For the one-sample MR analysis using individual-level data, instruments were selected from analysis of populations that did not overlap with those considered in the outcome estimates. Accordingly, we used 74 independent single-nucleotide polymorphisms (SNPs) that attained genome-wide significance (P<5x10^-8^) for education reported in main results from an earlier 2016 SSGAC GWAS meta-analysis of 293,723 individuals that did not include UK Biobank participants, to create a weighted allele score (6). The 77 reported genome-wide significant SNPs from the GIANT consortium’s meta-analysis of 322,154 individuals of European ancestry identified were used to create a weighted allele score for BMI (2). Five instruments for education were not available in UK Biobank and proxy SNPs in perfect LD (r^2^=1) were used (Supplementary Table 4).

Mendelian randomization studies require the SNP-exposure and the SNP-outcome associations to be estimated in independent samples, otherwise the MR estimates can be overestimated (7, 8). Existing SBP and lifetime smoking GWASs have been estimated using UK Biobank data (4, 9, 10). To avoid participant overlap for exposure and outcome genetic estimates in the UK Biobank (7), split sample GWASs of SBP and smoking respectively were performed using the University of Bristol MRC Integrative Epidemiology Unit GWAS Pipeline (11). A total of 318,147 UK Biobank participants were eligible for inclusion to the GWAS (see Supplementary Figure 2). All the eligible participants were randomly allocated into one of two halves (sample 1 and sample 2). A GWAS was performed on both samples 1 and 2 separately, adjusted for age, sex and the first 40 principle components in UK Biobank. A BOLT-LMM model was used to account for population stratification. The top hit SNPs were determined using the ‘clump_data’ command in the Two-Sample MR R package (R^2^ > 0.001) (12). This process was carried out for both SBP and lifetime smoking phenotypes.

The genetic score was created for each sample independently, by weighting each SNP by its relative effect size from the GWAS results of the opposing sample (i.e. the genome-wide significant SNPs and betas identified in the GWAS of sample 1 were used to generate the genetic score in sample 2 individuals). All genetic variants were summed together in an additive model. A total of 65 and 55 genome-wide significant SNPs were identified for SBP (with 10mmHg added for antihypertensive use) for sample 1 and sample 2 respectively (see Supplementary Table 5). In the split-sample GWAS for smoking, 17 SNPs were identified in the GWAS of sample 1 individuals and 16 SNPs in sample 2 individuals (see Supplementary Table 6).

Additionally, published SBP GWAS have been adjusted for BMI. Given the consideration for BMI as a mediator in this analysis and the potential for collider bias, we carried out a further GWAS of SBP in UK Biobank using the full eligible sample in a single GWAS to provide instruments for two-sample MR analysis. This used the same pipeline and adjustments as described previously for the split sample analysis. As with the split sample approach, the top hits were determined using the ‘clump_data’ command in the Two-Sample MR R package (13).

#### GWAS meta-analysis data: MI and stroke data sources

For the risk of MI, we used genetic association estimates were generated from the CARDIoGRAMplusC4D subgroup analysis of approximately 70% of the total cases from that had a reported history of MI (14). For risk of stroke risk, we used publicly available genetic association estimates from the MEGASTROKE consortium GWAS meta-analysis of 67,162 stroke cases (comprising of ischaemic stroke, intracerebral haemorrhage and stroke of unknown type) and 406,111 controls (15).

All genetic association estimates used in each two-sample MR analyses are provided in Supplementary Tables 7-21.

#### Statistical analysis for one-sample MR

In the one-sample MR of UK Biobank data, the total effect of education on CVD outcomes was investigated using two-stage least squares regression. In the first regression, we estimated the effect of the education weighted allele score on self-reported educational attainment. We used this estimate to generate a prediction of educational attainment. In the second stage, we estimate the effect of predicted educational attainment on the CVD outcome using robust standard errors (16). Both regression stages were adjusted for adjusted for age, sex, place of birth, birth distance from London, and TDI as well as the first ten principal components (PCs).

To estimate the effect of the education weighted allele score on each of BMI, SBP and smoking, the Stata IVREG2 package was used, adjusted covariates and PCs as above.

To then estimate the effect of each risk factor individually on the CVD outcomes, linear regression models were used to estimate the gene-exposure association between the weighted allele score for each risk factor and the observed value, whilst controlling for the weighted allele score of education. Additionally, the gene-education estimates whilst controlling for the allele score of the risk factor were calculated. For both models, the predicted values were stored for use in a second stage regression, where they were regressed against each CVD outcome risk using logistic regression. The final estimate of interest came from the predicted value of the mediator, controlling for education and all other covariates and PCs as previously described. Where split sample GWAS estimates were used to create the allele score in SBP and smoking the MR analyses were run separately for each 50% sample and meta-analysed to estimate an overall effect.

These estimates were then multiplied to estimate the indirect effect, that is the amount of the association between education and CVD going via each of the three risk factors individually.

#### Sensitivity analyses

MR estimates are prone to bias if the underlying assumptions of the analysis are violated (17). Horizontal pleiotropy, where a genetic variant is associated to the outcome of interest via an alternative pathway, can potentially bias the MR estimates (17). MR-Egger allows for directional horizontal pleiotropy under the assumption that the size of the effects of the variants on the exposure are independent of their direct effects on the outcome (18). Furthermore, the weighted median estimator is able to provide robust MR estimates when more than half of the information for the analysis comes from valid instruments (19). In the MR analysis of the total effect of education on CVD outcome risk, and the effect of education on each risk factor, we also perform these techniques to investigate the robustness of our findings when relaxing assumptions on horizontal pleiotropy. Incidentally, these techniques are not yet developed for application in MR mediation analysis.

For all analyses in UK Biobank, models were replicated on the risk difference scale using multivariable linear regression. For multivariable observational analyses multivariable linear regression was used and for the one-sample MR analyses the IVREG2 Stata package was used (20). Additionally, all analyses were replicated using unadjusted models and models adjusted for age and sex only.

### Supplementary Figures

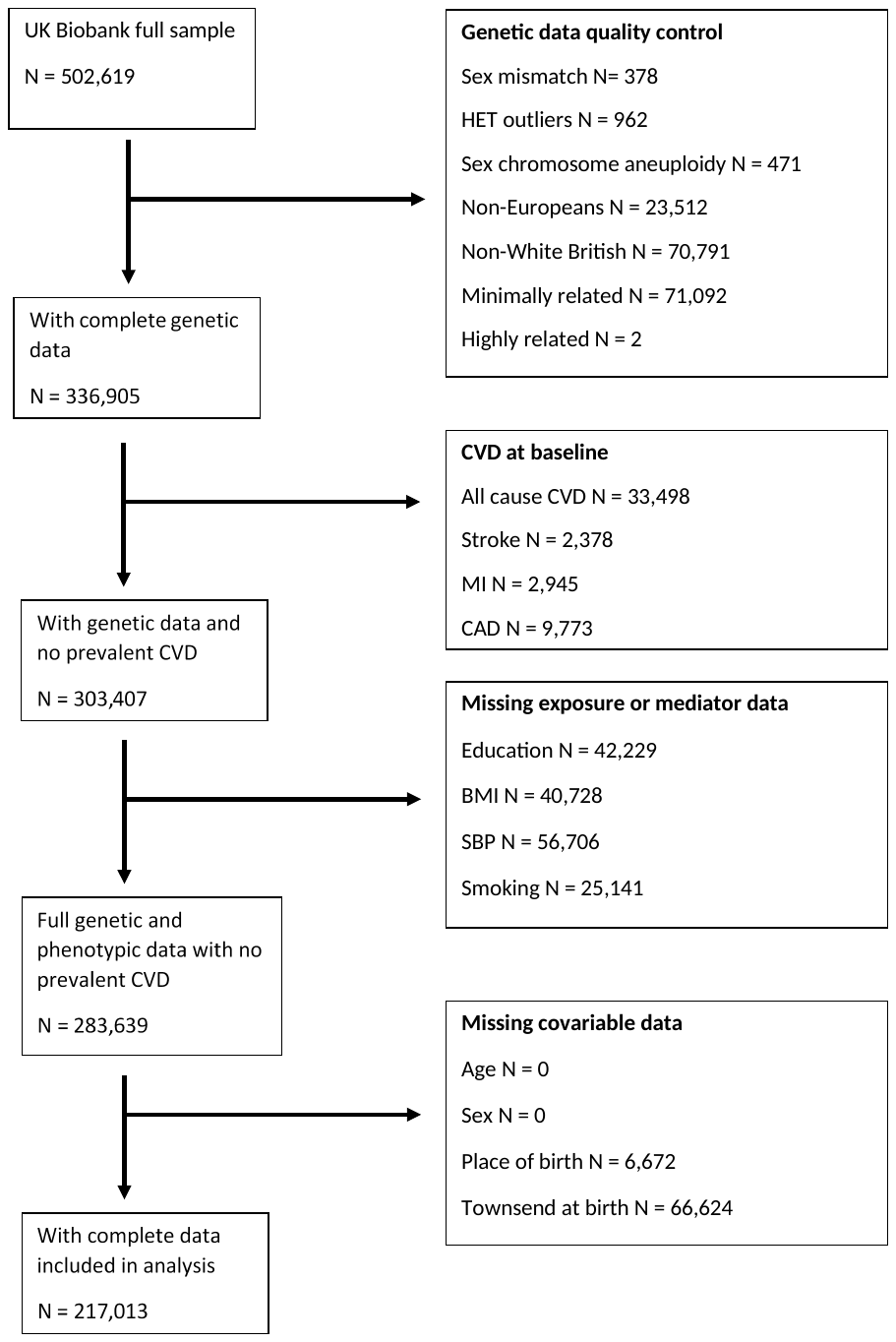

#### **Supplementary Figure 1**: Flow chart for exclusions made in UK Biobank for resultant sample for mediation analysis

#### **Supplementary Figure 2**: Flow chart for exclusions made in UK Biobank for use in SBP and smoking GWAS analyses

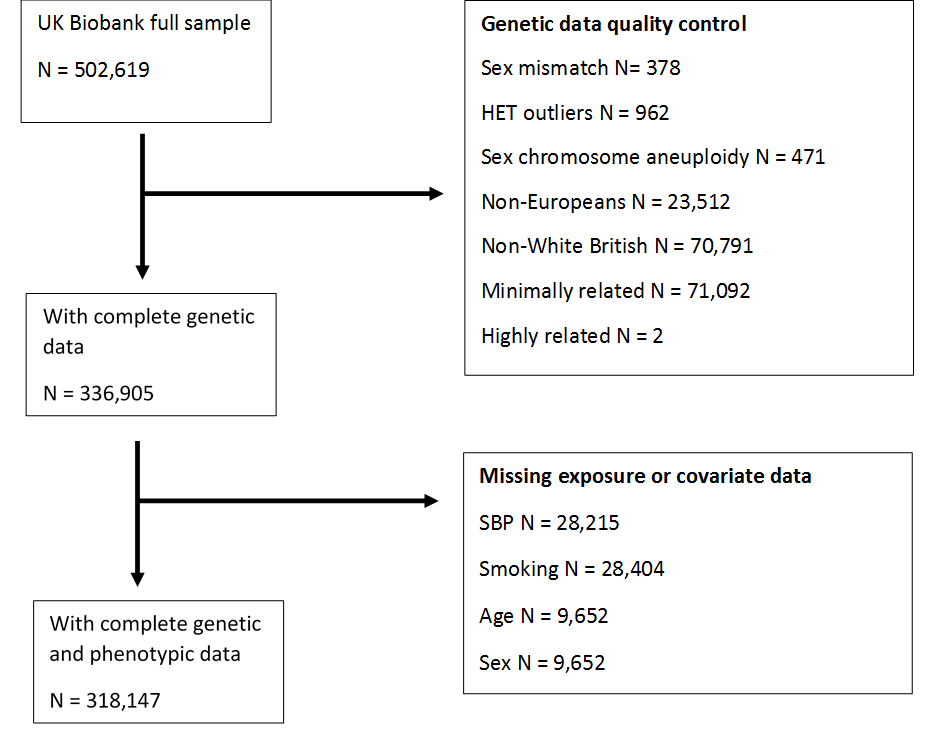

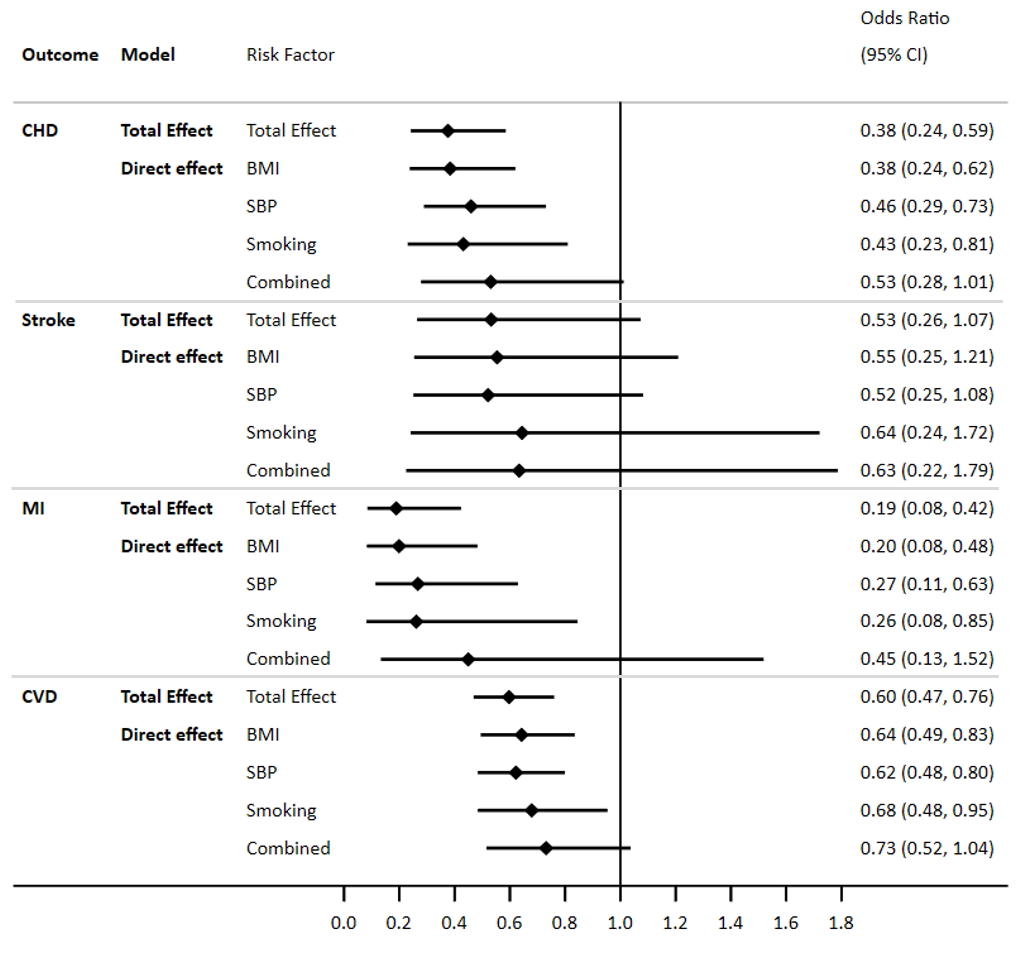

#### **Supplementary Figure 3**: Total and Direct effects for the association between 1-SD higher education and CVD outcomes estimated in one-sample MR analyses.

Total effect = overall effect of education on CVD, including that mediated by BMI, SBP, smoking, and all other risk factors on the causal pathway. Direct effect = the effect of education on CVD that is not mediated by the risk factor listed. E.g. direct effect of education on CHD with risk factor BMI is the effect of education on CHD that is not mediated by BMI. Analyses adjusted for: age, sex, place of birth and Townsend deprivation index at birth. BMI, SBP and smoking were measured in 1-SD units.

#### **Supplementary Figure 4**: Association between 1-SD higher education and BMI, SBP and smoking in one-sample MR.

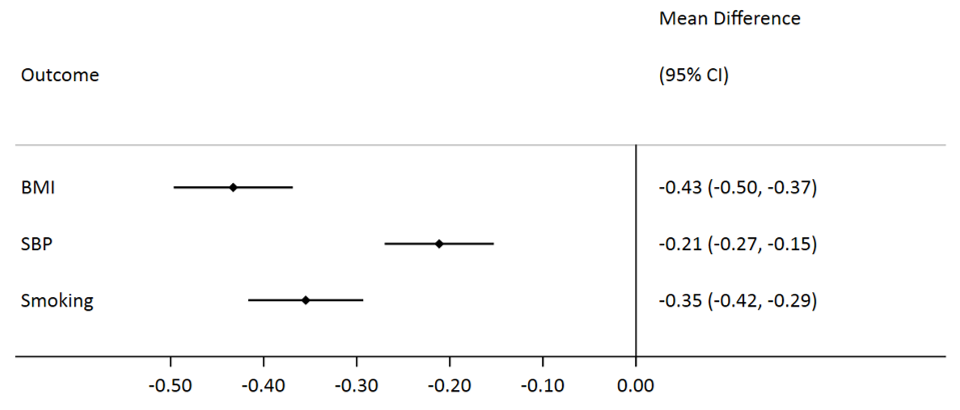

Analyses were adjusted for: age, sex, place of birth and Townsend deprivation index at birth. BMI, SBP and smoking were measured in 1-SD units.

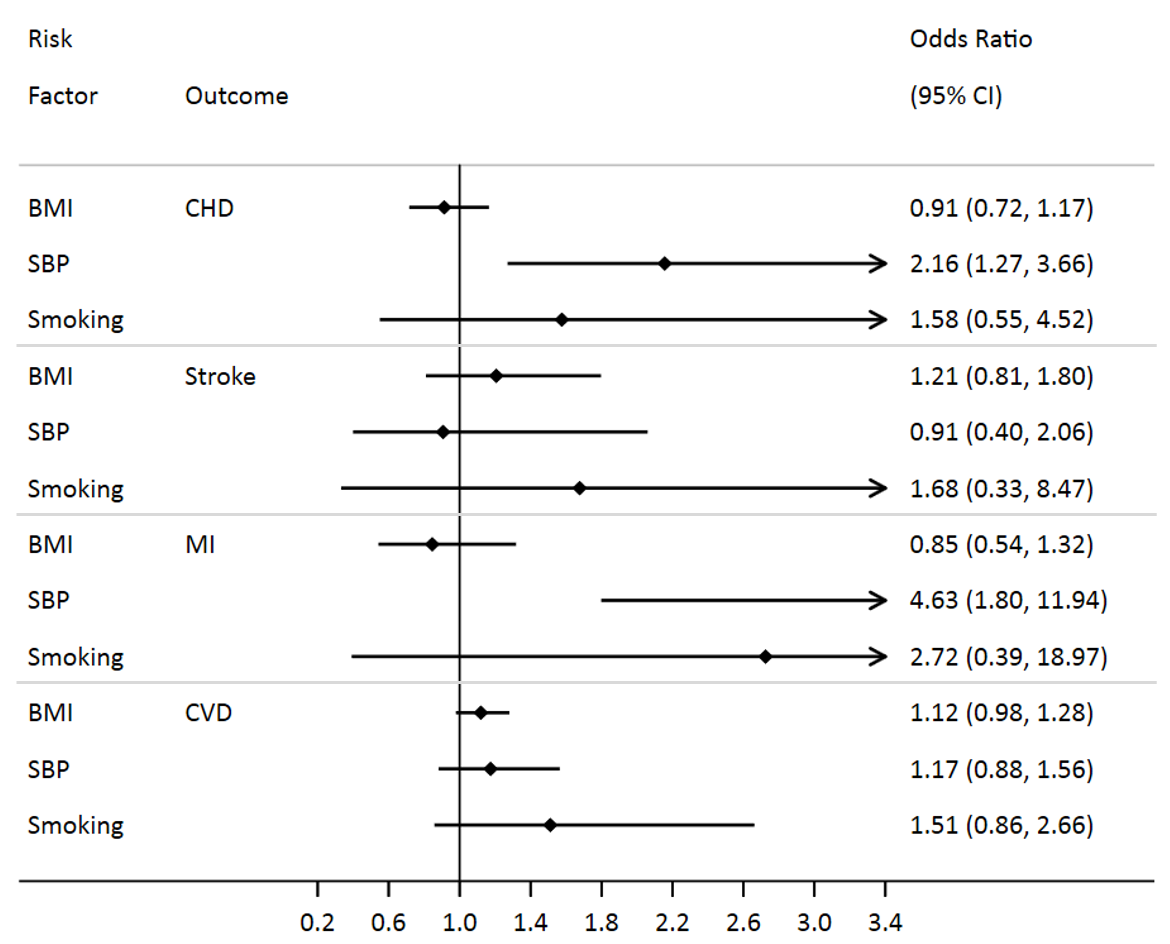

#### **Supplementary Figure 5**: Associations between BMI, SBP and smoking, and risk of the CVD outcomes in one-sample MR.

Analyses adjusted for: age, sex, place of birth and Townsend deprivation index at birth. BMI, SBP and smoking were measured in 1-SD units.

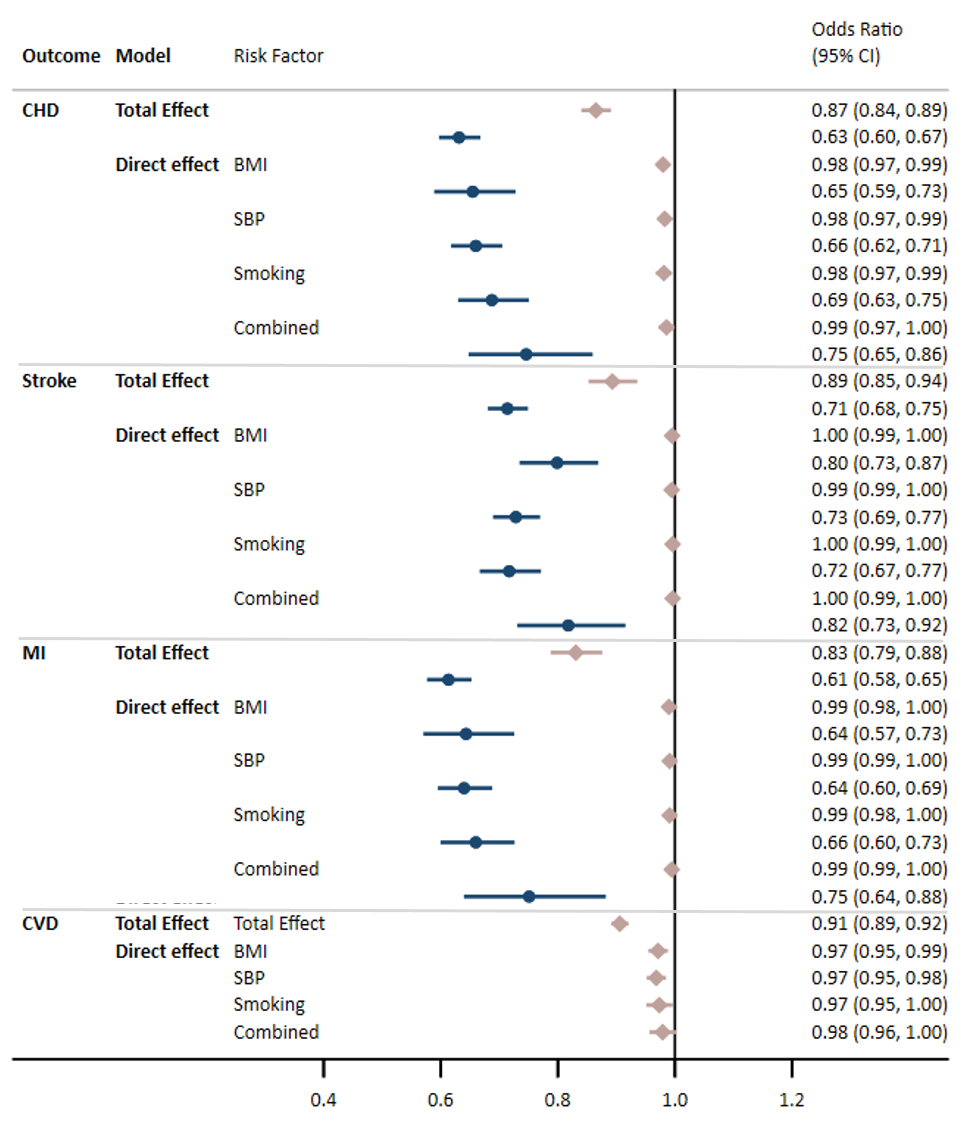

#### **Supplementary Figure 6**: Total and Direct effects for the association between 1-SD higher education and CVD outcomes estimated in multivaribale observational analyses and two-sample MR analyses.

Total effect = overall effect of education on CVD, including that mediated by BMI, SBP, smoking, and all other risk factors on the causal pathway. Direct effect = the effect of education on CVD that is not mediated by the risk factor listed. E.g. direct effect of education on CHD with risk factor BMI is the effect of education on CHD that is not mediated by BMI. Analyses adjusted for: age, sex, place of birth and Townsend deprivation index at birth. BMI, SBP and smoking were measured in 1-SD units.

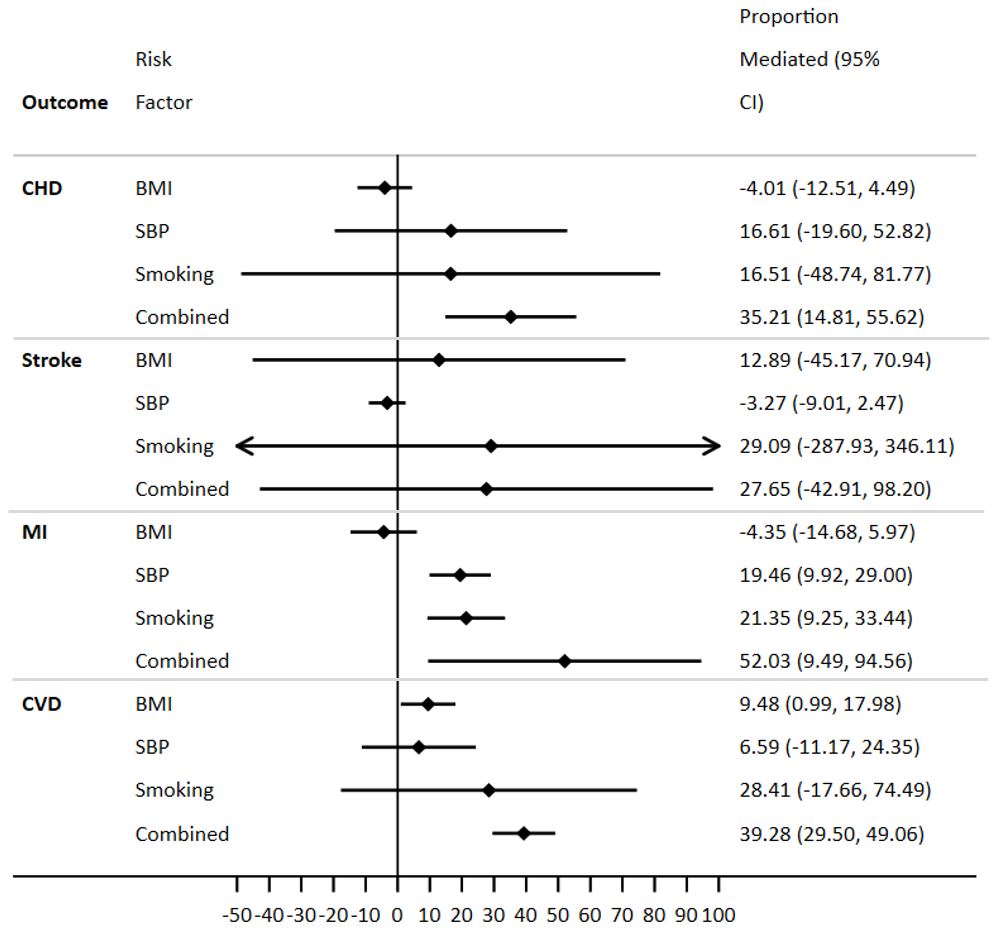

#### **Supplementary Figure 7:** Estimates for the effect of education on CVD and its subtypes explained by BMI, SBP and smoking respectively.

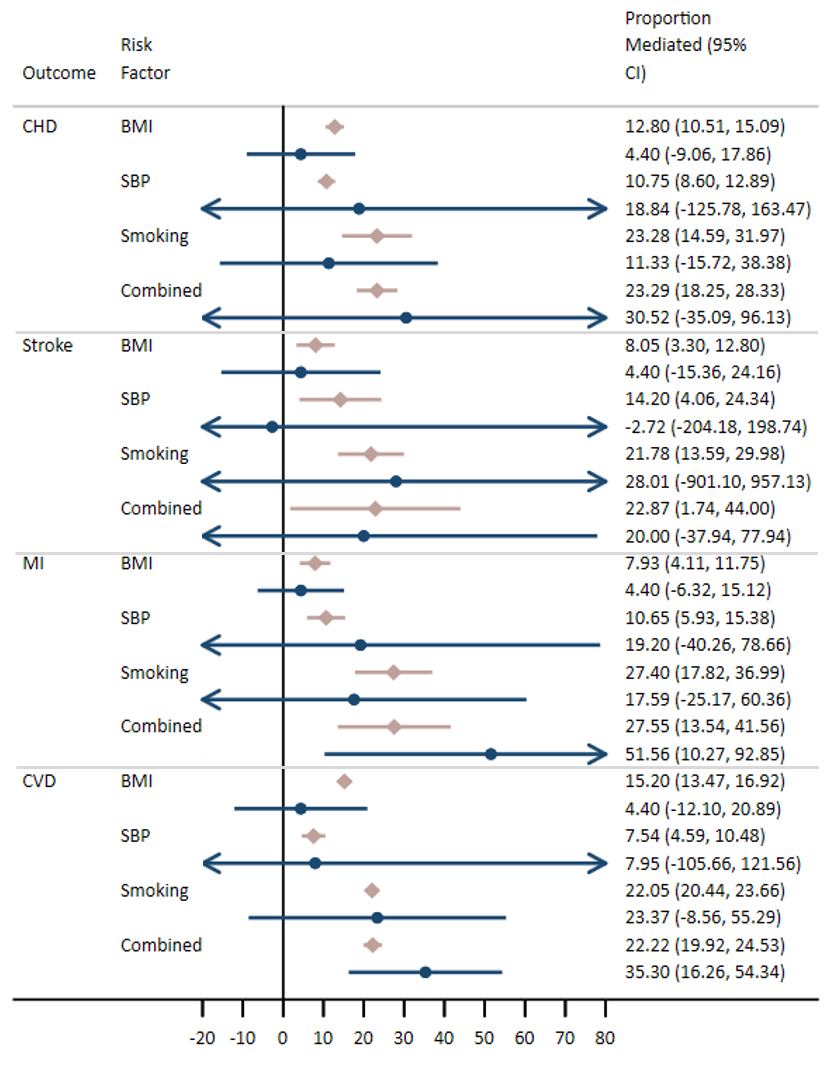

#### **Supplementary figure 8:** Estimates for the effect of education on CVD and its subtypes explained by BMI, SBP and smoking respectively, estimated on the risk difference scale using the difference method. Results are provided for the multivariable observational analysis (plotted in pink) and one-sample MR (plotted in navy).

### Supplementary Tables

#### Supplementary table 1: International Standard for Classification of Education codes mapped to UK Biobank self-report highest qualification to estimate years of education

| **Qualification (As reported in UK Biobank)** | **ISCED** | **Years of education** |
| --- | --- | --- |
| **College or University degree** | 5 | 20 |
| **NVQ or HND or HNC or equivalent** | 5 | 19 |
| **Other prof. qual. eg: nursing, teaching** | 4 | 15 |
| **A levels/AS levels or equivalent** | 3 | 13 |
| **O levels/GCSEs or equivalent** | 2 | 10 |
| **CSEs or equivalent** | 2 | 10 |
| **None of the above** | 1 | 7 |
| **Prefer not to answer** | Excluded | |

#### Supplementary table 2: ICD codes for cardiovascular outcomes in UK Biobank

| **Outcome** | **ICD-9 code** | **ICD-10 code** |
| --- | --- | --- |
| CVD (all subtypes) | 390-459 | I, G45 |
| Stroke | 434.91 | I6, G45 |
| MI | 410.9, 412.9 | I21, I22 |
| CHD | 410-414 | I20-I25 |

#### Supplementary table 3: Cohort descriptives for UK Biobank sample used in multivariable observational and one-sample MR analysis compared with the whole UK Biobank cohort

| **Variable** | **Level** | **N Analysis Sample** | **% Analysis Sample** |  | **N Full UKBB** | **% Full UKBB** |
| --- | --- | --- | --- | --- | --- | --- |
| **Sex** | Male | 108,563 | 46 |  | 229,164 | 45.63 |
|  | Female | 127,867 | 54 |  | 273,076 | 54.37 |
| **Age** | <40 | 2,342 | 0.99 |  | 5,424 | 1.08 |
|  | 41-50 | 56,888 | 24.06 |  | 126,426 | 25.15 |
|  | 51-60 | 83,219 | 35.2 |  | 177,264 | 35.27 |
|  | 61-70 | 93,976 | 39.75 |  | 193,119 | 38.42 |
|  | 71+ | 5 | <0.01 |  | 422 | 0.08 |
| **ISCED Category** | 1 | 39,396 | 16.77 |  | 84,895 | 17.23 |
|  | 2 | 41,501 | 28.27 |  | 82,757 | 16.79 |
|  | 3 | 12,787 | 11.62 |  | 27,008 | 5.48 |
|  | 4 | 29,298 | 5.13 |  | 58,680 | 11.91 |
|  | 5 | 113,448 | 38.22 |  | 239,464 | 48.59 |
| **BMI** | Underweight | 1,143 | 0.48 |  | 2,624 | 0.52 |
|  | Normal | 77,598 | 32.82 |  | 162,261 | 32.28 |
|  | Overweight | 101,288 | 42.84 |  | 212,071 | 42.19 |
|  | Obese | 56,401 | 23.86 |  | 125,699 | 25.01 |
| **SBP** | (Mean) | 136.39 |  |  | 135.95 |  |
| **SBP - Medication adjusted** | (Mean) | 137.3 |  |  | 136.86 |  |
| **Smoking initiation** | Never | 94,058 | 39.78 |  | 200,747 | 40.2 |
|  | Ever (inc current and former) | 142,372 | 60.22 |  | 298,665 | 59.8 |
| **CVD** | Case | 198,207 | 92.77 |  | 418,126 | 92.38 |
|  | Control | 15,448 | 7.23 |  | 34,513 | 7.62 |
| **Stroke** | Case | 198,207 | 99.18 |  | 418,126 | 99.09 |
|  | Control | 1,640 | 0.82 |  | 3,840 | 0.91 |
| **Acute Myocardial Infarction** | Case | 198,207 | 99.37 |  | 418,126 | 99.32 |
|  | Control | 1,265 | 0.63 |  | 2,860 | 0.68 |
| **Coronary Heart Disease** | Case | 198,207 | 97.82 |  | 418,126 | 97.74 |
|  | Control | 4,407 | 2.18 |  | 9,677 | 2.26 |
| **BMI** | R-square value for instrument | 0.017 |  |  |  |  |
| **SBP** | Sample 1 - R-square value for instrument | 0.0035 |  |  |  |  |
|  | Sample 2 - R-square value for instrument | 0.0027 |  |  |  |  |
| **Lifetime smoking** | Sample 1 - R-square value for instrument | 0.0012 |  |  |  |  |
|  | Sample 2 - R-square value for instrument | 0.0014 |  |  |  |  |

#### Supplementary table 4: Proxy SNPs for education instrument used in one-sample MR analysis

| **GWAS SNP (Okbay)** | **SNP in LD used (UKBB)** |
| --- | --- |
| rs114598875 | rs17538393 |
| rs148734725 | rs9878943 |
| rs9320913 | rs1487445 |
| rs8005528 | rs8008779 |
| rs192818565 | rs55943044 |

#### Supplementary table 5: Genome-wide significant SNPs for SBP from split sample GWAS analysis in UK Biobank

| **Chromosome** | **RSID** | **Position** | **Beta** | **SE** | **Other Allele** | **P Value** | **Sample** |
| --- | --- | --- | --- | --- | --- | --- | --- |
| 1 | rs5068 | 11905974 | 1.3283 | 0.1374 | A | 4.10E-22 | 1 |
| 1 | rs448385 | 25395133 | -0.3464 | 0.0630 | G | 3.80E-08 | 1 |
| 1 | rs3790604 | 113046879 | -0.8724 | 0.1204 | C | 4.30E-13 | 1 |
| 1 | rs2765524 | 89417695 | 0.3965 | 0.0641 | C | 6.00E-10 | 1 |
| 2 | rs953246 | 146335486 | -0.3924 | 0.0685 | T | 1.00E-08 | 1 |
| 2 | rs1344653 | 19730845 | -0.3481 | 0.0626 | A | 2.70E-08 | 1 |
| 2 | rs1009358 | 65276452 | 0.3693 | 0.0645 | T | 1.00E-08 | 1 |
| 2 | rs268263 | 164954174 | -0.5652 | 0.0735 | T | 1.50E-14 | 1 |
| 2 | rs35021474 | 26916844 | 0.4639 | 0.0645 | C | 6.30E-13 | 1 |
| 2 | rs2867114 | 651380 | 0.6149 | 0.1073 | C | 1.00E-08 | 1 |
| 3 | rs3821843 | 53558012 | -0.4129 | 0.0681 | G | 1.40E-09 | 1 |
| 3 | rs1343040 | 169186293 | -0.4317 | 0.0637 | G | 1.20E-11 | 1 |
| 3 | rs2643826 | 27562988 | -0.3780 | 0.0631 | C | 2.20E-09 | 1 |
| 3 | rs263016 | 183502559 | 0.3475 | 0.0631 | T | 3.60E-08 | 1 |
| 4 | rs10857147 | 81181072 | -0.8157 | 0.0690 | A | 3.00E-32 | 1 |
| 4 | rs6825268 | 26783453 | -0.3573 | 0.0631 | A | 1.50E-08 | 1 |
| 4 | rs13107325 | 103188709 | 0.7689 | 0.1190 | C | 1.00E-10 | 1 |
| 4 | rs1842896 | 156511459 | -0.4014 | 0.0626 | G | 1.40E-10 | 1 |
| 5 | rs13436194 | 157803588 | 0.4570 | 0.0632 | A | 4.80E-13 | 1 |
| 5 | rs12656497 | 32831939 | -0.7161 | 0.0638 | T | 3.00E-29 | 1 |
| 6 | rs2607015 | 31762843 | -0.4232 | 0.0636 | G | 2.90E-11 | 1 |
| 6 | rs2499801 | 96854594 | 0.4442 | 0.0806 | G | 3.60E-08 | 1 |
| 6 | rs13219548 | 127165290 | -0.4051 | 0.0630 | C | 1.30E-10 | 1 |
| 7 | rs62481856 | 106412082 | -0.8444 | 0.0789 | G | 9.30E-27 | 1 |
| 7 | rs2854747 | 45959917 | 0.4198 | 0.0637 | G | 4.40E-11 | 1 |
| 7 | rs10241964 | 19042114 | -0.5983 | 0.1056 | G | 1.40E-08 | 1 |
| 7 | rs10269774 | 92253972 | 0.3854 | 0.0669 | G | 8.40E-09 | 1 |
| 7 | rs891511 | 150704843 | 0.3765 | 0.0681 | G | 3.20E-08 | 1 |
| 7 | rs2023843 | 27243221 | -0.8627 | 0.1198 | C | 6.00E-13 | 1 |
| 7 | rs3823483 | 131010943 | -0.3546 | 0.0634 | T | 2.30E-08 | 1 |
| 8 | rs877116 | 10712945 | 0.4697 | 0.0636 | G | 1.50E-13 | 1 |
| 8 | rs7463212 | 143991858 | 0.3776 | 0.0629 | T | 2.00E-09 | 1 |
| 8 | rs73563812 | 25900405 | 0.4215 | 0.0737 | G | 1.10E-08 | 1 |
| 10 | rs56137952 | 134376691 | -0.5494 | 0.0985 | G | 2.40E-08 | 1 |
| 10 | rs10883543 | 102552752 | -0.6995 | 0.0996 | G | 2.20E-12 | 1 |
| 10 | rs10995311 | 64564934 | 0.3941 | 0.0634 | C | 5.00E-10 | 1 |
| 10 | rs11191580 | 104906211 | 1.0970 | 0.1180 | T | 1.50E-20 | 1 |
| 10 | rs7076938 | 115789375 | -0.4606 | 0.0711 | C | 9.30E-11 | 1 |
| 10 | rs12258967 | 18727959 | 0.6582 | 0.0686 | C | 8.00E-22 | 1 |
| 10 | rs7922049 | 63462365 | 0.5533 | 0.0869 | G | 1.90E-10 | 1 |
| 10 | rs10786156 | 96014622 | 0.3767 | 0.0632 | C | 2.50E-09 | 1 |
| 11 | rs55925664 | 10192809 | -0.6415 | 0.0805 | T | 1.60E-15 | 1 |
| 11 | rs7120737 | 47702395 | 0.6316 | 0.0891 | A | 1.30E-12 | 1 |
| 11 | rs10750766 | 65473798 | -0.3930 | 0.0691 | C | 1.30E-08 | 1 |
| 11 | rs633185 | 100593538 | -0.6945 | 0.0698 | G | 2.60E-23 | 1 |
| 11 | rs12418543 | 1894163 | 0.5668 | 0.0645 | A | 1.50E-18 | 1 |
| 11 | rs747249 | 130271647 | 0.3747 | 0.0656 | A | 1.10E-08 | 1 |
| 12 | rs73437338 | 90054619 | 0.7678 | 0.0843 | T | 8.80E-20 | 1 |
| 12 | rs4766578 | 111904371 | 0.4066 | 0.0626 | T | 8.60E-11 | 1 |
| 12 | rs35444 | 115552437 | 0.3909 | 0.0643 | A | 1.20E-09 | 1 |
| 12 | rs73073676 | 20351276 | 0.3773 | 0.0670 | A | 1.80E-08 | 1 |
| 15 | rs8039305 | 91422543 | -0.5943 | 0.0630 | T | 4.00E-21 | 1 |
| 15 | rs1717200 | 41368334 | -0.3912 | 0.0628 | A | 4.70E-10 | 1 |
| 15 | rs1543927 | 75063573 | 0.4218 | 0.0712 | T | 3.10E-09 | 1 |
| 15 | rs11634851 | 81028965 | -0.4216 | 0.0628 | C | 1.90E-11 | 1 |
| 16 | rs77870048 | 69965021 | -0.9117 | 0.1399 | C | 7.20E-11 | 1 |
| 16 | rs2188717 | 24730230 | -0.5361 | 0.0792 | T | 1.30E-11 | 1 |
| 17 | rs9907379 | 59489893 | -0.4257 | 0.0769 | T | 3.20E-08 | 1 |
| 17 | rs60289499 | 43218677 | -0.4564 | 0.0708 | G | 1.10E-10 | 1 |
| 17 | rs34710835 | 45146717 | 0.5223 | 0.0643 | C | 4.40E-16 | 1 |
| 17 | rs11650511 | 1337960 | -0.4085 | 0.0634 | C | 1.20E-10 | 1 |
| 19 | rs73046792 | 49605705 | 0.4714 | 0.0843 | G | 2.30E-08 | 1 |
| 19 | rs12978472 | 7257990 | 0.8468 | 0.0941 | C | 2.20E-19 | 1 |
| 20 | rs74729242 | 57718690 | -0.5901 | 0.1013 | T | 5.80E-09 | 1 |
| 20 | rs2423514 | 10693337 | 0.3709 | 0.0628 | A | 3.50E-09 | 1 |
| 1 | rs55857306 | 11895795 | 0.7687 | 0.0850 | G | 1.60E-19 | 2 |
| 1 | rs4648815 | 1687152 | 0.3807 | 0.0638 | G | 2.40E-09 | 2 |
| 1 | rs6541328 | 230833262 | -0.5551 | 0.1010 | A | 3.80E-08 | 2 |
| 1 | rs778121 | 56620268 | -0.3887 | 0.0659 | T | 3.60E-09 | 2 |
| 1 | rs6657049 | 115825531 | -0.3665 | 0.0658 | G | 2.50E-08 | 2 |
| 2 | rs268263 | 164954174 | -0.5602 | 0.0738 | T | 3.20E-14 | 2 |
| 2 | rs4666493 | 19765225 | -0.3641 | 0.0639 | G | 1.20E-08 | 2 |
| 2 | rs1275988 | 26914364 | 0.4835 | 0.0650 | C | 9.90E-14 | 2 |
| 2 | rs6724607 | 191466532 | 0.3656 | 0.0630 | A | 6.40E-09 | 2 |
| 3 | rs2307032 | 27432995 | 0.3879 | 0.0664 | T | 5.20E-09 | 2 |
| 3 | rs6442260 | 11590751 | 0.3595 | 0.0659 | G | 4.90E-08 | 2 |
| 4 | rs10024506 | 89764197 | 0.4173 | 0.0745 | G | 2.10E-08 | 2 |
| 4 | rs11099097 | 81167309 | -0.6371 | 0.0696 | C | 5.40E-20 | 2 |
| 4 | rs4690974 | 156393641 | -0.3741 | 0.0631 | T | 3.10E-09 | 2 |
| 4 | rs17010961 | 86723103 | -0.5775 | 0.0913 | T | 2.50E-10 | 2 |
| 5 | rs10059884 | 32832474 | -0.5859 | 0.0643 | C | 7.80E-20 | 2 |
| 5 | rs12652819 | 121244520 | 0.3775 | 0.0677 | A | 2.50E-08 | 2 |
| 5 | rs17715065 | 158261163 | -0.3662 | 0.0631 | C | 6.60E-09 | 2 |
| 5 | rs11241959 | 127787964 | -0.3715 | 0.0631 | A | 4.00E-09 | 2 |
| 5 | rs2964330 | 157743781 | -0.3645 | 0.0641 | G | 1.30E-08 | 2 |
| 6 | rs17080069 | 150989698 | 0.7199 | 0.1218 | A | 3.40E-09 | 2 |
| 6 | rs6923947 | 127098553 | -0.4974 | 0.0635 | G | 5.00E-15 | 2 |
| 6 | rs7889 | 31605448 | -0.3792 | 0.0657 | C | 7.80E-09 | 2 |
| 7 | rs891511 | 150704843 | 0.4141 | 0.0683 | G | 1.30E-09 | 2 |
| 7 | rs2392929 | 106414069 | -0.7165 | 0.0790 | T | 1.20E-19 | 2 |
| 7 | rs42032 | 92237426 | 0.3967 | 0.0720 | G | 3.60E-08 | 2 |
| 7 | rs57301765 | 19052733 | -0.5059 | 0.0866 | G | 5.20E-09 | 2 |
| 9 | rs2780072 | 9340831 | -0.5035 | 0.0902 | A | 2.40E-08 | 2 |
| 10 | rs76443711 | 75449789 | -0.5442 | 0.0915 | G | 2.70E-09 | 2 |
| 10 | rs7070797 | 63551773 | 0.6290 | 0.0905 | G | 3.60E-12 | 2 |
| 10 | rs11187838 | 96038686 | 0.5615 | 0.0637 | G | 1.20E-18 | 2 |
| 10 | rs732998 | 104897901 | 0.8365 | 0.1184 | T | 1.60E-12 | 2 |
| 10 | rs12258967 | 18727959 | 0.5893 | 0.0691 | C | 1.50E-17 | 2 |
| 11 | rs4980379 | 1888614 | -0.5896 | 0.0657 | C | 2.70E-19 | 2 |
| 11 | rs12807950 | 107057190 | -0.3790 | 0.0631 | T | 1.90E-09 | 2 |
| 11 | rs7107356 | 47676170 | -0.4793 | 0.0630 | A | 2.90E-14 | 2 |
| 11 | rs1216743 | 100573120 | -0.5662 | 0.0705 | G | 9.60E-16 | 2 |
| 12 | rs2681492 | 90013089 | 0.6765 | 0.0840 | T | 8.00E-16 | 2 |
| 12 | rs4767328 | 115929396 | -0.3575 | 0.0640 | G | 2.30E-08 | 2 |
| 12 | rs35427 | 115556307 | 0.4110 | 0.0664 | T | 6.00E-10 | 2 |
| 12 | rs4883481 | 50574311 | 0.4218 | 0.0652 | T | 1.00E-10 | 2 |
| 12 | rs597808 | 111973358 | 0.4310 | 0.0634 | A | 1.00E-11 | 2 |
| 15 | rs7176022 | 75107880 | 0.4365 | 0.0713 | A | 9.10E-10 | 2 |
| 15 | rs4932373 | 91429287 | -0.5144 | 0.0672 | A | 2.00E-14 | 2 |
| 15 | rs117539635 | 69682916 | 1.3387 | 0.2009 | A | 2.70E-11 | 2 |
| 16 | rs77870048 | 69965021 | -0.9216 | 0.1415 | C | 7.40E-11 | 2 |
| 16 | rs11646987 | 24832408 | 0.3922 | 0.0709 | G | 3.20E-08 | 2 |
| 17 | rs1436138 | 75316880 | 0.3615 | 0.0661 | A | 4.40E-08 | 2 |
| 17 | rs7217916 | 76769434 | 0.3944 | 0.0650 | A | 1.30E-09 | 2 |
| 17 | rs2301597 | 43173273 | 0.5081 | 0.0640 | T | 2.00E-15 | 2 |
| 17 | rs11874 | 45017193 | -0.5820 | 0.0917 | G | 2.20E-10 | 2 |
| 17 | rs4480845 | 1958609 | 0.4024 | 0.0662 | T | 1.20E-09 | 2 |
| 19 | rs167479 | 11526765 | 0.5743 | 0.0630 | G | 8.20E-20 | 2 |
| 20 | rs75777337 | 57702450 | -0.6125 | 0.1033 | T | 3.10E-09 | 2 |
| 20 | rs913220 | 10966476 | -0.4389 | 0.0649 | C | 1.40E-11 | 2 |

#### Supplementary table 6: Genome-wide significant SNPs for lifetime smoking from split sample GWAS analysis in UK Biobank

| **Chromosome** | **RSID** | **Position** | **Beta** | **SE** | **Other Allele** | **P Value** | **Sample** |
| --- | --- | --- | --- | --- | --- | --- | --- |
| 1 | rs499257 | 44078384 | 0.0146 | 0.0025 | T | 3.10E-09 | 1 |
| 2 | rs2890772 | 146175106 | -0.0139 | 0.0024 | G | 3.30E-09 | 1 |
| 3 | rs4856463 | 83638568 | 0.0155 | 0.0028 | C | 3.80E-08 | 1 |
| 3 | rs326341 | 107811142 | 0.0132 | 0.0023 | G | 1.30E-08 | 1 |
| 4 | rs6852351 | 28064697 | 0.0132 | 0.0024 | C | 4.10E-08 | 1 |
| 5 | rs986391 | 166993972 | 0.0151 | 0.0024 | G | 3.00E-10 | 1 |
| 5 | rs17159727 | 106632458 | 0.0241 | 0.0041 | T | 3.30E-09 | 1 |
| 6 | rs16879271 | 16822974 | -0.0323 | 0.0059 | A | 4.90E-08 | 1 |
| 7 | rs10233018 | 117523709 | -0.0130 | 0.0023 | A | 2.20E-08 | 1 |
| 7 | rs10226228 | 32315613 | -0.0142 | 0.0024 | A | 3.60E-09 | 1 |
| 8 | rs10093628 | 9393379 | -0.0165 | 0.0027 | T | 1.10E-09 | 1 |
| 9 | rs113382419 | 136463019 | -0.0244 | 0.0037 | C | 3.20E-11 | 1 |
| 11 | rs6590701 | 133315869 | -0.0145 | 0.0026 | G | 4.20E-08 | 1 |
| 11 | rs2155646 | 112912811 | -0.0160 | 0.0024 | T | 1.90E-11 | 1 |
| 11 | rs11030088 | 27646247 | -0.0159 | 0.0027 | G | 2.50E-09 | 1 |
| 12 | rs4763463 | 10355901 | 0.0132 | 0.0024 | G | 3.00E-08 | 1 |
| 15 | rs7173514 | 78849918 | 0.0223 | 0.0028 | C | 9.10E-16 | 1 |
| 1 | rs10922907 | 91193049 | 0.0133 | 0.0023 | A | 9.40E-09 | 2 |
| 2 | rs16824949 | 146168208 | -0.0144 | 0.0023 | G | 4.10E-10 | 2 |
| 2 | rs7559547 | 615627 | -0.0219 | 0.0030 | C | 6.10E-13 | 2 |
| 2 | rs1863161 | 60139524 | -0.0128 | 0.0023 | G | 3.60E-08 | 2 |
| 2 | rs75120545 | 44271496 | 0.0388 | 0.0071 | C | 4.70E-08 | 2 |
| 2 | rs263771 | 185921692 | -0.0151 | 0.0027 | C | 3.60E-08 | 2 |
| 3 | rs62261249 | 49594060 | -0.0158 | 0.0026 | T | 1.90E-09 | 2 |
| 7 | rs17657924 | 96625589 | 0.0132 | 0.0023 | C | 1.30E-08 | 2 |
| 9 | rs56116178 | 136460224 | -0.0306 | 0.0038 | A | 7.30E-16 | 2 |
| 9 | rs12553882 | 128195044 | -0.0142 | 0.0024 | G | 3.00E-09 | 2 |
| 11 | rs7948789 | 112839532 | -0.0163 | 0.0024 | A | 6.70E-12 | 2 |
| 14 | rs12897150 | 104319530 | -0.0136 | 0.0023 | A | 6.20E-09 | 2 |
| 15 | rs12901436 | 47683095 | -0.0196 | 0.0028 | C | 3.60E-12 | 2 |
| 15 | rs28669908 | 78910267 | 0.0238 | 0.0028 | C | 2.50E-17 | 2 |
| 20 | rs45577732 | 61983934 | -0.0386 | 0.0043 | C | 1.90E-19 | 2 |
| 20 | rs159058 | 31108108 | -0.0142 | 0.0025 | A | 1.90E-08 | 2 |

#### Supplementary table 7: Genetic association estimates for the two-sample regression-based MVMR analysis estimating the effect of BMI on CAD, adjusted for education. ea=effect allele; gx=BMI; gy=CAD; gz=education; se=standard error

| SNP | ea | gx | gx_se | gy | gy_se | gz | gz_se |
| --- | --- | --- | --- | --- | --- | --- | --- |
| rs10083803 | T | -0.0139 | 0.0022 | 0.0051 | 0.0107 | -0.0020 | 0.0019 |
| rs10124645 | A | 0.0124 | 0.0020 | 0.0061 | 0.0101 | -0.0041 | 0.0017 |
| rs10128597 | A | -0.0186 | 0.0022 | -0.0107 | 0.0101 | -0.0004 | 0.0019 |
| rs1013293 | A | -0.0177 | 0.0020 | -0.0252 | 0.0098 | 0.0030 | 0.0017 |
| rs10145154 | T | 0.0288 | 0.0024 | 0.0208 | 0.0120 | 0.0002 | 0.0021 |
| rs10160769 | C | -0.0154 | 0.0024 | -0.0082 | 0.0111 | 0.0125 | 0.0021 |
| rs10169594 | T | -0.0122 | 0.0020 | -0.0117 | 0.0100 | 0.0086 | 0.0018 |
| rs10206166 | T | 0.0168 | 0.0020 | 0.0367 | 0.0099 | -0.0111 | 0.0018 |
| rs10236214 | T | 0.0145 | 0.0021 | -0.0180 | 0.0106 | 0.0015 | 0.0018 |
| rs10259490 | A | -0.0129 | 0.0020 | -0.0088 | 0.0094 | -0.0023 | 0.0017 |
| rs10423928 | A | -0.0342 | 0.0025 | -0.0399 | 0.0125 | 0.0031 | 0.0021 |
| rs1045411 | T | -0.0163 | 0.0022 | -0.0011 | 0.0113 | 0.0032 | 0.0019 |
| rs1048932 | A | -0.0149 | 0.0020 | -0.0022 | 0.0094 | 0.0041 | 0.0017 |
| rs10514944 | A | 0.0247 | 0.0034 | 0.0150 | 0.0148 | -0.0074 | 0.0029 |
| rs10514975 | A | -0.0139 | 0.0020 | -0.0179 | 0.0095 | 0.0012 | 0.0017 |
| rs10515237 | A | -0.0151 | 0.0022 | 0.0008 | 0.0105 | -0.0003 | 0.0019 |
| rs1064213 | A | 0.0146 | 0.0019 | 0.0052 | 0.0096 | -0.0026 | 0.0017 |
| rs10742752 | T | -0.0119 | 0.0020 | 0.0067 | 0.0095 | -0.0026 | 0.0017 |
| rs10756714 | A | 0.0199 | 0.0020 | 0.0072 | 0.0093 | -0.0048 | 0.0017 |
| rs10768994 | T | 0.0125 | 0.0020 | 0.0059 | 0.0094 | -0.0017 | 0.0017 |
| rs10790519 | T | -0.0124 | 0.0020 | -0.0179 | 0.0096 | 0.0041 | 0.0018 |
| rs10818810 | A | 0.0125 | 0.0020 | 0.0248 | 0.0097 | -0.0051 | 0.0018 |
| rs10824218 | A | 0.0141 | 0.0020 | 0.0131 | 0.0100 | 0.0020 | 0.0017 |
| rs10841188 | A | -0.0135 | 0.0022 | -0.0202 | 0.0101 | 0.0010 | 0.0020 |
| rs10842240 | C | 0.0215 | 0.0031 | 0.0006 | 0.0120 | -0.0071 | 0.0027 |
| rs10878946 | T | -0.0148 | 0.0022 | -0.0110 | 0.0103 | -0.0015 | 0.0019 |
| rs10887578 | C | 0.0134 | 0.0020 | -0.0014 | 0.0096 | -0.0019 | 0.0017 |
| rs10899736 | A | -0.0116 | 0.0020 | -0.0109 | 0.0093 | -0.0020 | 0.0017 |
| rs10921743 | A | 0.0138 | 0.0020 | 0.0148 | 0.0097 | -0.0021 | 0.0018 |
| rs10938397 | A | -0.0292 | 0.0020 | -0.0306 | 0.0093 | 0.0066 | 0.0017 |
| rs10962550 | C | 0.0196 | 0.0025 | 0.0235 | 0.0115 | -0.0093 | 0.0022 |
| rs11075489 | T | -0.0115 | 0.0020 | -0.0069 | 0.0092 | 0.0023 | 0.0017 |
| rs11079849 | T | -0.0213 | 0.0021 | -0.0506 | 0.0109 | 0.0074 | 0.0018 |
| rs11084553 | A | 0.0236 | 0.0027 | 0.0239 | 0.0150 | 0.0045 | 0.0024 |
| rs11134679 | A | -0.0177 | 0.0021 | -0.0032 | 0.0104 | 0.0025 | 0.0018 |
| rs11150461 | C | 0.0148 | 0.0022 | 0.0099 | 0.0105 | 0.0017 | 0.0019 |
| rs11161044 | C | 0.0175 | 0.0025 | 0.0336 | 0.0111 | -0.0180 | 0.0021 |
| rs11165493 | A | 0.0143 | 0.0021 | 0.0068 | 0.0101 | 0.0053 | 0.0018 |
| rs11190658 | T | 0.0150 | 0.0024 | 0.0127 | 0.0112 | 0.0002 | 0.0021 |
| rs11223970 | A | -0.0193 | 0.0026 | -0.0017 | 0.0134 | 0.0049 | 0.0023 |
| rs11251352 | A | -0.0117 | 0.0020 | 0.0066 | 0.0094 | 0.0038 | 0.0017 |
| rs11496125 | T | 0.0189 | 0.0020 | 0.0102 | 0.0093 | 0.0025 | 0.0017 |
| rs11505821 | A | -0.0301 | 0.0040 | -0.0355 | 0.0176 | -0.0030 | 0.0036 |
| rs11609297 | T | -0.0172 | 0.0023 | -0.0174 | 0.0122 | 0.0058 | 0.0020 |
| rs11611246 | T | 0.0255 | 0.0024 | 0.0074 | 0.0114 | 0.0079 | 0.0021 |
| rs11615578 | T | 0.0137 | 0.0023 | -0.0068 | 0.0117 | 0.0022 | 0.0020 |
| rs11675464 | A | -0.0117 | 0.0020 | 0.0412 | 0.0096 | -0.0001 | 0.0017 |
| rs11713193 | A | 0.0266 | 0.0019 | 0.0301 | 0.0097 | -0.0225 | 0.0017 |
| rs11757278 | T | 0.0146 | 0.0021 | -0.0071 | 0.0101 | -0.0073 | 0.0018 |
| rs11773362 | T | -0.0125 | 0.0021 | 0.0073 | 0.0097 | 0.0033 | 0.0018 |
| rs11846126 | T | 0.0158 | 0.0020 | 0.0119 | 0.0095 | -0.0037 | 0.0017 |
| rs11874191 | T | -0.0151 | 0.0023 | 0.0109 | 0.0104 | 0.0069 | 0.0020 |
| rs11895606 | T | 0.0131 | 0.0020 | -0.0082 | 0.0099 | 0.0015 | 0.0017 |
| rs11915371 | A | -0.0148 | 0.0024 | -0.0132 | 0.0116 | 0.0102 | 0.0021 |
| rs11945861 | A | -0.0143 | 0.0023 | 0.0047 | 0.0106 | 0.0068 | 0.0020 |
| rs11976018 | A | -0.0228 | 0.0027 | -0.0100 | 0.0115 | 0.0041 | 0.0023 |
| rs12049202 | T | 0.0245 | 0.0024 | 0.0273 | 0.0116 | -0.0067 | 0.0021 |
| rs1218822 | A | 0.0176 | 0.0021 | 0.0061 | 0.0098 | -0.0024 | 0.0018 |
| rs12334877 | A | -0.0152 | 0.0025 | -0.0151 | 0.0111 | 0.0047 | 0.0021 |
| rs12369179 | T | -0.0372 | 0.0034 | -0.0101 | 0.0193 | -0.0143 | 0.0030 |
| rs12427047 | T | -0.0184 | 0.0023 | 0.0065 | 0.0109 | -0.0041 | 0.0020 |
| rs12439798 | T | 0.0139 | 0.0020 | 0.0003 | 0.0094 | -0.0038 | 0.0017 |
| rs12448257 | A | 0.0157 | 0.0024 | 0.0097 | 0.0113 | 0.0074 | 0.0021 |
| rs12512994 | T | -0.0127 | 0.0021 | -0.0067 | 0.0098 | 0.0140 | 0.0018 |
| rs12564848 | T | -0.0229 | 0.0031 | -0.0032 | 0.0136 | -0.0015 | 0.0027 |
| rs12595749 | A | 0.0149 | 0.0020 | -0.0037 | 0.0094 | -0.0022 | 0.0017 |
| rs12608738 | A | 0.0130 | 0.0022 | -0.0028 | 0.0118 | -0.0020 | 0.0019 |
| rs1266874 | A | -0.0147 | 0.0020 | -0.0037 | 0.0097 | 0.0023 | 0.0018 |
| rs12681792 | A | 0.0149 | 0.0025 | -0.0024 | 0.0115 | -0.0034 | 0.0021 |
| rs12705977 | T | 0.0134 | 0.0020 | 0.0045 | 0.0096 | -0.0079 | 0.0017 |
| rs12762034 | T | -0.0262 | 0.0037 | -0.0037 | 0.0160 | 0.0059 | 0.0032 |
| rs12878246 | T | 0.0137 | 0.0023 | 0.0172 | 0.0120 | -0.0017 | 0.0020 |
| rs12906208 | A | -0.0149 | 0.0024 | 0.0089 | 0.0123 | 0.0039 | 0.0021 |
| rs12936083 | A | -0.0137 | 0.0021 | -0.0002 | 0.0099 | 0.0014 | 0.0018 |
| rs12936319 | A | 0.0167 | 0.0020 | 0.0038 | 0.0095 | -0.0072 | 0.0017 |
| rs12949279 | T | 0.0180 | 0.0020 | 0.0179 | 0.0095 | -0.0031 | 0.0017 |
| rs1296328 | A | 0.0190 | 0.0020 | -0.0024 | 0.0099 | -0.0024 | 0.0017 |
| rs12981256 | A | 0.0140 | 0.0020 | 0.0139 | 0.0101 | -0.0021 | 0.0017 |
| rs13034320 | C | 0.0136 | 0.0023 | 0.0252 | 0.0108 | -0.0012 | 0.0020 |
| rs13047416 | C | 0.0143 | 0.0020 | -0.0023 | 0.0096 | 0.0015 | 0.0018 |
| rs13085002 | T | -0.0168 | 0.0027 | -0.0135 | 0.0124 | 0.0009 | 0.0023 |
| rs13107325 | T | 0.0469 | 0.0037 | -0.0067 | 0.0223 | -0.0188 | 0.0034 |
| rs13110266 | A | -0.0119 | 0.0020 | -0.0088 | 0.0093 | 0.0030 | 0.0017 |
| rs13132853 | A | 0.0138 | 0.0021 | -0.0105 | 0.0112 | -0.0096 | 0.0018 |
| rs13147390 | T | -0.0122 | 0.0020 | 0.0219 | 0.0099 | 0.0034 | 0.0018 |
| rs13174863 | A | -0.0210 | 0.0028 | -0.0098 | 0.0141 | -0.0014 | 0.0024 |
| rs13199223 | T | 0.0209 | 0.0028 | 0.0313 | 0.0140 | -0.0076 | 0.0025 |
| rs1320251 | T | -0.0168 | 0.0020 | -0.0188 | 0.0098 | 0.0038 | 0.0017 |
| rs1320903 | A | 0.0229 | 0.0021 | -0.0019 | 0.0101 | -0.0004 | 0.0018 |
| rs13209689 | A | -0.0132 | 0.0021 | -0.0120 | 0.0098 | 0.0056 | 0.0018 |
| rs13209918 | A | -0.0134 | 0.0020 | -0.0079 | 0.0099 | 0.0018 | 0.0018 |
| rs13218383 | C | 0.0135 | 0.0021 | 0.0133 | 0.0096 | -0.0014 | 0.0018 |
| rs13263601 | A | -0.0150 | 0.0020 | -0.0115 | 0.0105 | 0.0056 | 0.0018 |
| rs1327259 | A | 0.0155 | 0.0020 | 0.0041 | 0.0094 | -0.0031 | 0.0017 |
| rs1328448 | A | 0.0127 | 0.0021 | -0.0125 | 0.0100 | -0.0019 | 0.0019 |
| rs13288178 | A | 0.0141 | 0.0020 | 0.0175 | 0.0098 | 0.0012 | 0.0018 |
| rs13299074 | T | 0.0147 | 0.0025 | -0.0059 | 0.0126 | -0.0007 | 0.0022 |
| rs13427822 | A | 0.0178 | 0.0022 | 0.0284 | 0.0115 | -0.0053 | 0.0019 |
| rs1348716 | A | -0.0117 | 0.0020 | 0.0148 | 0.0095 | -0.0127 | 0.0018 |
| rs13665 | C | 0.0160 | 0.0021 | -0.0129 | 0.0100 | -0.0041 | 0.0018 |
| rs1406120 | A | -0.0182 | 0.0031 | -0.0250 | 0.0155 | 0.0123 | 0.0027 |
| rs1412235 | C | 0.0246 | 0.0021 | 0.0231 | 0.0103 | 0.0013 | 0.0018 |
| rs1421085 | T | -0.0767 | 0.0020 | -0.0297 | 0.0096 | -0.0027 | 0.0017 |
| rs1421334 | A | 0.0131 | 0.0020 | 0.0144 | 0.0096 | -0.0112 | 0.0017 |
| rs1431659 | A | 0.0222 | 0.0022 | 0.0010 | 0.0100 | -0.0001 | 0.0019 |
| rs1436344 | C | 0.0145 | 0.0020 | 0.0072 | 0.0093 | -0.0019 | 0.0017 |
| rs1454687 | C | 0.0226 | 0.0019 | 0.0068 | 0.0093 | -0.0072 | 0.0017 |
| rs1460940 | A | 0.0312 | 0.0024 | 0.0310 | 0.0133 | 0.0024 | 0.0022 |
| rs1470153 | A | -0.0138 | 0.0023 | 0.0212 | 0.0102 | 0.0067 | 0.0020 |
| rs1477890 | A | -0.0162 | 0.0020 | 0.0049 | 0.0091 | 0.0018 | 0.0017 |
| rs1503526 | T | -0.0144 | 0.0020 | -0.0056 | 0.0094 | 0.0105 | 0.0017 |
| rs1522569 | T | 0.0167 | 0.0025 | -0.0080 | 0.0125 | -0.0055 | 0.0022 |
| rs1574389 | T | 0.0161 | 0.0024 | 0.0168 | 0.0107 | -0.0031 | 0.0021 |
| rs1579557 | T | 0.0225 | 0.0021 | -0.0048 | 0.0104 | -0.0037 | 0.0019 |
| rs1582931 | A | -0.0142 | 0.0020 | -0.0161 | 0.0095 | -0.0074 | 0.0017 |
| rs1583123 | A | 0.0124 | 0.0020 | -0.0094 | 0.0095 | -0.0038 | 0.0018 |
| rs1681740 | A | 0.0122 | 0.0020 | -0.0040 | 0.0097 | -0.0027 | 0.0017 |
| rs16868443 | C | 0.0129 | 0.0020 | -0.0058 | 0.0098 | 0.0035 | 0.0018 |
| rs16882450 | A | 0.0132 | 0.0021 | 0.0210 | 0.0098 | -0.0057 | 0.0019 |
| rs16916303 | A | 0.0195 | 0.0031 | 0.0100 | 0.0131 | -0.0026 | 0.0027 |
| rs16973520 | T | -0.0138 | 0.0023 | -0.0234 | 0.0102 | 0.0045 | 0.0020 |
| rs16996657 | T | -0.0193 | 0.0029 | 0.0001 | 0.0156 | 0.0012 | 0.0025 |
| rs17014375 | T | -0.0187 | 0.0029 | 0.0075 | 0.0130 | 0.0004 | 0.0025 |
| rs1701827 | A | 0.0145 | 0.0022 | 0.0115 | 0.0108 | -0.0028 | 0.0019 |
| rs17024258 | T | 0.0648 | 0.0061 | -0.0490 | 0.0295 | 0.0039 | 0.0050 |
| rs17056301 | T | -0.0132 | 0.0023 | -0.0131 | 0.0102 | -0.0022 | 0.0019 |
| rs17066856 | T | 0.0392 | 0.0034 | 0.0379 | 0.0155 | -0.0042 | 0.0030 |
| rs1711171 | T | -0.0185 | 0.0023 | -0.0302 | 0.0113 | 0.0029 | 0.0020 |
| rs17132130 | C | -0.0173 | 0.0024 | -0.0218 | 0.0106 | 0.0108 | 0.0020 |
| rs17207196 | T | -0.0228 | 0.0020 | -0.0150 | 0.0111 | 0.0091 | 0.0018 |
| rs17379561 | A | -0.0181 | 0.0028 | -0.0186 | 0.0134 | 0.0190 | 0.0024 |
| rs17399237 | T | 0.0120 | 0.0020 | 0.0211 | 0.0100 | -0.0051 | 0.0017 |
| rs17446299 | C | -0.0157 | 0.0026 | -0.0097 | 0.0135 | -0.0014 | 0.0023 |
| rs17513613 | T | -0.0201 | 0.0021 | 0.0225 | 0.0104 | -0.0010 | 0.0018 |
| rs17636031 | T | -0.0164 | 0.0022 | -0.0026 | 0.0122 | 0.0041 | 0.0019 |
| rs17639546 | A | -0.0240 | 0.0027 | -0.0185 | 0.0131 | 0.0035 | 0.0024 |
| rs17724992 | A | 0.0177 | 0.0022 | 0.0139 | 0.0102 | -0.0060 | 0.0019 |
| rs17776719 | A | -0.0171 | 0.0028 | 0.0228 | 0.0122 | -0.0016 | 0.0024 |
| rs17806379 | T | -0.0267 | 0.0025 | -0.0060 | 0.0123 | 0.0079 | 0.0022 |
| rs17814208 | A | -0.0134 | 0.0023 | 0.0111 | 0.0113 | 0.0018 | 0.0020 |
| rs17827419 | A | -0.0213 | 0.0030 | -0.0080 | 0.0159 | 0.0033 | 0.0026 |
| rs1786141 | T | 0.0136 | 0.0020 | 0.0184 | 0.0097 | -0.0062 | 0.0017 |
| rs1814170 | A | 0.0198 | 0.0033 | -0.0142 | 0.0129 | 0.0040 | 0.0028 |
| rs1861410 | T | -0.0219 | 0.0020 | -0.0067 | 0.0095 | 0.0031 | 0.0017 |
| rs1884897 | A | -0.0216 | 0.0020 | 0.0002 | 0.0096 | -0.0010 | 0.0018 |
| rs1928538 | T | -0.0135 | 0.0020 | -0.0245 | 0.0093 | 0.0024 | 0.0017 |
| rs1951455 | T | -0.0150 | 0.0022 | 0.0236 | 0.0107 | 0.0018 | 0.0019 |
| rs1955540 | T | -0.0154 | 0.0025 | 0.0067 | 0.0114 | 0.0038 | 0.0022 |
| rs197374 | T | 0.0140 | 0.0020 | -0.0112 | 0.0094 | -0.0087 | 0.0018 |
| rs1976053 | C | -0.0186 | 0.0025 | -0.0242 | 0.0110 | 0.0034 | 0.0022 |
| rs2012286 | A | 0.0139 | 0.0020 | 0.0041 | 0.0094 | -0.0044 | 0.0017 |
| rs2051559 | T | -0.0192 | 0.0029 | -0.0111 | 0.0134 | 0.0152 | 0.0025 |
| rs2065418 | T | 0.0184 | 0.0020 | -0.0043 | 0.0097 | 0.0025 | 0.0018 |
| rs2072858 | T | 0.0164 | 0.0021 | -0.0085 | 0.0095 | 0.0045 | 0.0018 |
| rs214247 | T | 0.0136 | 0.0020 | -0.0043 | 0.0100 | -0.0035 | 0.0018 |
| rs2143253 | A | -0.0201 | 0.0030 | -0.0118 | 0.0125 | 0.0125 | 0.0026 |
| rs215634 | A | 0.0158 | 0.0020 | 0.0193 | 0.0094 | -0.0095 | 0.0018 |
| rs2161796 | T | 0.0141 | 0.0020 | -0.0018 | 0.0096 | 0.0005 | 0.0018 |
| rs217671 | A | -0.0169 | 0.0022 | -0.0082 | 0.0102 | 0.0070 | 0.0019 |
| rs2181375 | A | -0.0201 | 0.0020 | -0.0052 | 0.0093 | 0.0021 | 0.0017 |
| rs2183947 | A | -0.0162 | 0.0023 | 0.0019 | 0.0115 | 0.0106 | 0.0020 |
| rs2253310 | C | -0.0169 | 0.0020 | -0.0009 | 0.0095 | -0.0093 | 0.0018 |
| rs2271189 | A | -0.0161 | 0.0020 | -0.0228 | 0.0100 | 0.0121 | 0.0018 |
| rs2281819 | A | -0.0162 | 0.0023 | -0.0100 | 0.0112 | 0.0065 | 0.0020 |
| rs2284746 | C | 0.0112 | 0.0019 | 0.0189 | 0.0094 | -0.0026 | 0.0018 |
| rs2306540 | T | 0.0137 | 0.0021 | 0.0023 | 0.0108 | -0.0031 | 0.0018 |
| rs2307111 | T | 0.0276 | 0.0020 | 0.0075 | 0.0095 | -0.0052 | 0.0017 |
| rs2414128 | T | 0.0131 | 0.0020 | -0.0017 | 0.0097 | 0.0026 | 0.0017 |
| rs241472 | T | -0.0190 | 0.0021 | -0.0063 | 0.0097 | 0.0021 | 0.0018 |
| rs2425847 | A | -0.0126 | 0.0020 | -0.0038 | 0.0096 | 0.0031 | 0.0017 |
| rs2433733 | A | -0.0169 | 0.0021 | 0.0086 | 0.0101 | 0.0057 | 0.0018 |
| rs244415 | A | -0.0226 | 0.0020 | -0.0265 | 0.0096 | 0.0011 | 0.0017 |
| rs2444253 | C | 0.0138 | 0.0020 | 0.0033 | 0.0095 | -0.0038 | 0.0017 |
| rs2450444 | A | -0.0127 | 0.0021 | -0.0026 | 0.0101 | 0.0008 | 0.0018 |
| rs2472297 | T | 0.0147 | 0.0022 | -0.0030 | 0.0129 | -0.0002 | 0.0020 |
| rs2479958 | A | 0.0158 | 0.0020 | 0.0155 | 0.0097 | -0.0021 | 0.0017 |
| rs248139 | A | 0.0151 | 0.0025 | -0.0116 | 0.0115 | 0.0021 | 0.0021 |
| rs2591175 | A | -0.0144 | 0.0025 | 0.0017 | 0.0117 | -0.0013 | 0.0022 |
| rs2605570 | A | 0.0151 | 0.0026 | -0.0082 | 0.0123 | -0.0019 | 0.0023 |
| rs2616143 | A | -0.0133 | 0.0021 | 0.0039 | 0.0106 | 0.0005 | 0.0018 |
| rs2644112 | T | -0.0241 | 0.0021 | -0.0389 | 0.0100 | 0.0109 | 0.0018 |
| rs2693826 | A | -0.0136 | 0.0020 | -0.0231 | 0.0092 | 0.0022 | 0.0017 |
| rs2694047 | A | -0.0211 | 0.0023 | -0.0088 | 0.0107 | 0.0013 | 0.0020 |
| rs2733287 | A | -0.0155 | 0.0020 | -0.0210 | 0.0091 | -0.0067 | 0.0017 |
| rs273512 | T | 0.0171 | 0.0020 | 0.0011 | 0.0098 | -0.0054 | 0.0017 |
| rs2781668 | T | 0.0159 | 0.0026 | 0.0217 | 0.0122 | 0.0048 | 0.0023 |
| rs2791643 | T | -0.0143 | 0.0023 | 0.0033 | 0.0112 | 0.0040 | 0.0020 |
| rs2803316 | A | -0.0148 | 0.0020 | 0.0156 | 0.0103 | -0.0062 | 0.0017 |
| rs2814942 | A | 0.0269 | 0.0021 | 0.0233 | 0.0100 | -0.0038 | 0.0019 |
| rs28350 | A | 0.0186 | 0.0026 | 0.0025 | 0.0124 | 0.0022 | 0.0022 |
| rs2836968 | A | 0.0122 | 0.0020 | 0.0054 | 0.0102 | -0.0052 | 0.0018 |
| rs2861683 | A | 0.0171 | 0.0020 | -0.0028 | 0.0095 | 0.0003 | 0.0017 |
| rs2865384 | A | -0.0121 | 0.0020 | -0.0104 | 0.0095 | 0.0047 | 0.0018 |
| rs2875762 | C | 0.0151 | 0.0023 | -0.0035 | 0.0109 | -0.0012 | 0.0020 |
| rs2906132 | T | 0.0119 | 0.0020 | 0.0102 | 0.0100 | -0.0016 | 0.0017 |
| rs2920502 | C | -0.0147 | 0.0021 | -0.0058 | 0.0100 | 0.0030 | 0.0018 |
| rs294704 | T | -0.0146 | 0.0022 | -0.0157 | 0.0108 | -0.0007 | 0.0019 |
| rs2952862 | T | 0.0143 | 0.0021 | 0.0211 | 0.0095 | -0.0043 | 0.0018 |
| rs326896 | T | -0.0123 | 0.0020 | -0.0138 | 0.0093 | 0.0038 | 0.0017 |
| rs329651 | T | 0.0169 | 0.0025 | -0.0006 | 0.0141 | -0.0075 | 0.0022 |
| rs355748 | T | 0.0162 | 0.0020 | 0.0128 | 0.0093 | -0.0037 | 0.0018 |
| rs3730071 | A | -0.0340 | 0.0057 | 0.0064 | 0.0369 | -0.0043 | 0.0052 |
| rs3743950 | A | -0.0119 | 0.0020 | 0.0048 | 0.0105 | 0.0013 | 0.0018 |
| rs3754963 | A | 0.0139 | 0.0022 | 0.0187 | 0.0108 | -0.0106 | 0.0019 |
| rs3762444 | T | -0.0144 | 0.0020 | -0.0119 | 0.0097 | 0.0017 | 0.0017 |
| rs3770451 | T | 0.0144 | 0.0022 | -0.0077 | 0.0109 | -0.0104 | 0.0019 |
| rs3784710 | T | 0.0287 | 0.0023 | 0.0135 | 0.0107 | -0.0005 | 0.0021 |
| rs3803286 | A | 0.0193 | 0.0021 | 0.0212 | 0.0098 | -0.0070 | 0.0018 |
| rs3806114 | A | -0.0125 | 0.0021 | -0.0092 | 0.0108 | 0.0000 | 0.0018 |
| rs3806572 | A | -0.0154 | 0.0021 | 0.0200 | 0.0110 | -0.0004 | 0.0019 |
| rs3814883 | T | 0.0249 | 0.0020 | -0.0097 | 0.0099 | -0.0005 | 0.0017 |
| rs3821709 | T | -0.0338 | 0.0039 | -0.0133 | 0.0165 | 0.0128 | 0.0035 |
| rs38314 | A | -0.0116 | 0.0020 | 0.0015 | 0.0097 | -0.0057 | 0.0017 |
| rs394608 | T | -0.0177 | 0.0020 | -0.0170 | 0.0092 | 0.0044 | 0.0017 |
| rs40067 | A | -0.0272 | 0.0026 | -0.0389 | 0.0115 | 0.0011 | 0.0023 |
| rs4072917 | A | 0.0118 | 0.0020 | 0.0108 | 0.0098 | -0.0024 | 0.0017 |
| rs4148155 | A | 0.0203 | 0.0031 | -0.0238 | 0.0149 | -0.0004 | 0.0027 |
| rs4148866 | T | 0.0124 | 0.0020 | -0.0028 | 0.0100 | -0.0072 | 0.0017 |
| rs4237195 | A | 0.0177 | 0.0021 | 0.0089 | 0.0098 | -0.0046 | 0.0018 |
| rs4237643 | T | 0.0237 | 0.0021 | 0.0282 | 0.0103 | -0.0024 | 0.0019 |
| rs4240673 | T | 0.0201 | 0.0020 | 0.0098 | 0.0100 | -0.0026 | 0.0017 |
| rs4284600 | T | -0.0115 | 0.0020 | -0.0005 | 0.0096 | 0.0019 | 0.0017 |
| rs429343 | A | 0.0172 | 0.0020 | 0.0107 | 0.0097 | -0.0006 | 0.0017 |
| rs4307239 | A | -0.0120 | 0.0020 | -0.0017 | 0.0094 | 0.0000 | 0.0017 |
| rs4343164 | T | 0.0165 | 0.0022 | 0.0031 | 0.0107 | -0.0066 | 0.0019 |
| rs4477562 | T | 0.0317 | 0.0030 | 0.0101 | 0.0130 | -0.0001 | 0.0026 |
| rs4482463 | A | -0.0346 | 0.0037 | 0.0028 | 0.0161 | -0.0043 | 0.0032 |
| rs4490752 | A | 0.0172 | 0.0020 | -0.0050 | 0.0097 | 0.0000 | 0.0018 |
| rs4516268 | A | -0.0241 | 0.0025 | -0.0073 | 0.0129 | 0.0096 | 0.0022 |
| rs4639527 | A | -0.0174 | 0.0021 | -0.0018 | 0.0102 | -0.0012 | 0.0018 |
| rs4652839 | C | -0.0132 | 0.0021 | -0.0005 | 0.0099 | 0.0027 | 0.0018 |
| rs4654831 | T | 0.0188 | 0.0026 | 0.0091 | 0.0114 | -0.0021 | 0.0023 |
| rs4658403 | T | -0.0186 | 0.0026 | -0.0098 | 0.0114 | 0.0104 | 0.0023 |
| rs4660443 | T | 0.0182 | 0.0024 | 0.0217 | 0.0118 | -0.0010 | 0.0021 |
| rs4670169 | A | -0.0120 | 0.0020 | -0.0163 | 0.0096 | -0.0056 | 0.0018 |
| rs4722398 | T | 0.0178 | 0.0028 | 0.0047 | 0.0139 | 0.0040 | 0.0026 |
| rs4762369 | A | 0.0151 | 0.0025 | 0.0255 | 0.0120 | 0.0018 | 0.0022 |
| rs4764949 | A | 0.0196 | 0.0021 | 0.0072 | 0.0096 | 0.0010 | 0.0018 |
| rs4851716 | A | 0.0162 | 0.0020 | 0.0117 | 0.0101 | 0.0018 | 0.0018 |
| rs4911382 | T | 0.0147 | 0.0020 | 0.0039 | 0.0096 | -0.0034 | 0.0017 |
| rs4936175 | T | -0.0124 | 0.0020 | -0.0005 | 0.0100 | 0.0036 | 0.0017 |
| rs4942924 | T | 0.0158 | 0.0020 | 0.0338 | 0.0096 | -0.0022 | 0.0017 |
| rs4952843 | A | 0.0151 | 0.0020 | 0.0165 | 0.0099 | 0.0006 | 0.0017 |
| rs4969387 | C | -0.0146 | 0.0022 | -0.0061 | 0.0123 | -0.0027 | 0.0020 |
| rs500764 | T | 0.0164 | 0.0020 | 0.0257 | 0.0093 | 0.0001 | 0.0017 |
| rs538656 | T | 0.0544 | 0.0023 | 0.0562 | 0.0105 | 0.0056 | 0.0020 |
| rs538740 | T | -0.0145 | 0.0024 | -0.0028 | 0.0114 | 0.0074 | 0.0021 |
| rs5396 | T | -0.0175 | 0.0022 | 0.0138 | 0.0103 | 0.0088 | 0.0019 |
| rs543874 | A | -0.0466 | 0.0024 | -0.0076 | 0.0117 | 0.0047 | 0.0021 |
| rs559231 | T | 0.0142 | 0.0020 | 0.0117 | 0.0094 | 0.0015 | 0.0017 |
| rs577525 | T | -0.0187 | 0.0020 | -0.0179 | 0.0093 | 0.0066 | 0.0017 |
| rs592483 | T | -0.0152 | 0.0020 | 0.0231 | 0.0102 | 0.0029 | 0.0017 |
| rs6011457 | A | -0.0116 | 0.0020 | -0.0054 | 0.0094 | -0.0073 | 0.0017 |
| rs6023649 | A | 0.0153 | 0.0023 | 0.0012 | 0.0109 | -0.0074 | 0.0019 |
| rs6050446 | A | -0.0375 | 0.0056 | -0.0342 | 0.0320 | 0.0121 | 0.0048 |
| rs6098868 | A | -0.0117 | 0.0020 | 0.0015 | 0.0092 | -0.0010 | 0.0017 |
| rs6265 | T | -0.0405 | 0.0025 | -0.0304 | 0.0115 | 0.0097 | 0.0022 |
| rs6443750 | T | -0.0152 | 0.0025 | -0.0190 | 0.0149 | 0.0067 | 0.0022 |
| rs6448587 | A | 0.0196 | 0.0026 | 0.0108 | 0.0110 | -0.0004 | 0.0022 |
| rs6479905 | A | -0.0126 | 0.0020 | 0.0066 | 0.0092 | 0.0143 | 0.0017 |
| rs6502482 | C | -0.0133 | 0.0020 | -0.0069 | 0.0094 | -0.0041 | 0.0017 |
| rs6542924 | A | -0.0174 | 0.0021 | -0.0163 | 0.0110 | 0.0174 | 0.0018 |
| rs6551304 | A | -0.0186 | 0.0026 | -0.0063 | 0.0133 | 0.0047 | 0.0023 |
| rs6577584 | T | -0.0131 | 0.0021 | -0.0025 | 0.0097 | 0.0023 | 0.0018 |
| rs6587552 | A | 0.0175 | 0.0023 | 0.0128 | 0.0108 | -0.0076 | 0.0020 |
| rs6604866 | C | -0.0161 | 0.0020 | 0.0032 | 0.0094 | 0.0003 | 0.0017 |
| rs6674079 | A | -0.0127 | 0.0020 | 0.0105 | 0.0095 | 0.0055 | 0.0018 |
| rs6682 | A | -0.0121 | 0.0020 | -0.0043 | 0.0101 | 0.0038 | 0.0017 |
| rs671722 | A | -0.0126 | 0.0020 | -0.0064 | 0.0105 | 0.0093 | 0.0017 |
| rs6725931 | T | 0.0187 | 0.0027 | 0.0257 | 0.0134 | -0.0115 | 0.0024 |
| rs6728037 | A | 0.0198 | 0.0020 | 0.0081 | 0.0098 | -0.0046 | 0.0017 |
| rs6731688 | A | -0.0564 | 0.0026 | -0.0372 | 0.0124 | 0.0047 | 0.0023 |
| rs6752378 | A | 0.0332 | 0.0019 | 0.0190 | 0.0093 | 0.0012 | 0.0017 |
| rs6764533 | A | 0.0141 | 0.0020 | 0.0122 | 0.0101 | -0.0073 | 0.0018 |
| rs6831020 | A | -0.0161 | 0.0021 | -0.0056 | 0.0111 | 0.0027 | 0.0019 |
| rs6870983 | T | -0.0219 | 0.0024 | -0.0186 | 0.0110 | 0.0141 | 0.0021 |
| rs6905544 | A | -0.0164 | 0.0020 | 0.0051 | 0.0097 | 0.0233 | 0.0017 |
| rs698147 | A | 0.0130 | 0.0020 | 0.0022 | 0.0092 | -0.0045 | 0.0017 |
| rs7038943 | T | 0.0143 | 0.0021 | 0.0125 | 0.0096 | -0.0075 | 0.0018 |
| rs705158 | A | 0.0162 | 0.0023 | 0.0098 | 0.0103 | 0.0007 | 0.0020 |
| rs705217 | T | 0.0123 | 0.0020 | -0.0008 | 0.0096 | -0.0040 | 0.0018 |
| rs7084454 | A | 0.0210 | 0.0021 | 0.0035 | 0.0105 | -0.0068 | 0.0018 |
| rs7096307 | T | -0.0117 | 0.0020 | 0.0105 | 0.0098 | 0.0024 | 0.0017 |
| rs7102454 | T | -0.0176 | 0.0020 | 0.0009 | 0.0104 | 0.0071 | 0.0018 |
| rs7124681 | A | 0.0269 | 0.0020 | -0.0007 | 0.0096 | 0.0002 | 0.0017 |
| rs7132908 | A | 0.0296 | 0.0020 | 0.0107 | 0.0096 | -0.0005 | 0.0018 |
| rs7158822 | T | -0.0162 | 0.0021 | -0.0253 | 0.0097 | 0.0034 | 0.0018 |
| rs7181498 | T | 0.0160 | 0.0021 | 0.0041 | 0.0097 | -0.0052 | 0.0018 |
| rs7196720 | T | 0.0131 | 0.0020 | -0.0050 | 0.0095 | -0.0045 | 0.0017 |
| rs7258722 | A | -0.0200 | 0.0020 | -0.0139 | 0.0099 | 0.0015 | 0.0017 |
| rs725959 | T | -0.0133 | 0.0020 | -0.0047 | 0.0096 | 0.0054 | 0.0017 |
| rs7313220 | A | 0.0129 | 0.0020 | -0.0057 | 0.0092 | 0.0020 | 0.0017 |
| rs7334078 | T | 0.0125 | 0.0022 | -0.0024 | 0.0104 | -0.0103 | 0.0019 |
| rs733758 | T | -0.0313 | 0.0045 | 0.0091 | 0.0236 | -0.0068 | 0.0039 |
| rs738140 | A | 0.0147 | 0.0021 | 0.0037 | 0.0108 | -0.0033 | 0.0018 |
| rs741959 | A | 0.0173 | 0.0020 | 0.0145 | 0.0096 | -0.0001 | 0.0017 |
| rs7498665 | A | -0.0256 | 0.0020 | -0.0190 | 0.0101 | 0.0166 | 0.0017 |
| rs7518221 | T | 0.0126 | 0.0020 | -0.0031 | 0.0098 | 0.0003 | 0.0018 |
| rs7519259 | A | 0.0137 | 0.0020 | 0.0196 | 0.0094 | -0.0064 | 0.0017 |
| rs754635 | C | -0.0198 | 0.0031 | -0.0058 | 0.0134 | -0.0032 | 0.0027 |
| rs7557796 | T | 0.0171 | 0.0020 | 0.0250 | 0.0099 | -0.0030 | 0.0018 |
| rs756717 | A | -0.0158 | 0.0020 | -0.0242 | 0.0099 | -0.0006 | 0.0017 |
| rs7568228 | C | -0.0116 | 0.0019 | -0.0024 | 0.0097 | 0.0050 | 0.0017 |
| rs7619139 | A | 0.0146 | 0.0020 | 0.0086 | 0.0094 | 0.0007 | 0.0017 |
| rs7626079 | T | 0.0121 | 0.0021 | 0.0021 | 0.0096 | 0.0052 | 0.0018 |
| rs7637852 | A | 0.0143 | 0.0021 | 0.0150 | 0.0103 | -0.0013 | 0.0019 |
| rs7640424 | T | -0.0128 | 0.0021 | 0.0023 | 0.0108 | 0.0107 | 0.0018 |
| rs7646113 | T | -0.0125 | 0.0020 | 0.0101 | 0.0095 | 0.0010 | 0.0018 |
| rs765874 | A | -0.0125 | 0.0020 | 0.0152 | 0.0093 | 0.0022 | 0.0017 |
| rs7683836 | A | -0.0135 | 0.0020 | -0.0148 | 0.0094 | 0.0003 | 0.0017 |
| rs769449 | A | -0.0235 | 0.0030 | 0.0896 | 0.0165 | 0.0067 | 0.0026 |
| rs7707981 | A | 0.0142 | 0.0021 | -0.0012 | 0.0106 | 0.0009 | 0.0018 |
| rs7715256 | T | -0.0165 | 0.0020 | -0.0024 | 0.0096 | -0.0007 | 0.0017 |
| rs7733438 | T | -0.0365 | 0.0029 | -0.0257 | 0.0129 | -0.0173 | 0.0025 |
| rs774246 | A | -0.0169 | 0.0028 | -0.0088 | 0.0143 | 0.0012 | 0.0024 |
| rs7747583 | A | -0.0136 | 0.0022 | 0.0047 | 0.0096 | 0.0074 | 0.0019 |
| rs775403 | T | 0.0117 | 0.0020 | 0.0045 | 0.0094 | -0.0044 | 0.0017 |
| rs7792703 | A | -0.0166 | 0.0024 | -0.0072 | 0.0111 | 0.0010 | 0.0020 |
| rs7869771 | A | 0.0156 | 0.0022 | 0.0001 | 0.0102 | -0.0047 | 0.0019 |
| rs7871866 | C | 0.0177 | 0.0027 | 0.0226 | 0.0128 | -0.0062 | 0.0024 |
| rs7893571 | T | 0.0146 | 0.0021 | -0.0068 | 0.0102 | 0.0024 | 0.0018 |
| rs7903146 | T | -0.0157 | 0.0022 | 0.0331 | 0.0104 | -0.0021 | 0.0019 |
| rs7925214 | T | 0.0157 | 0.0020 | 0.0100 | 0.0095 | -0.0046 | 0.0017 |
| rs799449 | T | 0.0141 | 0.0020 | 0.0175 | 0.0097 | -0.0070 | 0.0017 |
| rs8008285 | T | -0.0175 | 0.0028 | 0.0007 | 0.0135 | -0.0046 | 0.0024 |
| rs8015400 | A | 0.0226 | 0.0021 | 0.0092 | 0.0099 | -0.0022 | 0.0018 |
| rs8089514 | A | 0.0133 | 0.0021 | 0.0234 | 0.0103 | -0.0038 | 0.0018 |
| rs8181823 | A | -0.0139 | 0.0023 | -0.0058 | 0.0111 | 0.0061 | 0.0020 |
| rs879620 | T | 0.0227 | 0.0020 | -0.0029 | 0.0097 | -0.0008 | 0.0018 |
| rs890156 | A | 0.0125 | 0.0020 | 0.0066 | 0.0092 | 0.0023 | 0.0017 |
| rs891387 | T | 0.0205 | 0.0020 | 0.0085 | 0.0094 | -0.0137 | 0.0017 |
| rs895330 | C | 0.0205 | 0.0025 | 0.0140 | 0.0148 | -0.0034 | 0.0022 |
| rs900144 | T | 0.0174 | 0.0020 | 0.0374 | 0.0095 | 0.0015 | 0.0017 |
| rs902503 | A | 0.0130 | 0.0020 | 0.0020 | 0.0095 | -0.0013 | 0.0017 |
| rs903959 | A | 0.0116 | 0.0020 | -0.0005 | 0.0100 | -0.0157 | 0.0017 |
| rs912690 | C | -0.0152 | 0.0020 | 0.0068 | 0.0092 | 0.0030 | 0.0017 |
| rs9294260 | A | 0.0158 | 0.0020 | -0.0019 | 0.0094 | -0.0054 | 0.0017 |
| rs9296389 | C | 0.0122 | 0.0020 | 0.0051 | 0.0093 | -0.0012 | 0.0017 |
| rs930295 | A | 0.0214 | 0.0026 | -0.0010 | 0.0132 | 0.0000 | 0.0023 |
| rs9304665 | A | 0.0225 | 0.0023 | 0.0332 | 0.0106 | -0.0003 | 0.0020 |
| rs942065 | A | 0.0204 | 0.0020 | 0.0043 | 0.0098 | -0.0071 | 0.0018 |
| rs9446149 | T | 0.0145 | 0.0024 | 0.0137 | 0.0115 | 0.0017 | 0.0021 |
| rs9478496 | T | -0.0173 | 0.0026 | -0.0009 | 0.0131 | 0.0013 | 0.0023 |
| rs9496567 | A | 0.0163 | 0.0023 | -0.0041 | 0.0119 | 0.0008 | 0.0020 |
| rs9515448 | A | -0.0141 | 0.0020 | -0.0051 | 0.0095 | 0.0066 | 0.0017 |
| rs9530843 | A | 0.0134 | 0.0020 | -0.0052 | 0.0097 | -0.0027 | 0.0017 |
| rs9538162 | T | 0.0168 | 0.0020 | -0.0070 | 0.0094 | -0.0018 | 0.0017 |
| rs9571687 | A | -0.0126 | 0.0021 | -0.0013 | 0.0099 | -0.0016 | 0.0018 |
| rs960923 | A | -0.0210 | 0.0020 | -0.0004 | 0.0093 | -0.0029 | 0.0017 |
| rs9630985 | A | -0.0173 | 0.0021 | -0.0205 | 0.0102 | 0.0058 | 0.0018 |
| rs9650755 | A | -0.0159 | 0.0022 | 0.0023 | 0.0101 | 0.0069 | 0.0019 |
| rs9688431 | T | 0.0242 | 0.0042 | -0.0041 | 0.0182 | 0.0000 | 0.0036 |
| rs9816226 | A | -0.0292 | 0.0025 | 0.0025 | 0.0123 | -0.0078 | 0.0022 |
| rs9827823 | T | 0.0219 | 0.0027 | -0.0149 | 0.0128 | -0.0017 | 0.0024 |
| rs987237 | A | -0.0395 | 0.0025 | -0.0252 | 0.0116 | 0.0033 | 0.0022 |
| rs9876664 | T | -0.0181 | 0.0020 | -0.0012 | 0.0095 | 0.0139 | 0.0018 |
| rs9926784 | T | 0.0262 | 0.0025 | -0.0034 | 0.0113 | 0.0032 | 0.0022 |
| rs9927848 | A | -0.0129 | 0.0022 | 0.0148 | 0.0103 | -0.0022 | 0.0019 |
| rs9951619 | T | -0.0160 | 0.0024 | -0.0021 | 0.0107 | -0.0052 | 0.0020 |
| rs9958909 | T | -0.0214 | 0.0029 | 0.0045 | 0.0143 | 0.0027 | 0.0024 |

#### Supplementary table 8: Genetic association estimates for the two-sample regression-based MVMR analysis estimating the effect of BMI on MI, adjusted for education. ea=effect allele; gx=BMI; gy=MI; gz=education; se=standard error

| SNP | ea | gx | gx_se | gy | gy_se | gz | gz_se |
| --- | --- | --- | --- | --- | --- | --- | --- |
| rs10083803 | T | -0.0139 | 0.0022 | -0.0168 | 0.0119 | -0.0020 | 0.0019 |
| rs10124645 | A | 0.0124 | 0.0020 | 0.0228 | 0.0111 | -0.0041 | 0.0017 |
| rs10128597 | A | -0.0186 | 0.0022 | -0.0044 | 0.0111 | -0.0004 | 0.0019 |
| rs1013293 | A | -0.0177 | 0.0020 | -0.0256 | 0.0108 | 0.0030 | 0.0017 |
| rs10145154 | T | 0.0288 | 0.0024 | 0.0221 | 0.0132 | 0.0002 | 0.0021 |
| rs10160769 | C | -0.0154 | 0.0024 | -0.0154 | 0.0125 | 0.0125 | 0.0021 |
| rs10169594 | T | -0.0122 | 0.0020 | -0.0208 | 0.0110 | 0.0086 | 0.0018 |
| rs10206166 | T | 0.0168 | 0.0020 | 0.0366 | 0.0107 | -0.0111 | 0.0018 |
| rs10236214 | T | 0.0145 | 0.0021 | -0.0056 | 0.0117 | 0.0015 | 0.0018 |
| rs10259490 | A | -0.0129 | 0.0020 | 0.0117 | 0.0104 | -0.0023 | 0.0017 |
| rs10423928 | A | -0.0342 | 0.0025 | -0.0355 | 0.0141 | 0.0031 | 0.0021 |
| rs1045411 | T | -0.0163 | 0.0022 | -0.0081 | 0.0126 | 0.0032 | 0.0019 |
| rs1048932 | A | -0.0149 | 0.0020 | -0.0015 | 0.0104 | 0.0041 | 0.0017 |
| rs10514944 | A | 0.0247 | 0.0034 | 0.0038 | 0.0164 | -0.0074 | 0.0029 |
| rs10514975 | A | -0.0139 | 0.0020 | -0.0210 | 0.0105 | 0.0012 | 0.0017 |
| rs10515237 | A | -0.0151 | 0.0022 | -0.0015 | 0.0115 | -0.0003 | 0.0019 |
| rs1064213 | A | 0.0146 | 0.0019 | 0.0109 | 0.0103 | -0.0026 | 0.0017 |
| rs10742752 | T | -0.0119 | 0.0020 | 0.0198 | 0.0107 | -0.0026 | 0.0017 |
| rs10756714 | A | 0.0199 | 0.0020 | 0.0108 | 0.0103 | -0.0048 | 0.0017 |
| rs10768994 | T | 0.0125 | 0.0020 | 0.0119 | 0.0104 | -0.0017 | 0.0017 |
| rs10790519 | T | -0.0124 | 0.0020 | -0.0135 | 0.0107 | 0.0041 | 0.0018 |
| rs10818810 | A | 0.0125 | 0.0020 | 0.0218 | 0.0110 | -0.0051 | 0.0018 |
| rs10824218 | A | 0.0141 | 0.0020 | 0.0078 | 0.0112 | 0.0020 | 0.0017 |
| rs10841188 | A | -0.0135 | 0.0022 | -0.0157 | 0.0113 | 0.0010 | 0.0020 |
| rs10842240 | C | 0.0215 | 0.0031 | 0.0101 | 0.0133 | -0.0071 | 0.0027 |
| rs10878946 | T | -0.0148 | 0.0022 | -0.0175 | 0.0114 | -0.0015 | 0.0019 |
| rs10887578 | C | 0.0134 | 0.0020 | -0.0042 | 0.0108 | -0.0019 | 0.0017 |
| rs10899736 | A | -0.0116 | 0.0020 | -0.0066 | 0.0104 | -0.0020 | 0.0017 |
| rs10921743 | A | 0.0138 | 0.0020 | 0.0065 | 0.0107 | -0.0021 | 0.0018 |
| rs10938397 | A | -0.0292 | 0.0020 | -0.0225 | 0.0103 | 0.0066 | 0.0017 |
| rs10962550 | C | 0.0196 | 0.0025 | 0.0226 | 0.0128 | -0.0093 | 0.0022 |
| rs11075489 | T | -0.0115 | 0.0020 | -0.0016 | 0.0102 | 0.0023 | 0.0017 |
| rs11079849 | T | -0.0213 | 0.0021 | -0.0505 | 0.0119 | 0.0074 | 0.0018 |
| rs11084553 | A | 0.0236 | 0.0027 | 0.0174 | 0.0165 | 0.0045 | 0.0024 |
| rs11134679 | A | -0.0177 | 0.0021 | -0.0007 | 0.0118 | 0.0025 | 0.0018 |
| rs11150461 | C | 0.0148 | 0.0022 | 0.0064 | 0.0119 | 0.0017 | 0.0019 |
| rs11161044 | C | 0.0175 | 0.0025 | 0.0256 | 0.0123 | -0.0180 | 0.0021 |
| rs11165493 | A | 0.0143 | 0.0021 | 0.0135 | 0.0111 | 0.0053 | 0.0018 |
| rs11190658 | T | 0.0150 | 0.0024 | 0.0044 | 0.0125 | 0.0002 | 0.0021 |
| rs11223970 | A | -0.0193 | 0.0026 | -0.0082 | 0.0147 | 0.0049 | 0.0023 |
| rs11251352 | A | -0.0117 | 0.0020 | 0.0056 | 0.0105 | 0.0038 | 0.0017 |
| rs11496125 | T | 0.0189 | 0.0020 | -0.0077 | 0.0105 | 0.0025 | 0.0017 |
| rs11505821 | A | -0.0301 | 0.0040 | -0.0338 | 0.0197 | -0.0030 | 0.0036 |
| rs11609297 | T | -0.0172 | 0.0023 | -0.0129 | 0.0134 | 0.0058 | 0.0020 |
| rs11611246 | T | 0.0255 | 0.0024 | 0.0095 | 0.0128 | 0.0079 | 0.0021 |
| rs11615578 | T | 0.0137 | 0.0023 | -0.0052 | 0.0130 | 0.0022 | 0.0020 |
| rs11675464 | A | -0.0117 | 0.0020 | 0.0351 | 0.0103 | -0.0001 | 0.0017 |
| rs11713193 | A | 0.0266 | 0.0019 | 0.0270 | 0.0108 | -0.0225 | 0.0017 |
| rs11757278 | T | 0.0146 | 0.0021 | -0.0073 | 0.0113 | -0.0073 | 0.0018 |
| rs11773362 | T | -0.0125 | 0.0021 | 0.0058 | 0.0107 | 0.0033 | 0.0018 |
| rs11846126 | T | 0.0158 | 0.0020 | 0.0098 | 0.0105 | -0.0037 | 0.0017 |
| rs11874191 | T | -0.0151 | 0.0023 | 0.0002 | 0.0116 | 0.0069 | 0.0020 |
| rs11895606 | T | 0.0131 | 0.0020 | 0.0027 | 0.0105 | 0.0015 | 0.0017 |
| rs11915371 | A | -0.0148 | 0.0024 | -0.0169 | 0.0128 | 0.0102 | 0.0021 |
| rs11945861 | A | -0.0143 | 0.0023 | 0.0100 | 0.0117 | 0.0068 | 0.0020 |
| rs11976018 | A | -0.0228 | 0.0027 | -0.0102 | 0.0129 | 0.0041 | 0.0023 |
| rs12049202 | T | 0.0245 | 0.0024 | 0.0270 | 0.0131 | -0.0067 | 0.0021 |
| rs1218822 | A | 0.0176 | 0.0021 | 0.0063 | 0.0110 | -0.0024 | 0.0018 |
| rs12334877 | A | -0.0152 | 0.0025 | -0.0214 | 0.0123 | 0.0047 | 0.0021 |
| rs12369179 | T | -0.0372 | 0.0034 | 0.0076 | 0.0219 | -0.0143 | 0.0030 |
| rs12427047 | T | -0.0184 | 0.0023 | -0.0007 | 0.0122 | -0.0041 | 0.0020 |
| rs12439798 | T | 0.0139 | 0.0020 | -0.0057 | 0.0104 | -0.0038 | 0.0017 |
| rs12448257 | A | 0.0157 | 0.0024 | 0.0073 | 0.0126 | 0.0074 | 0.0021 |
| rs12512994 | T | -0.0127 | 0.0021 | -0.0116 | 0.0110 | 0.0140 | 0.0018 |
| rs12564848 | T | -0.0229 | 0.0031 | 0.0060 | 0.0152 | -0.0015 | 0.0027 |
| rs12595749 | A | 0.0149 | 0.0020 | -0.0141 | 0.0105 | -0.0022 | 0.0017 |
| rs12608738 | A | 0.0130 | 0.0022 | -0.0041 | 0.0130 | -0.0020 | 0.0019 |
| rs1266874 | A | -0.0147 | 0.0020 | -0.0098 | 0.0106 | 0.0023 | 0.0018 |
| rs12681792 | A | 0.0149 | 0.0025 | -0.0010 | 0.0128 | -0.0034 | 0.0021 |
| rs12705977 | T | 0.0134 | 0.0020 | 0.0114 | 0.0106 | -0.0079 | 0.0017 |
| rs12762034 | T | -0.0262 | 0.0037 | -0.0095 | 0.0178 | 0.0059 | 0.0032 |
| rs12878246 | T | 0.0137 | 0.0023 | 0.0142 | 0.0132 | -0.0017 | 0.0020 |
| rs12906208 | A | -0.0149 | 0.0024 | 0.0095 | 0.0139 | 0.0039 | 0.0021 |
| rs12936083 | A | -0.0137 | 0.0021 | -0.0114 | 0.0110 | 0.0014 | 0.0018 |
| rs12936319 | A | 0.0167 | 0.0020 | -0.0004 | 0.0106 | -0.0072 | 0.0017 |
| rs12949279 | T | 0.0180 | 0.0020 | 0.0129 | 0.0106 | -0.0031 | 0.0017 |
| rs1296328 | A | 0.0190 | 0.0020 | 0.0020 | 0.0108 | -0.0024 | 0.0017 |
| rs12981256 | A | 0.0140 | 0.0020 | 0.0159 | 0.0112 | -0.0021 | 0.0017 |
| rs13034320 | C | 0.0136 | 0.0023 | 0.0163 | 0.0117 | -0.0012 | 0.0020 |
| rs13047416 | C | 0.0143 | 0.0020 | -0.0031 | 0.0107 | 0.0015 | 0.0018 |
| rs13085002 | T | -0.0168 | 0.0027 | -0.0082 | 0.0137 | 0.0009 | 0.0023 |
| rs13107325 | T | 0.0469 | 0.0037 | -0.0015 | 0.0244 | -0.0188 | 0.0034 |
| rs13110266 | A | -0.0119 | 0.0020 | -0.0064 | 0.0103 | 0.0030 | 0.0017 |
| rs13132853 | A | 0.0138 | 0.0021 | 0.0059 | 0.0122 | -0.0096 | 0.0018 |
| rs13147390 | T | -0.0122 | 0.0020 | 0.0132 | 0.0110 | 0.0034 | 0.0018 |
| rs13174863 | A | -0.0210 | 0.0028 | -0.0139 | 0.0157 | -0.0014 | 0.0024 |
| rs13199223 | T | 0.0209 | 0.0028 | 0.0280 | 0.0152 | -0.0076 | 0.0025 |
| rs1320251 | T | -0.0168 | 0.0020 | -0.0133 | 0.0109 | 0.0038 | 0.0017 |
| rs1320903 | A | 0.0229 | 0.0021 | 0.0018 | 0.0112 | -0.0004 | 0.0018 |
| rs13209689 | A | -0.0132 | 0.0021 | -0.0051 | 0.0108 | 0.0056 | 0.0018 |
| rs13209918 | A | -0.0134 | 0.0020 | -0.0140 | 0.0109 | 0.0018 | 0.0018 |
| rs13218383 | C | 0.0135 | 0.0021 | 0.0202 | 0.0107 | -0.0014 | 0.0018 |
| rs13263601 | A | -0.0150 | 0.0020 | -0.0001 | 0.0116 | 0.0056 | 0.0018 |
| rs1327259 | A | 0.0155 | 0.0020 | 0.0024 | 0.0104 | -0.0031 | 0.0017 |
| rs1328448 | A | 0.0127 | 0.0021 | -0.0093 | 0.0112 | -0.0019 | 0.0019 |
| rs13288178 | A | 0.0141 | 0.0020 | 0.0278 | 0.0108 | 0.0012 | 0.0018 |
| rs13299074 | T | 0.0147 | 0.0025 | 0.0023 | 0.0140 | -0.0007 | 0.0022 |
| rs13427822 | A | 0.0178 | 0.0022 | 0.0129 | 0.0124 | -0.0053 | 0.0019 |
| rs1348716 | A | -0.0117 | 0.0020 | 0.0091 | 0.0105 | -0.0127 | 0.0018 |
| rs13665 | C | 0.0160 | 0.0021 | -0.0308 | 0.0110 | -0.0041 | 0.0018 |
| rs1406120 | A | -0.0182 | 0.0031 | -0.0236 | 0.0168 | 0.0123 | 0.0027 |
| rs1412235 | C | 0.0246 | 0.0021 | 0.0279 | 0.0114 | 0.0013 | 0.0018 |
| rs1421085 | T | -0.0767 | 0.0020 | -0.0266 | 0.0107 | -0.0027 | 0.0017 |
| rs1421334 | A | 0.0131 | 0.0020 | 0.0161 | 0.0106 | -0.0112 | 0.0017 |
| rs1431659 | A | 0.0222 | 0.0022 | 0.0061 | 0.0112 | -0.0001 | 0.0019 |
| rs1436344 | C | 0.0145 | 0.0020 | 0.0039 | 0.0103 | -0.0019 | 0.0017 |
| rs1454687 | C | 0.0226 | 0.0019 | 0.0033 | 0.0103 | -0.0072 | 0.0017 |
| rs1460940 | A | 0.0312 | 0.0024 | 0.0189 | 0.0147 | 0.0024 | 0.0022 |
| rs1470153 | A | -0.0138 | 0.0023 | 0.0262 | 0.0114 | 0.0067 | 0.0020 |
| rs1477890 | A | -0.0162 | 0.0020 | -0.0014 | 0.0101 | 0.0018 | 0.0017 |
| rs1503526 | T | -0.0144 | 0.0020 | -0.0097 | 0.0104 | 0.0105 | 0.0017 |
| rs1522569 | T | 0.0167 | 0.0025 | -0.0087 | 0.0137 | -0.0055 | 0.0022 |
| rs1574389 | T | 0.0161 | 0.0024 | 0.0176 | 0.0120 | -0.0031 | 0.0021 |
| rs1579557 | T | 0.0225 | 0.0021 | -0.0021 | 0.0117 | -0.0037 | 0.0019 |
| rs1582931 | A | -0.0142 | 0.0020 | -0.0217 | 0.0106 | -0.0074 | 0.0017 |
| rs1583123 | A | 0.0124 | 0.0020 | -0.0129 | 0.0106 | -0.0038 | 0.0018 |
| rs1681740 | A | 0.0122 | 0.0020 | -0.0040 | 0.0107 | -0.0027 | 0.0017 |
| rs16868443 | C | 0.0129 | 0.0020 | -0.0081 | 0.0108 | 0.0035 | 0.0018 |
| rs16882450 | A | 0.0132 | 0.0021 | 0.0202 | 0.0109 | -0.0057 | 0.0019 |
| rs16916303 | A | 0.0195 | 0.0031 | 0.0184 | 0.0148 | -0.0026 | 0.0027 |
| rs16973520 | T | -0.0138 | 0.0023 | -0.0316 | 0.0114 | 0.0045 | 0.0020 |
| rs16996657 | T | -0.0193 | 0.0029 | -0.0222 | 0.0173 | 0.0012 | 0.0025 |
| rs17014375 | T | -0.0187 | 0.0029 | 0.0056 | 0.0143 | 0.0004 | 0.0025 |
| rs1701827 | A | 0.0145 | 0.0022 | 0.0106 | 0.0122 | -0.0028 | 0.0019 |
| rs17024258 | T | 0.0648 | 0.0061 | -0.0360 | 0.0331 | 0.0039 | 0.0050 |
| rs17056301 | T | -0.0132 | 0.0023 | -0.0167 | 0.0112 | -0.0022 | 0.0019 |
| rs17066856 | T | 0.0392 | 0.0034 | 0.0357 | 0.0174 | -0.0042 | 0.0030 |
| rs1711171 | T | -0.0185 | 0.0023 | -0.0274 | 0.0124 | 0.0029 | 0.0020 |
| rs17132130 | C | -0.0173 | 0.0024 | -0.0281 | 0.0117 | 0.0108 | 0.0020 |
| rs17207196 | T | -0.0228 | 0.0020 | -0.0126 | 0.0121 | 0.0091 | 0.0018 |
| rs17379561 | A | -0.0181 | 0.0028 | -0.0218 | 0.0148 | 0.0190 | 0.0024 |
| rs17399237 | T | 0.0120 | 0.0020 | 0.0108 | 0.0110 | -0.0051 | 0.0017 |
| rs17446299 | C | -0.0157 | 0.0026 | -0.0176 | 0.0149 | -0.0014 | 0.0023 |
| rs17513613 | T | -0.0201 | 0.0021 | 0.0123 | 0.0114 | -0.0010 | 0.0018 |
| rs17636031 | T | -0.0164 | 0.0022 | 0.0130 | 0.0133 | 0.0041 | 0.0019 |
| rs17639546 | A | -0.0240 | 0.0027 | -0.0297 | 0.0146 | 0.0035 | 0.0024 |
| rs17724992 | A | 0.0177 | 0.0022 | 0.0103 | 0.0113 | -0.0060 | 0.0019 |
| rs17776719 | A | -0.0171 | 0.0028 | 0.0249 | 0.0135 | -0.0016 | 0.0024 |
| rs17806379 | T | -0.0267 | 0.0025 | -0.0097 | 0.0135 | 0.0079 | 0.0022 |
| rs17814208 | A | -0.0134 | 0.0023 | 0.0000 | 0.0121 | 0.0018 | 0.0020 |
| rs17827419 | A | -0.0213 | 0.0030 | -0.0107 | 0.0173 | 0.0033 | 0.0026 |
| rs1786141 | T | 0.0136 | 0.0020 | 0.0089 | 0.0108 | -0.0062 | 0.0017 |
| rs1814170 | A | 0.0198 | 0.0033 | -0.0059 | 0.0145 | 0.0040 | 0.0028 |
| rs1861410 | T | -0.0219 | 0.0020 | -0.0095 | 0.0102 | 0.0031 | 0.0017 |
| rs1884897 | A | -0.0216 | 0.0020 | -0.0058 | 0.0107 | -0.0010 | 0.0018 |
| rs1928538 | T | -0.0135 | 0.0020 | -0.0368 | 0.0103 | 0.0024 | 0.0017 |
| rs1951455 | T | -0.0150 | 0.0022 | 0.0158 | 0.0119 | 0.0018 | 0.0019 |
| rs1955540 | T | -0.0154 | 0.0025 | 0.0039 | 0.0129 | 0.0038 | 0.0022 |
| rs197374 | T | 0.0140 | 0.0020 | -0.0092 | 0.0105 | -0.0087 | 0.0018 |
| rs1976053 | C | -0.0186 | 0.0025 | -0.0364 | 0.0124 | 0.0034 | 0.0022 |
| rs2012286 | A | 0.0139 | 0.0020 | -0.0010 | 0.0104 | -0.0044 | 0.0017 |
| rs2051559 | T | -0.0192 | 0.0029 | -0.0177 | 0.0150 | 0.0152 | 0.0025 |
| rs2065418 | T | 0.0184 | 0.0020 | -0.0043 | 0.0108 | 0.0025 | 0.0018 |
| rs2072858 | T | 0.0164 | 0.0021 | -0.0057 | 0.0106 | 0.0045 | 0.0018 |
| rs214247 | T | 0.0136 | 0.0020 | 0.0027 | 0.0112 | -0.0035 | 0.0018 |
| rs2143253 | A | -0.0201 | 0.0030 | 0.0023 | 0.0138 | 0.0125 | 0.0026 |
| rs215634 | A | 0.0158 | 0.0020 | 0.0242 | 0.0106 | -0.0095 | 0.0018 |
| rs2161796 | T | 0.0141 | 0.0020 | 0.0005 | 0.0106 | 0.0005 | 0.0018 |
| rs217671 | A | -0.0169 | 0.0022 | -0.0053 | 0.0113 | 0.0070 | 0.0019 |
| rs2181375 | A | -0.0201 | 0.0020 | -0.0077 | 0.0102 | 0.0021 | 0.0017 |
| rs2183947 | A | -0.0162 | 0.0023 | 0.0160 | 0.0127 | 0.0106 | 0.0020 |
| rs2253310 | C | -0.0169 | 0.0020 | -0.0209 | 0.0106 | -0.0093 | 0.0018 |
| rs2271189 | A | -0.0161 | 0.0020 | -0.0298 | 0.0114 | 0.0121 | 0.0018 |
| rs2281819 | A | -0.0162 | 0.0023 | -0.0045 | 0.0124 | 0.0065 | 0.0020 |
| rs2284746 | C | 0.0112 | 0.0019 | 0.0174 | 0.0104 | -0.0026 | 0.0018 |
| rs2306540 | T | 0.0137 | 0.0021 | -0.0037 | 0.0120 | -0.0031 | 0.0018 |
| rs2307111 | T | 0.0276 | 0.0020 | 0.0086 | 0.0105 | -0.0052 | 0.0017 |
| rs2414128 | T | 0.0131 | 0.0020 | 0.0093 | 0.0110 | 0.0026 | 0.0017 |
| rs241472 | T | -0.0190 | 0.0021 | -0.0042 | 0.0108 | 0.0021 | 0.0018 |
| rs2425847 | A | -0.0126 | 0.0020 | 0.0024 | 0.0106 | 0.0031 | 0.0017 |
| rs2433733 | A | -0.0169 | 0.0021 | 0.0108 | 0.0108 | 0.0057 | 0.0018 |
| rs244415 | A | -0.0226 | 0.0020 | -0.0185 | 0.0107 | 0.0011 | 0.0017 |
| rs2444253 | C | 0.0138 | 0.0020 | 0.0060 | 0.0105 | -0.0038 | 0.0017 |
| rs2450444 | A | -0.0127 | 0.0021 | -0.0065 | 0.0114 | 0.0008 | 0.0018 |
| rs2472297 | T | 0.0147 | 0.0022 | -0.0120 | 0.0146 | -0.0002 | 0.0020 |
| rs2479958 | A | 0.0158 | 0.0020 | 0.0172 | 0.0108 | -0.0021 | 0.0017 |
| rs248139 | A | 0.0151 | 0.0025 | -0.0132 | 0.0129 | 0.0021 | 0.0021 |
| rs2591175 | A | -0.0144 | 0.0025 | -0.0098 | 0.0127 | -0.0013 | 0.0022 |
| rs2605570 | A | 0.0151 | 0.0026 | -0.0071 | 0.0136 | -0.0019 | 0.0023 |
| rs2616143 | A | -0.0133 | 0.0021 | 0.0108 | 0.0119 | 0.0005 | 0.0018 |
| rs2644112 | T | -0.0241 | 0.0021 | -0.0429 | 0.0111 | 0.0109 | 0.0018 |
| rs2693826 | A | -0.0136 | 0.0020 | -0.0162 | 0.0103 | 0.0022 | 0.0017 |
| rs2694047 | A | -0.0211 | 0.0023 | -0.0135 | 0.0119 | 0.0013 | 0.0020 |
| rs2733287 | A | -0.0155 | 0.0020 | -0.0198 | 0.0102 | -0.0067 | 0.0017 |
| rs273512 | T | 0.0171 | 0.0020 | -0.0112 | 0.0109 | -0.0054 | 0.0017 |
| rs2781668 | T | 0.0159 | 0.0026 | 0.0354 | 0.0136 | 0.0048 | 0.0023 |
| rs2791643 | T | -0.0143 | 0.0023 | 0.0033 | 0.0123 | 0.0040 | 0.0020 |
| rs2803316 | A | -0.0148 | 0.0020 | -0.0023 | 0.0115 | -0.0062 | 0.0017 |
| rs2814942 | A | 0.0269 | 0.0021 | 0.0332 | 0.0111 | -0.0038 | 0.0019 |
| rs28350 | A | 0.0186 | 0.0026 | 0.0004 | 0.0138 | 0.0022 | 0.0022 |
| rs2836968 | A | 0.0122 | 0.0020 | 0.0098 | 0.0112 | -0.0052 | 0.0018 |
| rs2861683 | A | 0.0171 | 0.0020 | -0.0054 | 0.0103 | 0.0003 | 0.0017 |
| rs2865384 | A | -0.0121 | 0.0020 | -0.0149 | 0.0105 | 0.0047 | 0.0018 |
| rs2875762 | C | 0.0151 | 0.0023 | -0.0029 | 0.0120 | -0.0012 | 0.0020 |
| rs2906132 | T | 0.0119 | 0.0020 | 0.0133 | 0.0110 | -0.0016 | 0.0017 |
| rs2920502 | C | -0.0147 | 0.0021 | -0.0111 | 0.0110 | 0.0030 | 0.0018 |
| rs294704 | T | -0.0146 | 0.0022 | -0.0136 | 0.0119 | -0.0007 | 0.0019 |
| rs2952862 | T | 0.0143 | 0.0021 | 0.0204 | 0.0106 | -0.0043 | 0.0018 |
| rs326896 | T | -0.0123 | 0.0020 | -0.0176 | 0.0103 | 0.0038 | 0.0017 |
| rs329651 | T | 0.0169 | 0.0025 | 0.0008 | 0.0154 | -0.0075 | 0.0022 |
| rs355748 | T | 0.0162 | 0.0020 | 0.0046 | 0.0104 | -0.0037 | 0.0018 |
| rs3730071 | A | -0.0340 | 0.0057 | 0.0353 | 0.0415 | -0.0043 | 0.0052 |
| rs3743950 | A | -0.0119 | 0.0020 | 0.0031 | 0.0116 | 0.0013 | 0.0018 |
| rs3754963 | A | 0.0139 | 0.0022 | 0.0113 | 0.0117 | -0.0106 | 0.0019 |
| rs3762444 | T | -0.0144 | 0.0020 | -0.0233 | 0.0107 | 0.0017 | 0.0017 |
| rs3770451 | T | 0.0144 | 0.0022 | -0.0084 | 0.0118 | -0.0104 | 0.0019 |
| rs3784710 | T | 0.0287 | 0.0023 | 0.0156 | 0.0119 | -0.0005 | 0.0021 |
| rs3803286 | A | 0.0193 | 0.0021 | 0.0137 | 0.0110 | -0.0070 | 0.0018 |
| rs3806114 | A | -0.0125 | 0.0021 | -0.0062 | 0.0121 | 0.0000 | 0.0018 |
| rs3806572 | A | -0.0154 | 0.0021 | 0.0108 | 0.0117 | -0.0004 | 0.0019 |
| rs3814883 | T | 0.0249 | 0.0020 | -0.0085 | 0.0111 | -0.0005 | 0.0017 |
| rs3821709 | T | -0.0338 | 0.0039 | -0.0183 | 0.0184 | 0.0128 | 0.0035 |
| rs38314 | A | -0.0116 | 0.0020 | -0.0004 | 0.0108 | -0.0057 | 0.0017 |
| rs394608 | T | -0.0177 | 0.0020 | -0.0139 | 0.0102 | 0.0044 | 0.0017 |
| rs40067 | A | -0.0272 | 0.0026 | -0.0322 | 0.0126 | 0.0011 | 0.0023 |
| rs4072917 | A | 0.0118 | 0.0020 | 0.0068 | 0.0109 | -0.0024 | 0.0017 |
| rs4148155 | A | 0.0203 | 0.0031 | -0.0218 | 0.0168 | -0.0004 | 0.0027 |
| rs4148866 | T | 0.0124 | 0.0020 | -0.0003 | 0.0113 | -0.0072 | 0.0017 |
| rs4237195 | A | 0.0177 | 0.0021 | 0.0098 | 0.0110 | -0.0046 | 0.0018 |
| rs4237643 | T | 0.0237 | 0.0021 | 0.0370 | 0.0113 | -0.0024 | 0.0019 |
| rs4240673 | T | 0.0201 | 0.0020 | 0.0007 | 0.0110 | -0.0026 | 0.0017 |
| rs4284600 | T | -0.0115 | 0.0020 | 0.0063 | 0.0105 | 0.0019 | 0.0017 |
| rs429343 | A | 0.0172 | 0.0020 | 0.0139 | 0.0104 | -0.0006 | 0.0017 |
| rs4307239 | A | -0.0120 | 0.0020 | -0.0089 | 0.0105 | 0.0000 | 0.0017 |
| rs4343164 | T | 0.0165 | 0.0022 | -0.0007 | 0.0120 | -0.0066 | 0.0019 |
| rs4477562 | T | 0.0317 | 0.0030 | 0.0134 | 0.0145 | -0.0001 | 0.0026 |
| rs4482463 | A | -0.0346 | 0.0037 | 0.0038 | 0.0178 | -0.0043 | 0.0032 |
| rs4490752 | A | 0.0172 | 0.0020 | -0.0027 | 0.0108 | 0.0000 | 0.0018 |
| rs4516268 | A | -0.0241 | 0.0025 | 0.0092 | 0.0145 | 0.0096 | 0.0022 |
| rs4639527 | A | -0.0174 | 0.0021 | -0.0035 | 0.0114 | -0.0012 | 0.0018 |
| rs4652839 | C | -0.0132 | 0.0021 | -0.0017 | 0.0109 | 0.0027 | 0.0018 |
| rs4654831 | T | 0.0188 | 0.0026 | -0.0003 | 0.0125 | -0.0021 | 0.0023 |
| rs4658403 | T | -0.0186 | 0.0026 | -0.0040 | 0.0127 | 0.0104 | 0.0023 |
| rs4660443 | T | 0.0182 | 0.0024 | 0.0169 | 0.0131 | -0.0010 | 0.0021 |
| rs4670169 | A | -0.0120 | 0.0020 | -0.0104 | 0.0107 | -0.0056 | 0.0018 |
| rs4722398 | T | 0.0178 | 0.0028 | -0.0086 | 0.0151 | 0.0040 | 0.0026 |
| rs4762369 | A | 0.0151 | 0.0025 | 0.0280 | 0.0134 | 0.0018 | 0.0022 |
| rs4764949 | A | 0.0196 | 0.0021 | 0.0045 | 0.0107 | 0.0010 | 0.0018 |
| rs4851716 | A | 0.0162 | 0.0020 | 0.0030 | 0.0108 | 0.0018 | 0.0018 |
| rs4911382 | T | 0.0147 | 0.0020 | 0.0088 | 0.0106 | -0.0034 | 0.0017 |
| rs4936175 | T | -0.0124 | 0.0020 | 0.0001 | 0.0112 | 0.0036 | 0.0017 |
| rs4942924 | T | 0.0158 | 0.0020 | 0.0428 | 0.0107 | -0.0022 | 0.0017 |
| rs4952843 | A | 0.0151 | 0.0020 | 0.0192 | 0.0109 | 0.0006 | 0.0017 |
| rs4969387 | C | -0.0146 | 0.0022 | -0.0033 | 0.0138 | -0.0027 | 0.0020 |
| rs500764 | T | 0.0164 | 0.0020 | 0.0273 | 0.0103 | 0.0001 | 0.0017 |
| rs538656 | T | 0.0544 | 0.0023 | 0.0520 | 0.0116 | 0.0056 | 0.0020 |
| rs538740 | T | -0.0145 | 0.0024 | -0.0097 | 0.0128 | 0.0074 | 0.0021 |
| rs5396 | T | -0.0175 | 0.0022 | 0.0269 | 0.0115 | 0.0088 | 0.0019 |
| rs543874 | A | -0.0466 | 0.0024 | -0.0238 | 0.0132 | 0.0047 | 0.0021 |
| rs559231 | T | 0.0142 | 0.0020 | 0.0170 | 0.0104 | 0.0015 | 0.0017 |
| rs577525 | T | -0.0187 | 0.0020 | -0.0181 | 0.0104 | 0.0066 | 0.0017 |
| rs592483 | T | -0.0152 | 0.0020 | 0.0137 | 0.0113 | 0.0029 | 0.0017 |
| rs6011457 | A | -0.0116 | 0.0020 | 0.0049 | 0.0104 | -0.0073 | 0.0017 |
| rs6023649 | A | 0.0153 | 0.0023 | 0.0137 | 0.0121 | -0.0074 | 0.0019 |
| rs6050446 | A | -0.0375 | 0.0056 | -0.0113 | 0.0350 | 0.0121 | 0.0048 |
| rs6098868 | A | -0.0117 | 0.0020 | 0.0041 | 0.0101 | -0.0010 | 0.0017 |
| rs6265 | T | -0.0405 | 0.0025 | -0.0303 | 0.0129 | 0.0097 | 0.0022 |
| rs6443750 | T | -0.0152 | 0.0025 | -0.0286 | 0.0169 | 0.0067 | 0.0022 |
| rs6448587 | A | 0.0196 | 0.0026 | 0.0098 | 0.0122 | -0.0004 | 0.0022 |
| rs6479905 | A | -0.0126 | 0.0020 | 0.0026 | 0.0101 | 0.0143 | 0.0017 |
| rs6502482 | C | -0.0133 | 0.0020 | -0.0031 | 0.0103 | -0.0041 | 0.0017 |
| rs6542924 | A | -0.0174 | 0.0021 | -0.0094 | 0.0119 | 0.0174 | 0.0018 |
| rs6551304 | A | -0.0186 | 0.0026 | -0.0074 | 0.0148 | 0.0047 | 0.0023 |
| rs6577584 | T | -0.0131 | 0.0021 | 0.0065 | 0.0108 | 0.0023 | 0.0018 |
| rs6587552 | A | 0.0175 | 0.0023 | 0.0056 | 0.0121 | -0.0076 | 0.0020 |
| rs6604866 | C | -0.0161 | 0.0020 | 0.0045 | 0.0104 | 0.0003 | 0.0017 |
| rs6674079 | A | -0.0127 | 0.0020 | 0.0048 | 0.0105 | 0.0055 | 0.0018 |
| rs6682 | A | -0.0121 | 0.0020 | -0.0131 | 0.0114 | 0.0038 | 0.0017 |
| rs671722 | A | -0.0126 | 0.0020 | -0.0080 | 0.0113 | 0.0093 | 0.0017 |
| rs6725931 | T | 0.0187 | 0.0027 | 0.0226 | 0.0140 | -0.0115 | 0.0024 |
| rs6728037 | A | 0.0198 | 0.0020 | 0.0101 | 0.0106 | -0.0046 | 0.0017 |
| rs6731688 | A | -0.0564 | 0.0026 | -0.0143 | 0.0136 | 0.0047 | 0.0023 |
| rs6752378 | A | 0.0332 | 0.0019 | 0.0329 | 0.0102 | 0.0012 | 0.0017 |
| rs6764533 | A | 0.0141 | 0.0020 | 0.0138 | 0.0112 | -0.0073 | 0.0018 |
| rs6831020 | A | -0.0161 | 0.0021 | -0.0085 | 0.0122 | 0.0027 | 0.0019 |
| rs6870983 | T | -0.0219 | 0.0024 | -0.0143 | 0.0120 | 0.0141 | 0.0021 |
| rs6905544 | A | -0.0164 | 0.0020 | -0.0036 | 0.0107 | 0.0233 | 0.0017 |
| rs698147 | A | 0.0130 | 0.0020 | 0.0063 | 0.0102 | -0.0045 | 0.0017 |
| rs7038943 | T | 0.0143 | 0.0021 | 0.0209 | 0.0107 | -0.0075 | 0.0018 |
| rs705158 | A | 0.0162 | 0.0023 | -0.0014 | 0.0114 | 0.0007 | 0.0020 |
| rs705217 | T | 0.0123 | 0.0020 | -0.0048 | 0.0106 | -0.0040 | 0.0018 |
| rs7084454 | A | 0.0210 | 0.0021 | 0.0018 | 0.0115 | -0.0068 | 0.0018 |
| rs7096307 | T | -0.0117 | 0.0020 | 0.0101 | 0.0108 | 0.0024 | 0.0017 |
| rs7102454 | T | -0.0176 | 0.0020 | -0.0130 | 0.0116 | 0.0071 | 0.0018 |
| rs7124681 | A | 0.0269 | 0.0020 | -0.0092 | 0.0108 | 0.0002 | 0.0017 |
| rs7132908 | A | 0.0296 | 0.0020 | 0.0041 | 0.0107 | -0.0005 | 0.0018 |
| rs7158822 | T | -0.0162 | 0.0021 | -0.0184 | 0.0107 | 0.0034 | 0.0018 |
| rs7181498 | T | 0.0160 | 0.0021 | 0.0053 | 0.0108 | -0.0052 | 0.0018 |
| rs7196720 | T | 0.0131 | 0.0020 | -0.0088 | 0.0106 | -0.0045 | 0.0017 |
| rs7258722 | A | -0.0200 | 0.0020 | -0.0252 | 0.0109 | 0.0015 | 0.0017 |
| rs725959 | T | -0.0133 | 0.0020 | -0.0169 | 0.0108 | 0.0054 | 0.0017 |
| rs7313220 | A | 0.0129 | 0.0020 | -0.0068 | 0.0102 | 0.0020 | 0.0017 |
| rs7334078 | T | 0.0125 | 0.0022 | 0.0059 | 0.0116 | -0.0103 | 0.0019 |
| rs733758 | T | -0.0313 | 0.0045 | 0.0066 | 0.0260 | -0.0068 | 0.0039 |
| rs738140 | A | 0.0147 | 0.0021 | -0.0050 | 0.0118 | -0.0033 | 0.0018 |
| rs741959 | A | 0.0173 | 0.0020 | 0.0100 | 0.0106 | -0.0001 | 0.0017 |
| rs7498665 | A | -0.0256 | 0.0020 | -0.0180 | 0.0113 | 0.0166 | 0.0017 |
| rs7518221 | T | 0.0126 | 0.0020 | 0.0062 | 0.0108 | 0.0003 | 0.0018 |
| rs7519259 | A | 0.0137 | 0.0020 | 0.0148 | 0.0104 | -0.0064 | 0.0017 |
| rs754635 | C | -0.0198 | 0.0031 | 0.0026 | 0.0152 | -0.0032 | 0.0027 |
| rs7557796 | T | 0.0171 | 0.0020 | 0.0265 | 0.0106 | -0.0030 | 0.0018 |
| rs756717 | A | -0.0158 | 0.0020 | -0.0209 | 0.0112 | -0.0006 | 0.0017 |
| rs7568228 | C | -0.0116 | 0.0019 | -0.0120 | 0.0105 | 0.0050 | 0.0017 |
| rs7619139 | A | 0.0146 | 0.0020 | 0.0115 | 0.0104 | 0.0007 | 0.0017 |
| rs7626079 | T | 0.0121 | 0.0021 | -0.0016 | 0.0107 | 0.0052 | 0.0018 |
| rs7637852 | A | 0.0143 | 0.0021 | 0.0195 | 0.0114 | -0.0013 | 0.0019 |
| rs7640424 | T | -0.0128 | 0.0021 | -0.0074 | 0.0119 | 0.0107 | 0.0018 |
| rs7646113 | T | -0.0125 | 0.0020 | 0.0117 | 0.0106 | 0.0010 | 0.0018 |
| rs765874 | A | -0.0125 | 0.0020 | 0.0188 | 0.0104 | 0.0022 | 0.0017 |
| rs7683836 | A | -0.0135 | 0.0020 | -0.0086 | 0.0104 | 0.0003 | 0.0017 |
| rs769449 | A | -0.0235 | 0.0030 | 0.0855 | 0.0183 | 0.0067 | 0.0026 |
| rs7707981 | A | 0.0142 | 0.0021 | -0.0128 | 0.0117 | 0.0009 | 0.0018 |
| rs7715256 | T | -0.0165 | 0.0020 | -0.0047 | 0.0105 | -0.0007 | 0.0017 |
| rs7733438 | T | -0.0365 | 0.0029 | -0.0278 | 0.0142 | -0.0173 | 0.0025 |
| rs774246 | A | -0.0169 | 0.0028 | 0.0087 | 0.0161 | 0.0012 | 0.0024 |
| rs7747583 | A | -0.0136 | 0.0022 | 0.0046 | 0.0108 | 0.0074 | 0.0019 |
| rs775403 | T | 0.0117 | 0.0020 | 0.0054 | 0.0104 | -0.0044 | 0.0017 |
| rs7792703 | A | -0.0166 | 0.0024 | -0.0133 | 0.0122 | 0.0010 | 0.0020 |
| rs7869771 | A | 0.0156 | 0.0022 | 0.0124 | 0.0114 | -0.0047 | 0.0019 |
| rs7871866 | C | 0.0177 | 0.0027 | 0.0155 | 0.0145 | -0.0062 | 0.0024 |
| rs7893571 | T | 0.0146 | 0.0021 | 0.0012 | 0.0111 | 0.0024 | 0.0018 |
| rs7903146 | T | -0.0157 | 0.0022 | 0.0241 | 0.0114 | -0.0021 | 0.0019 |
| rs7925214 | T | 0.0157 | 0.0020 | 0.0162 | 0.0106 | -0.0046 | 0.0017 |
| rs799449 | T | 0.0141 | 0.0020 | 0.0227 | 0.0108 | -0.0070 | 0.0017 |
| rs8008285 | T | -0.0175 | 0.0028 | 0.0076 | 0.0150 | -0.0046 | 0.0024 |
| rs8015400 | A | 0.0226 | 0.0021 | -0.0017 | 0.0110 | -0.0022 | 0.0018 |
| rs8089514 | A | 0.0133 | 0.0021 | 0.0203 | 0.0115 | -0.0038 | 0.0018 |
| rs8181823 | A | -0.0139 | 0.0023 | -0.0042 | 0.0125 | 0.0061 | 0.0020 |
| rs879620 | T | 0.0227 | 0.0020 | 0.0013 | 0.0110 | -0.0008 | 0.0018 |
| rs890156 | A | 0.0125 | 0.0020 | 0.0004 | 0.0103 | 0.0023 | 0.0017 |
| rs891387 | T | 0.0205 | 0.0020 | 0.0173 | 0.0104 | -0.0137 | 0.0017 |
| rs895330 | C | 0.0205 | 0.0025 | 0.0371 | 0.0168 | -0.0034 | 0.0022 |
| rs900144 | T | 0.0174 | 0.0020 | 0.0227 | 0.0105 | 0.0015 | 0.0017 |
| rs902503 | A | 0.0130 | 0.0020 | 0.0003 | 0.0105 | -0.0013 | 0.0017 |
| rs903959 | A | 0.0116 | 0.0020 | -0.0169 | 0.0111 | -0.0157 | 0.0017 |
| rs912690 | C | -0.0152 | 0.0020 | 0.0146 | 0.0102 | 0.0030 | 0.0017 |
| rs9294260 | A | 0.0158 | 0.0020 | -0.0028 | 0.0103 | -0.0054 | 0.0017 |
| rs9296389 | C | 0.0122 | 0.0020 | 0.0107 | 0.0103 | -0.0012 | 0.0017 |
| rs930295 | A | 0.0214 | 0.0026 | 0.0112 | 0.0142 | 0.0000 | 0.0023 |
| rs9304665 | A | 0.0225 | 0.0023 | 0.0241 | 0.0119 | -0.0003 | 0.0020 |
| rs942065 | A | 0.0204 | 0.0020 | 0.0150 | 0.0108 | -0.0071 | 0.0018 |
| rs9446149 | T | 0.0145 | 0.0024 | 0.0119 | 0.0128 | 0.0017 | 0.0021 |
| rs9478496 | T | -0.0173 | 0.0026 | -0.0178 | 0.0145 | 0.0013 | 0.0023 |
| rs9496567 | A | 0.0163 | 0.0023 | 0.0004 | 0.0130 | 0.0008 | 0.0020 |
| rs9515448 | A | -0.0141 | 0.0020 | -0.0008 | 0.0105 | 0.0066 | 0.0017 |
| rs9530843 | A | 0.0134 | 0.0020 | -0.0050 | 0.0107 | -0.0027 | 0.0017 |
| rs9538162 | T | 0.0168 | 0.0020 | -0.0082 | 0.0104 | -0.0018 | 0.0017 |
| rs9571687 | A | -0.0126 | 0.0021 | 0.0035 | 0.0110 | -0.0016 | 0.0018 |
| rs960923 | A | -0.0210 | 0.0020 | 0.0064 | 0.0102 | -0.0029 | 0.0017 |
| rs9630985 | A | -0.0173 | 0.0021 | -0.0214 | 0.0110 | 0.0058 | 0.0018 |
| rs9650755 | A | -0.0159 | 0.0022 | -0.0003 | 0.0112 | 0.0069 | 0.0019 |
| rs9688431 | T | 0.0242 | 0.0042 | -0.0092 | 0.0209 | 0.0000 | 0.0036 |
| rs9816226 | A | -0.0292 | 0.0025 | 0.0009 | 0.0134 | -0.0078 | 0.0022 |
| rs9827823 | T | 0.0219 | 0.0027 | -0.0225 | 0.0141 | -0.0017 | 0.0024 |
| rs987237 | A | -0.0395 | 0.0025 | -0.0177 | 0.0130 | 0.0033 | 0.0022 |
| rs9876664 | T | -0.0181 | 0.0020 | -0.0014 | 0.0105 | 0.0139 | 0.0018 |
| rs9926784 | T | 0.0262 | 0.0025 | 0.0041 | 0.0128 | 0.0032 | 0.0022 |
| rs9927848 | A | -0.0129 | 0.0022 | 0.0177 | 0.0115 | -0.0022 | 0.0019 |
| rs9951619 | T | -0.0160 | 0.0024 | -0.0103 | 0.0119 | -0.0052 | 0.0020 |
| rs9958909 | T | -0.0214 | 0.0029 | -0.0123 | 0.0157 | 0.0027 | 0.0024 |

#### Supplementary table 9: Genetic association estimates for the two-sample regression-based MVMR analysis estimating the effect of BMI on stroke, adjusted for education. ea=effect allele; gx=BMI; gy=stroke; gz=education; se=standard error

| SNP | ea | gx | gx_se | gy | gy_se | gz | gz_se |
| --- | --- | --- | --- | --- | --- | --- | --- |
| rs10083803 | T | -0.0139 | 0.0022 | -0.0112 | 0.0089 | -0.0020 | 0.0019 |
| rs10124645 | A | 0.0124 | 0.0020 | 0.0094 | 0.0088 | -0.0041 | 0.0017 |
| rs10128597 | A | -0.0186 | 0.0022 | -0.0061 | 0.0087 | -0.0004 | 0.0019 |
| rs1013293 | A | -0.0177 | 0.0020 | -0.0050 | 0.0087 | 0.0030 | 0.0017 |
| rs10145154 | T | 0.0288 | 0.0024 | 0.0139 | 0.0102 | 0.0002 | 0.0021 |
| rs10160769 | C | -0.0154 | 0.0024 | -0.0120 | 0.0093 | 0.0125 | 0.0021 |
| rs10169594 | T | -0.0122 | 0.0020 | -0.0175 | 0.0091 | 0.0086 | 0.0018 |
| rs10206166 | T | 0.0168 | 0.0020 | 0.0089 | 0.0081 | -0.0111 | 0.0018 |
| rs10236214 | T | 0.0145 | 0.0021 | -0.0288 | 0.0090 | 0.0015 | 0.0018 |
| rs10259490 | A | -0.0129 | 0.0020 | -0.0030 | 0.0080 | -0.0023 | 0.0017 |
| rs10423928 | A | -0.0342 | 0.0025 | 0.0034 | 0.0098 | 0.0031 | 0.0021 |
| rs1045411 | T | -0.0163 | 0.0022 | -0.0125 | 0.0096 | 0.0032 | 0.0019 |
| rs1048932 | A | -0.0149 | 0.0020 | -0.0034 | 0.0079 | 0.0041 | 0.0017 |
| rs10514944 | A | 0.0247 | 0.0034 | -0.0106 | 0.0124 | -0.0074 | 0.0029 |
| rs10514975 | A | -0.0139 | 0.0020 | 0.0120 | 0.0080 | 0.0012 | 0.0017 |
| rs10515237 | A | -0.0151 | 0.0022 | -0.0078 | 0.0087 | -0.0003 | 0.0019 |
| rs1064213 | A | 0.0146 | 0.0019 | 0.0033 | 0.0081 | -0.0026 | 0.0017 |
| rs10742752 | T | -0.0119 | 0.0020 | -0.0068 | 0.0081 | -0.0026 | 0.0017 |
| rs10756714 | A | 0.0199 | 0.0020 | 0.0023 | 0.0080 | -0.0048 | 0.0017 |
| rs10768994 | T | 0.0125 | 0.0020 | 0.0004 | 0.0078 | -0.0017 | 0.0017 |
| rs10790519 | T | -0.0124 | 0.0020 | 0.0023 | 0.0079 | 0.0041 | 0.0018 |
| rs10818810 | A | 0.0125 | 0.0020 | -0.0016 | 0.0082 | -0.0051 | 0.0018 |
| rs10824218 | A | 0.0141 | 0.0020 | 0.0033 | 0.0085 | 0.0020 | 0.0017 |
| rs10841188 | A | -0.0135 | 0.0022 | -0.0048 | 0.0088 | 0.0010 | 0.0020 |
| rs10842240 | C | 0.0215 | 0.0031 | -0.0081 | 0.0108 | -0.0071 | 0.0027 |
| rs10878946 | T | -0.0148 | 0.0022 | 0.0105 | 0.0089 | -0.0015 | 0.0019 |
| rs10887578 | C | 0.0134 | 0.0020 | -0.0143 | 0.0081 | -0.0019 | 0.0017 |
| rs10899736 | A | -0.0116 | 0.0020 | -0.0016 | 0.0078 | -0.0020 | 0.0017 |
| rs10921743 | A | 0.0138 | 0.0020 | 0.0094 | 0.0083 | -0.0021 | 0.0018 |
| rs10938397 | A | -0.0292 | 0.0020 | -0.0112 | 0.0083 | 0.0066 | 0.0017 |
| rs10962550 | C | 0.0196 | 0.0025 | -0.0043 | 0.0098 | -0.0093 | 0.0022 |
| rs11075489 | T | -0.0115 | 0.0020 | -0.0069 | 0.0081 | 0.0023 | 0.0017 |
| rs11079849 | T | -0.0213 | 0.0021 | -0.0058 | 0.0094 | 0.0074 | 0.0018 |
| rs11084553 | A | 0.0236 | 0.0027 | 0.0000 | 0.0129 | 0.0045 | 0.0024 |
| rs11134679 | A | -0.0177 | 0.0021 | -0.0131 | 0.0091 | 0.0025 | 0.0018 |
| rs11150461 | C | 0.0148 | 0.0022 | 0.0097 | 0.0090 | 0.0017 | 0.0019 |
| rs11161044 | C | 0.0175 | 0.0025 | 0.0071 | 0.0094 | -0.0180 | 0.0021 |
| rs11165493 | A | 0.0143 | 0.0021 | -0.0132 | 0.0095 | 0.0053 | 0.0018 |
| rs11190658 | T | 0.0150 | 0.0024 | 0.0100 | 0.0095 | 0.0002 | 0.0021 |
| rs11223970 | A | -0.0193 | 0.0026 | 0.0048 | 0.0126 | 0.0049 | 0.0023 |
| rs11251352 | A | -0.0117 | 0.0020 | 0.0088 | 0.0081 | 0.0038 | 0.0017 |
| rs11496125 | T | 0.0189 | 0.0020 | -0.0053 | 0.0079 | 0.0025 | 0.0017 |
| rs11505821 | A | -0.0301 | 0.0040 | 0.0137 | 0.0156 | -0.0030 | 0.0036 |
| rs11609297 | T | -0.0172 | 0.0023 | -0.0010 | 0.0106 | 0.0058 | 0.0020 |
| rs11611246 | T | 0.0255 | 0.0024 | -0.0014 | 0.0097 | 0.0079 | 0.0021 |
| rs11615578 | T | 0.0137 | 0.0023 | -0.0047 | 0.0104 | 0.0022 | 0.0020 |
| rs11675464 | A | -0.0117 | 0.0020 | 0.0126 | 0.0081 | -0.0001 | 0.0017 |
| rs11713193 | A | 0.0266 | 0.0019 | 0.0168 | 0.0084 | -0.0225 | 0.0017 |
| rs11757278 | T | 0.0146 | 0.0021 | 0.0058 | 0.0084 | -0.0073 | 0.0018 |
| rs11773362 | T | -0.0125 | 0.0021 | -0.0104 | 0.0082 | 0.0033 | 0.0018 |
| rs11846126 | T | 0.0158 | 0.0020 | 0.0147 | 0.0081 | -0.0037 | 0.0017 |
| rs11874191 | T | -0.0151 | 0.0023 | 0.0167 | 0.0088 | 0.0069 | 0.0020 |
| rs11895606 | T | 0.0131 | 0.0020 | 0.0087 | 0.0086 | 0.0015 | 0.0017 |
| rs11915371 | A | -0.0148 | 0.0024 | -0.0082 | 0.0101 | 0.0102 | 0.0021 |
| rs11945861 | A | -0.0143 | 0.0023 | -0.0008 | 0.0092 | 0.0068 | 0.0020 |
| rs11976018 | A | -0.0228 | 0.0027 | -0.0138 | 0.0103 | 0.0041 | 0.0023 |
| rs12049202 | T | 0.0245 | 0.0024 | 0.0200 | 0.0095 | -0.0067 | 0.0021 |
| rs1218822 | A | 0.0176 | 0.0021 | -0.0172 | 0.0083 | -0.0024 | 0.0018 |
| rs12334877 | A | -0.0152 | 0.0025 | -0.0076 | 0.0095 | 0.0047 | 0.0021 |
| rs12369179 | T | -0.0372 | 0.0034 | 0.0270 | 0.0168 | -0.0143 | 0.0030 |
| rs12427047 | T | -0.0184 | 0.0023 | -0.0199 | 0.0091 | -0.0041 | 0.0020 |
| rs12439798 | T | 0.0139 | 0.0020 | 0.0027 | 0.0080 | -0.0038 | 0.0017 |
| rs12448257 | A | 0.0157 | 0.0024 | 0.0042 | 0.0097 | 0.0074 | 0.0021 |
| rs12512994 | T | -0.0127 | 0.0021 | -0.0051 | 0.0092 | 0.0140 | 0.0018 |
| rs12564848 | T | -0.0229 | 0.0031 | 0.0365 | 0.0120 | -0.0015 | 0.0027 |
| rs12595749 | A | 0.0149 | 0.0020 | -0.0119 | 0.0079 | -0.0022 | 0.0017 |
| rs12608738 | A | 0.0130 | 0.0022 | 0.0079 | 0.0098 | -0.0020 | 0.0019 |
| rs1266874 | A | -0.0147 | 0.0020 | 0.0088 | 0.0084 | 0.0023 | 0.0018 |
| rs12681792 | A | 0.0149 | 0.0025 | 0.0260 | 0.0096 | -0.0034 | 0.0021 |
| rs12705977 | T | 0.0134 | 0.0020 | 0.0011 | 0.0085 | -0.0079 | 0.0017 |
| rs12762034 | T | -0.0262 | 0.0037 | -0.0372 | 0.0140 | 0.0059 | 0.0032 |
| rs12878246 | T | 0.0137 | 0.0023 | 0.0023 | 0.0101 | -0.0017 | 0.0020 |
| rs12906208 | A | -0.0149 | 0.0024 | 0.0011 | 0.0104 | 0.0039 | 0.0021 |
| rs12936083 | A | -0.0137 | 0.0021 | 0.0000 | 0.0083 | 0.0014 | 0.0018 |
| rs12936319 | A | 0.0167 | 0.0020 | -0.0143 | 0.0084 | -0.0072 | 0.0017 |
| rs12949279 | T | 0.0180 | 0.0020 | 0.0070 | 0.0080 | -0.0031 | 0.0017 |
| rs1296328 | A | 0.0190 | 0.0020 | 0.0086 | 0.0090 | -0.0024 | 0.0017 |
| rs12981256 | A | 0.0140 | 0.0020 | 0.0036 | 0.0086 | -0.0021 | 0.0017 |
| rs13034320 | C | 0.0136 | 0.0023 | 0.0102 | 0.0088 | -0.0012 | 0.0020 |
| rs13047416 | C | 0.0143 | 0.0020 | -0.0065 | 0.0083 | 0.0015 | 0.0018 |
| rs13085002 | T | -0.0168 | 0.0027 | 0.0041 | 0.0114 | 0.0009 | 0.0023 |
| rs13107325 | T | 0.0469 | 0.0037 | -0.0005 | 0.0193 | -0.0188 | 0.0034 |
| rs13110266 | A | -0.0119 | 0.0020 | 0.0031 | 0.0080 | 0.0030 | 0.0017 |
| rs13132853 | A | 0.0138 | 0.0021 | -0.0083 | 0.0096 | -0.0096 | 0.0018 |
| rs13147390 | T | -0.0122 | 0.0020 | -0.0059 | 0.0082 | 0.0034 | 0.0018 |
| rs13174863 | A | -0.0210 | 0.0028 | 0.0084 | 0.0125 | -0.0014 | 0.0024 |
| rs13199223 | T | 0.0209 | 0.0028 | -0.0213 | 0.0124 | -0.0076 | 0.0025 |
| rs1320251 | T | -0.0168 | 0.0020 | -0.0104 | 0.0084 | 0.0038 | 0.0017 |
| rs1320903 | A | 0.0229 | 0.0021 | -0.0012 | 0.0087 | -0.0004 | 0.0018 |
| rs13209689 | A | -0.0132 | 0.0021 | 0.0018 | 0.0086 | 0.0056 | 0.0018 |
| rs13209918 | A | -0.0134 | 0.0020 | -0.0093 | 0.0084 | 0.0018 | 0.0018 |
| rs13218383 | C | 0.0135 | 0.0021 | 0.0175 | 0.0082 | -0.0014 | 0.0018 |
| rs13263601 | A | -0.0150 | 0.0020 | -0.0074 | 0.0086 | 0.0056 | 0.0018 |
| rs1327259 | A | 0.0155 | 0.0020 | -0.0101 | 0.0083 | -0.0031 | 0.0017 |
| rs1328448 | A | 0.0127 | 0.0021 | -0.0007 | 0.0085 | -0.0019 | 0.0019 |
| rs13288178 | A | 0.0141 | 0.0020 | -0.0014 | 0.0083 | 0.0012 | 0.0018 |
| rs13299074 | T | 0.0147 | 0.0025 | -0.0260 | 0.0106 | -0.0007 | 0.0022 |
| rs13427822 | A | 0.0178 | 0.0022 | -0.0069 | 0.0096 | -0.0053 | 0.0019 |
| rs1348716 | A | -0.0117 | 0.0020 | 0.0022 | 0.0083 | -0.0127 | 0.0018 |
| rs13665 | C | 0.0160 | 0.0021 | -0.0064 | 0.0086 | -0.0041 | 0.0018 |
| rs1406120 | A | -0.0182 | 0.0031 | -0.0206 | 0.0130 | 0.0123 | 0.0027 |
| rs1412235 | C | 0.0246 | 0.0021 | 0.0051 | 0.0088 | 0.0013 | 0.0018 |
| rs1421085 | T | -0.0767 | 0.0020 | -0.0071 | 0.0085 | -0.0027 | 0.0017 |
| rs1421334 | A | 0.0131 | 0.0020 | -0.0076 | 0.0082 | -0.0112 | 0.0017 |
| rs1431659 | A | 0.0222 | 0.0022 | 0.0142 | 0.0088 | -0.0001 | 0.0019 |
| rs1436344 | C | 0.0145 | 0.0020 | 0.0077 | 0.0081 | -0.0019 | 0.0017 |
| rs1454687 | C | 0.0226 | 0.0019 | 0.0066 | 0.0081 | -0.0072 | 0.0017 |
| rs1460940 | A | 0.0312 | 0.0024 | -0.0167 | 0.0108 | 0.0024 | 0.0022 |
| rs1470153 | A | -0.0138 | 0.0023 | -0.0008 | 0.0088 | 0.0067 | 0.0020 |
| rs1477890 | A | -0.0162 | 0.0020 | -0.0065 | 0.0079 | 0.0018 | 0.0017 |
| rs1503526 | T | -0.0144 | 0.0020 | -0.0187 | 0.0083 | 0.0105 | 0.0017 |
| rs1522569 | T | 0.0167 | 0.0025 | 0.0074 | 0.0112 | -0.0055 | 0.0022 |
| rs1574389 | T | 0.0161 | 0.0024 | -0.0093 | 0.0091 | -0.0031 | 0.0021 |
| rs1579557 | T | 0.0225 | 0.0021 | 0.0008 | 0.0087 | -0.0037 | 0.0019 |
| rs1582931 | A | -0.0142 | 0.0020 | -0.0283 | 0.0079 | -0.0074 | 0.0017 |
| rs1583123 | A | 0.0124 | 0.0020 | -0.0054 | 0.0082 | -0.0038 | 0.0018 |
| rs1681740 | A | 0.0122 | 0.0020 | -0.0020 | 0.0085 | -0.0027 | 0.0017 |
| rs16868443 | C | 0.0129 | 0.0020 | -0.0012 | 0.0086 | 0.0035 | 0.0018 |
| rs16882450 | A | 0.0132 | 0.0021 | 0.0154 | 0.0088 | -0.0057 | 0.0019 |
| rs16916303 | A | 0.0195 | 0.0031 | 0.0028 | 0.0111 | -0.0026 | 0.0027 |
| rs16973520 | T | -0.0138 | 0.0023 | -0.0173 | 0.0087 | 0.0045 | 0.0020 |
| rs16996657 | T | -0.0193 | 0.0029 | -0.0177 | 0.0126 | 0.0012 | 0.0025 |
| rs17014375 | T | -0.0187 | 0.0029 | -0.0209 | 0.0114 | 0.0004 | 0.0025 |
| rs1701827 | A | 0.0145 | 0.0022 | 0.0112 | 0.0091 | -0.0028 | 0.0019 |
| rs17024258 | T | 0.0648 | 0.0061 | -0.0185 | 0.0239 | 0.0039 | 0.0050 |
| rs17056301 | T | -0.0132 | 0.0023 | 0.0075 | 0.0090 | -0.0022 | 0.0019 |
| rs17066856 | T | 0.0392 | 0.0034 | -0.0159 | 0.0121 | -0.0042 | 0.0030 |
| rs1711171 | T | -0.0185 | 0.0023 | 0.0003 | 0.0093 | 0.0029 | 0.0020 |
| rs17132130 | C | -0.0173 | 0.0024 | -0.0094 | 0.0094 | 0.0108 | 0.0020 |
| rs17207196 | T | -0.0228 | 0.0020 | -0.0356 | 0.0096 | 0.0091 | 0.0018 |
| rs17379561 | A | -0.0181 | 0.0028 | -0.0013 | 0.0121 | 0.0190 | 0.0024 |
| rs17399237 | T | 0.0120 | 0.0020 | -0.0114 | 0.0087 | -0.0051 | 0.0017 |
| rs17446299 | C | -0.0157 | 0.0026 | 0.0028 | 0.0116 | -0.0014 | 0.0023 |
| rs17513613 | T | -0.0201 | 0.0021 | 0.0067 | 0.0090 | -0.0010 | 0.0018 |
| rs17636031 | T | -0.0164 | 0.0022 | -0.0031 | 0.0107 | 0.0041 | 0.0019 |
| rs17639546 | A | -0.0240 | 0.0027 | -0.0011 | 0.0110 | 0.0035 | 0.0024 |
| rs17724992 | A | 0.0177 | 0.0022 | 0.0233 | 0.0087 | -0.0060 | 0.0019 |
| rs17776719 | A | -0.0171 | 0.0028 | 0.0131 | 0.0108 | -0.0016 | 0.0024 |
| rs17806379 | T | -0.0267 | 0.0025 | -0.0203 | 0.0106 | 0.0079 | 0.0022 |
| rs17814208 | A | -0.0134 | 0.0023 | -0.0049 | 0.0095 | 0.0018 | 0.0020 |
| rs17827419 | A | -0.0213 | 0.0030 | -0.0329 | 0.0139 | 0.0033 | 0.0026 |
| rs1786141 | T | 0.0136 | 0.0020 | 0.0026 | 0.0087 | -0.0062 | 0.0017 |
| rs1814170 | A | 0.0198 | 0.0033 | -0.0108 | 0.0110 | 0.0040 | 0.0028 |
| rs1861410 | T | -0.0219 | 0.0020 | 0.0020 | 0.0079 | 0.0031 | 0.0017 |
| rs1884897 | A | -0.0216 | 0.0020 | -0.0024 | 0.0086 | -0.0010 | 0.0018 |
| rs1928538 | T | -0.0135 | 0.0020 | -0.0138 | 0.0078 | 0.0024 | 0.0017 |
| rs1951455 | T | -0.0150 | 0.0022 | 0.0162 | 0.0093 | 0.0018 | 0.0019 |
| rs1955540 | T | -0.0154 | 0.0025 | -0.0013 | 0.0098 | 0.0038 | 0.0022 |
| rs197374 | T | 0.0140 | 0.0020 | -0.0064 | 0.0081 | -0.0087 | 0.0018 |
| rs1976053 | C | -0.0186 | 0.0025 | -0.0143 | 0.0093 | 0.0034 | 0.0022 |
| rs2012286 | A | 0.0139 | 0.0020 | 0.0083 | 0.0081 | -0.0044 | 0.0017 |
| rs2051559 | T | -0.0192 | 0.0029 | -0.0268 | 0.0112 | 0.0152 | 0.0025 |
| rs2065418 | T | 0.0184 | 0.0020 | 0.0183 | 0.0083 | 0.0025 | 0.0018 |
| rs2072858 | T | 0.0164 | 0.0021 | -0.0143 | 0.0083 | 0.0045 | 0.0018 |
| rs214247 | T | 0.0136 | 0.0020 | 0.0026 | 0.0083 | -0.0035 | 0.0018 |
| rs2143253 | A | -0.0201 | 0.0030 | -0.0191 | 0.0108 | 0.0125 | 0.0026 |
| rs215634 | A | 0.0158 | 0.0020 | 0.0100 | 0.0082 | -0.0095 | 0.0018 |
| rs2161796 | T | 0.0141 | 0.0020 | 0.0069 | 0.0080 | 0.0005 | 0.0018 |
| rs217671 | A | -0.0169 | 0.0022 | 0.0116 | 0.0088 | 0.0070 | 0.0019 |
| rs2181375 | A | -0.0201 | 0.0020 | 0.0039 | 0.0081 | 0.0021 | 0.0017 |
| rs2183947 | A | -0.0162 | 0.0023 | -0.0064 | 0.0096 | 0.0106 | 0.0020 |
| rs2253310 | C | -0.0169 | 0.0020 | 0.0085 | 0.0081 | -0.0093 | 0.0018 |
| rs2271189 | A | -0.0161 | 0.0020 | -0.0095 | 0.0081 | 0.0121 | 0.0018 |
| rs2281819 | A | -0.0162 | 0.0023 | -0.0122 | 0.0095 | 0.0065 | 0.0020 |
| rs2284746 | C | 0.0112 | 0.0019 | -0.0020 | 0.0083 | -0.0026 | 0.0018 |
| rs2306540 | T | 0.0137 | 0.0021 | 0.0096 | 0.0098 | -0.0031 | 0.0018 |
| rs2307111 | T | 0.0276 | 0.0020 | 0.0103 | 0.0083 | -0.0052 | 0.0017 |
| rs2414128 | T | 0.0131 | 0.0020 | 0.0025 | 0.0081 | 0.0026 | 0.0017 |
| rs241472 | T | -0.0190 | 0.0021 | 0.0049 | 0.0084 | 0.0021 | 0.0018 |
| rs2425847 | A | -0.0126 | 0.0020 | -0.0147 | 0.0083 | 0.0031 | 0.0017 |
| rs2433733 | A | -0.0169 | 0.0021 | -0.0070 | 0.0085 | 0.0057 | 0.0018 |
| rs244415 | A | -0.0226 | 0.0020 | -0.0091 | 0.0083 | 0.0011 | 0.0017 |
| rs2444253 | C | 0.0138 | 0.0020 | -0.0029 | 0.0079 | -0.0038 | 0.0017 |
| rs2450444 | A | -0.0127 | 0.0021 | 0.0220 | 0.0087 | 0.0008 | 0.0018 |
| rs2472297 | T | 0.0147 | 0.0022 | -0.0003 | 0.0111 | -0.0002 | 0.0020 |
| rs2479958 | A | 0.0158 | 0.0020 | 0.0084 | 0.0085 | -0.0021 | 0.0017 |
| rs248139 | A | 0.0151 | 0.0025 | -0.0118 | 0.0097 | 0.0021 | 0.0021 |
| rs2591175 | A | -0.0144 | 0.0025 | 0.0001 | 0.0095 | -0.0013 | 0.0022 |
| rs2605570 | A | 0.0151 | 0.0026 | 0.0133 | 0.0104 | -0.0019 | 0.0023 |
| rs2616143 | A | -0.0133 | 0.0021 | -0.0117 | 0.0093 | 0.0005 | 0.0018 |
| rs2644112 | T | -0.0241 | 0.0021 | -0.0187 | 0.0088 | 0.0109 | 0.0018 |
| rs2693826 | A | -0.0136 | 0.0020 | -0.0150 | 0.0079 | 0.0022 | 0.0017 |
| rs2694047 | A | -0.0211 | 0.0023 | -0.0057 | 0.0089 | 0.0013 | 0.0020 |
| rs2733287 | A | -0.0155 | 0.0020 | -0.0068 | 0.0078 | -0.0067 | 0.0017 |
| rs273512 | T | 0.0171 | 0.0020 | 0.0131 | 0.0082 | -0.0054 | 0.0017 |
| rs2781668 | T | 0.0159 | 0.0026 | 0.0017 | 0.0102 | 0.0048 | 0.0023 |
| rs2791643 | T | -0.0143 | 0.0023 | 0.0014 | 0.0097 | 0.0040 | 0.0020 |
| rs2803316 | A | -0.0148 | 0.0020 | 0.0079 | 0.0088 | -0.0062 | 0.0017 |
| rs2814942 | A | 0.0269 | 0.0021 | 0.0111 | 0.0082 | -0.0038 | 0.0019 |
| rs28350 | A | 0.0186 | 0.0026 | -0.0133 | 0.0108 | 0.0022 | 0.0022 |
| rs2836968 | A | 0.0122 | 0.0020 | -0.0040 | 0.0089 | -0.0052 | 0.0018 |
| rs2861683 | A | 0.0171 | 0.0020 | 0.0085 | 0.0080 | 0.0003 | 0.0017 |
| rs2865384 | A | -0.0121 | 0.0020 | -0.0112 | 0.0082 | 0.0047 | 0.0018 |
| rs2875762 | C | 0.0151 | 0.0023 | -0.0001 | 0.0094 | -0.0012 | 0.0020 |
| rs2906132 | T | 0.0119 | 0.0020 | 0.0094 | 0.0088 | -0.0016 | 0.0017 |
| rs2920502 | C | -0.0147 | 0.0021 | 0.0091 | 0.0086 | 0.0030 | 0.0018 |
| rs294704 | T | -0.0146 | 0.0022 | 0.0091 | 0.0096 | -0.0007 | 0.0019 |
| rs2952862 | T | 0.0143 | 0.0021 | 0.0189 | 0.0090 | -0.0043 | 0.0018 |
| rs326896 | T | -0.0123 | 0.0020 | 0.0001 | 0.0082 | 0.0038 | 0.0017 |
| rs329651 | T | 0.0169 | 0.0025 | -0.0174 | 0.0121 | -0.0075 | 0.0022 |
| rs355748 | T | 0.0162 | 0.0020 | 0.0009 | 0.0079 | -0.0037 | 0.0018 |
| rs3730071 | A | -0.0340 | 0.0057 | 0.0424 | 0.0294 | -0.0043 | 0.0052 |
| rs3743950 | A | -0.0119 | 0.0020 | -0.0001 | 0.0090 | 0.0013 | 0.0018 |
| rs3754963 | A | 0.0139 | 0.0022 | 0.0208 | 0.0090 | -0.0106 | 0.0019 |
| rs3762444 | T | -0.0144 | 0.0020 | -0.0035 | 0.0081 | 0.0017 | 0.0017 |
| rs3770451 | T | 0.0144 | 0.0022 | -0.0019 | 0.0088 | -0.0104 | 0.0019 |
| rs3784710 | T | 0.0287 | 0.0023 | 0.0241 | 0.0089 | -0.0005 | 0.0021 |
| rs3803286 | A | 0.0193 | 0.0021 | -0.0074 | 0.0081 | -0.0070 | 0.0018 |
| rs3806114 | A | -0.0125 | 0.0021 | -0.0045 | 0.0088 | 0.0000 | 0.0018 |
| rs3806572 | A | -0.0154 | 0.0021 | -0.0079 | 0.0094 | -0.0004 | 0.0019 |
| rs3814883 | T | 0.0249 | 0.0020 | -0.0208 | 0.0083 | -0.0005 | 0.0017 |
| rs3821709 | T | -0.0338 | 0.0039 | -0.0134 | 0.0134 | 0.0128 | 0.0035 |
| rs38314 | A | -0.0116 | 0.0020 | -0.0231 | 0.0085 | -0.0057 | 0.0017 |
| rs394608 | T | -0.0177 | 0.0020 | 0.0028 | 0.0080 | 0.0044 | 0.0017 |
| rs40067 | A | -0.0272 | 0.0026 | -0.0266 | 0.0099 | 0.0011 | 0.0023 |
| rs4072917 | A | 0.0118 | 0.0020 | -0.0051 | 0.0079 | -0.0024 | 0.0017 |
| rs4148155 | A | 0.0203 | 0.0031 | 0.0039 | 0.0119 | -0.0004 | 0.0027 |
| rs4148866 | T | 0.0124 | 0.0020 | 0.0129 | 0.0084 | -0.0072 | 0.0017 |
| rs4237195 | A | 0.0177 | 0.0021 | 0.0094 | 0.0088 | -0.0046 | 0.0018 |
| rs4237643 | T | 0.0237 | 0.0021 | 0.0259 | 0.0086 | -0.0024 | 0.0019 |
| rs4240673 | T | 0.0201 | 0.0020 | 0.0153 | 0.0088 | -0.0026 | 0.0017 |
| rs4284600 | T | -0.0115 | 0.0020 | -0.0125 | 0.0087 | 0.0019 | 0.0017 |
| rs429343 | A | 0.0172 | 0.0020 | 0.0194 | 0.0083 | -0.0006 | 0.0017 |
| rs4307239 | A | -0.0120 | 0.0020 | 0.0087 | 0.0082 | 0.0000 | 0.0017 |
| rs4343164 | T | 0.0165 | 0.0022 | 0.0086 | 0.0097 | -0.0066 | 0.0019 |
| rs4477562 | T | 0.0317 | 0.0030 | 0.0258 | 0.0112 | -0.0001 | 0.0026 |
| rs4482463 | A | -0.0346 | 0.0037 | -0.0381 | 0.0120 | -0.0043 | 0.0032 |
| rs4490752 | A | 0.0172 | 0.0020 | 0.0101 | 0.0082 | 0.0000 | 0.0018 |
| rs4516268 | A | -0.0241 | 0.0025 | -0.0193 | 0.0102 | 0.0096 | 0.0022 |
| rs4639527 | A | -0.0174 | 0.0021 | 0.0214 | 0.0087 | -0.0012 | 0.0018 |
| rs4652839 | C | -0.0132 | 0.0021 | -0.0116 | 0.0086 | 0.0027 | 0.0018 |
| rs4654831 | T | 0.0188 | 0.0026 | 0.0089 | 0.0098 | -0.0021 | 0.0023 |
| rs4658403 | T | -0.0186 | 0.0026 | -0.0075 | 0.0098 | 0.0104 | 0.0023 |
| rs4660443 | T | 0.0182 | 0.0024 | -0.0003 | 0.0098 | -0.0010 | 0.0021 |
| rs4670169 | A | -0.0120 | 0.0020 | -0.0061 | 0.0089 | -0.0056 | 0.0018 |
| rs4722398 | T | 0.0178 | 0.0028 | 0.0102 | 0.0122 | 0.0040 | 0.0026 |
| rs4762369 | A | 0.0151 | 0.0025 | -0.0053 | 0.0105 | 0.0018 | 0.0022 |
| rs4764949 | A | 0.0196 | 0.0021 | -0.0013 | 0.0083 | 0.0010 | 0.0018 |
| rs4851716 | A | 0.0162 | 0.0020 | -0.0024 | 0.0082 | 0.0018 | 0.0018 |
| rs4911382 | T | 0.0147 | 0.0020 | 0.0061 | 0.0080 | -0.0034 | 0.0017 |
| rs4936175 | T | -0.0124 | 0.0020 | 0.0000 | 0.0090 | 0.0036 | 0.0017 |
| rs4942924 | T | 0.0158 | 0.0020 | 0.0036 | 0.0082 | -0.0022 | 0.0017 |
| rs4952843 | A | 0.0151 | 0.0020 | -0.0022 | 0.0087 | 0.0006 | 0.0017 |
| rs4969387 | C | -0.0146 | 0.0022 | 0.0065 | 0.0095 | -0.0027 | 0.0020 |
| rs500764 | T | 0.0164 | 0.0020 | -0.0035 | 0.0080 | 0.0001 | 0.0017 |
| rs538656 | T | 0.0544 | 0.0023 | 0.0077 | 0.0089 | 0.0056 | 0.0020 |
| rs538740 | T | -0.0145 | 0.0024 | -0.0082 | 0.0095 | 0.0074 | 0.0021 |
| rs5396 | T | -0.0175 | 0.0022 | -0.0135 | 0.0096 | 0.0088 | 0.0019 |
| rs543874 | A | -0.0466 | 0.0024 | 0.0008 | 0.0098 | 0.0047 | 0.0021 |
| rs559231 | T | 0.0142 | 0.0020 | 0.0029 | 0.0082 | 0.0015 | 0.0017 |
| rs577525 | T | -0.0187 | 0.0020 | 0.0122 | 0.0081 | 0.0066 | 0.0017 |
| rs592483 | T | -0.0152 | 0.0020 | 0.0148 | 0.0084 | 0.0029 | 0.0017 |
| rs6011457 | A | -0.0116 | 0.0020 | 0.0051 | 0.0081 | -0.0073 | 0.0017 |
| rs6023649 | A | 0.0153 | 0.0023 | -0.0076 | 0.0095 | -0.0074 | 0.0019 |
| rs6050446 | A | -0.0375 | 0.0056 | -0.0066 | 0.0292 | 0.0121 | 0.0048 |
| rs6098868 | A | -0.0117 | 0.0020 | 0.0145 | 0.0079 | -0.0010 | 0.0017 |
| rs6265 | T | -0.0405 | 0.0025 | -0.0136 | 0.0100 | 0.0097 | 0.0022 |
| rs6443750 | T | -0.0152 | 0.0025 | -0.0048 | 0.0127 | 0.0067 | 0.0022 |
| rs6448587 | A | 0.0196 | 0.0026 | 0.0057 | 0.0097 | -0.0004 | 0.0022 |
| rs6479905 | A | -0.0126 | 0.0020 | -0.0062 | 0.0078 | 0.0143 | 0.0017 |
| rs6502482 | C | -0.0133 | 0.0020 | 0.0112 | 0.0081 | -0.0041 | 0.0017 |
| rs6542924 | A | -0.0174 | 0.0021 | -0.0119 | 0.0092 | 0.0174 | 0.0018 |
| rs6551304 | A | -0.0186 | 0.0026 | 0.0148 | 0.0106 | 0.0047 | 0.0023 |
| rs6577584 | T | -0.0131 | 0.0021 | 0.0000 | 0.0085 | 0.0023 | 0.0018 |
| rs6587552 | A | 0.0175 | 0.0023 | -0.0040 | 0.0093 | -0.0076 | 0.0020 |
| rs6604866 | C | -0.0161 | 0.0020 | 0.0064 | 0.0090 | 0.0003 | 0.0017 |
| rs6674079 | A | -0.0127 | 0.0020 | 0.0062 | 0.0082 | 0.0055 | 0.0018 |
| rs6682 | A | -0.0121 | 0.0020 | -0.0155 | 0.0083 | 0.0038 | 0.0017 |
| rs671722 | A | -0.0126 | 0.0020 | -0.0073 | 0.0090 | 0.0093 | 0.0017 |
| rs6725931 | T | 0.0187 | 0.0027 | 0.0096 | 0.0112 | -0.0115 | 0.0024 |
| rs6728037 | A | 0.0198 | 0.0020 | 0.0125 | 0.0081 | -0.0046 | 0.0017 |
| rs6731688 | A | -0.0564 | 0.0026 | -0.0037 | 0.0109 | 0.0047 | 0.0023 |
| rs6752378 | A | 0.0332 | 0.0019 | 0.0002 | 0.0081 | 0.0012 | 0.0017 |
| rs6764533 | A | 0.0141 | 0.0020 | 0.0176 | 0.0087 | -0.0073 | 0.0018 |
| rs6831020 | A | -0.0161 | 0.0021 | 0.0015 | 0.0099 | 0.0027 | 0.0019 |
| rs6870983 | T | -0.0219 | 0.0024 | -0.0170 | 0.0101 | 0.0141 | 0.0021 |
| rs6905544 | A | -0.0164 | 0.0020 | -0.0018 | 0.0080 | 0.0233 | 0.0017 |
| rs698147 | A | 0.0130 | 0.0020 | 0.0005 | 0.0081 | -0.0045 | 0.0017 |
| rs7038943 | T | 0.0143 | 0.0021 | 0.0012 | 0.0080 | -0.0075 | 0.0018 |
| rs705158 | A | 0.0162 | 0.0023 | -0.0001 | 0.0091 | 0.0007 | 0.0020 |
| rs705217 | T | 0.0123 | 0.0020 | 0.0059 | 0.0083 | -0.0040 | 0.0018 |
| rs7084454 | A | 0.0210 | 0.0021 | -0.0183 | 0.0090 | -0.0068 | 0.0018 |
| rs7096307 | T | -0.0117 | 0.0020 | -0.0063 | 0.0085 | 0.0024 | 0.0017 |
| rs7102454 | T | -0.0176 | 0.0020 | -0.0097 | 0.0084 | 0.0071 | 0.0018 |
| rs7124681 | A | 0.0269 | 0.0020 | 0.0177 | 0.0080 | 0.0002 | 0.0017 |
| rs7132908 | A | 0.0296 | 0.0020 | 0.0057 | 0.0088 | -0.0005 | 0.0018 |
| rs7158822 | T | -0.0162 | 0.0021 | -0.0052 | 0.0079 | 0.0034 | 0.0018 |
| rs7181498 | T | 0.0160 | 0.0021 | 0.0006 | 0.0082 | -0.0052 | 0.0018 |
| rs7196720 | T | 0.0131 | 0.0020 | 0.0087 | 0.0081 | -0.0045 | 0.0017 |
| rs7258722 | A | -0.0200 | 0.0020 | -0.0155 | 0.0084 | 0.0015 | 0.0017 |
| rs725959 | T | -0.0133 | 0.0020 | -0.0088 | 0.0081 | 0.0054 | 0.0017 |
| rs7313220 | A | 0.0129 | 0.0020 | -0.0079 | 0.0080 | 0.0020 | 0.0017 |
| rs7334078 | T | 0.0125 | 0.0022 | 0.0090 | 0.0091 | -0.0103 | 0.0019 |
| rs733758 | T | -0.0313 | 0.0045 | 0.0108 | 0.0192 | -0.0068 | 0.0039 |
| rs738140 | A | 0.0147 | 0.0021 | 0.0103 | 0.0096 | -0.0033 | 0.0018 |
| rs741959 | A | 0.0173 | 0.0020 | 0.0070 | 0.0087 | -0.0001 | 0.0017 |
| rs7498665 | A | -0.0256 | 0.0020 | -0.0014 | 0.0085 | 0.0166 | 0.0017 |
| rs7518221 | T | 0.0126 | 0.0020 | 0.0056 | 0.0084 | 0.0003 | 0.0018 |
| rs7519259 | A | 0.0137 | 0.0020 | 0.0246 | 0.0083 | -0.0064 | 0.0017 |
| rs754635 | C | -0.0198 | 0.0031 | 0.0059 | 0.0110 | -0.0032 | 0.0027 |
| rs7557796 | T | 0.0171 | 0.0020 | -0.0021 | 0.0083 | -0.0030 | 0.0018 |
| rs756717 | A | -0.0158 | 0.0020 | 0.0037 | 0.0083 | -0.0006 | 0.0017 |
| rs7568228 | C | -0.0116 | 0.0019 | -0.0030 | 0.0086 | 0.0050 | 0.0017 |
| rs7619139 | A | 0.0146 | 0.0020 | 0.0052 | 0.0079 | 0.0007 | 0.0017 |
| rs7626079 | T | 0.0121 | 0.0021 | 0.0013 | 0.0082 | 0.0052 | 0.0018 |
| rs7637852 | A | 0.0143 | 0.0021 | 0.0050 | 0.0092 | -0.0013 | 0.0019 |
| rs7640424 | T | -0.0128 | 0.0021 | -0.0127 | 0.0097 | 0.0107 | 0.0018 |
| rs7646113 | T | -0.0125 | 0.0020 | -0.0033 | 0.0080 | 0.0010 | 0.0018 |
| rs765874 | A | -0.0125 | 0.0020 | 0.0130 | 0.0082 | 0.0022 | 0.0017 |
| rs7683836 | A | -0.0135 | 0.0020 | -0.0020 | 0.0082 | 0.0003 | 0.0017 |
| rs769449 | A | -0.0235 | 0.0030 | -0.0060 | 0.0137 | 0.0067 | 0.0026 |
| rs7707981 | A | 0.0142 | 0.0021 | 0.0014 | 0.0091 | 0.0009 | 0.0018 |
| rs7715256 | T | -0.0165 | 0.0020 | -0.0063 | 0.0083 | -0.0007 | 0.0017 |
| rs7733438 | T | -0.0365 | 0.0029 | -0.0154 | 0.0107 | -0.0173 | 0.0025 |
| rs774246 | A | -0.0169 | 0.0028 | 0.0113 | 0.0123 | 0.0012 | 0.0024 |
| rs7747583 | A | -0.0136 | 0.0022 | -0.0127 | 0.0084 | 0.0074 | 0.0019 |
| rs775403 | T | 0.0117 | 0.0020 | 0.0094 | 0.0088 | -0.0044 | 0.0017 |
| rs7792703 | A | -0.0166 | 0.0024 | 0.0059 | 0.0097 | 0.0010 | 0.0020 |
| rs7869771 | A | 0.0156 | 0.0022 | -0.0033 | 0.0087 | -0.0047 | 0.0019 |
| rs7871866 | C | 0.0177 | 0.0027 | 0.0023 | 0.0108 | -0.0062 | 0.0024 |
| rs7893571 | T | 0.0146 | 0.0021 | 0.0007 | 0.0089 | 0.0024 | 0.0018 |
| rs7903146 | T | -0.0157 | 0.0022 | 0.0324 | 0.0097 | -0.0021 | 0.0019 |
| rs7925214 | T | 0.0157 | 0.0020 | 0.0070 | 0.0079 | -0.0046 | 0.0017 |
| rs799449 | T | 0.0141 | 0.0020 | 0.0082 | 0.0086 | -0.0070 | 0.0017 |
| rs8008285 | T | -0.0175 | 0.0028 | 0.0101 | 0.0119 | -0.0046 | 0.0024 |
| rs8015400 | A | 0.0226 | 0.0021 | -0.0134 | 0.0084 | -0.0022 | 0.0018 |
| rs8089514 | A | 0.0133 | 0.0021 | 0.0071 | 0.0090 | -0.0038 | 0.0018 |
| rs8181823 | A | -0.0139 | 0.0023 | 0.0058 | 0.0094 | 0.0061 | 0.0020 |
| rs879620 | T | 0.0227 | 0.0020 | 0.0069 | 0.0084 | -0.0008 | 0.0018 |
| rs890156 | A | 0.0125 | 0.0020 | -0.0022 | 0.0078 | 0.0023 | 0.0017 |
| rs891387 | T | 0.0205 | 0.0020 | 0.0197 | 0.0083 | -0.0137 | 0.0017 |
| rs895330 | C | 0.0205 | 0.0025 | 0.0319 | 0.0113 | -0.0034 | 0.0022 |
| rs900144 | T | 0.0174 | 0.0020 | -0.0007 | 0.0083 | 0.0015 | 0.0017 |
| rs902503 | A | 0.0130 | 0.0020 | -0.0009 | 0.0080 | -0.0013 | 0.0017 |
| rs903959 | A | 0.0116 | 0.0020 | 0.0087 | 0.0085 | -0.0157 | 0.0017 |
| rs912690 | C | -0.0152 | 0.0020 | -0.0134 | 0.0078 | 0.0030 | 0.0017 |
| rs9294260 | A | 0.0158 | 0.0020 | 0.0138 | 0.0082 | -0.0054 | 0.0017 |
| rs9296389 | C | 0.0122 | 0.0020 | -0.0146 | 0.0078 | -0.0012 | 0.0017 |
| rs930295 | A | 0.0214 | 0.0026 | 0.0040 | 0.0111 | 0.0000 | 0.0023 |
| rs9304665 | A | 0.0225 | 0.0023 | 0.0058 | 0.0098 | -0.0003 | 0.0020 |
| rs942065 | A | 0.0204 | 0.0020 | 0.0047 | 0.0084 | -0.0071 | 0.0018 |
| rs9446149 | T | 0.0145 | 0.0024 | -0.0007 | 0.0100 | 0.0017 | 0.0021 |
| rs9478496 | T | -0.0173 | 0.0026 | -0.0364 | 0.0111 | 0.0013 | 0.0023 |
| rs9496567 | A | 0.0163 | 0.0023 | 0.0146 | 0.0104 | 0.0008 | 0.0020 |
| rs9515448 | A | -0.0141 | 0.0020 | -0.0135 | 0.0081 | 0.0066 | 0.0017 |
| rs9530843 | A | 0.0134 | 0.0020 | -0.0048 | 0.0083 | -0.0027 | 0.0017 |
| rs9538162 | T | 0.0168 | 0.0020 | 0.0005 | 0.0082 | -0.0018 | 0.0017 |
| rs9571687 | A | -0.0126 | 0.0021 | -0.0013 | 0.0082 | -0.0016 | 0.0018 |
| rs960923 | A | -0.0210 | 0.0020 | 0.0035 | 0.0080 | -0.0029 | 0.0017 |
| rs9630985 | A | -0.0173 | 0.0021 | 0.0045 | 0.0085 | 0.0058 | 0.0018 |
| rs9650755 | A | -0.0159 | 0.0022 | 0.0034 | 0.0087 | 0.0069 | 0.0019 |
| rs9688431 | T | 0.0242 | 0.0042 | 0.0130 | 0.0136 | 0.0000 | 0.0036 |
| rs9816226 | A | -0.0292 | 0.0025 | -0.0253 | 0.0111 | -0.0078 | 0.0022 |
| rs9827823 | T | 0.0219 | 0.0027 | -0.0070 | 0.0111 | -0.0017 | 0.0024 |
| rs987237 | A | -0.0395 | 0.0025 | -0.0172 | 0.0102 | 0.0033 | 0.0022 |
| rs9876664 | T | -0.0181 | 0.0020 | -0.0042 | 0.0082 | 0.0139 | 0.0018 |
| rs9926784 | T | 0.0262 | 0.0025 | -0.0014 | 0.0097 | 0.0032 | 0.0022 |
| rs9927848 | A | -0.0129 | 0.0022 | -0.0031 | 0.0089 | -0.0022 | 0.0019 |
| rs9951619 | T | -0.0160 | 0.0024 | 0.0026 | 0.0094 | -0.0052 | 0.0020 |
| rs9958909 | T | -0.0214 | 0.0029 | -0.0027 | 0.0120 | 0.0027 | 0.0024 |

#### Supplementary table 10: Genetic association estimates for the IVW ratio-method MR analysis estimating the effect of education on BMI. ea=effect allele; gx=education; gy=BMI; se=standard error

| SNP | ea | gx | gx_se | gy | gy_se |
| --- | --- | --- | --- | --- | --- |
| rs10032941 | T | -0.0080 | 0.0015 | -0.0026 | 0.0020 |
| rs10041403 | T | 0.0151 | 0.0019 | -0.0047 | 0.0026 |
| rs10042828 | A | -0.0145 | 0.0022 | 0.0051 | 0.0030 |
| rs10057590 | A | 0.0106 | 0.0014 | -0.0108 | 0.0020 |
| rs10061420 | A | 0.0082 | 0.0014 | 0.0019 | 0.0020 |
| rs1007731 | A | -0.0143 | 0.0022 | -0.0028 | 0.0031 |
| rs10080647 | A | 0.0124 | 0.0020 | 0.0022 | 0.0028 |
| rs1009470 | T | 0.0086 | 0.0014 | 0.0033 | 0.0020 |
| rs10098073 | A | -0.0091 | 0.0014 | -0.0064 | 0.0020 |
| rs10169002 | A | -0.0080 | 0.0014 | -0.0033 | 0.0020 |
| rs10181071 | A | 0.0102 | 0.0019 | -0.0035 | 0.0026 |
| rs10189857 | A | 0.0158 | 0.0014 | -0.0076 | 0.0020 |
| rs10192369 | A | -0.0078 | 0.0014 | 0.0025 | 0.0019 |
| rs10192834 | T | 0.0083 | 0.0015 | 0.0020 | 0.0020 |
| rs10204051 | T | -0.0101 | 0.0014 | 0.0059 | 0.0020 |
| rs10205057 | T | 0.0090 | 0.0014 | -0.0012 | 0.0020 |
| rs10208 | T | 0.0091 | 0.0015 | 0.0100 | 0.0021 |
| rs10251438 | A | 0.0078 | 0.0014 | -0.0058 | 0.0020 |
| rs1035578 | A | -0.0094 | 0.0014 | 0.0024 | 0.0020 |
| rs10480450 | A | 0.0158 | 0.0028 | -0.0021 | 0.0042 |
| rs10496091 | A | -0.0131 | 0.0016 | 0.0028 | 0.0022 |
| rs10499535 | A | 0.0080 | 0.0014 | -0.0047 | 0.0020 |
| rs1050847 | T | 0.0092 | 0.0014 | 0.0016 | 0.0020 |
| rs10509251 | T | -0.0093 | 0.0017 | 0.0023 | 0.0024 |
| rs10515007 | T | 0.0143 | 0.0019 | 0.0035 | 0.0027 |
| rs1051860 | A | -0.0079 | 0.0014 | 0.0047 | 0.0020 |
| rs1054442 | A | -0.0136 | 0.0015 | -0.0098 | 0.0020 |
| rs1066769 | A | -0.0230 | 0.0041 | -0.0006 | 0.0053 |
| rs10750539 | A | -0.0095 | 0.0015 | -0.0047 | 0.0021 |
| rs10761202 | T | 0.0088 | 0.0014 | -0.0010 | 0.0020 |
| rs10782651 | T | -0.0082 | 0.0014 | 0.0125 | 0.0020 |
| rs10783243 | A | -0.0092 | 0.0014 | -0.0011 | 0.0020 |
| rs10789285 | T | 0.0091 | 0.0016 | -0.0020 | 0.0023 |
| rs10795831 | T | -0.0099 | 0.0017 | 0.0039 | 0.0024 |
| rs10798888 | T | -0.0140 | 0.0019 | 0.0126 | 0.0026 |
| rs10799615 | A | -0.0089 | 0.0016 | 0.0069 | 0.0022 |
| rs10805383 | A | -0.0103 | 0.0014 | 0.0142 | 0.0020 |
| rs10810099 | A | -0.0146 | 0.0016 | 0.0062 | 0.0022 |
| rs10831656 | T | 0.0090 | 0.0015 | -0.0017 | 0.0021 |
| rs10845051 | A | 0.0083 | 0.0014 | -0.0019 | 0.0020 |
| rs10875121 | C | 0.0174 | 0.0019 | -0.0164 | 0.0026 |
| rs10877283 | T | 0.0085 | 0.0014 | -0.0010 | 0.0020 |
| rs10879676 | T | -0.0083 | 0.0014 | 0.0046 | 0.0020 |
| rs10928190 | T | -0.0087 | 0.0014 | 0.0002 | 0.0020 |
| rs10937240 | T | -0.0123 | 0.0018 | -0.0246 | 0.0025 |
| rs10940921 | T | 0.0111 | 0.0014 | 0.0002 | 0.0020 |
| rs10947439 | T | 0.0090 | 0.0015 | -0.0021 | 0.0021 |
| rs10974256 | A | -0.0093 | 0.0015 | -0.0028 | 0.0021 |
| rs1097784 | T | -0.0083 | 0.0014 | -0.0031 | 0.0020 |
| rs10979613 | T | -0.0117 | 0.0015 | 0.0000 | 0.0020 |
| rs10985402 | T | 0.0089 | 0.0016 | -0.0010 | 0.0023 |
| rs11030084 | T | 0.0106 | 0.0018 | -0.0385 | 0.0025 |
| rs11030102 | C | 0.0101 | 0.0016 | -0.0286 | 0.0022 |
| rs1106090 | A | 0.0094 | 0.0014 | -0.0128 | 0.0020 |
| rs11076962 | T | 0.0086 | 0.0016 | -0.0044 | 0.0022 |
| rs11081529 | T | 0.0126 | 0.0015 | -0.0009 | 0.0022 |
| rs11082011 | T | 0.0194 | 0.0015 | 0.0039 | 0.0021 |
| rs11100237 | A | -0.0101 | 0.0014 | 0.0000 | 0.0020 |
| rs11115056 | A | 0.0187 | 0.0023 | -0.0123 | 0.0033 |
| rs11130380 | T | 0.0090 | 0.0014 | 0.0042 | 0.0020 |
| rs11138947 | T | 0.0105 | 0.0016 | 0.0034 | 0.0022 |
| rs11155813 | T | -0.0129 | 0.0023 | 0.0026 | 0.0033 |
| rs11158800 | A | -0.0091 | 0.0014 | 0.0085 | 0.0020 |
| rs1117152 | T | 0.0106 | 0.0015 | -0.0069 | 0.0022 |
| rs11174399 | A | 0.0111 | 0.0018 | -0.0021 | 0.0024 |
| rs11190955 | T | -0.0083 | 0.0015 | -0.0058 | 0.0020 |
| rs1120924 | T | 0.0108 | 0.0015 | -0.0057 | 0.0021 |
| rs11210400 | A | 0.0120 | 0.0014 | 0.0032 | 0.0020 |
| rs11213482 | A | -0.0106 | 0.0019 | 0.0006 | 0.0026 |
| rs1123285 | C | -0.0094 | 0.0015 | 0.0016 | 0.0021 |
| rs11265191 | T | 0.0083 | 0.0015 | 0.0001 | 0.0021 |
| rs1146079 | C | -0.0087 | 0.0016 | 0.0050 | 0.0022 |
| rs11588857 | A | 0.0199 | 0.0017 | -0.0030 | 0.0024 |
| rs11599236 | T | -0.0106 | 0.0014 | 0.0080 | 0.0020 |
| rs11613431 | A | -0.0094 | 0.0016 | 0.0014 | 0.0022 |
| rs11623285 | T | -0.0135 | 0.0021 | -0.0009 | 0.0029 |
| rs11644446 | A | 0.0117 | 0.0019 | -0.0037 | 0.0028 |
| rs11663602 | A | -0.0130 | 0.0016 | 0.0051 | 0.0022 |
| rs11663678 | T | -0.0121 | 0.0022 | 0.0129 | 0.0030 |
| rs11675476 | T | -0.0127 | 0.0014 | 0.0068 | 0.0019 |
| rs1167827 | A | 0.0104 | 0.0014 | -0.0204 | 0.0020 |
| rs11693885 | A | -0.0091 | 0.0014 | 0.0045 | 0.0020 |
| rs1171040 | A | -0.0113 | 0.0017 | 0.0064 | 0.0024 |
| rs11724690 | T | 0.0093 | 0.0015 | 0.0001 | 0.0022 |
| rs11732160 | A | 0.0105 | 0.0016 | -0.0079 | 0.0023 |
| rs11780023 | T | 0.0085 | 0.0014 | 0.0005 | 0.0020 |
| rs11789013 | T | 0.0106 | 0.0016 | 0.0020 | 0.0023 |
| rs11845781 | T | -0.0079 | 0.0014 | 0.0017 | 0.0020 |
| rs11871429 | A | 0.0128 | 0.0017 | -0.0040 | 0.0023 |
| rs11876620 | T | 0.0143 | 0.0024 | 0.0150 | 0.0033 |
| rs11894424 | A | 0.0209 | 0.0033 | -0.0135 | 0.0047 |
| rs11917701 | A | -0.0087 | 0.0014 | -0.0025 | 0.0020 |
| rs11919835 | C | 0.0089 | 0.0015 | 0.0002 | 0.0020 |
| rs1196760 | C | 0.0139 | 0.0025 | -0.0105 | 0.0034 |
| rs12028010 | T | 0.0164 | 0.0017 | -0.0014 | 0.0023 |
| rs12113634 | T | 0.0088 | 0.0015 | 0.0004 | 0.0021 |
| rs12123293 | T | 0.0086 | 0.0015 | 0.0003 | 0.0020 |
| rs12151248 | T | -0.0149 | 0.0023 | -0.0055 | 0.0031 |
| rs12170452 | A | 0.0105 | 0.0014 | -0.0033 | 0.0020 |
| rs12234369 | A | 0.0087 | 0.0015 | -0.0048 | 0.0021 |
| rs12238011 | T | -0.0141 | 0.0026 | 0.0055 | 0.0036 |
| rs12342546 | A | -0.0128 | 0.0017 | 0.0009 | 0.0023 |
| rs12375949 | T | -0.0138 | 0.0014 | 0.0087 | 0.0020 |
| rs12405889 | T | -0.0098 | 0.0014 | -0.0056 | 0.0019 |
| rs12431682 | T | -0.0093 | 0.0015 | -0.0132 | 0.0020 |
| rs12438177 | A | 0.0093 | 0.0015 | -0.0006 | 0.0020 |
| rs12453682 | T | 0.0102 | 0.0015 | -0.0020 | 0.0022 |
| rs1245829 | A | -0.0099 | 0.0014 | -0.0040 | 0.0020 |
| rs12467175 | T | -0.0081 | 0.0014 | 0.0099 | 0.0019 |
| rs12477385 | T | 0.0109 | 0.0017 | -0.0144 | 0.0023 |
| rs12478156 | T | 0.0108 | 0.0015 | -0.0033 | 0.0020 |
| rs12503522 | T | -0.0113 | 0.0016 | 0.0037 | 0.0022 |
| rs12516990 | C | -0.0099 | 0.0017 | 0.0042 | 0.0025 |
| rs12519073 | T | -0.0107 | 0.0017 | 0.0042 | 0.0024 |
| rs12571549 | A | 0.0151 | 0.0020 | -0.0072 | 0.0028 |
| rs12601380 | A | -0.0082 | 0.0014 | 0.0158 | 0.0020 |
| rs12602286 | T | 0.0160 | 0.0021 | -0.0043 | 0.0030 |
| rs12613500 | C | 0.0098 | 0.0014 | -0.0082 | 0.0020 |
| rs12646523 | T | -0.0132 | 0.0016 | 0.0084 | 0.0022 |
| rs12670376 | A | 0.0092 | 0.0014 | -0.0063 | 0.0020 |
| rs12682775 | T | -0.0119 | 0.0017 | 0.0010 | 0.0024 |
| rs12789313 | T | 0.0092 | 0.0014 | -0.0035 | 0.0020 |
| rs12810587 | T | 0.0094 | 0.0015 | 0.0127 | 0.0021 |
| rs12891042 | T | 0.0092 | 0.0014 | 0.0039 | 0.0020 |
| rs12891191 | C | 0.0080 | 0.0014 | -0.0022 | 0.0020 |
| rs12952191 | T | -0.0081 | 0.0014 | 0.0073 | 0.0020 |
| rs12957463 | A | 0.0146 | 0.0018 | 0.0027 | 0.0024 |
| rs12962845 | C | -0.0123 | 0.0021 | 0.0055 | 0.0029 |
| rs13009915 | T | 0.0212 | 0.0037 | -0.0072 | 0.0052 |
| rs13010288 | T | 0.0198 | 0.0021 | -0.0080 | 0.0029 |
| rs13010566 | A | -0.0091 | 0.0014 | 0.0019 | 0.0019 |
| rs13015496 | C | 0.0119 | 0.0018 | 0.0015 | 0.0025 |
| rs13016316 | A | -0.0138 | 0.0024 | -0.0078 | 0.0034 |
| rs13029602 | T | -0.0092 | 0.0014 | 0.0060 | 0.0020 |
| rs13034349 | T | -0.0082 | 0.0014 | 0.0048 | 0.0020 |
| rs13050131 | A | -0.0081 | 0.0015 | 0.0014 | 0.0021 |
| rs13107325 | T | -0.0239 | 0.0027 | 0.0469 | 0.0037 |
| rs13117856 | A | 0.0084 | 0.0015 | 0.0022 | 0.0022 |
| rs13133213 | A | 0.0096 | 0.0014 | 0.0005 | 0.0019 |
| rs13163845 | T | -0.0149 | 0.0020 | -0.0015 | 0.0028 |
| rs13169187 | A | -0.0079 | 0.0014 | 0.0013 | 0.0020 |
| rs13177031 | A | -0.0085 | 0.0015 | -0.0077 | 0.0021 |
| rs13190235 | T | -0.0138 | 0.0024 | -0.0009 | 0.0034 |
| rs13212041 | T | 0.0101 | 0.0018 | -0.0071 | 0.0024 |
| rs13261773 | C | 0.0122 | 0.0019 | 0.0099 | 0.0027 |
| rs13284516 | C | -0.0082 | 0.0014 | -0.0019 | 0.0020 |
| rs1329044 | T | 0.0134 | 0.0019 | -0.0056 | 0.0026 |
| rs1329125 | T | -0.0097 | 0.0015 | 0.0036 | 0.0021 |
| rs13296345 | T | 0.0090 | 0.0015 | -0.0088 | 0.0020 |
| rs1334297 | A | 0.0257 | 0.0016 | -0.0113 | 0.0022 |
| rs13388333 | A | 0.0085 | 0.0015 | 0.0005 | 0.0020 |
| rs13397529 | C | -0.0097 | 0.0017 | 0.0098 | 0.0023 |
| rs13398860 | A | 0.0086 | 0.0015 | -0.0119 | 0.0020 |
| rs13422673 | T | -0.0110 | 0.0014 | -0.0006 | 0.0020 |
| rs1392816 | T | 0.0082 | 0.0014 | -0.0124 | 0.0020 |
| rs1404549 | A | -0.0089 | 0.0015 | 0.0026 | 0.0021 |
| rs1408430 | C | 0.0096 | 0.0016 | 0.0052 | 0.0022 |
| rs1426619 | T | 0.0096 | 0.0014 | 0.0023 | 0.0020 |
| rs1427298 | T | 0.0086 | 0.0014 | -0.0058 | 0.0020 |
| rs1427829 | A | -0.0083 | 0.0014 | 0.0137 | 0.0020 |
| rs1434630 | T | -0.0137 | 0.0020 | 0.0106 | 0.0027 |
| rs1464297 | T | -0.0109 | 0.0015 | 0.0029 | 0.0021 |
| rs1467737 | T | -0.0084 | 0.0014 | 0.0017 | 0.0020 |
| rs1485300 | C | 0.0098 | 0.0015 | 0.0041 | 0.0021 |
| rs1490612 | T | 0.0193 | 0.0026 | 0.0054 | 0.0037 |
| rs1527878 | A | -0.0105 | 0.0016 | 0.0058 | 0.0023 |
| rs1529597 | A | 0.0223 | 0.0038 | -0.0122 | 0.0058 |
| rs1538389 | T | -0.0109 | 0.0018 | 0.0097 | 0.0025 |
| rs1542354 | A | -0.0084 | 0.0014 | 0.0075 | 0.0020 |
| rs1544 | A | -0.0096 | 0.0016 | 0.0063 | 0.0022 |
| rs1564347 | T | 0.0088 | 0.0015 | -0.0073 | 0.0021 |
| rs1569092 | A | 0.0160 | 0.0020 | 0.0009 | 0.0027 |
| rs1603460 | T | 0.0090 | 0.0014 | 0.0009 | 0.0020 |
| rs162445 | A | 0.0152 | 0.0026 | 0.0032 | 0.0037 |
| rs1637770 | T | 0.0191 | 0.0028 | -0.0133 | 0.0041 |
| rs164938 | T | -0.0088 | 0.0014 | 0.0078 | 0.0020 |
| rs1671269 | T | -0.0100 | 0.0016 | -0.0042 | 0.0023 |
| rs16851779 | T | -0.0153 | 0.0027 | 0.0000 | 0.0035 |
| rs16901689 | T | 0.0111 | 0.0019 | -0.0021 | 0.0026 |
| rs1693584 | T | -0.0087 | 0.0015 | 0.0028 | 0.0021 |
| rs16958559 | T | 0.0096 | 0.0017 | -0.0069 | 0.0023 |
| rs16966271 | T | 0.0092 | 0.0016 | 0.0011 | 0.0022 |
| rs16975275 | A | -0.0102 | 0.0017 | 0.0004 | 0.0024 |
| rs17048801 | A | -0.0106 | 0.0014 | 0.0012 | 0.0021 |
| rs17048855 | A | 0.0108 | 0.0015 | -0.0010 | 0.0020 |
| rs17069646 | T | 0.0085 | 0.0015 | 0.0074 | 0.0021 |
| rs17110109 | T | -0.0093 | 0.0014 | 0.0124 | 0.0020 |
| rs17113730 | A | -0.0170 | 0.0027 | -0.0122 | 0.0038 |
| rs17133297 | A | -0.0146 | 0.0023 | 0.0007 | 0.0034 |
| rs17144467 | A | 0.0098 | 0.0015 | -0.0093 | 0.0021 |
| rs17148998 | A | 0.0101 | 0.0017 | -0.0060 | 0.0024 |
| rs17170519 | C | 0.0132 | 0.0023 | -0.0054 | 0.0032 |
| rs17190418 | T | -0.0213 | 0.0031 | -0.0028 | 0.0042 |
| rs17205908 | T | -0.0092 | 0.0015 | 0.0068 | 0.0021 |
| rs17248751 | A | -0.0136 | 0.0017 | 0.0024 | 0.0023 |
| rs1728118 | A | 0.0089 | 0.0016 | -0.0078 | 0.0022 |
| rs1729412 | T | -0.0107 | 0.0014 | -0.0058 | 0.0020 |
| rs1734370 | A | 0.0099 | 0.0016 | 0.0012 | 0.0021 |
| rs1738050 | C | -0.0093 | 0.0014 | -0.0003 | 0.0020 |
| rs17428076 | C | 0.0137 | 0.0016 | -0.0126 | 0.0023 |
| rs17440885 | A | 0.0160 | 0.0029 | -0.0125 | 0.0040 |
| rs1747817 | T | -0.0092 | 0.0016 | 0.0022 | 0.0023 |
| rs17489649 | A | 0.0127 | 0.0015 | -0.0018 | 0.0021 |
| rs17502934 | T | -0.0167 | 0.0020 | 0.0116 | 0.0028 |
| rs17536059 | C | -0.0108 | 0.0019 | 0.0075 | 0.0026 |
| rs17551064 | A | 0.0141 | 0.0019 | -0.0045 | 0.0027 |
| rs17563464 | A | -0.0147 | 0.0017 | 0.0015 | 0.0024 |
| rs17565975 | A | -0.0106 | 0.0014 | -0.0010 | 0.0020 |
| rs17568389 | A | 0.0127 | 0.0014 | -0.0025 | 0.0020 |
| rs17574007 | A | -0.0129 | 0.0021 | 0.0122 | 0.0031 |
| rs1758747 | A | 0.0092 | 0.0015 | 0.0005 | 0.0021 |
| rs17598675 | T | -0.0116 | 0.0014 | -0.0011 | 0.0020 |
| rs17604349 | A | 0.0127 | 0.0018 | 0.0076 | 0.0026 |
| rs17609255 | T | -0.0087 | 0.0014 | 0.0025 | 0.0020 |
| rs176218 | T | 0.0200 | 0.0018 | -0.0154 | 0.0025 |
| rs17650634 | T | -0.0104 | 0.0019 | -0.0012 | 0.0026 |
| rs17667540 | A | -0.0087 | 0.0015 | 0.0126 | 0.0021 |
| rs17669337 | T | -0.0110 | 0.0014 | -0.0048 | 0.0020 |
| rs17686649 | T | -0.0098 | 0.0017 | -0.0049 | 0.0024 |
| rs17732878 | T | -0.0095 | 0.0017 | 0.0080 | 0.0024 |
| rs17742342 | A | -0.0119 | 0.0018 | -0.0057 | 0.0024 |
| rs17747544 | A | -0.0090 | 0.0014 | 0.0089 | 0.0020 |
| rs1779549 | A | 0.0081 | 0.0014 | -0.0026 | 0.0020 |
| rs1792602 | A | -0.0097 | 0.0014 | 0.0041 | 0.0020 |
| rs1861786 | A | -0.0081 | 0.0014 | -0.0004 | 0.0020 |
| rs1865955 | T | 0.0122 | 0.0019 | -0.0126 | 0.0026 |
| rs1866823 | A | 0.0107 | 0.0014 | -0.0015 | 0.0020 |
| rs187580 | T | -0.0104 | 0.0017 | -0.0046 | 0.0023 |
| rs1880692 | A | 0.0086 | 0.0014 | 0.0037 | 0.0020 |
| rs1890132 | T | -0.0089 | 0.0015 | 0.0024 | 0.0021 |
| rs1898111 | A | 0.0101 | 0.0018 | -0.0062 | 0.0026 |
| rs1910005 | T | -0.0090 | 0.0015 | 0.0021 | 0.0021 |
| rs1918394 | T | 0.0119 | 0.0019 | -0.0034 | 0.0026 |
| rs1933264 | T | 0.0100 | 0.0016 | -0.0129 | 0.0023 |
| rs1952183 | A | -0.0089 | 0.0014 | 0.0031 | 0.0020 |
| rs1955250 | A | -0.0140 | 0.0025 | 0.0033 | 0.0035 |
| rs1991585 | T | -0.0101 | 0.0015 | 0.0074 | 0.0021 |
| rs2011603 | A | -0.0091 | 0.0016 | 0.0028 | 0.0022 |
| rs2014830 | T | 0.0110 | 0.0015 | -0.0209 | 0.0021 |
| rs2023016 | C | 0.0107 | 0.0017 | 0.0000 | 0.0024 |
| rs2029401 | A | -0.0091 | 0.0014 | 0.0012 | 0.0020 |
| rs2034631 | T | 0.0089 | 0.0015 | -0.0003 | 0.0021 |
| rs2039204 | A | 0.0082 | 0.0014 | -0.0012 | 0.0020 |
| rs2055940 | A | 0.0082 | 0.0015 | 0.0010 | 0.0021 |
| rs2098526 | A | -0.0241 | 0.0043 | 0.0150 | 0.0060 |
| rs2179152 | T | -0.0131 | 0.0015 | 0.0017 | 0.0020 |
| rs2183271 | T | 0.0084 | 0.0015 | -0.0183 | 0.0020 |
| rs2199409 | T | 0.0103 | 0.0017 | -0.0051 | 0.0025 |
| rs2212430 | T | -0.0100 | 0.0016 | 0.0083 | 0.0022 |
| rs2216144 | T | 0.0098 | 0.0014 | 0.0017 | 0.0020 |
| rs2220926 | T | -0.0095 | 0.0014 | -0.0038 | 0.0020 |
| rs2252098 | T | -0.0077 | 0.0014 | 0.0044 | 0.0020 |
| rs2256965 | A | 0.0106 | 0.0014 | -0.0061 | 0.0020 |
| rs2276209 | A | 0.0085 | 0.0015 | 0.0037 | 0.0022 |
| rs2283076 | A | 0.0115 | 0.0017 | -0.0043 | 0.0023 |
| rs2287838 | A | -0.0104 | 0.0014 | -0.0030 | 0.0020 |
| rs2290601 | T | -0.0149 | 0.0017 | 0.0098 | 0.0023 |
| rs2297293 | C | 0.0099 | 0.0015 | -0.0001 | 0.0021 |
| rs2299156 | T | 0.0099 | 0.0018 | 0.0022 | 0.0026 |
| rs2321157 | A | -0.0081 | 0.0014 | 0.0056 | 0.0020 |
| rs232496 | T | 0.0090 | 0.0015 | 0.0004 | 0.0021 |
| rs2358628 | A | -0.0091 | 0.0015 | 0.0052 | 0.0021 |
| rs2365376 | A | 0.0092 | 0.0015 | -0.0041 | 0.0021 |
| rs2373124 | A | -0.0096 | 0.0016 | -0.0037 | 0.0023 |
| rs2406253 | A | 0.0135 | 0.0018 | -0.0038 | 0.0025 |
| rs2416214 | A | 0.0079 | 0.0014 | -0.0072 | 0.0020 |
| rs2416845 | T | 0.0263 | 0.0048 | -0.0141 | 0.0068 |
| rs2436760 | T | 0.0232 | 0.0042 | -0.0090 | 0.0057 |
| rs2447097 | T | 0.0096 | 0.0014 | -0.0026 | 0.0020 |
| rs2470966 | C | -0.0103 | 0.0015 | 0.0045 | 0.0021 |
| rs2478208 | C | -0.0103 | 0.0014 | 0.0016 | 0.0020 |
| rs2496482 | T | 0.0121 | 0.0015 | -0.0030 | 0.0020 |
| rs2517086 | C | -0.0088 | 0.0014 | 0.0018 | 0.0020 |
| rs2522545 | T | -0.0083 | 0.0014 | -0.0023 | 0.0020 |
| rs252991 | A | 0.0095 | 0.0015 | -0.0046 | 0.0020 |
| rs2570497 | T | -0.0121 | 0.0015 | 0.0115 | 0.0020 |
| rs2718277 | T | 0.0142 | 0.0026 | -0.0080 | 0.0035 |
| rs27220 | A | -0.0114 | 0.0015 | -0.0016 | 0.0020 |
| rs2736752 | T | 0.0095 | 0.0017 | -0.0058 | 0.0024 |
| rs2764684 | T | 0.0143 | 0.0019 | 0.0058 | 0.0026 |
| rs2805064 | C | 0.0093 | 0.0016 | 0.0015 | 0.0022 |
| rs2838006 | T | 0.0132 | 0.0015 | -0.0125 | 0.0021 |
| rs2852349 | T | -0.0093 | 0.0014 | 0.0046 | 0.0020 |
| rs2885198 | A | 0.0088 | 0.0014 | -0.0042 | 0.0020 |
| rs2926702 | T | -0.0124 | 0.0021 | -0.0027 | 0.0030 |
| rs2929032 | A | -0.0081 | 0.0014 | 0.0046 | 0.0020 |
| rs2929860 | T | 0.0110 | 0.0018 | -0.0048 | 0.0025 |
| rs2942884 | A | -0.0090 | 0.0014 | 0.0041 | 0.0020 |
| rs2958182 | A | 0.0082 | 0.0015 | 0.0009 | 0.0021 |
| rs2964255 | A | 0.0088 | 0.0015 | 0.0004 | 0.0021 |
| rs2976397 | T | 0.0081 | 0.0014 | -0.0080 | 0.0020 |
| rs2977464 | T | 0.0102 | 0.0018 | -0.0010 | 0.0025 |
| rs29792 | A | -0.0093 | 0.0015 | 0.0033 | 0.0021 |
| rs2980813 | A | 0.0088 | 0.0014 | 0.0011 | 0.0020 |
| rs2989476 | C | 0.0078 | 0.0014 | -0.0033 | 0.0020 |
| rs3111251 | T | -0.0090 | 0.0014 | 0.0014 | 0.0020 |
| rs312927 | A | -0.0147 | 0.0022 | 0.0033 | 0.0031 |
| rs322614 | A | -0.0087 | 0.0014 | -0.0002 | 0.0020 |
| rs339057 | A | -0.0081 | 0.0014 | 0.0019 | 0.0020 |
| rs341504 | A | 0.0099 | 0.0015 | -0.0034 | 0.0021 |
| rs34309 | A | 0.0080 | 0.0014 | 0.0069 | 0.0020 |
| rs34410 | C | -0.0083 | 0.0014 | 0.0090 | 0.0020 |
| rs347661 | T | -0.0081 | 0.0014 | 0.0084 | 0.0020 |
| rs350281 | T | -0.0117 | 0.0015 | 0.0027 | 0.0021 |
| rs356999 | A | -0.0101 | 0.0014 | 0.0080 | 0.0020 |
| rs363096 | T | -0.0143 | 0.0014 | 0.0043 | 0.0020 |
| rs3735478 | T | 0.0108 | 0.0016 | -0.0124 | 0.0021 |
| rs3768480 | C | 0.0101 | 0.0014 | -0.0042 | 0.0020 |
| rs3781339 | T | -0.0104 | 0.0018 | 0.0054 | 0.0025 |
| rs3790609 | T | 0.0107 | 0.0019 | -0.0043 | 0.0026 |
| rs3809634 | A | -0.0109 | 0.0015 | -0.0050 | 0.0022 |
| rs3812281 | T | 0.0113 | 0.0014 | 0.0034 | 0.0020 |
| rs382196 | T | -0.0082 | 0.0014 | -0.0032 | 0.0020 |
| rs3847228 | T | 0.0083 | 0.0014 | -0.0028 | 0.0020 |
| rs3848715 | A | -0.0080 | 0.0014 | -0.0037 | 0.0020 |
| rs3859523 | T | 0.0124 | 0.0018 | -0.0027 | 0.0026 |
| rs3895736 | A | 0.0144 | 0.0019 | 0.0050 | 0.0026 |
| rs3897821 | A | 0.0154 | 0.0015 | -0.0058 | 0.0021 |
| rs3948495 | T | -0.0089 | 0.0014 | 0.0091 | 0.0020 |
| rs399821 | A | -0.0078 | 0.0014 | 0.0037 | 0.0020 |
| rs39998 | A | 0.0090 | 0.0015 | 0.0008 | 0.0022 |
| rs4127499 | A | 0.0087 | 0.0015 | -0.0042 | 0.0020 |
| rs42302 | A | 0.0086 | 0.0015 | -0.0268 | 0.0021 |
| rs4255791 | A | -0.0084 | 0.0015 | 0.0057 | 0.0021 |
| rs429150 | T | 0.0085 | 0.0014 | 0.0002 | 0.0020 |
| rs4298514 | T | 0.0104 | 0.0015 | 0.0091 | 0.0021 |
| rs4320563 | A | 0.0109 | 0.0014 | -0.0062 | 0.0020 |
| rs4328757 | T | 0.0096 | 0.0014 | -0.0058 | 0.0020 |
| rs4358081 | A | -0.0095 | 0.0014 | -0.0098 | 0.0019 |
| rs4396896 | A | -0.0107 | 0.0014 | 0.0137 | 0.0020 |
| rs4423373 | A | -0.0087 | 0.0014 | 0.0007 | 0.0020 |
| rs4434676 | A | -0.0089 | 0.0014 | 0.0005 | 0.0020 |
| rs4458044 | C | 0.0101 | 0.0016 | -0.0027 | 0.0023 |
| rs4480339 | A | -0.0097 | 0.0015 | -0.0052 | 0.0021 |
| rs4490539 | A | 0.0142 | 0.0015 | 0.0060 | 0.0021 |
| rs4500930 | T | -0.0109 | 0.0015 | 0.0168 | 0.0020 |
| rs4500960 | T | -0.0131 | 0.0014 | -0.0045 | 0.0020 |
| rs4502401 | T | 0.0085 | 0.0015 | -0.0008 | 0.0021 |
| rs4658019 | T | 0.0084 | 0.0014 | -0.0077 | 0.0020 |
| rs4673840 | T | -0.0138 | 0.0019 | 0.0078 | 0.0027 |
| rs4712371 | A | -0.0124 | 0.0021 | 0.0051 | 0.0030 |
| rs4719944 | T | -0.0121 | 0.0014 | -0.0058 | 0.0020 |
| rs4724083 | T | 0.0083 | 0.0015 | 0.0071 | 0.0021 |
| rs4726070 | A | 0.0122 | 0.0014 | -0.0043 | 0.0020 |
| rs4731413 | A | 0.0111 | 0.0017 | -0.0072 | 0.0024 |
| rs4735297 | A | -0.0086 | 0.0015 | 0.0126 | 0.0021 |
| rs4739235 | A | 0.0109 | 0.0019 | -0.0107 | 0.0026 |
| rs4757957 | C | 0.0133 | 0.0015 | -0.0031 | 0.0021 |
| rs4766424 | C | 0.0142 | 0.0021 | -0.0111 | 0.0029 |
| rs4785187 | A | -0.0100 | 0.0017 | 0.0032 | 0.0024 |
| rs4787028 | T | 0.0086 | 0.0015 | 0.0000 | 0.0021 |
| rs481940 | T | 0.0100 | 0.0016 | 0.0062 | 0.0022 |
| rs482787 | T | 0.0100 | 0.0015 | -0.0144 | 0.0021 |
| rs4839155 | T | 0.0115 | 0.0017 | -0.0059 | 0.0023 |
| rs4846010 | A | 0.0099 | 0.0018 | 0.0000 | 0.0025 |
| rs4846724 | A | 0.0098 | 0.0014 | -0.0072 | 0.0020 |
| rs4851263 | A | -0.0151 | 0.0023 | 0.0065 | 0.0032 |
| rs4877516 | A | -0.0101 | 0.0014 | 0.0046 | 0.0020 |
| rs4881269 | A | 0.0100 | 0.0014 | -0.0042 | 0.0020 |
| rs4904523 | A | -0.0082 | 0.0014 | 0.0047 | 0.0020 |
| rs4919624 | A | 0.0193 | 0.0018 | 0.0067 | 0.0024 |
| rs4925065 | T | -0.0078 | 0.0014 | -0.0032 | 0.0020 |
| rs4925109 | A | -0.0096 | 0.0015 | -0.0054 | 0.0021 |
| rs4938815 | T | 0.0085 | 0.0015 | 0.0019 | 0.0021 |
| rs4941735 | T | 0.0092 | 0.0014 | -0.0006 | 0.0020 |
| rs4972748 | T | 0.0105 | 0.0018 | -0.0010 | 0.0025 |
| rs4984541 | A | -0.0138 | 0.0017 | 0.0048 | 0.0023 |
| rs4984613 | T | 0.0174 | 0.0028 | -0.0004 | 0.0040 |
| rs4984682 | C | -0.0138 | 0.0017 | 0.0045 | 0.0023 |
| rs548897 | A | 0.0081 | 0.0014 | -0.0013 | 0.0020 |
| rs563954 | A | -0.0081 | 0.0014 | -0.0023 | 0.0020 |
| rs567003 | A | 0.0094 | 0.0015 | -0.0002 | 0.0021 |
| rs5754581 | T | -0.0082 | 0.0014 | 0.0057 | 0.0020 |
| rs5754762 | A | 0.0197 | 0.0030 | -0.0073 | 0.0043 |
| rs6020560 | T | -0.0083 | 0.0014 | 0.0037 | 0.0020 |
| rs6065080 | T | -0.0130 | 0.0015 | -0.0052 | 0.0021 |
| rs6091570 | A | 0.0081 | 0.0015 | -0.0119 | 0.0021 |
| rs613872 | T | -0.0163 | 0.0019 | 0.0141 | 0.0026 |
| rs631287 | A | 0.0084 | 0.0014 | -0.0038 | 0.0020 |
| rs635754 | A | 0.0119 | 0.0014 | 0.0081 | 0.0020 |
| rs6428587 | T | 0.0084 | 0.0015 | 0.0004 | 0.0020 |
| rs6435326 | A | -0.0091 | 0.0014 | 0.0017 | 0.0020 |
| rs6442126 | A | -0.0121 | 0.0015 | 0.0047 | 0.0021 |
| rs6445633 | A | 0.0088 | 0.0015 | -0.0012 | 0.0021 |
| rs6461536 | A | 0.0088 | 0.0015 | 0.0002 | 0.0022 |
| rs6469654 | C | -0.0103 | 0.0017 | 0.0011 | 0.0024 |
| rs6480234 | T | -0.0084 | 0.0014 | 0.0040 | 0.0020 |
| rs6493265 | T | -0.0122 | 0.0014 | 0.0011 | 0.0020 |
| rs6493275 | A | 0.0131 | 0.0017 | 0.0063 | 0.0024 |
| rs6503409 | T | 0.0087 | 0.0015 | -0.0067 | 0.0021 |
| rs6534338 | T | 0.0103 | 0.0015 | -0.0031 | 0.0022 |
| rs6557171 | T | -0.0155 | 0.0015 | 0.0016 | 0.0021 |
| rs6567288 | A | -0.0079 | 0.0014 | 0.0061 | 0.0020 |
| rs6689641 | A | 0.0078 | 0.0014 | 0.0030 | 0.0020 |
| rs6695132 | T | 0.0097 | 0.0017 | -0.0069 | 0.0024 |
| rs6696068 | T | -0.0088 | 0.0014 | 0.0044 | 0.0020 |
| rs6704768 | A | -0.0116 | 0.0014 | -0.0008 | 0.0020 |
| rs6706275 | T | 0.0083 | 0.0015 | -0.0035 | 0.0021 |
| rs6715321 | T | -0.0097 | 0.0014 | 0.0048 | 0.0020 |
| rs6720515 | A | 0.0103 | 0.0017 | -0.0092 | 0.0024 |
| rs6729612 | T | 0.0092 | 0.0015 | 0.0002 | 0.0021 |
| rs6731373 | A | -0.0115 | 0.0015 | 0.0032 | 0.0020 |
| rs6736025 | T | 0.0079 | 0.0014 | -0.0156 | 0.0020 |
| rs6743032 | A | -0.0165 | 0.0024 | 0.0027 | 0.0033 |
| rs6744040 | A | -0.0080 | 0.0014 | 0.0049 | 0.0020 |
| rs6752228 | T | -0.0080 | 0.0014 | 0.0061 | 0.0020 |
| rs6757087 | T | -0.0085 | 0.0014 | 0.0034 | 0.0020 |
| rs6780414 | T | 0.0105 | 0.0018 | -0.0003 | 0.0025 |
| rs6792805 | A | -0.0090 | 0.0016 | -0.0016 | 0.0022 |
| rs6824567 | T | 0.0130 | 0.0016 | -0.0081 | 0.0022 |
| rs6861925 | C | -0.0090 | 0.0014 | -0.0007 | 0.0020 |
| rs6871635 | A | -0.0087 | 0.0014 | 0.0084 | 0.0020 |
| rs6881733 | T | -0.0086 | 0.0016 | -0.0082 | 0.0022 |
| rs6917154 | T | 0.0127 | 0.0021 | -0.0081 | 0.0029 |
| rs6917204 | T | 0.0119 | 0.0017 | 0.0026 | 0.0025 |
| rs6918506 | C | 0.0113 | 0.0014 | -0.0032 | 0.0020 |
| rs6932108 | C | -0.0210 | 0.0027 | 0.0014 | 0.0035 |
| rs6946136 | A | 0.0081 | 0.0014 | 0.0054 | 0.0020 |
| rs6969783 | A | -0.0098 | 0.0014 | 0.0026 | 0.0020 |
| rs6977237 | T | 0.0082 | 0.0014 | -0.0063 | 0.0020 |
| rs6994287 | A | 0.0102 | 0.0014 | 0.0005 | 0.0020 |
| rs7016302 | C | -0.0125 | 0.0019 | -0.0014 | 0.0026 |
| rs7030373 | A | -0.0098 | 0.0017 | 0.0022 | 0.0024 |
| rs7035315 | A | -0.0090 | 0.0015 | -0.0039 | 0.0020 |
| rs7041702 | A | -0.0111 | 0.0016 | -0.0024 | 0.0022 |
| rs7108020 | A | 0.0111 | 0.0015 | -0.0014 | 0.0021 |
| rs7127580 | C | -0.0150 | 0.0023 | 0.0126 | 0.0031 |
| rs713584 | A | -0.0080 | 0.0014 | 0.0010 | 0.0020 |
| rs7158218 | A | -0.0090 | 0.0015 | 0.0033 | 0.0021 |
| rs716513 | A | 0.0084 | 0.0014 | 0.0043 | 0.0020 |
| rs7171405 | A | -0.0101 | 0.0016 | 0.0062 | 0.0023 |
| rs717996 | T | -0.0122 | 0.0014 | 0.0048 | 0.0020 |
| rs7182216 | C | -0.0096 | 0.0017 | 0.0017 | 0.0023 |
| rs721579 | T | -0.0110 | 0.0016 | 0.0051 | 0.0022 |
| rs7226824 | T | -0.0078 | 0.0014 | 0.0060 | 0.0020 |
| rs7254263 | T | -0.0119 | 0.0016 | 0.0015 | 0.0022 |
| rs7255223 | A | 0.0097 | 0.0016 | -0.0110 | 0.0022 |
| rs728054 | A | -0.0128 | 0.0015 | 0.0060 | 0.0021 |
| rs730384 | A | 0.0101 | 0.0014 | -0.0020 | 0.0020 |
| rs7321274 | A | 0.0114 | 0.0018 | 0.0027 | 0.0024 |
| rs7323027 | A | 0.0130 | 0.0015 | 0.0041 | 0.0021 |
| rs736281 | T | 0.0086 | 0.0014 | -0.0038 | 0.0020 |
| rs736471 | T | 0.0083 | 0.0014 | 0.0043 | 0.0020 |
| rs737945 | C | -0.0115 | 0.0014 | -0.0022 | 0.0020 |
| rs743316 | T | 0.0112 | 0.0017 | -0.0034 | 0.0024 |
| rs7560871 | A | -0.0193 | 0.0027 | 0.0236 | 0.0038 |
| rs7561705 | A | 0.0078 | 0.0014 | 0.0093 | 0.0019 |
| rs7575637 | A | 0.0107 | 0.0014 | -0.0028 | 0.0020 |
| rs7597412 | C | 0.0088 | 0.0015 | -0.0021 | 0.0020 |
| rs7625428 | T | 0.0098 | 0.0014 | -0.0007 | 0.0020 |
| rs763553 | A | 0.0087 | 0.0014 | -0.0054 | 0.0020 |
| rs7640424 | T | 0.0113 | 0.0015 | -0.0128 | 0.0021 |
| rs7672622 | A | -0.0088 | 0.0016 | 0.0001 | 0.0023 |
| rs767943 | A | -0.0122 | 0.0016 | 0.0076 | 0.0023 |
| rs7683416 | T | 0.0139 | 0.0014 | -0.0001 | 0.0020 |
| rs7692359 | T | -0.0109 | 0.0017 | 0.0009 | 0.0024 |
| rs7772172 | A | 0.0091 | 0.0014 | 0.0001 | 0.0020 |
| rs7803932 | A | 0.0127 | 0.0019 | -0.0002 | 0.0026 |
| rs780569 | A | -0.0085 | 0.0016 | 0.0004 | 0.0022 |
| rs7849480 | A | -0.0084 | 0.0014 | 0.0081 | 0.0020 |
| rs7849487 | T | -0.0167 | 0.0015 | -0.0013 | 0.0021 |
| rs7855503 | C | 0.0088 | 0.0015 | 0.0020 | 0.0020 |
| rs7875078 | A | -0.0078 | 0.0014 | 0.0061 | 0.0020 |
| rs7894722 | T | 0.0085 | 0.0014 | -0.0019 | 0.0020 |
| rs7899270 | A | -0.0084 | 0.0015 | 0.0030 | 0.0021 |
| rs790647 | A | -0.0145 | 0.0017 | 0.0007 | 0.0023 |
| rs7910403 | T | 0.0122 | 0.0018 | -0.0057 | 0.0025 |
| rs7920624 | A | 0.0120 | 0.0014 | -0.0050 | 0.0020 |
| rs7931563 | T | -0.0109 | 0.0014 | 0.0016 | 0.0020 |
| rs795230 | T | 0.0081 | 0.0014 | 0.0024 | 0.0020 |
| rs7958371 | A | -0.0085 | 0.0015 | 0.0053 | 0.0021 |
| rs7965154 | A | -0.0085 | 0.0015 | 0.0033 | 0.0021 |
| rs7974852 | A | 0.0117 | 0.0014 | -0.0075 | 0.0020 |
| rs7977614 | A | -0.0134 | 0.0016 | -0.0001 | 0.0021 |
| rs8009933 | A | 0.0088 | 0.0015 | 0.0033 | 0.0021 |
| rs8024 | A | -0.0115 | 0.0015 | 0.0237 | 0.0021 |
| rs8052523 | T | -0.0084 | 0.0014 | 0.0069 | 0.0020 |
| rs8097125 | T | 0.0086 | 0.0014 | -0.0036 | 0.0020 |
| rs852771 | T | 0.0092 | 0.0015 | -0.0034 | 0.0021 |
| rs870589 | A | -0.0079 | 0.0014 | 0.0009 | 0.0020 |
| rs884108 | A | -0.0094 | 0.0017 | 0.0006 | 0.0023 |
| rs893522 | A | 0.0149 | 0.0025 | 0.0019 | 0.0034 |
| rs902820 | A | -0.0096 | 0.0014 | 0.0016 | 0.0020 |
| rs911149 | T | 0.0091 | 0.0017 | -0.0033 | 0.0023 |
| rs912883 | T | 0.0087 | 0.0015 | 0.0031 | 0.0021 |
| rs913509 | A | 0.0096 | 0.0015 | -0.0077 | 0.0021 |
| rs9267658 | T | 0.0138 | 0.0020 | -0.0083 | 0.0027 |
| rs9267677 | T | 0.0180 | 0.0024 | -0.0236 | 0.0033 |
| rs9291437 | C | -0.0081 | 0.0014 | -0.0002 | 0.0020 |
| rs9294770 | T | 0.0083 | 0.0014 | -0.0059 | 0.0020 |
| rs9300612 | T | 0.0084 | 0.0015 | -0.0005 | 0.0021 |
| rs9349956 | A | -0.0159 | 0.0018 | -0.0016 | 0.0025 |
| rs9373363 | A | -0.0111 | 0.0016 | 0.0064 | 0.0023 |
| rs9375403 | C | 0.0126 | 0.0016 | -0.0090 | 0.0023 |
| rs9386110 | T | -0.0163 | 0.0023 | -0.0030 | 0.0032 |
| rs9388490 | T | 0.0091 | 0.0014 | -0.0034 | 0.0020 |
| rs939400 | T | -0.0095 | 0.0015 | 0.0050 | 0.0020 |
| rs9411331 | A | 0.0145 | 0.0015 | -0.0106 | 0.0021 |
| rs9442750 | A | -0.0089 | 0.0016 | -0.0043 | 0.0022 |
| rs9490512 | A | -0.0136 | 0.0014 | 0.0069 | 0.0020 |
| rs9492774 | C | 0.0080 | 0.0015 | -0.0039 | 0.0020 |
| rs9513416 | A | -0.0116 | 0.0019 | 0.0084 | 0.0027 |
| rs9527905 | A | -0.0094 | 0.0014 | 0.0104 | 0.0020 |
| rs9540718 | A | -0.0089 | 0.0014 | 0.0067 | 0.0020 |
| rs9545395 | T | 0.0138 | 0.0021 | -0.0026 | 0.0029 |
| rs9556958 | T | -0.0110 | 0.0014 | 0.0100 | 0.0020 |
| rs9563168 | A | 0.0101 | 0.0017 | -0.0094 | 0.0024 |
| rs9568798 | T | -0.0093 | 0.0016 | -0.0055 | 0.0022 |
| rs9597907 | T | 0.0152 | 0.0027 | -0.0089 | 0.0039 |
| rs9611597 | A | -0.0107 | 0.0019 | 0.0108 | 0.0027 |
| rs9616906 | A | 0.0122 | 0.0014 | 0.0035 | 0.0020 |
| rs9649 | T | 0.0108 | 0.0019 | -0.0040 | 0.0026 |
| rs9655780 | A | -0.0153 | 0.0019 | 0.0031 | 0.0027 |
| rs9666728 | T | -0.0082 | 0.0014 | 0.0078 | 0.0020 |
| rs9771228 | T | 0.0096 | 0.0015 | -0.0116 | 0.0020 |
| rs977143 | A | 0.0085 | 0.0015 | 0.0038 | 0.0020 |
| rs978807 | A | 0.0139 | 0.0018 | -0.0092 | 0.0025 |
| rs981230 | T | 0.0100 | 0.0014 | -0.0073 | 0.0020 |
| rs981883 | A | 0.0099 | 0.0014 | -0.0022 | 0.0020 |
| rs9820604 | T | 0.0115 | 0.0018 | 0.0021 | 0.0026 |
| rs9853928 | T | -0.0127 | 0.0017 | 0.0063 | 0.0024 |
| rs9858921 | A | -0.0087 | 0.0014 | 0.0004 | 0.0020 |
| rs9870317 | A | 0.0086 | 0.0015 | -0.0048 | 0.0021 |
| rs9882532 | T | 0.0124 | 0.0015 | -0.0033 | 0.0021 |
| rs9886703 | A | -0.0136 | 0.0019 | 0.0063 | 0.0026 |
| rs9926649 | T | 0.0177 | 0.0031 | -0.0116 | 0.0045 |
| rs9927842 | T | -0.0130 | 0.0020 | 0.0093 | 0.0027 |
| rs9929762 | A | 0.0096 | 0.0014 | -0.0030 | 0.0020 |
| rs9929993 | T | -0.0102 | 0.0015 | 0.0055 | 0.0021 |
| rs995698 | A | -0.0099 | 0.0014 | -0.0035 | 0.0020 |

#### Supplementary table 11: Genetic association estimates for the IVW ratio-method MR analysis estimating the effect of education on CAD. ea=effect allele; gx=education; gy=CAD; se=standard error

| SNP | ea | gx | gx_se | gy | gy_se |
| --- | --- | --- | --- | --- | --- |
| rs10006235 | T | -0.0093 | 0.0016 | 0.0000 | 0.0101 |
| rs10009513 | A | 0.0088 | 0.0014 | -0.0071 | 0.0093 |
| rs10021733 | T | -0.0164 | 0.0027 | 0.0039 | 0.0141 |
| rs10032941 | T | -0.0080 | 0.0015 | 0.0149 | 0.0095 |
| rs10041403 | T | 0.0151 | 0.0019 | -0.0031 | 0.0131 |
| rs10042828 | A | -0.0145 | 0.0022 | 0.0149 | 0.0158 |
| rs10046069 | A | 0.0127 | 0.0022 | 0.0008 | 0.0141 |
| rs10057590 | A | 0.0106 | 0.0014 | -0.0050 | 0.0095 |
| rs10060023 | T | 0.0110 | 0.0015 | -0.0028 | 0.0104 |
| rs10061420 | A | 0.0082 | 0.0014 | 0.0014 | 0.0095 |
| rs1006749 | A | 0.0079 | 0.0014 | -0.0042 | 0.0097 |
| rs10074178 | A | 0.0079 | 0.0014 | 0.0011 | 0.0094 |
| rs1007731 | A | -0.0143 | 0.0022 | 0.0114 | 0.0174 |
| rs10080647 | A | 0.0124 | 0.0020 | -0.0121 | 0.0125 |
| rs1008078 | T | -0.0182 | 0.0014 | 0.0027 | 0.0096 |
| rs1009470 | T | 0.0086 | 0.0014 | -0.0134 | 0.0092 |
| rs10098073 | A | -0.0091 | 0.0014 | 0.0002 | 0.0098 |
| rs10122669 | A | -0.0079 | 0.0014 | -0.0017 | 0.0094 |
| rs10128888 | A | -0.0083 | 0.0015 | 0.0217 | 0.0105 |
| rs10145520 | T | -0.0122 | 0.0018 | -0.0095 | 0.0119 |
| rs10166286 | T | 0.0151 | 0.0019 | -0.0065 | 0.0116 |
| rs10169002 | A | -0.0080 | 0.0014 | -0.0030 | 0.0097 |
| rs10181071 | A | 0.0102 | 0.0019 | 0.0033 | 0.0132 |
| rs10189857 | A | 0.0158 | 0.0014 | -0.0216 | 0.0097 |
| rs10192369 | A | -0.0078 | 0.0014 | 0.0024 | 0.0099 |
| rs10192834 | T | 0.0083 | 0.0015 | 0.0021 | 0.0101 |
| rs10193498 | A | 0.0097 | 0.0016 | 0.0031 | 0.0108 |
| rs10204051 | T | -0.0101 | 0.0014 | 0.0043 | 0.0101 |
| rs10205057 | T | 0.0090 | 0.0014 | -0.0110 | 0.0095 |
| rs10208 | T | 0.0091 | 0.0015 | -0.0022 | 0.0100 |
| rs10215082 | A | -0.0149 | 0.0014 | 0.0148 | 0.0098 |
| rs1024268 | T | -0.0084 | 0.0014 | 0.0060 | 0.0093 |
| rs10251438 | A | 0.0078 | 0.0014 | -0.0268 | 0.0093 |
| rs10264573 | A | 0.0175 | 0.0032 | 0.0124 | 0.0216 |
| rs1035578 | A | -0.0094 | 0.0014 | -0.0080 | 0.0095 |
| rs10402747 | T | 0.0077 | 0.0014 | -0.0350 | 0.0098 |
| rs10411759 | A | 0.0116 | 0.0020 | -0.0050 | 0.0135 |
| rs10444280 | T | 0.0100 | 0.0018 | -0.0005 | 0.0119 |
| rs10460095 | A | -0.0124 | 0.0014 | 0.0065 | 0.0095 |
| rs10480450 | A | 0.0158 | 0.0028 | 0.0051 | 0.0172 |
| rs10496091 | A | -0.0131 | 0.0016 | 0.0171 | 0.0113 |
| rs10499535 | A | 0.0080 | 0.0014 | 0.0085 | 0.0093 |
| rs1050847 | T | 0.0092 | 0.0014 | -0.0262 | 0.0103 |
| rs10509251 | T | -0.0093 | 0.0017 | 0.0215 | 0.0105 |
| rs10515007 | T | 0.0143 | 0.0019 | -0.0127 | 0.0120 |
| rs1051860 | A | -0.0079 | 0.0014 | 0.0172 | 0.0096 |
| rs10519504 | T | 0.0119 | 0.0019 | -0.0111 | 0.0126 |
| rs1054442 | A | -0.0136 | 0.0015 | 0.0006 | 0.0097 |
| rs1061801 | A | -0.0110 | 0.0018 | 0.0157 | 0.0125 |
| rs1066769 | A | -0.0230 | 0.0041 | -0.0249 | 0.0279 |
| rs10742591 | A | -0.0084 | 0.0014 | 0.0054 | 0.0095 |
| rs10748841 | A | -0.0111 | 0.0020 | 0.0085 | 0.0099 |
| rs10750539 | A | -0.0095 | 0.0015 | 0.0000 | 0.0098 |
| rs10752262 | T | 0.0102 | 0.0014 | 0.0032 | 0.0097 |
| rs1075228 | A | -0.0081 | 0.0014 | -0.0064 | 0.0101 |
| rs10761202 | T | 0.0088 | 0.0014 | 0.0210 | 0.0094 |
| rs10761251 | A | 0.0113 | 0.0015 | 0.0091 | 0.0096 |
| rs10772644 | C | 0.0142 | 0.0022 | -0.0022 | 0.0141 |
| rs10773002 | A | 0.0185 | 0.0016 | -0.0060 | 0.0104 |
| rs10773208 | T | -0.0110 | 0.0016 | -0.0133 | 0.0110 |
| rs10782651 | T | -0.0082 | 0.0014 | 0.0070 | 0.0094 |
| rs10783243 | A | -0.0092 | 0.0014 | 0.0142 | 0.0093 |
| rs10789285 | T | 0.0091 | 0.0016 | -0.0128 | 0.0104 |
| rs10793903 | A | 0.0126 | 0.0022 | -0.0222 | 0.0140 |
| rs10795831 | T | -0.0099 | 0.0017 | 0.0186 | 0.0107 |
| rs10798888 | T | -0.0140 | 0.0019 | 0.0219 | 0.0117 |
| rs10799615 | A | -0.0089 | 0.0016 | 0.0190 | 0.0101 |
| rs10805383 | A | -0.0103 | 0.0014 | 0.0062 | 0.0094 |
| rs10810099 | A | -0.0146 | 0.0016 | -0.0016 | 0.0101 |
| rs10810145 | A | 0.0078 | 0.0014 | -0.0081 | 0.0094 |
| rs10830858 | T | 0.0092 | 0.0014 | -0.0107 | 0.0093 |
| rs10831656 | T | 0.0090 | 0.0015 | -0.0069 | 0.0103 |
| rs10844179 | A | 0.0104 | 0.0017 | -0.0107 | 0.0110 |
| rs10845051 | A | 0.0083 | 0.0014 | -0.0052 | 0.0096 |
| rs10875121 | C | 0.0174 | 0.0019 | -0.0198 | 0.0127 |
| rs10877283 | T | 0.0085 | 0.0014 | 0.0005 | 0.0096 |
| rs10879676 | T | -0.0083 | 0.0014 | -0.0037 | 0.0095 |
| rs10886012 | C | 0.0078 | 0.0014 | 0.0011 | 0.0093 |
| rs10892807 | T | 0.0109 | 0.0014 | 0.0093 | 0.0099 |
| rs10899282 | A | 0.0140 | 0.0017 | -0.0071 | 0.0114 |
| rs10927053 | A | -0.0125 | 0.0022 | 0.0099 | 0.0138 |
| rs10928190 | T | -0.0087 | 0.0014 | 0.0083 | 0.0096 |
| rs10931821 | A | -0.0141 | 0.0014 | 0.0111 | 0.0096 |
| rs10937240 | T | -0.0123 | 0.0018 | -0.0047 | 0.0110 |
| rs10940540 | A | 0.0078 | 0.0014 | 0.0162 | 0.0096 |
| rs10940921 | T | 0.0111 | 0.0014 | 0.0057 | 0.0093 |
| rs10943588 | A | 0.0084 | 0.0015 | -0.0224 | 0.0095 |
| rs10947439 | T | 0.0090 | 0.0015 | -0.0067 | 0.0100 |
| rs10949263 | A | -0.0091 | 0.0016 | -0.0098 | 0.0104 |
| rs10951590 | T | -0.0109 | 0.0015 | 0.0154 | 0.0099 |
| rs10974256 | A | -0.0093 | 0.0015 | -0.0046 | 0.0106 |
| rs1097784 | T | -0.0083 | 0.0014 | 0.0066 | 0.0094 |
| rs10979613 | T | -0.0117 | 0.0015 | 0.0037 | 0.0096 |
| rs10983324 | A | 0.0084 | 0.0015 | -0.0141 | 0.0100 |
| rs10984445 | A | 0.0115 | 0.0014 | -0.0191 | 0.0092 |
| rs10985402 | T | 0.0089 | 0.0016 | -0.0078 | 0.0103 |
| rs10995639 | T | 0.0078 | 0.0014 | -0.0155 | 0.0094 |
| rs10996167 | C | 0.0113 | 0.0020 | 0.0077 | 0.0098 |
| rs11003463 | T | 0.0091 | 0.0015 | -0.0096 | 0.0112 |
| rs11021432 | A | 0.0103 | 0.0015 | 0.0147 | 0.0102 |
| rs11023764 | A | 0.0091 | 0.0015 | 0.0202 | 0.0102 |
| rs11030084 | T | 0.0106 | 0.0018 | -0.0306 | 0.0116 |
| rs11030102 | C | 0.0101 | 0.0016 | -0.0261 | 0.0118 |
| rs1106090 | A | 0.0094 | 0.0014 | -0.0052 | 0.0096 |
| rs11076962 | T | 0.0086 | 0.0016 | 0.0059 | 0.0101 |
| rs1107871 | A | -0.0083 | 0.0014 | 0.0098 | 0.0092 |
| rs11081529 | T | 0.0126 | 0.0015 | -0.0138 | 0.0103 |
| rs11082011 | T | 0.0194 | 0.0015 | 0.0090 | 0.0097 |
| rs11100237 | A | -0.0101 | 0.0014 | 0.0104 | 0.0092 |
| rs11115056 | A | 0.0187 | 0.0023 | -0.0033 | 0.0182 |
| rs11121177 | A | 0.0151 | 0.0018 | -0.0083 | 0.0118 |
| rs111226181 | T | -0.0201 | 0.0018 | 0.0246 | 0.0132 |
| rs111235962 | T | 0.0133 | 0.0024 | -0.0029 | 0.0165 |
| rs11130380 | T | 0.0090 | 0.0014 | 0.0138 | 0.0098 |
| rs111370527 | T | 0.0151 | 0.0027 | 0.0289 | 0.0245 |
| rs11138947 | T | 0.0105 | 0.0016 | -0.0204 | 0.0099 |
| rs111530150 | T | 0.0269 | 0.0048 | 0.0530 | 0.0324 |
| rs11155813 | T | -0.0129 | 0.0023 | -0.0298 | 0.0133 |
| rs11157931 | A | -0.0139 | 0.0014 | -0.0210 | 0.0098 |
| rs11158800 | A | -0.0091 | 0.0014 | 0.0158 | 0.0096 |
| rs1117152 | T | 0.0106 | 0.0015 | -0.0090 | 0.0099 |
| rs11174399 | A | 0.0111 | 0.0018 | -0.0033 | 0.0116 |
| rs111821073 | T | 0.0124 | 0.0019 | -0.0141 | 0.0138 |
| rs111852224 | T | 0.0156 | 0.0022 | 0.0105 | 0.0174 |
| rs11190955 | T | -0.0083 | 0.0015 | 0.0018 | 0.0101 |
| rs11191237 | T | 0.0132 | 0.0024 | -0.0119 | 0.0112 |
| rs1120924 | T | 0.0108 | 0.0015 | -0.0031 | 0.0102 |
| rs11209894 | A | 0.0119 | 0.0021 | -0.0090 | 0.0130 |
| rs11210228 | T | -0.0105 | 0.0019 | -0.0001 | 0.0098 |
| rs11210400 | A | 0.0120 | 0.0014 | 0.0095 | 0.0096 |
| rs11211123 | A | 0.0104 | 0.0017 | 0.0060 | 0.0110 |
| rs11213482 | A | -0.0106 | 0.0019 | 0.0016 | 0.0147 |
| rs112210983 | A | 0.0105 | 0.0017 | 0.0012 | 0.0119 |
| rs1123285 | C | -0.0094 | 0.0015 | 0.0055 | 0.0100 |
| rs112375785 | A | 0.0184 | 0.0021 | -0.0084 | 0.0180 |
| rs112379405 | C | -0.0082 | 0.0014 | -0.0123 | 0.0095 |
| rs112512729 | T | -0.0166 | 0.0027 | 0.0142 | 0.0208 |
| rs112603734 | A | -0.0131 | 0.0022 | 0.0041 | 0.0117 |
| rs11265191 | T | 0.0083 | 0.0015 | -0.0042 | 0.0096 |
| rs112806496 | C | -0.0160 | 0.0025 | 0.0019 | 0.0191 |
| rs1128956 | T | -0.0140 | 0.0019 | 0.0188 | 0.0134 |
| rs113205706 | A | -0.0135 | 0.0024 | -0.0106 | 0.0184 |
| rs113520408 | A | 0.0155 | 0.0016 | -0.0100 | 0.0118 |
| rs113552169 | C | 0.0294 | 0.0053 | -0.0428 | 0.0383 |
| rs113588399 | T | -0.0104 | 0.0017 | 0.0052 | 0.0124 |
| rs113615161 | T | -0.0123 | 0.0021 | 0.0210 | 0.0134 |
| rs113720505 | A | 0.0164 | 0.0029 | 0.0022 | 0.0264 |
| rs113731629 | T | 0.0339 | 0.0061 | -0.0205 | 0.0497 |
| rs114192810 | T | -0.0279 | 0.0051 | 0.0375 | 0.0353 |
| rs114468556 | A | 0.0293 | 0.0036 | -0.0511 | 0.0320 |
| rs114593137 | A | -0.0122 | 0.0017 | 0.0080 | 0.0122 |
| rs1146079 | C | -0.0087 | 0.0016 | 0.0050 | 0.0109 |
| rs114810763 | A | -0.0121 | 0.0020 | -0.0014 | 0.0162 |
| rs114952970 | T | 0.0265 | 0.0040 | 0.0289 | 0.0304 |
| rs114958262 | T | -0.0220 | 0.0039 | -0.0194 | 0.0338 |
| rs115000530 | A | -0.0250 | 0.0031 | 0.0537 | 0.0237 |
| rs115017135 | A | -0.0206 | 0.0038 | 0.0049 | 0.0309 |
| rs11542663 | A | 0.0112 | 0.0015 | 0.0065 | 0.0098 |
| rs115438240 | T | -0.0169 | 0.0030 | 0.0315 | 0.0244 |
| rs115693355 | A | 0.0309 | 0.0053 | -0.0115 | 0.0428 |
| rs115732722 | A | -0.0415 | 0.0072 | -0.0805 | 0.0530 |
| rs11588857 | A | 0.0199 | 0.0017 | 0.0190 | 0.0110 |
| rs11591870 | T | 0.0098 | 0.0017 | -0.0100 | 0.0128 |
| rs11592299 | A | 0.0097 | 0.0018 | 0.0220 | 0.0129 |
| rs11596387 | T | 0.0258 | 0.0046 | 0.0273 | 0.0404 |
| rs11598765 | A | -0.0106 | 0.0018 | 0.0012 | 0.0112 |
| rs11599236 | T | -0.0106 | 0.0014 | 0.0126 | 0.0094 |
| rs11613431 | A | -0.0094 | 0.0016 | -0.0097 | 0.0114 |
| rs11620355 | A | 0.0182 | 0.0025 | -0.0175 | 0.0176 |
| rs11623285 | T | -0.0135 | 0.0021 | 0.0199 | 0.0146 |
| rs116386746 | C | 0.0249 | 0.0045 | -0.0102 | 0.0397 |
| rs11640569 | A | 0.0112 | 0.0020 | 0.0023 | 0.0144 |
| rs11644446 | A | 0.0117 | 0.0019 | 0.0156 | 0.0145 |
| rs11647188 | A | 0.0079 | 0.0014 | 0.0022 | 0.0097 |
| rs11652522 | A | -0.0154 | 0.0023 | 0.0168 | 0.0156 |
| rs11657342 | A | 0.0140 | 0.0019 | -0.0018 | 0.0131 |
| rs11657979 | A | -0.0091 | 0.0017 | 0.0316 | 0.0113 |
| rs11663602 | A | -0.0130 | 0.0016 | -0.0032 | 0.0100 |
| rs11663678 | T | -0.0121 | 0.0022 | -0.0047 | 0.0149 |
| rs116656374 | T | -0.0242 | 0.0038 | 0.0022 | 0.0302 |
| rs11675476 | T | -0.0127 | 0.0014 | 0.0131 | 0.0094 |
| rs1167827 | A | 0.0104 | 0.0014 | -0.0116 | 0.0098 |
| rs11678980 | A | -0.0166 | 0.0014 | 0.0092 | 0.0106 |
| rs11687736 | A | -0.0105 | 0.0015 | -0.0004 | 0.0102 |
| rs11693885 | A | -0.0091 | 0.0014 | 0.0104 | 0.0097 |
| rs116967397 | A | 0.0315 | 0.0056 | -0.0109 | 0.0439 |
| rs117005905 | T | 0.0150 | 0.0022 | 0.0007 | 0.0140 |
| rs11703948 | A | -0.0168 | 0.0024 | 0.0081 | 0.0156 |
| rs1171040 | A | -0.0113 | 0.0017 | 0.0302 | 0.0113 |
| rs11716398 | A | 0.0159 | 0.0021 | 0.0144 | 0.0131 |
| rs11720093 | T | 0.0150 | 0.0026 | 0.0168 | 0.0177 |
| rs11720985 | T | 0.0085 | 0.0015 | -0.0046 | 0.0106 |
| rs11724690 | T | 0.0093 | 0.0015 | 0.0187 | 0.0107 |
| rs117273411 | T | -0.0211 | 0.0034 | 0.0317 | 0.0297 |
| rs11732160 | A | 0.0105 | 0.0016 | 0.0071 | 0.0106 |
| rs11732657 | A | -0.0126 | 0.0016 | 0.0074 | 0.0105 |
| rs11733439 | A | -0.0096 | 0.0017 | 0.0127 | 0.0108 |
| rs11736863 | A | 0.0142 | 0.0018 | 0.0075 | 0.0122 |
| rs117398064 | C | -0.0141 | 0.0025 | 0.0045 | 0.0179 |
| rs117468730 | A | -0.0310 | 0.0050 | 0.0154 | 0.0322 |
| rs117623407 | A | -0.0118 | 0.0020 | -0.0102 | 0.0147 |
| rs11765387 | T | 0.0086 | 0.0015 | 0.0001 | 0.0095 |
| rs117668569 | T | 0.0234 | 0.0042 | -0.0318 | 0.0293 |
| rs117677995 | A | -0.0180 | 0.0032 | 0.0345 | 0.0300 |
| rs11772232 | T | 0.0177 | 0.0019 | -0.0276 | 0.0132 |
| rs11774212 | T | 0.0128 | 0.0014 | -0.0057 | 0.0099 |
| rs117799466 | C | 0.0106 | 0.0016 | -0.0111 | 0.0124 |
| rs11780023 | T | 0.0085 | 0.0014 | -0.0012 | 0.0094 |
| rs11789013 | T | 0.0106 | 0.0016 | -0.0157 | 0.0106 |
| rs117895796 | T | 0.0362 | 0.0064 | -0.0004 | 0.0517 |
| rs118040169 | A | -0.0248 | 0.0038 | -0.0110 | 0.0313 |
| rs118093058 | T | 0.0146 | 0.0021 | 0.0149 | 0.0163 |
| rs118134876 | T | -0.0223 | 0.0030 | 0.0521 | 0.0239 |
| rs11845781 | T | -0.0079 | 0.0014 | -0.0050 | 0.0094 |
| rs11871429 | A | 0.0128 | 0.0017 | 0.0050 | 0.0107 |
| rs11876620 | T | 0.0143 | 0.0024 | 0.0054 | 0.0165 |
| rs11894424 | A | 0.0209 | 0.0033 | -0.0161 | 0.0174 |
| rs11917701 | A | -0.0087 | 0.0014 | 0.0068 | 0.0094 |
| rs11919835 | C | 0.0089 | 0.0015 | -0.0215 | 0.0099 |
| rs1196760 | C | 0.0139 | 0.0025 | 0.0113 | 0.0192 |
| rs12005151 | A | 0.0087 | 0.0015 | 0.0051 | 0.0103 |
| rs12028010 | T | 0.0164 | 0.0017 | -0.0118 | 0.0114 |
| rs12028229 | T | 0.0099 | 0.0016 | 0.0225 | 0.0123 |
| rs12054166 | C | 0.0096 | 0.0016 | -0.0113 | 0.0110 |
| rs12076635 | C | 0.0207 | 0.0017 | -0.0143 | 0.0106 |
| rs12113634 | T | 0.0088 | 0.0015 | -0.0117 | 0.0095 |
| rs12123293 | T | 0.0086 | 0.0015 | -0.0026 | 0.0098 |
| rs12126231 | A | 0.0095 | 0.0014 | -0.0097 | 0.0094 |
| rs12127928 | T | 0.0108 | 0.0018 | -0.0100 | 0.0128 |
| rs12145078 | A | -0.0127 | 0.0017 | 0.0063 | 0.0113 |
| rs12151248 | T | -0.0149 | 0.0023 | -0.0062 | 0.0185 |
| rs12170452 | A | 0.0105 | 0.0014 | -0.0009 | 0.0097 |
| rs1220779 | A | -0.0079 | 0.0014 | 0.0149 | 0.0095 |
| rs12234369 | A | 0.0087 | 0.0015 | 0.0070 | 0.0101 |
| rs12238011 | T | -0.0141 | 0.0026 | 0.0081 | 0.0157 |
| rs12273435 | A | -0.0132 | 0.0017 | 0.0216 | 0.0120 |
| rs12285074 | A | 0.0161 | 0.0023 | 0.0164 | 0.0127 |
| rs12304188 | A | 0.0111 | 0.0019 | -0.0035 | 0.0117 |
| rs12325727 | A | 0.0088 | 0.0014 | -0.0330 | 0.0096 |
| rs12332731 | A | 0.0135 | 0.0018 | -0.0142 | 0.0131 |
| rs12342546 | A | -0.0128 | 0.0017 | -0.0058 | 0.0109 |
| rs12359372 | T | -0.0092 | 0.0015 | 0.0232 | 0.0101 |
| rs12375949 | T | -0.0138 | 0.0014 | 0.0044 | 0.0094 |
| rs12405889 | T | -0.0098 | 0.0014 | 0.0002 | 0.0092 |
| rs12431682 | T | -0.0093 | 0.0015 | -0.0043 | 0.0098 |
| rs12438177 | A | 0.0093 | 0.0015 | -0.0192 | 0.0094 |
| rs12453682 | T | 0.0102 | 0.0015 | -0.0246 | 0.0103 |
| rs1245829 | A | -0.0099 | 0.0014 | 0.0179 | 0.0101 |
| rs12467175 | T | -0.0081 | 0.0014 | 0.0250 | 0.0096 |
| rs12468040 | T | 0.0137 | 0.0014 | 0.0026 | 0.0094 |
| rs12473986 | T | -0.0085 | 0.0015 | 0.0139 | 0.0102 |
| rs12477385 | T | 0.0109 | 0.0017 | -0.0231 | 0.0117 |
| rs12478156 | T | 0.0108 | 0.0015 | 0.0129 | 0.0098 |
| rs12503522 | T | -0.0113 | 0.0016 | 0.0172 | 0.0102 |
| rs12506221 | T | -0.0130 | 0.0014 | 0.0165 | 0.0092 |
| rs12516990 | C | -0.0099 | 0.0017 | 0.0103 | 0.0122 |
| rs12519073 | T | -0.0107 | 0.0017 | -0.0098 | 0.0112 |
| rs12524795 | T | 0.0102 | 0.0014 | -0.0111 | 0.0097 |
| rs12568153 | T | -0.0091 | 0.0015 | 0.0117 | 0.0104 |
| rs12571549 | A | 0.0151 | 0.0020 | -0.0072 | 0.0119 |
| rs12574281 | A | -0.0080 | 0.0015 | -0.0070 | 0.0102 |
| rs12591647 | T | -0.0152 | 0.0018 | 0.0175 | 0.0112 |
| rs12601380 | A | -0.0082 | 0.0014 | 0.0061 | 0.0096 |
| rs12602286 | T | 0.0160 | 0.0021 | -0.0306 | 0.0121 |
| rs12613500 | C | 0.0098 | 0.0014 | -0.0058 | 0.0096 |
| rs12614263 | A | 0.0083 | 0.0014 | -0.0080 | 0.0095 |
| rs12620796 | A | 0.0121 | 0.0019 | -0.0140 | 0.0102 |
| rs12638072 | A | -0.0107 | 0.0015 | 0.0017 | 0.0105 |
| rs12643771 | T | 0.0130 | 0.0015 | -0.0079 | 0.0105 |
| rs12646216 | T | 0.0084 | 0.0014 | -0.0173 | 0.0098 |
| rs12646523 | T | -0.0132 | 0.0016 | 0.0036 | 0.0104 |
| rs12659776 | T | 0.0085 | 0.0014 | -0.0186 | 0.0092 |
| rs12670376 | A | 0.0092 | 0.0014 | -0.0129 | 0.0092 |
| rs1267062 | C | 0.0113 | 0.0018 | -0.0045 | 0.0128 |
| rs12682775 | T | -0.0119 | 0.0017 | 0.0034 | 0.0107 |
| rs12694681 | T | 0.0102 | 0.0015 | -0.0062 | 0.0105 |
| rs12709186 | A | -0.0091 | 0.0015 | 0.0062 | 0.0103 |
| rs12712269 | T | -0.0107 | 0.0014 | 0.0078 | 0.0097 |
| rs12746551 | C | 0.0248 | 0.0041 | 0.0243 | 0.0301 |
| rs12750688 | C | 0.0116 | 0.0016 | 0.0249 | 0.0117 |
| rs12761729 | T | -0.0098 | 0.0016 | -0.0018 | 0.0118 |
| rs12761761 | T | 0.0153 | 0.0017 | -0.0011 | 0.0119 |
| rs12764593 | C | 0.0202 | 0.0029 | 0.0015 | 0.0221 |
| rs12765185 | A | -0.0088 | 0.0016 | 0.0170 | 0.0114 |
| rs12774577 | T | 0.0142 | 0.0020 | -0.0175 | 0.0142 |
| rs12789313 | T | 0.0092 | 0.0014 | -0.0042 | 0.0094 |
| rs12790196 | T | -0.0101 | 0.0015 | -0.0090 | 0.0103 |
| rs12810587 | T | 0.0094 | 0.0015 | -0.0080 | 0.0100 |
| rs12875339 | A | -0.0108 | 0.0015 | -0.0007 | 0.0094 |
| rs12888615 | T | 0.0120 | 0.0018 | -0.0093 | 0.0122 |
| rs12891042 | T | 0.0092 | 0.0014 | -0.0041 | 0.0094 |
| rs12891191 | C | 0.0080 | 0.0014 | -0.0086 | 0.0099 |
| rs12912465 | T | -0.0104 | 0.0016 | 0.0348 | 0.0122 |
| rs12952191 | T | -0.0081 | 0.0014 | 0.0055 | 0.0094 |
| rs12953422 | A | -0.0080 | 0.0014 | 0.0152 | 0.0097 |
| rs12957463 | A | 0.0146 | 0.0018 | -0.0158 | 0.0115 |
| rs12962845 | C | -0.0123 | 0.0021 | 0.0138 | 0.0136 |
| rs12967010 | T | 0.0114 | 0.0017 | 0.0002 | 0.0122 |
| rs12970264 | A | -0.0085 | 0.0014 | 0.0009 | 0.0098 |
| rs12981405 | T | -0.0112 | 0.0019 | 0.0003 | 0.0120 |
| rs13009915 | T | 0.0212 | 0.0037 | 0.0401 | 0.0343 |
| rs13010288 | T | 0.0198 | 0.0021 | -0.0061 | 0.0154 |
| rs13010566 | A | -0.0091 | 0.0014 | -0.0002 | 0.0097 |
| rs13015496 | C | 0.0119 | 0.0018 | -0.0113 | 0.0115 |
| rs13016316 | A | -0.0138 | 0.0024 | -0.0091 | 0.0145 |
| rs1301838 | T | -0.0096 | 0.0015 | 0.0109 | 0.0105 |
| rs13018640 | T | -0.0215 | 0.0014 | 0.0133 | 0.0099 |
| rs13026611 | A | -0.0084 | 0.0015 | -0.0221 | 0.0105 |
| rs13029602 | T | -0.0092 | 0.0014 | -0.0050 | 0.0094 |
| rs13034349 | T | -0.0082 | 0.0014 | 0.0161 | 0.0103 |
| rs13050131 | A | -0.0081 | 0.0015 | -0.0019 | 0.0099 |
| rs13085461 | C | 0.0103 | 0.0014 | -0.0051 | 0.0092 |
| rs13099165 | T | -0.0118 | 0.0017 | -0.0218 | 0.0121 |
| rs13107325 | T | -0.0239 | 0.0027 | -0.0067 | 0.0223 |
| rs13117856 | A | 0.0084 | 0.0015 | -0.0037 | 0.0105 |
| rs13130765 | C | -0.0091 | 0.0014 | 0.0363 | 0.0105 |
| rs13133213 | A | 0.0096 | 0.0014 | -0.0206 | 0.0092 |
| rs13145650 | T | -0.0173 | 0.0025 | 0.0246 | 0.0170 |
| rs13154429 | C | -0.0136 | 0.0023 | 0.0023 | 0.0153 |
| rs13163062 | T | 0.0115 | 0.0014 | 0.0014 | 0.0095 |
| rs13163845 | T | -0.0149 | 0.0020 | -0.0106 | 0.0136 |
| rs13168136 | A | -0.0092 | 0.0017 | -0.0069 | 0.0105 |
| rs13169187 | A | -0.0079 | 0.0014 | -0.0008 | 0.0094 |
| rs13177031 | A | -0.0085 | 0.0015 | 0.0059 | 0.0097 |
| rs13190235 | T | -0.0138 | 0.0024 | -0.0034 | 0.0149 |
| rs13197257 | T | 0.0109 | 0.0016 | 0.0027 | 0.0108 |
| rs1320139 | C | -0.0149 | 0.0014 | 0.0270 | 0.0096 |
| rs13212041 | T | 0.0101 | 0.0018 | -0.0069 | 0.0120 |
| rs13240401 | T | 0.0176 | 0.0017 | -0.0116 | 0.0113 |
| rs13246220 | T | -0.0093 | 0.0014 | 0.0027 | 0.0096 |
| rs13261773 | C | 0.0122 | 0.0019 | -0.0031 | 0.0120 |
| rs13266287 | A | -0.0085 | 0.0015 | -0.0025 | 0.0099 |
| rs13281564 | A | 0.0088 | 0.0014 | 0.0019 | 0.0094 |
| rs13284516 | C | -0.0082 | 0.0014 | 0.0024 | 0.0099 |
| rs1329044 | T | 0.0134 | 0.0019 | -0.0229 | 0.0120 |
| rs1329125 | T | -0.0097 | 0.0015 | 0.0186 | 0.0109 |
| rs13296345 | T | 0.0090 | 0.0015 | -0.0181 | 0.0094 |
| rs13318986 | A | 0.0124 | 0.0015 | -0.0055 | 0.0099 |
| rs13327482 | A | -0.0121 | 0.0018 | -0.0006 | 0.0131 |
| rs1334297 | A | 0.0257 | 0.0016 | -0.0001 | 0.0111 |
| rs1335482 | T | 0.0096 | 0.0014 | -0.0066 | 0.0098 |
| rs13361043 | T | -0.0158 | 0.0028 | 0.0188 | 0.0164 |
| rs13381557 | A | -0.0087 | 0.0014 | -0.0019 | 0.0093 |
| rs13388333 | A | 0.0085 | 0.0015 | -0.0050 | 0.0104 |
| rs13397529 | C | -0.0097 | 0.0017 | 0.0269 | 0.0116 |
| rs13398860 | A | 0.0086 | 0.0015 | -0.0140 | 0.0102 |
| rs13402497 | A | -0.0080 | 0.0014 | -0.0087 | 0.0095 |
| rs13422673 | T | -0.0110 | 0.0014 | 0.0114 | 0.0098 |
| rs13425585 | C | -0.0089 | 0.0014 | 0.0109 | 0.0096 |
| rs1368250 | T | 0.0250 | 0.0045 | -0.0197 | 0.0416 |
| rs137079 | T | 0.0128 | 0.0020 | 0.0185 | 0.0140 |
| rs137858393 | A | 0.0193 | 0.0029 | 0.0343 | 0.0218 |
| rs138096147 | A | 0.0234 | 0.0042 | 0.0714 | 0.0366 |
| rs138484388 | T | -0.0306 | 0.0051 | 0.0404 | 0.0419 |
| rs1391513 | T | -0.0095 | 0.0015 | 0.0032 | 0.0101 |
| rs139244147 | A | -0.0193 | 0.0030 | 0.0062 | 0.0186 |
| rs1392816 | T | 0.0082 | 0.0014 | -0.0101 | 0.0097 |
| rs139980871 | T | 0.0180 | 0.0029 | -0.0091 | 0.0210 |
| rs1404549 | A | -0.0089 | 0.0015 | 0.0127 | 0.0101 |
| rs1405876 | T | 0.0109 | 0.0015 | -0.0007 | 0.0102 |
| rs140711597 | C | 0.0366 | 0.0055 | 0.0285 | 0.0425 |
| rs1408284 | C | -0.0127 | 0.0020 | 0.0146 | 0.0145 |
| rs1408430 | C | 0.0096 | 0.0016 | 0.0068 | 0.0102 |
| rs141586924 | A | -0.0090 | 0.0015 | 0.0037 | 0.0109 |
| rs141729694 | T | 0.0195 | 0.0026 | 0.0433 | 0.0175 |
| rs142014757 | A | 0.0102 | 0.0018 | 0.0162 | 0.0137 |
| rs1426619 | T | 0.0096 | 0.0014 | 0.0186 | 0.0092 |
| rs1427298 | T | 0.0086 | 0.0014 | -0.0091 | 0.0098 |
| rs142747148 | A | 0.0329 | 0.0052 | -0.0160 | 0.0432 |
| rs1427829 | A | -0.0083 | 0.0014 | 0.0146 | 0.0092 |
| rs143148393 | T | -0.0211 | 0.0039 | 0.0006 | 0.0307 |
| rs1434630 | T | -0.0137 | 0.0020 | -0.0037 | 0.0124 |
| rs143743568 | A | 0.0118 | 0.0020 | -0.0119 | 0.0136 |
| rs143812851 | A | -0.0113 | 0.0019 | -0.0071 | 0.0126 |
| rs1440930 | C | 0.0081 | 0.0014 | 0.0137 | 0.0094 |
| rs144336753 | A | 0.0388 | 0.0053 | -0.0402 | 0.0396 |
| rs1445591 | A | 0.0085 | 0.0015 | 0.0033 | 0.0111 |
| rs145018899 | A | 0.0095 | 0.0015 | 0.0082 | 0.0109 |
| rs1464297 | T | -0.0109 | 0.0015 | -0.0164 | 0.0098 |
| rs1467737 | T | -0.0084 | 0.0014 | 0.0167 | 0.0095 |
| rs1485300 | C | 0.0098 | 0.0015 | -0.0059 | 0.0100 |
| rs1490612 | T | 0.0193 | 0.0026 | 0.0065 | 0.0173 |
| rs150252215 | A | 0.0204 | 0.0035 | 0.0032 | 0.0263 |
| rs150537577 | A | -0.0151 | 0.0026 | 0.0265 | 0.0182 |
| rs1505676 | C | 0.0084 | 0.0015 | -0.0157 | 0.0094 |
| rs151381 | T | -0.0087 | 0.0014 | 0.0192 | 0.0094 |
| rs1518890 | T | -0.0090 | 0.0016 | -0.0024 | 0.0104 |
| rs1527878 | A | -0.0105 | 0.0016 | 0.0266 | 0.0116 |
| rs1529597 | A | 0.0223 | 0.0038 | -0.0413 | 0.0250 |
| rs1538389 | T | -0.0109 | 0.0018 | 0.0179 | 0.0120 |
| rs1542354 | A | -0.0084 | 0.0014 | -0.0115 | 0.0098 |
| rs1544 | A | -0.0096 | 0.0016 | 0.0064 | 0.0104 |
| rs1554798 | A | -0.0080 | 0.0014 | 0.0161 | 0.0098 |
| rs1564347 | T | 0.0088 | 0.0015 | -0.0246 | 0.0097 |
| rs1569092 | A | 0.0160 | 0.0020 | -0.0055 | 0.0134 |
| rs1593022 | T | -0.0099 | 0.0017 | 0.0061 | 0.0115 |
| rs1603460 | T | 0.0090 | 0.0014 | -0.0048 | 0.0093 |
| rs162445 | A | 0.0152 | 0.0026 | 0.0037 | 0.0150 |
| rs163229 | C | -0.0260 | 0.0047 | 0.0412 | 0.0315 |
| rs1637770 | T | 0.0191 | 0.0028 | -0.0141 | 0.0192 |
| rs164938 | T | -0.0088 | 0.0014 | 0.0068 | 0.0095 |
| rs165633 | A | -0.0116 | 0.0016 | 0.0180 | 0.0116 |
| rs1656614 | C | 0.0122 | 0.0015 | -0.0016 | 0.0106 |
| rs1671269 | T | -0.0100 | 0.0016 | 0.0093 | 0.0101 |
| rs1671770 | A | 0.0125 | 0.0018 | 0.0260 | 0.0118 |
| rs16851779 | T | -0.0153 | 0.0027 | 0.0285 | 0.0196 |
| rs16871807 | T | 0.0090 | 0.0015 | -0.0086 | 0.0098 |
| rs1689510 | C | 0.0194 | 0.0015 | 0.0017 | 0.0103 |
| rs16901689 | T | 0.0111 | 0.0019 | -0.0238 | 0.0125 |
| rs1693584 | T | -0.0087 | 0.0015 | 0.0147 | 0.0100 |
| rs16958559 | T | 0.0096 | 0.0017 | 0.0163 | 0.0128 |
| rs16966271 | T | 0.0092 | 0.0016 | -0.0066 | 0.0105 |
| rs16975275 | A | -0.0102 | 0.0017 | -0.0210 | 0.0114 |
| rs17048801 | A | -0.0106 | 0.0014 | 0.0378 | 0.0101 |
| rs17048855 | A | 0.0108 | 0.0015 | -0.0013 | 0.0097 |
| rs17069646 | T | 0.0085 | 0.0015 | 0.0058 | 0.0099 |
| rs17110109 | T | -0.0093 | 0.0014 | 0.0132 | 0.0095 |
| rs17113730 | A | -0.0170 | 0.0027 | 0.0090 | 0.0184 |
| rs17131123 | A | -0.0243 | 0.0039 | -0.0364 | 0.0370 |
| rs17133297 | A | -0.0146 | 0.0023 | 0.0121 | 0.0137 |
| rs17144467 | A | 0.0098 | 0.0015 | -0.0032 | 0.0098 |
| rs17148998 | A | 0.0101 | 0.0017 | -0.0202 | 0.0124 |
| rs171697 | C | 0.0133 | 0.0015 | -0.0055 | 0.0101 |
| rs17170519 | C | 0.0132 | 0.0023 | -0.0066 | 0.0137 |
| rs1717204 | A | -0.0117 | 0.0018 | 0.0052 | 0.0127 |
| rs1718188 | A | 0.0096 | 0.0014 | -0.0215 | 0.0094 |
| rs17186106 | T | 0.0108 | 0.0018 | -0.0171 | 0.0129 |
| rs17190418 | T | -0.0213 | 0.0031 | 0.0198 | 0.0252 |
| rs17205908 | T | -0.0092 | 0.0015 | 0.0000 | 0.0110 |
| rs17248751 | A | -0.0136 | 0.0017 | 0.0250 | 0.0129 |
| rs1728118 | A | 0.0089 | 0.0016 | -0.0082 | 0.0101 |
| rs1729412 | T | -0.0107 | 0.0014 | 0.0012 | 0.0096 |
| rs173003 | A | -0.0082 | 0.0014 | -0.0083 | 0.0096 |
| rs1734370 | A | 0.0099 | 0.0016 | -0.0034 | 0.0101 |
| rs1738050 | C | -0.0093 | 0.0014 | -0.0024 | 0.0096 |
| rs17411339 | A | 0.0140 | 0.0014 | -0.0132 | 0.0100 |
| rs17428076 | C | 0.0137 | 0.0016 | 0.0030 | 0.0114 |
| rs17440885 | A | 0.0160 | 0.0029 | 0.0226 | 0.0222 |
| rs1747714 | T | 0.0106 | 0.0014 | 0.0205 | 0.0093 |
| rs1747817 | T | -0.0092 | 0.0016 | 0.0317 | 0.0114 |
| rs17489649 | A | 0.0127 | 0.0015 | -0.0008 | 0.0112 |
| rs17502934 | T | -0.0167 | 0.0020 | -0.0001 | 0.0134 |
| rs175325 | A | -0.0100 | 0.0014 | 0.0018 | 0.0095 |
| rs17536059 | C | -0.0108 | 0.0019 | -0.0010 | 0.0130 |
| rs17551064 | A | 0.0141 | 0.0019 | 0.0190 | 0.0132 |
| rs17563464 | A | -0.0147 | 0.0017 | -0.0065 | 0.0145 |
| rs17565975 | A | -0.0106 | 0.0014 | -0.0260 | 0.0093 |
| rs17568389 | A | 0.0127 | 0.0014 | -0.0073 | 0.0094 |
| rs17574007 | A | -0.0129 | 0.0021 | -0.0027 | 0.0160 |
| rs1758747 | A | 0.0092 | 0.0015 | -0.0143 | 0.0100 |
| rs17598675 | T | -0.0116 | 0.0014 | 0.0084 | 0.0096 |
| rs17604349 | A | 0.0127 | 0.0018 | -0.0067 | 0.0122 |
| rs17609255 | T | -0.0087 | 0.0014 | 0.0225 | 0.0094 |
| rs176218 | T | 0.0200 | 0.0018 | -0.0340 | 0.0111 |
| rs17622379 | T | -0.0113 | 0.0018 | -0.0106 | 0.0116 |
| rs17650634 | T | -0.0104 | 0.0019 | 0.0177 | 0.0138 |
| rs17667540 | A | -0.0087 | 0.0015 | 0.0034 | 0.0101 |
| rs17669337 | T | -0.0110 | 0.0014 | 0.0035 | 0.0097 |
| rs17680712 | T | -0.0092 | 0.0014 | 0.0083 | 0.0093 |
| rs17686649 | T | -0.0098 | 0.0017 | 0.0272 | 0.0113 |
| rs17732878 | T | -0.0095 | 0.0017 | -0.0234 | 0.0115 |
| rs17742342 | A | -0.0119 | 0.0018 | 0.0192 | 0.0126 |
| rs17747544 | A | -0.0090 | 0.0014 | 0.0141 | 0.0095 |
| rs1779549 | A | 0.0081 | 0.0014 | 0.0012 | 0.0096 |
| rs178183 | T | 0.0145 | 0.0016 | -0.0150 | 0.0106 |
| rs17882802 | A | 0.0084 | 0.0014 | -0.0017 | 0.0101 |
| rs17883331 | A | -0.0108 | 0.0018 | 0.0459 | 0.0129 |
| rs1792602 | A | -0.0097 | 0.0014 | -0.0042 | 0.0097 |
| rs181214 | T | -0.0104 | 0.0017 | 0.0229 | 0.0132 |
| rs182355396 | A | -0.0525 | 0.0084 | 0.0223 | 0.0653 |
| rs182902112 | A | -0.0300 | 0.0053 | 0.0255 | 0.0335 |
| rs1835339 | T | 0.0084 | 0.0014 | 0.0161 | 0.0098 |
| rs183869217 | C | 0.0174 | 0.0030 | -0.0025 | 0.0219 |
| rs1841023 | A | -0.0092 | 0.0015 | -0.0064 | 0.0106 |
| rs1842713 | A | -0.0117 | 0.0017 | 0.0016 | 0.0108 |
| rs1843815 | A | 0.0077 | 0.0014 | -0.0185 | 0.0093 |
| rs1861786 | A | -0.0081 | 0.0014 | 0.0001 | 0.0094 |
| rs1865955 | T | 0.0122 | 0.0019 | 0.0024 | 0.0114 |
| rs1866823 | A | 0.0107 | 0.0014 | -0.0172 | 0.0096 |
| rs187580 | T | -0.0104 | 0.0017 | 0.0255 | 0.0123 |
| rs187951956 | A | -0.0315 | 0.0058 | 0.0025 | 0.0376 |
| rs1880088 | A | -0.0097 | 0.0016 | 0.0236 | 0.0101 |
| rs1880692 | A | 0.0086 | 0.0014 | -0.0007 | 0.0093 |
| rs188251563 | A | 0.0428 | 0.0070 | -0.1372 | 0.0593 |
| rs1890132 | T | -0.0089 | 0.0015 | 0.0036 | 0.0106 |
| rs1898111 | A | 0.0101 | 0.0018 | -0.0010 | 0.0115 |
| rs190102446 | T | -0.0210 | 0.0037 | -0.0366 | 0.0270 |
| rs1905616 | A | 0.0083 | 0.0015 | -0.0222 | 0.0098 |
| rs1910005 | T | -0.0090 | 0.0015 | 0.0004 | 0.0103 |
| rs1918394 | T | 0.0119 | 0.0019 | -0.0099 | 0.0121 |
| rs192436652 | T | -0.0319 | 0.0045 | -0.0637 | 0.0385 |
| rs1933264 | T | 0.0100 | 0.0016 | 0.0015 | 0.0105 |
| rs1952183 | A | -0.0089 | 0.0014 | -0.0017 | 0.0095 |
| rs1955250 | A | -0.0140 | 0.0025 | 0.0145 | 0.0171 |
| rs1963381 | A | 0.0099 | 0.0016 | 0.0081 | 0.0106 |
| rs1980129 | A | 0.0085 | 0.0014 | 0.0055 | 0.0092 |
| rs198262 | T | -0.0223 | 0.0035 | 0.0309 | 0.0262 |
| rs1991585 | T | -0.0101 | 0.0015 | -0.0013 | 0.0098 |
| rs1995181 | A | -0.0078 | 0.0014 | -0.0143 | 0.0093 |
| rs2007655 | T | 0.0086 | 0.0014 | -0.0034 | 0.0092 |
| rs2011603 | A | -0.0091 | 0.0016 | 0.0314 | 0.0108 |
| rs2014830 | T | 0.0110 | 0.0015 | -0.0253 | 0.0102 |
| rs201495 | T | -0.0112 | 0.0019 | 0.0160 | 0.0101 |
| rs2023016 | C | 0.0107 | 0.0017 | -0.0097 | 0.0109 |
| rs2024568 | T | -0.0097 | 0.0016 | 0.0055 | 0.0105 |
| rs2029401 | A | -0.0091 | 0.0014 | 0.0223 | 0.0097 |
| rs2034631 | T | 0.0089 | 0.0015 | -0.0090 | 0.0096 |
| rs2039204 | A | 0.0082 | 0.0014 | -0.0052 | 0.0096 |
| rs2043187 | A | 0.0107 | 0.0015 | -0.0016 | 0.0108 |
| rs2052285 | A | 0.0112 | 0.0014 | -0.0100 | 0.0104 |
| rs2055940 | A | 0.0082 | 0.0015 | -0.0069 | 0.0109 |
| rs2077235 | T | 0.0097 | 0.0017 | 0.0122 | 0.0107 |
| rs2081652 | A | 0.0123 | 0.0015 | 0.0121 | 0.0102 |
| rs2082317 | T | -0.0078 | 0.0014 | -0.0106 | 0.0095 |
| rs2088913 | A | 0.0081 | 0.0014 | 0.0047 | 0.0092 |
| rs2092248 | T | -0.0083 | 0.0015 | 0.0054 | 0.0103 |
| rs2098526 | A | -0.0241 | 0.0043 | 0.0387 | 0.0255 |
| rs2100249 | T | -0.0089 | 0.0015 | 0.0108 | 0.0096 |
| rs2126069 | T | 0.0080 | 0.0014 | 0.0021 | 0.0095 |
| rs2131167 | A | 0.0080 | 0.0014 | -0.0151 | 0.0097 |
| rs214626 | A | 0.0101 | 0.0018 | -0.0059 | 0.0112 |
| rs2160514 | A | -0.0094 | 0.0014 | 0.0040 | 0.0094 |
| rs2179152 | T | -0.0131 | 0.0015 | -0.0022 | 0.0099 |
| rs2183271 | T | 0.0084 | 0.0015 | 0.0002 | 0.0100 |
| rs2199409 | T | 0.0103 | 0.0017 | -0.0104 | 0.0117 |
| rs2212430 | T | -0.0100 | 0.0016 | 0.0018 | 0.0101 |
| rs2216144 | T | 0.0098 | 0.0014 | -0.0096 | 0.0093 |
| rs2220926 | T | -0.0095 | 0.0014 | 0.0021 | 0.0093 |
| rs2250660 | C | 0.0080 | 0.0014 | -0.0115 | 0.0098 |
| rs2252098 | T | -0.0077 | 0.0014 | -0.0108 | 0.0096 |
| rs2254681 | A | 0.0101 | 0.0016 | -0.0104 | 0.0106 |
| rs2256965 | A | 0.0106 | 0.0014 | -0.0126 | 0.0101 |
| rs2276209 | A | 0.0085 | 0.0015 | -0.0043 | 0.0097 |
| rs2283076 | A | 0.0115 | 0.0017 | 0.0021 | 0.0115 |
| rs2287838 | A | -0.0104 | 0.0014 | -0.0010 | 0.0095 |
| rs2290601 | T | -0.0149 | 0.0017 | 0.0020 | 0.0106 |
| rs2297293 | C | 0.0099 | 0.0015 | 0.0067 | 0.0100 |
| rs2299156 | T | 0.0099 | 0.0018 | -0.0065 | 0.0114 |
| rs2314338 | T | 0.0093 | 0.0016 | -0.0157 | 0.0121 |
| rs2321157 | A | -0.0081 | 0.0014 | -0.0040 | 0.0093 |
| rs232496 | T | 0.0090 | 0.0015 | 0.0301 | 0.0094 |
| rs2332179 | A | 0.0121 | 0.0020 | -0.0156 | 0.0128 |
| rs2336721 | T | 0.0095 | 0.0015 | -0.0409 | 0.0099 |
| rs2358628 | A | -0.0091 | 0.0015 | 0.0095 | 0.0100 |
| rs236318 | C | -0.0102 | 0.0017 | -0.0042 | 0.0105 |
| rs2364544 | A | -0.0105 | 0.0014 | 0.0116 | 0.0097 |
| rs2365376 | A | 0.0092 | 0.0015 | 0.0039 | 0.0098 |
| rs2368831 | T | -0.0092 | 0.0014 | 0.0017 | 0.0103 |
| rs2373124 | A | -0.0096 | 0.0016 | 0.0171 | 0.0116 |
| rs2406253 | A | 0.0135 | 0.0018 | 0.0193 | 0.0129 |
| rs2414072 | A | -0.0083 | 0.0014 | -0.0028 | 0.0093 |
| rs2416214 | A | 0.0079 | 0.0014 | -0.0021 | 0.0095 |
| rs2416759 | A | -0.0093 | 0.0015 | 0.0090 | 0.0102 |
| rs2416845 | T | 0.0263 | 0.0048 | -0.0091 | 0.0301 |
| rs2431023 | A | 0.0093 | 0.0014 | -0.0020 | 0.0093 |
| rs2434672 | A | 0.0091 | 0.0014 | 0.0075 | 0.0095 |
| rs2436760 | T | 0.0232 | 0.0042 | 0.0233 | 0.0265 |
| rs2447097 | T | 0.0096 | 0.0014 | 0.0131 | 0.0097 |
| rs2458370 | T | -0.0086 | 0.0015 | 0.0173 | 0.0095 |
| rs2469226 | A | 0.0096 | 0.0017 | -0.0166 | 0.0115 |
| rs2470966 | C | -0.0103 | 0.0015 | 0.0059 | 0.0102 |
| rs2478208 | C | -0.0103 | 0.0014 | -0.0101 | 0.0094 |
| rs2496482 | T | 0.0121 | 0.0015 | -0.0160 | 0.0104 |
| rs2517086 | C | -0.0088 | 0.0014 | -0.0108 | 0.0099 |
| rs2521602 | A | -0.0266 | 0.0049 | 0.0022 | 0.0313 |
| rs2522545 | T | -0.0083 | 0.0014 | 0.0033 | 0.0094 |
| rs2526398 | C | -0.0222 | 0.0015 | 0.0174 | 0.0108 |
| rs2529069 | T | -0.0106 | 0.0016 | 0.0143 | 0.0101 |
| rs252991 | A | 0.0095 | 0.0015 | -0.0054 | 0.0098 |
| rs2542673 | A | 0.0088 | 0.0015 | 0.0100 | 0.0100 |
| rs2545795 | A | 0.0125 | 0.0014 | 0.0017 | 0.0094 |
| rs255053 | A | -0.0111 | 0.0018 | -0.0014 | 0.0115 |
| rs2554835 | A | 0.0088 | 0.0014 | 0.0060 | 0.0099 |
| rs2568955 | T | -0.0165 | 0.0016 | 0.0069 | 0.0112 |
| rs2569041 | T | 0.0077 | 0.0014 | -0.0061 | 0.0097 |
| rs2570497 | T | -0.0121 | 0.0015 | 0.0087 | 0.0098 |
| rs2588959 | T | -0.0080 | 0.0014 | 0.0056 | 0.0092 |
| rs2657283 | T | 0.0085 | 0.0014 | -0.0110 | 0.0097 |
| rs2665668 | A | -0.0094 | 0.0015 | 0.0227 | 0.0107 |
| rs2702576 | A | -0.0090 | 0.0014 | 0.0053 | 0.0094 |
| rs2718277 | T | 0.0142 | 0.0026 | -0.0179 | 0.0186 |
| rs2718791 | T | -0.0081 | 0.0015 | -0.0148 | 0.0101 |
| rs27220 | A | -0.0114 | 0.0015 | 0.0027 | 0.0095 |
| rs2725370 | T | -0.0141 | 0.0015 | 0.0165 | 0.0102 |
| rs2736752 | T | 0.0095 | 0.0017 | -0.0032 | 0.0124 |
| rs2740795 | A | 0.0098 | 0.0016 | -0.0040 | 0.0102 |
| rs2761438 | A | 0.0098 | 0.0015 | -0.0145 | 0.0098 |
| rs2764684 | T | 0.0143 | 0.0019 | -0.0132 | 0.0141 |
| rs2805064 | C | 0.0093 | 0.0016 | 0.0002 | 0.0111 |
| rs2838006 | T | 0.0132 | 0.0015 | -0.0117 | 0.0101 |
| rs28482086 | A | 0.0086 | 0.0015 | -0.0135 | 0.0097 |
| rs28505285 | C | 0.0106 | 0.0018 | 0.0217 | 0.0130 |
| rs28512462 | C | 0.0112 | 0.0014 | -0.0021 | 0.0094 |
| rs28513882 | A | -0.0110 | 0.0018 | -0.0014 | 0.0136 |
| rs2852349 | T | -0.0093 | 0.0014 | 0.0077 | 0.0092 |
| rs28587776 | T | 0.0088 | 0.0015 | 0.0214 | 0.0102 |
| rs28669886 | A | -0.0084 | 0.0015 | 0.0128 | 0.0099 |
| rs2870281 | A | -0.0083 | 0.0014 | -0.0050 | 0.0093 |
| rs28735993 | A | 0.0162 | 0.0022 | -0.0319 | 0.0158 |
| rs2885198 | A | 0.0088 | 0.0014 | -0.0207 | 0.0095 |
| rs2898191 | A | 0.0093 | 0.0015 | 0.0116 | 0.0099 |
| rs2910823 | T | 0.0115 | 0.0014 | 0.0215 | 0.0096 |
| rs2916490 | A | -0.0092 | 0.0015 | 0.0070 | 0.0103 |
| rs2923424 | A | 0.0120 | 0.0014 | 0.0073 | 0.0095 |
| rs2926702 | T | -0.0124 | 0.0021 | -0.0210 | 0.0122 |
| rs2929032 | A | -0.0081 | 0.0014 | -0.0060 | 0.0092 |
| rs2929860 | T | 0.0110 | 0.0018 | 0.0128 | 0.0132 |
| rs2942884 | A | -0.0090 | 0.0014 | 0.0047 | 0.0097 |
| rs2954114 | A | -0.0081 | 0.0014 | 0.0090 | 0.0099 |
| rs2958182 | A | 0.0082 | 0.0015 | -0.0216 | 0.0101 |
| rs2962378 | A | 0.0101 | 0.0017 | -0.0084 | 0.0110 |
| rs2964199 | T | -0.0099 | 0.0015 | 0.0172 | 0.0100 |
| rs2964255 | A | 0.0088 | 0.0015 | 0.0012 | 0.0100 |
| rs2976397 | T | 0.0081 | 0.0014 | 0.0117 | 0.0094 |
| rs2977464 | T | 0.0102 | 0.0018 | 0.0009 | 0.0115 |
| rs29792 | A | -0.0093 | 0.0015 | 0.0016 | 0.0104 |
| rs2980813 | A | 0.0088 | 0.0014 | 0.0037 | 0.0094 |
| rs2989476 | C | 0.0078 | 0.0014 | -0.0080 | 0.0093 |
| rs2989751 | A | -0.0092 | 0.0017 | 0.0104 | 0.0105 |
| rs2992037 | A | 0.0149 | 0.0015 | 0.0170 | 0.0109 |
| rs2994326 | T | -0.0100 | 0.0018 | -0.0036 | 0.0124 |
| rs2998299 | T | -0.0136 | 0.0017 | -0.0019 | 0.0118 |
| rs3026996 | A | 0.0138 | 0.0016 | -0.0163 | 0.0122 |
| rs303752 | A | -0.0110 | 0.0014 | 0.0235 | 0.0104 |
| rs3111251 | T | -0.0090 | 0.0014 | 0.0000 | 0.0096 |
| rs312927 | A | -0.0147 | 0.0022 | -0.0193 | 0.0169 |
| rs312945 | A | -0.0091 | 0.0015 | -0.0306 | 0.0101 |
| rs317050 | T | 0.0078 | 0.0014 | -0.0071 | 0.0093 |
| rs322614 | A | -0.0087 | 0.0014 | 0.0102 | 0.0095 |
| rs322744 | T | -0.0090 | 0.0016 | -0.0035 | 0.0102 |
| rs324885 | A | 0.0142 | 0.0014 | -0.0084 | 0.0096 |
| rs32940 | T | -0.0096 | 0.0015 | 0.0261 | 0.0102 |
| rs339057 | A | -0.0081 | 0.0014 | -0.0052 | 0.0094 |
| rs34067381 | T | -0.0104 | 0.0015 | 0.0094 | 0.0098 |
| rs34098770 | A | -0.0160 | 0.0020 | 0.0023 | 0.0124 |
| rs34106693 | C | 0.0159 | 0.0019 | -0.0248 | 0.0154 |
| rs34122915 | C | 0.0084 | 0.0014 | -0.0006 | 0.0109 |
| rs341504 | A | 0.0099 | 0.0015 | -0.0048 | 0.0097 |
| rs34155847 | A | 0.0135 | 0.0016 | -0.0133 | 0.0112 |
| rs34262657 | T | -0.0240 | 0.0038 | -0.0183 | 0.0214 |
| rs34286836 | T | -0.0089 | 0.0014 | 0.0080 | 0.0094 |
| rs34298584 | A | 0.0111 | 0.0019 | -0.0231 | 0.0128 |
| rs34305371 | A | 0.0314 | 0.0024 | 0.0156 | 0.0182 |
| rs34309 | A | 0.0080 | 0.0014 | 0.0056 | 0.0096 |
| rs34316274 | A | 0.0097 | 0.0017 | 0.0140 | 0.0109 |
| rs34363861 | A | 0.0087 | 0.0014 | -0.0189 | 0.0104 |
| rs34394051 | A | -0.0117 | 0.0020 | 0.0166 | 0.0122 |
| rs34410 | C | -0.0083 | 0.0014 | 0.0204 | 0.0097 |
| rs34624793 | T | 0.0116 | 0.0021 | 0.0200 | 0.0171 |
| rs34720381 | T | -0.0180 | 0.0024 | -0.0028 | 0.0176 |
| rs34748029 | A | 0.0185 | 0.0025 | -0.0268 | 0.0227 |
| rs347661 | T | -0.0081 | 0.0014 | -0.0124 | 0.0097 |
| rs34780702 | A | 0.0111 | 0.0017 | -0.0002 | 0.0106 |
| rs34807077 | A | 0.0128 | 0.0019 | -0.0060 | 0.0122 |
| rs34811474 | A | 0.0104 | 0.0017 | -0.0108 | 0.0150 |
| rs34967082 | A | -0.0080 | 0.0014 | -0.0009 | 0.0098 |
| rs35016816 | T | 0.0176 | 0.0023 | -0.0186 | 0.0146 |
| rs350281 | T | -0.0117 | 0.0015 | 0.0181 | 0.0116 |
| rs35104491 | A | 0.0124 | 0.0018 | 0.0009 | 0.0114 |
| rs35111506 | C | -0.0175 | 0.0030 | 0.0269 | 0.0214 |
| rs35192107 | T | -0.0091 | 0.0017 | -0.0078 | 0.0119 |
| rs35319653 | T | 0.0119 | 0.0015 | 0.0044 | 0.0101 |
| rs35417702 | T | -0.0152 | 0.0014 | 0.0188 | 0.0092 |
| rs35493937 | C | 0.0124 | 0.0017 | -0.0133 | 0.0114 |
| rs35606437 | A | 0.0105 | 0.0016 | -0.0144 | 0.0104 |
| rs356999 | A | -0.0101 | 0.0014 | 0.0148 | 0.0101 |
| rs35745455 | A | 0.0080 | 0.0014 | -0.0088 | 0.0095 |
| rs35751693 | T | -0.0246 | 0.0040 | 0.0328 | 0.0389 |
| rs35754740 | T | 0.0083 | 0.0014 | -0.0127 | 0.0094 |
| rs35811586 | T | 0.0217 | 0.0031 | -0.0516 | 0.0283 |
| rs35929923 | A | -0.0114 | 0.0016 | 0.0293 | 0.0108 |
| rs36085856 | C | 0.0201 | 0.0036 | -0.0029 | 0.0246 |
| rs36120534 | T | 0.0100 | 0.0018 | 0.0119 | 0.0131 |
| rs362307 | T | -0.0226 | 0.0027 | 0.0177 | 0.0199 |
| rs363096 | T | -0.0143 | 0.0014 | 0.0182 | 0.0093 |
| rs3735478 | T | 0.0108 | 0.0016 | -0.0317 | 0.0109 |
| rs3751331 | A | -0.0097 | 0.0014 | 0.0212 | 0.0096 |
| rs3766979 | A | 0.0085 | 0.0015 | -0.0050 | 0.0097 |
| rs3768480 | C | 0.0101 | 0.0014 | -0.0218 | 0.0096 |
| rs3781339 | T | -0.0104 | 0.0018 | 0.0398 | 0.0123 |
| rs3790609 | T | 0.0107 | 0.0019 | 0.0021 | 0.0125 |
| rs3809169 | T | 0.0140 | 0.0025 | 0.0054 | 0.0162 |
| rs3809634 | A | -0.0109 | 0.0015 | 0.0074 | 0.0107 |
| rs3812281 | T | 0.0113 | 0.0014 | -0.0136 | 0.0094 |
| rs3817923 | A | -0.0121 | 0.0022 | 0.0320 | 0.0143 |
| rs382196 | T | -0.0082 | 0.0014 | 0.0082 | 0.0095 |
| rs3847228 | T | 0.0083 | 0.0014 | 0.0078 | 0.0098 |
| rs3848715 | A | -0.0080 | 0.0014 | 0.0108 | 0.0092 |
| rs3859523 | T | 0.0124 | 0.0018 | -0.0257 | 0.0122 |
| rs387027 | A | -0.0083 | 0.0015 | 0.0230 | 0.0097 |
| rs3895736 | A | 0.0144 | 0.0019 | 0.0170 | 0.0126 |
| rs3897821 | A | 0.0154 | 0.0015 | -0.0086 | 0.0098 |
| rs3948495 | T | -0.0089 | 0.0014 | 0.0135 | 0.0098 |
| rs399821 | A | -0.0078 | 0.0014 | -0.0038 | 0.0093 |
| rs39998 | A | 0.0090 | 0.0015 | -0.0164 | 0.0108 |
| rs401526 | T | -0.0104 | 0.0014 | 0.0117 | 0.0093 |
| rs401966 | C | -0.0097 | 0.0014 | -0.0216 | 0.0100 |
| rs4127499 | A | 0.0087 | 0.0015 | -0.0037 | 0.0107 |
| rs41282553 | A | 0.0237 | 0.0040 | -0.0510 | 0.0294 |
| rs42210 | C | -0.0098 | 0.0016 | -0.0257 | 0.0101 |
| rs42302 | A | 0.0086 | 0.0015 | 0.0006 | 0.0097 |
| rs4255791 | A | -0.0084 | 0.0015 | -0.0018 | 0.0113 |
| rs4263475 | A | 0.0078 | 0.0014 | 0.0076 | 0.0102 |
| rs4283754 | A | -0.0084 | 0.0014 | -0.0055 | 0.0115 |
| rs429150 | T | 0.0085 | 0.0014 | -0.0129 | 0.0100 |
| rs4298514 | T | 0.0104 | 0.0015 | -0.0015 | 0.0099 |
| rs4320563 | A | 0.0109 | 0.0014 | -0.0206 | 0.0092 |
| rs4328757 | T | 0.0096 | 0.0014 | 0.0058 | 0.0096 |
| rs4352658 | T | -0.0190 | 0.0026 | 0.0295 | 0.0167 |
| rs4358081 | A | -0.0095 | 0.0014 | 0.0104 | 0.0092 |
| rs4358358 | A | 0.0109 | 0.0015 | -0.0141 | 0.0101 |
| rs4369924 | A | 0.0125 | 0.0019 | -0.0103 | 0.0123 |
| rs4384309 | A | 0.0102 | 0.0014 | -0.0060 | 0.0098 |
| rs4396896 | A | -0.0107 | 0.0014 | -0.0060 | 0.0094 |
| rs4423373 | A | -0.0087 | 0.0014 | 0.0060 | 0.0098 |
| rs4426420 | T | -0.0140 | 0.0019 | 0.0214 | 0.0124 |
| rs4434676 | A | -0.0089 | 0.0014 | 0.0157 | 0.0095 |
| rs4458044 | C | 0.0101 | 0.0016 | 0.0077 | 0.0127 |
| rs4467547 | T | 0.0121 | 0.0014 | -0.0030 | 0.0104 |
| rs4469771 | T | 0.0101 | 0.0015 | -0.0129 | 0.0098 |
| rs4480339 | A | -0.0097 | 0.0015 | 0.0114 | 0.0108 |
| rs4490539 | A | 0.0142 | 0.0015 | -0.0002 | 0.0106 |
| rs4497562 | A | 0.0112 | 0.0016 | -0.0083 | 0.0105 |
| rs4500930 | T | -0.0109 | 0.0015 | 0.0377 | 0.0100 |
| rs4500960 | T | -0.0131 | 0.0014 | 0.0183 | 0.0096 |
| rs4502401 | T | 0.0085 | 0.0015 | -0.0074 | 0.0102 |
| rs4652135 | A | 0.0103 | 0.0016 | 0.0057 | 0.0102 |
| rs4658019 | T | 0.0084 | 0.0014 | -0.0131 | 0.0094 |
| rs4663617 | A | 0.0104 | 0.0017 | -0.0018 | 0.0116 |
| rs4664983 | T | -0.0103 | 0.0018 | 0.0012 | 0.0115 |
| rs4673840 | T | -0.0138 | 0.0019 | 0.0077 | 0.0126 |
| rs4675248 | A | -0.0094 | 0.0014 | -0.0034 | 0.0097 |
| rs4685405 | T | -0.0111 | 0.0018 | -0.0020 | 0.0108 |
| rs4687735 | T | -0.0200 | 0.0033 | -0.0421 | 0.0262 |
| rs4691601 | A | -0.0115 | 0.0014 | 0.0097 | 0.0093 |
| rs4705763 | C | 0.0089 | 0.0014 | 0.0170 | 0.0095 |
| rs4712371 | A | -0.0124 | 0.0021 | 0.0114 | 0.0129 |
| rs4719460 | T | -0.0088 | 0.0015 | 0.0116 | 0.0100 |
| rs4719944 | T | -0.0121 | 0.0014 | -0.0045 | 0.0092 |
| rs4724083 | T | 0.0083 | 0.0015 | -0.0098 | 0.0098 |
| rs4726070 | A | 0.0122 | 0.0014 | -0.0087 | 0.0094 |
| rs4730020 | T | 0.0090 | 0.0016 | -0.0003 | 0.0116 |
| rs4731413 | A | 0.0111 | 0.0017 | -0.0017 | 0.0127 |
| rs4735297 | A | -0.0086 | 0.0015 | -0.0089 | 0.0097 |
| rs4739235 | A | 0.0109 | 0.0019 | -0.0087 | 0.0119 |
| rs4741571 | A | -0.0102 | 0.0015 | 0.0034 | 0.0107 |
| rs4757957 | C | 0.0133 | 0.0015 | -0.0131 | 0.0100 |
| rs4766424 | C | 0.0142 | 0.0021 | -0.0407 | 0.0161 |
| rs4778058 | T | -0.0092 | 0.0014 | 0.0038 | 0.0098 |
| rs4785187 | A | -0.0100 | 0.0017 | -0.0155 | 0.0106 |
| rs4787028 | T | 0.0086 | 0.0015 | 0.0053 | 0.0099 |
| rs4787457 | A | 0.0162 | 0.0015 | -0.0142 | 0.0100 |
| rs4788115 | A | 0.0111 | 0.0019 | -0.0242 | 0.0157 |
| rs479018 | A | 0.0109 | 0.0015 | -0.0002 | 0.0112 |
| rs4793090 | A | 0.0084 | 0.0015 | -0.0391 | 0.0096 |
| rs4810894 | A | -0.0085 | 0.0015 | 0.0165 | 0.0096 |
| rs4812697 | T | 0.0171 | 0.0027 | -0.0065 | 0.0188 |
| rs481940 | T | 0.0100 | 0.0016 | -0.0047 | 0.0113 |
| rs482787 | T | 0.0100 | 0.0015 | -0.0146 | 0.0100 |
| rs4839155 | T | 0.0115 | 0.0017 | -0.0089 | 0.0110 |
| rs4846010 | A | 0.0099 | 0.0018 | -0.0313 | 0.0130 |
| rs4846724 | A | 0.0098 | 0.0014 | -0.0108 | 0.0092 |
| rs4848924 | A | -0.0117 | 0.0015 | -0.0051 | 0.0100 |
| rs4850954 | T | 0.0077 | 0.0014 | 0.0030 | 0.0100 |
| rs4851263 | A | -0.0151 | 0.0023 | 0.0342 | 0.0157 |
| rs4877516 | A | -0.0101 | 0.0014 | 0.0157 | 0.0093 |
| rs4881269 | A | 0.0100 | 0.0014 | 0.0199 | 0.0095 |
| rs488476 | C | -0.0113 | 0.0015 | 0.0014 | 0.0103 |
| rs4894658 | C | 0.0091 | 0.0016 | -0.0145 | 0.0102 |
| rs4895650 | T | 0.0093 | 0.0014 | 0.0003 | 0.0101 |
| rs4899012 | C | -0.0103 | 0.0014 | 0.0144 | 0.0096 |
| rs4904523 | A | -0.0082 | 0.0014 | 0.0087 | 0.0093 |
| rs4915735 | A | 0.0116 | 0.0020 | 0.0128 | 0.0142 |
| rs4919624 | A | 0.0193 | 0.0018 | -0.0195 | 0.0119 |
| rs4925065 | T | -0.0078 | 0.0014 | 0.0007 | 0.0095 |
| rs4925109 | A | -0.0096 | 0.0015 | 0.0296 | 0.0100 |
| rs4938815 | T | 0.0085 | 0.0015 | -0.0034 | 0.0102 |
| rs4941735 | T | 0.0092 | 0.0014 | -0.0027 | 0.0096 |
| rs4972748 | T | 0.0105 | 0.0018 | 0.0039 | 0.0116 |
| rs4977885 | A | -0.0084 | 0.0014 | 0.0203 | 0.0097 |
| rs4981245 | A | 0.0080 | 0.0014 | -0.0107 | 0.0099 |
| rs4984541 | A | -0.0138 | 0.0017 | 0.0026 | 0.0115 |
| rs4984613 | T | 0.0174 | 0.0028 | -0.0009 | 0.0174 |
| rs4984682 | C | -0.0138 | 0.0017 | -0.0042 | 0.0110 |
| rs548897 | A | 0.0081 | 0.0014 | -0.0128 | 0.0095 |
| rs55675587 | T | -0.0103 | 0.0018 | 0.0143 | 0.0115 |
| rs55736314 | C | -0.0154 | 0.0014 | 0.0216 | 0.0102 |
| rs55826493 | T | -0.0127 | 0.0021 | 0.0272 | 0.0145 |
| rs55986781 | T | 0.0187 | 0.0026 | -0.0499 | 0.0209 |
| rs56085180 | A | -0.0269 | 0.0037 | 0.0019 | 0.0238 |
| rs56099375 | T | 0.0120 | 0.0017 | -0.0083 | 0.0125 |
| rs56171318 | T | -0.0137 | 0.0020 | -0.0096 | 0.0128 |
| rs56174996 | A | -0.0143 | 0.0020 | 0.0079 | 0.0136 |
| rs56306882 | A | 0.0086 | 0.0016 | -0.0188 | 0.0115 |
| rs56319902 | T | -0.0206 | 0.0017 | 0.0433 | 0.0136 |
| rs56391344 | A | 0.0148 | 0.0016 | -0.0328 | 0.0117 |
| rs563954 | A | -0.0081 | 0.0014 | 0.0044 | 0.0096 |
| rs56405138 | A | 0.0153 | 0.0022 | -0.0020 | 0.0139 |
| rs56408528 | T | -0.0082 | 0.0014 | 0.0115 | 0.0096 |
| rs567003 | A | 0.0094 | 0.0015 | 0.0113 | 0.0103 |
| rs56794817 | A | 0.0122 | 0.0019 | -0.0068 | 0.0121 |
| rs57016874 | T | 0.0209 | 0.0037 | 0.0050 | 0.0293 |
| rs57148205 | A | 0.0095 | 0.0016 | 0.0069 | 0.0115 |
| rs57204268 | A | 0.0114 | 0.0020 | 0.0000 | 0.0138 |
| rs57349798 | A | 0.0093 | 0.0014 | -0.0462 | 0.0100 |
| rs57352738 | A | -0.0152 | 0.0017 | -0.0002 | 0.0123 |
| rs57437407 | A | -0.0145 | 0.0023 | -0.0140 | 0.0181 |
| rs575113 | A | 0.0133 | 0.0015 | -0.0159 | 0.0104 |
| rs5754581 | T | -0.0082 | 0.0014 | 0.0127 | 0.0093 |
| rs5754753 | T | -0.0126 | 0.0016 | 0.0001 | 0.0106 |
| rs5754762 | A | 0.0197 | 0.0030 | -0.0235 | 0.0174 |
| rs57661533 | T | -0.0114 | 0.0021 | 0.0029 | 0.0120 |
| rs580652 | T | -0.0147 | 0.0024 | -0.0112 | 0.0135 |
| rs585557 | A | 0.0128 | 0.0018 | -0.0220 | 0.0121 |
| rs58779949 | A | 0.0110 | 0.0019 | -0.0319 | 0.0137 |
| rs58859557 | T | 0.0187 | 0.0028 | 0.0124 | 0.0192 |
| rs58921703 | T | 0.0109 | 0.0015 | -0.0045 | 0.0099 |
| rs58950082 | T | 0.0095 | 0.0017 | -0.0058 | 0.0112 |
| rs58996896 | A | -0.0094 | 0.0016 | -0.0071 | 0.0113 |
| rs590013 | T | 0.0100 | 0.0015 | -0.0062 | 0.0099 |
| rs59300999 | T | -0.0164 | 0.0028 | 0.0078 | 0.0147 |
| rs59480703 | C | -0.0123 | 0.0018 | -0.0235 | 0.0114 |
| rs59813324 | T | 0.0097 | 0.0018 | -0.0053 | 0.0111 |
| rs59967356 | T | -0.0127 | 0.0021 | 0.0258 | 0.0180 |
| rs60096640 | A | 0.0159 | 0.0023 | 0.0005 | 0.0125 |
| rs6020560 | T | -0.0083 | 0.0014 | -0.0057 | 0.0092 |
| rs6043521 | T | -0.0083 | 0.0014 | 0.0159 | 0.0100 |
| rs60589532 | A | 0.0168 | 0.0028 | -0.0078 | 0.0188 |
| rs6060308 | A | 0.0095 | 0.0016 | -0.0184 | 0.0103 |
| rs6065080 | T | -0.0130 | 0.0015 | 0.0042 | 0.0095 |
| rs6065784 | C | 0.0110 | 0.0015 | -0.0012 | 0.0104 |
| rs60717745 | C | -0.0118 | 0.0019 | 0.0148 | 0.0121 |
| rs60726488 | T | -0.0090 | 0.0016 | 0.0002 | 0.0111 |
| rs6091570 | A | 0.0081 | 0.0015 | -0.0086 | 0.0096 |
| rs61104616 | A | -0.0172 | 0.0014 | -0.0007 | 0.0094 |
| rs6123924 | A | 0.0129 | 0.0019 | -0.0348 | 0.0119 |
| rs613872 | T | -0.0163 | 0.0019 | 0.0232 | 0.0131 |
| rs61387839 | A | 0.0088 | 0.0015 | -0.0123 | 0.0111 |
| rs61527214 | A | 0.0110 | 0.0014 | -0.0121 | 0.0101 |
| rs61739710 | A | -0.0099 | 0.0016 | -0.0133 | 0.0106 |
| rs61748951 | A | -0.0273 | 0.0048 | -0.0138 | 0.0458 |
| rs61755388 | T | 0.0125 | 0.0017 | -0.0005 | 0.0121 |
| rs61757207 | A | 0.0359 | 0.0063 | 0.0794 | 0.0445 |
| rs61798586 | A | -0.0112 | 0.0020 | 0.0160 | 0.0141 |
| rs61853335 | T | -0.0109 | 0.0018 | -0.0084 | 0.0116 |
| rs61958175 | A | -0.0198 | 0.0034 | 0.0285 | 0.0299 |
| rs61997667 | T | -0.0144 | 0.0020 | 0.0074 | 0.0135 |
| rs62007304 | A | 0.0264 | 0.0047 | -0.0001 | 0.0457 |
| rs62018216 | T | 0.0113 | 0.0019 | 0.0087 | 0.0137 |
| rs62051146 | A | -0.0134 | 0.0022 | 0.0248 | 0.0139 |
| rs62090515 | A | -0.0090 | 0.0015 | 0.0059 | 0.0098 |
| rs62092949 | T | -0.0087 | 0.0014 | -0.0132 | 0.0096 |
| rs62103198 | T | -0.0147 | 0.0025 | 0.0077 | 0.0208 |
| rs62109862 | A | 0.0136 | 0.0020 | 0.0155 | 0.0129 |
| rs62142891 | A | 0.0108 | 0.0016 | 0.0029 | 0.0117 |
| rs62155350 | A | 0.0338 | 0.0053 | -0.0257 | 0.0431 |
| rs62155770 | A | -0.0196 | 0.0032 | 0.0468 | 0.0270 |
| rs62155873 | T | -0.0151 | 0.0021 | -0.0192 | 0.0158 |
| rs62174974 | A | -0.0105 | 0.0018 | 0.0115 | 0.0114 |
| rs62177359 | A | -0.0275 | 0.0038 | 0.0148 | 0.0300 |
| rs62179650 | A | 0.0112 | 0.0016 | 0.0055 | 0.0107 |
| rs62182994 | T | 0.0137 | 0.0015 | -0.0077 | 0.0108 |
| rs62190914 | T | 0.0089 | 0.0015 | -0.0062 | 0.0103 |
| rs62194170 | A | 0.0111 | 0.0017 | -0.0125 | 0.0119 |
| rs622169 | T | 0.0086 | 0.0014 | -0.0114 | 0.0112 |
| rs62247449 | C | 0.0114 | 0.0014 | -0.0062 | 0.0097 |
| rs62252819 | A | -0.0094 | 0.0017 | -0.0032 | 0.0113 |
| rs62256284 | T | 0.0111 | 0.0017 | 0.0079 | 0.0113 |
| rs62260764 | C | -0.0132 | 0.0016 | -0.0168 | 0.0109 |
| rs62262671 | A | 0.0213 | 0.0020 | -0.0199 | 0.0156 |
| rs62340636 | T | 0.0103 | 0.0014 | -0.0043 | 0.0094 |
| rs62370510 | A | -0.0109 | 0.0019 | -0.0070 | 0.0133 |
| rs62379838 | T | 0.0121 | 0.0015 | -0.0192 | 0.0103 |
| rs62409395 | T | 0.0101 | 0.0018 | -0.0111 | 0.0167 |
| rs62420387 | T | -0.0184 | 0.0027 | 0.0316 | 0.0204 |
| rs62506074 | T | 0.0091 | 0.0015 | 0.0058 | 0.0110 |
| rs62506104 | A | -0.0098 | 0.0018 | 0.0124 | 0.0115 |
| rs628993 | A | 0.0128 | 0.0023 | -0.0068 | 0.0145 |
| rs631287 | A | 0.0084 | 0.0014 | -0.0255 | 0.0092 |
| rs633279 | A | -0.0089 | 0.0015 | -0.0061 | 0.0102 |
| rs635754 | A | 0.0119 | 0.0014 | -0.0004 | 0.0092 |
| rs6428587 | T | 0.0084 | 0.0015 | -0.0039 | 0.0095 |
| rs6435326 | A | -0.0091 | 0.0014 | -0.0057 | 0.0097 |
| rs6442126 | A | -0.0121 | 0.0015 | 0.0301 | 0.0103 |
| rs6445633 | A | 0.0088 | 0.0015 | -0.0003 | 0.0112 |
| rs6449503 | A | 0.0185 | 0.0014 | 0.0042 | 0.0095 |
| rs6450476 | A | -0.0116 | 0.0015 | 0.0075 | 0.0102 |
| rs6452793 | T | 0.0175 | 0.0017 | -0.0176 | 0.0111 |
| rs6461536 | A | 0.0088 | 0.0015 | 0.0054 | 0.0102 |
| rs6469654 | C | -0.0103 | 0.0017 | -0.0092 | 0.0104 |
| rs6472208 | T | -0.0091 | 0.0014 | 0.0039 | 0.0097 |
| rs6480234 | T | -0.0084 | 0.0014 | 0.0043 | 0.0094 |
| rs648044 | A | 0.0083 | 0.0015 | 0.0155 | 0.0104 |
| rs6490618 | T | 0.0088 | 0.0015 | -0.0020 | 0.0098 |
| rs6493265 | T | -0.0122 | 0.0014 | 0.0137 | 0.0097 |
| rs6493275 | A | 0.0131 | 0.0017 | 0.0108 | 0.0120 |
| rs6503409 | T | 0.0087 | 0.0015 | 0.0047 | 0.0105 |
| rs6534338 | T | 0.0103 | 0.0015 | -0.0153 | 0.0105 |
| rs6543810 | T | 0.0082 | 0.0015 | 0.0137 | 0.0095 |
| rs6557171 | T | -0.0155 | 0.0015 | -0.0031 | 0.0096 |
| rs6567288 | A | -0.0079 | 0.0014 | 0.0047 | 0.0093 |
| rs6569077 | T | 0.0118 | 0.0014 | -0.0033 | 0.0094 |
| rs6573552 | T | -0.0090 | 0.0014 | 0.0031 | 0.0092 |
| rs6573559 | T | 0.0105 | 0.0015 | 0.0105 | 0.0100 |
| rs660001 | A | -0.0164 | 0.0017 | 0.0210 | 0.0114 |
| rs663251 | T | 0.0081 | 0.0015 | 0.0064 | 0.0095 |
| rs66482320 | C | 0.0124 | 0.0022 | 0.0104 | 0.0149 |
| rs66568921 | T | -0.0181 | 0.0015 | 0.0078 | 0.0097 |
| rs66671632 | T | -0.0133 | 0.0021 | 0.0084 | 0.0133 |
| rs66721975 | A | -0.0087 | 0.0015 | 0.0008 | 0.0107 |
| rs6672986 | C | 0.0094 | 0.0014 | 0.0020 | 0.0092 |
| rs6689641 | A | 0.0078 | 0.0014 | -0.0002 | 0.0094 |
| rs6690195 | T | -0.0113 | 0.0014 | 0.0110 | 0.0098 |
| rs6695132 | T | 0.0097 | 0.0017 | -0.0148 | 0.0105 |
| rs6696068 | T | -0.0088 | 0.0014 | 0.0036 | 0.0098 |
| rs6697584 | T | -0.0126 | 0.0017 | 0.0258 | 0.0121 |
| rs6704768 | A | -0.0116 | 0.0014 | 0.0237 | 0.0096 |
| rs6706275 | T | 0.0083 | 0.0015 | -0.0075 | 0.0108 |
| rs6711399 | T | -0.0107 | 0.0019 | 0.0175 | 0.0114 |
| rs6715321 | T | -0.0097 | 0.0014 | 0.0099 | 0.0099 |
| rs6715849 | A | -0.0122 | 0.0014 | 0.0021 | 0.0097 |
| rs6720515 | A | 0.0103 | 0.0017 | -0.0042 | 0.0111 |
| rs67224963 | A | 0.0113 | 0.0017 | -0.0037 | 0.0113 |
| rs6729612 | T | 0.0092 | 0.0015 | 0.0056 | 0.0101 |
| rs6731373 | A | -0.0115 | 0.0015 | -0.0103 | 0.0113 |
| rs6736025 | T | 0.0079 | 0.0014 | -0.0164 | 0.0095 |
| rs6738860 | A | -0.0088 | 0.0014 | 0.0040 | 0.0096 |
| rs6743032 | A | -0.0165 | 0.0024 | -0.0253 | 0.0180 |
| rs6744040 | A | -0.0080 | 0.0014 | 0.0074 | 0.0093 |
| rs67456868 | A | 0.0119 | 0.0018 | -0.0031 | 0.0132 |
| rs6752228 | T | -0.0080 | 0.0014 | -0.0016 | 0.0094 |
| rs6757087 | T | -0.0085 | 0.0014 | 0.0024 | 0.0100 |
| rs6760887 | T | 0.0155 | 0.0022 | -0.0152 | 0.0119 |
| rs67661275 | A | 0.0134 | 0.0025 | -0.0104 | 0.0177 |
| rs6774533 | T | 0.0090 | 0.0015 | 0.0059 | 0.0099 |
| rs67790232 | A | 0.0101 | 0.0017 | 0.0036 | 0.0126 |
| rs6780414 | T | 0.0105 | 0.0018 | 0.0058 | 0.0119 |
| rs6792805 | A | -0.0090 | 0.0016 | -0.0034 | 0.0101 |
| rs6812533 | T | 0.0106 | 0.0016 | 0.0107 | 0.0103 |
| rs68145588 | T | -0.0127 | 0.0021 | 0.0021 | 0.0148 |
| rs6824567 | T | 0.0130 | 0.0016 | -0.0049 | 0.0111 |
| rs6853599 | T | 0.0108 | 0.0019 | -0.0116 | 0.0140 |
| rs6861925 | C | -0.0090 | 0.0014 | 0.0049 | 0.0092 |
| rs6871635 | A | -0.0087 | 0.0014 | -0.0151 | 0.0099 |
| rs6881581 | A | -0.0096 | 0.0015 | 0.0301 | 0.0101 |
| rs6881733 | T | -0.0086 | 0.0016 | 0.0023 | 0.0104 |
| rs6917154 | T | 0.0127 | 0.0021 | 0.0033 | 0.0159 |
| rs6917204 | T | 0.0119 | 0.0017 | -0.0270 | 0.0105 |
| rs6918506 | C | 0.0113 | 0.0014 | 0.0132 | 0.0093 |
| rs6924023 | A | 0.0107 | 0.0016 | -0.0012 | 0.0106 |
| rs6932108 | C | -0.0210 | 0.0027 | 0.0182 | 0.0195 |
| rs6946136 | A | 0.0081 | 0.0014 | 0.0001 | 0.0097 |
| rs6946362 | T | 0.0091 | 0.0015 | -0.0193 | 0.0109 |
| rs6951996 | A | -0.0182 | 0.0032 | 0.0352 | 0.0205 |
| rs6956283 | T | 0.0098 | 0.0016 | -0.0102 | 0.0105 |
| rs6969783 | A | -0.0098 | 0.0014 | 0.0209 | 0.0093 |
| rs6977237 | T | 0.0082 | 0.0014 | -0.0061 | 0.0094 |
| rs6989141 | A | 0.0100 | 0.0017 | -0.0077 | 0.0131 |
| rs6994287 | A | 0.0102 | 0.0014 | -0.0039 | 0.0096 |
| rs7012546 | T | 0.0092 | 0.0014 | -0.0101 | 0.0095 |
| rs7016302 | C | -0.0125 | 0.0019 | 0.0080 | 0.0130 |
| rs702606 | T | 0.0118 | 0.0021 | -0.0001 | 0.0134 |
| rs7029718 | A | 0.0244 | 0.0014 | -0.0030 | 0.0095 |
| rs7030373 | A | -0.0098 | 0.0017 | 0.0092 | 0.0107 |
| rs7035315 | A | -0.0090 | 0.0015 | -0.0177 | 0.0096 |
| rs7040995 | C | 0.0097 | 0.0014 | -0.0240 | 0.0093 |
| rs7041702 | A | -0.0111 | 0.0016 | 0.0149 | 0.0110 |
| rs7084508 | T | -0.0085 | 0.0015 | -0.0067 | 0.0103 |
| rs710629 | A | 0.0082 | 0.0015 | -0.0081 | 0.0096 |
| rs7108020 | A | 0.0111 | 0.0015 | -0.0089 | 0.0101 |
| rs711793 | T | 0.0103 | 0.0015 | -0.0046 | 0.0117 |
| rs7127580 | C | -0.0150 | 0.0023 | 0.0042 | 0.0170 |
| rs713584 | A | -0.0080 | 0.0014 | 0.0011 | 0.0093 |
| rs71413877 | A | 0.0351 | 0.0036 | -0.0326 | 0.0307 |
| rs7146625 | A | 0.0091 | 0.0016 | -0.0215 | 0.0109 |
| rs7147473 | A | -0.0103 | 0.0015 | 0.0161 | 0.0106 |
| rs7158218 | A | -0.0090 | 0.0015 | -0.0015 | 0.0106 |
| rs71646142 | T | 0.0104 | 0.0018 | -0.0003 | 0.0124 |
| rs716513 | A | 0.0084 | 0.0014 | -0.0092 | 0.0094 |
| rs7167688 | T | -0.0094 | 0.0014 | 0.0023 | 0.0093 |
| rs7171405 | A | -0.0101 | 0.0016 | 0.0135 | 0.0107 |
| rs717996 | T | -0.0122 | 0.0014 | 0.0217 | 0.0099 |
| rs7182216 | C | -0.0096 | 0.0017 | -0.0125 | 0.0122 |
| rs7190 | A | -0.0095 | 0.0015 | -0.0009 | 0.0110 |
| rs721579 | T | -0.0110 | 0.0016 | 0.0102 | 0.0104 |
| rs7218235 | A | -0.0097 | 0.0017 | 0.0198 | 0.0124 |
| rs7223311 | C | -0.0079 | 0.0014 | 0.0109 | 0.0100 |
| rs7226824 | T | -0.0078 | 0.0014 | -0.0031 | 0.0093 |
| rs72482130 | T | 0.0141 | 0.0019 | -0.0249 | 0.0128 |
| rs72486027 | T | -0.0104 | 0.0016 | 0.0021 | 0.0105 |
| rs7254263 | T | -0.0119 | 0.0016 | 0.0052 | 0.0102 |
| rs7255223 | A | 0.0097 | 0.0016 | -0.0110 | 0.0103 |
| rs72622559 | T | -0.0120 | 0.0017 | 0.0086 | 0.0110 |
| rs72624911 | T | 0.0282 | 0.0034 | -0.0947 | 0.0277 |
| rs72636697 | T | -0.0140 | 0.0020 | 0.0175 | 0.0129 |
| rs72667460 | T | 0.0197 | 0.0031 | -0.0122 | 0.0215 |
| rs72671456 | A | -0.0174 | 0.0020 | 0.0294 | 0.0147 |
| rs72672052 | A | -0.0101 | 0.0018 | 0.0198 | 0.0120 |
| rs72673097 | A | 0.0257 | 0.0042 | 0.0444 | 0.0410 |
| rs72677177 | A | 0.0100 | 0.0014 | -0.0009 | 0.0098 |
| rs72694479 | T | 0.0127 | 0.0020 | 0.0005 | 0.0142 |
| rs72709560 | A | 0.0085 | 0.0015 | -0.0143 | 0.0112 |
| rs72771860 | T | -0.0211 | 0.0029 | -0.0217 | 0.0253 |
| rs72792395 | T | -0.0145 | 0.0022 | 0.0051 | 0.0146 |
| rs728054 | A | -0.0128 | 0.0015 | 0.0109 | 0.0096 |
| rs72819118 | T | -0.0231 | 0.0024 | -0.0043 | 0.0195 |
| rs72824753 | T | 0.0133 | 0.0024 | -0.0158 | 0.0152 |
| rs72828517 | T | -0.0168 | 0.0019 | 0.0066 | 0.0127 |
| rs72829857 | A | -0.0149 | 0.0016 | 0.0162 | 0.0109 |
| rs72834698 | A | 0.0117 | 0.0020 | -0.0140 | 0.0160 |
| rs72881110 | A | 0.0146 | 0.0027 | -0.0056 | 0.0176 |
| rs72883760 | A | 0.0101 | 0.0018 | -0.0363 | 0.0121 |
| rs72896637 | T | 0.0162 | 0.0028 | -0.0202 | 0.0245 |
| rs72902523 | T | -0.0103 | 0.0016 | -0.0192 | 0.0104 |
| rs72906124 | T | 0.0185 | 0.0030 | -0.0254 | 0.0227 |
| rs72917504 | T | -0.0178 | 0.0030 | -0.0167 | 0.0217 |
| rs72919450 | T | -0.0132 | 0.0023 | 0.0009 | 0.0156 |
| rs72944064 | T | -0.0098 | 0.0016 | -0.0132 | 0.0115 |
| rs72962169 | T | -0.0174 | 0.0019 | -0.0069 | 0.0125 |
| rs72972965 | A | 0.0083 | 0.0015 | 0.0037 | 0.0106 |
| rs72993796 | T | 0.0127 | 0.0022 | 0.0106 | 0.0181 |
| rs73034295 | A | 0.0099 | 0.0018 | -0.0151 | 0.0129 |
| rs730384 | A | 0.0101 | 0.0014 | -0.0108 | 0.0096 |
| rs73039077 | C | 0.0092 | 0.0016 | -0.0097 | 0.0119 |
| rs73055556 | A | 0.0111 | 0.0020 | -0.0058 | 0.0134 |
| rs73082325 | C | -0.0252 | 0.0044 | 0.0696 | 0.0393 |
| rs73106136 | T | -0.0147 | 0.0026 | -0.0057 | 0.0192 |
| rs73154546 | A | 0.0197 | 0.0036 | -0.0070 | 0.0291 |
| rs7317761 | C | 0.0109 | 0.0016 | 0.0071 | 0.0100 |
| rs73191311 | A | -0.0094 | 0.0015 | -0.0093 | 0.0103 |
| rs7321274 | A | 0.0114 | 0.0018 | -0.0181 | 0.0115 |
| rs73219806 | A | 0.0106 | 0.0019 | -0.0096 | 0.0133 |
| rs7323027 | A | 0.0130 | 0.0015 | -0.0081 | 0.0096 |
| rs7326331 | A | -0.0128 | 0.0016 | 0.0060 | 0.0103 |
| rs73344830 | A | 0.0170 | 0.0014 | -0.0020 | 0.0098 |
| rs73405293 | A | 0.0113 | 0.0020 | -0.0022 | 0.0137 |
| rs73457936 | A | 0.0144 | 0.0025 | 0.0203 | 0.0194 |
| rs73518807 | C | -0.0128 | 0.0021 | 0.0179 | 0.0109 |
| rs7356536 | T | -0.0129 | 0.0021 | 0.0021 | 0.0143 |
| rs736281 | T | 0.0086 | 0.0014 | 0.0161 | 0.0098 |
| rs73643713 | T | -0.0165 | 0.0027 | 0.0046 | 0.0177 |
| rs736471 | T | 0.0083 | 0.0014 | 0.0049 | 0.0101 |
| rs73648455 | T | -0.0183 | 0.0026 | 0.0148 | 0.0167 |
| rs737945 | C | -0.0115 | 0.0014 | 0.0044 | 0.0094 |
| rs73874335 | T | -0.0178 | 0.0029 | -0.0046 | 0.0170 |
| rs73961845 | A | -0.0114 | 0.0014 | 0.0035 | 0.0102 |
| rs74091672 | A | -0.0153 | 0.0022 | 0.0101 | 0.0159 |
| rs7430651 | T | -0.0111 | 0.0016 | 0.0088 | 0.0101 |
| rs743316 | T | 0.0112 | 0.0017 | 0.0071 | 0.0119 |
| rs74415461 | T | 0.0176 | 0.0026 | -0.0227 | 0.0211 |
| rs74453875 | A | 0.0214 | 0.0037 | -0.0112 | 0.0219 |
| rs74462621 | T | 0.0125 | 0.0020 | 0.0129 | 0.0146 |
| rs7449561 | A | 0.0102 | 0.0017 | 0.0192 | 0.0110 |
| rs7451726 | A | 0.0109 | 0.0017 | 0.0070 | 0.0112 |
| rs74545339 | A | -0.0158 | 0.0022 | -0.0141 | 0.0149 |
| rs7460106 | T | -0.0123 | 0.0017 | 0.0137 | 0.0120 |
| rs74615093 | A | 0.0145 | 0.0026 | -0.0002 | 0.0190 |
| rs746839 | C | 0.0135 | 0.0015 | 0.0021 | 0.0103 |
| rs74747621 | T | 0.0153 | 0.0022 | -0.0274 | 0.0159 |
| rs74787922 | A | 0.0182 | 0.0028 | -0.0133 | 0.0208 |
| rs74944275 | T | 0.0231 | 0.0035 | -0.0363 | 0.0287 |
| rs74944857 | T | -0.0089 | 0.0015 | 0.0078 | 0.0105 |
| rs7495033 | A | 0.0095 | 0.0015 | -0.0116 | 0.0100 |
| rs75033012 | C | -0.0258 | 0.0038 | -0.0127 | 0.0343 |
| rs75177132 | T | 0.0298 | 0.0036 | -0.0102 | 0.0386 |
| rs75203411 | T | 0.0135 | 0.0019 | -0.0070 | 0.0132 |
| rs7526112 | T | 0.0124 | 0.0015 | 0.0036 | 0.0096 |
| rs75308819 | A | 0.0287 | 0.0047 | 0.0006 | 0.0375 |
| rs75434274 | A | 0.0151 | 0.0026 | -0.0382 | 0.0201 |
| rs75500877 | T | 0.0207 | 0.0033 | -0.0145 | 0.0255 |
| rs7552964 | A | 0.0148 | 0.0023 | -0.0223 | 0.0158 |
| rs7560871 | A | -0.0193 | 0.0027 | 0.0067 | 0.0198 |
| rs7561705 | A | 0.0078 | 0.0014 | -0.0076 | 0.0098 |
| rs7575637 | A | 0.0107 | 0.0014 | 0.0301 | 0.0096 |
| rs75756843 | A | 0.0288 | 0.0049 | -0.0137 | 0.0353 |
| rs7575938 | A | 0.0090 | 0.0015 | -0.0033 | 0.0099 |
| rs7590368 | T | -0.0146 | 0.0016 | -0.0031 | 0.0108 |
| rs7597126 | T | -0.0094 | 0.0014 | -0.0010 | 0.0101 |
| rs7597412 | C | 0.0088 | 0.0015 | 0.0093 | 0.0103 |
| rs7603132 | A | 0.0137 | 0.0018 | -0.0240 | 0.0116 |
| rs76076331 | T | 0.0206 | 0.0021 | -0.0116 | 0.0144 |
| rs76077165 | A | 0.0165 | 0.0025 | -0.0169 | 0.0186 |
| rs76167224 | T | -0.0187 | 0.0033 | -0.0182 | 0.0310 |
| rs7617204 | A | -0.0097 | 0.0014 | -0.0058 | 0.0093 |
| rs76235882 | A | 0.0241 | 0.0039 | -0.0220 | 0.0278 |
| rs76241605 | A | 0.0139 | 0.0022 | 0.0092 | 0.0186 |
| rs76246107 | A | -0.0162 | 0.0026 | -0.0253 | 0.0227 |
| rs7625428 | T | 0.0098 | 0.0014 | -0.0008 | 0.0096 |
| rs76267866 | A | 0.0104 | 0.0017 | -0.0124 | 0.0118 |
| rs7630133 | A | 0.0082 | 0.0015 | -0.0118 | 0.0094 |
| rs763553 | A | 0.0087 | 0.0014 | -0.0026 | 0.0093 |
| rs7640424 | T | 0.0113 | 0.0015 | 0.0023 | 0.0108 |
| rs7650602 | T | -0.0089 | 0.0014 | 0.0224 | 0.0097 |
| rs76552497 | T | -0.0120 | 0.0019 | 0.0192 | 0.0128 |
| rs76577427 | C | 0.0203 | 0.0024 | -0.0196 | 0.0201 |
| rs7672622 | A | -0.0088 | 0.0016 | 0.0111 | 0.0102 |
| rs767943 | A | -0.0122 | 0.0016 | 0.0224 | 0.0102 |
| rs7683416 | T | 0.0139 | 0.0014 | -0.0046 | 0.0094 |
| rs76876592 | A | 0.0126 | 0.0020 | 0.0062 | 0.0138 |
| rs76878669 | C | 0.0141 | 0.0017 | -0.0052 | 0.0112 |
| rs7692359 | T | -0.0109 | 0.0017 | 0.0264 | 0.0117 |
| rs76957677 | T | -0.0297 | 0.0054 | 0.0441 | 0.0619 |
| rs77025239 | A | -0.0140 | 0.0019 | -0.0016 | 0.0127 |
| rs7714719 | C | 0.0116 | 0.0020 | 0.0314 | 0.0133 |
| rs7716161 | C | 0.0105 | 0.0019 | 0.0237 | 0.0128 |
| rs77201694 | A | 0.0156 | 0.0022 | -0.0284 | 0.0188 |
| rs77370942 | A | -0.0147 | 0.0027 | -0.0055 | 0.0216 |
| rs77554090 | T | -0.0186 | 0.0027 | 0.0396 | 0.0209 |
| rs77609760 | A | 0.0236 | 0.0028 | -0.0159 | 0.0218 |
| rs7766240 | T | -0.0134 | 0.0024 | 0.0205 | 0.0168 |
| rs77702622 | A | -0.0213 | 0.0029 | 0.0268 | 0.0230 |
| rs77702819 | T | 0.0187 | 0.0025 | -0.0072 | 0.0198 |
| rs7772172 | A | 0.0091 | 0.0014 | -0.0081 | 0.0096 |
| rs7775100 | T | -0.0087 | 0.0014 | -0.0210 | 0.0096 |
| rs77826402 | T | 0.0151 | 0.0027 | 0.0042 | 0.0180 |
| rs7796103 | C | -0.0077 | 0.0014 | 0.0058 | 0.0092 |
| rs7799141 | A | 0.0099 | 0.0015 | -0.0139 | 0.0098 |
| rs77999825 | A | 0.0294 | 0.0039 | -0.0297 | 0.0332 |
| rs7803932 | A | 0.0127 | 0.0019 | -0.0150 | 0.0135 |
| rs780569 | A | -0.0085 | 0.0016 | 0.0061 | 0.0104 |
| rs78116078 | C | 0.0086 | 0.0016 | 0.0049 | 0.0111 |
| rs7816777 | T | -0.0088 | 0.0015 | -0.0031 | 0.0100 |
| rs78193153 | A | 0.0191 | 0.0029 | -0.0338 | 0.0164 |
| rs7823700 | T | 0.0132 | 0.0024 | -0.0152 | 0.0180 |
| rs78365243 | T | 0.0181 | 0.0033 | 0.0190 | 0.0268 |
| rs78440611 | A | -0.0147 | 0.0024 | 0.0228 | 0.0156 |
| rs7849480 | A | -0.0084 | 0.0014 | 0.0180 | 0.0094 |
| rs7849487 | T | -0.0167 | 0.0015 | -0.0149 | 0.0098 |
| rs7855503 | C | 0.0088 | 0.0015 | 0.0048 | 0.0098 |
| rs78648104 | T | -0.0141 | 0.0025 | 0.0118 | 0.0168 |
| rs78702390 | C | 0.0167 | 0.0030 | -0.0148 | 0.0226 |
| rs78714229 | T | 0.0172 | 0.0030 | 0.0320 | 0.0217 |
| rs7875078 | A | -0.0078 | 0.0014 | 0.0082 | 0.0095 |
| rs78918150 | T | -0.0080 | 0.0015 | 0.0048 | 0.0099 |
| rs7894722 | T | 0.0085 | 0.0014 | 0.0007 | 0.0099 |
| rs7899270 | A | -0.0084 | 0.0015 | 0.0078 | 0.0101 |
| rs7905192 | T | 0.0084 | 0.0014 | 0.0284 | 0.0093 |
| rs790647 | A | -0.0145 | 0.0017 | -0.0077 | 0.0107 |
| rs7910403 | T | 0.0122 | 0.0018 | -0.0048 | 0.0128 |
| rs7914674 | T | -0.0139 | 0.0022 | 0.0041 | 0.0119 |
| rs7920624 | A | 0.0120 | 0.0014 | -0.0094 | 0.0092 |
| rs7924036 | T | 0.0133 | 0.0014 | 0.0061 | 0.0092 |
| rs79265434 | A | -0.0197 | 0.0022 | -0.0005 | 0.0162 |
| rs7928017 | A | 0.0096 | 0.0014 | -0.0081 | 0.0095 |
| rs7931563 | T | -0.0109 | 0.0014 | 0.0028 | 0.0096 |
| rs795230 | T | 0.0081 | 0.0014 | -0.0045 | 0.0094 |
| rs7958371 | A | -0.0085 | 0.0015 | 0.0145 | 0.0096 |
| rs79585412 | T | -0.0178 | 0.0031 | -0.0166 | 0.0214 |
| rs7965154 | A | -0.0085 | 0.0015 | -0.0005 | 0.0099 |
| rs7967550 | A | -0.0086 | 0.0014 | 0.0068 | 0.0094 |
| rs7972246 | T | 0.0110 | 0.0015 | -0.0179 | 0.0097 |
| rs79728014 | A | -0.0134 | 0.0020 | 0.0073 | 0.0145 |
| rs7974852 | A | 0.0117 | 0.0014 | -0.0130 | 0.0093 |
| rs7977614 | A | -0.0134 | 0.0016 | 0.0310 | 0.0113 |
| rs79798166 | A | 0.0172 | 0.0025 | 0.0039 | 0.0199 |
| rs79855925 | T | -0.0194 | 0.0033 | -0.0026 | 0.0262 |
| rs79994730 | T | 0.0240 | 0.0035 | 0.0198 | 0.0283 |
| rs79997166 | A | 0.0172 | 0.0030 | -0.0110 | 0.0254 |
| rs80037907 | T | -0.0107 | 0.0019 | -0.0010 | 0.0138 |
| rs8008382 | T | -0.0106 | 0.0015 | 0.0000 | 0.0099 |
| rs8009933 | A | 0.0088 | 0.0015 | 0.0092 | 0.0097 |
| rs8016504 | A | -0.0087 | 0.0014 | 0.0051 | 0.0097 |
| rs80223410 | T | -0.0118 | 0.0020 | 0.0172 | 0.0179 |
| rs8024 | A | -0.0115 | 0.0015 | 0.0416 | 0.0101 |
| rs80257979 | T | -0.0252 | 0.0040 | -0.0033 | 0.0285 |
| rs8030487 | A | 0.0085 | 0.0015 | -0.0276 | 0.0104 |
| rs803619 | T | -0.0173 | 0.0023 | 0.0223 | 0.0153 |
| rs8046072 | A | 0.0101 | 0.0018 | -0.0054 | 0.0116 |
| rs8052523 | T | -0.0084 | 0.0014 | -0.0032 | 0.0101 |
| rs8066044 | A | 0.0090 | 0.0016 | -0.0187 | 0.0105 |
| rs806816 | A | 0.0101 | 0.0016 | -0.0034 | 0.0113 |
| rs8097125 | T | 0.0086 | 0.0014 | -0.0015 | 0.0099 |
| rs8103741 | A | -0.0108 | 0.0019 | 0.0075 | 0.0133 |
| rs818415 | T | -0.0110 | 0.0018 | 0.0032 | 0.0112 |
| rs8192465 | A | -0.0272 | 0.0049 | -0.0044 | 0.0428 |
| rs852771 | T | 0.0092 | 0.0015 | 0.0074 | 0.0104 |
| rs853286 | T | 0.0124 | 0.0023 | -0.0049 | 0.0131 |
| rs854796 | A | 0.0090 | 0.0015 | 0.0043 | 0.0100 |
| rs868456 | A | 0.0089 | 0.0015 | 0.0044 | 0.0104 |
| rs870589 | A | -0.0079 | 0.0014 | 0.0156 | 0.0093 |
| rs884108 | A | -0.0094 | 0.0017 | -0.0138 | 0.0117 |
| rs891793 | C | -0.0099 | 0.0014 | -0.0071 | 0.0094 |
| rs893522 | A | 0.0149 | 0.0025 | -0.0183 | 0.0168 |
| rs902820 | A | -0.0096 | 0.0014 | -0.0078 | 0.0096 |
| rs911149 | T | 0.0091 | 0.0017 | -0.0094 | 0.0109 |
| rs912883 | T | 0.0087 | 0.0015 | -0.0037 | 0.0098 |
| rs913509 | A | 0.0096 | 0.0015 | -0.0077 | 0.0098 |
| rs925161 | C | -0.0078 | 0.0014 | -0.0033 | 0.0094 |
| rs9267658 | T | 0.0138 | 0.0020 | -0.0496 | 0.0170 |
| rs9267677 | T | 0.0180 | 0.0024 | 0.0231 | 0.0154 |
| rs9289300 | T | -0.0164 | 0.0019 | 0.0247 | 0.0129 |
| rs9291437 | C | -0.0081 | 0.0014 | 0.0000 | 0.0094 |
| rs9294770 | T | 0.0083 | 0.0014 | -0.0117 | 0.0095 |
| rs929511 | T | -0.0160 | 0.0021 | 0.0046 | 0.0138 |
| rs9300612 | T | 0.0084 | 0.0015 | 0.0066 | 0.0097 |
| rs933738 | A | -0.0110 | 0.0018 | 0.0132 | 0.0122 |
| rs9349956 | A | -0.0159 | 0.0018 | -0.0089 | 0.0126 |
| rs9359939 | A | -0.0105 | 0.0016 | -0.0118 | 0.0105 |
| rs936496 | A | -0.0079 | 0.0014 | 0.0015 | 0.0096 |
| rs9371883 | C | -0.0087 | 0.0015 | 0.0124 | 0.0098 |
| rs9373363 | A | -0.0111 | 0.0016 | -0.0097 | 0.0107 |
| rs9375188 | T | 0.0212 | 0.0014 | -0.0012 | 0.0094 |
| rs9375403 | C | 0.0126 | 0.0016 | -0.0134 | 0.0113 |
| rs9386110 | T | -0.0163 | 0.0023 | 0.0106 | 0.0152 |
| rs9388490 | T | 0.0091 | 0.0014 | -0.0356 | 0.0097 |
| rs939400 | T | -0.0095 | 0.0015 | 0.0047 | 0.0098 |
| rs9411331 | A | 0.0145 | 0.0015 | -0.0148 | 0.0097 |
| rs9435340 | A | 0.0086 | 0.0015 | -0.0111 | 0.0097 |
| rs9442750 | A | -0.0089 | 0.0016 | -0.0017 | 0.0111 |
| rs9446060 | A | 0.0077 | 0.0014 | -0.0004 | 0.0094 |
| rs9465509 | A | -0.0084 | 0.0014 | 0.0109 | 0.0092 |
| rs9490512 | A | -0.0136 | 0.0014 | 0.0008 | 0.0094 |
| rs9492774 | C | 0.0080 | 0.0015 | -0.0240 | 0.0094 |
| rs9513416 | A | -0.0116 | 0.0019 | 0.0026 | 0.0137 |
| rs9513754 | T | 0.0096 | 0.0016 | 0.0062 | 0.0111 |
| rs9527662 | A | 0.0094 | 0.0014 | 0.0083 | 0.0101 |
| rs9527905 | A | -0.0094 | 0.0014 | -0.0121 | 0.0093 |
| rs9529146 | T | 0.0127 | 0.0017 | -0.0037 | 0.0105 |
| rs9536462 | A | 0.0133 | 0.0020 | -0.0102 | 0.0130 |
| rs9540718 | A | -0.0089 | 0.0014 | 0.0191 | 0.0093 |
| rs9545395 | T | 0.0138 | 0.0021 | 0.0037 | 0.0130 |
| rs9556958 | T | -0.0110 | 0.0014 | -0.0053 | 0.0094 |
| rs9563168 | A | 0.0101 | 0.0017 | -0.0100 | 0.0116 |
| rs9568798 | T | -0.0093 | 0.0016 | 0.0157 | 0.0103 |
| rs9597907 | T | 0.0152 | 0.0027 | 0.0065 | 0.0163 |
| rs9611597 | A | -0.0107 | 0.0019 | 0.0147 | 0.0151 |
| rs9616906 | A | 0.0122 | 0.0014 | -0.0136 | 0.0098 |
| rs9616947 | T | -0.0134 | 0.0015 | 0.0233 | 0.0106 |
| rs9633970 | T | 0.0091 | 0.0016 | -0.0037 | 0.0109 |
| rs9649 | T | 0.0108 | 0.0019 | -0.0150 | 0.0122 |
| rs9655780 | A | -0.0153 | 0.0019 | -0.0204 | 0.0139 |
| rs9666728 | T | -0.0082 | 0.0014 | -0.0051 | 0.0094 |
| rs9683585 | C | 0.0084 | 0.0014 | 0.0019 | 0.0092 |
| rs969512 | A | -0.0106 | 0.0015 | 0.0054 | 0.0095 |
| rs9771228 | T | 0.0096 | 0.0015 | -0.0181 | 0.0095 |
| rs977143 | A | 0.0085 | 0.0015 | 0.0024 | 0.0101 |
| rs978807 | A | 0.0139 | 0.0018 | -0.0074 | 0.0116 |
| rs981230 | T | 0.0100 | 0.0014 | 0.0109 | 0.0094 |
| rs981883 | A | 0.0099 | 0.0014 | -0.0162 | 0.0095 |
| rs9820604 | T | 0.0115 | 0.0018 | -0.0055 | 0.0116 |
| rs9821664 | A | 0.0094 | 0.0017 | -0.0100 | 0.0105 |
| rs9830359 | T | -0.0147 | 0.0022 | 0.0091 | 0.0145 |
| rs9844755 | A | 0.0083 | 0.0015 | 0.0075 | 0.0099 |
| rs9849884 | A | -0.0090 | 0.0014 | 0.0076 | 0.0098 |
| rs9853928 | T | -0.0127 | 0.0017 | 0.0220 | 0.0105 |
| rs9858921 | A | -0.0087 | 0.0014 | 0.0031 | 0.0093 |
| rs9859556 | T | 0.0290 | 0.0015 | -0.0431 | 0.0103 |
| rs9866123 | A | 0.0086 | 0.0014 | -0.0128 | 0.0096 |
| rs9870317 | A | 0.0086 | 0.0015 | -0.0290 | 0.0097 |
| rs9877225 | T | 0.0090 | 0.0016 | -0.0080 | 0.0106 |
| rs9882532 | T | 0.0124 | 0.0015 | 0.0021 | 0.0100 |
| rs9886703 | A | -0.0136 | 0.0019 | 0.0333 | 0.0121 |
| rs9888796 | T | 0.0113 | 0.0016 | -0.0066 | 0.0107 |
| rs9916901 | T | 0.0092 | 0.0016 | -0.0218 | 0.0100 |
| rs9924031 | C | 0.0093 | 0.0015 | 0.0066 | 0.0104 |
| rs9926649 | T | 0.0177 | 0.0031 | -0.0121 | 0.0223 |
| rs9927049 | A | 0.0099 | 0.0016 | 0.0127 | 0.0117 |
| rs9927137 | A | 0.0082 | 0.0014 | 0.0003 | 0.0104 |
| rs9927842 | T | -0.0130 | 0.0020 | 0.0031 | 0.0127 |
| rs9929556 | T | -0.0099 | 0.0014 | 0.0123 | 0.0095 |
| rs9929762 | A | 0.0096 | 0.0014 | -0.0007 | 0.0093 |
| rs9929993 | T | -0.0102 | 0.0015 | 0.0011 | 0.0097 |
| rs9933256 | A | 0.0119 | 0.0014 | 0.0077 | 0.0110 |
| rs995698 | A | -0.0099 | 0.0014 | 0.0001 | 0.0096 |
| rs9974899 | T | -0.0093 | 0.0016 | 0.0022 | 0.0107 |
| rs9977825 | T | -0.0092 | 0.0015 | 0.0260 | 0.0104 |

#### Supplementary table 12: Genetic association estimates for the IVW ratio-method MR analysis estimating the effect of education on MI. ea=effect allele; gx=education; gy=MI; se=standard error

| SNP | ea | gx | gx_se | gy | gy_se |
| --- | --- | --- | --- | --- | --- |
| rs10006235 | T | -0.0093 | 0.0016 | -0.0077 | 0.0112 |
| rs10009513 | A | 0.0088 | 0.0014 | -0.0174 | 0.0103 |
| rs10021733 | T | -0.0164 | 0.0027 | -0.0023 | 0.0154 |
| rs10032941 | T | -0.0080 | 0.0015 | 0.0127 | 0.0106 |
| rs10041403 | T | 0.0151 | 0.0019 | -0.0142 | 0.0142 |
| rs10042828 | A | -0.0145 | 0.0022 | 0.0007 | 0.0172 |
| rs10046069 | A | 0.0127 | 0.0022 | 0.0075 | 0.0154 |
| rs10057590 | A | 0.0106 | 0.0014 | -0.0049 | 0.0106 |
| rs10060023 | T | 0.0110 | 0.0015 | 0.0019 | 0.0116 |
| rs10061420 | A | 0.0082 | 0.0014 | -0.0027 | 0.0104 |
| rs1006749 | A | 0.0079 | 0.0014 | -0.0013 | 0.0108 |
| rs10074178 | A | 0.0079 | 0.0014 | -0.0025 | 0.0104 |
| rs1007731 | A | -0.0143 | 0.0022 | 0.0031 | 0.0192 |
| rs10080647 | A | 0.0124 | 0.0020 | -0.0220 | 0.0141 |
| rs1008078 | T | -0.0182 | 0.0014 | 0.0058 | 0.0106 |
| rs1009470 | T | 0.0086 | 0.0014 | -0.0198 | 0.0103 |
| rs10098073 | A | -0.0091 | 0.0014 | -0.0033 | 0.0109 |
| rs10122669 | A | -0.0079 | 0.0014 | 0.0039 | 0.0104 |
| rs10128888 | A | -0.0083 | 0.0015 | 0.0334 | 0.0119 |
| rs10145520 | T | -0.0122 | 0.0018 | -0.0182 | 0.0133 |
| rs10166286 | T | 0.0151 | 0.0019 | -0.0035 | 0.0125 |
| rs10169002 | A | -0.0080 | 0.0014 | 0.0044 | 0.0104 |
| rs10181071 | A | 0.0102 | 0.0019 | -0.0008 | 0.0144 |
| rs10189857 | A | 0.0158 | 0.0014 | -0.0086 | 0.0105 |
| rs10192369 | A | -0.0078 | 0.0014 | -0.0010 | 0.0106 |
| rs10192834 | T | 0.0083 | 0.0015 | -0.0087 | 0.0111 |
| rs10193498 | A | 0.0097 | 0.0016 | 0.0107 | 0.0116 |
| rs10204051 | T | -0.0101 | 0.0014 | 0.0000 | 0.0111 |
| rs10205057 | T | 0.0090 | 0.0014 | -0.0191 | 0.0102 |
| rs10208 | T | 0.0091 | 0.0015 | -0.0124 | 0.0111 |
| rs10215082 | A | -0.0149 | 0.0014 | 0.0030 | 0.0109 |
| rs1024268 | T | -0.0084 | 0.0014 | 0.0025 | 0.0103 |
| rs10251438 | A | 0.0078 | 0.0014 | -0.0195 | 0.0103 |
| rs10264573 | A | 0.0175 | 0.0032 | 0.0262 | 0.0238 |
| rs1035578 | A | -0.0094 | 0.0014 | -0.0111 | 0.0106 |
| rs10402747 | T | 0.0077 | 0.0014 | -0.0253 | 0.0108 |
| rs10411759 | A | 0.0116 | 0.0020 | -0.0069 | 0.0153 |
| rs10444280 | T | 0.0100 | 0.0018 | 0.0044 | 0.0134 |
| rs10460095 | A | -0.0124 | 0.0014 | 0.0057 | 0.0105 |
| rs10480450 | A | 0.0158 | 0.0028 | 0.0190 | 0.0197 |
| rs10496091 | A | -0.0131 | 0.0016 | 0.0213 | 0.0124 |
| rs10499535 | A | 0.0080 | 0.0014 | 0.0059 | 0.0104 |
| rs1050847 | T | 0.0092 | 0.0014 | -0.0304 | 0.0116 |
| rs10509251 | T | -0.0093 | 0.0017 | 0.0227 | 0.0116 |
| rs10515007 | T | 0.0143 | 0.0019 | -0.0287 | 0.0134 |
| rs1051860 | A | -0.0079 | 0.0014 | 0.0240 | 0.0108 |
| rs10519504 | T | 0.0119 | 0.0019 | -0.0104 | 0.0141 |
| rs1054442 | A | -0.0136 | 0.0015 | -0.0026 | 0.0109 |
| rs1061801 | A | -0.0110 | 0.0018 | 0.0112 | 0.0137 |
| rs1066769 | A | -0.0230 | 0.0041 | -0.0222 | 0.0319 |
| rs10742591 | A | -0.0084 | 0.0014 | 0.0022 | 0.0105 |
| rs10748841 | A | -0.0111 | 0.0020 | 0.0084 | 0.0110 |
| rs10750539 | A | -0.0095 | 0.0015 | 0.0025 | 0.0109 |
| rs10752262 | T | 0.0102 | 0.0014 | -0.0032 | 0.0108 |
| rs1075228 | A | -0.0081 | 0.0014 | -0.0029 | 0.0112 |
| rs10761202 | T | 0.0088 | 0.0014 | 0.0211 | 0.0104 |
| rs10761251 | A | 0.0113 | 0.0015 | 0.0111 | 0.0107 |
| rs10772644 | C | 0.0142 | 0.0022 | -0.0278 | 0.0157 |
| rs10773002 | A | 0.0185 | 0.0016 | -0.0135 | 0.0117 |
| rs10773208 | T | -0.0110 | 0.0016 | -0.0124 | 0.0121 |
| rs10782651 | T | -0.0082 | 0.0014 | 0.0037 | 0.0104 |
| rs10783243 | A | -0.0092 | 0.0014 | 0.0076 | 0.0105 |
| rs10789285 | T | 0.0091 | 0.0016 | -0.0160 | 0.0116 |
| rs10793903 | A | 0.0126 | 0.0022 | -0.0248 | 0.0159 |
| rs10795831 | T | -0.0099 | 0.0017 | 0.0152 | 0.0118 |
| rs10798888 | T | -0.0140 | 0.0019 | 0.0197 | 0.0129 |
| rs10799615 | A | -0.0089 | 0.0016 | 0.0225 | 0.0112 |
| rs10805383 | A | -0.0103 | 0.0014 | 0.0088 | 0.0103 |
| rs10810099 | A | -0.0146 | 0.0016 | 0.0015 | 0.0112 |
| rs10810145 | A | 0.0078 | 0.0014 | -0.0123 | 0.0104 |
| rs10830858 | T | 0.0092 | 0.0014 | -0.0100 | 0.0103 |
| rs10831656 | T | 0.0090 | 0.0015 | -0.0006 | 0.0114 |
| rs10844179 | A | 0.0104 | 0.0017 | -0.0208 | 0.0124 |
| rs10845051 | A | 0.0083 | 0.0014 | 0.0010 | 0.0107 |
| rs10875121 | C | 0.0174 | 0.0019 | -0.0192 | 0.0142 |
| rs10877283 | T | 0.0085 | 0.0014 | 0.0014 | 0.0109 |
| rs10879676 | T | -0.0083 | 0.0014 | 0.0064 | 0.0106 |
| rs10886012 | C | 0.0078 | 0.0014 | -0.0027 | 0.0103 |
| rs10892807 | T | 0.0109 | 0.0014 | 0.0098 | 0.0110 |
| rs10899282 | A | 0.0140 | 0.0017 | -0.0165 | 0.0128 |
| rs10927053 | A | -0.0125 | 0.0022 | 0.0159 | 0.0152 |
| rs10928190 | T | -0.0087 | 0.0014 | 0.0058 | 0.0104 |
| rs10931821 | A | -0.0141 | 0.0014 | 0.0109 | 0.0103 |
| rs10937240 | T | -0.0123 | 0.0018 | -0.0028 | 0.0122 |
| rs10940540 | A | 0.0078 | 0.0014 | 0.0149 | 0.0106 |
| rs10940921 | T | 0.0111 | 0.0014 | 0.0105 | 0.0102 |
| rs10943588 | A | 0.0084 | 0.0015 | -0.0196 | 0.0106 |
| rs10947439 | T | 0.0090 | 0.0015 | -0.0173 | 0.0111 |
| rs10949263 | A | -0.0091 | 0.0016 | -0.0131 | 0.0115 |
| rs10951590 | T | -0.0109 | 0.0015 | 0.0069 | 0.0110 |
| rs10974256 | A | -0.0093 | 0.0015 | -0.0030 | 0.0119 |
| rs1097784 | T | -0.0083 | 0.0014 | -0.0012 | 0.0105 |
| rs10979613 | T | -0.0117 | 0.0015 | -0.0031 | 0.0108 |
| rs10983324 | A | 0.0084 | 0.0015 | -0.0259 | 0.0112 |
| rs10984445 | A | 0.0115 | 0.0014 | -0.0217 | 0.0103 |
| rs10985402 | T | 0.0089 | 0.0016 | -0.0058 | 0.0115 |
| rs10995639 | T | 0.0078 | 0.0014 | -0.0117 | 0.0103 |
| rs10996167 | C | 0.0113 | 0.0020 | 0.0085 | 0.0108 |
| rs11003463 | T | 0.0091 | 0.0015 | -0.0086 | 0.0123 |
| rs11021432 | A | 0.0103 | 0.0015 | 0.0086 | 0.0113 |
| rs11023764 | A | 0.0091 | 0.0015 | 0.0148 | 0.0112 |
| rs11030084 | T | 0.0106 | 0.0018 | -0.0276 | 0.0128 |
| rs11030102 | C | 0.0101 | 0.0016 | -0.0239 | 0.0131 |
| rs1106090 | A | 0.0094 | 0.0014 | -0.0005 | 0.0103 |
| rs11076962 | T | 0.0086 | 0.0016 | 0.0006 | 0.0113 |
| rs1107871 | A | -0.0083 | 0.0014 | 0.0158 | 0.0103 |
| rs11081529 | T | 0.0126 | 0.0015 | -0.0199 | 0.0116 |
| rs11082011 | T | 0.0194 | 0.0015 | 0.0141 | 0.0107 |
| rs11100237 | A | -0.0101 | 0.0014 | 0.0028 | 0.0102 |
| rs11115056 | A | 0.0187 | 0.0023 | -0.0012 | 0.0203 |
| rs11121177 | A | 0.0151 | 0.0018 | -0.0106 | 0.0129 |
| rs111226181 | T | -0.0201 | 0.0018 | 0.0140 | 0.0147 |
| rs111235962 | T | 0.0133 | 0.0024 | -0.0034 | 0.0182 |
| rs11130380 | T | 0.0090 | 0.0014 | 0.0181 | 0.0109 |
| rs111370527 | T | 0.0151 | 0.0027 | 0.0077 | 0.0275 |
| rs11138947 | T | 0.0105 | 0.0016 | -0.0148 | 0.0110 |
| rs111530150 | T | 0.0269 | 0.0048 | 0.0620 | 0.0354 |
| rs11155813 | T | -0.0129 | 0.0023 | -0.0401 | 0.0151 |
| rs11157931 | A | -0.0139 | 0.0014 | -0.0191 | 0.0109 |
| rs11158800 | A | -0.0091 | 0.0014 | 0.0117 | 0.0108 |
| rs1117152 | T | 0.0106 | 0.0015 | -0.0225 | 0.0111 |
| rs11174399 | A | 0.0111 | 0.0018 | -0.0123 | 0.0129 |
| rs111821073 | T | 0.0124 | 0.0019 | -0.0162 | 0.0152 |
| rs111852224 | T | 0.0156 | 0.0022 | 0.0197 | 0.0194 |
| rs11190955 | T | -0.0083 | 0.0015 | 0.0044 | 0.0111 |
| rs11191237 | T | 0.0132 | 0.0024 | -0.0154 | 0.0123 |
| rs1120924 | T | 0.0108 | 0.0015 | -0.0029 | 0.0113 |
| rs11209894 | A | 0.0119 | 0.0021 | -0.0125 | 0.0144 |
| rs11210228 | T | -0.0105 | 0.0019 | 0.0070 | 0.0108 |
| rs11210400 | A | 0.0120 | 0.0014 | 0.0299 | 0.0106 |
| rs11211123 | A | 0.0104 | 0.0017 | 0.0081 | 0.0121 |
| rs11213482 | A | -0.0106 | 0.0019 | -0.0006 | 0.0161 |
| rs112210983 | A | 0.0105 | 0.0017 | -0.0005 | 0.0131 |
| rs1123285 | C | -0.0094 | 0.0015 | -0.0100 | 0.0112 |
| rs112375785 | A | 0.0184 | 0.0021 | -0.0098 | 0.0194 |
| rs112379405 | C | -0.0082 | 0.0014 | -0.0143 | 0.0105 |
| rs112512729 | T | -0.0166 | 0.0027 | -0.0019 | 0.0230 |
| rs112603734 | A | -0.0131 | 0.0022 | -0.0021 | 0.0128 |
| rs11265191 | T | 0.0083 | 0.0015 | 0.0078 | 0.0107 |
| rs112806496 | C | -0.0160 | 0.0025 | 0.0160 | 0.0205 |
| rs1128956 | T | -0.0140 | 0.0019 | 0.0265 | 0.0149 |
| rs113205706 | A | -0.0135 | 0.0024 | -0.0305 | 0.0203 |
| rs113520408 | A | 0.0155 | 0.0016 | -0.0102 | 0.0132 |
| rs113552169 | C | 0.0294 | 0.0053 | -0.0360 | 0.0415 |
| rs113588399 | T | -0.0104 | 0.0017 | 0.0019 | 0.0137 |
| rs113615161 | T | -0.0123 | 0.0021 | 0.0300 | 0.0149 |
| rs113720505 | A | 0.0164 | 0.0029 | -0.0029 | 0.0292 |
| rs113731629 | T | 0.0339 | 0.0061 | -0.0889 | 0.0544 |
| rs114192810 | T | -0.0279 | 0.0051 | 0.0809 | 0.0387 |
| rs114468556 | A | 0.0293 | 0.0036 | -0.0472 | 0.0344 |
| rs114593137 | A | -0.0122 | 0.0017 | 0.0131 | 0.0133 |
| rs1146079 | C | -0.0087 | 0.0016 | 0.0129 | 0.0121 |
| rs114810763 | A | -0.0121 | 0.0020 | 0.0027 | 0.0175 |
| rs114952970 | T | 0.0265 | 0.0040 | 0.0274 | 0.0336 |
| rs114958262 | T | -0.0220 | 0.0039 | 0.0180 | 0.0375 |
| rs115000530 | A | -0.0250 | 0.0031 | 0.0557 | 0.0259 |
| rs115017135 | A | -0.0206 | 0.0038 | 0.0112 | 0.0348 |
| rs11542663 | A | 0.0112 | 0.0015 | 0.0000 | 0.0109 |
| rs115438240 | T | -0.0169 | 0.0030 | 0.0126 | 0.0261 |
| rs115693355 | A | 0.0309 | 0.0053 | 0.0185 | 0.0484 |
| rs115732722 | A | -0.0415 | 0.0072 | -0.0209 | 0.0590 |
| rs11588857 | A | 0.0199 | 0.0017 | 0.0202 | 0.0122 |
| rs11591870 | T | 0.0098 | 0.0017 | -0.0161 | 0.0140 |
| rs11592299 | A | 0.0097 | 0.0018 | 0.0180 | 0.0141 |
| rs11596387 | T | 0.0258 | 0.0046 | 0.0192 | 0.0456 |
| rs11598765 | A | -0.0106 | 0.0018 | 0.0029 | 0.0123 |
| rs11599236 | T | -0.0106 | 0.0014 | 0.0171 | 0.0105 |
| rs11613431 | A | -0.0094 | 0.0016 | -0.0062 | 0.0126 |
| rs11620355 | A | 0.0182 | 0.0025 | -0.0312 | 0.0193 |
| rs11623285 | T | -0.0135 | 0.0021 | 0.0294 | 0.0162 |
| rs11640569 | A | 0.0112 | 0.0020 | -0.0063 | 0.0160 |
| rs11644446 | A | 0.0117 | 0.0019 | 0.0067 | 0.0163 |
| rs11647188 | A | 0.0079 | 0.0014 | -0.0036 | 0.0110 |
| rs11652522 | A | -0.0154 | 0.0023 | 0.0086 | 0.0173 |
| rs11657342 | A | 0.0140 | 0.0019 | 0.0043 | 0.0147 |
| rs11657979 | A | -0.0091 | 0.0017 | 0.0301 | 0.0126 |
| rs11663602 | A | -0.0130 | 0.0016 | 0.0020 | 0.0112 |
| rs11663678 | T | -0.0121 | 0.0022 | -0.0054 | 0.0166 |
| rs116656374 | T | -0.0242 | 0.0038 | -0.0131 | 0.0332 |
| rs11675476 | T | -0.0127 | 0.0014 | 0.0062 | 0.0102 |
| rs1167827 | A | 0.0104 | 0.0014 | -0.0050 | 0.0110 |
| rs11678980 | A | -0.0166 | 0.0014 | 0.0175 | 0.0114 |
| rs11687736 | A | -0.0105 | 0.0015 | 0.0067 | 0.0110 |
| rs11693885 | A | -0.0091 | 0.0014 | 0.0154 | 0.0104 |
| rs116967397 | A | 0.0315 | 0.0056 | -0.0204 | 0.0474 |
| rs117005905 | T | 0.0150 | 0.0022 | -0.0034 | 0.0156 |
| rs11703948 | A | -0.0168 | 0.0024 | 0.0054 | 0.0174 |
| rs1171040 | A | -0.0113 | 0.0017 | 0.0303 | 0.0126 |
| rs11716398 | A | 0.0159 | 0.0021 | 0.0042 | 0.0149 |
| rs11720093 | T | 0.0150 | 0.0026 | 0.0065 | 0.0197 |
| rs11720985 | T | 0.0085 | 0.0015 | -0.0094 | 0.0117 |
| rs11724690 | T | 0.0093 | 0.0015 | 0.0079 | 0.0119 |
| rs117273411 | T | -0.0211 | 0.0034 | 0.0376 | 0.0327 |
| rs11732160 | A | 0.0105 | 0.0016 | 0.0229 | 0.0117 |
| rs11732657 | A | -0.0126 | 0.0016 | 0.0039 | 0.0117 |
| rs11733439 | A | -0.0096 | 0.0017 | 0.0089 | 0.0119 |
| rs11736863 | A | 0.0142 | 0.0018 | -0.0020 | 0.0135 |
| rs117398064 | C | -0.0141 | 0.0025 | 0.0003 | 0.0196 |
| rs117468730 | A | -0.0310 | 0.0050 | 0.0376 | 0.0348 |
| rs117623407 | A | -0.0118 | 0.0020 | -0.0137 | 0.0161 |
| rs11765387 | T | 0.0086 | 0.0015 | 0.0065 | 0.0106 |
| rs117668569 | T | 0.0234 | 0.0042 | -0.0060 | 0.0321 |
| rs117677995 | A | -0.0180 | 0.0032 | 0.0296 | 0.0329 |
| rs11772232 | T | 0.0177 | 0.0019 | -0.0407 | 0.0145 |
| rs11774212 | T | 0.0128 | 0.0014 | -0.0003 | 0.0110 |
| rs117799466 | C | 0.0106 | 0.0016 | 0.0017 | 0.0136 |
| rs11780023 | T | 0.0085 | 0.0014 | -0.0047 | 0.0104 |
| rs11789013 | T | 0.0106 | 0.0016 | -0.0185 | 0.0118 |
| rs117895796 | T | 0.0362 | 0.0064 | -0.0286 | 0.0601 |
| rs118040169 | A | -0.0248 | 0.0038 | 0.0163 | 0.0346 |
| rs118093058 | T | 0.0146 | 0.0021 | -0.0028 | 0.0181 |
| rs118134876 | T | -0.0223 | 0.0030 | 0.0424 | 0.0266 |
| rs11845781 | T | -0.0079 | 0.0014 | -0.0032 | 0.0105 |
| rs11871429 | A | 0.0128 | 0.0017 | 0.0135 | 0.0119 |
| rs11876620 | T | 0.0143 | 0.0024 | -0.0077 | 0.0180 |
| rs11894424 | A | 0.0209 | 0.0033 | -0.0015 | 0.0188 |
| rs11917701 | A | -0.0087 | 0.0014 | 0.0102 | 0.0105 |
| rs11919835 | C | 0.0089 | 0.0015 | -0.0233 | 0.0110 |
| rs1196760 | C | 0.0139 | 0.0025 | 0.0274 | 0.0212 |
| rs12005151 | A | 0.0087 | 0.0015 | 0.0067 | 0.0114 |
| rs12028010 | T | 0.0164 | 0.0017 | -0.0139 | 0.0127 |
| rs12028229 | T | 0.0099 | 0.0016 | 0.0195 | 0.0135 |
| rs12054166 | C | 0.0096 | 0.0016 | -0.0156 | 0.0121 |
| rs12076635 | C | 0.0207 | 0.0017 | -0.0195 | 0.0117 |
| rs12113634 | T | 0.0088 | 0.0015 | -0.0149 | 0.0106 |
| rs12123293 | T | 0.0086 | 0.0015 | 0.0029 | 0.0110 |
| rs12126231 | A | 0.0095 | 0.0014 | -0.0109 | 0.0104 |
| rs12127928 | T | 0.0108 | 0.0018 | -0.0115 | 0.0140 |
| rs12145078 | A | -0.0127 | 0.0017 | 0.0143 | 0.0124 |
| rs12151248 | T | -0.0149 | 0.0023 | -0.0240 | 0.0207 |
| rs12170452 | A | 0.0105 | 0.0014 | -0.0064 | 0.0106 |
| rs1220779 | A | -0.0079 | 0.0014 | 0.0139 | 0.0105 |
| rs12234369 | A | 0.0087 | 0.0015 | 0.0178 | 0.0112 |
| rs12238011 | T | -0.0141 | 0.0026 | 0.0124 | 0.0174 |
| rs12273435 | A | -0.0132 | 0.0017 | 0.0193 | 0.0133 |
| rs12285074 | A | 0.0161 | 0.0023 | 0.0135 | 0.0142 |
| rs12304188 | A | 0.0111 | 0.0019 | -0.0063 | 0.0132 |
| rs12325727 | A | 0.0088 | 0.0014 | -0.0268 | 0.0107 |
| rs12332731 | A | 0.0135 | 0.0018 | -0.0186 | 0.0143 |
| rs12342546 | A | -0.0128 | 0.0017 | -0.0038 | 0.0123 |
| rs12359372 | T | -0.0092 | 0.0015 | 0.0218 | 0.0111 |
| rs12375949 | T | -0.0138 | 0.0014 | 0.0056 | 0.0105 |
| rs12405889 | T | -0.0098 | 0.0014 | 0.0013 | 0.0102 |
| rs12431682 | T | -0.0093 | 0.0015 | -0.0004 | 0.0108 |
| rs12438177 | A | 0.0093 | 0.0015 | -0.0208 | 0.0105 |
| rs12453682 | T | 0.0102 | 0.0015 | -0.0184 | 0.0114 |
| rs1245829 | A | -0.0099 | 0.0014 | 0.0269 | 0.0112 |
| rs12467175 | T | -0.0081 | 0.0014 | 0.0281 | 0.0104 |
| rs12468040 | T | 0.0137 | 0.0014 | -0.0035 | 0.0104 |
| rs12473986 | T | -0.0085 | 0.0015 | 0.0070 | 0.0110 |
| rs12477385 | T | 0.0109 | 0.0017 | -0.0158 | 0.0128 |
| rs12478156 | T | 0.0108 | 0.0015 | 0.0074 | 0.0107 |
| rs12503522 | T | -0.0113 | 0.0016 | 0.0175 | 0.0114 |
| rs12506221 | T | -0.0130 | 0.0014 | 0.0156 | 0.0103 |
| rs12516990 | C | -0.0099 | 0.0017 | 0.0031 | 0.0135 |
| rs12519073 | T | -0.0107 | 0.0017 | -0.0101 | 0.0123 |
| rs12524795 | T | 0.0102 | 0.0014 | -0.0070 | 0.0107 |
| rs12568153 | T | -0.0091 | 0.0015 | -0.0028 | 0.0115 |
| rs12571549 | A | 0.0151 | 0.0020 | -0.0051 | 0.0131 |
| rs12574281 | A | -0.0080 | 0.0015 | -0.0123 | 0.0113 |
| rs12591647 | T | -0.0152 | 0.0018 | 0.0070 | 0.0125 |
| rs12601380 | A | -0.0082 | 0.0014 | 0.0011 | 0.0108 |
| rs12602286 | T | 0.0160 | 0.0021 | -0.0415 | 0.0133 |
| rs12613500 | C | 0.0098 | 0.0014 | -0.0169 | 0.0104 |
| rs12614263 | A | 0.0083 | 0.0014 | -0.0190 | 0.0105 |
| rs12620796 | A | 0.0121 | 0.0019 | -0.0237 | 0.0111 |
| rs12638072 | A | -0.0107 | 0.0015 | -0.0008 | 0.0116 |
| rs12643771 | T | 0.0130 | 0.0015 | -0.0137 | 0.0117 |
| rs12646216 | T | 0.0084 | 0.0014 | -0.0087 | 0.0108 |
| rs12646523 | T | -0.0132 | 0.0016 | 0.0035 | 0.0116 |
| rs12659776 | T | 0.0085 | 0.0014 | -0.0233 | 0.0102 |
| rs12670376 | A | 0.0092 | 0.0014 | -0.0154 | 0.0102 |
| rs1267062 | C | 0.0113 | 0.0018 | 0.0105 | 0.0137 |
| rs12682775 | T | -0.0119 | 0.0017 | 0.0024 | 0.0119 |
| rs12694681 | T | 0.0102 | 0.0015 | -0.0172 | 0.0115 |
| rs12709186 | A | -0.0091 | 0.0015 | 0.0111 | 0.0114 |
| rs12712269 | T | -0.0107 | 0.0014 | 0.0057 | 0.0104 |
| rs12746551 | C | 0.0248 | 0.0041 | -0.0033 | 0.0329 |
| rs12750688 | C | 0.0116 | 0.0016 | 0.0293 | 0.0129 |
| rs12761729 | T | -0.0098 | 0.0016 | -0.0106 | 0.0129 |
| rs12761761 | T | 0.0153 | 0.0017 | 0.0017 | 0.0131 |
| rs12764593 | C | 0.0202 | 0.0029 | 0.0057 | 0.0243 |
| rs12765185 | A | -0.0088 | 0.0016 | 0.0184 | 0.0125 |
| rs12774577 | T | 0.0142 | 0.0020 | -0.0218 | 0.0158 |
| rs12789313 | T | 0.0092 | 0.0014 | -0.0076 | 0.0104 |
| rs12790196 | T | -0.0101 | 0.0015 | -0.0005 | 0.0113 |
| rs12810587 | T | 0.0094 | 0.0015 | -0.0072 | 0.0112 |
| rs12875339 | A | -0.0108 | 0.0015 | -0.0050 | 0.0104 |
| rs12888615 | T | 0.0120 | 0.0018 | -0.0059 | 0.0134 |
| rs12891042 | T | 0.0092 | 0.0014 | -0.0054 | 0.0105 |
| rs12891191 | C | 0.0080 | 0.0014 | -0.0127 | 0.0112 |
| rs12912465 | T | -0.0104 | 0.0016 | 0.0385 | 0.0136 |
| rs12952191 | T | -0.0081 | 0.0014 | 0.0020 | 0.0104 |
| rs12953422 | A | -0.0080 | 0.0014 | 0.0055 | 0.0107 |
| rs12957463 | A | 0.0146 | 0.0018 | -0.0061 | 0.0127 |
| rs12962845 | C | -0.0123 | 0.0021 | 0.0067 | 0.0150 |
| rs12967010 | T | 0.0114 | 0.0017 | -0.0157 | 0.0136 |
| rs12970264 | A | -0.0085 | 0.0014 | 0.0049 | 0.0108 |
| rs12981405 | T | -0.0112 | 0.0019 | 0.0072 | 0.0134 |
| rs13009915 | T | 0.0212 | 0.0037 | -0.0288 | 0.0370 |
| rs13010288 | T | 0.0198 | 0.0021 | -0.0164 | 0.0167 |
| rs13010566 | A | -0.0091 | 0.0014 | 0.0020 | 0.0104 |
| rs13015496 | C | 0.0119 | 0.0018 | -0.0163 | 0.0124 |
| rs13016316 | A | -0.0138 | 0.0024 | -0.0211 | 0.0155 |
| rs1301838 | T | -0.0096 | 0.0015 | 0.0291 | 0.0115 |
| rs13018640 | T | -0.0215 | 0.0014 | 0.0226 | 0.0108 |
| rs13026611 | A | -0.0084 | 0.0015 | -0.0224 | 0.0114 |
| rs13029602 | T | -0.0092 | 0.0014 | -0.0008 | 0.0104 |
| rs13034349 | T | -0.0082 | 0.0014 | 0.0044 | 0.0113 |
| rs13050131 | A | -0.0081 | 0.0015 | -0.0022 | 0.0109 |
| rs13085461 | C | 0.0103 | 0.0014 | -0.0080 | 0.0104 |
| rs13099165 | T | -0.0118 | 0.0017 | -0.0165 | 0.0134 |
| rs13107325 | T | -0.0239 | 0.0027 | -0.0015 | 0.0244 |
| rs13117856 | A | 0.0084 | 0.0015 | -0.0144 | 0.0116 |
| rs13130765 | C | -0.0091 | 0.0014 | 0.0212 | 0.0116 |
| rs13133213 | A | 0.0096 | 0.0014 | -0.0301 | 0.0102 |
| rs13145650 | T | -0.0173 | 0.0025 | 0.0092 | 0.0190 |
| rs13154429 | C | -0.0136 | 0.0023 | 0.0045 | 0.0167 |
| rs13163062 | T | 0.0115 | 0.0014 | -0.0130 | 0.0103 |
| rs13163845 | T | -0.0149 | 0.0020 | -0.0024 | 0.0151 |
| rs13168136 | A | -0.0092 | 0.0017 | -0.0032 | 0.0116 |
| rs13169187 | A | -0.0079 | 0.0014 | -0.0055 | 0.0104 |
| rs13177031 | A | -0.0085 | 0.0015 | -0.0114 | 0.0108 |
| rs13190235 | T | -0.0138 | 0.0024 | 0.0090 | 0.0163 |
| rs13197257 | T | 0.0109 | 0.0016 | -0.0063 | 0.0118 |
| rs1320139 | C | -0.0149 | 0.0014 | 0.0234 | 0.0103 |
| rs13212041 | T | 0.0101 | 0.0018 | -0.0065 | 0.0136 |
| rs13240401 | T | 0.0176 | 0.0017 | -0.0032 | 0.0124 |
| rs13246220 | T | -0.0093 | 0.0014 | 0.0056 | 0.0107 |
| rs13261773 | C | 0.0122 | 0.0019 | 0.0127 | 0.0131 |
| rs13266287 | A | -0.0085 | 0.0015 | -0.0008 | 0.0111 |
| rs13281564 | A | 0.0088 | 0.0014 | -0.0004 | 0.0104 |
| rs13284516 | C | -0.0082 | 0.0014 | -0.0024 | 0.0110 |
| rs1329044 | T | 0.0134 | 0.0019 | -0.0283 | 0.0135 |
| rs1329125 | T | -0.0097 | 0.0015 | 0.0127 | 0.0121 |
| rs13296345 | T | 0.0090 | 0.0015 | -0.0192 | 0.0105 |
| rs13318986 | A | 0.0124 | 0.0015 | -0.0101 | 0.0110 |
| rs13327482 | A | -0.0121 | 0.0018 | -0.0073 | 0.0145 |
| rs1334297 | A | 0.0257 | 0.0016 | -0.0058 | 0.0124 |
| rs1335482 | T | 0.0096 | 0.0014 | -0.0094 | 0.0107 |
| rs13361043 | T | -0.0158 | 0.0028 | 0.0180 | 0.0186 |
| rs13381557 | A | -0.0087 | 0.0014 | -0.0035 | 0.0102 |
| rs13388333 | A | 0.0085 | 0.0015 | -0.0203 | 0.0113 |
| rs13397529 | C | -0.0097 | 0.0017 | 0.0123 | 0.0124 |
| rs13398860 | A | 0.0086 | 0.0015 | -0.0095 | 0.0111 |
| rs13402497 | A | -0.0080 | 0.0014 | -0.0094 | 0.0103 |
| rs13422673 | T | -0.0110 | 0.0014 | 0.0070 | 0.0107 |
| rs13425585 | C | -0.0089 | 0.0014 | 0.0178 | 0.0104 |
| rs1368250 | T | 0.0250 | 0.0045 | 0.0132 | 0.0437 |
| rs137079 | T | 0.0128 | 0.0020 | 0.0144 | 0.0154 |
| rs137858393 | A | 0.0193 | 0.0029 | 0.0456 | 0.0239 |
| rs138096147 | A | 0.0234 | 0.0042 | 0.0607 | 0.0408 |
| rs138484388 | T | -0.0306 | 0.0051 | 0.0580 | 0.0462 |
| rs1391513 | T | -0.0095 | 0.0015 | 0.0048 | 0.0111 |
| rs139244147 | A | -0.0193 | 0.0030 | -0.0136 | 0.0208 |
| rs1392816 | T | 0.0082 | 0.0014 | -0.0158 | 0.0107 |
| rs139980871 | T | 0.0180 | 0.0029 | -0.0098 | 0.0228 |
| rs1404549 | A | -0.0089 | 0.0015 | 0.0086 | 0.0112 |
| rs1405876 | T | 0.0109 | 0.0015 | -0.0018 | 0.0113 |
| rs140711597 | C | 0.0366 | 0.0055 | 0.0637 | 0.0466 |
| rs1408284 | C | -0.0127 | 0.0020 | 0.0193 | 0.0159 |
| rs1408430 | C | 0.0096 | 0.0016 | 0.0108 | 0.0115 |
| rs141586924 | A | -0.0090 | 0.0015 | 0.0024 | 0.0120 |
| rs141729694 | T | 0.0195 | 0.0026 | 0.0461 | 0.0189 |
| rs142014757 | A | 0.0102 | 0.0018 | 0.0024 | 0.0150 |
| rs1426619 | T | 0.0096 | 0.0014 | 0.0123 | 0.0104 |
| rs1427298 | T | 0.0086 | 0.0014 | -0.0138 | 0.0106 |
| rs142747148 | A | 0.0329 | 0.0052 | -0.0431 | 0.0486 |
| rs1427829 | A | -0.0083 | 0.0014 | 0.0147 | 0.0102 |
| rs143148393 | T | -0.0211 | 0.0039 | -0.0054 | 0.0351 |
| rs1434630 | T | -0.0137 | 0.0020 | -0.0133 | 0.0137 |
| rs143743568 | A | 0.0118 | 0.0020 | -0.0085 | 0.0151 |
| rs143812851 | A | -0.0113 | 0.0019 | 0.0070 | 0.0141 |
| rs1440930 | C | 0.0081 | 0.0014 | 0.0120 | 0.0104 |
| rs144336753 | A | 0.0388 | 0.0053 | -0.0411 | 0.0425 |
| rs1445591 | A | 0.0085 | 0.0015 | 0.0045 | 0.0123 |
| rs145018899 | A | 0.0095 | 0.0015 | 0.0085 | 0.0117 |
| rs1464297 | T | -0.0109 | 0.0015 | -0.0156 | 0.0106 |
| rs1467737 | T | -0.0084 | 0.0014 | 0.0146 | 0.0107 |
| rs1485300 | C | 0.0098 | 0.0015 | -0.0100 | 0.0111 |
| rs1490612 | T | 0.0193 | 0.0026 | 0.0000 | 0.0190 |
| rs150252215 | A | 0.0204 | 0.0035 | -0.0102 | 0.0289 |
| rs150537577 | A | -0.0151 | 0.0026 | 0.0165 | 0.0203 |
| rs1505676 | C | 0.0084 | 0.0015 | -0.0204 | 0.0104 |
| rs151381 | T | -0.0087 | 0.0014 | 0.0285 | 0.0105 |
| rs1518890 | T | -0.0090 | 0.0016 | 0.0015 | 0.0117 |
| rs1527878 | A | -0.0105 | 0.0016 | 0.0203 | 0.0126 |
| rs1529597 | A | 0.0223 | 0.0038 | -0.0332 | 0.0281 |
| rs1538389 | T | -0.0109 | 0.0018 | 0.0152 | 0.0133 |
| rs1542354 | A | -0.0084 | 0.0014 | -0.0091 | 0.0106 |
| rs1544 | A | -0.0096 | 0.0016 | 0.0091 | 0.0115 |
| rs1554798 | A | -0.0080 | 0.0014 | 0.0173 | 0.0106 |
| rs1564347 | T | 0.0088 | 0.0015 | -0.0250 | 0.0108 |
| rs1569092 | A | 0.0160 | 0.0020 | -0.0086 | 0.0149 |
| rs1593022 | T | -0.0099 | 0.0017 | 0.0084 | 0.0126 |
| rs1603460 | T | 0.0090 | 0.0014 | -0.0024 | 0.0105 |
| rs162445 | A | 0.0152 | 0.0026 | 0.0111 | 0.0170 |
| rs163229 | C | -0.0260 | 0.0047 | 0.0044 | 0.0344 |
| rs1637770 | T | 0.0191 | 0.0028 | -0.0091 | 0.0210 |
| rs164938 | T | -0.0088 | 0.0014 | 0.0037 | 0.0106 |
| rs165633 | A | -0.0116 | 0.0016 | 0.0053 | 0.0127 |
| rs1656614 | C | 0.0122 | 0.0015 | -0.0142 | 0.0118 |
| rs1671269 | T | -0.0100 | 0.0016 | 0.0035 | 0.0112 |
| rs1671770 | A | 0.0125 | 0.0018 | 0.0119 | 0.0132 |
| rs16851779 | T | -0.0153 | 0.0027 | 0.0295 | 0.0218 |
| rs16871807 | T | 0.0090 | 0.0015 | -0.0080 | 0.0109 |
| rs1689510 | C | 0.0194 | 0.0015 | -0.0077 | 0.0115 |
| rs16901689 | T | 0.0111 | 0.0019 | -0.0294 | 0.0138 |
| rs1693584 | T | -0.0087 | 0.0015 | 0.0074 | 0.0112 |
| rs16958559 | T | 0.0096 | 0.0017 | 0.0234 | 0.0142 |
| rs16966271 | T | 0.0092 | 0.0016 | -0.0020 | 0.0116 |
| rs16975275 | A | -0.0102 | 0.0017 | -0.0006 | 0.0129 |
| rs17048801 | A | -0.0106 | 0.0014 | 0.0322 | 0.0112 |
| rs17048855 | A | 0.0108 | 0.0015 | 0.0007 | 0.0108 |
| rs17069646 | T | 0.0085 | 0.0015 | 0.0149 | 0.0109 |
| rs17110109 | T | -0.0093 | 0.0014 | 0.0140 | 0.0107 |
| rs17113730 | A | -0.0170 | 0.0027 | 0.0296 | 0.0200 |
| rs17131123 | A | -0.0243 | 0.0039 | -0.0059 | 0.0404 |
| rs17133297 | A | -0.0146 | 0.0023 | 0.0121 | 0.0155 |
| rs17144467 | A | 0.0098 | 0.0015 | -0.0116 | 0.0110 |
| rs17148998 | A | 0.0101 | 0.0017 | -0.0109 | 0.0136 |
| rs171697 | C | 0.0133 | 0.0015 | -0.0096 | 0.0111 |
| rs17170519 | C | 0.0132 | 0.0023 | 0.0045 | 0.0158 |
| rs1717204 | A | -0.0117 | 0.0018 | -0.0088 | 0.0142 |
| rs1718188 | A | 0.0096 | 0.0014 | -0.0203 | 0.0105 |
| rs17186106 | T | 0.0108 | 0.0018 | -0.0191 | 0.0145 |
| rs17190418 | T | -0.0213 | 0.0031 | 0.0071 | 0.0276 |
| rs17205908 | T | -0.0092 | 0.0015 | 0.0139 | 0.0120 |
| rs17248751 | A | -0.0136 | 0.0017 | 0.0340 | 0.0141 |
| rs1728118 | A | 0.0089 | 0.0016 | -0.0146 | 0.0113 |
| rs1729412 | T | -0.0107 | 0.0014 | 0.0097 | 0.0103 |
| rs173003 | A | -0.0082 | 0.0014 | -0.0048 | 0.0107 |
| rs1734370 | A | 0.0099 | 0.0016 | -0.0036 | 0.0112 |
| rs1738050 | C | -0.0093 | 0.0014 | 0.0014 | 0.0106 |
| rs17411339 | A | 0.0140 | 0.0014 | -0.0117 | 0.0110 |
| rs17428076 | C | 0.0137 | 0.0016 | 0.0020 | 0.0123 |
| rs17440885 | A | 0.0160 | 0.0029 | 0.0044 | 0.0246 |
| rs1747714 | T | 0.0106 | 0.0014 | 0.0251 | 0.0103 |
| rs1747817 | T | -0.0092 | 0.0016 | 0.0387 | 0.0126 |
| rs17489649 | A | 0.0127 | 0.0015 | 0.0000 | 0.0123 |
| rs17502934 | T | -0.0167 | 0.0020 | 0.0147 | 0.0146 |
| rs175325 | A | -0.0100 | 0.0014 | -0.0049 | 0.0105 |
| rs17536059 | C | -0.0108 | 0.0019 | 0.0103 | 0.0143 |
| rs17551064 | A | 0.0141 | 0.0019 | 0.0153 | 0.0149 |
| rs17563464 | A | -0.0147 | 0.0017 | -0.0008 | 0.0159 |
| rs17565975 | A | -0.0106 | 0.0014 | -0.0254 | 0.0103 |
| rs17568389 | A | 0.0127 | 0.0014 | -0.0038 | 0.0104 |
| rs17574007 | A | -0.0129 | 0.0021 | 0.0125 | 0.0177 |
| rs1758747 | A | 0.0092 | 0.0015 | -0.0155 | 0.0111 |
| rs17598675 | T | -0.0116 | 0.0014 | 0.0152 | 0.0105 |
| rs17604349 | A | 0.0127 | 0.0018 | -0.0100 | 0.0136 |
| rs17609255 | T | -0.0087 | 0.0014 | 0.0198 | 0.0104 |
| rs176218 | T | 0.0200 | 0.0018 | -0.0211 | 0.0124 |
| rs17622379 | T | -0.0113 | 0.0018 | 0.0002 | 0.0128 |
| rs17650634 | T | -0.0104 | 0.0019 | 0.0264 | 0.0150 |
| rs17667540 | A | -0.0087 | 0.0015 | 0.0066 | 0.0109 |
| rs17669337 | T | -0.0110 | 0.0014 | -0.0050 | 0.0106 |
| rs17680712 | T | -0.0092 | 0.0014 | 0.0106 | 0.0104 |
| rs17686649 | T | -0.0098 | 0.0017 | 0.0403 | 0.0124 |
| rs17732878 | T | -0.0095 | 0.0017 | -0.0216 | 0.0128 |
| rs17742342 | A | -0.0119 | 0.0018 | 0.0146 | 0.0135 |
| rs17747544 | A | -0.0090 | 0.0014 | 0.0079 | 0.0105 |
| rs1779549 | A | 0.0081 | 0.0014 | -0.0046 | 0.0108 |
| rs178183 | T | 0.0145 | 0.0016 | -0.0105 | 0.0118 |
| rs17882802 | A | 0.0084 | 0.0014 | -0.0141 | 0.0111 |
| rs17883331 | A | -0.0108 | 0.0018 | 0.0421 | 0.0141 |
| rs1792602 | A | -0.0097 | 0.0014 | -0.0014 | 0.0109 |
| rs181214 | T | -0.0104 | 0.0017 | 0.0150 | 0.0144 |
| rs182355396 | A | -0.0525 | 0.0084 | -0.0035 | 0.0774 |
| rs182902112 | A | -0.0300 | 0.0053 | 0.0217 | 0.0363 |
| rs1835339 | T | 0.0084 | 0.0014 | 0.0170 | 0.0105 |
| rs183869217 | C | 0.0174 | 0.0030 | -0.0113 | 0.0236 |
| rs1841023 | A | -0.0092 | 0.0015 | -0.0054 | 0.0117 |
| rs1842713 | A | -0.0117 | 0.0017 | -0.0037 | 0.0120 |
| rs1843815 | A | 0.0077 | 0.0014 | -0.0202 | 0.0102 |
| rs1861786 | A | -0.0081 | 0.0014 | 0.0027 | 0.0105 |
| rs1865955 | T | 0.0122 | 0.0019 | -0.0126 | 0.0128 |
| rs1866823 | A | 0.0107 | 0.0014 | -0.0176 | 0.0107 |
| rs187580 | T | -0.0104 | 0.0017 | 0.0286 | 0.0136 |
| rs187951956 | A | -0.0315 | 0.0058 | 0.0060 | 0.0418 |
| rs1880088 | A | -0.0097 | 0.0016 | 0.0264 | 0.0112 |
| rs1880692 | A | 0.0086 | 0.0014 | -0.0012 | 0.0105 |
| rs188251563 | A | 0.0428 | 0.0070 | -0.1369 | 0.0654 |
| rs1890132 | T | -0.0089 | 0.0015 | 0.0106 | 0.0115 |
| rs1898111 | A | 0.0101 | 0.0018 | -0.0131 | 0.0128 |
| rs190102446 | T | -0.0210 | 0.0037 | -0.0608 | 0.0299 |
| rs1905616 | A | 0.0083 | 0.0015 | -0.0220 | 0.0108 |
| rs1910005 | T | -0.0090 | 0.0015 | -0.0041 | 0.0113 |
| rs1918394 | T | 0.0119 | 0.0019 | -0.0214 | 0.0136 |
| rs192436652 | T | -0.0319 | 0.0045 | -0.0647 | 0.0430 |
| rs1933264 | T | 0.0100 | 0.0016 | -0.0009 | 0.0118 |
| rs1952183 | A | -0.0089 | 0.0014 | 0.0005 | 0.0106 |
| rs1955250 | A | -0.0140 | 0.0025 | 0.0331 | 0.0194 |
| rs1963381 | A | 0.0099 | 0.0016 | -0.0100 | 0.0117 |
| rs1980129 | A | 0.0085 | 0.0014 | -0.0009 | 0.0102 |
| rs198262 | T | -0.0223 | 0.0035 | 0.0036 | 0.0294 |
| rs1991585 | T | -0.0101 | 0.0015 | -0.0039 | 0.0109 |
| rs1995181 | A | -0.0078 | 0.0014 | -0.0156 | 0.0103 |
| rs2007655 | T | 0.0086 | 0.0014 | 0.0064 | 0.0101 |
| rs2011603 | A | -0.0091 | 0.0016 | 0.0425 | 0.0121 |
| rs2014830 | T | 0.0110 | 0.0015 | -0.0259 | 0.0113 |
| rs201495 | T | -0.0112 | 0.0019 | 0.0275 | 0.0112 |
| rs2023016 | C | 0.0107 | 0.0017 | -0.0137 | 0.0120 |
| rs2024568 | T | -0.0097 | 0.0016 | 0.0079 | 0.0116 |
| rs2029401 | A | -0.0091 | 0.0014 | 0.0183 | 0.0107 |
| rs2034631 | T | 0.0089 | 0.0015 | -0.0083 | 0.0106 |
| rs2039204 | A | 0.0082 | 0.0014 | 0.0031 | 0.0108 |
| rs2043187 | A | 0.0107 | 0.0015 | -0.0089 | 0.0120 |
| rs2052285 | A | 0.0112 | 0.0014 | -0.0152 | 0.0117 |
| rs2055940 | A | 0.0082 | 0.0015 | -0.0027 | 0.0120 |
| rs2077235 | T | 0.0097 | 0.0017 | 0.0055 | 0.0119 |
| rs2081652 | A | 0.0123 | 0.0015 | 0.0157 | 0.0112 |
| rs2082317 | T | -0.0078 | 0.0014 | -0.0123 | 0.0105 |
| rs2088913 | A | 0.0081 | 0.0014 | 0.0038 | 0.0103 |
| rs2092248 | T | -0.0083 | 0.0015 | 0.0125 | 0.0113 |
| rs2098526 | A | -0.0241 | 0.0043 | 0.0511 | 0.0277 |
| rs2100249 | T | -0.0089 | 0.0015 | 0.0139 | 0.0108 |
| rs2126069 | T | 0.0080 | 0.0014 | -0.0094 | 0.0105 |
| rs2131167 | A | 0.0080 | 0.0014 | -0.0229 | 0.0109 |
| rs214626 | A | 0.0101 | 0.0018 | -0.0096 | 0.0126 |
| rs2160514 | A | -0.0094 | 0.0014 | 0.0024 | 0.0105 |
| rs2179152 | T | -0.0131 | 0.0015 | -0.0079 | 0.0108 |
| rs2183271 | T | 0.0084 | 0.0015 | -0.0049 | 0.0110 |
| rs2199409 | T | 0.0103 | 0.0017 | -0.0155 | 0.0130 |
| rs2212430 | T | -0.0100 | 0.0016 | 0.0125 | 0.0113 |
| rs2216144 | T | 0.0098 | 0.0014 | -0.0139 | 0.0105 |
| rs2220926 | T | -0.0095 | 0.0014 | -0.0015 | 0.0103 |
| rs2250660 | C | 0.0080 | 0.0014 | -0.0103 | 0.0106 |
| rs2252098 | T | -0.0077 | 0.0014 | -0.0106 | 0.0105 |
| rs2254681 | A | 0.0101 | 0.0016 | 0.0046 | 0.0117 |
| rs2256965 | A | 0.0106 | 0.0014 | -0.0038 | 0.0111 |
| rs2276209 | A | 0.0085 | 0.0015 | -0.0171 | 0.0109 |
| rs2283076 | A | 0.0115 | 0.0017 | -0.0071 | 0.0127 |
| rs2287838 | A | -0.0104 | 0.0014 | -0.0037 | 0.0105 |
| rs2290601 | T | -0.0149 | 0.0017 | 0.0055 | 0.0119 |
| rs2297293 | C | 0.0099 | 0.0015 | 0.0022 | 0.0110 |
| rs2299156 | T | 0.0099 | 0.0018 | -0.0009 | 0.0128 |
| rs2314338 | T | 0.0093 | 0.0016 | -0.0110 | 0.0137 |
| rs2321157 | A | -0.0081 | 0.0014 | -0.0110 | 0.0103 |
| rs232496 | T | 0.0090 | 0.0015 | 0.0273 | 0.0105 |
| rs2332179 | A | 0.0121 | 0.0020 | -0.0123 | 0.0141 |
| rs2336721 | T | 0.0095 | 0.0015 | -0.0327 | 0.0109 |
| rs2358628 | A | -0.0091 | 0.0015 | -0.0025 | 0.0110 |
| rs236318 | C | -0.0102 | 0.0017 | -0.0032 | 0.0116 |
| rs2364544 | A | -0.0105 | 0.0014 | 0.0036 | 0.0107 |
| rs2365376 | A | 0.0092 | 0.0015 | 0.0070 | 0.0108 |
| rs2368831 | T | -0.0092 | 0.0014 | 0.0041 | 0.0113 |
| rs2373124 | A | -0.0096 | 0.0016 | 0.0154 | 0.0132 |
| rs2406253 | A | 0.0135 | 0.0018 | 0.0075 | 0.0145 |
| rs2414072 | A | -0.0083 | 0.0014 | -0.0078 | 0.0104 |
| rs2416214 | A | 0.0079 | 0.0014 | 0.0016 | 0.0104 |
| rs2416759 | A | -0.0093 | 0.0015 | 0.0075 | 0.0114 |
| rs2416845 | T | 0.0263 | 0.0048 | -0.0002 | 0.0341 |
| rs2431023 | A | 0.0093 | 0.0014 | -0.0046 | 0.0103 |
| rs2434672 | A | 0.0091 | 0.0014 | -0.0044 | 0.0105 |
| rs2436760 | T | 0.0232 | 0.0042 | 0.0072 | 0.0306 |
| rs2447097 | T | 0.0096 | 0.0014 | 0.0147 | 0.0109 |
| rs2458370 | T | -0.0086 | 0.0015 | 0.0134 | 0.0105 |
| rs2469226 | A | 0.0096 | 0.0017 | -0.0140 | 0.0128 |
| rs2470966 | C | -0.0103 | 0.0015 | 0.0113 | 0.0114 |
| rs2478208 | C | -0.0103 | 0.0014 | -0.0031 | 0.0105 |
| rs2496482 | T | 0.0121 | 0.0015 | -0.0287 | 0.0115 |
| rs2517086 | C | -0.0088 | 0.0014 | -0.0093 | 0.0112 |
| rs2521602 | A | -0.0266 | 0.0049 | 0.0104 | 0.0356 |
| rs2522545 | T | -0.0083 | 0.0014 | 0.0055 | 0.0105 |
| rs2526398 | C | -0.0222 | 0.0015 | 0.0150 | 0.0120 |
| rs2529069 | T | -0.0106 | 0.0016 | 0.0149 | 0.0113 |
| rs252991 | A | 0.0095 | 0.0015 | -0.0049 | 0.0107 |
| rs2542673 | A | 0.0088 | 0.0015 | 0.0133 | 0.0113 |
| rs2545795 | A | 0.0125 | 0.0014 | 0.0060 | 0.0104 |
| rs255053 | A | -0.0111 | 0.0018 | 0.0007 | 0.0127 |
| rs2554835 | A | 0.0088 | 0.0014 | -0.0026 | 0.0111 |
| rs2568955 | T | -0.0165 | 0.0016 | 0.0041 | 0.0123 |
| rs2569041 | T | 0.0077 | 0.0014 | -0.0058 | 0.0106 |
| rs2570497 | T | -0.0121 | 0.0015 | 0.0071 | 0.0106 |
| rs2588959 | T | -0.0080 | 0.0014 | 0.0027 | 0.0101 |
| rs2657283 | T | 0.0085 | 0.0014 | -0.0119 | 0.0108 |
| rs2665668 | A | -0.0094 | 0.0015 | 0.0231 | 0.0116 |
| rs2702576 | A | -0.0090 | 0.0014 | 0.0131 | 0.0103 |
| rs2718277 | T | 0.0142 | 0.0026 | -0.0176 | 0.0210 |
| rs2718791 | T | -0.0081 | 0.0015 | -0.0095 | 0.0112 |
| rs27220 | A | -0.0114 | 0.0015 | 0.0072 | 0.0105 |
| rs2725370 | T | -0.0141 | 0.0015 | 0.0221 | 0.0111 |
| rs2736752 | T | 0.0095 | 0.0017 | 0.0098 | 0.0140 |
| rs2740795 | A | 0.0098 | 0.0016 | -0.0079 | 0.0113 |
| rs2761438 | A | 0.0098 | 0.0015 | -0.0182 | 0.0109 |
| rs2764684 | T | 0.0143 | 0.0019 | -0.0185 | 0.0154 |
| rs2805064 | C | 0.0093 | 0.0016 | -0.0196 | 0.0124 |
| rs2838006 | T | 0.0132 | 0.0015 | -0.0175 | 0.0112 |
| rs28482086 | A | 0.0086 | 0.0015 | -0.0119 | 0.0107 |
| rs28505285 | C | 0.0106 | 0.0018 | 0.0219 | 0.0144 |
| rs28512462 | C | 0.0112 | 0.0014 | -0.0095 | 0.0104 |
| rs28513882 | A | -0.0110 | 0.0018 | 0.0184 | 0.0150 |
| rs2852349 | T | -0.0093 | 0.0014 | -0.0014 | 0.0102 |
| rs28587776 | T | 0.0088 | 0.0015 | 0.0104 | 0.0114 |
| rs28669886 | A | -0.0084 | 0.0015 | 0.0167 | 0.0109 |
| rs2870281 | A | -0.0083 | 0.0014 | -0.0090 | 0.0104 |
| rs28735993 | A | 0.0162 | 0.0022 | -0.0270 | 0.0177 |
| rs2885198 | A | 0.0088 | 0.0014 | -0.0210 | 0.0106 |
| rs2898191 | A | 0.0093 | 0.0015 | 0.0073 | 0.0109 |
| rs2910823 | T | 0.0115 | 0.0014 | 0.0081 | 0.0106 |
| rs2916490 | A | -0.0092 | 0.0015 | 0.0037 | 0.0111 |
| rs2923424 | A | 0.0120 | 0.0014 | 0.0149 | 0.0105 |
| rs2926702 | T | -0.0124 | 0.0021 | -0.0201 | 0.0136 |
| rs2929032 | A | -0.0081 | 0.0014 | 0.0070 | 0.0104 |
| rs2929860 | T | 0.0110 | 0.0018 | 0.0268 | 0.0147 |
| rs2942884 | A | -0.0090 | 0.0014 | 0.0093 | 0.0105 |
| rs2954114 | A | -0.0081 | 0.0014 | 0.0040 | 0.0110 |
| rs2958182 | A | 0.0082 | 0.0015 | -0.0131 | 0.0112 |
| rs2962378 | A | 0.0101 | 0.0017 | -0.0133 | 0.0120 |
| rs2964199 | T | -0.0099 | 0.0015 | 0.0117 | 0.0110 |
| rs2964255 | A | 0.0088 | 0.0015 | 0.0010 | 0.0111 |
| rs2976397 | T | 0.0081 | 0.0014 | 0.0146 | 0.0104 |
| rs2977464 | T | 0.0102 | 0.0018 | -0.0046 | 0.0128 |
| rs29792 | A | -0.0093 | 0.0015 | 0.0075 | 0.0115 |
| rs2980813 | A | 0.0088 | 0.0014 | 0.0189 | 0.0104 |
| rs2989476 | C | 0.0078 | 0.0014 | -0.0116 | 0.0104 |
| rs2989751 | A | -0.0092 | 0.0017 | 0.0127 | 0.0118 |
| rs2992037 | A | 0.0149 | 0.0015 | 0.0178 | 0.0121 |
| rs2994326 | T | -0.0100 | 0.0018 | 0.0005 | 0.0135 |
| rs2998299 | T | -0.0136 | 0.0017 | -0.0066 | 0.0134 |
| rs3026996 | A | 0.0138 | 0.0016 | -0.0077 | 0.0134 |
| rs303752 | A | -0.0110 | 0.0014 | 0.0236 | 0.0114 |
| rs3111251 | T | -0.0090 | 0.0014 | 0.0038 | 0.0108 |
| rs312927 | A | -0.0147 | 0.0022 | -0.0203 | 0.0188 |
| rs312945 | A | -0.0091 | 0.0015 | -0.0173 | 0.0111 |
| rs317050 | T | 0.0078 | 0.0014 | -0.0144 | 0.0103 |
| rs322614 | A | -0.0087 | 0.0014 | 0.0195 | 0.0105 |
| rs322744 | T | -0.0090 | 0.0016 | -0.0123 | 0.0114 |
| rs324885 | A | 0.0142 | 0.0014 | 0.0074 | 0.0106 |
| rs32940 | T | -0.0096 | 0.0015 | 0.0267 | 0.0114 |
| rs339057 | A | -0.0081 | 0.0014 | -0.0067 | 0.0106 |
| rs34067381 | T | -0.0104 | 0.0015 | -0.0058 | 0.0107 |
| rs34098770 | A | -0.0160 | 0.0020 | 0.0130 | 0.0138 |
| rs34106693 | C | 0.0159 | 0.0019 | -0.0118 | 0.0165 |
| rs34122915 | C | 0.0084 | 0.0014 | 0.0014 | 0.0120 |
| rs341504 | A | 0.0099 | 0.0015 | -0.0091 | 0.0108 |
| rs34155847 | A | 0.0135 | 0.0016 | -0.0264 | 0.0124 |
| rs34262657 | T | -0.0240 | 0.0038 | -0.0080 | 0.0245 |
| rs34286836 | T | -0.0089 | 0.0014 | 0.0050 | 0.0103 |
| rs34298584 | A | 0.0111 | 0.0019 | -0.0180 | 0.0140 |
| rs34305371 | A | 0.0314 | 0.0024 | 0.0100 | 0.0198 |
| rs34309 | A | 0.0080 | 0.0014 | 0.0130 | 0.0105 |
| rs34316274 | A | 0.0097 | 0.0017 | 0.0217 | 0.0121 |
| rs34363861 | A | 0.0087 | 0.0014 | -0.0184 | 0.0113 |
| rs34394051 | A | -0.0117 | 0.0020 | 0.0139 | 0.0136 |
| rs34410 | C | -0.0083 | 0.0014 | 0.0185 | 0.0106 |
| rs34624793 | T | 0.0116 | 0.0021 | 0.0177 | 0.0181 |
| rs34720381 | T | -0.0180 | 0.0024 | 0.0172 | 0.0196 |
| rs34748029 | A | 0.0185 | 0.0025 | -0.0422 | 0.0241 |
| rs347661 | T | -0.0081 | 0.0014 | -0.0020 | 0.0107 |
| rs34780702 | A | 0.0111 | 0.0017 | 0.0026 | 0.0117 |
| rs34807077 | A | 0.0128 | 0.0019 | -0.0157 | 0.0137 |
| rs34811474 | A | 0.0104 | 0.0017 | 0.0070 | 0.0164 |
| rs34967082 | A | -0.0080 | 0.0014 | -0.0042 | 0.0105 |
| rs35016816 | T | 0.0176 | 0.0023 | -0.0101 | 0.0162 |
| rs350281 | T | -0.0117 | 0.0015 | 0.0166 | 0.0130 |
| rs35104491 | A | 0.0124 | 0.0018 | 0.0071 | 0.0126 |
| rs35111506 | C | -0.0175 | 0.0030 | 0.0366 | 0.0232 |
| rs35192107 | T | -0.0091 | 0.0017 | 0.0044 | 0.0131 |
| rs35319653 | T | 0.0119 | 0.0015 | -0.0005 | 0.0113 |
| rs35417702 | T | -0.0152 | 0.0014 | 0.0157 | 0.0103 |
| rs35493937 | C | 0.0124 | 0.0017 | -0.0209 | 0.0123 |
| rs35606437 | A | 0.0105 | 0.0016 | -0.0059 | 0.0115 |
| rs356999 | A | -0.0101 | 0.0014 | 0.0129 | 0.0110 |
| rs35745455 | A | 0.0080 | 0.0014 | -0.0095 | 0.0105 |
| rs35754740 | T | 0.0083 | 0.0014 | -0.0132 | 0.0105 |
| rs35811586 | T | 0.0217 | 0.0031 | -0.0251 | 0.0297 |
| rs35929923 | A | -0.0114 | 0.0016 | 0.0257 | 0.0119 |
| rs36085856 | C | 0.0201 | 0.0036 | 0.0010 | 0.0278 |
| rs36120534 | T | 0.0100 | 0.0018 | 0.0149 | 0.0144 |
| rs362307 | T | -0.0226 | 0.0027 | 0.0286 | 0.0219 |
| rs363096 | T | -0.0143 | 0.0014 | 0.0251 | 0.0103 |
| rs3735478 | T | 0.0108 | 0.0016 | -0.0356 | 0.0122 |
| rs3751331 | A | -0.0097 | 0.0014 | 0.0089 | 0.0108 |
| rs3766979 | A | 0.0085 | 0.0015 | -0.0189 | 0.0107 |
| rs3768480 | C | 0.0101 | 0.0014 | -0.0284 | 0.0107 |
| rs3781339 | T | -0.0104 | 0.0018 | 0.0403 | 0.0138 |
| rs3790609 | T | 0.0107 | 0.0019 | -0.0013 | 0.0138 |
| rs3809169 | T | 0.0140 | 0.0025 | -0.0148 | 0.0180 |
| rs3809634 | A | -0.0109 | 0.0015 | 0.0162 | 0.0119 |
| rs3812281 | T | 0.0113 | 0.0014 | -0.0116 | 0.0104 |
| rs3817923 | A | -0.0121 | 0.0022 | 0.0304 | 0.0155 |
| rs382196 | T | -0.0082 | 0.0014 | 0.0121 | 0.0107 |
| rs3847228 | T | 0.0083 | 0.0014 | 0.0089 | 0.0110 |
| rs3848715 | A | -0.0080 | 0.0014 | 0.0070 | 0.0102 |
| rs3859523 | T | 0.0124 | 0.0018 | -0.0159 | 0.0135 |
| rs387027 | A | -0.0083 | 0.0015 | 0.0188 | 0.0108 |
| rs3895736 | A | 0.0144 | 0.0019 | 0.0279 | 0.0139 |
| rs3897821 | A | 0.0154 | 0.0015 | -0.0149 | 0.0107 |
| rs3948495 | T | -0.0089 | 0.0014 | 0.0104 | 0.0105 |
| rs399821 | A | -0.0078 | 0.0014 | -0.0121 | 0.0104 |
| rs39998 | A | 0.0090 | 0.0015 | -0.0173 | 0.0120 |
| rs401526 | T | -0.0104 | 0.0014 | 0.0047 | 0.0103 |
| rs401966 | C | -0.0097 | 0.0014 | -0.0087 | 0.0111 |
| rs4127499 | A | 0.0087 | 0.0015 | -0.0125 | 0.0117 |
| rs41282553 | A | 0.0237 | 0.0040 | -0.0331 | 0.0319 |
| rs42210 | C | -0.0098 | 0.0016 | -0.0179 | 0.0112 |
| rs42302 | A | 0.0086 | 0.0015 | -0.0011 | 0.0106 |
| rs4255791 | A | -0.0084 | 0.0015 | -0.0038 | 0.0127 |
| rs4263475 | A | 0.0078 | 0.0014 | 0.0104 | 0.0113 |
| rs4283754 | A | -0.0084 | 0.0014 | 0.0007 | 0.0127 |
| rs429150 | T | 0.0085 | 0.0014 | -0.0078 | 0.0111 |
| rs4298514 | T | 0.0104 | 0.0015 | 0.0003 | 0.0110 |
| rs4320563 | A | 0.0109 | 0.0014 | -0.0219 | 0.0103 |
| rs4328757 | T | 0.0096 | 0.0014 | 0.0018 | 0.0108 |
| rs4352658 | T | -0.0190 | 0.0026 | 0.0238 | 0.0185 |
| rs4358081 | A | -0.0095 | 0.0014 | 0.0093 | 0.0102 |
| rs4358358 | A | 0.0109 | 0.0015 | -0.0215 | 0.0112 |
| rs4369924 | A | 0.0125 | 0.0019 | -0.0023 | 0.0136 |
| rs4384309 | A | 0.0102 | 0.0014 | -0.0040 | 0.0109 |
| rs4396896 | A | -0.0107 | 0.0014 | -0.0069 | 0.0104 |
| rs4423373 | A | -0.0087 | 0.0014 | 0.0042 | 0.0108 |
| rs4426420 | T | -0.0140 | 0.0019 | 0.0108 | 0.0136 |
| rs4434676 | A | -0.0089 | 0.0014 | 0.0097 | 0.0106 |
| rs4458044 | C | 0.0101 | 0.0016 | 0.0085 | 0.0139 |
| rs4467547 | T | 0.0121 | 0.0014 | -0.0016 | 0.0114 |
| rs4469771 | T | 0.0101 | 0.0015 | 0.0004 | 0.0109 |
| rs4480339 | A | -0.0097 | 0.0015 | 0.0191 | 0.0118 |
| rs4490539 | A | 0.0142 | 0.0015 | -0.0030 | 0.0118 |
| rs4497562 | A | 0.0112 | 0.0016 | -0.0115 | 0.0117 |
| rs4500930 | T | -0.0109 | 0.0015 | 0.0382 | 0.0108 |
| rs4500960 | T | -0.0131 | 0.0014 | 0.0068 | 0.0103 |
| rs4502401 | T | 0.0085 | 0.0015 | -0.0125 | 0.0109 |
| rs4652135 | A | 0.0103 | 0.0016 | -0.0123 | 0.0113 |
| rs4658019 | T | 0.0084 | 0.0014 | -0.0040 | 0.0103 |
| rs4663617 | A | 0.0104 | 0.0017 | -0.0052 | 0.0123 |
| rs4664983 | T | -0.0103 | 0.0018 | 0.0063 | 0.0125 |
| rs4673840 | T | -0.0138 | 0.0019 | 0.0065 | 0.0137 |
| rs4675248 | A | -0.0094 | 0.0014 | -0.0129 | 0.0105 |
| rs4685405 | T | -0.0111 | 0.0018 | -0.0063 | 0.0122 |
| rs4687735 | T | -0.0200 | 0.0033 | -0.0428 | 0.0290 |
| rs4691601 | A | -0.0115 | 0.0014 | 0.0121 | 0.0103 |
| rs4705763 | C | 0.0089 | 0.0014 | 0.0140 | 0.0105 |
| rs4712371 | A | -0.0124 | 0.0021 | 0.0221 | 0.0143 |
| rs4719460 | T | -0.0088 | 0.0015 | 0.0007 | 0.0112 |
| rs4719944 | T | -0.0121 | 0.0014 | -0.0055 | 0.0103 |
| rs4724083 | T | 0.0083 | 0.0015 | -0.0082 | 0.0109 |
| rs4726070 | A | 0.0122 | 0.0014 | -0.0117 | 0.0104 |
| rs4730020 | T | 0.0090 | 0.0016 | -0.0005 | 0.0130 |
| rs4731413 | A | 0.0111 | 0.0017 | 0.0036 | 0.0142 |
| rs4735297 | A | -0.0086 | 0.0015 | -0.0153 | 0.0108 |
| rs4739235 | A | 0.0109 | 0.0019 | -0.0091 | 0.0134 |
| rs4741571 | A | -0.0102 | 0.0015 | -0.0125 | 0.0119 |
| rs4757957 | C | 0.0133 | 0.0015 | -0.0164 | 0.0112 |
| rs4766424 | C | 0.0142 | 0.0021 | -0.0364 | 0.0177 |
| rs4778058 | T | -0.0092 | 0.0014 | 0.0174 | 0.0111 |
| rs4785187 | A | -0.0100 | 0.0017 | -0.0151 | 0.0118 |
| rs4787028 | T | 0.0086 | 0.0015 | 0.0071 | 0.0112 |
| rs4787457 | A | 0.0162 | 0.0015 | -0.0148 | 0.0111 |
| rs4788115 | A | 0.0111 | 0.0019 | -0.0316 | 0.0182 |
| rs479018 | A | 0.0109 | 0.0015 | 0.0089 | 0.0122 |
| rs4793090 | A | 0.0084 | 0.0015 | -0.0421 | 0.0106 |
| rs4810894 | A | -0.0085 | 0.0015 | 0.0189 | 0.0106 |
| rs4812697 | T | 0.0171 | 0.0027 | -0.0095 | 0.0204 |
| rs481940 | T | 0.0100 | 0.0016 | -0.0168 | 0.0125 |
| rs482787 | T | 0.0100 | 0.0015 | -0.0228 | 0.0112 |
| rs4839155 | T | 0.0115 | 0.0017 | -0.0106 | 0.0122 |
| rs4846010 | A | 0.0099 | 0.0018 | -0.0425 | 0.0145 |
| rs4846724 | A | 0.0098 | 0.0014 | -0.0108 | 0.0102 |
| rs4848924 | A | -0.0117 | 0.0015 | -0.0102 | 0.0108 |
| rs4850954 | T | 0.0077 | 0.0014 | 0.0161 | 0.0108 |
| rs4851263 | A | -0.0151 | 0.0023 | 0.0562 | 0.0176 |
| rs4877516 | A | -0.0101 | 0.0014 | 0.0178 | 0.0103 |
| rs4881269 | A | 0.0100 | 0.0014 | 0.0193 | 0.0105 |
| rs488476 | C | -0.0113 | 0.0015 | -0.0003 | 0.0114 |
| rs4894658 | C | 0.0091 | 0.0016 | 0.0087 | 0.0113 |
| rs4895650 | T | 0.0093 | 0.0014 | 0.0045 | 0.0111 |
| rs4899012 | C | -0.0103 | 0.0014 | 0.0170 | 0.0105 |
| rs4904523 | A | -0.0082 | 0.0014 | 0.0185 | 0.0103 |
| rs4915735 | A | 0.0116 | 0.0020 | 0.0054 | 0.0155 |
| rs4919624 | A | 0.0193 | 0.0018 | -0.0250 | 0.0133 |
| rs4925065 | T | -0.0078 | 0.0014 | -0.0119 | 0.0105 |
| rs4925109 | A | -0.0096 | 0.0015 | 0.0308 | 0.0111 |
| rs4938815 | T | 0.0085 | 0.0015 | -0.0020 | 0.0115 |
| rs4941735 | T | 0.0092 | 0.0014 | -0.0037 | 0.0105 |
| rs4972748 | T | 0.0105 | 0.0018 | 0.0119 | 0.0126 |
| rs4977885 | A | -0.0084 | 0.0014 | 0.0319 | 0.0107 |
| rs4981245 | A | 0.0080 | 0.0014 | 0.0015 | 0.0109 |
| rs4984541 | A | -0.0138 | 0.0017 | -0.0010 | 0.0129 |
| rs4984613 | T | 0.0174 | 0.0028 | 0.0010 | 0.0198 |
| rs4984682 | C | -0.0138 | 0.0017 | -0.0049 | 0.0123 |
| rs548897 | A | 0.0081 | 0.0014 | -0.0199 | 0.0106 |
| rs55675587 | T | -0.0103 | 0.0018 | 0.0001 | 0.0128 |
| rs55736314 | C | -0.0154 | 0.0014 | 0.0105 | 0.0113 |
| rs55826493 | T | -0.0127 | 0.0021 | 0.0423 | 0.0157 |
| rs55986781 | T | 0.0187 | 0.0026 | -0.0410 | 0.0224 |
| rs56085180 | A | -0.0269 | 0.0037 | 0.0096 | 0.0257 |
| rs56099375 | T | 0.0120 | 0.0017 | -0.0059 | 0.0137 |
| rs56171318 | T | -0.0137 | 0.0020 | -0.0086 | 0.0139 |
| rs56174996 | A | -0.0143 | 0.0020 | 0.0060 | 0.0151 |
| rs56306882 | A | 0.0086 | 0.0016 | -0.0147 | 0.0127 |
| rs56319902 | T | -0.0206 | 0.0017 | 0.0484 | 0.0147 |
| rs56391344 | A | 0.0148 | 0.0016 | -0.0267 | 0.0129 |
| rs563954 | A | -0.0081 | 0.0014 | -0.0015 | 0.0106 |
| rs56405138 | A | 0.0153 | 0.0022 | -0.0026 | 0.0153 |
| rs56408528 | T | -0.0082 | 0.0014 | 0.0082 | 0.0104 |
| rs567003 | A | 0.0094 | 0.0015 | 0.0197 | 0.0114 |
| rs56794817 | A | 0.0122 | 0.0019 | -0.0097 | 0.0136 |
| rs57016874 | T | 0.0209 | 0.0037 | 0.0016 | 0.0316 |
| rs57148205 | A | 0.0095 | 0.0016 | -0.0140 | 0.0125 |
| rs57204268 | A | 0.0114 | 0.0020 | 0.0050 | 0.0154 |
| rs57349798 | A | 0.0093 | 0.0014 | -0.0474 | 0.0110 |
| rs57352738 | A | -0.0152 | 0.0017 | -0.0044 | 0.0138 |
| rs57437407 | A | -0.0145 | 0.0023 | -0.0293 | 0.0201 |
| rs575113 | A | 0.0133 | 0.0015 | -0.0182 | 0.0115 |
| rs5754581 | T | -0.0082 | 0.0014 | 0.0106 | 0.0103 |
| rs5754753 | T | -0.0126 | 0.0016 | 0.0046 | 0.0116 |
| rs5754762 | A | 0.0197 | 0.0030 | -0.0141 | 0.0198 |
| rs57661533 | T | -0.0114 | 0.0021 | -0.0105 | 0.0135 |
| rs580652 | T | -0.0147 | 0.0024 | -0.0109 | 0.0151 |
| rs585557 | A | 0.0128 | 0.0018 | -0.0295 | 0.0136 |
| rs58779949 | A | 0.0110 | 0.0019 | -0.0262 | 0.0151 |
| rs58859557 | T | 0.0187 | 0.0028 | 0.0074 | 0.0214 |
| rs58921703 | T | 0.0109 | 0.0015 | -0.0101 | 0.0110 |
| rs58950082 | T | 0.0095 | 0.0017 | -0.0069 | 0.0124 |
| rs58996896 | A | -0.0094 | 0.0016 | -0.0009 | 0.0124 |
| rs590013 | T | 0.0100 | 0.0015 | -0.0054 | 0.0110 |
| rs59300999 | T | -0.0164 | 0.0028 | 0.0128 | 0.0165 |
| rs59480703 | C | -0.0123 | 0.0018 | -0.0152 | 0.0126 |
| rs59813324 | T | 0.0097 | 0.0018 | -0.0057 | 0.0124 |
| rs59967356 | T | -0.0127 | 0.0021 | 0.0319 | 0.0199 |
| rs60096640 | A | 0.0159 | 0.0023 | 0.0084 | 0.0140 |
| rs6020560 | T | -0.0083 | 0.0014 | -0.0094 | 0.0102 |
| rs6043521 | T | -0.0083 | 0.0014 | 0.0109 | 0.0111 |
| rs60589532 | A | 0.0168 | 0.0028 | -0.0019 | 0.0207 |
| rs6060308 | A | 0.0095 | 0.0016 | -0.0205 | 0.0113 |
| rs6065080 | T | -0.0130 | 0.0015 | 0.0101 | 0.0105 |
| rs6065784 | C | 0.0110 | 0.0015 | 0.0004 | 0.0113 |
| rs60717745 | C | -0.0118 | 0.0019 | 0.0248 | 0.0137 |
| rs60726488 | T | -0.0090 | 0.0016 | 0.0027 | 0.0124 |
| rs6091570 | A | 0.0081 | 0.0015 | -0.0043 | 0.0107 |
| rs61104616 | A | -0.0172 | 0.0014 | -0.0022 | 0.0104 |
| rs6123924 | A | 0.0129 | 0.0019 | -0.0413 | 0.0132 |
| rs613872 | T | -0.0163 | 0.0019 | 0.0305 | 0.0143 |
| rs61387839 | A | 0.0088 | 0.0015 | -0.0181 | 0.0122 |
| rs61527214 | A | 0.0110 | 0.0014 | 0.0054 | 0.0115 |
| rs61739710 | A | -0.0099 | 0.0016 | -0.0132 | 0.0116 |
| rs61748951 | A | -0.0273 | 0.0048 | -0.0520 | 0.0502 |
| rs61755388 | T | 0.0125 | 0.0017 | -0.0045 | 0.0135 |
| rs61757207 | A | 0.0359 | 0.0063 | 0.0761 | 0.0476 |
| rs61798586 | A | -0.0112 | 0.0020 | 0.0360 | 0.0158 |
| rs61853335 | T | -0.0109 | 0.0018 | -0.0061 | 0.0128 |
| rs61958175 | A | -0.0198 | 0.0034 | 0.0284 | 0.0329 |
| rs61997667 | T | -0.0144 | 0.0020 | 0.0046 | 0.0148 |
| rs62007304 | A | 0.0264 | 0.0047 | 0.0143 | 0.0500 |
| rs62018216 | T | 0.0113 | 0.0019 | 0.0173 | 0.0151 |
| rs62051146 | A | -0.0134 | 0.0022 | 0.0189 | 0.0156 |
| rs62090515 | A | -0.0090 | 0.0015 | 0.0085 | 0.0108 |
| rs62092949 | T | -0.0087 | 0.0014 | -0.0085 | 0.0106 |
| rs62103198 | T | -0.0147 | 0.0025 | 0.0159 | 0.0225 |
| rs62109862 | A | 0.0136 | 0.0020 | 0.0114 | 0.0144 |
| rs62142891 | A | 0.0108 | 0.0016 | 0.0065 | 0.0127 |
| rs62155350 | A | 0.0338 | 0.0053 | -0.0429 | 0.0473 |
| rs62155770 | A | -0.0196 | 0.0032 | 0.0430 | 0.0284 |
| rs62155873 | T | -0.0151 | 0.0021 | -0.0189 | 0.0171 |
| rs62174974 | A | -0.0105 | 0.0018 | 0.0204 | 0.0122 |
| rs62177359 | A | -0.0275 | 0.0038 | 0.0004 | 0.0311 |
| rs62179650 | A | 0.0112 | 0.0016 | 0.0074 | 0.0114 |
| rs62182994 | T | 0.0137 | 0.0015 | -0.0043 | 0.0115 |
| rs62190914 | T | 0.0089 | 0.0015 | 0.0075 | 0.0112 |
| rs62194170 | A | 0.0111 | 0.0017 | -0.0157 | 0.0130 |
| rs622169 | T | 0.0086 | 0.0014 | -0.0068 | 0.0124 |
| rs62247449 | C | 0.0114 | 0.0014 | -0.0186 | 0.0107 |
| rs62252819 | A | -0.0094 | 0.0017 | -0.0160 | 0.0125 |
| rs62256284 | T | 0.0111 | 0.0017 | 0.0023 | 0.0126 |
| rs62260764 | C | -0.0132 | 0.0016 | -0.0167 | 0.0120 |
| rs62262671 | A | 0.0213 | 0.0020 | -0.0280 | 0.0171 |
| rs62340636 | T | 0.0103 | 0.0014 | -0.0033 | 0.0104 |
| rs62370510 | A | -0.0109 | 0.0019 | -0.0151 | 0.0146 |
| rs62379838 | T | 0.0121 | 0.0015 | -0.0163 | 0.0114 |
| rs62409395 | T | 0.0101 | 0.0018 | -0.0259 | 0.0184 |
| rs62420387 | T | -0.0184 | 0.0027 | 0.0143 | 0.0224 |
| rs62506074 | T | 0.0091 | 0.0015 | -0.0067 | 0.0122 |
| rs62506104 | A | -0.0098 | 0.0018 | 0.0128 | 0.0126 |
| rs628993 | A | 0.0128 | 0.0023 | 0.0103 | 0.0161 |
| rs631287 | A | 0.0084 | 0.0014 | -0.0281 | 0.0104 |
| rs633279 | A | -0.0089 | 0.0015 | -0.0096 | 0.0113 |
| rs635754 | A | 0.0119 | 0.0014 | 0.0052 | 0.0102 |
| rs6428587 | T | 0.0084 | 0.0015 | -0.0022 | 0.0105 |
| rs6435326 | A | -0.0091 | 0.0014 | -0.0097 | 0.0105 |
| rs6442126 | A | -0.0121 | 0.0015 | 0.0306 | 0.0114 |
| rs6445633 | A | 0.0088 | 0.0015 | 0.0107 | 0.0124 |
| rs6449503 | A | 0.0185 | 0.0014 | -0.0035 | 0.0105 |
| rs6450476 | A | -0.0116 | 0.0015 | 0.0097 | 0.0114 |
| rs6452793 | T | 0.0175 | 0.0017 | -0.0074 | 0.0121 |
| rs6461536 | A | 0.0088 | 0.0015 | 0.0123 | 0.0113 |
| rs6469654 | C | -0.0103 | 0.0017 | -0.0059 | 0.0116 |
| rs6472208 | T | -0.0091 | 0.0014 | 0.0036 | 0.0107 |
| rs6480234 | T | -0.0084 | 0.0014 | 0.0160 | 0.0104 |
| rs648044 | A | 0.0083 | 0.0015 | 0.0074 | 0.0115 |
| rs6490618 | T | 0.0088 | 0.0015 | -0.0098 | 0.0109 |
| rs6493265 | T | -0.0122 | 0.0014 | 0.0129 | 0.0108 |
| rs6493275 | A | 0.0131 | 0.0017 | 0.0074 | 0.0135 |
| rs6503409 | T | 0.0087 | 0.0015 | 0.0235 | 0.0117 |
| rs6534338 | T | 0.0103 | 0.0015 | -0.0146 | 0.0116 |
| rs6543810 | T | 0.0082 | 0.0015 | 0.0075 | 0.0105 |
| rs6557171 | T | -0.0155 | 0.0015 | -0.0008 | 0.0107 |
| rs6567288 | A | -0.0079 | 0.0014 | 0.0018 | 0.0103 |
| rs6569077 | T | 0.0118 | 0.0014 | -0.0032 | 0.0106 |
| rs6573552 | T | -0.0090 | 0.0014 | 0.0040 | 0.0102 |
| rs6573559 | T | 0.0105 | 0.0015 | 0.0023 | 0.0111 |
| rs660001 | A | -0.0164 | 0.0017 | 0.0062 | 0.0126 |
| rs663251 | T | 0.0081 | 0.0015 | -0.0023 | 0.0105 |
| rs66482320 | C | 0.0124 | 0.0022 | 0.0079 | 0.0165 |
| rs66568921 | T | -0.0181 | 0.0015 | 0.0164 | 0.0108 |
| rs66671632 | T | -0.0133 | 0.0021 | 0.0067 | 0.0148 |
| rs66721975 | A | -0.0087 | 0.0015 | 0.0016 | 0.0113 |
| rs6672986 | C | 0.0094 | 0.0014 | -0.0021 | 0.0102 |
| rs6689641 | A | 0.0078 | 0.0014 | 0.0034 | 0.0105 |
| rs6690195 | T | -0.0113 | 0.0014 | 0.0103 | 0.0108 |
| rs6695132 | T | 0.0097 | 0.0017 | -0.0139 | 0.0117 |
| rs6696068 | T | -0.0088 | 0.0014 | 0.0093 | 0.0109 |
| rs6697584 | T | -0.0126 | 0.0017 | 0.0215 | 0.0135 |
| rs6704768 | A | -0.0116 | 0.0014 | 0.0215 | 0.0106 |
| rs6706275 | T | 0.0083 | 0.0015 | -0.0090 | 0.0117 |
| rs6711399 | T | -0.0107 | 0.0019 | 0.0135 | 0.0123 |
| rs6715321 | T | -0.0097 | 0.0014 | 0.0100 | 0.0107 |
| rs6715849 | A | -0.0122 | 0.0014 | -0.0039 | 0.0105 |
| rs6720515 | A | 0.0103 | 0.0017 | 0.0042 | 0.0122 |
| rs67224963 | A | 0.0113 | 0.0017 | -0.0060 | 0.0126 |
| rs6729612 | T | 0.0092 | 0.0015 | -0.0017 | 0.0109 |
| rs6731373 | A | -0.0115 | 0.0015 | -0.0326 | 0.0121 |
| rs6736025 | T | 0.0079 | 0.0014 | -0.0158 | 0.0102 |
| rs6738860 | A | -0.0088 | 0.0014 | 0.0066 | 0.0105 |
| rs6743032 | A | -0.0165 | 0.0024 | -0.0377 | 0.0195 |
| rs6744040 | A | -0.0080 | 0.0014 | 0.0100 | 0.0103 |
| rs67456868 | A | 0.0119 | 0.0018 | -0.0099 | 0.0146 |
| rs6752228 | T | -0.0080 | 0.0014 | 0.0072 | 0.0102 |
| rs6757087 | T | -0.0085 | 0.0014 | 0.0005 | 0.0108 |
| rs6760887 | T | 0.0155 | 0.0022 | -0.0095 | 0.0129 |
| rs67661275 | A | 0.0134 | 0.0025 | -0.0307 | 0.0195 |
| rs6774533 | T | 0.0090 | 0.0015 | -0.0082 | 0.0112 |
| rs67790232 | A | 0.0101 | 0.0017 | 0.0059 | 0.0142 |
| rs6780414 | T | 0.0105 | 0.0018 | 0.0007 | 0.0132 |
| rs6792805 | A | -0.0090 | 0.0016 | -0.0116 | 0.0112 |
| rs6812533 | T | 0.0106 | 0.0016 | 0.0111 | 0.0114 |
| rs68145588 | T | -0.0127 | 0.0021 | -0.0063 | 0.0166 |
| rs6824567 | T | 0.0130 | 0.0016 | -0.0166 | 0.0122 |
| rs6853599 | T | 0.0108 | 0.0019 | -0.0113 | 0.0154 |
| rs6861925 | C | -0.0090 | 0.0014 | 0.0085 | 0.0102 |
| rs6871635 | A | -0.0087 | 0.0014 | -0.0095 | 0.0110 |
| rs6881581 | A | -0.0096 | 0.0015 | 0.0297 | 0.0113 |
| rs6881733 | T | -0.0086 | 0.0016 | -0.0056 | 0.0114 |
| rs6917154 | T | 0.0127 | 0.0021 | 0.0202 | 0.0175 |
| rs6917204 | T | 0.0119 | 0.0017 | -0.0330 | 0.0117 |
| rs6918506 | C | 0.0113 | 0.0014 | 0.0137 | 0.0103 |
| rs6924023 | A | 0.0107 | 0.0016 | 0.0011 | 0.0116 |
| rs6932108 | C | -0.0210 | 0.0027 | 0.0211 | 0.0215 |
| rs6946136 | A | 0.0081 | 0.0014 | 0.0031 | 0.0108 |
| rs6946362 | T | 0.0091 | 0.0015 | -0.0134 | 0.0121 |
| rs6951996 | A | -0.0182 | 0.0032 | 0.0262 | 0.0229 |
| rs6956283 | T | 0.0098 | 0.0016 | -0.0052 | 0.0117 |
| rs6969783 | A | -0.0098 | 0.0014 | 0.0230 | 0.0103 |
| rs6977237 | T | 0.0082 | 0.0014 | -0.0023 | 0.0104 |
| rs6989141 | A | 0.0100 | 0.0017 | -0.0018 | 0.0145 |
| rs6994287 | A | 0.0102 | 0.0014 | 0.0007 | 0.0106 |
| rs7012546 | T | 0.0092 | 0.0014 | -0.0141 | 0.0104 |
| rs7016302 | C | -0.0125 | 0.0019 | 0.0050 | 0.0146 |
| rs702606 | T | 0.0118 | 0.0021 | -0.0115 | 0.0148 |
| rs7029718 | A | 0.0244 | 0.0014 | -0.0012 | 0.0105 |
| rs7030373 | A | -0.0098 | 0.0017 | 0.0058 | 0.0120 |
| rs7035315 | A | -0.0090 | 0.0015 | -0.0092 | 0.0107 |
| rs7040995 | C | 0.0097 | 0.0014 | -0.0275 | 0.0104 |
| rs7041702 | A | -0.0111 | 0.0016 | 0.0247 | 0.0122 |
| rs7084508 | T | -0.0085 | 0.0015 | -0.0139 | 0.0115 |
| rs710629 | A | 0.0082 | 0.0015 | -0.0130 | 0.0106 |
| rs7108020 | A | 0.0111 | 0.0015 | -0.0134 | 0.0112 |
| rs711793 | T | 0.0103 | 0.0015 | -0.0021 | 0.0128 |
| rs7127580 | C | -0.0150 | 0.0023 | 0.0098 | 0.0191 |
| rs713584 | A | -0.0080 | 0.0014 | 0.0030 | 0.0102 |
| rs71413877 | A | 0.0351 | 0.0036 | -0.0293 | 0.0334 |
| rs7146625 | A | 0.0091 | 0.0016 | -0.0150 | 0.0123 |
| rs7147473 | A | -0.0103 | 0.0015 | 0.0207 | 0.0118 |
| rs7158218 | A | -0.0090 | 0.0015 | -0.0132 | 0.0117 |
| rs71646142 | T | 0.0104 | 0.0018 | -0.0002 | 0.0136 |
| rs716513 | A | 0.0084 | 0.0014 | -0.0086 | 0.0104 |
| rs7167688 | T | -0.0094 | 0.0014 | -0.0020 | 0.0103 |
| rs7171405 | A | -0.0101 | 0.0016 | 0.0117 | 0.0119 |
| rs717996 | T | -0.0122 | 0.0014 | 0.0210 | 0.0113 |
| rs7182216 | C | -0.0096 | 0.0017 | -0.0191 | 0.0136 |
| rs7190 | A | -0.0095 | 0.0015 | 0.0054 | 0.0122 |
| rs721579 | T | -0.0110 | 0.0016 | 0.0224 | 0.0115 |
| rs7218235 | A | -0.0097 | 0.0017 | 0.0080 | 0.0137 |
| rs7223311 | C | -0.0079 | 0.0014 | 0.0090 | 0.0112 |
| rs7226824 | T | -0.0078 | 0.0014 | -0.0042 | 0.0104 |
| rs72482130 | T | 0.0141 | 0.0019 | -0.0104 | 0.0138 |
| rs72486027 | T | -0.0104 | 0.0016 | 0.0044 | 0.0116 |
| rs7254263 | T | -0.0119 | 0.0016 | 0.0026 | 0.0113 |
| rs7255223 | A | 0.0097 | 0.0016 | -0.0006 | 0.0114 |
| rs72622559 | T | -0.0120 | 0.0017 | 0.0105 | 0.0123 |
| rs72624911 | T | 0.0282 | 0.0034 | -0.0660 | 0.0306 |
| rs72636697 | T | -0.0140 | 0.0020 | 0.0016 | 0.0142 |
| rs72667460 | T | 0.0197 | 0.0031 | -0.0450 | 0.0233 |
| rs72671456 | A | -0.0174 | 0.0020 | 0.0334 | 0.0160 |
| rs72672052 | A | -0.0101 | 0.0018 | 0.0095 | 0.0131 |
| rs72673097 | A | 0.0257 | 0.0042 | -0.0065 | 0.0446 |
| rs72677177 | A | 0.0100 | 0.0014 | 0.0101 | 0.0108 |
| rs72694479 | T | 0.0127 | 0.0020 | -0.0018 | 0.0155 |
| rs72709560 | A | 0.0085 | 0.0015 | -0.0085 | 0.0123 |
| rs72771860 | T | -0.0211 | 0.0029 | -0.0046 | 0.0278 |
| rs72792395 | T | -0.0145 | 0.0022 | 0.0043 | 0.0162 |
| rs728054 | A | -0.0128 | 0.0015 | 0.0135 | 0.0108 |
| rs72819118 | T | -0.0231 | 0.0024 | -0.0188 | 0.0210 |
| rs72824753 | T | 0.0133 | 0.0024 | -0.0103 | 0.0167 |
| rs72828517 | T | -0.0168 | 0.0019 | 0.0090 | 0.0140 |
| rs72829857 | A | -0.0149 | 0.0016 | 0.0066 | 0.0120 |
| rs72834698 | A | 0.0117 | 0.0020 | 0.0009 | 0.0176 |
| rs72881110 | A | 0.0146 | 0.0027 | 0.0159 | 0.0196 |
| rs72883760 | A | 0.0101 | 0.0018 | -0.0324 | 0.0133 |
| rs72896637 | T | 0.0162 | 0.0028 | -0.0358 | 0.0267 |
| rs72902523 | T | -0.0103 | 0.0016 | -0.0121 | 0.0115 |
| rs72906124 | T | 0.0185 | 0.0030 | -0.0447 | 0.0245 |
| rs72917504 | T | -0.0178 | 0.0030 | -0.0259 | 0.0228 |
| rs72919450 | T | -0.0132 | 0.0023 | 0.0119 | 0.0169 |
| rs72944064 | T | -0.0098 | 0.0016 | -0.0079 | 0.0126 |
| rs72962169 | T | -0.0174 | 0.0019 | -0.0209 | 0.0138 |
| rs72972965 | A | 0.0083 | 0.0015 | 0.0031 | 0.0114 |
| rs72993796 | T | 0.0127 | 0.0022 | 0.0184 | 0.0191 |
| rs73034295 | A | 0.0099 | 0.0018 | -0.0158 | 0.0143 |
| rs730384 | A | 0.0101 | 0.0014 | -0.0081 | 0.0106 |
| rs73039077 | C | 0.0092 | 0.0016 | -0.0093 | 0.0131 |
| rs73055556 | A | 0.0111 | 0.0020 | -0.0038 | 0.0151 |
| rs73082325 | C | -0.0252 | 0.0044 | 0.1306 | 0.0434 |
| rs73106136 | T | -0.0147 | 0.0026 | -0.0008 | 0.0209 |
| rs73154546 | A | 0.0197 | 0.0036 | -0.0032 | 0.0316 |
| rs7317761 | C | 0.0109 | 0.0016 | 0.0180 | 0.0111 |
| rs73191311 | A | -0.0094 | 0.0015 | -0.0197 | 0.0116 |
| rs7321274 | A | 0.0114 | 0.0018 | -0.0111 | 0.0127 |
| rs73219806 | A | 0.0106 | 0.0019 | 0.0059 | 0.0149 |
| rs7323027 | A | 0.0130 | 0.0015 | -0.0121 | 0.0106 |
| rs7326331 | A | -0.0128 | 0.0016 | -0.0010 | 0.0114 |
| rs73344830 | A | 0.0170 | 0.0014 | 0.0012 | 0.0108 |
| rs73405293 | A | 0.0113 | 0.0020 | -0.0079 | 0.0154 |
| rs73457936 | A | 0.0144 | 0.0025 | 0.0110 | 0.0216 |
| rs73518807 | C | -0.0128 | 0.0021 | 0.0191 | 0.0120 |
| rs7356536 | T | -0.0129 | 0.0021 | -0.0012 | 0.0156 |
| rs736281 | T | 0.0086 | 0.0014 | 0.0194 | 0.0108 |
| rs73643713 | T | -0.0165 | 0.0027 | 0.0213 | 0.0198 |
| rs736471 | T | 0.0083 | 0.0014 | 0.0042 | 0.0112 |
| rs73648455 | T | -0.0183 | 0.0026 | 0.0016 | 0.0185 |
| rs737945 | C | -0.0115 | 0.0014 | 0.0024 | 0.0105 |
| rs73874335 | T | -0.0178 | 0.0029 | 0.0158 | 0.0188 |
| rs73961845 | A | -0.0114 | 0.0014 | 0.0045 | 0.0110 |
| rs74091672 | A | -0.0153 | 0.0022 | 0.0115 | 0.0175 |
| rs7430651 | T | -0.0111 | 0.0016 | 0.0119 | 0.0111 |
| rs743316 | T | 0.0112 | 0.0017 | 0.0130 | 0.0136 |
| rs74415461 | T | 0.0176 | 0.0026 | -0.0249 | 0.0240 |
| rs74453875 | A | 0.0214 | 0.0037 | 0.0007 | 0.0249 |
| rs74462621 | T | 0.0125 | 0.0020 | 0.0044 | 0.0160 |
| rs7449561 | A | 0.0102 | 0.0017 | 0.0199 | 0.0122 |
| rs7451726 | A | 0.0109 | 0.0017 | 0.0014 | 0.0124 |
| rs74545339 | A | -0.0158 | 0.0022 | -0.0287 | 0.0164 |
| rs7460106 | T | -0.0123 | 0.0017 | 0.0117 | 0.0134 |
| rs74615093 | A | 0.0145 | 0.0026 | -0.0284 | 0.0214 |
| rs746839 | C | 0.0135 | 0.0015 | 0.0120 | 0.0115 |
| rs74747621 | T | 0.0153 | 0.0022 | -0.0287 | 0.0172 |
| rs74787922 | A | 0.0182 | 0.0028 | 0.0202 | 0.0219 |
| rs74944275 | T | 0.0231 | 0.0035 | -0.0524 | 0.0314 |
| rs74944857 | T | -0.0089 | 0.0015 | 0.0126 | 0.0114 |
| rs7495033 | A | 0.0095 | 0.0015 | -0.0238 | 0.0111 |
| rs75033012 | C | -0.0258 | 0.0038 | 0.0045 | 0.0379 |
| rs75177132 | T | 0.0298 | 0.0036 | -0.0008 | 0.0425 |
| rs75203411 | T | 0.0135 | 0.0019 | -0.0058 | 0.0142 |
| rs7526112 | T | 0.0124 | 0.0015 | 0.0028 | 0.0106 |
| rs75308819 | A | 0.0287 | 0.0047 | 0.0006 | 0.0398 |
| rs75434274 | A | 0.0151 | 0.0026 | -0.0563 | 0.0215 |
| rs75500877 | T | 0.0207 | 0.0033 | -0.0139 | 0.0279 |
| rs7552964 | A | 0.0148 | 0.0023 | -0.0178 | 0.0174 |
| rs7560871 | A | -0.0193 | 0.0027 | 0.0050 | 0.0214 |
| rs7561705 | A | 0.0078 | 0.0014 | -0.0024 | 0.0106 |
| rs7575637 | A | 0.0107 | 0.0014 | 0.0230 | 0.0103 |
| rs75756843 | A | 0.0288 | 0.0049 | 0.0029 | 0.0382 |
| rs7575938 | A | 0.0090 | 0.0015 | -0.0048 | 0.0106 |
| rs7590368 | T | -0.0146 | 0.0016 | 0.0115 | 0.0119 |
| rs7597126 | T | -0.0094 | 0.0014 | 0.0010 | 0.0108 |
| rs7597412 | C | 0.0088 | 0.0015 | 0.0107 | 0.0109 |
| rs7603132 | A | 0.0137 | 0.0018 | -0.0239 | 0.0128 |
| rs76076331 | T | 0.0206 | 0.0021 | -0.0319 | 0.0158 |
| rs76077165 | A | 0.0165 | 0.0025 | 0.0112 | 0.0203 |
| rs76167224 | T | -0.0187 | 0.0033 | -0.0233 | 0.0345 |
| rs7617204 | A | -0.0097 | 0.0014 | -0.0141 | 0.0103 |
| rs76235882 | A | 0.0241 | 0.0039 | -0.0273 | 0.0308 |
| rs76241605 | A | 0.0139 | 0.0022 | -0.0054 | 0.0206 |
| rs76246107 | A | -0.0162 | 0.0026 | -0.0266 | 0.0259 |
| rs7625428 | T | 0.0098 | 0.0014 | 0.0060 | 0.0107 |
| rs76267866 | A | 0.0104 | 0.0017 | -0.0175 | 0.0130 |
| rs7630133 | A | 0.0082 | 0.0015 | -0.0033 | 0.0105 |
| rs763553 | A | 0.0087 | 0.0014 | 0.0035 | 0.0104 |
| rs7640424 | T | 0.0113 | 0.0015 | -0.0074 | 0.0119 |
| rs7650602 | T | -0.0089 | 0.0014 | 0.0207 | 0.0108 |
| rs76552497 | T | -0.0120 | 0.0019 | 0.0290 | 0.0143 |
| rs76577427 | C | 0.0203 | 0.0024 | -0.0135 | 0.0221 |
| rs7672622 | A | -0.0088 | 0.0016 | 0.0167 | 0.0114 |
| rs767943 | A | -0.0122 | 0.0016 | 0.0202 | 0.0114 |
| rs7683416 | T | 0.0139 | 0.0014 | -0.0011 | 0.0104 |
| rs76876592 | A | 0.0126 | 0.0020 | 0.0193 | 0.0151 |
| rs76878669 | C | 0.0141 | 0.0017 | -0.0014 | 0.0124 |
| rs7692359 | T | -0.0109 | 0.0017 | 0.0298 | 0.0129 |
| rs76957677 | T | -0.0297 | 0.0054 | 0.0244 | 0.0718 |
| rs77025239 | A | -0.0140 | 0.0019 | 0.0068 | 0.0142 |
| rs7714719 | C | 0.0116 | 0.0020 | 0.0219 | 0.0147 |
| rs7716161 | C | 0.0105 | 0.0019 | 0.0259 | 0.0140 |
| rs77201694 | A | 0.0156 | 0.0022 | -0.0048 | 0.0205 |
| rs77370942 | A | -0.0147 | 0.0027 | 0.0027 | 0.0235 |
| rs77554090 | T | -0.0186 | 0.0027 | 0.0373 | 0.0230 |
| rs77609760 | A | 0.0236 | 0.0028 | -0.0051 | 0.0237 |
| rs7766240 | T | -0.0134 | 0.0024 | 0.0160 | 0.0189 |
| rs77702622 | A | -0.0213 | 0.0029 | -0.0030 | 0.0251 |
| rs77702819 | T | 0.0187 | 0.0025 | -0.0358 | 0.0218 |
| rs7772172 | A | 0.0091 | 0.0014 | -0.0189 | 0.0106 |
| rs7775100 | T | -0.0087 | 0.0014 | -0.0165 | 0.0106 |
| rs77826402 | T | 0.0151 | 0.0027 | -0.0092 | 0.0198 |
| rs7796103 | C | -0.0077 | 0.0014 | 0.0011 | 0.0103 |
| rs7799141 | A | 0.0099 | 0.0015 | -0.0094 | 0.0111 |
| rs77999825 | A | 0.0294 | 0.0039 | -0.0171 | 0.0368 |
| rs7803932 | A | 0.0127 | 0.0019 | -0.0155 | 0.0150 |
| rs780569 | A | -0.0085 | 0.0016 | 0.0141 | 0.0114 |
| rs78116078 | C | 0.0086 | 0.0016 | -0.0021 | 0.0122 |
| rs7816777 | T | -0.0088 | 0.0015 | 0.0078 | 0.0112 |
| rs78193153 | A | 0.0191 | 0.0029 | -0.0206 | 0.0190 |
| rs7823700 | T | 0.0132 | 0.0024 | -0.0310 | 0.0196 |
| rs78365243 | T | 0.0181 | 0.0033 | 0.0167 | 0.0298 |
| rs78440611 | A | -0.0147 | 0.0024 | 0.0196 | 0.0172 |
| rs7849480 | A | -0.0084 | 0.0014 | 0.0211 | 0.0105 |
| rs7849487 | T | -0.0167 | 0.0015 | -0.0154 | 0.0110 |
| rs7855503 | C | 0.0088 | 0.0015 | 0.0089 | 0.0109 |
| rs78648104 | T | -0.0141 | 0.0025 | 0.0156 | 0.0188 |
| rs78702390 | C | 0.0167 | 0.0030 | -0.0066 | 0.0263 |
| rs78714229 | T | 0.0172 | 0.0030 | 0.0263 | 0.0236 |
| rs7875078 | A | -0.0078 | 0.0014 | 0.0258 | 0.0105 |
| rs78918150 | T | -0.0080 | 0.0015 | 0.0025 | 0.0107 |
| rs7894722 | T | 0.0085 | 0.0014 | 0.0117 | 0.0110 |
| rs7899270 | A | -0.0084 | 0.0015 | 0.0088 | 0.0112 |
| rs7905192 | T | 0.0084 | 0.0014 | 0.0363 | 0.0103 |
| rs790647 | A | -0.0145 | 0.0017 | -0.0079 | 0.0118 |
| rs7910403 | T | 0.0122 | 0.0018 | 0.0075 | 0.0141 |
| rs7914674 | T | -0.0139 | 0.0022 | 0.0115 | 0.0131 |
| rs7920624 | A | 0.0120 | 0.0014 | -0.0128 | 0.0102 |
| rs7924036 | T | 0.0133 | 0.0014 | 0.0049 | 0.0101 |
| rs79265434 | A | -0.0197 | 0.0022 | 0.0136 | 0.0181 |
| rs7928017 | A | 0.0096 | 0.0014 | -0.0018 | 0.0104 |
| rs7931563 | T | -0.0109 | 0.0014 | -0.0040 | 0.0106 |
| rs795230 | T | 0.0081 | 0.0014 | -0.0028 | 0.0104 |
| rs7958371 | A | -0.0085 | 0.0015 | 0.0061 | 0.0107 |
| rs79585412 | T | -0.0178 | 0.0031 | -0.0453 | 0.0241 |
| rs7965154 | A | -0.0085 | 0.0015 | 0.0034 | 0.0110 |
| rs7967550 | A | -0.0086 | 0.0014 | 0.0120 | 0.0104 |
| rs7972246 | T | 0.0110 | 0.0015 | -0.0226 | 0.0108 |
| rs79728014 | A | -0.0134 | 0.0020 | 0.0040 | 0.0161 |
| rs7974852 | A | 0.0117 | 0.0014 | -0.0056 | 0.0104 |
| rs7977614 | A | -0.0134 | 0.0016 | 0.0255 | 0.0128 |
| rs79798166 | A | 0.0172 | 0.0025 | -0.0047 | 0.0215 |
| rs79855925 | T | -0.0194 | 0.0033 | 0.0143 | 0.0289 |
| rs79994730 | T | 0.0240 | 0.0035 | -0.0109 | 0.0310 |
| rs79997166 | A | 0.0172 | 0.0030 | -0.0260 | 0.0275 |
| rs80037907 | T | -0.0107 | 0.0019 | -0.0092 | 0.0152 |
| rs8008382 | T | -0.0106 | 0.0015 | -0.0005 | 0.0110 |
| rs8009933 | A | 0.0088 | 0.0015 | -0.0075 | 0.0108 |
| rs8016504 | A | -0.0087 | 0.0014 | 0.0083 | 0.0109 |
| rs80223410 | T | -0.0118 | 0.0020 | 0.0250 | 0.0202 |
| rs8024 | A | -0.0115 | 0.0015 | 0.0462 | 0.0111 |
| rs80257979 | T | -0.0252 | 0.0040 | -0.0556 | 0.0324 |
| rs8030487 | A | 0.0085 | 0.0015 | -0.0265 | 0.0115 |
| rs803619 | T | -0.0173 | 0.0023 | 0.0269 | 0.0169 |
| rs8046072 | A | 0.0101 | 0.0018 | -0.0227 | 0.0128 |
| rs8052523 | T | -0.0084 | 0.0014 | -0.0076 | 0.0112 |
| rs8066044 | A | 0.0090 | 0.0016 | -0.0142 | 0.0116 |
| rs806816 | A | 0.0101 | 0.0016 | -0.0040 | 0.0123 |
| rs8097125 | T | 0.0086 | 0.0014 | 0.0033 | 0.0109 |
| rs8103741 | A | -0.0108 | 0.0019 | -0.0022 | 0.0149 |
| rs818415 | T | -0.0110 | 0.0018 | 0.0029 | 0.0125 |
| rs8192465 | A | -0.0272 | 0.0049 | -0.0187 | 0.0474 |
| rs852771 | T | 0.0092 | 0.0015 | 0.0104 | 0.0115 |
| rs853286 | T | 0.0124 | 0.0023 | -0.0106 | 0.0148 |
| rs854796 | A | 0.0090 | 0.0015 | -0.0001 | 0.0110 |
| rs868456 | A | 0.0089 | 0.0015 | -0.0059 | 0.0115 |
| rs870589 | A | -0.0079 | 0.0014 | 0.0143 | 0.0104 |
| rs884108 | A | -0.0094 | 0.0017 | -0.0059 | 0.0129 |
| rs891793 | C | -0.0099 | 0.0014 | 0.0020 | 0.0104 |
| rs893522 | A | 0.0149 | 0.0025 | -0.0183 | 0.0186 |
| rs902820 | A | -0.0096 | 0.0014 | 0.0025 | 0.0106 |
| rs911149 | T | 0.0091 | 0.0017 | -0.0109 | 0.0119 |
| rs912883 | T | 0.0087 | 0.0015 | 0.0006 | 0.0109 |
| rs913509 | A | 0.0096 | 0.0015 | -0.0065 | 0.0109 |
| rs925161 | C | -0.0078 | 0.0014 | -0.0109 | 0.0104 |
| rs9267658 | T | 0.0138 | 0.0020 | -0.0457 | 0.0185 |
| rs9267677 | T | 0.0180 | 0.0024 | 0.0229 | 0.0168 |
| rs9289300 | T | -0.0164 | 0.0019 | 0.0233 | 0.0142 |
| rs9291437 | C | -0.0081 | 0.0014 | 0.0085 | 0.0104 |
| rs9294770 | T | 0.0083 | 0.0014 | -0.0089 | 0.0105 |
| rs929511 | T | -0.0160 | 0.0021 | 0.0041 | 0.0154 |
| rs9300612 | T | 0.0084 | 0.0015 | 0.0076 | 0.0108 |
| rs933738 | A | -0.0110 | 0.0018 | 0.0030 | 0.0138 |
| rs9349956 | A | -0.0159 | 0.0018 | -0.0252 | 0.0139 |
| rs9359939 | A | -0.0105 | 0.0016 | -0.0185 | 0.0116 |
| rs936496 | A | -0.0079 | 0.0014 | 0.0048 | 0.0107 |
| rs9371883 | C | -0.0087 | 0.0015 | 0.0040 | 0.0108 |
| rs9373363 | A | -0.0111 | 0.0016 | -0.0062 | 0.0120 |
| rs9375188 | T | 0.0212 | 0.0014 | 0.0014 | 0.0104 |
| rs9375403 | C | 0.0126 | 0.0016 | -0.0154 | 0.0125 |
| rs9386110 | T | -0.0163 | 0.0023 | 0.0021 | 0.0170 |
| rs9388490 | T | 0.0091 | 0.0014 | -0.0333 | 0.0106 |
| rs939400 | T | -0.0095 | 0.0015 | 0.0068 | 0.0107 |
| rs9411331 | A | 0.0145 | 0.0015 | -0.0220 | 0.0109 |
| rs9435340 | A | 0.0086 | 0.0015 | -0.0137 | 0.0107 |
| rs9442750 | A | -0.0089 | 0.0016 | -0.0001 | 0.0122 |
| rs9446060 | A | 0.0077 | 0.0014 | -0.0004 | 0.0103 |
| rs9465509 | A | -0.0084 | 0.0014 | 0.0044 | 0.0102 |
| rs9490512 | A | -0.0136 | 0.0014 | 0.0125 | 0.0104 |
| rs9492774 | C | 0.0080 | 0.0015 | -0.0197 | 0.0104 |
| rs9513416 | A | -0.0116 | 0.0019 | -0.0063 | 0.0149 |
| rs9513754 | T | 0.0096 | 0.0016 | 0.0086 | 0.0124 |
| rs9527662 | A | 0.0094 | 0.0014 | 0.0038 | 0.0110 |
| rs9527905 | A | -0.0094 | 0.0014 | -0.0126 | 0.0104 |
| rs9529146 | T | 0.0127 | 0.0017 | -0.0107 | 0.0116 |
| rs9536462 | A | 0.0133 | 0.0020 | -0.0133 | 0.0144 |
| rs9540718 | A | -0.0089 | 0.0014 | 0.0159 | 0.0104 |
| rs9545395 | T | 0.0138 | 0.0021 | -0.0039 | 0.0146 |
| rs9556958 | T | -0.0110 | 0.0014 | -0.0014 | 0.0104 |
| rs9563168 | A | 0.0101 | 0.0017 | -0.0020 | 0.0128 |
| rs9568798 | T | -0.0093 | 0.0016 | 0.0064 | 0.0115 |
| rs9597907 | T | 0.0152 | 0.0027 | -0.0066 | 0.0176 |
| rs9611597 | A | -0.0107 | 0.0019 | 0.0076 | 0.0166 |
| rs9616906 | A | 0.0122 | 0.0014 | -0.0094 | 0.0109 |
| rs9616947 | T | -0.0134 | 0.0015 | 0.0275 | 0.0118 |
| rs9633970 | T | 0.0091 | 0.0016 | 0.0000 | 0.0123 |
| rs9649 | T | 0.0108 | 0.0019 | -0.0012 | 0.0134 |
| rs9655780 | A | -0.0153 | 0.0019 | -0.0271 | 0.0155 |
| rs9666728 | T | -0.0082 | 0.0014 | -0.0017 | 0.0104 |
| rs9683585 | C | 0.0084 | 0.0014 | 0.0079 | 0.0102 |
| rs969512 | A | -0.0106 | 0.0015 | 0.0086 | 0.0107 |
| rs9771228 | T | 0.0096 | 0.0015 | -0.0197 | 0.0106 |
| rs977143 | A | 0.0085 | 0.0015 | -0.0008 | 0.0110 |
| rs978807 | A | 0.0139 | 0.0018 | -0.0233 | 0.0131 |
| rs981230 | T | 0.0100 | 0.0014 | 0.0136 | 0.0103 |
| rs981883 | A | 0.0099 | 0.0014 | -0.0187 | 0.0105 |
| rs9820604 | T | 0.0115 | 0.0018 | -0.0103 | 0.0128 |
| rs9821664 | A | 0.0094 | 0.0017 | -0.0057 | 0.0118 |
| rs9830359 | T | -0.0147 | 0.0022 | 0.0253 | 0.0159 |
| rs9844755 | A | 0.0083 | 0.0015 | -0.0028 | 0.0110 |
| rs9849884 | A | -0.0090 | 0.0014 | 0.0033 | 0.0109 |
| rs9853928 | T | -0.0127 | 0.0017 | 0.0273 | 0.0118 |
| rs9858921 | A | -0.0087 | 0.0014 | -0.0018 | 0.0104 |
| rs9859556 | T | 0.0290 | 0.0015 | -0.0423 | 0.0115 |
| rs9866123 | A | 0.0086 | 0.0014 | -0.0178 | 0.0108 |
| rs9870317 | A | 0.0086 | 0.0015 | -0.0306 | 0.0109 |
| rs9877225 | T | 0.0090 | 0.0016 | -0.0093 | 0.0118 |
| rs9882532 | T | 0.0124 | 0.0015 | 0.0019 | 0.0111 |
| rs9886703 | A | -0.0136 | 0.0019 | 0.0175 | 0.0137 |
| rs9888796 | T | 0.0113 | 0.0016 | -0.0086 | 0.0117 |
| rs9916901 | T | 0.0092 | 0.0016 | -0.0310 | 0.0111 |
| rs9924031 | C | 0.0093 | 0.0015 | 0.0036 | 0.0115 |
| rs9926649 | T | 0.0177 | 0.0031 | -0.0302 | 0.0254 |
| rs9927049 | A | 0.0099 | 0.0016 | 0.0109 | 0.0129 |
| rs9927137 | A | 0.0082 | 0.0014 | 0.0052 | 0.0115 |
| rs9927842 | T | -0.0130 | 0.0020 | -0.0067 | 0.0142 |
| rs9929556 | T | -0.0099 | 0.0014 | 0.0150 | 0.0105 |
| rs9929762 | A | 0.0096 | 0.0014 | 0.0046 | 0.0105 |
| rs9929993 | T | -0.0102 | 0.0015 | 0.0000 | 0.0109 |
| rs9933256 | A | 0.0119 | 0.0014 | 0.0003 | 0.0123 |
| rs995698 | A | -0.0099 | 0.0014 | -0.0070 | 0.0106 |
| rs9974899 | T | -0.0093 | 0.0016 | -0.0011 | 0.0119 |
| rs9977825 | T | -0.0092 | 0.0015 | 0.0307 | 0.0114 |

#### Supplementary table 13: Genetic association estimates for the IVW ratio-method MR analysis estimating the effect of education on SBP. ea=effect allele; gx=education; gy=SBP; se=standard error

| SNP | ea | gx | gx_se | gy | gy_se |
| --- | --- | --- | --- | --- | --- |
| rs10006235 | T | -0.0093 | 0.0016 | -0.0395 | 0.0497 |
| rs10009513 | A | 0.0088 | 0.0014 | 0.0018 | 0.0442 |
| rs10032941 | T | -0.0080 | 0.0015 | 0.0894 | 0.0460 |
| rs10041403 | T | 0.0151 | 0.0019 | 0.0401 | 0.0586 |
| rs10042828 | A | -0.0145 | 0.0022 | 0.1308 | 0.0682 |
| rs10046069 | A | 0.0127 | 0.0022 | 0.0832 | 0.0700 |
| rs10057590 | A | 0.0106 | 0.0014 | -0.0603 | 0.0447 |
| rs10060023 | T | 0.0110 | 0.0015 | -0.0364 | 0.0467 |
| rs10061420 | A | 0.0082 | 0.0014 | -0.0057 | 0.0443 |
| rs1006749 | A | 0.0079 | 0.0014 | -0.0187 | 0.0442 |
| rs10074178 | A | 0.0079 | 0.0014 | -0.0822 | 0.0443 |
| rs1007731 | A | -0.0143 | 0.0022 | 0.1084 | 0.0691 |
| rs10080647 | A | 0.0124 | 0.0020 | -0.1050 | 0.0641 |
| rs1008078 | T | -0.0182 | 0.0014 | 0.0074 | 0.0452 |
| rs1009470 | T | 0.0086 | 0.0014 | -0.0915 | 0.0445 |
| rs10098073 | A | -0.0091 | 0.0014 | 0.1493 | 0.0444 |
| rs10122669 | A | -0.0079 | 0.0014 | -0.0819 | 0.0448 |
| rs10128888 | A | -0.0083 | 0.0015 | -0.0346 | 0.0486 |
| rs10145520 | T | -0.0122 | 0.0018 | 0.0325 | 0.0554 |
| rs10166286 | T | 0.0151 | 0.0019 | -0.1156 | 0.0586 |
| rs10169002 | A | -0.0080 | 0.0014 | -0.0320 | 0.0444 |
| rs10181071 | A | 0.0102 | 0.0019 | -0.0164 | 0.0584 |
| rs10189857 | A | 0.0158 | 0.0014 | -0.0370 | 0.0446 |
| rs10192369 | A | -0.0078 | 0.0014 | 0.0122 | 0.0442 |
| rs10192834 | T | 0.0083 | 0.0015 | -0.0126 | 0.0464 |
| rs10193498 | A | 0.0097 | 0.0016 | 0.1208 | 0.0522 |
| rs10204051 | T | -0.0101 | 0.0014 | -0.0294 | 0.0453 |
| rs10205057 | T | 0.0090 | 0.0014 | -0.0305 | 0.0445 |
| rs10208 | T | 0.0091 | 0.0015 | 0.0797 | 0.0478 |
| rs10215082 | A | -0.0149 | 0.0014 | 0.0204 | 0.0443 |
| rs1024268 | T | -0.0084 | 0.0014 | -0.0620 | 0.0448 |
| rs10251438 | A | 0.0078 | 0.0014 | -0.0583 | 0.0445 |
| rs10264573 | A | 0.0175 | 0.0032 | -0.0713 | 0.1022 |
| rs1035578 | A | -0.0094 | 0.0014 | -0.0513 | 0.0445 |
| rs10402747 | T | 0.0077 | 0.0014 | -0.1542 | 0.0444 |
| rs10411759 | A | 0.0116 | 0.0020 | -0.3387 | 0.0624 |
| rs10444280 | T | 0.0100 | 0.0018 | 0.0137 | 0.0549 |
| rs10460095 | A | -0.0124 | 0.0014 | -0.0277 | 0.0447 |
| rs10480450 | A | 0.0158 | 0.0028 | 0.0624 | 0.0938 |
| rs10496091 | A | -0.0131 | 0.0016 | 0.1681 | 0.0492 |
| rs10499535 | A | 0.0080 | 0.0014 | -0.0400 | 0.0442 |
| rs1050847 | T | 0.0092 | 0.0014 | 0.0610 | 0.0449 |
| rs10509251 | T | -0.0093 | 0.0017 | 0.0569 | 0.0532 |
| rs10515007 | T | 0.0143 | 0.0019 | -0.0922 | 0.0605 |
| rs1051860 | A | -0.0079 | 0.0014 | -0.0216 | 0.0449 |
| rs10519504 | T | 0.0119 | 0.0019 | 0.0685 | 0.0610 |
| rs1054442 | A | -0.0136 | 0.0015 | 0.0070 | 0.0457 |
| rs1061801 | A | -0.0110 | 0.0018 | 0.0654 | 0.0562 |
| rs1066769 | A | -0.0230 | 0.0041 | 0.1637 | 0.1200 |
| rs10742591 | A | -0.0084 | 0.0014 | 0.0245 | 0.0454 |
| rs10750539 | A | -0.0095 | 0.0015 | -0.0672 | 0.0464 |
| rs10752262 | T | 0.0102 | 0.0014 | -0.0716 | 0.0446 |
| rs1075228 | A | -0.0081 | 0.0014 | -0.0255 | 0.0448 |
| rs10761202 | T | 0.0088 | 0.0014 | -0.0086 | 0.0442 |
| rs10761251 | A | 0.0113 | 0.0015 | 0.0800 | 0.0471 |
| rs10772644 | C | 0.0142 | 0.0022 | -0.1412 | 0.0715 |
| rs10773002 | A | 0.0185 | 0.0016 | -0.1224 | 0.0513 |
| rs10773208 | T | -0.0110 | 0.0016 | -0.0001 | 0.0506 |
| rs10782651 | T | -0.0082 | 0.0014 | -0.0246 | 0.0446 |
| rs10783243 | A | -0.0092 | 0.0014 | 0.0080 | 0.0443 |
| rs10789285 | T | 0.0091 | 0.0016 | -0.0654 | 0.0527 |
| rs10793903 | A | 0.0126 | 0.0022 | -0.0109 | 0.0719 |
| rs10795831 | T | -0.0099 | 0.0017 | -0.0020 | 0.0546 |
| rs10798888 | T | -0.0140 | 0.0019 | 0.0737 | 0.0586 |
| rs10799615 | A | -0.0089 | 0.0016 | -0.0256 | 0.0507 |
| rs10805383 | A | -0.0103 | 0.0014 | 0.0683 | 0.0443 |
| rs10810099 | A | -0.0146 | 0.0016 | 0.0260 | 0.0494 |
| rs10810145 | A | 0.0078 | 0.0014 | -0.1067 | 0.0449 |
| rs10830858 | T | 0.0092 | 0.0014 | -0.1799 | 0.0442 |
| rs10831656 | T | 0.0090 | 0.0015 | -0.0114 | 0.0481 |
| rs10844179 | A | 0.0104 | 0.0017 | -0.0852 | 0.0534 |
| rs10845051 | A | 0.0083 | 0.0014 | -0.0334 | 0.0444 |
| rs10875121 | C | 0.0174 | 0.0019 | 0.1168 | 0.0596 |
| rs10877283 | T | 0.0085 | 0.0014 | -0.0001 | 0.0451 |
| rs10879676 | T | -0.0083 | 0.0014 | 0.0993 | 0.0449 |
| rs10886012 | C | 0.0078 | 0.0014 | 0.0591 | 0.0448 |
| rs10892807 | T | 0.0109 | 0.0014 | -0.1401 | 0.0448 |
| rs10899282 | A | 0.0140 | 0.0017 | -0.1262 | 0.0539 |
| rs10927053 | A | -0.0125 | 0.0022 | 0.0073 | 0.0714 |
| rs10928190 | T | -0.0087 | 0.0014 | -0.0018 | 0.0445 |
| rs10931821 | A | -0.0141 | 0.0014 | 0.0703 | 0.0443 |
| rs10937240 | T | -0.0123 | 0.0018 | -0.0590 | 0.0564 |
| rs10940540 | A | 0.0078 | 0.0014 | 0.0789 | 0.0445 |
| rs10940921 | T | 0.0111 | 0.0014 | 0.0207 | 0.0450 |
| rs10943588 | A | 0.0084 | 0.0015 | -0.2603 | 0.0456 |
| rs10947439 | T | 0.0090 | 0.0015 | -0.0366 | 0.0472 |
| rs10949263 | A | -0.0091 | 0.0016 | 0.0543 | 0.0519 |
| rs10951590 | T | -0.0109 | 0.0015 | 0.0719 | 0.0470 |
| rs10974256 | A | -0.0093 | 0.0015 | -0.0697 | 0.0468 |
| rs1097784 | T | -0.0083 | 0.0014 | -0.0926 | 0.0443 |
| rs10979613 | T | -0.0117 | 0.0015 | 0.0375 | 0.0470 |
| rs10983324 | A | 0.0084 | 0.0015 | 0.1482 | 0.0485 |
| rs10984445 | A | 0.0115 | 0.0014 | -0.0293 | 0.0445 |
| rs10985402 | T | 0.0089 | 0.0016 | 0.0152 | 0.0517 |
| rs10995639 | T | 0.0078 | 0.0014 | -0.1465 | 0.0448 |
| rs11003463 | T | 0.0091 | 0.0015 | 0.0430 | 0.0453 |
| rs11021432 | A | 0.0103 | 0.0015 | 0.0266 | 0.0455 |
| rs11023764 | A | 0.0091 | 0.0015 | -0.0675 | 0.0472 |
| rs11030084 | T | 0.0106 | 0.0018 | -0.0168 | 0.0566 |
| rs11030102 | C | 0.0101 | 0.0016 | 0.0157 | 0.0501 |
| rs1106090 | A | 0.0094 | 0.0014 | -0.0769 | 0.0462 |
| rs11076962 | T | 0.0086 | 0.0016 | 0.0373 | 0.0489 |
| rs1107871 | A | -0.0083 | 0.0014 | -0.0425 | 0.0445 |
| rs11081529 | T | 0.0126 | 0.0015 | -0.0574 | 0.0486 |
| rs11082011 | T | 0.0194 | 0.0015 | -0.0672 | 0.0471 |
| rs11100237 | A | -0.0101 | 0.0014 | 0.0863 | 0.0442 |
| rs11115056 | A | 0.0187 | 0.0023 | -0.2173 | 0.0746 |
| rs11121177 | A | 0.0151 | 0.0018 | -0.1586 | 0.0585 |
| rs111226181 | T | -0.0201 | 0.0018 | 0.1488 | 0.0580 |
| rs111235962 | T | 0.0133 | 0.0024 | 0.1370 | 0.0741 |
| rs11130380 | T | 0.0090 | 0.0014 | 0.0977 | 0.0451 |
| rs111370527 | T | 0.0151 | 0.0027 | 0.0489 | 0.0814 |
| rs11138947 | T | 0.0105 | 0.0016 | -0.0895 | 0.0494 |
| rs111530150 | T | 0.0269 | 0.0048 | -0.1564 | 0.1882 |
| rs11155813 | T | -0.0129 | 0.0023 | 0.3294 | 0.0741 |
| rs11157931 | A | -0.0139 | 0.0014 | 0.1772 | 0.0452 |
| rs11158800 | A | -0.0091 | 0.0014 | -0.0267 | 0.0443 |
| rs1117152 | T | 0.0106 | 0.0015 | -0.0607 | 0.0486 |
| rs11174399 | A | 0.0111 | 0.0018 | -0.0974 | 0.0547 |
| rs111821073 | T | 0.0124 | 0.0019 | -0.0615 | 0.0612 |
| rs111852224 | T | 0.0156 | 0.0022 | -0.0815 | 0.0675 |
| rs11190955 | T | -0.0083 | 0.0015 | 0.1176 | 0.0462 |
| rs1120924 | T | 0.0108 | 0.0015 | -0.0207 | 0.0465 |
| rs11209894 | A | 0.0119 | 0.0021 | 0.0636 | 0.0642 |
| rs11210400 | A | 0.0120 | 0.0014 | 0.0281 | 0.0445 |
| rs11211123 | A | 0.0104 | 0.0017 | -0.1296 | 0.0524 |
| rs11213482 | A | -0.0106 | 0.0019 | 0.0640 | 0.0582 |
| rs112210983 | A | 0.0105 | 0.0017 | -0.0031 | 0.0537 |
| rs1123285 | C | -0.0094 | 0.0015 | 0.0421 | 0.0475 |
| rs112375785 | A | 0.0184 | 0.0021 | -0.1935 | 0.0632 |
| rs112379405 | C | -0.0082 | 0.0014 | -0.0325 | 0.0446 |
| rs112512729 | T | -0.0166 | 0.0027 | 0.1393 | 0.0879 |
| rs11265191 | T | 0.0083 | 0.0015 | 0.0248 | 0.0467 |
| rs112806496 | C | -0.0160 | 0.0025 | -0.0852 | 0.0768 |
| rs1128956 | T | -0.0140 | 0.0019 | -0.1189 | 0.0577 |
| rs113205706 | A | -0.0135 | 0.0024 | -0.1382 | 0.0746 |
| rs113520408 | A | 0.0155 | 0.0016 | -0.1223 | 0.0491 |
| rs113552169 | C | 0.0294 | 0.0053 | 0.1768 | 0.1662 |
| rs113588399 | T | -0.0104 | 0.0017 | 0.0842 | 0.0542 |
| rs113615161 | T | -0.0123 | 0.0021 | 0.0547 | 0.0649 |
| rs113720505 | A | 0.0164 | 0.0029 | -0.0808 | 0.0917 |
| rs113731629 | T | 0.0339 | 0.0061 | -0.2119 | 0.1908 |
| rs114192810 | T | -0.0279 | 0.0051 | 0.1908 | 0.1567 |
| rs114468556 | A | 0.0293 | 0.0036 | -0.1341 | 0.1060 |
| rs114593137 | A | -0.0122 | 0.0017 | 0.1157 | 0.0548 |
| rs1146079 | C | -0.0087 | 0.0016 | 0.0946 | 0.0503 |
| rs114810763 | A | -0.0121 | 0.0020 | 0.0681 | 0.0589 |
| rs114952970 | T | 0.0265 | 0.0040 | 0.0205 | 0.1313 |
| rs114958262 | T | -0.0220 | 0.0039 | -0.0636 | 0.1214 |
| rs115000530 | A | -0.0250 | 0.0031 | 0.2215 | 0.0973 |
| rs115017135 | A | -0.0206 | 0.0038 | 0.2565 | 0.1196 |
| rs11542663 | A | 0.0112 | 0.0015 | -0.1018 | 0.0486 |
| rs115438240 | T | -0.0169 | 0.0030 | -0.0608 | 0.1000 |
| rs115693355 | A | 0.0309 | 0.0053 | -0.2054 | 0.1635 |
| rs115732722 | A | -0.0415 | 0.0072 | -0.4482 | 0.2584 |
| rs11588857 | A | 0.0199 | 0.0017 | -0.0519 | 0.0542 |
| rs11591870 | T | 0.0098 | 0.0017 | -0.0696 | 0.0541 |
| rs11592299 | A | 0.0097 | 0.0018 | -0.1175 | 0.0547 |
| rs11596387 | T | 0.0258 | 0.0046 | -0.0052 | 0.1510 |
| rs11598765 | A | -0.0106 | 0.0018 | 0.0630 | 0.0564 |
| rs11599236 | T | -0.0106 | 0.0014 | -0.0839 | 0.0455 |
| rs11613431 | A | -0.0094 | 0.0016 | -0.0867 | 0.0500 |
| rs11620355 | A | 0.0182 | 0.0025 | -0.1094 | 0.0793 |
| rs11623285 | T | -0.0135 | 0.0021 | 0.1113 | 0.0643 |
| rs116386746 | C | 0.0249 | 0.0045 | -0.0148 | 0.1433 |
| rs11640569 | A | 0.0112 | 0.0020 | -0.0580 | 0.0642 |
| rs11644446 | A | 0.0117 | 0.0019 | 0.1033 | 0.0629 |
| rs11647188 | A | 0.0079 | 0.0014 | -0.0143 | 0.0452 |
| rs11652522 | A | -0.0154 | 0.0023 | -0.2832 | 0.0738 |
| rs11657342 | A | 0.0140 | 0.0019 | -0.2108 | 0.0458 |
| rs11657979 | A | -0.0091 | 0.0017 | 0.0449 | 0.0522 |
| rs11663602 | A | -0.0130 | 0.0016 | -0.0399 | 0.0496 |
| rs11663678 | T | -0.0121 | 0.0022 | 0.1341 | 0.0681 |
| rs116656374 | T | -0.0242 | 0.0038 | -0.0390 | 0.1157 |
| rs11675476 | T | -0.0127 | 0.0014 | 0.1318 | 0.0442 |
| rs1167827 | A | 0.0104 | 0.0014 | -0.2630 | 0.0446 |
| rs11678980 | A | -0.0166 | 0.0014 | 0.1217 | 0.0458 |
| rs11687736 | A | -0.0105 | 0.0015 | -0.1214 | 0.0474 |
| rs11693885 | A | -0.0091 | 0.0014 | 0.0204 | 0.0446 |
| rs116967397 | A | 0.0315 | 0.0056 | -0.2708 | 0.1686 |
| rs117005905 | T | 0.0150 | 0.0022 | -0.0204 | 0.0719 |
| rs11703948 | A | -0.0168 | 0.0024 | 0.1094 | 0.0726 |
| rs1171040 | A | -0.0113 | 0.0017 | 0.0728 | 0.0546 |
| rs11716398 | A | 0.0159 | 0.0021 | -0.1567 | 0.0660 |
| rs11720093 | T | 0.0150 | 0.0026 | 0.0006 | 0.0807 |
| rs11720985 | T | 0.0085 | 0.0015 | -0.0300 | 0.0479 |
| rs11724690 | T | 0.0093 | 0.0015 | 0.0033 | 0.0494 |
| rs117273411 | T | -0.0211 | 0.0034 | 0.1545 | 0.1111 |
| rs11732160 | A | 0.0105 | 0.0016 | -0.0739 | 0.0504 |
| rs11732657 | A | -0.0126 | 0.0016 | -0.0588 | 0.0506 |
| rs11733439 | A | -0.0096 | 0.0017 | 0.1345 | 0.0548 |
| rs11736863 | A | 0.0142 | 0.0018 | -0.0198 | 0.0574 |
| rs117398064 | C | -0.0141 | 0.0025 | -0.0412 | 0.0753 |
| rs117468730 | A | -0.0310 | 0.0050 | -0.2872 | 0.1565 |
| rs117623407 | A | -0.0118 | 0.0020 | 0.2018 | 0.0624 |
| rs11765387 | T | 0.0086 | 0.0015 | 0.0553 | 0.0471 |
| rs117668569 | T | 0.0234 | 0.0042 | 0.0391 | 0.1346 |
| rs117677995 | A | -0.0180 | 0.0032 | 0.2530 | 0.1001 |
| rs11772232 | T | 0.0177 | 0.0019 | -0.0771 | 0.0609 |
| rs11774212 | T | 0.0128 | 0.0014 | -0.0260 | 0.0443 |
| rs117799466 | C | 0.0106 | 0.0016 | -0.0442 | 0.0483 |
| rs11780023 | T | 0.0085 | 0.0014 | 0.1344 | 0.0442 |
| rs11789013 | T | 0.0106 | 0.0016 | -0.0170 | 0.0519 |
| rs117895796 | T | 0.0362 | 0.0064 | -0.1861 | 0.2169 |
| rs118040169 | A | -0.0248 | 0.0038 | 0.0451 | 0.1183 |
| rs118093058 | T | 0.0146 | 0.0021 | -0.1332 | 0.0640 |
| rs118134876 | T | -0.0223 | 0.0030 | 0.4038 | 0.0928 |
| rs11845781 | T | -0.0079 | 0.0014 | -0.0347 | 0.0445 |
| rs11871429 | A | 0.0128 | 0.0017 | 0.2242 | 0.0527 |
| rs11876620 | T | 0.0143 | 0.0024 | 0.0733 | 0.0739 |
| rs11894424 | A | 0.0209 | 0.0033 | 0.0340 | 0.1072 |
| rs11917701 | A | -0.0087 | 0.0014 | 0.0501 | 0.0453 |
| rs11919835 | C | 0.0089 | 0.0015 | -0.1669 | 0.0459 |
| rs1196760 | C | 0.0139 | 0.0025 | -0.2964 | 0.0760 |
| rs12005151 | A | 0.0087 | 0.0015 | -0.0459 | 0.0466 |
| rs12028010 | T | 0.0164 | 0.0017 | 0.1085 | 0.0524 |
| rs12028229 | T | 0.0099 | 0.0016 | -0.0008 | 0.0505 |
| rs12054166 | C | 0.0096 | 0.0016 | -0.1526 | 0.0507 |
| rs12076635 | C | 0.0207 | 0.0017 | 0.0879 | 0.0535 |
| rs12113634 | T | 0.0088 | 0.0015 | -0.0521 | 0.0463 |
| rs12123293 | T | 0.0086 | 0.0015 | 0.0069 | 0.0460 |
| rs12126231 | A | 0.0095 | 0.0014 | -0.2249 | 0.0450 |
| rs12127928 | T | 0.0108 | 0.0018 | 0.1129 | 0.0554 |
| rs12145078 | A | -0.0127 | 0.0017 | 0.0137 | 0.0552 |
| rs12151248 | T | -0.0149 | 0.0023 | -0.0953 | 0.0688 |
| rs12170452 | A | 0.0105 | 0.0014 | -0.0316 | 0.0445 |
| rs1220779 | A | -0.0079 | 0.0014 | 0.0220 | 0.0442 |
| rs12234369 | A | 0.0087 | 0.0015 | -0.0205 | 0.0479 |
| rs12238011 | T | -0.0141 | 0.0026 | 0.0706 | 0.0803 |
| rs12273435 | A | -0.0132 | 0.0017 | 0.1307 | 0.0546 |
| rs12285074 | A | 0.0161 | 0.0023 | 0.3027 | 0.0772 |
| rs12304188 | A | 0.0111 | 0.0019 | -0.0315 | 0.0617 |
| rs12325727 | A | 0.0088 | 0.0014 | -0.1377 | 0.0443 |
| rs12332731 | A | 0.0135 | 0.0018 | 0.0566 | 0.0554 |
| rs12342546 | A | -0.0128 | 0.0017 | 0.0201 | 0.0522 |
| rs12359372 | T | -0.0092 | 0.0015 | 0.0698 | 0.0473 |
| rs12375949 | T | -0.0138 | 0.0014 | 0.0686 | 0.0445 |
| rs12405889 | T | -0.0098 | 0.0014 | 0.0328 | 0.0440 |
| rs12431682 | T | -0.0093 | 0.0015 | 0.0124 | 0.0459 |
| rs12438177 | A | 0.0093 | 0.0015 | -0.0783 | 0.0459 |
| rs12453682 | T | 0.0102 | 0.0015 | 0.0471 | 0.0483 |
| rs1245829 | A | -0.0099 | 0.0014 | 0.2447 | 0.0447 |
| rs12467175 | T | -0.0081 | 0.0014 | 0.0752 | 0.0442 |
| rs12468040 | T | 0.0137 | 0.0014 | -0.1166 | 0.0455 |
| rs12473986 | T | -0.0085 | 0.0015 | 0.1137 | 0.0487 |
| rs12477385 | T | 0.0109 | 0.0017 | -0.0171 | 0.0528 |
| rs12478156 | T | 0.0108 | 0.0015 | 0.0611 | 0.0463 |
| rs12503522 | T | -0.0113 | 0.0016 | 0.0315 | 0.0489 |
| rs12506221 | T | -0.0130 | 0.0014 | 0.0966 | 0.0448 |
| rs12516990 | C | -0.0099 | 0.0017 | 0.0613 | 0.0547 |
| rs12519073 | T | -0.0107 | 0.0017 | 0.0375 | 0.0530 |
| rs12524795 | T | 0.0102 | 0.0014 | -0.0163 | 0.0444 |
| rs12568153 | T | -0.0091 | 0.0015 | 0.1598 | 0.0488 |
| rs12571549 | A | 0.0151 | 0.0020 | -0.0086 | 0.0644 |
| rs12574281 | A | -0.0080 | 0.0015 | -0.0456 | 0.0456 |
| rs12591647 | T | -0.0152 | 0.0018 | 0.0515 | 0.0578 |
| rs12601380 | A | -0.0082 | 0.0014 | 0.1563 | 0.0447 |
| rs12602286 | T | 0.0160 | 0.0021 | 0.0653 | 0.0668 |
| rs12613500 | C | 0.0098 | 0.0014 | 0.0076 | 0.0448 |
| rs12614263 | A | 0.0083 | 0.0014 | -0.0323 | 0.0443 |
| rs12638072 | A | -0.0107 | 0.0015 | -0.1098 | 0.0491 |
| rs12643771 | T | 0.0130 | 0.0015 | -0.0068 | 0.0479 |
| rs12646216 | T | 0.0084 | 0.0014 | -0.1530 | 0.0454 |
| rs12646523 | T | -0.0132 | 0.0016 | 0.0686 | 0.0508 |
| rs12659776 | T | 0.0085 | 0.0014 | -0.1029 | 0.0444 |
| rs12670376 | A | 0.0092 | 0.0014 | -0.0962 | 0.0444 |
| rs1267062 | C | 0.0113 | 0.0018 | 0.0835 | 0.0578 |
| rs12682775 | T | -0.0119 | 0.0017 | 0.1087 | 0.0537 |
| rs12694681 | T | 0.0102 | 0.0015 | -0.1316 | 0.0476 |
| rs12709186 | A | -0.0091 | 0.0015 | 0.0050 | 0.0468 |
| rs12712269 | T | -0.0107 | 0.0014 | 0.1305 | 0.0455 |
| rs12746551 | C | 0.0248 | 0.0041 | 0.0941 | 0.1251 |
| rs12750688 | C | 0.0116 | 0.0016 | -0.0707 | 0.0496 |
| rs12761729 | T | -0.0098 | 0.0016 | -0.0036 | 0.0509 |
| rs12761761 | T | 0.0153 | 0.0017 | -0.0578 | 0.0508 |
| rs12764593 | C | 0.0202 | 0.0029 | -0.1161 | 0.0952 |
| rs12765185 | A | -0.0088 | 0.0016 | 0.0837 | 0.0498 |
| rs12774577 | T | 0.0142 | 0.0020 | -0.1128 | 0.0630 |
| rs12789313 | T | 0.0092 | 0.0014 | -0.1019 | 0.0441 |
| rs12790196 | T | -0.0101 | 0.0015 | -0.2739 | 0.0470 |
| rs12810587 | T | 0.0094 | 0.0015 | -0.0637 | 0.0476 |
| rs12875339 | A | -0.0108 | 0.0015 | -0.0828 | 0.0463 |
| rs12888615 | T | 0.0120 | 0.0018 | 0.1015 | 0.0552 |
| rs12891042 | T | 0.0092 | 0.0014 | -0.0625 | 0.0453 |
| rs12891191 | C | 0.0080 | 0.0014 | -0.0685 | 0.0443 |
| rs12912465 | T | -0.0104 | 0.0016 | 0.0435 | 0.0508 |
| rs12952191 | T | -0.0081 | 0.0014 | 0.1622 | 0.0442 |
| rs12953422 | A | -0.0080 | 0.0014 | 0.0330 | 0.0449 |
| rs12957463 | A | 0.0146 | 0.0018 | -0.0001 | 0.0550 |
| rs12962845 | C | -0.0123 | 0.0021 | -0.0356 | 0.0651 |
| rs12967010 | T | 0.0114 | 0.0017 | 0.0300 | 0.0534 |
| rs12970264 | A | -0.0085 | 0.0014 | 0.0413 | 0.0445 |
| rs12981405 | T | -0.0112 | 0.0019 | -0.0259 | 0.0594 |
| rs13009915 | T | 0.0212 | 0.0037 | -0.0801 | 0.1148 |
| rs13010288 | T | 0.0198 | 0.0021 | -0.0054 | 0.0647 |
| rs13010566 | A | -0.0091 | 0.0014 | 0.1162 | 0.0451 |
| rs13015496 | C | 0.0119 | 0.0018 | -0.1061 | 0.0568 |
| rs13016316 | A | -0.0138 | 0.0024 | -0.2364 | 0.0763 |
| rs1301838 | T | -0.0096 | 0.0015 | 0.0598 | 0.0470 |
| rs13018640 | T | -0.0215 | 0.0014 | 0.0231 | 0.0450 |
| rs13026611 | A | -0.0084 | 0.0015 | 0.1297 | 0.0463 |
| rs13029602 | T | -0.0092 | 0.0014 | 0.0780 | 0.0453 |
| rs13034349 | T | -0.0082 | 0.0014 | 0.0469 | 0.0451 |
| rs13050131 | A | -0.0081 | 0.0015 | -0.0386 | 0.0474 |
| rs13085461 | C | 0.0103 | 0.0014 | -0.1156 | 0.0446 |
| rs13099165 | T | -0.0118 | 0.0017 | 0.0368 | 0.0548 |
| rs13107325 | T | -0.0239 | 0.0027 | -0.6251 | 0.0841 |
| rs13117856 | A | 0.0084 | 0.0015 | -0.0593 | 0.0489 |
| rs13130765 | C | -0.0091 | 0.0014 | 0.0644 | 0.0442 |
| rs13133213 | A | 0.0096 | 0.0014 | -0.0596 | 0.0441 |
| rs13145650 | T | -0.0173 | 0.0025 | -0.0322 | 0.0786 |
| rs13154429 | C | -0.0136 | 0.0023 | 0.0872 | 0.0704 |
| rs13163062 | T | 0.0115 | 0.0014 | -0.0055 | 0.0447 |
| rs13163845 | T | -0.0149 | 0.0020 | 0.1133 | 0.0630 |
| rs13168136 | A | -0.0092 | 0.0017 | 0.1000 | 0.0525 |
| rs13169187 | A | -0.0079 | 0.0014 | 0.1781 | 0.0443 |
| rs13177031 | A | -0.0085 | 0.0015 | 0.0255 | 0.0461 |
| rs13190235 | T | -0.0138 | 0.0024 | 0.1305 | 0.0760 |
| rs13197257 | T | 0.0109 | 0.0016 | -0.0353 | 0.0500 |
| rs1320139 | C | -0.0149 | 0.0014 | 0.0513 | 0.0446 |
| rs13212041 | T | 0.0101 | 0.0018 | -0.0382 | 0.0546 |
| rs13240401 | T | 0.0176 | 0.0017 | -0.0056 | 0.0530 |
| rs13246220 | T | -0.0093 | 0.0014 | -0.0518 | 0.0453 |
| rs13261773 | C | 0.0122 | 0.0019 | 0.0346 | 0.0619 |
| rs13266287 | A | -0.0085 | 0.0015 | 0.0308 | 0.0472 |
| rs13281564 | A | 0.0088 | 0.0014 | -0.0218 | 0.0443 |
| rs13284516 | C | -0.0082 | 0.0014 | 0.0070 | 0.0443 |
| rs1329044 | T | 0.0134 | 0.0019 | -0.0356 | 0.0579 |
| rs1329125 | T | -0.0097 | 0.0015 | 0.0761 | 0.0472 |
| rs13296345 | T | 0.0090 | 0.0015 | -0.1369 | 0.0460 |
| rs13318986 | A | 0.0124 | 0.0015 | 0.0393 | 0.0466 |
| rs13327482 | A | -0.0121 | 0.0018 | -0.0052 | 0.0584 |
| rs1334297 | A | 0.0257 | 0.0016 | 0.0467 | 0.0503 |
| rs1335482 | T | 0.0096 | 0.0014 | 0.0275 | 0.0443 |
| rs13361043 | T | -0.0158 | 0.0028 | -0.0030 | 0.0892 |
| rs13381557 | A | -0.0087 | 0.0014 | 0.1282 | 0.0445 |
| rs13388333 | A | 0.0085 | 0.0015 | -0.1161 | 0.0457 |
| rs13397529 | C | -0.0097 | 0.0017 | 0.0281 | 0.0522 |
| rs13398860 | A | 0.0086 | 0.0015 | -0.0060 | 0.0463 |
| rs13402497 | A | -0.0080 | 0.0014 | 0.0778 | 0.0442 |
| rs13422673 | T | -0.0110 | 0.0014 | 0.0927 | 0.0443 |
| rs13425585 | C | -0.0089 | 0.0014 | 0.0025 | 0.0443 |
| rs1368250 | T | 0.0250 | 0.0045 | 0.0328 | 0.1276 |
| rs137079 | T | 0.0128 | 0.0020 | 0.0188 | 0.0632 |
| rs137858393 | A | 0.0193 | 0.0029 | 0.2457 | 0.0886 |
| rs138096147 | A | 0.0234 | 0.0042 | 0.0837 | 0.1312 |
| rs138484388 | T | -0.0306 | 0.0051 | -0.2900 | 0.1504 |
| rs1391513 | T | -0.0095 | 0.0015 | 0.0262 | 0.0485 |
| rs139244147 | A | -0.0193 | 0.0030 | 0.1445 | 0.0973 |
| rs1392816 | T | 0.0082 | 0.0014 | 0.0162 | 0.0456 |
| rs139980871 | T | 0.0180 | 0.0029 | -0.1954 | 0.0888 |
| rs1404549 | A | -0.0089 | 0.0015 | -0.0131 | 0.0474 |
| rs1405876 | T | 0.0109 | 0.0015 | 0.0327 | 0.0461 |
| rs140711597 | C | 0.0366 | 0.0055 | -0.4791 | 0.1548 |
| rs1408284 | C | -0.0127 | 0.0020 | -0.0562 | 0.0646 |
| rs1408430 | C | 0.0096 | 0.0016 | 0.0129 | 0.0492 |
| rs141729694 | T | 0.0195 | 0.0026 | 0.2035 | 0.0855 |
| rs142014757 | A | 0.0102 | 0.0018 | 0.0216 | 0.0552 |
| rs1426619 | T | 0.0096 | 0.0014 | 0.0120 | 0.0447 |
| rs1427298 | T | 0.0086 | 0.0014 | -0.0731 | 0.0451 |
| rs142747148 | A | 0.0329 | 0.0052 | 0.1229 | 0.1674 |
| rs1427829 | A | -0.0083 | 0.0014 | -0.1141 | 0.0444 |
| rs143148393 | T | -0.0211 | 0.0039 | -0.0203 | 0.1316 |
| rs1434630 | T | -0.0137 | 0.0020 | 0.0530 | 0.0616 |
| rs143743568 | A | 0.0118 | 0.0020 | -0.0589 | 0.0616 |
| rs143812851 | A | -0.0113 | 0.0019 | 0.1158 | 0.0583 |
| rs1440930 | C | 0.0081 | 0.0014 | 0.0094 | 0.0446 |
| rs144336753 | A | 0.0388 | 0.0053 | -0.3430 | 0.1594 |
| rs1445591 | A | 0.0085 | 0.0015 | -0.0256 | 0.0486 |
| rs1464297 | T | -0.0109 | 0.0015 | -0.0096 | 0.0466 |
| rs1467737 | T | -0.0084 | 0.0014 | -0.0034 | 0.0446 |
| rs1485300 | C | 0.0098 | 0.0015 | 0.0626 | 0.0472 |
| rs1490612 | T | 0.0193 | 0.0026 | -0.1830 | 0.0840 |
| rs150252215 | A | 0.0204 | 0.0035 | -0.3593 | 0.1116 |
| rs150421637 | A | 0.0349 | 0.0061 | -0.3692 | 0.1823 |
| rs150537577 | A | -0.0151 | 0.0026 | 0.1759 | 0.0848 |
| rs1505676 | C | 0.0084 | 0.0015 | 0.1290 | 0.0456 |
| rs151381 | T | -0.0087 | 0.0014 | -0.0257 | 0.0441 |
| rs1518890 | T | -0.0090 | 0.0016 | 0.1485 | 0.0503 |
| rs1527878 | A | -0.0105 | 0.0016 | 0.1536 | 0.0517 |
| rs1529597 | A | 0.0223 | 0.0038 | 0.0154 | 0.1304 |
| rs1538389 | T | -0.0109 | 0.0018 | -0.0022 | 0.0559 |
| rs1542354 | A | -0.0084 | 0.0014 | -0.0718 | 0.0446 |
| rs1544 | A | -0.0096 | 0.0016 | 0.0688 | 0.0501 |
| rs1554798 | A | -0.0080 | 0.0014 | -0.0072 | 0.0454 |
| rs1564347 | T | 0.0088 | 0.0015 | 0.0413 | 0.0466 |
| rs1569092 | A | 0.0160 | 0.0020 | -0.0940 | 0.0608 |
| rs1593022 | T | -0.0099 | 0.0017 | -0.0322 | 0.0519 |
| rs1603460 | T | 0.0090 | 0.0014 | -0.1323 | 0.0448 |
| rs162445 | A | 0.0152 | 0.0026 | 0.0697 | 0.0832 |
| rs163229 | C | -0.0260 | 0.0047 | 0.0712 | 0.1483 |
| rs1637770 | T | 0.0191 | 0.0028 | -0.1866 | 0.0919 |
| rs164938 | T | -0.0088 | 0.0014 | 0.0787 | 0.0453 |
| rs165633 | A | -0.0116 | 0.0016 | -0.0467 | 0.0531 |
| rs1656614 | C | 0.0122 | 0.0015 | -0.1916 | 0.0482 |
| rs1671269 | T | -0.0100 | 0.0016 | -0.0102 | 0.0523 |
| rs1671770 | A | 0.0125 | 0.0018 | -0.0571 | 0.0589 |
| rs16851779 | T | -0.0153 | 0.0027 | -0.0166 | 0.0804 |
| rs16871807 | T | 0.0090 | 0.0015 | -0.0100 | 0.0467 |
| rs1689510 | C | 0.0194 | 0.0015 | -0.1921 | 0.0466 |
| rs16901689 | T | 0.0111 | 0.0019 | -0.1317 | 0.0597 |
| rs1693584 | T | -0.0087 | 0.0015 | 0.1117 | 0.0479 |
| rs16958559 | T | 0.0096 | 0.0017 | -0.1098 | 0.0528 |
| rs16966271 | T | 0.0092 | 0.0016 | -0.0971 | 0.0494 |
| rs16975275 | A | -0.0102 | 0.0017 | -0.0081 | 0.0523 |
| rs17048801 | A | -0.0106 | 0.0014 | -0.0080 | 0.0461 |
| rs17048855 | A | 0.0108 | 0.0015 | -0.0095 | 0.0463 |
| rs17069646 | T | 0.0085 | 0.0015 | -0.0231 | 0.0478 |
| rs17110109 | T | -0.0093 | 0.0014 | 0.0493 | 0.0453 |
| rs17113730 | A | -0.0170 | 0.0027 | -0.0737 | 0.0869 |
| rs17131123 | A | -0.0243 | 0.0039 | 0.0299 | 0.1123 |
| rs17133297 | A | -0.0146 | 0.0023 | 0.0811 | 0.0761 |
| rs17144467 | A | 0.0098 | 0.0015 | -0.0951 | 0.0476 |
| rs17148998 | A | 0.0101 | 0.0017 | -0.1533 | 0.0545 |
| rs171697 | C | 0.0133 | 0.0015 | 0.1485 | 0.0470 |
| rs17170519 | C | 0.0132 | 0.0023 | 0.0051 | 0.0719 |
| rs1717204 | A | -0.0117 | 0.0018 | 0.0545 | 0.0567 |
| rs1718188 | A | 0.0096 | 0.0014 | 0.1476 | 0.0446 |
| rs17186106 | T | 0.0108 | 0.0018 | -0.1258 | 0.0580 |
| rs17190418 | T | -0.0213 | 0.0031 | 0.0034 | 0.0938 |
| rs17205908 | T | -0.0092 | 0.0015 | -0.0103 | 0.0465 |
| rs17248751 | A | -0.0136 | 0.0017 | 0.0791 | 0.0528 |
| rs1728118 | A | 0.0089 | 0.0016 | -0.0617 | 0.0496 |
| rs1729412 | T | -0.0107 | 0.0014 | 0.1045 | 0.0448 |
| rs173003 | A | -0.0082 | 0.0014 | -0.0291 | 0.0443 |
| rs1734370 | A | 0.0099 | 0.0016 | -0.0226 | 0.0486 |
| rs1738050 | C | -0.0093 | 0.0014 | -0.0241 | 0.0454 |
| rs17411339 | A | 0.0140 | 0.0014 | -0.0735 | 0.0459 |
| rs17428076 | C | 0.0137 | 0.0016 | -0.0841 | 0.0515 |
| rs17440885 | A | 0.0160 | 0.0029 | 0.0289 | 0.0907 |
| rs1747714 | T | 0.0106 | 0.0014 | 0.0123 | 0.0444 |
| rs1747817 | T | -0.0092 | 0.0016 | -0.0563 | 0.0514 |
| rs17489649 | A | 0.0127 | 0.0015 | -0.0135 | 0.0468 |
| rs17502934 | T | -0.0167 | 0.0020 | 0.0277 | 0.0631 |
| rs175325 | A | -0.0100 | 0.0014 | -0.0084 | 0.0454 |
| rs17536059 | C | -0.0108 | 0.0019 | 0.1281 | 0.0578 |
| rs17551064 | A | 0.0141 | 0.0019 | -0.0641 | 0.0596 |
| rs17563464 | A | -0.0147 | 0.0017 | 0.1319 | 0.0538 |
| rs17565975 | A | -0.0106 | 0.0014 | -0.1532 | 0.0448 |
| rs17568389 | A | 0.0127 | 0.0014 | -0.0617 | 0.0443 |
| rs17574007 | A | -0.0129 | 0.0021 | 0.0481 | 0.0685 |
| rs1758747 | A | 0.0092 | 0.0015 | -0.0602 | 0.0479 |
| rs17598675 | T | -0.0116 | 0.0014 | 0.0829 | 0.0445 |
| rs17604349 | A | 0.0127 | 0.0018 | -0.1064 | 0.0581 |
| rs17609255 | T | -0.0087 | 0.0014 | 0.1060 | 0.0448 |
| rs176218 | T | 0.0200 | 0.0018 | 0.0685 | 0.0566 |
| rs17622379 | T | -0.0113 | 0.0018 | 0.0490 | 0.0562 |
| rs17650634 | T | -0.0104 | 0.0019 | -0.0544 | 0.0587 |
| rs17667540 | A | -0.0087 | 0.0015 | 0.0816 | 0.0481 |
| rs17669337 | T | -0.0110 | 0.0014 | -0.0272 | 0.0447 |
| rs17680712 | T | -0.0092 | 0.0014 | 0.0783 | 0.0446 |
| rs17686649 | T | -0.0098 | 0.0017 | -0.0879 | 0.0537 |
| rs17732878 | T | -0.0095 | 0.0017 | 0.1779 | 0.0539 |
| rs17742342 | A | -0.0119 | 0.0018 | 0.2674 | 0.0553 |
| rs17747544 | A | -0.0090 | 0.0014 | 0.1002 | 0.0442 |
| rs1779549 | A | 0.0081 | 0.0014 | -0.0334 | 0.0449 |
| rs178183 | T | 0.0145 | 0.0016 | -0.0815 | 0.0511 |
| rs17882802 | A | 0.0084 | 0.0014 | 0.0440 | 0.0444 |
| rs17883331 | A | -0.0108 | 0.0018 | 0.1992 | 0.0556 |
| rs1792602 | A | -0.0097 | 0.0014 | 0.0117 | 0.0450 |
| rs181214 | T | -0.0104 | 0.0017 | 0.1669 | 0.0543 |
| rs182355396 | A | -0.0525 | 0.0084 | -0.2252 | 0.2589 |
| rs182902112 | A | -0.0300 | 0.0053 | 0.0453 | 0.1663 |
| rs1835339 | T | 0.0084 | 0.0014 | 0.0346 | 0.0455 |
| rs183869217 | C | 0.0174 | 0.0030 | 0.0262 | 0.0876 |
| rs1841023 | A | -0.0092 | 0.0015 | 0.0085 | 0.0478 |
| rs1842713 | A | -0.0117 | 0.0017 | 0.0034 | 0.0540 |
| rs1843815 | A | 0.0077 | 0.0014 | 0.0595 | 0.0443 |
| rs1861786 | A | -0.0081 | 0.0014 | 0.0264 | 0.0454 |
| rs186456786 | A | 0.0455 | 0.0079 | -0.2867 | 0.2313 |
| rs1865955 | T | 0.0122 | 0.0019 | 0.0053 | 0.0582 |
| rs1866823 | A | 0.0107 | 0.0014 | -0.0744 | 0.0448 |
| rs187580 | T | -0.0104 | 0.0017 | 0.0553 | 0.0518 |
| rs187951956 | A | -0.0315 | 0.0058 | 0.1045 | 0.1952 |
| rs1880088 | A | -0.0097 | 0.0016 | -0.0196 | 0.0509 |
| rs1880692 | A | 0.0086 | 0.0014 | -0.0584 | 0.0444 |
| rs188251563 | A | 0.0428 | 0.0070 | 0.2065 | 0.2256 |
| rs1890132 | T | -0.0089 | 0.0015 | -0.0717 | 0.0484 |
| rs1898111 | A | 0.0101 | 0.0018 | -0.0572 | 0.0581 |
| rs190102446 | T | -0.0210 | 0.0037 | -0.1061 | 0.1211 |
| rs1905616 | A | 0.0083 | 0.0015 | 0.1076 | 0.0470 |
| rs1910005 | T | -0.0090 | 0.0015 | -0.0481 | 0.0486 |
| rs1918394 | T | 0.0119 | 0.0019 | 0.0513 | 0.0596 |
| rs191903670 | A | 0.0335 | 0.0061 | -0.1809 | 0.2023 |
| rs192436652 | T | -0.0319 | 0.0045 | 0.0699 | 0.1355 |
| rs1933264 | T | 0.0100 | 0.0016 | 0.1101 | 0.0513 |
| rs1952183 | A | -0.0089 | 0.0014 | -0.0619 | 0.0442 |
| rs1955250 | A | -0.0140 | 0.0025 | 0.0678 | 0.0806 |
| rs1963381 | A | 0.0099 | 0.0016 | -0.1165 | 0.0517 |
| rs1980129 | A | 0.0085 | 0.0014 | -0.0041 | 0.0442 |
| rs198262 | T | -0.0223 | 0.0035 | 0.1961 | 0.1097 |
| rs1991585 | T | -0.0101 | 0.0015 | 0.1295 | 0.0485 |
| rs1995181 | A | -0.0078 | 0.0014 | -0.1169 | 0.0443 |
| rs2007655 | T | 0.0086 | 0.0014 | -0.0387 | 0.0442 |
| rs2011603 | A | -0.0091 | 0.0016 | 0.3345 | 0.0502 |
| rs2014830 | T | 0.0110 | 0.0015 | -0.2347 | 0.0484 |
| rs2023016 | C | 0.0107 | 0.0017 | -0.0468 | 0.0538 |
| rs2024568 | T | -0.0097 | 0.0016 | -0.0695 | 0.0515 |
| rs2029401 | A | -0.0091 | 0.0014 | 0.0047 | 0.0450 |
| rs2034631 | T | 0.0089 | 0.0015 | 0.0853 | 0.0465 |
| rs2039204 | A | 0.0082 | 0.0014 | -0.1387 | 0.0443 |
| rs2043187 | A | 0.0107 | 0.0015 | 0.0044 | 0.0489 |
| rs2052285 | A | 0.0112 | 0.0014 | -0.0030 | 0.0453 |
| rs2055940 | A | 0.0082 | 0.0015 | -0.0353 | 0.0472 |
| rs2077235 | T | 0.0097 | 0.0017 | 0.0124 | 0.0522 |
| rs2081652 | A | 0.0123 | 0.0015 | -0.0670 | 0.0466 |
| rs2082317 | T | -0.0078 | 0.0014 | 0.0101 | 0.0451 |
| rs2088913 | A | 0.0081 | 0.0014 | 0.0224 | 0.0447 |
| rs2092248 | T | -0.0083 | 0.0015 | -0.0471 | 0.0484 |
| rs2098526 | A | -0.0241 | 0.0043 | 0.0717 | 0.1352 |
| rs2100249 | T | -0.0089 | 0.0015 | 0.0301 | 0.0468 |
| rs2126069 | T | 0.0080 | 0.0014 | 0.0502 | 0.0445 |
| rs2131167 | A | 0.0080 | 0.0014 | -0.0110 | 0.0445 |
| rs214626 | A | 0.0101 | 0.0018 | 0.0448 | 0.0571 |
| rs2160514 | A | -0.0094 | 0.0014 | -0.0090 | 0.0452 |
| rs2179152 | T | -0.0131 | 0.0015 | -0.1236 | 0.0455 |
| rs2183271 | T | 0.0084 | 0.0015 | -0.1625 | 0.0461 |
| rs2199409 | T | 0.0103 | 0.0017 | -0.0017 | 0.0552 |
| rs2212430 | T | -0.0100 | 0.0016 | 0.0134 | 0.0484 |
| rs2216144 | T | 0.0098 | 0.0014 | -0.0167 | 0.0444 |
| rs2220926 | T | -0.0095 | 0.0014 | -0.0636 | 0.0448 |
| rs2250660 | C | 0.0080 | 0.0014 | 0.0227 | 0.0442 |
| rs2252098 | T | -0.0077 | 0.0014 | -0.0695 | 0.0443 |
| rs2254681 | A | 0.0101 | 0.0016 | 0.0864 | 0.0505 |
| rs2256965 | A | 0.0106 | 0.0014 | -0.1265 | 0.0449 |
| rs2276209 | A | 0.0085 | 0.0015 | 0.0562 | 0.0488 |
| rs2283076 | A | 0.0115 | 0.0017 | -0.0077 | 0.0527 |
| rs2287838 | A | -0.0104 | 0.0014 | -0.0413 | 0.0445 |
| rs2290601 | T | -0.0149 | 0.0017 | 0.0510 | 0.0525 |
| rs2297293 | C | 0.0099 | 0.0015 | -0.0733 | 0.0481 |
| rs2299156 | T | 0.0099 | 0.0018 | -0.0999 | 0.0584 |
| rs2314338 | T | 0.0093 | 0.0016 | -0.0152 | 0.0498 |
| rs2321157 | A | -0.0081 | 0.0014 | -0.0933 | 0.0445 |
| rs232496 | T | 0.0090 | 0.0015 | 0.0862 | 0.0462 |
| rs2332179 | A | 0.0121 | 0.0020 | 0.1331 | 0.0652 |
| rs2336721 | T | 0.0095 | 0.0015 | -0.1737 | 0.0469 |
| rs2358628 | A | -0.0091 | 0.0015 | -0.0249 | 0.0478 |
| rs236318 | C | -0.0102 | 0.0017 | 0.0095 | 0.0522 |
| rs2364544 | A | -0.0105 | 0.0014 | -0.0021 | 0.0454 |
| rs2365376 | A | 0.0092 | 0.0015 | -0.0685 | 0.0471 |
| rs2368831 | T | -0.0092 | 0.0014 | 0.1108 | 0.0451 |
| rs2373124 | A | -0.0096 | 0.0016 | -0.1531 | 0.0520 |
| rs2406253 | A | 0.0135 | 0.0018 | -0.1352 | 0.0556 |
| rs2414072 | A | -0.0083 | 0.0014 | -0.2248 | 0.0445 |
| rs2416214 | A | 0.0079 | 0.0014 | 0.0060 | 0.0447 |
| rs2416759 | A | -0.0093 | 0.0015 | -0.1499 | 0.0483 |
| rs2416845 | T | 0.0263 | 0.0048 | -0.1896 | 0.1529 |
| rs2431023 | A | 0.0093 | 0.0014 | 0.0408 | 0.0447 |
| rs2434672 | A | 0.0091 | 0.0014 | -0.0018 | 0.0448 |
| rs2436760 | T | 0.0232 | 0.0042 | -0.0837 | 0.1271 |
| rs2447097 | T | 0.0096 | 0.0014 | 0.0345 | 0.0446 |
| rs2458370 | T | -0.0086 | 0.0015 | 0.1403 | 0.0463 |
| rs2469226 | A | 0.0096 | 0.0017 | 0.0369 | 0.0531 |
| rs2470966 | C | -0.0103 | 0.0015 | 0.0683 | 0.0479 |
| rs2478208 | C | -0.0103 | 0.0014 | 0.0086 | 0.0443 |
| rs2496482 | T | 0.0121 | 0.0015 | -0.0429 | 0.0452 |
| rs2517086 | C | -0.0088 | 0.0014 | 0.0061 | 0.0450 |
| rs2521602 | A | -0.0266 | 0.0049 | -0.1953 | 0.1509 |
| rs2522545 | T | -0.0083 | 0.0014 | 0.1148 | 0.0450 |
| rs2526398 | C | -0.0222 | 0.0015 | 0.0856 | 0.0462 |
| rs2529069 | T | -0.0106 | 0.0016 | 0.0820 | 0.0509 |
| rs252991 | A | 0.0095 | 0.0015 | -0.0703 | 0.0458 |
| rs2542673 | A | 0.0088 | 0.0015 | 0.0134 | 0.0472 |
| rs2545795 | A | 0.0125 | 0.0014 | 0.0245 | 0.0446 |
| rs255053 | A | -0.0111 | 0.0018 | -0.0681 | 0.0565 |
| rs2554835 | A | 0.0088 | 0.0014 | -0.0193 | 0.0453 |
| rs2568955 | T | -0.0165 | 0.0016 | 0.0351 | 0.0534 |
| rs2569041 | T | 0.0077 | 0.0014 | -0.0984 | 0.0451 |
| rs2570497 | T | -0.0121 | 0.0015 | -0.0756 | 0.0462 |
| rs2588959 | T | -0.0080 | 0.0014 | 0.0466 | 0.0442 |
| rs2657283 | T | 0.0085 | 0.0014 | -0.1156 | 0.0449 |
| rs2665668 | A | -0.0094 | 0.0015 | -0.0426 | 0.0466 |
| rs2702576 | A | -0.0090 | 0.0014 | 0.1262 | 0.0456 |
| rs2718277 | T | 0.0142 | 0.0026 | -0.1554 | 0.0782 |
| rs2718791 | T | -0.0081 | 0.0015 | -0.0372 | 0.0462 |
| rs27220 | A | -0.0114 | 0.0015 | 0.0181 | 0.0460 |
| rs2725370 | T | -0.0141 | 0.0015 | 0.1240 | 0.0482 |
| rs2736752 | T | 0.0095 | 0.0017 | -0.1088 | 0.0542 |
| rs2740795 | A | 0.0098 | 0.0016 | 0.1492 | 0.0503 |
| rs2761438 | A | 0.0098 | 0.0015 | -0.0167 | 0.0455 |
| rs2764684 | T | 0.0143 | 0.0019 | -0.0428 | 0.0577 |
| rs2805064 | C | 0.0093 | 0.0016 | 0.0227 | 0.0493 |
| rs2838006 | T | 0.0132 | 0.0015 | -0.0361 | 0.0463 |
| rs28482086 | A | 0.0086 | 0.0015 | -0.0731 | 0.0467 |
| rs28505285 | C | 0.0106 | 0.0018 | -0.0099 | 0.0573 |
| rs28512462 | C | 0.0112 | 0.0014 | -0.2510 | 0.0458 |
| rs28513882 | A | -0.0110 | 0.0018 | -0.0768 | 0.0570 |
| rs2852349 | T | -0.0093 | 0.0014 | -0.0042 | 0.0444 |
| rs28587776 | T | 0.0088 | 0.0015 | -0.1027 | 0.0460 |
| rs28669886 | A | -0.0084 | 0.0015 | -0.0345 | 0.0463 |
| rs2870281 | A | -0.0083 | 0.0014 | 0.0529 | 0.0445 |
| rs28735993 | A | 0.0162 | 0.0022 | -0.0615 | 0.0690 |
| rs2885198 | A | 0.0088 | 0.0014 | -0.0557 | 0.0446 |
| rs2898191 | A | 0.0093 | 0.0015 | -0.1006 | 0.0476 |
| rs2910823 | T | 0.0115 | 0.0014 | -0.0744 | 0.0449 |
| rs2916490 | A | -0.0092 | 0.0015 | 0.1374 | 0.0487 |
| rs2923424 | A | 0.0120 | 0.0014 | -0.1555 | 0.0452 |
| rs2926702 | T | -0.0124 | 0.0021 | 0.1792 | 0.0667 |
| rs2929032 | A | -0.0081 | 0.0014 | 0.0505 | 0.0443 |
| rs2929860 | T | 0.0110 | 0.0018 | 0.0285 | 0.0556 |
| rs2942884 | A | -0.0090 | 0.0014 | 0.1015 | 0.0444 |
| rs2954114 | A | -0.0081 | 0.0014 | 0.0047 | 0.0444 |
| rs2958182 | A | 0.0082 | 0.0015 | 0.0301 | 0.0470 |
| rs2962378 | A | 0.0101 | 0.0017 | -0.0780 | 0.0536 |
| rs2964199 | T | -0.0099 | 0.0015 | -0.0175 | 0.0479 |
| rs2964255 | A | 0.0088 | 0.0015 | 0.1047 | 0.0479 |
| rs2976397 | T | 0.0081 | 0.0014 | 0.0158 | 0.0448 |
| rs2977464 | T | 0.0102 | 0.0018 | -0.1001 | 0.0567 |
| rs29792 | A | -0.0093 | 0.0015 | 0.1476 | 0.0482 |
| rs2980813 | A | 0.0088 | 0.0014 | -0.0822 | 0.0445 |
| rs2989476 | C | 0.0078 | 0.0014 | 0.0127 | 0.0446 |
| rs2989751 | A | -0.0092 | 0.0017 | -0.0121 | 0.0521 |
| rs2992037 | A | 0.0149 | 0.0015 | 0.1665 | 0.0478 |
| rs2994326 | T | -0.0100 | 0.0018 | -0.0758 | 0.0572 |
| rs2998299 | T | -0.0136 | 0.0017 | -0.0206 | 0.0542 |
| rs3026996 | A | 0.0138 | 0.0016 | 0.1277 | 0.0515 |
| rs303752 | A | -0.0110 | 0.0014 | 0.1068 | 0.0455 |
| rs3111251 | T | -0.0090 | 0.0014 | -0.0350 | 0.0447 |
| rs312927 | A | -0.0147 | 0.0022 | -0.0902 | 0.0688 |
| rs312945 | A | -0.0091 | 0.0015 | -0.0811 | 0.0483 |
| rs317050 | T | 0.0078 | 0.0014 | -0.0294 | 0.0442 |
| rs322614 | A | -0.0087 | 0.0014 | 0.0037 | 0.0454 |
| rs322744 | T | -0.0090 | 0.0016 | 0.0415 | 0.0502 |
| rs324885 | A | 0.0142 | 0.0014 | -0.0298 | 0.0443 |
| rs32940 | T | -0.0096 | 0.0015 | 0.0790 | 0.0483 |
| rs339057 | A | -0.0081 | 0.0014 | 0.0940 | 0.0447 |
| rs34067381 | T | -0.0104 | 0.0015 | -0.0154 | 0.0463 |
| rs34098770 | A | -0.0160 | 0.0020 | 0.1059 | 0.0615 |
| rs34106693 | C | 0.0159 | 0.0019 | -0.0557 | 0.0599 |
| rs34122915 | C | 0.0084 | 0.0014 | -0.0580 | 0.0457 |
| rs341504 | A | 0.0099 | 0.0015 | 0.0477 | 0.0462 |
| rs34155847 | A | 0.0135 | 0.0016 | -0.1774 | 0.0516 |
| rs34262657 | T | -0.0240 | 0.0038 | -0.1359 | 0.1274 |
| rs34286836 | T | -0.0089 | 0.0014 | -0.1135 | 0.0446 |
| rs34298584 | A | 0.0111 | 0.0019 | -0.0748 | 0.0590 |
| rs34305371 | A | 0.0314 | 0.0024 | -0.1364 | 0.0730 |
| rs34309 | A | 0.0080 | 0.0014 | -0.0102 | 0.0457 |
| rs34316274 | A | 0.0097 | 0.0017 | 0.1355 | 0.0517 |
| rs34363861 | A | 0.0087 | 0.0014 | -0.2093 | 0.0453 |
| rs34394051 | A | -0.0117 | 0.0020 | 0.1590 | 0.0606 |
| rs34410 | C | -0.0083 | 0.0014 | 0.0875 | 0.0443 |
| rs34624793 | T | 0.0116 | 0.0021 | -0.0045 | 0.0641 |
| rs34720381 | T | -0.0180 | 0.0024 | 0.2131 | 0.0764 |
| rs34748029 | A | 0.0185 | 0.0025 | -0.0060 | 0.0768 |
| rs347661 | T | -0.0081 | 0.0014 | 0.1000 | 0.0450 |
| rs34780702 | A | 0.0111 | 0.0017 | -0.0300 | 0.0518 |
| rs34807077 | A | 0.0128 | 0.0019 | -0.0460 | 0.0598 |
| rs34811474 | A | 0.0104 | 0.0017 | -0.2049 | 0.0523 |
| rs34967082 | A | -0.0080 | 0.0014 | 0.1451 | 0.0449 |
| rs35016816 | T | 0.0176 | 0.0023 | 0.1088 | 0.0741 |
| rs350281 | T | -0.0117 | 0.0015 | 0.0945 | 0.0471 |
| rs35104491 | A | 0.0124 | 0.0018 | 0.0190 | 0.0564 |
| rs35111506 | C | -0.0175 | 0.0030 | 0.1860 | 0.0954 |
| rs35192107 | T | -0.0091 | 0.0017 | 0.0509 | 0.0518 |
| rs35319653 | T | 0.0119 | 0.0015 | -0.1378 | 0.0465 |
| rs35417702 | T | -0.0152 | 0.0014 | 0.0479 | 0.0442 |
| rs35493937 | C | 0.0124 | 0.0017 | 0.0426 | 0.0530 |
| rs35606437 | A | 0.0105 | 0.0016 | -0.0005 | 0.0501 |
| rs356999 | A | -0.0101 | 0.0014 | -0.0168 | 0.0456 |
| rs35745455 | A | 0.0080 | 0.0014 | -0.0311 | 0.0441 |
| rs35751693 | T | -0.0246 | 0.0040 | 0.4313 | 0.1167 |
| rs35754740 | T | 0.0083 | 0.0014 | -0.0872 | 0.0449 |
| rs35811586 | T | 0.0217 | 0.0031 | 0.0811 | 0.0951 |
| rs35929923 | A | -0.0114 | 0.0016 | 0.0499 | 0.0508 |
| rs36085856 | C | 0.0201 | 0.0036 | -0.1558 | 0.0882 |
| rs36120534 | T | 0.0100 | 0.0018 | -0.0322 | 0.0567 |
| rs362307 | T | -0.0226 | 0.0027 | 0.2642 | 0.0843 |
| rs363096 | T | -0.0143 | 0.0014 | 0.1230 | 0.0447 |
| rs3735478 | T | 0.0108 | 0.0016 | -0.1233 | 0.0486 |
| rs3751331 | A | -0.0097 | 0.0014 | -0.0574 | 0.0457 |
| rs3766979 | A | 0.0085 | 0.0015 | -0.0932 | 0.0471 |
| rs3768480 | C | 0.0101 | 0.0014 | -0.1056 | 0.0443 |
| rs3781339 | T | -0.0104 | 0.0018 | -0.0337 | 0.0559 |
| rs3790609 | T | 0.0107 | 0.0019 | 0.1439 | 0.0576 |
| rs3809169 | T | 0.0140 | 0.0025 | 0.1836 | 0.0775 |
| rs3809634 | A | -0.0109 | 0.0015 | 0.0637 | 0.0480 |
| rs3812281 | T | 0.0113 | 0.0014 | -0.0564 | 0.0451 |
| rs3817923 | A | -0.0121 | 0.0022 | 0.0926 | 0.0690 |
| rs382196 | T | -0.0082 | 0.0014 | -0.0421 | 0.0455 |
| rs3847228 | T | 0.0083 | 0.0014 | 0.0049 | 0.0451 |
| rs3848715 | A | -0.0080 | 0.0014 | 0.0596 | 0.0443 |
| rs3859523 | T | 0.0124 | 0.0018 | -0.0850 | 0.0575 |
| rs387027 | A | -0.0083 | 0.0015 | 0.1790 | 0.0464 |
| rs3895736 | A | 0.0144 | 0.0019 | 0.1848 | 0.0578 |
| rs3897821 | A | 0.0154 | 0.0015 | -0.0647 | 0.0468 |
| rs3948495 | T | -0.0089 | 0.0014 | 0.0485 | 0.0453 |
| rs399821 | A | -0.0078 | 0.0014 | 0.0034 | 0.0446 |
| rs39998 | A | 0.0090 | 0.0015 | 0.0424 | 0.0489 |
| rs401526 | T | -0.0104 | 0.0014 | -0.0131 | 0.0443 |
| rs401966 | C | -0.0097 | 0.0014 | 0.0389 | 0.0456 |
| rs4127499 | A | 0.0087 | 0.0015 | 0.0270 | 0.0460 |
| rs41282553 | A | 0.0237 | 0.0040 | -0.0504 | 0.1250 |
| rs42210 | C | -0.0098 | 0.0016 | -0.0054 | 0.0490 |
| rs42302 | A | 0.0086 | 0.0015 | -0.0562 | 0.0467 |
| rs4255791 | A | -0.0084 | 0.0015 | 0.0733 | 0.0469 |
| rs4263475 | A | 0.0078 | 0.0014 | -0.0777 | 0.0454 |
| rs4283754 | A | -0.0084 | 0.0014 | -0.0218 | 0.0468 |
| rs429150 | T | 0.0085 | 0.0014 | -0.2606 | 0.0444 |
| rs4298514 | T | 0.0104 | 0.0015 | -0.0023 | 0.0483 |
| rs4320563 | A | 0.0109 | 0.0014 | -0.0416 | 0.0443 |
| rs4328757 | T | 0.0096 | 0.0014 | -0.0402 | 0.0454 |
| rs4352658 | T | -0.0190 | 0.0026 | 0.0531 | 0.0809 |
| rs4358081 | A | -0.0095 | 0.0014 | -0.1201 | 0.0442 |
| rs4358358 | A | 0.0109 | 0.0015 | -0.0341 | 0.0480 |
| rs4369924 | A | 0.0125 | 0.0019 | -0.0931 | 0.0606 |
| rs4384309 | A | 0.0102 | 0.0014 | 0.0009 | 0.0446 |
| rs4396896 | A | -0.0107 | 0.0014 | 0.0005 | 0.0453 |
| rs4423373 | A | -0.0087 | 0.0014 | 0.0315 | 0.0445 |
| rs4426420 | T | -0.0140 | 0.0019 | 0.0897 | 0.0606 |
| rs4434676 | A | -0.0089 | 0.0014 | -0.0454 | 0.0446 |
| rs4458044 | C | 0.0101 | 0.0016 | 0.1886 | 0.0514 |
| rs4467547 | T | 0.0121 | 0.0014 | 0.0247 | 0.0449 |
| rs4469771 | T | 0.0101 | 0.0015 | -0.0132 | 0.0472 |
| rs4480339 | A | -0.0097 | 0.0015 | -0.0546 | 0.0477 |
| rs4490539 | A | 0.0142 | 0.0015 | 0.0726 | 0.0480 |
| rs4497562 | A | 0.0112 | 0.0016 | -0.0503 | 0.0508 |
| rs4500930 | T | -0.0109 | 0.0015 | 0.0721 | 0.0464 |
| rs4500960 | T | -0.0131 | 0.0014 | 0.0409 | 0.0443 |
| rs4502401 | T | 0.0085 | 0.0015 | -0.0434 | 0.0468 |
| rs4652135 | A | 0.0103 | 0.0016 | 0.0711 | 0.0497 |
| rs4658019 | T | 0.0084 | 0.0014 | -0.1452 | 0.0443 |
| rs4663617 | A | 0.0104 | 0.0017 | -0.1114 | 0.0529 |
| rs4664983 | T | -0.0103 | 0.0018 | 0.1239 | 0.0558 |
| rs4673840 | T | -0.0138 | 0.0019 | 0.1415 | 0.0602 |
| rs4675248 | A | -0.0094 | 0.0014 | -0.0281 | 0.0451 |
| rs4685405 | T | -0.0111 | 0.0018 | 0.0367 | 0.0587 |
| rs4687735 | T | -0.0200 | 0.0033 | 0.0345 | 0.1013 |
| rs4691601 | A | -0.0115 | 0.0014 | -0.0028 | 0.0443 |
| rs4705763 | C | 0.0089 | 0.0014 | -0.1175 | 0.0443 |
| rs4712371 | A | -0.0124 | 0.0021 | 0.0376 | 0.0678 |
| rs4719460 | T | -0.0088 | 0.0015 | 0.0872 | 0.0467 |
| rs4719944 | T | -0.0121 | 0.0014 | 0.0248 | 0.0445 |
| rs4724083 | T | 0.0083 | 0.0015 | -0.0377 | 0.0479 |
| rs4726070 | A | 0.0122 | 0.0014 | -0.0637 | 0.0452 |
| rs4730020 | T | 0.0090 | 0.0016 | -0.0630 | 0.0500 |
| rs4731413 | A | 0.0111 | 0.0017 | -0.0913 | 0.0543 |
| rs4735297 | A | -0.0086 | 0.0015 | -0.1124 | 0.0475 |
| rs4739235 | A | 0.0109 | 0.0019 | -0.0171 | 0.0596 |
| rs4741571 | A | -0.0102 | 0.0015 | -0.0956 | 0.0477 |
| rs4757957 | C | 0.0133 | 0.0015 | 0.1122 | 0.0474 |
| rs4766424 | C | 0.0142 | 0.0021 | -0.0325 | 0.0662 |
| rs4778058 | T | -0.0092 | 0.0014 | -0.0370 | 0.0449 |
| rs4785187 | A | -0.0100 | 0.0017 | -0.0842 | 0.0532 |
| rs4787028 | T | 0.0086 | 0.0015 | -0.0766 | 0.0467 |
| rs4787457 | A | 0.0162 | 0.0015 | -0.1380 | 0.0456 |
| rs4788115 | A | 0.0111 | 0.0019 | 0.0085 | 0.0592 |
| rs479018 | A | 0.0109 | 0.0015 | -0.0189 | 0.0476 |
| rs4793090 | A | 0.0084 | 0.0015 | 0.0353 | 0.0466 |
| rs4810894 | A | -0.0085 | 0.0015 | 0.0976 | 0.0459 |
| rs4812697 | T | 0.0171 | 0.0027 | 0.1107 | 0.0849 |
| rs481940 | T | 0.0100 | 0.0016 | -0.0488 | 0.0500 |
| rs482787 | T | 0.0100 | 0.0015 | -0.0519 | 0.0471 |
| rs4839155 | T | 0.0115 | 0.0017 | -0.0093 | 0.0519 |
| rs4846010 | A | 0.0099 | 0.0018 | -0.1588 | 0.0554 |
| rs4846724 | A | 0.0098 | 0.0014 | -0.0470 | 0.0444 |
| rs4848924 | A | -0.0117 | 0.0015 | 0.0286 | 0.0492 |
| rs4850954 | T | 0.0077 | 0.0014 | -0.0052 | 0.0446 |
| rs4851263 | A | -0.0151 | 0.0023 | 0.0596 | 0.0725 |
| rs4877516 | A | -0.0101 | 0.0014 | 0.0003 | 0.0445 |
| rs4881269 | A | 0.0100 | 0.0014 | -0.0072 | 0.0456 |
| rs488476 | C | -0.0113 | 0.0015 | 0.0945 | 0.0455 |
| rs4894658 | C | 0.0091 | 0.0016 | -0.0806 | 0.0506 |
| rs4895650 | T | 0.0093 | 0.0014 | -0.0792 | 0.0449 |
| rs4899012 | C | -0.0103 | 0.0014 | -0.0950 | 0.0453 |
| rs4904523 | A | -0.0082 | 0.0014 | -0.0018 | 0.0442 |
| rs4915735 | A | 0.0116 | 0.0020 | -0.0579 | 0.0632 |
| rs4919624 | A | 0.0193 | 0.0018 | -0.1195 | 0.0546 |
| rs4925065 | T | -0.0078 | 0.0014 | 0.2212 | 0.0443 |
| rs4925109 | A | -0.0096 | 0.0015 | 0.1350 | 0.0483 |
| rs4938815 | T | 0.0085 | 0.0015 | -0.1291 | 0.0483 |
| rs4941735 | T | 0.0092 | 0.0014 | -0.1394 | 0.0447 |
| rs4972748 | T | 0.0105 | 0.0018 | -0.1418 | 0.0574 |
| rs4977885 | A | -0.0084 | 0.0014 | -0.0047 | 0.0451 |
| rs4981245 | A | 0.0080 | 0.0014 | -0.1122 | 0.0447 |
| rs4984541 | A | -0.0138 | 0.0017 | 0.0941 | 0.0528 |
| rs4984613 | T | 0.0174 | 0.0028 | 0.0119 | 0.0916 |
| rs4984682 | C | -0.0138 | 0.0017 | -0.0256 | 0.0532 |
| rs548897 | A | 0.0081 | 0.0014 | -0.0712 | 0.0445 |
| rs55675587 | T | -0.0103 | 0.0018 | 0.0258 | 0.0563 |
| rs55736314 | C | -0.0154 | 0.0014 | 0.0776 | 0.0452 |
| rs55826493 | T | -0.0127 | 0.0021 | -0.0440 | 0.0666 |
| rs55986781 | T | 0.0187 | 0.0026 | 0.1182 | 0.0780 |
| rs56085180 | A | -0.0269 | 0.0037 | 0.1163 | 0.1151 |
| rs56099375 | T | 0.0120 | 0.0017 | -0.0108 | 0.0517 |
| rs56171318 | T | -0.0137 | 0.0020 | 0.0954 | 0.0633 |
| rs56174996 | A | -0.0143 | 0.0020 | -0.0248 | 0.0649 |
| rs56306882 | A | 0.0086 | 0.0016 | -0.1570 | 0.0498 |
| rs56319902 | T | -0.0206 | 0.0017 | -0.2734 | 0.0528 |
| rs56391344 | A | 0.0148 | 0.0016 | -0.0948 | 0.0508 |
| rs563954 | A | -0.0081 | 0.0014 | 0.0216 | 0.0443 |
| rs56405138 | A | 0.0153 | 0.0022 | 0.0689 | 0.0676 |
| rs56408528 | T | -0.0082 | 0.0014 | -0.0077 | 0.0443 |
| rs567003 | A | 0.0094 | 0.0015 | 0.0430 | 0.0465 |
| rs56794817 | A | 0.0122 | 0.0019 | -0.1145 | 0.0608 |
| rs57016874 | T | 0.0209 | 0.0037 | -0.1524 | 0.1130 |
| rs57148205 | A | 0.0095 | 0.0016 | -0.1135 | 0.0523 |
| rs57204268 | A | 0.0114 | 0.0020 | -0.1395 | 0.0614 |
| rs57349798 | A | 0.0093 | 0.0014 | -0.0172 | 0.0455 |
| rs57352738 | A | -0.0152 | 0.0017 | 0.0922 | 0.0545 |
| rs57437407 | A | -0.0145 | 0.0023 | -0.0849 | 0.0703 |
| rs575113 | A | 0.0133 | 0.0015 | 0.0688 | 0.0488 |
| rs5754581 | T | -0.0082 | 0.0014 | 0.1115 | 0.0446 |
| rs5754753 | T | -0.0126 | 0.0016 | -0.0103 | 0.0490 |
| rs5754762 | A | 0.0197 | 0.0030 | -0.1614 | 0.0978 |
| rs57661533 | T | -0.0114 | 0.0021 | 0.0176 | 0.0660 |
| rs580652 | T | -0.0147 | 0.0024 | 0.0377 | 0.0745 |
| rs585557 | A | 0.0128 | 0.0018 | -0.2410 | 0.0553 |
| rs58779949 | A | 0.0110 | 0.0019 | -0.0743 | 0.0573 |
| rs58859557 | T | 0.0187 | 0.0028 | 0.1457 | 0.0864 |
| rs58921703 | T | 0.0109 | 0.0015 | 0.0851 | 0.0466 |
| rs58950082 | T | 0.0095 | 0.0017 | -0.0722 | 0.0547 |
| rs58996896 | A | -0.0094 | 0.0016 | -0.0404 | 0.0499 |
| rs590013 | T | 0.0100 | 0.0015 | 0.0456 | 0.0473 |
| rs59300999 | T | -0.0164 | 0.0028 | -0.0979 | 0.0890 |
| rs59480703 | C | -0.0123 | 0.0018 | 0.1603 | 0.0556 |
| rs59813324 | T | 0.0097 | 0.0018 | -0.0260 | 0.0556 |
| rs59967356 | T | -0.0127 | 0.0021 | 0.0778 | 0.0657 |
| rs60096640 | A | 0.0159 | 0.0023 | -0.2028 | 0.0729 |
| rs6020560 | T | -0.0083 | 0.0014 | -0.0675 | 0.0443 |
| rs6043521 | T | -0.0083 | 0.0014 | 0.0538 | 0.0457 |
| rs60589532 | A | 0.0168 | 0.0028 | -0.1437 | 0.0933 |
| rs6060308 | A | 0.0095 | 0.0016 | 0.0899 | 0.0505 |
| rs6065080 | T | -0.0130 | 0.0015 | -0.0393 | 0.0463 |
| rs6065784 | C | 0.0110 | 0.0015 | 0.0062 | 0.0489 |
| rs60717745 | C | -0.0118 | 0.0019 | -0.0562 | 0.0612 |
| rs60726488 | T | -0.0090 | 0.0016 | -0.0428 | 0.0503 |
| rs6091570 | A | 0.0081 | 0.0015 | -0.0257 | 0.0468 |
| rs61104616 | A | -0.0172 | 0.0014 | -0.0867 | 0.0442 |
| rs6123924 | A | 0.0129 | 0.0019 | -0.0087 | 0.0614 |
| rs613872 | T | -0.0163 | 0.0019 | -0.1163 | 0.0583 |
| rs61387839 | A | 0.0088 | 0.0015 | -0.0244 | 0.0476 |
| rs61527214 | A | 0.0110 | 0.0014 | -0.0485 | 0.0451 |
| rs61739710 | A | -0.0099 | 0.0016 | -0.1245 | 0.0491 |
| rs61748951 | A | -0.0273 | 0.0048 | 0.2476 | 0.1474 |
| rs61755388 | T | 0.0125 | 0.0017 | -0.0475 | 0.0536 |
| rs61757207 | A | 0.0359 | 0.0063 | 0.0887 | 0.1829 |
| rs61798586 | A | -0.0112 | 0.0020 | -0.0076 | 0.0620 |
| rs61853335 | T | -0.0109 | 0.0018 | 0.0021 | 0.0581 |
| rs61958175 | A | -0.0198 | 0.0034 | 0.0525 | 0.1039 |
| rs61997667 | T | -0.0144 | 0.0020 | 0.0861 | 0.0613 |
| rs62007304 | A | 0.0264 | 0.0047 | -0.1417 | 0.1352 |
| rs62018216 | T | 0.0113 | 0.0019 | 0.0309 | 0.0614 |
| rs62051146 | A | -0.0134 | 0.0022 | -0.3401 | 0.0693 |
| rs62090515 | A | -0.0090 | 0.0015 | 0.0593 | 0.0461 |
| rs62092949 | T | -0.0087 | 0.0014 | -0.0058 | 0.0445 |
| rs62103198 | T | -0.0147 | 0.0025 | 0.0375 | 0.0775 |
| rs62109862 | A | 0.0136 | 0.0020 | -0.0401 | 0.0642 |
| rs62142891 | A | 0.0108 | 0.0016 | -0.0566 | 0.0503 |
| rs62155350 | A | 0.0338 | 0.0053 | 0.1493 | 0.1572 |
| rs62155770 | A | -0.0196 | 0.0032 | 0.0842 | 0.0951 |
| rs62155873 | T | -0.0151 | 0.0021 | 0.1928 | 0.0664 |
| rs62174974 | A | -0.0105 | 0.0018 | 0.1616 | 0.0556 |
| rs62177359 | A | -0.0275 | 0.0038 | 0.1214 | 0.1222 |
| rs62179650 | A | 0.0112 | 0.0016 | 0.0011 | 0.0479 |
| rs62182994 | T | 0.0137 | 0.0015 | -0.0259 | 0.0474 |
| rs62190914 | T | 0.0089 | 0.0015 | -0.1062 | 0.0460 |
| rs62194170 | A | 0.0111 | 0.0017 | -0.1683 | 0.0552 |
| rs622169 | T | 0.0086 | 0.0014 | -0.0385 | 0.0462 |
| rs62247449 | C | 0.0114 | 0.0014 | 0.0209 | 0.0446 |
| rs62252819 | A | -0.0094 | 0.0017 | -0.0659 | 0.0521 |
| rs62256284 | T | 0.0111 | 0.0017 | 0.1543 | 0.0542 |
| rs62260764 | C | -0.0132 | 0.0016 | -0.1516 | 0.0501 |
| rs62262671 | A | 0.0213 | 0.0020 | -0.2061 | 0.0638 |
| rs62340636 | T | 0.0103 | 0.0014 | 0.0248 | 0.0459 |
| rs62370510 | A | -0.0109 | 0.0019 | 0.0809 | 0.0578 |
| rs62379838 | T | 0.0121 | 0.0015 | 0.1173 | 0.0481 |
| rs62409395 | T | 0.0101 | 0.0018 | -0.0551 | 0.0548 |
| rs62420387 | T | -0.0184 | 0.0027 | 0.3290 | 0.0836 |
| rs62506074 | T | 0.0091 | 0.0015 | -0.0737 | 0.0469 |
| rs62506104 | A | -0.0098 | 0.0018 | 0.1261 | 0.0567 |
| rs628993 | A | 0.0128 | 0.0023 | -0.0147 | 0.0753 |
| rs631287 | A | 0.0084 | 0.0014 | 0.1939 | 0.0445 |
| rs633279 | A | -0.0089 | 0.0015 | 0.0533 | 0.0486 |
| rs635754 | A | 0.0119 | 0.0014 | 0.0571 | 0.0450 |
| rs6428587 | T | 0.0084 | 0.0015 | -0.0559 | 0.0459 |
| rs6435326 | A | -0.0091 | 0.0014 | 0.1079 | 0.0444 |
| rs6442126 | A | -0.0121 | 0.0015 | 0.0687 | 0.0468 |
| rs6445633 | A | 0.0088 | 0.0015 | 0.0351 | 0.0477 |
| rs6449503 | A | 0.0185 | 0.0014 | -0.0326 | 0.0445 |
| rs6450476 | A | -0.0116 | 0.0015 | -0.1081 | 0.0486 |
| rs6452793 | T | 0.0175 | 0.0017 | -0.1754 | 0.0534 |
| rs6461536 | A | 0.0088 | 0.0015 | -0.0952 | 0.0489 |
| rs6469654 | C | -0.0103 | 0.0017 | 0.1858 | 0.0547 |
| rs6472208 | T | -0.0091 | 0.0014 | 0.0117 | 0.0451 |
| rs6480234 | T | -0.0084 | 0.0014 | 0.1402 | 0.0443 |
| rs648044 | A | 0.0083 | 0.0015 | 0.0109 | 0.0463 |
| rs6490618 | T | 0.0088 | 0.0015 | -0.0278 | 0.0472 |
| rs6493265 | T | -0.0122 | 0.0014 | 0.0724 | 0.0452 |
| rs6493275 | A | 0.0131 | 0.0017 | -0.0215 | 0.0541 |
| rs6503409 | T | 0.0087 | 0.0015 | 0.0500 | 0.0464 |
| rs6534338 | T | 0.0103 | 0.0015 | -0.0199 | 0.0485 |
| rs6543810 | T | 0.0082 | 0.0015 | -0.0251 | 0.0454 |
| rs6557171 | T | -0.0155 | 0.0015 | 0.0818 | 0.0466 |
| rs6567288 | A | -0.0079 | 0.0014 | 0.1406 | 0.0448 |
| rs6569077 | T | 0.0118 | 0.0014 | -0.0673 | 0.0473 |
| rs6573552 | T | -0.0090 | 0.0014 | 0.0486 | 0.0442 |
| rs6573559 | T | 0.0105 | 0.0015 | 0.0043 | 0.0476 |
| rs660001 | A | -0.0164 | 0.0017 | 0.1957 | 0.0546 |
| rs663251 | T | 0.0081 | 0.0015 | -0.1603 | 0.0465 |
| rs66482320 | C | 0.0124 | 0.0022 | -0.0232 | 0.0705 |
| rs66568921 | T | -0.0181 | 0.0015 | -0.0472 | 0.0465 |
| rs66671632 | T | -0.0133 | 0.0021 | -0.1108 | 0.0665 |
| rs66721975 | A | -0.0087 | 0.0015 | 0.2263 | 0.0492 |
| rs6672986 | C | 0.0094 | 0.0014 | -0.0822 | 0.0444 |
| rs6689641 | A | 0.0078 | 0.0014 | -0.0711 | 0.0443 |
| rs6690195 | T | -0.0113 | 0.0014 | 0.0825 | 0.0443 |
| rs6695132 | T | 0.0097 | 0.0017 | -0.0470 | 0.0536 |
| rs6696068 | T | -0.0088 | 0.0014 | 0.1698 | 0.0453 |
| rs6697584 | T | -0.0126 | 0.0017 | 0.0539 | 0.0532 |
| rs6704768 | A | -0.0116 | 0.0014 | -0.0445 | 0.0445 |
| rs6706275 | T | 0.0083 | 0.0015 | -0.0660 | 0.0468 |
| rs6711399 | T | -0.0107 | 0.0019 | -0.0744 | 0.0583 |
| rs6715321 | T | -0.0097 | 0.0014 | 0.0222 | 0.0446 |
| rs6715849 | A | -0.0122 | 0.0014 | 0.0497 | 0.0446 |
| rs6720515 | A | 0.0103 | 0.0017 | -0.0073 | 0.0533 |
| rs67224963 | A | 0.0113 | 0.0017 | -0.1065 | 0.0551 |
| rs6729612 | T | 0.0092 | 0.0015 | 0.0360 | 0.0476 |
| rs6731373 | A | -0.0115 | 0.0015 | 0.2127 | 0.0460 |
| rs6736025 | T | 0.0079 | 0.0014 | -0.0281 | 0.0447 |
| rs6738860 | A | -0.0088 | 0.0014 | 0.0397 | 0.0444 |
| rs6743032 | A | -0.0165 | 0.0024 | -0.0926 | 0.0750 |
| rs6744040 | A | -0.0080 | 0.0014 | -0.0169 | 0.0445 |
| rs67456868 | A | 0.0119 | 0.0018 | -0.0335 | 0.0568 |
| rs6752228 | T | -0.0080 | 0.0014 | 0.1231 | 0.0443 |
| rs6757087 | T | -0.0085 | 0.0014 | 0.0443 | 0.0450 |
| rs67661275 | A | 0.0134 | 0.0025 | 0.1133 | 0.0780 |
| rs6774533 | T | 0.0090 | 0.0015 | -0.0767 | 0.0493 |
| rs67790232 | A | 0.0101 | 0.0017 | 0.0570 | 0.0547 |
| rs6780414 | T | 0.0105 | 0.0018 | 0.0470 | 0.0575 |
| rs6792805 | A | -0.0090 | 0.0016 | -0.0601 | 0.0499 |
| rs6812533 | T | 0.0106 | 0.0016 | 0.0686 | 0.0512 |
| rs68145588 | T | -0.0127 | 0.0021 | 0.0142 | 0.0625 |
| rs6824567 | T | 0.0130 | 0.0016 | -0.1792 | 0.0490 |
| rs6853599 | T | 0.0108 | 0.0019 | -0.1579 | 0.0576 |
| rs6861925 | C | -0.0090 | 0.0014 | 0.0301 | 0.0444 |
| rs6871635 | A | -0.0087 | 0.0014 | 0.1806 | 0.0449 |
| rs6881581 | A | -0.0096 | 0.0015 | 0.1074 | 0.0467 |
| rs6881733 | T | -0.0086 | 0.0016 | -0.0823 | 0.0491 |
| rs6917154 | T | 0.0127 | 0.0021 | -0.0727 | 0.0662 |
| rs6917204 | T | 0.0119 | 0.0017 | -0.0491 | 0.0557 |
| rs6918506 | C | 0.0113 | 0.0014 | 0.1136 | 0.0443 |
| rs6924023 | A | 0.0107 | 0.0016 | 0.0321 | 0.0521 |
| rs6932108 | C | -0.0210 | 0.0027 | -0.0901 | 0.0792 |
| rs6946136 | A | 0.0081 | 0.0014 | -0.0249 | 0.0452 |
| rs6946362 | T | 0.0091 | 0.0015 | -0.0112 | 0.0487 |
| rs6951996 | A | -0.0182 | 0.0032 | 0.1361 | 0.1022 |
| rs6956283 | T | 0.0098 | 0.0016 | 0.0804 | 0.0500 |
| rs6969783 | A | -0.0098 | 0.0014 | 0.0598 | 0.0445 |
| rs6977237 | T | 0.0082 | 0.0014 | 0.0983 | 0.0443 |
| rs6989141 | A | 0.0100 | 0.0017 | -0.0785 | 0.0553 |
| rs6994287 | A | 0.0102 | 0.0014 | -0.0428 | 0.0448 |
| rs7012546 | T | 0.0092 | 0.0014 | 0.0065 | 0.0449 |
| rs7016302 | C | -0.0125 | 0.0019 | 0.0316 | 0.0589 |
| rs702606 | T | 0.0118 | 0.0021 | -0.1376 | 0.0648 |
| rs7029718 | A | 0.0244 | 0.0014 | 0.0054 | 0.0448 |
| rs7030373 | A | -0.0098 | 0.0017 | 0.0598 | 0.0548 |
| rs7035315 | A | -0.0090 | 0.0015 | 0.1421 | 0.0460 |
| rs7040995 | C | 0.0097 | 0.0014 | -0.0667 | 0.0447 |
| rs7041702 | A | -0.0111 | 0.0016 | -0.0265 | 0.0503 |
| rs7084508 | T | -0.0085 | 0.0015 | 0.1212 | 0.0464 |
| rs710629 | A | 0.0082 | 0.0015 | -0.1144 | 0.0465 |
| rs7108020 | A | 0.0111 | 0.0015 | 0.0116 | 0.0461 |
| rs711793 | T | 0.0103 | 0.0015 | 0.0842 | 0.0474 |
| rs7127580 | C | -0.0150 | 0.0023 | 0.0852 | 0.0705 |
| rs713584 | A | -0.0080 | 0.0014 | 0.0217 | 0.0448 |
| rs71413877 | A | 0.0351 | 0.0036 | -0.1507 | 0.1134 |
| rs7146625 | A | 0.0091 | 0.0016 | 0.0647 | 0.0504 |
| rs7147473 | A | -0.0103 | 0.0015 | -0.0656 | 0.0472 |
| rs7158218 | A | -0.0090 | 0.0015 | 0.0454 | 0.0483 |
| rs71646142 | T | 0.0104 | 0.0018 | -0.0279 | 0.0563 |
| rs716513 | A | 0.0084 | 0.0014 | -0.0389 | 0.0444 |
| rs7167688 | T | -0.0094 | 0.0014 | 0.0252 | 0.0443 |
| rs7171405 | A | -0.0101 | 0.0016 | 0.1533 | 0.0521 |
| rs717996 | T | -0.0122 | 0.0014 | 0.1049 | 0.0453 |
| rs7182216 | C | -0.0096 | 0.0017 | -0.0982 | 0.0547 |
| rs7190 | A | -0.0095 | 0.0015 | -0.1158 | 0.0462 |
| rs721579 | T | -0.0110 | 0.0016 | -0.0302 | 0.0506 |
| rs7218235 | A | -0.0097 | 0.0017 | 0.0535 | 0.0546 |
| rs7223311 | C | -0.0079 | 0.0014 | 0.0280 | 0.0445 |
| rs7226824 | T | -0.0078 | 0.0014 | 0.0484 | 0.0443 |
| rs72482130 | T | 0.0141 | 0.0019 | -0.1144 | 0.0618 |
| rs7254263 | T | -0.0119 | 0.0016 | 0.1165 | 0.0494 |
| rs7255223 | A | 0.0097 | 0.0016 | -0.1611 | 0.0502 |
| rs72622559 | T | -0.0120 | 0.0017 | 0.0836 | 0.0567 |
| rs72624911 | T | 0.0282 | 0.0034 | -0.4305 | 0.1068 |
| rs72636697 | T | -0.0140 | 0.0020 | 0.0993 | 0.0652 |
| rs72667460 | T | 0.0197 | 0.0031 | -0.0523 | 0.0966 |
| rs72671456 | A | -0.0174 | 0.0020 | 0.0509 | 0.0646 |
| rs72672052 | A | -0.0101 | 0.0018 | 0.1032 | 0.0587 |
| rs72673097 | A | 0.0257 | 0.0042 | 0.3710 | 0.1213 |
| rs72677177 | A | 0.0100 | 0.0014 | 0.0298 | 0.0451 |
| rs72694479 | T | 0.0127 | 0.0020 | 0.0107 | 0.0632 |
| rs72709560 | A | 0.0085 | 0.0015 | -0.0272 | 0.0472 |
| rs72771860 | T | -0.0211 | 0.0029 | -0.0516 | 0.0924 |
| rs72792395 | T | -0.0145 | 0.0022 | 0.0301 | 0.0701 |
| rs728054 | A | -0.0128 | 0.0015 | 0.1003 | 0.0463 |
| rs72819118 | T | -0.0231 | 0.0024 | 0.0329 | 0.0752 |
| rs72824753 | T | 0.0133 | 0.0024 | 0.0771 | 0.0724 |
| rs72828517 | T | -0.0168 | 0.0019 | 0.1766 | 0.0575 |
| rs72829857 | A | -0.0149 | 0.0016 | 0.0756 | 0.0524 |
| rs72834698 | A | 0.0117 | 0.0020 | -0.1791 | 0.0633 |
| rs72881110 | A | 0.0146 | 0.0027 | 0.0048 | 0.0836 |
| rs72883760 | A | 0.0101 | 0.0018 | -0.1216 | 0.0555 |
| rs72896637 | T | 0.0162 | 0.0028 | -0.2336 | 0.0840 |
| rs72902523 | T | -0.0103 | 0.0016 | -0.0099 | 0.0513 |
| rs72906124 | T | 0.0185 | 0.0030 | -0.0710 | 0.0956 |
| rs72917504 | T | -0.0178 | 0.0030 | -0.2355 | 0.0938 |
| rs72919450 | T | -0.0132 | 0.0023 | 0.1270 | 0.0734 |
| rs72944064 | T | -0.0098 | 0.0016 | 0.1244 | 0.0506 |
| rs72962169 | T | -0.0174 | 0.0019 | 0.0667 | 0.0604 |
| rs72972965 | A | 0.0083 | 0.0015 | -0.1212 | 0.0462 |
| rs72993796 | T | 0.0127 | 0.0022 | -0.0831 | 0.0687 |
| rs73034295 | A | 0.0099 | 0.0018 | -0.0929 | 0.0567 |
| rs730384 | A | 0.0101 | 0.0014 | -0.0717 | 0.0448 |
| rs73039077 | C | 0.0092 | 0.0016 | -0.0356 | 0.0506 |
| rs73055556 | A | 0.0111 | 0.0020 | -0.0230 | 0.0638 |
| rs73082325 | C | -0.0252 | 0.0044 | 0.4356 | 0.1301 |
| rs73106136 | T | -0.0147 | 0.0026 | -0.2982 | 0.0823 |
| rs73154546 | A | 0.0197 | 0.0036 | -0.2548 | 0.1120 |
| rs7317761 | C | 0.0109 | 0.0016 | -0.0111 | 0.0504 |
| rs73191311 | A | -0.0094 | 0.0015 | 0.0898 | 0.0469 |
| rs7321274 | A | 0.0114 | 0.0018 | 0.0058 | 0.0550 |
| rs73219806 | A | 0.0106 | 0.0019 | 0.0698 | 0.0606 |
| rs7323027 | A | 0.0130 | 0.0015 | 0.0154 | 0.0471 |
| rs7326331 | A | -0.0128 | 0.0016 | 0.0372 | 0.0496 |
| rs73344830 | A | 0.0170 | 0.0014 | -0.1032 | 0.0448 |
| rs73405293 | A | 0.0113 | 0.0020 | 0.0968 | 0.0623 |
| rs73457936 | A | 0.0144 | 0.0025 | 0.0387 | 0.0778 |
| rs7356536 | T | -0.0129 | 0.0021 | 0.1913 | 0.0642 |
| rs73581580 | A | -0.0181 | 0.0022 | -0.0355 | 0.0669 |
| rs736281 | T | 0.0086 | 0.0014 | -0.1172 | 0.0452 |
| rs73643713 | T | -0.0165 | 0.0027 | 0.0499 | 0.0849 |
| rs736471 | T | 0.0083 | 0.0014 | -0.0578 | 0.0446 |
| rs73648455 | T | -0.0183 | 0.0026 | 0.0022 | 0.0840 |
| rs737945 | C | -0.0115 | 0.0014 | 0.0870 | 0.0445 |
| rs73874335 | T | -0.0178 | 0.0029 | 0.1351 | 0.0970 |
| rs73961845 | A | -0.0114 | 0.0014 | 0.0141 | 0.0454 |
| rs74091672 | A | -0.0153 | 0.0022 | 0.0257 | 0.0677 |
| rs7430651 | T | -0.0111 | 0.0016 | 0.0177 | 0.0483 |
| rs743316 | T | 0.0112 | 0.0017 | 0.1110 | 0.0543 |
| rs74415461 | T | 0.0176 | 0.0026 | -0.0280 | 0.0783 |
| rs74453875 | A | 0.0214 | 0.0037 | -0.0958 | 0.1195 |
| rs74462621 | T | 0.0125 | 0.0020 | -0.0452 | 0.0647 |
| rs7449561 | A | 0.0102 | 0.0017 | 0.0139 | 0.0528 |
| rs7451726 | A | 0.0109 | 0.0017 | -0.0638 | 0.0547 |
| rs74545339 | A | -0.0158 | 0.0022 | -0.0475 | 0.0703 |
| rs7460106 | T | -0.0123 | 0.0017 | 0.1675 | 0.0529 |
| rs74615093 | A | 0.0145 | 0.0026 | 0.0582 | 0.0810 |
| rs746839 | C | 0.0135 | 0.0015 | -0.0711 | 0.0463 |
| rs74747621 | T | 0.0153 | 0.0022 | -0.1544 | 0.0703 |
| rs74787922 | A | 0.0182 | 0.0028 | 0.0079 | 0.0877 |
| rs74944275 | T | 0.0231 | 0.0035 | 0.1961 | 0.1050 |
| rs74944857 | T | -0.0089 | 0.0015 | 0.0714 | 0.0469 |
| rs7495033 | A | 0.0095 | 0.0015 | -0.3547 | 0.0489 |
| rs75033012 | C | -0.0258 | 0.0038 | -0.2226 | 0.1127 |
| rs75177132 | T | 0.0298 | 0.0036 | -0.1252 | 0.1020 |
| rs75203411 | T | 0.0135 | 0.0019 | -0.0764 | 0.0590 |
| rs7526112 | T | 0.0124 | 0.0015 | 0.0260 | 0.0459 |
| rs75308819 | A | 0.0287 | 0.0047 | 0.1488 | 0.1494 |
| rs75434274 | A | 0.0151 | 0.0026 | -0.2109 | 0.0790 |
| rs75500877 | T | 0.0207 | 0.0033 | -0.2212 | 0.1020 |
| rs7552964 | A | 0.0148 | 0.0023 | -0.1681 | 0.0749 |
| rs7560871 | A | -0.0193 | 0.0027 | 0.1493 | 0.0866 |
| rs7561705 | A | 0.0078 | 0.0014 | 0.0234 | 0.0443 |
| rs7575637 | A | 0.0107 | 0.0014 | -0.1676 | 0.0443 |
| rs75756843 | A | 0.0288 | 0.0049 | -0.0513 | 0.1665 |
| rs7575938 | A | 0.0090 | 0.0015 | 0.0499 | 0.0467 |
| rs7590368 | T | -0.0146 | 0.0016 | 0.0811 | 0.0503 |
| rs7597126 | T | -0.0094 | 0.0014 | 0.0506 | 0.0443 |
| rs7597412 | C | 0.0088 | 0.0015 | -0.0321 | 0.0460 |
| rs7603132 | A | 0.0137 | 0.0018 | -0.1234 | 0.0556 |
| rs76076331 | T | 0.0206 | 0.0021 | -0.1192 | 0.0675 |
| rs76077165 | A | 0.0165 | 0.0025 | -0.2660 | 0.0799 |
| rs76167224 | T | -0.0187 | 0.0033 | 0.3284 | 0.0997 |
| rs7617204 | A | -0.0097 | 0.0014 | 0.1134 | 0.0443 |
| rs76235882 | A | 0.0241 | 0.0039 | -0.0173 | 0.1315 |
| rs76241605 | A | 0.0139 | 0.0022 | -0.1209 | 0.0690 |
| rs76246107 | A | -0.0162 | 0.0026 | 0.0092 | 0.0834 |
| rs7625428 | T | 0.0098 | 0.0014 | 0.0803 | 0.0452 |
| rs76267866 | A | 0.0104 | 0.0017 | 0.0159 | 0.0546 |
| rs7630133 | A | 0.0082 | 0.0015 | -0.0754 | 0.0464 |
| rs763553 | A | 0.0087 | 0.0014 | -0.0924 | 0.0445 |
| rs7640424 | T | 0.0113 | 0.0015 | 0.0910 | 0.0477 |
| rs7650602 | T | -0.0089 | 0.0014 | -0.0383 | 0.0446 |
| rs76552497 | T | -0.0120 | 0.0019 | 0.1559 | 0.0590 |
| rs76577427 | C | 0.0203 | 0.0024 | -0.0850 | 0.0722 |
| rs7672622 | A | -0.0088 | 0.0016 | 0.1055 | 0.0512 |
| rs767943 | A | -0.0122 | 0.0016 | 0.1687 | 0.0505 |
| rs7683416 | T | 0.0139 | 0.0014 | -0.1420 | 0.0445 |
| rs76876592 | A | 0.0126 | 0.0020 | 0.0314 | 0.0648 |
| rs76878669 | C | 0.0141 | 0.0017 | -0.0414 | 0.0524 |
| rs7692359 | T | -0.0109 | 0.0017 | 0.0476 | 0.0538 |
| rs76957677 | T | -0.0297 | 0.0054 | 0.3804 | 0.1577 |
| rs77025239 | A | -0.0140 | 0.0019 | 0.0126 | 0.0618 |
| rs7714719 | C | 0.0116 | 0.0020 | -0.0845 | 0.0630 |
| rs7716161 | C | 0.0105 | 0.0019 | 0.1367 | 0.0581 |
| rs77201694 | A | 0.0156 | 0.0022 | 0.0167 | 0.0684 |
| rs77370942 | A | -0.0147 | 0.0027 | -0.0610 | 0.0830 |
| rs77554090 | T | -0.0186 | 0.0027 | 0.0448 | 0.0841 |
| rs77609760 | A | 0.0236 | 0.0028 | -0.1124 | 0.0880 |
| rs7766240 | T | -0.0134 | 0.0024 | 0.0535 | 0.0746 |
| rs77702622 | A | -0.0213 | 0.0029 | -0.0274 | 0.0890 |
| rs77702819 | T | 0.0187 | 0.0025 | -0.1398 | 0.0760 |
| rs77719387 | A | -0.0403 | 0.0058 | 0.7345 | 0.1775 |
| rs7772172 | A | 0.0091 | 0.0014 | 0.0013 | 0.0450 |
| rs7775100 | T | -0.0087 | 0.0014 | 0.0873 | 0.0452 |
| rs77826402 | T | 0.0151 | 0.0027 | 0.0683 | 0.0858 |
| rs7796103 | C | -0.0077 | 0.0014 | -0.0284 | 0.0447 |
| rs7799141 | A | 0.0099 | 0.0015 | -0.0093 | 0.0474 |
| rs77999825 | A | 0.0294 | 0.0039 | -0.3624 | 0.1206 |
| rs7803932 | A | 0.0127 | 0.0019 | -0.0470 | 0.0591 |
| rs780569 | A | -0.0085 | 0.0016 | 0.0032 | 0.0489 |
| rs78116078 | C | 0.0086 | 0.0016 | 0.0114 | 0.0488 |
| rs7816777 | T | -0.0088 | 0.0015 | 0.0473 | 0.0465 |
| rs78193153 | A | 0.0191 | 0.0029 | -0.2361 | 0.0913 |
| rs7823700 | T | 0.0132 | 0.0024 | -0.0325 | 0.0745 |
| rs78365243 | T | 0.0181 | 0.0033 | -0.0634 | 0.1010 |
| rs78440611 | A | -0.0147 | 0.0024 | -0.0103 | 0.0761 |
| rs7849480 | A | -0.0084 | 0.0014 | 0.1135 | 0.0447 |
| rs7849487 | T | -0.0167 | 0.0015 | -0.0176 | 0.0467 |
| rs7855503 | C | 0.0088 | 0.0015 | 0.0300 | 0.0461 |
| rs78648104 | T | -0.0141 | 0.0025 | -0.2874 | 0.0787 |
| rs78702390 | C | 0.0167 | 0.0030 | -0.1621 | 0.0929 |
| rs78714229 | T | 0.0172 | 0.0030 | 0.1572 | 0.0947 |
| rs7875078 | A | -0.0078 | 0.0014 | 0.0303 | 0.0446 |
| rs78918150 | T | -0.0080 | 0.0015 | 0.0430 | 0.0456 |
| rs7894722 | T | 0.0085 | 0.0014 | -0.0025 | 0.0455 |
| rs7899270 | A | -0.0084 | 0.0015 | -0.0422 | 0.0465 |
| rs7905192 | T | 0.0084 | 0.0014 | 0.0859 | 0.0444 |
| rs790647 | A | -0.0145 | 0.0017 | 0.0758 | 0.0524 |
| rs7910403 | T | 0.0122 | 0.0018 | -0.1157 | 0.0554 |
| rs7914674 | T | -0.0139 | 0.0022 | -0.0073 | 0.0549 |
| rs7920624 | A | 0.0120 | 0.0014 | -0.0910 | 0.0445 |
| rs7924036 | T | 0.0133 | 0.0014 | -0.1268 | 0.0442 |
| rs79265434 | A | -0.0197 | 0.0022 | 0.3693 | 0.0691 |
| rs7928017 | A | 0.0096 | 0.0014 | -0.0834 | 0.0445 |
| rs7931563 | T | -0.0109 | 0.0014 | 0.0410 | 0.0457 |
| rs795230 | T | 0.0081 | 0.0014 | 0.1249 | 0.0447 |
| rs7958371 | A | -0.0085 | 0.0015 | 0.0506 | 0.0463 |
| rs79585412 | T | -0.0178 | 0.0031 | 0.2009 | 0.0945 |
| rs7965154 | A | -0.0085 | 0.0015 | 0.0832 | 0.0477 |
| rs7967550 | A | -0.0086 | 0.0014 | -0.0129 | 0.0452 |
| rs7972246 | T | 0.0110 | 0.0015 | -0.2111 | 0.0467 |
| rs79728014 | A | -0.0134 | 0.0020 | 0.0659 | 0.0620 |
| rs7974852 | A | 0.0117 | 0.0014 | -0.1255 | 0.0442 |
| rs7977614 | A | -0.0134 | 0.0016 | 0.0936 | 0.0485 |
| rs79798166 | A | 0.0172 | 0.0025 | -0.0345 | 0.0800 |
| rs79855925 | T | -0.0194 | 0.0033 | -0.0210 | 0.1082 |
| rs79994730 | T | 0.0240 | 0.0035 | -0.0523 | 0.1017 |
| rs79997166 | A | 0.0172 | 0.0030 | -0.2574 | 0.0926 |
| rs80037907 | T | -0.0107 | 0.0019 | 0.1139 | 0.0599 |
| rs8008382 | T | -0.0106 | 0.0015 | -0.0225 | 0.0484 |
| rs8009933 | A | 0.0088 | 0.0015 | -0.0532 | 0.0477 |
| rs8016504 | A | -0.0087 | 0.0014 | 0.0150 | 0.0447 |
| rs80223410 | T | -0.0118 | 0.0020 | 0.0153 | 0.0641 |
| rs8024 | A | -0.0115 | 0.0015 | 0.1466 | 0.0466 |
| rs80257979 | T | -0.0252 | 0.0040 | 0.3318 | 0.1316 |
| rs8030487 | A | 0.0085 | 0.0015 | -0.1296 | 0.0478 |
| rs803619 | T | -0.0173 | 0.0023 | 0.1388 | 0.0743 |
| rs8046072 | A | 0.0101 | 0.0018 | -0.2254 | 0.0556 |
| rs8052523 | T | -0.0084 | 0.0014 | -0.0496 | 0.0452 |
| rs8066044 | A | 0.0090 | 0.0016 | -0.2723 | 0.0501 |
| rs806816 | A | 0.0101 | 0.0016 | -0.2070 | 0.0511 |
| rs8097125 | T | 0.0086 | 0.0014 | -0.0636 | 0.0454 |
| rs8103741 | A | -0.0108 | 0.0019 | 0.0676 | 0.0576 |
| rs818415 | T | -0.0110 | 0.0018 | 0.0449 | 0.0566 |
| rs8192465 | A | -0.0272 | 0.0049 | 0.0320 | 0.1496 |
| rs852771 | T | 0.0092 | 0.0015 | -0.0804 | 0.0480 |
| rs853286 | T | 0.0124 | 0.0023 | -0.0205 | 0.0736 |
| rs854796 | A | 0.0090 | 0.0015 | 0.0410 | 0.0476 |
| rs868456 | A | 0.0089 | 0.0015 | 0.0084 | 0.0485 |
| rs870589 | A | -0.0079 | 0.0014 | 0.0490 | 0.0444 |
| rs884108 | A | -0.0094 | 0.0017 | 0.0101 | 0.0525 |
| rs891793 | C | -0.0099 | 0.0014 | 0.0584 | 0.0444 |
| rs893522 | A | 0.0149 | 0.0025 | -0.1039 | 0.0764 |
| rs902820 | A | -0.0096 | 0.0014 | -0.0380 | 0.0448 |
| rs911149 | T | 0.0091 | 0.0017 | 0.0304 | 0.0522 |
| rs912883 | T | 0.0087 | 0.0015 | 0.0670 | 0.0475 |
| rs913509 | A | 0.0096 | 0.0015 | -0.0025 | 0.0469 |
| rs925161 | C | -0.0078 | 0.0014 | 0.0074 | 0.0443 |
| rs9267658 | T | 0.0138 | 0.0020 | -0.3224 | 0.0623 |
| rs9267677 | T | 0.0180 | 0.0024 | -0.0001 | 0.0758 |
| rs9289300 | T | -0.0164 | 0.0019 | 0.1380 | 0.0609 |
| rs9291437 | C | -0.0081 | 0.0014 | 0.0740 | 0.0455 |
| rs9294770 | T | 0.0083 | 0.0014 | 0.0086 | 0.0459 |
| rs929511 | T | -0.0160 | 0.0021 | -0.0372 | 0.0670 |
| rs9300612 | T | 0.0084 | 0.0015 | -0.0079 | 0.0483 |
| rs933738 | A | -0.0110 | 0.0018 | 0.3409 | 0.0588 |
| rs9349956 | A | -0.0159 | 0.0018 | 0.0147 | 0.0571 |
| rs9359939 | A | -0.0105 | 0.0016 | 0.0533 | 0.0523 |
| rs936496 | A | -0.0079 | 0.0014 | -0.0423 | 0.0455 |
| rs9371883 | C | -0.0087 | 0.0015 | 0.0107 | 0.0461 |
| rs9373363 | A | -0.0111 | 0.0016 | 0.2332 | 0.0511 |
| rs9375188 | T | 0.0212 | 0.0014 | -0.1275 | 0.0443 |
| rs9375403 | C | 0.0126 | 0.0016 | -0.1313 | 0.0522 |
| rs9386110 | T | -0.0163 | 0.0023 | 0.0043 | 0.0713 |
| rs9388490 | T | 0.0091 | 0.0014 | 0.0311 | 0.0446 |
| rs939400 | T | -0.0095 | 0.0015 | 0.0745 | 0.0461 |
| rs9411331 | A | 0.0145 | 0.0015 | -0.0490 | 0.0474 |
| rs9435340 | A | 0.0086 | 0.0015 | -0.0389 | 0.0467 |
| rs9442750 | A | -0.0089 | 0.0016 | -0.0236 | 0.0502 |
| rs9446060 | A | 0.0077 | 0.0014 | 0.0530 | 0.0456 |
| rs9465509 | A | -0.0084 | 0.0014 | -0.0116 | 0.0442 |
| rs9490512 | A | -0.0136 | 0.0014 | 0.1525 | 0.0445 |
| rs9492774 | C | 0.0080 | 0.0015 | -0.0334 | 0.0459 |
| rs9513416 | A | -0.0116 | 0.0019 | 0.0626 | 0.0603 |
| rs9513754 | T | 0.0096 | 0.0016 | -0.0690 | 0.0489 |
| rs9527662 | A | 0.0094 | 0.0014 | 0.0442 | 0.0466 |
| rs9527905 | A | -0.0094 | 0.0014 | 0.0940 | 0.0451 |
| rs9529146 | T | 0.0127 | 0.0017 | -0.0959 | 0.0528 |
| rs9536462 | A | 0.0133 | 0.0020 | -0.0822 | 0.0636 |
| rs9540718 | A | -0.0089 | 0.0014 | 0.0953 | 0.0444 |
| rs9545395 | T | 0.0138 | 0.0021 | 0.0189 | 0.0663 |
| rs9556958 | T | -0.0110 | 0.0014 | 0.1348 | 0.0445 |
| rs9563168 | A | 0.0101 | 0.0017 | -0.1964 | 0.0546 |
| rs9568798 | T | -0.0093 | 0.0016 | 0.0421 | 0.0491 |
| rs9597907 | T | 0.0152 | 0.0027 | -0.0906 | 0.0872 |
| rs9611597 | A | -0.0107 | 0.0019 | 0.1415 | 0.0610 |
| rs9616906 | A | 0.0122 | 0.0014 | 0.0272 | 0.0446 |
| rs9616947 | T | -0.0134 | 0.0015 | 0.0289 | 0.0456 |
| rs9633970 | T | 0.0091 | 0.0016 | -0.0251 | 0.0531 |
| rs9649 | T | 0.0108 | 0.0019 | -0.0675 | 0.0590 |
| rs9655780 | A | -0.0153 | 0.0019 | 0.0544 | 0.0593 |
| rs9666728 | T | -0.0082 | 0.0014 | 0.1334 | 0.0443 |
| rs9683585 | C | 0.0084 | 0.0014 | 0.0636 | 0.0445 |
| rs969512 | A | -0.0106 | 0.0015 | -0.0385 | 0.0469 |
| rs9771228 | T | 0.0096 | 0.0015 | 0.0263 | 0.0462 |
| rs977143 | A | 0.0085 | 0.0015 | -0.1333 | 0.0459 |
| rs978807 | A | 0.0139 | 0.0018 | 0.0005 | 0.0559 |
| rs981230 | T | 0.0100 | 0.0014 | 0.0123 | 0.0443 |
| rs981883 | A | 0.0099 | 0.0014 | -0.1676 | 0.0451 |
| rs9820604 | T | 0.0115 | 0.0018 | -0.0636 | 0.0585 |
| rs9821664 | A | 0.0094 | 0.0017 | 0.0067 | 0.0552 |
| rs9830359 | T | -0.0147 | 0.0022 | -0.1921 | 0.0701 |
| rs9844755 | A | 0.0083 | 0.0015 | -0.0452 | 0.0470 |
| rs9849884 | A | -0.0090 | 0.0014 | -0.1602 | 0.0449 |
| rs9853928 | T | -0.0127 | 0.0017 | 0.0596 | 0.0550 |
| rs9858921 | A | -0.0087 | 0.0014 | 0.1422 | 0.0446 |
| rs9859556 | T | 0.0290 | 0.0015 | -0.2416 | 0.0480 |
| rs9866123 | A | 0.0086 | 0.0014 | 0.0268 | 0.0446 |
| rs9870317 | A | 0.0086 | 0.0015 | 0.0057 | 0.0477 |
| rs9877225 | T | 0.0090 | 0.0016 | -0.0917 | 0.0517 |
| rs9882532 | T | 0.0124 | 0.0015 | -0.1023 | 0.0462 |
| rs9886703 | A | -0.0136 | 0.0019 | -0.1061 | 0.0591 |
| rs9888796 | T | 0.0113 | 0.0016 | 0.0796 | 0.0507 |
| rs9916901 | T | 0.0092 | 0.0016 | 0.0039 | 0.0504 |
| rs9924031 | C | 0.0093 | 0.0015 | -0.1213 | 0.0455 |
| rs9926649 | T | 0.0177 | 0.0031 | 0.3493 | 0.1024 |
| rs9927049 | A | 0.0099 | 0.0016 | 0.1250 | 0.0508 |
| rs9927137 | A | 0.0082 | 0.0014 | -0.1498 | 0.0446 |
| rs9927842 | T | -0.0130 | 0.0020 | 0.0506 | 0.0609 |
| rs9929556 | T | -0.0099 | 0.0014 | -0.0599 | 0.0448 |
| rs9929762 | A | 0.0096 | 0.0014 | -0.0378 | 0.0447 |
| rs9929993 | T | -0.0102 | 0.0015 | 0.1178 | 0.0464 |
| rs9933256 | A | 0.0119 | 0.0014 | -0.0576 | 0.0444 |
| rs995698 | A | -0.0099 | 0.0014 | 0.0732 | 0.0445 |
| rs9974899 | T | -0.0093 | 0.0016 | 0.0045 | 0.0514 |
| rs9977825 | T | -0.0092 | 0.0015 | 0.0773 | 0.0461 |

#### Supplementary table 14: Genetic association estimates for the IVW ratio-method MR analysis estimating the effect of education on smoking. ea=effect allele; gx=education; gy=smoking; se=standard error

| SNP | ea | gx | gx_se | gy | gy_se |
| --- | --- | --- | --- | --- | --- |
| rs10006235 | T | -0.0093 | 0.0016 | 0.0052 | 0.0022 |
| rs10009513 | A | 0.0088 | 0.0014 | 0.0016 | 0.0020 |
| rs10032941 | T | -0.0080 | 0.0015 | 0.0047 | 0.0021 |
| rs10041403 | T | 0.0151 | 0.0019 | -0.0103 | 0.0026 |
| rs10042828 | A | -0.0145 | 0.0022 | 0.0055 | 0.0031 |
| rs10046069 | A | 0.0127 | 0.0022 | -0.0020 | 0.0031 |
| rs10057590 | A | 0.0106 | 0.0014 | -0.0050 | 0.0020 |
| rs10060023 | T | 0.0110 | 0.0015 | -0.0051 | 0.0021 |
| rs10061420 | A | 0.0082 | 0.0014 | 0.0010 | 0.0020 |
| rs1006749 | A | 0.0079 | 0.0014 | -0.0027 | 0.0020 |
| rs10074178 | A | 0.0079 | 0.0014 | 0.0034 | 0.0020 |
| rs1007731 | A | -0.0143 | 0.0022 | -0.0013 | 0.0031 |
| rs10080647 | A | 0.0124 | 0.0020 | -0.0073 | 0.0029 |
| rs1008078 | T | -0.0182 | 0.0014 | 0.0132 | 0.0020 |
| rs1009470 | T | 0.0086 | 0.0014 | -0.0002 | 0.0020 |
| rs10098073 | A | -0.0091 | 0.0014 | -0.0003 | 0.0020 |
| rs10122669 | A | -0.0079 | 0.0014 | -0.0008 | 0.0020 |
| rs10128888 | A | -0.0083 | 0.0015 | 0.0061 | 0.0022 |
| rs10145520 | T | -0.0122 | 0.0018 | 0.0107 | 0.0025 |
| rs10166286 | T | 0.0151 | 0.0019 | -0.0019 | 0.0026 |
| rs10169002 | A | -0.0080 | 0.0014 | 0.0006 | 0.0020 |
| rs10181071 | A | 0.0102 | 0.0019 | -0.0068 | 0.0026 |
| rs10189857 | A | 0.0158 | 0.0014 | -0.0046 | 0.0020 |
| rs10192369 | A | -0.0078 | 0.0014 | 0.0021 | 0.0020 |
| rs10192834 | T | 0.0083 | 0.0015 | -0.0036 | 0.0021 |
| rs10193498 | A | 0.0097 | 0.0016 | -0.0121 | 0.0023 |
| rs10204051 | T | -0.0101 | 0.0014 | 0.0093 | 0.0020 |
| rs10205057 | T | 0.0090 | 0.0014 | -0.0066 | 0.0020 |
| rs10208 | T | 0.0091 | 0.0015 | -0.0003 | 0.0021 |
| rs10215082 | A | -0.0149 | 0.0014 | 0.0047 | 0.0020 |
| rs1024268 | T | -0.0084 | 0.0014 | 0.0048 | 0.0020 |
| rs10251438 | A | 0.0078 | 0.0014 | 0.0001 | 0.0020 |
| rs10264573 | A | 0.0175 | 0.0032 | 0.0002 | 0.0046 |
| rs1035578 | A | -0.0094 | 0.0014 | 0.0084 | 0.0020 |
| rs10402747 | T | 0.0077 | 0.0014 | -0.0070 | 0.0020 |
| rs10411759 | A | 0.0116 | 0.0020 | -0.0068 | 0.0028 |
| rs10444280 | T | 0.0100 | 0.0018 | -0.0088 | 0.0025 |
| rs10460095 | A | -0.0124 | 0.0014 | 0.0056 | 0.0020 |
| rs10480450 | A | 0.0158 | 0.0028 | -0.0057 | 0.0042 |
| rs10496091 | A | -0.0131 | 0.0016 | 0.0075 | 0.0022 |
| rs10499535 | A | 0.0080 | 0.0014 | -0.0047 | 0.0020 |
| rs1050847 | T | 0.0092 | 0.0014 | -0.0114 | 0.0020 |
| rs10509251 | T | -0.0093 | 0.0017 | 0.0024 | 0.0024 |
| rs10515007 | T | 0.0143 | 0.0019 | 0.0001 | 0.0027 |
| rs1051860 | A | -0.0079 | 0.0014 | 0.0062 | 0.0020 |
| rs10519504 | T | 0.0119 | 0.0019 | -0.0018 | 0.0027 |
| rs1054442 | A | -0.0136 | 0.0015 | 0.0053 | 0.0021 |
| rs1061801 | A | -0.0110 | 0.0018 | 0.0135 | 0.0025 |
| rs1066769 | A | -0.0230 | 0.0041 | 0.0049 | 0.0054 |
| rs10742591 | A | -0.0084 | 0.0014 | 0.0081 | 0.0021 |
| rs10750539 | A | -0.0095 | 0.0015 | 0.0062 | 0.0021 |
| rs10752262 | T | 0.0102 | 0.0014 | -0.0037 | 0.0020 |
| rs1075228 | A | -0.0081 | 0.0014 | 0.0035 | 0.0020 |
| rs10761202 | T | 0.0088 | 0.0014 | 0.0037 | 0.0020 |
| rs10761251 | A | 0.0113 | 0.0015 | -0.0012 | 0.0021 |
| rs10772644 | C | 0.0142 | 0.0022 | -0.0069 | 0.0032 |
| rs10773002 | A | 0.0185 | 0.0016 | -0.0008 | 0.0023 |
| rs10773208 | T | -0.0110 | 0.0016 | 0.0013 | 0.0023 |
| rs10782651 | T | -0.0082 | 0.0014 | 0.0065 | 0.0020 |
| rs10783243 | A | -0.0092 | 0.0014 | 0.0066 | 0.0020 |
| rs10789285 | T | 0.0091 | 0.0016 | 0.0001 | 0.0024 |
| rs10793903 | A | 0.0126 | 0.0022 | -0.0089 | 0.0032 |
| rs10795831 | T | -0.0099 | 0.0017 | 0.0074 | 0.0025 |
| rs10798888 | T | -0.0140 | 0.0019 | 0.0143 | 0.0026 |
| rs10799615 | A | -0.0089 | 0.0016 | -0.0013 | 0.0023 |
| rs10805383 | A | -0.0103 | 0.0014 | 0.0081 | 0.0020 |
| rs10810099 | A | -0.0146 | 0.0016 | 0.0052 | 0.0022 |
| rs10810145 | A | 0.0078 | 0.0014 | -0.0065 | 0.0020 |
| rs10830858 | T | 0.0092 | 0.0014 | -0.0025 | 0.0020 |
| rs10831656 | T | 0.0090 | 0.0015 | -0.0056 | 0.0022 |
| rs10844179 | A | 0.0104 | 0.0017 | -0.0032 | 0.0024 |
| rs10845051 | A | 0.0083 | 0.0014 | -0.0050 | 0.0020 |
| rs10875121 | C | 0.0174 | 0.0019 | -0.0068 | 0.0027 |
| rs10877283 | T | 0.0085 | 0.0014 | -0.0015 | 0.0020 |
| rs10879676 | T | -0.0083 | 0.0014 | 0.0036 | 0.0020 |
| rs10886012 | C | 0.0078 | 0.0014 | -0.0045 | 0.0020 |
| rs10892807 | T | 0.0109 | 0.0014 | -0.0039 | 0.0020 |
| rs10899282 | A | 0.0140 | 0.0017 | -0.0049 | 0.0024 |
| rs10927053 | A | -0.0125 | 0.0022 | 0.0160 | 0.0032 |
| rs10928190 | T | -0.0087 | 0.0014 | 0.0044 | 0.0020 |
| rs10931821 | A | -0.0141 | 0.0014 | 0.0059 | 0.0020 |
| rs10937240 | T | -0.0123 | 0.0018 | -0.0006 | 0.0025 |
| rs10940540 | A | 0.0078 | 0.0014 | -0.0023 | 0.0020 |
| rs10940921 | T | 0.0111 | 0.0014 | -0.0049 | 0.0020 |
| rs10943588 | A | 0.0084 | 0.0015 | -0.0017 | 0.0021 |
| rs10947439 | T | 0.0090 | 0.0015 | -0.0051 | 0.0021 |
| rs10949263 | A | -0.0091 | 0.0016 | 0.0073 | 0.0023 |
| rs10951590 | T | -0.0109 | 0.0015 | 0.0060 | 0.0021 |
| rs10974256 | A | -0.0093 | 0.0015 | 0.0036 | 0.0021 |
| rs1097784 | T | -0.0083 | 0.0014 | 0.0014 | 0.0020 |
| rs10979613 | T | -0.0117 | 0.0015 | 0.0077 | 0.0021 |
| rs10983324 | A | 0.0084 | 0.0015 | -0.0037 | 0.0022 |
| rs10984445 | A | 0.0115 | 0.0014 | 0.0003 | 0.0020 |
| rs10985402 | T | 0.0089 | 0.0016 | -0.0017 | 0.0023 |
| rs10995639 | T | 0.0078 | 0.0014 | -0.0023 | 0.0020 |
| rs11003463 | T | 0.0091 | 0.0015 | 0.0041 | 0.0020 |
| rs11021432 | A | 0.0103 | 0.0015 | -0.0014 | 0.0021 |
| rs11023764 | A | 0.0091 | 0.0015 | -0.0007 | 0.0021 |
| rs11030084 | T | 0.0106 | 0.0018 | -0.0171 | 0.0026 |
| rs11030102 | C | 0.0101 | 0.0016 | -0.0159 | 0.0023 |
| rs1106090 | A | 0.0094 | 0.0014 | -0.0003 | 0.0021 |
| rs11076962 | T | 0.0086 | 0.0016 | -0.0049 | 0.0022 |
| rs1107871 | A | -0.0083 | 0.0014 | 0.0023 | 0.0020 |
| rs11081529 | T | 0.0126 | 0.0015 | 0.0055 | 0.0022 |
| rs11082011 | T | 0.0194 | 0.0015 | -0.0114 | 0.0021 |
| rs11100237 | A | -0.0101 | 0.0014 | -0.0001 | 0.0020 |
| rs11115056 | A | 0.0187 | 0.0023 | 0.0006 | 0.0034 |
| rs11121177 | A | 0.0151 | 0.0018 | -0.0118 | 0.0026 |
| rs111226181 | T | -0.0201 | 0.0018 | -0.0001 | 0.0026 |
| rs111235962 | T | 0.0133 | 0.0024 | -0.0050 | 0.0034 |
| rs11130380 | T | 0.0090 | 0.0014 | -0.0072 | 0.0020 |
| rs111370527 | T | 0.0151 | 0.0027 | 0.0058 | 0.0037 |
| rs11138947 | T | 0.0105 | 0.0016 | -0.0029 | 0.0022 |
| rs111530150 | T | 0.0269 | 0.0048 | -0.0064 | 0.0085 |
| rs11155813 | T | -0.0129 | 0.0023 | 0.0020 | 0.0033 |
| rs11157931 | A | -0.0139 | 0.0014 | 0.0018 | 0.0020 |
| rs11158800 | A | -0.0091 | 0.0014 | 0.0083 | 0.0020 |
| rs1117152 | T | 0.0106 | 0.0015 | -0.0042 | 0.0022 |
| rs11174399 | A | 0.0111 | 0.0018 | -0.0001 | 0.0025 |
| rs111821073 | T | 0.0124 | 0.0019 | -0.0054 | 0.0028 |
| rs111852224 | T | 0.0156 | 0.0022 | -0.0049 | 0.0030 |
| rs11190955 | T | -0.0083 | 0.0015 | 0.0065 | 0.0021 |
| rs1120924 | T | 0.0108 | 0.0015 | -0.0055 | 0.0021 |
| rs11209894 | A | 0.0119 | 0.0021 | -0.0007 | 0.0029 |
| rs11210400 | A | 0.0120 | 0.0014 | -0.0030 | 0.0020 |
| rs11211123 | A | 0.0104 | 0.0017 | -0.0082 | 0.0024 |
| rs11213482 | A | -0.0106 | 0.0019 | 0.0049 | 0.0026 |
| rs112210983 | A | 0.0105 | 0.0017 | -0.0040 | 0.0024 |
| rs1123285 | C | -0.0094 | 0.0015 | 0.0040 | 0.0021 |
| rs112375785 | A | 0.0184 | 0.0021 | -0.0065 | 0.0029 |
| rs112379405 | C | -0.0082 | 0.0014 | 0.0033 | 0.0020 |
| rs112512729 | T | -0.0166 | 0.0027 | 0.0126 | 0.0040 |
| rs11265191 | T | 0.0083 | 0.0015 | -0.0002 | 0.0021 |
| rs112806496 | C | -0.0160 | 0.0025 | 0.0016 | 0.0035 |
| rs1128956 | T | -0.0140 | 0.0019 | 0.0079 | 0.0026 |
| rs113205706 | A | -0.0135 | 0.0024 | 0.0052 | 0.0034 |
| rs113520408 | A | 0.0155 | 0.0016 | -0.0006 | 0.0022 |
| rs113552169 | C | 0.0294 | 0.0053 | -0.0078 | 0.0076 |
| rs113588399 | T | -0.0104 | 0.0017 | -0.0017 | 0.0025 |
| rs113615161 | T | -0.0123 | 0.0021 | 0.0091 | 0.0029 |
| rs113720505 | A | 0.0164 | 0.0029 | -0.0039 | 0.0041 |
| rs113731629 | T | 0.0339 | 0.0061 | -0.0047 | 0.0087 |
| rs114192810 | T | -0.0279 | 0.0051 | -0.0117 | 0.0071 |
| rs114468556 | A | 0.0293 | 0.0036 | -0.0026 | 0.0048 |
| rs114593137 | A | -0.0122 | 0.0017 | 0.0059 | 0.0025 |
| rs1146079 | C | -0.0087 | 0.0016 | 0.0036 | 0.0023 |
| rs114810763 | A | -0.0121 | 0.0020 | 0.0073 | 0.0027 |
| rs114952970 | T | 0.0265 | 0.0040 | -0.0268 | 0.0060 |
| rs114958262 | T | -0.0220 | 0.0039 | -0.0021 | 0.0055 |
| rs115000530 | A | -0.0250 | 0.0031 | 0.0112 | 0.0044 |
| rs115017135 | A | -0.0206 | 0.0038 | 0.0012 | 0.0055 |
| rs11542663 | A | 0.0112 | 0.0015 | -0.0012 | 0.0022 |
| rs115438240 | T | -0.0169 | 0.0030 | 0.0111 | 0.0045 |
| rs115693355 | A | 0.0309 | 0.0053 | -0.0084 | 0.0075 |
| rs11588857 | A | 0.0199 | 0.0017 | -0.0106 | 0.0024 |
| rs11591870 | T | 0.0098 | 0.0017 | -0.0041 | 0.0024 |
| rs11592299 | A | 0.0097 | 0.0018 | 0.0096 | 0.0025 |
| rs11596387 | T | 0.0258 | 0.0046 | 0.0039 | 0.0067 |
| rs11598765 | A | -0.0106 | 0.0018 | 0.0050 | 0.0025 |
| rs11599236 | T | -0.0106 | 0.0014 | 0.0135 | 0.0020 |
| rs11613431 | A | -0.0094 | 0.0016 | -0.0008 | 0.0023 |
| rs11620355 | A | 0.0182 | 0.0025 | -0.0077 | 0.0036 |
| rs11623285 | T | -0.0135 | 0.0021 | 0.0029 | 0.0029 |
| rs116386746 | C | 0.0249 | 0.0045 | -0.0295 | 0.0065 |
| rs11640569 | A | 0.0112 | 0.0020 | -0.0036 | 0.0029 |
| rs11644446 | A | 0.0117 | 0.0019 | -0.0060 | 0.0028 |
| rs11647188 | A | 0.0079 | 0.0014 | -0.0021 | 0.0020 |
| rs11652522 | A | -0.0154 | 0.0023 | 0.0010 | 0.0033 |
| rs11657342 | A | 0.0140 | 0.0019 | -0.0051 | 0.0021 |
| rs11657979 | A | -0.0091 | 0.0017 | 0.0092 | 0.0024 |
| rs11663602 | A | -0.0130 | 0.0016 | 0.0099 | 0.0022 |
| rs11663678 | T | -0.0121 | 0.0022 | 0.0080 | 0.0031 |
| rs116656374 | T | -0.0242 | 0.0038 | -0.0005 | 0.0052 |
| rs11675476 | T | -0.0127 | 0.0014 | -0.0009 | 0.0020 |
| rs1167827 | A | 0.0104 | 0.0014 | -0.0092 | 0.0020 |
| rs11678980 | A | -0.0166 | 0.0014 | 0.0095 | 0.0021 |
| rs11687736 | A | -0.0105 | 0.0015 | 0.0083 | 0.0021 |
| rs11693885 | A | -0.0091 | 0.0014 | 0.0134 | 0.0020 |
| rs116967397 | A | 0.0315 | 0.0056 | -0.0139 | 0.0077 |
| rs117005905 | T | 0.0150 | 0.0022 | -0.0007 | 0.0032 |
| rs11703948 | A | -0.0168 | 0.0024 | -0.0034 | 0.0033 |
| rs1171040 | A | -0.0113 | 0.0017 | 0.0058 | 0.0025 |
| rs11716398 | A | 0.0159 | 0.0021 | 0.0018 | 0.0030 |
| rs11720093 | T | 0.0150 | 0.0026 | -0.0008 | 0.0036 |
| rs11720985 | T | 0.0085 | 0.0015 | -0.0001 | 0.0022 |
| rs11724690 | T | 0.0093 | 0.0015 | -0.0003 | 0.0022 |
| rs117273411 | T | -0.0211 | 0.0034 | 0.0065 | 0.0050 |
| rs11732160 | A | 0.0105 | 0.0016 | -0.0068 | 0.0023 |
| rs11732657 | A | -0.0126 | 0.0016 | 0.0047 | 0.0023 |
| rs11733439 | A | -0.0096 | 0.0017 | 0.0049 | 0.0025 |
| rs11736863 | A | 0.0142 | 0.0018 | -0.0021 | 0.0026 |
| rs117398064 | C | -0.0141 | 0.0025 | 0.0097 | 0.0034 |
| rs117468730 | A | -0.0310 | 0.0050 | -0.0015 | 0.0071 |
| rs117623407 | A | -0.0118 | 0.0020 | 0.0019 | 0.0028 |
| rs11765387 | T | 0.0086 | 0.0015 | -0.0070 | 0.0021 |
| rs117668569 | T | 0.0234 | 0.0042 | -0.0130 | 0.0062 |
| rs117677995 | A | -0.0180 | 0.0032 | 0.0025 | 0.0046 |
| rs11772232 | T | 0.0177 | 0.0019 | -0.0110 | 0.0027 |
| rs11774212 | T | 0.0128 | 0.0014 | -0.0045 | 0.0020 |
| rs117799466 | C | 0.0106 | 0.0016 | -0.0063 | 0.0022 |
| rs11780023 | T | 0.0085 | 0.0014 | -0.0057 | 0.0020 |
| rs11789013 | T | 0.0106 | 0.0016 | -0.0047 | 0.0023 |
| rs117895796 | T | 0.0362 | 0.0064 | -0.0177 | 0.0097 |
| rs118040169 | A | -0.0248 | 0.0038 | -0.0011 | 0.0053 |
| rs118093058 | T | 0.0146 | 0.0021 | 0.0079 | 0.0029 |
| rs118134876 | T | -0.0223 | 0.0030 | 0.0122 | 0.0042 |
| rs11845781 | T | -0.0079 | 0.0014 | 0.0060 | 0.0020 |
| rs11871429 | A | 0.0128 | 0.0017 | -0.0026 | 0.0024 |
| rs11876620 | T | 0.0143 | 0.0024 | -0.0026 | 0.0033 |
| rs11894424 | A | 0.0209 | 0.0033 | -0.0047 | 0.0048 |
| rs11917701 | A | -0.0087 | 0.0014 | 0.0029 | 0.0020 |
| rs11919835 | C | 0.0089 | 0.0015 | -0.0049 | 0.0021 |
| rs1196760 | C | 0.0139 | 0.0025 | -0.0036 | 0.0035 |
| rs12005151 | A | 0.0087 | 0.0015 | -0.0022 | 0.0021 |
| rs12028010 | T | 0.0164 | 0.0017 | 0.0059 | 0.0024 |
| rs12028229 | T | 0.0099 | 0.0016 | -0.0014 | 0.0023 |
| rs12054166 | C | 0.0096 | 0.0016 | -0.0044 | 0.0023 |
| rs12076635 | C | 0.0207 | 0.0017 | -0.0130 | 0.0024 |
| rs12113634 | T | 0.0088 | 0.0015 | -0.0039 | 0.0021 |
| rs12123293 | T | 0.0086 | 0.0015 | -0.0048 | 0.0021 |
| rs12126231 | A | 0.0095 | 0.0014 | -0.0039 | 0.0020 |
| rs12127928 | T | 0.0108 | 0.0018 | -0.0037 | 0.0025 |
| rs12145078 | A | -0.0127 | 0.0017 | 0.0072 | 0.0025 |
| rs12151248 | T | -0.0149 | 0.0023 | 0.0052 | 0.0031 |
| rs12170452 | A | 0.0105 | 0.0014 | -0.0023 | 0.0020 |
| rs1220779 | A | -0.0079 | 0.0014 | -0.0022 | 0.0020 |
| rs12234369 | A | 0.0087 | 0.0015 | 0.0010 | 0.0022 |
| rs12238011 | T | -0.0141 | 0.0026 | 0.0102 | 0.0036 |
| rs12273435 | A | -0.0132 | 0.0017 | 0.0106 | 0.0025 |
| rs12285074 | A | 0.0161 | 0.0023 | -0.0083 | 0.0035 |
| rs12304188 | A | 0.0111 | 0.0019 | -0.0038 | 0.0028 |
| rs12325727 | A | 0.0088 | 0.0014 | -0.0064 | 0.0020 |
| rs12332731 | A | 0.0135 | 0.0018 | -0.0062 | 0.0025 |
| rs12342546 | A | -0.0128 | 0.0017 | 0.0004 | 0.0024 |
| rs12359372 | T | -0.0092 | 0.0015 | 0.0038 | 0.0021 |
| rs12375949 | T | -0.0138 | 0.0014 | 0.0072 | 0.0020 |
| rs12405889 | T | -0.0098 | 0.0014 | 0.0000 | 0.0020 |
| rs12431682 | T | -0.0093 | 0.0015 | 0.0017 | 0.0021 |
| rs12438177 | A | 0.0093 | 0.0015 | -0.0012 | 0.0021 |
| rs12453682 | T | 0.0102 | 0.0015 | -0.0111 | 0.0022 |
| rs1245829 | A | -0.0099 | 0.0014 | 0.0047 | 0.0020 |
| rs12467175 | T | -0.0081 | 0.0014 | 0.0020 | 0.0020 |
| rs12468040 | T | 0.0137 | 0.0014 | -0.0057 | 0.0021 |
| rs12473986 | T | -0.0085 | 0.0015 | 0.0071 | 0.0022 |
| rs12477385 | T | 0.0109 | 0.0017 | -0.0103 | 0.0024 |
| rs12478156 | T | 0.0108 | 0.0015 | -0.0006 | 0.0021 |
| rs12503522 | T | -0.0113 | 0.0016 | 0.0082 | 0.0022 |
| rs12506221 | T | -0.0130 | 0.0014 | 0.0076 | 0.0020 |
| rs12516990 | C | -0.0099 | 0.0017 | 0.0032 | 0.0025 |
| rs12519073 | T | -0.0107 | 0.0017 | 0.0065 | 0.0024 |
| rs12524795 | T | 0.0102 | 0.0014 | -0.0025 | 0.0020 |
| rs12568153 | T | -0.0091 | 0.0015 | -0.0078 | 0.0022 |
| rs12571549 | A | 0.0151 | 0.0020 | 0.0010 | 0.0029 |
| rs12574281 | A | -0.0080 | 0.0015 | -0.0003 | 0.0021 |
| rs12591647 | T | -0.0152 | 0.0018 | 0.0030 | 0.0026 |
| rs12601380 | A | -0.0082 | 0.0014 | 0.0050 | 0.0020 |
| rs12602286 | T | 0.0160 | 0.0021 | -0.0051 | 0.0030 |
| rs12613500 | C | 0.0098 | 0.0014 | -0.0006 | 0.0020 |
| rs12614263 | A | 0.0083 | 0.0014 | -0.0025 | 0.0020 |
| rs12638072 | A | -0.0107 | 0.0015 | 0.0038 | 0.0022 |
| rs12643771 | T | 0.0130 | 0.0015 | -0.0034 | 0.0022 |
| rs12646216 | T | 0.0084 | 0.0014 | -0.0042 | 0.0020 |
| rs12646523 | T | -0.0132 | 0.0016 | 0.0003 | 0.0023 |
| rs12659776 | T | 0.0085 | 0.0014 | -0.0097 | 0.0020 |
| rs12670376 | A | 0.0092 | 0.0014 | -0.0014 | 0.0020 |
| rs1267062 | C | 0.0113 | 0.0018 | -0.0008 | 0.0026 |
| rs12682775 | T | -0.0119 | 0.0017 | 0.0072 | 0.0024 |
| rs12694681 | T | 0.0102 | 0.0015 | -0.0029 | 0.0021 |
| rs12709186 | A | -0.0091 | 0.0015 | 0.0063 | 0.0021 |
| rs12712269 | T | -0.0107 | 0.0014 | 0.0018 | 0.0020 |
| rs12746551 | C | 0.0248 | 0.0041 | -0.0099 | 0.0057 |
| rs12750688 | C | 0.0116 | 0.0016 | 0.0006 | 0.0022 |
| rs12761729 | T | -0.0098 | 0.0016 | 0.0047 | 0.0023 |
| rs12761761 | T | 0.0153 | 0.0017 | -0.0036 | 0.0023 |
| rs12764593 | C | 0.0202 | 0.0029 | -0.0167 | 0.0043 |
| rs12765185 | A | -0.0088 | 0.0016 | 0.0070 | 0.0022 |
| rs12774577 | T | 0.0142 | 0.0020 | -0.0175 | 0.0028 |
| rs12789313 | T | 0.0092 | 0.0014 | 0.0003 | 0.0020 |
| rs12790196 | T | -0.0101 | 0.0015 | 0.0063 | 0.0021 |
| rs12810587 | T | 0.0094 | 0.0015 | 0.0045 | 0.0021 |
| rs12875339 | A | -0.0108 | 0.0015 | 0.0073 | 0.0021 |
| rs12888615 | T | 0.0120 | 0.0018 | -0.0010 | 0.0025 |
| rs12891042 | T | 0.0092 | 0.0014 | 0.0000 | 0.0020 |
| rs12891191 | C | 0.0080 | 0.0014 | -0.0020 | 0.0020 |
| rs12912465 | T | -0.0104 | 0.0016 | 0.0016 | 0.0023 |
| rs12952191 | T | -0.0081 | 0.0014 | 0.0033 | 0.0020 |
| rs12953422 | A | -0.0080 | 0.0014 | 0.0024 | 0.0020 |
| rs12957463 | A | 0.0146 | 0.0018 | -0.0053 | 0.0025 |
| rs12962845 | C | -0.0123 | 0.0021 | 0.0008 | 0.0029 |
| rs12967010 | T | 0.0114 | 0.0017 | -0.0058 | 0.0024 |
| rs12970264 | A | -0.0085 | 0.0014 | 0.0086 | 0.0020 |
| rs12981405 | T | -0.0112 | 0.0019 | 0.0065 | 0.0027 |
| rs13009915 | T | 0.0212 | 0.0037 | 0.0018 | 0.0052 |
| rs13010288 | T | 0.0198 | 0.0021 | -0.0098 | 0.0029 |
| rs13010566 | A | -0.0091 | 0.0014 | -0.0001 | 0.0020 |
| rs13015496 | C | 0.0119 | 0.0018 | -0.0112 | 0.0026 |
| rs13016316 | A | -0.0138 | 0.0024 | 0.0090 | 0.0034 |
| rs1301838 | T | -0.0096 | 0.0015 | 0.0006 | 0.0021 |
| rs13018640 | T | -0.0215 | 0.0014 | 0.0050 | 0.0020 |
| rs13026611 | A | -0.0084 | 0.0015 | 0.0009 | 0.0021 |
| rs13029602 | T | -0.0092 | 0.0014 | -0.0033 | 0.0020 |
| rs13034349 | T | -0.0082 | 0.0014 | 0.0060 | 0.0020 |
| rs13050131 | A | -0.0081 | 0.0015 | 0.0062 | 0.0021 |
| rs13085461 | C | 0.0103 | 0.0014 | -0.0050 | 0.0020 |
| rs13099165 | T | -0.0118 | 0.0017 | 0.0026 | 0.0025 |
| rs13107325 | T | -0.0239 | 0.0027 | -0.0057 | 0.0038 |
| rs13117856 | A | 0.0084 | 0.0015 | -0.0030 | 0.0022 |
| rs13130765 | C | -0.0091 | 0.0014 | 0.0013 | 0.0020 |
| rs13133213 | A | 0.0096 | 0.0014 | -0.0044 | 0.0020 |
| rs13145650 | T | -0.0173 | 0.0025 | 0.0113 | 0.0036 |
| rs13154429 | C | -0.0136 | 0.0023 | 0.0050 | 0.0032 |
| rs13163062 | T | 0.0115 | 0.0014 | -0.0056 | 0.0020 |
| rs13163845 | T | -0.0149 | 0.0020 | 0.0127 | 0.0029 |
| rs13168136 | A | -0.0092 | 0.0017 | 0.0025 | 0.0024 |
| rs13169187 | A | -0.0079 | 0.0014 | 0.0037 | 0.0020 |
| rs13177031 | A | -0.0085 | 0.0015 | 0.0017 | 0.0021 |
| rs13190235 | T | -0.0138 | 0.0024 | 0.0003 | 0.0034 |
| rs13197257 | T | 0.0109 | 0.0016 | -0.0052 | 0.0023 |
| rs1320139 | C | -0.0149 | 0.0014 | 0.0013 | 0.0020 |
| rs13212041 | T | 0.0101 | 0.0018 | -0.0007 | 0.0025 |
| rs13240401 | T | 0.0176 | 0.0017 | -0.0025 | 0.0024 |
| rs13246220 | T | -0.0093 | 0.0014 | -0.0002 | 0.0020 |
| rs13261773 | C | 0.0122 | 0.0019 | -0.0025 | 0.0028 |
| rs13266287 | A | -0.0085 | 0.0015 | 0.0074 | 0.0021 |
| rs13281564 | A | 0.0088 | 0.0014 | -0.0032 | 0.0020 |
| rs13284516 | C | -0.0082 | 0.0014 | 0.0042 | 0.0020 |
| rs1329044 | T | 0.0134 | 0.0019 | -0.0049 | 0.0026 |
| rs1329125 | T | -0.0097 | 0.0015 | 0.0065 | 0.0021 |
| rs13296345 | T | 0.0090 | 0.0015 | -0.0035 | 0.0021 |
| rs13318986 | A | 0.0124 | 0.0015 | -0.0033 | 0.0021 |
| rs13327482 | A | -0.0121 | 0.0018 | 0.0049 | 0.0026 |
| rs1334297 | A | 0.0257 | 0.0016 | -0.0101 | 0.0023 |
| rs1335482 | T | 0.0096 | 0.0014 | -0.0057 | 0.0020 |
| rs13361043 | T | -0.0158 | 0.0028 | 0.0037 | 0.0040 |
| rs13381557 | A | -0.0087 | 0.0014 | 0.0014 | 0.0020 |
| rs13388333 | A | 0.0085 | 0.0015 | 0.0002 | 0.0021 |
| rs13397529 | C | -0.0097 | 0.0017 | 0.0059 | 0.0024 |
| rs13398860 | A | 0.0086 | 0.0015 | -0.0086 | 0.0021 |
| rs13402497 | A | -0.0080 | 0.0014 | 0.0006 | 0.0020 |
| rs13422673 | T | -0.0110 | 0.0014 | 0.0021 | 0.0020 |
| rs13425585 | C | -0.0089 | 0.0014 | 0.0014 | 0.0020 |
| rs1368250 | T | 0.0250 | 0.0045 | -0.0061 | 0.0058 |
| rs137079 | T | 0.0128 | 0.0020 | -0.0080 | 0.0028 |
| rs137858393 | A | 0.0193 | 0.0029 | 0.0015 | 0.0040 |
| rs138096147 | A | 0.0234 | 0.0042 | -0.0066 | 0.0060 |
| rs138484388 | T | -0.0306 | 0.0051 | 0.0001 | 0.0069 |
| rs1391513 | T | -0.0095 | 0.0015 | 0.0031 | 0.0022 |
| rs139244147 | A | -0.0193 | 0.0030 | 0.0059 | 0.0044 |
| rs1392816 | T | 0.0082 | 0.0014 | -0.0142 | 0.0021 |
| rs139980871 | T | 0.0180 | 0.0029 | 0.0045 | 0.0040 |
| rs1404549 | A | -0.0089 | 0.0015 | 0.0006 | 0.0021 |
| rs1405876 | T | 0.0109 | 0.0015 | 0.0069 | 0.0021 |
| rs140711597 | C | 0.0366 | 0.0055 | -0.0140 | 0.0071 |
| rs1408284 | C | -0.0127 | 0.0020 | 0.0038 | 0.0029 |
| rs1408430 | C | 0.0096 | 0.0016 | -0.0032 | 0.0022 |
| rs141729694 | T | 0.0195 | 0.0026 | 0.0023 | 0.0038 |
| rs142014757 | A | 0.0102 | 0.0018 | -0.0092 | 0.0025 |
| rs1426619 | T | 0.0096 | 0.0014 | -0.0047 | 0.0020 |
| rs1427298 | T | 0.0086 | 0.0014 | -0.0031 | 0.0020 |
| rs142747148 | A | 0.0329 | 0.0052 | -0.0059 | 0.0076 |
| rs1427829 | A | -0.0083 | 0.0014 | 0.0078 | 0.0020 |
| rs143148393 | T | -0.0211 | 0.0039 | 0.0028 | 0.0059 |
| rs1434630 | T | -0.0137 | 0.0020 | 0.0055 | 0.0028 |
| rs143743568 | A | 0.0118 | 0.0020 | -0.0052 | 0.0028 |
| rs143812851 | A | -0.0113 | 0.0019 | -0.0011 | 0.0026 |
| rs1440930 | C | 0.0081 | 0.0014 | -0.0015 | 0.0020 |
| rs144336753 | A | 0.0388 | 0.0053 | -0.0274 | 0.0073 |
| rs1445591 | A | 0.0085 | 0.0015 | -0.0001 | 0.0022 |
| rs1464297 | T | -0.0109 | 0.0015 | -0.0003 | 0.0021 |
| rs1467737 | T | -0.0084 | 0.0014 | 0.0038 | 0.0020 |
| rs1485300 | C | 0.0098 | 0.0015 | 0.0029 | 0.0021 |
| rs1490612 | T | 0.0193 | 0.0026 | 0.0049 | 0.0038 |
| rs150252215 | A | 0.0204 | 0.0035 | 0.0023 | 0.0050 |
| rs150421637 | A | 0.0349 | 0.0061 | -0.0142 | 0.0082 |
| rs150537577 | A | -0.0151 | 0.0026 | -0.0020 | 0.0038 |
| rs1505676 | C | 0.0084 | 0.0015 | -0.0031 | 0.0021 |
| rs151381 | T | -0.0087 | 0.0014 | -0.0003 | 0.0020 |
| rs1518890 | T | -0.0090 | 0.0016 | 0.0002 | 0.0023 |
| rs1527878 | A | -0.0105 | 0.0016 | 0.0033 | 0.0023 |
| rs1529597 | A | 0.0223 | 0.0038 | -0.0103 | 0.0058 |
| rs1538389 | T | -0.0109 | 0.0018 | 0.0072 | 0.0025 |
| rs1542354 | A | -0.0084 | 0.0014 | 0.0032 | 0.0020 |
| rs1544 | A | -0.0096 | 0.0016 | 0.0052 | 0.0023 |
| rs1554798 | A | -0.0080 | 0.0014 | -0.0022 | 0.0020 |
| rs1564347 | T | 0.0088 | 0.0015 | -0.0044 | 0.0021 |
| rs1569092 | A | 0.0160 | 0.0020 | -0.0088 | 0.0027 |
| rs1593022 | T | -0.0099 | 0.0017 | 0.0032 | 0.0024 |
| rs1603460 | T | 0.0090 | 0.0014 | -0.0012 | 0.0020 |
| rs162445 | A | 0.0152 | 0.0026 | -0.0064 | 0.0037 |
| rs163229 | C | -0.0260 | 0.0047 | 0.0083 | 0.0067 |
| rs1637770 | T | 0.0191 | 0.0028 | -0.0063 | 0.0041 |
| rs164938 | T | -0.0088 | 0.0014 | 0.0028 | 0.0020 |
| rs165633 | A | -0.0116 | 0.0016 | -0.0004 | 0.0024 |
| rs1656614 | C | 0.0122 | 0.0015 | -0.0106 | 0.0022 |
| rs1671269 | T | -0.0100 | 0.0016 | 0.0062 | 0.0023 |
| rs1671770 | A | 0.0125 | 0.0018 | -0.0041 | 0.0026 |
| rs16851779 | T | -0.0153 | 0.0027 | 0.0014 | 0.0036 |
| rs16871807 | T | 0.0090 | 0.0015 | -0.0085 | 0.0021 |
| rs1689510 | C | 0.0194 | 0.0015 | -0.0095 | 0.0021 |
| rs16901689 | T | 0.0111 | 0.0019 | -0.0073 | 0.0027 |
| rs1693584 | T | -0.0087 | 0.0015 | 0.0031 | 0.0022 |
| rs16958559 | T | 0.0096 | 0.0017 | -0.0085 | 0.0024 |
| rs16966271 | T | 0.0092 | 0.0016 | -0.0020 | 0.0022 |
| rs16975275 | A | -0.0102 | 0.0017 | 0.0057 | 0.0024 |
| rs17048801 | A | -0.0106 | 0.0014 | 0.0049 | 0.0021 |
| rs17048855 | A | 0.0108 | 0.0015 | -0.0017 | 0.0021 |
| rs17069646 | T | 0.0085 | 0.0015 | -0.0069 | 0.0022 |
| rs17110109 | T | -0.0093 | 0.0014 | 0.0055 | 0.0020 |
| rs17113730 | A | -0.0170 | 0.0027 | 0.0032 | 0.0039 |
| rs17131123 | A | -0.0243 | 0.0039 | 0.0115 | 0.0052 |
| rs17133297 | A | -0.0146 | 0.0023 | 0.0147 | 0.0034 |
| rs17144467 | A | 0.0098 | 0.0015 | 0.0020 | 0.0021 |
| rs17148998 | A | 0.0101 | 0.0017 | -0.0047 | 0.0025 |
| rs171697 | C | 0.0133 | 0.0015 | -0.0039 | 0.0021 |
| rs17170519 | C | 0.0132 | 0.0023 | -0.0069 | 0.0032 |
| rs1717204 | A | -0.0117 | 0.0018 | 0.0072 | 0.0026 |
| rs1718188 | A | 0.0096 | 0.0014 | -0.0051 | 0.0020 |
| rs17186106 | T | 0.0108 | 0.0018 | -0.0036 | 0.0026 |
| rs17190418 | T | -0.0213 | 0.0031 | 0.0044 | 0.0043 |
| rs17205908 | T | -0.0092 | 0.0015 | 0.0032 | 0.0021 |
| rs17248751 | A | -0.0136 | 0.0017 | 0.0045 | 0.0024 |
| rs1728118 | A | 0.0089 | 0.0016 | -0.0025 | 0.0022 |
| rs1729412 | T | -0.0107 | 0.0014 | 0.0015 | 0.0020 |
| rs173003 | A | -0.0082 | 0.0014 | 0.0075 | 0.0020 |
| rs1734370 | A | 0.0099 | 0.0016 | -0.0012 | 0.0022 |
| rs1738050 | C | -0.0093 | 0.0014 | 0.0055 | 0.0021 |
| rs17411339 | A | 0.0140 | 0.0014 | -0.0087 | 0.0020 |
| rs17428076 | C | 0.0137 | 0.0016 | -0.0007 | 0.0023 |
| rs17440885 | A | 0.0160 | 0.0029 | -0.0085 | 0.0041 |
| rs1747714 | T | 0.0106 | 0.0014 | -0.0032 | 0.0020 |
| rs1747817 | T | -0.0092 | 0.0016 | 0.0092 | 0.0023 |
| rs17489649 | A | 0.0127 | 0.0015 | -0.0071 | 0.0021 |
| rs17502934 | T | -0.0167 | 0.0020 | 0.0080 | 0.0028 |
| rs175325 | A | -0.0100 | 0.0014 | 0.0019 | 0.0020 |
| rs17536059 | C | -0.0108 | 0.0019 | 0.0018 | 0.0026 |
| rs17551064 | A | 0.0141 | 0.0019 | -0.0075 | 0.0027 |
| rs17563464 | A | -0.0147 | 0.0017 | 0.0027 | 0.0024 |
| rs17565975 | A | -0.0106 | 0.0014 | 0.0035 | 0.0020 |
| rs17568389 | A | 0.0127 | 0.0014 | -0.0012 | 0.0020 |
| rs17574007 | A | -0.0129 | 0.0021 | 0.0089 | 0.0031 |
| rs1758747 | A | 0.0092 | 0.0015 | -0.0009 | 0.0022 |
| rs17598675 | T | -0.0116 | 0.0014 | 0.0019 | 0.0020 |
| rs17604349 | A | 0.0127 | 0.0018 | 0.0082 | 0.0026 |
| rs17609255 | T | -0.0087 | 0.0014 | 0.0102 | 0.0020 |
| rs176218 | T | 0.0200 | 0.0018 | -0.0054 | 0.0025 |
| rs17622379 | T | -0.0113 | 0.0018 | -0.0013 | 0.0025 |
| rs17650634 | T | -0.0104 | 0.0019 | -0.0005 | 0.0027 |
| rs17667540 | A | -0.0087 | 0.0015 | 0.0043 | 0.0022 |
| rs17669337 | T | -0.0110 | 0.0014 | 0.0009 | 0.0020 |
| rs17680712 | T | -0.0092 | 0.0014 | 0.0028 | 0.0020 |
| rs17686649 | T | -0.0098 | 0.0017 | -0.0014 | 0.0024 |
| rs17732878 | T | -0.0095 | 0.0017 | 0.0053 | 0.0024 |
| rs17742342 | A | -0.0119 | 0.0018 | 0.0059 | 0.0025 |
| rs17747544 | A | -0.0090 | 0.0014 | -0.0001 | 0.0020 |
| rs1779549 | A | 0.0081 | 0.0014 | -0.0026 | 0.0020 |
| rs178183 | T | 0.0145 | 0.0016 | -0.0058 | 0.0023 |
| rs17882802 | A | 0.0084 | 0.0014 | -0.0037 | 0.0020 |
| rs17883331 | A | -0.0108 | 0.0018 | 0.0089 | 0.0025 |
| rs1792602 | A | -0.0097 | 0.0014 | 0.0026 | 0.0020 |
| rs181214 | T | -0.0104 | 0.0017 | 0.0055 | 0.0025 |
| rs182902112 | A | -0.0300 | 0.0053 | 0.0111 | 0.0076 |
| rs1835339 | T | 0.0084 | 0.0014 | -0.0024 | 0.0021 |
| rs183869217 | C | 0.0174 | 0.0030 | -0.0114 | 0.0039 |
| rs1841023 | A | -0.0092 | 0.0015 | 0.0034 | 0.0022 |
| rs1842713 | A | -0.0117 | 0.0017 | 0.0053 | 0.0024 |
| rs1843815 | A | 0.0077 | 0.0014 | -0.0080 | 0.0020 |
| rs1861786 | A | -0.0081 | 0.0014 | -0.0016 | 0.0021 |
| rs186456786 | A | 0.0455 | 0.0079 | -0.0167 | 0.0104 |
| rs1865955 | T | 0.0122 | 0.0019 | -0.0093 | 0.0026 |
| rs1866823 | A | 0.0107 | 0.0014 | -0.0096 | 0.0020 |
| rs187580 | T | -0.0104 | 0.0017 | 0.0002 | 0.0023 |
| rs187951956 | A | -0.0315 | 0.0058 | -0.0035 | 0.0089 |
| rs1880088 | A | -0.0097 | 0.0016 | 0.0006 | 0.0023 |
| rs1880692 | A | 0.0086 | 0.0014 | -0.0024 | 0.0020 |
| rs1890132 | T | -0.0089 | 0.0015 | 0.0050 | 0.0022 |
| rs1898111 | A | 0.0101 | 0.0018 | -0.0138 | 0.0026 |
| rs190102446 | T | -0.0210 | 0.0037 | 0.0018 | 0.0054 |
| rs1905616 | A | 0.0083 | 0.0015 | -0.0034 | 0.0021 |
| rs1910005 | T | -0.0090 | 0.0015 | 0.0014 | 0.0022 |
| rs1918394 | T | 0.0119 | 0.0019 | -0.0070 | 0.0027 |
| rs191903670 | A | 0.0335 | 0.0061 | -0.0238 | 0.0094 |
| rs192436652 | T | -0.0319 | 0.0045 | 0.0162 | 0.0061 |
| rs1933264 | T | 0.0100 | 0.0016 | -0.0084 | 0.0023 |
| rs1952183 | A | -0.0089 | 0.0014 | 0.0052 | 0.0020 |
| rs1955250 | A | -0.0140 | 0.0025 | 0.0028 | 0.0036 |
| rs1963381 | A | 0.0099 | 0.0016 | 0.0007 | 0.0023 |
| rs1980129 | A | 0.0085 | 0.0014 | -0.0013 | 0.0020 |
| rs198262 | T | -0.0223 | 0.0035 | 0.0010 | 0.0050 |
| rs1991585 | T | -0.0101 | 0.0015 | 0.0003 | 0.0022 |
| rs1995181 | A | -0.0078 | 0.0014 | 0.0054 | 0.0020 |
| rs2007655 | T | 0.0086 | 0.0014 | -0.0024 | 0.0020 |
| rs2011603 | A | -0.0091 | 0.0016 | -0.0018 | 0.0023 |
| rs2014830 | T | 0.0110 | 0.0015 | -0.0047 | 0.0022 |
| rs2023016 | C | 0.0107 | 0.0017 | -0.0038 | 0.0024 |
| rs2024568 | T | -0.0097 | 0.0016 | 0.0051 | 0.0023 |
| rs2029401 | A | -0.0091 | 0.0014 | -0.0016 | 0.0020 |
| rs2034631 | T | 0.0089 | 0.0015 | -0.0052 | 0.0021 |
| rs2039204 | A | 0.0082 | 0.0014 | -0.0014 | 0.0020 |
| rs2043187 | A | 0.0107 | 0.0015 | -0.0065 | 0.0022 |
| rs2052285 | A | 0.0112 | 0.0014 | -0.0043 | 0.0020 |
| rs2055940 | A | 0.0082 | 0.0015 | -0.0017 | 0.0021 |
| rs2077235 | T | 0.0097 | 0.0017 | -0.0034 | 0.0023 |
| rs2081652 | A | 0.0123 | 0.0015 | -0.0013 | 0.0021 |
| rs2082317 | T | -0.0078 | 0.0014 | 0.0014 | 0.0020 |
| rs2088913 | A | 0.0081 | 0.0014 | -0.0021 | 0.0020 |
| rs2092248 | T | -0.0083 | 0.0015 | 0.0030 | 0.0022 |
| rs2098526 | A | -0.0241 | 0.0043 | 0.0073 | 0.0061 |
| rs2100249 | T | -0.0089 | 0.0015 | 0.0037 | 0.0021 |
| rs2126069 | T | 0.0080 | 0.0014 | 0.0036 | 0.0020 |
| rs2131167 | A | 0.0080 | 0.0014 | -0.0108 | 0.0020 |
| rs214626 | A | 0.0101 | 0.0018 | -0.0048 | 0.0026 |
| rs2160514 | A | -0.0094 | 0.0014 | 0.0100 | 0.0020 |
| rs2179152 | T | -0.0131 | 0.0015 | 0.0130 | 0.0021 |
| rs2183271 | T | 0.0084 | 0.0015 | -0.0092 | 0.0021 |
| rs2199409 | T | 0.0103 | 0.0017 | -0.0059 | 0.0025 |
| rs2212430 | T | -0.0100 | 0.0016 | 0.0029 | 0.0022 |
| rs2216144 | T | 0.0098 | 0.0014 | -0.0018 | 0.0020 |
| rs2220926 | T | -0.0095 | 0.0014 | 0.0040 | 0.0020 |
| rs2250660 | C | 0.0080 | 0.0014 | -0.0016 | 0.0020 |
| rs2252098 | T | -0.0077 | 0.0014 | 0.0044 | 0.0020 |
| rs2254681 | A | 0.0101 | 0.0016 | 0.0005 | 0.0023 |
| rs2256965 | A | 0.0106 | 0.0014 | -0.0020 | 0.0020 |
| rs2276209 | A | 0.0085 | 0.0015 | -0.0037 | 0.0022 |
| rs2283076 | A | 0.0115 | 0.0017 | -0.0076 | 0.0024 |
| rs2287838 | A | -0.0104 | 0.0014 | 0.0017 | 0.0020 |
| rs2290601 | T | -0.0149 | 0.0017 | 0.0063 | 0.0024 |
| rs2297293 | C | 0.0099 | 0.0015 | 0.0009 | 0.0022 |
| rs2299156 | T | 0.0099 | 0.0018 | -0.0018 | 0.0026 |
| rs2314338 | T | 0.0093 | 0.0016 | -0.0066 | 0.0022 |
| rs2321157 | A | -0.0081 | 0.0014 | 0.0008 | 0.0020 |
| rs232496 | T | 0.0090 | 0.0015 | 0.0019 | 0.0021 |
| rs2332179 | A | 0.0121 | 0.0020 | -0.0025 | 0.0029 |
| rs2336721 | T | 0.0095 | 0.0015 | -0.0089 | 0.0021 |
| rs2358628 | A | -0.0091 | 0.0015 | 0.0034 | 0.0022 |
| rs236318 | C | -0.0102 | 0.0017 | 0.0051 | 0.0024 |
| rs2364544 | A | -0.0105 | 0.0014 | 0.0002 | 0.0020 |
| rs2365376 | A | 0.0092 | 0.0015 | -0.0038 | 0.0021 |
| rs2368831 | T | -0.0092 | 0.0014 | 0.0048 | 0.0020 |
| rs2373124 | A | -0.0096 | 0.0016 | 0.0014 | 0.0024 |
| rs2406253 | A | 0.0135 | 0.0018 | -0.0016 | 0.0025 |
| rs2414072 | A | -0.0083 | 0.0014 | 0.0067 | 0.0020 |
| rs2416214 | A | 0.0079 | 0.0014 | -0.0021 | 0.0020 |
| rs2416759 | A | -0.0093 | 0.0015 | 0.0069 | 0.0022 |
| rs2416845 | T | 0.0263 | 0.0048 | -0.0077 | 0.0068 |
| rs2431023 | A | 0.0093 | 0.0014 | 0.0005 | 0.0020 |
| rs2434672 | A | 0.0091 | 0.0014 | -0.0051 | 0.0020 |
| rs2436760 | T | 0.0232 | 0.0042 | -0.0190 | 0.0058 |
| rs2447097 | T | 0.0096 | 0.0014 | -0.0070 | 0.0020 |
| rs2458370 | T | -0.0086 | 0.0015 | 0.0037 | 0.0021 |
| rs2469226 | A | 0.0096 | 0.0017 | -0.0075 | 0.0024 |
| rs2470966 | C | -0.0103 | 0.0015 | 0.0078 | 0.0022 |
| rs2478208 | C | -0.0103 | 0.0014 | 0.0044 | 0.0020 |
| rs2496482 | T | 0.0121 | 0.0015 | -0.0059 | 0.0020 |
| rs2517086 | C | -0.0088 | 0.0014 | 0.0057 | 0.0020 |
| rs2521602 | A | -0.0266 | 0.0049 | 0.0219 | 0.0067 |
| rs2522545 | T | -0.0083 | 0.0014 | -0.0024 | 0.0020 |
| rs2526398 | C | -0.0222 | 0.0015 | 0.0105 | 0.0021 |
| rs2529069 | T | -0.0106 | 0.0016 | 0.0056 | 0.0023 |
| rs252991 | A | 0.0095 | 0.0015 | -0.0049 | 0.0021 |
| rs2542673 | A | 0.0088 | 0.0015 | 0.0007 | 0.0021 |
| rs2545795 | A | 0.0125 | 0.0014 | -0.0064 | 0.0020 |
| rs255053 | A | -0.0111 | 0.0018 | -0.0006 | 0.0025 |
| rs2554835 | A | 0.0088 | 0.0014 | -0.0020 | 0.0020 |
| rs2568955 | T | -0.0165 | 0.0016 | 0.0055 | 0.0024 |
| rs2569041 | T | 0.0077 | 0.0014 | -0.0014 | 0.0020 |
| rs2570497 | T | -0.0121 | 0.0015 | 0.0089 | 0.0021 |
| rs2588959 | T | -0.0080 | 0.0014 | 0.0119 | 0.0020 |
| rs2657283 | T | 0.0085 | 0.0014 | -0.0046 | 0.0020 |
| rs2665668 | A | -0.0094 | 0.0015 | 0.0055 | 0.0021 |
| rs2702576 | A | -0.0090 | 0.0014 | -0.0005 | 0.0021 |
| rs2718277 | T | 0.0142 | 0.0026 | -0.0117 | 0.0035 |
| rs2718791 | T | -0.0081 | 0.0015 | 0.0082 | 0.0021 |
| rs27220 | A | -0.0114 | 0.0015 | 0.0036 | 0.0021 |
| rs2725370 | T | -0.0141 | 0.0015 | 0.0060 | 0.0022 |
| rs2736752 | T | 0.0095 | 0.0017 | 0.0006 | 0.0024 |
| rs2740795 | A | 0.0098 | 0.0016 | -0.0098 | 0.0023 |
| rs2761438 | A | 0.0098 | 0.0015 | -0.0049 | 0.0021 |
| rs2764684 | T | 0.0143 | 0.0019 | -0.0048 | 0.0026 |
| rs2805064 | C | 0.0093 | 0.0016 | -0.0022 | 0.0022 |
| rs2838006 | T | 0.0132 | 0.0015 | -0.0082 | 0.0021 |
| rs28482086 | A | 0.0086 | 0.0015 | -0.0037 | 0.0021 |
| rs28505285 | C | 0.0106 | 0.0018 | -0.0096 | 0.0026 |
| rs28512462 | C | 0.0112 | 0.0014 | -0.0111 | 0.0021 |
| rs28513882 | A | -0.0110 | 0.0018 | 0.0000 | 0.0026 |
| rs2852349 | T | -0.0093 | 0.0014 | 0.0048 | 0.0020 |
| rs28587776 | T | 0.0088 | 0.0015 | -0.0033 | 0.0021 |
| rs28669886 | A | -0.0084 | 0.0015 | 0.0003 | 0.0021 |
| rs2870281 | A | -0.0083 | 0.0014 | -0.0003 | 0.0020 |
| rs28735993 | A | 0.0162 | 0.0022 | -0.0003 | 0.0031 |
| rs2885198 | A | 0.0088 | 0.0014 | 0.0015 | 0.0020 |
| rs2898191 | A | 0.0093 | 0.0015 | 0.0016 | 0.0021 |
| rs2910823 | T | 0.0115 | 0.0014 | -0.0021 | 0.0020 |
| rs2916490 | A | -0.0092 | 0.0015 | 0.0017 | 0.0022 |
| rs2923424 | A | 0.0120 | 0.0014 | -0.0078 | 0.0020 |
| rs2926702 | T | -0.0124 | 0.0021 | -0.0051 | 0.0030 |
| rs2929032 | A | -0.0081 | 0.0014 | 0.0006 | 0.0020 |
| rs2929860 | T | 0.0110 | 0.0018 | -0.0076 | 0.0025 |
| rs2942884 | A | -0.0090 | 0.0014 | 0.0033 | 0.0020 |
| rs2954114 | A | -0.0081 | 0.0014 | 0.0052 | 0.0020 |
| rs2958182 | A | 0.0082 | 0.0015 | -0.0048 | 0.0021 |
| rs2962378 | A | 0.0101 | 0.0017 | 0.0032 | 0.0024 |
| rs2964199 | T | -0.0099 | 0.0015 | -0.0006 | 0.0021 |
| rs2964255 | A | 0.0088 | 0.0015 | 0.0029 | 0.0022 |
| rs2976397 | T | 0.0081 | 0.0014 | -0.0007 | 0.0020 |
| rs2977464 | T | 0.0102 | 0.0018 | -0.0015 | 0.0026 |
| rs29792 | A | -0.0093 | 0.0015 | -0.0002 | 0.0022 |
| rs2980813 | A | 0.0088 | 0.0014 | -0.0047 | 0.0020 |
| rs2989476 | C | 0.0078 | 0.0014 | -0.0025 | 0.0020 |
| rs2989751 | A | -0.0092 | 0.0017 | 0.0032 | 0.0023 |
| rs2992037 | A | 0.0149 | 0.0015 | -0.0094 | 0.0022 |
| rs2994326 | T | -0.0100 | 0.0018 | 0.0023 | 0.0026 |
| rs2998299 | T | -0.0136 | 0.0017 | 0.0021 | 0.0024 |
| rs3026996 | A | 0.0138 | 0.0016 | -0.0051 | 0.0023 |
| rs303752 | A | -0.0110 | 0.0014 | -0.0017 | 0.0021 |
| rs3111251 | T | -0.0090 | 0.0014 | 0.0024 | 0.0020 |
| rs312927 | A | -0.0147 | 0.0022 | 0.0051 | 0.0031 |
| rs312945 | A | -0.0091 | 0.0015 | 0.0029 | 0.0022 |
| rs317050 | T | 0.0078 | 0.0014 | -0.0122 | 0.0020 |
| rs322614 | A | -0.0087 | 0.0014 | 0.0046 | 0.0021 |
| rs322744 | T | -0.0090 | 0.0016 | 0.0010 | 0.0023 |
| rs324885 | A | 0.0142 | 0.0014 | -0.0066 | 0.0020 |
| rs32940 | T | -0.0096 | 0.0015 | 0.0062 | 0.0022 |
| rs339057 | A | -0.0081 | 0.0014 | 0.0005 | 0.0020 |
| rs34067381 | T | -0.0104 | 0.0015 | -0.0009 | 0.0021 |
| rs34098770 | A | -0.0160 | 0.0020 | 0.0021 | 0.0028 |
| rs34106693 | C | 0.0159 | 0.0019 | -0.0049 | 0.0027 |
| rs34122915 | C | 0.0084 | 0.0014 | -0.0026 | 0.0021 |
| rs341504 | A | 0.0099 | 0.0015 | 0.0006 | 0.0021 |
| rs34155847 | A | 0.0135 | 0.0016 | -0.0059 | 0.0023 |
| rs34262657 | T | -0.0240 | 0.0038 | 0.0008 | 0.0057 |
| rs34286836 | T | -0.0089 | 0.0014 | 0.0039 | 0.0020 |
| rs34298584 | A | 0.0111 | 0.0019 | -0.0065 | 0.0027 |
| rs34305371 | A | 0.0314 | 0.0024 | -0.0063 | 0.0033 |
| rs34309 | A | 0.0080 | 0.0014 | -0.0015 | 0.0021 |
| rs34316274 | A | 0.0097 | 0.0017 | -0.0010 | 0.0023 |
| rs34363861 | A | 0.0087 | 0.0014 | -0.0041 | 0.0020 |
| rs34394051 | A | -0.0117 | 0.0020 | 0.0049 | 0.0027 |
| rs34410 | C | -0.0083 | 0.0014 | 0.0013 | 0.0020 |
| rs34624793 | T | 0.0116 | 0.0021 | -0.0068 | 0.0029 |
| rs34720381 | T | -0.0180 | 0.0024 | 0.0094 | 0.0035 |
| rs34748029 | A | 0.0185 | 0.0025 | -0.0064 | 0.0035 |
| rs347661 | T | -0.0081 | 0.0014 | 0.0033 | 0.0020 |
| rs34780702 | A | 0.0111 | 0.0017 | -0.0067 | 0.0023 |
| rs34807077 | A | 0.0128 | 0.0019 | -0.0088 | 0.0027 |
| rs34811474 | A | 0.0104 | 0.0017 | -0.0068 | 0.0024 |
| rs34967082 | A | -0.0080 | 0.0014 | 0.0026 | 0.0020 |
| rs35016816 | T | 0.0176 | 0.0023 | -0.0006 | 0.0033 |
| rs350281 | T | -0.0117 | 0.0015 | 0.0068 | 0.0021 |
| rs35104491 | A | 0.0124 | 0.0018 | -0.0011 | 0.0026 |
| rs35111506 | C | -0.0175 | 0.0030 | 0.0081 | 0.0043 |
| rs35192107 | T | -0.0091 | 0.0017 | 0.0054 | 0.0023 |
| rs35319653 | T | 0.0119 | 0.0015 | -0.0106 | 0.0021 |
| rs35417702 | T | -0.0152 | 0.0014 | -0.0055 | 0.0020 |
| rs35493937 | C | 0.0124 | 0.0017 | 0.0061 | 0.0024 |
| rs35606437 | A | 0.0105 | 0.0016 | -0.0028 | 0.0023 |
| rs356999 | A | -0.0101 | 0.0014 | 0.0050 | 0.0021 |
| rs35745455 | A | 0.0080 | 0.0014 | 0.0016 | 0.0020 |
| rs35751693 | T | -0.0246 | 0.0040 | 0.0068 | 0.0053 |
| rs35754740 | T | 0.0083 | 0.0014 | -0.0075 | 0.0020 |
| rs35811586 | T | 0.0217 | 0.0031 | -0.0038 | 0.0043 |
| rs35929923 | A | -0.0114 | 0.0016 | 0.0040 | 0.0023 |
| rs36085856 | C | 0.0201 | 0.0036 | -0.0049 | 0.0039 |
| rs36120534 | T | 0.0100 | 0.0018 | -0.0002 | 0.0026 |
| rs362307 | T | -0.0226 | 0.0027 | 0.0103 | 0.0038 |
| rs363096 | T | -0.0143 | 0.0014 | 0.0065 | 0.0020 |
| rs3735478 | T | 0.0108 | 0.0016 | -0.0087 | 0.0022 |
| rs3751331 | A | -0.0097 | 0.0014 | 0.0065 | 0.0021 |
| rs3766979 | A | 0.0085 | 0.0015 | 0.0047 | 0.0021 |
| rs3768480 | C | 0.0101 | 0.0014 | -0.0047 | 0.0020 |
| rs3781339 | T | -0.0104 | 0.0018 | 0.0051 | 0.0025 |
| rs3790609 | T | 0.0107 | 0.0019 | -0.0095 | 0.0026 |
| rs3809169 | T | 0.0140 | 0.0025 | 0.0030 | 0.0035 |
| rs3809634 | A | -0.0109 | 0.0015 | 0.0044 | 0.0022 |
| rs3812281 | T | 0.0113 | 0.0014 | -0.0059 | 0.0020 |
| rs3817923 | A | -0.0121 | 0.0022 | 0.0020 | 0.0031 |
| rs382196 | T | -0.0082 | 0.0014 | 0.0046 | 0.0021 |
| rs3847228 | T | 0.0083 | 0.0014 | -0.0022 | 0.0020 |
| rs3848715 | A | -0.0080 | 0.0014 | -0.0014 | 0.0020 |
| rs3859523 | T | 0.0124 | 0.0018 | -0.0055 | 0.0026 |
| rs387027 | A | -0.0083 | 0.0015 | 0.0049 | 0.0021 |
| rs3895736 | A | 0.0144 | 0.0019 | -0.0038 | 0.0026 |
| rs3897821 | A | 0.0154 | 0.0015 | -0.0007 | 0.0021 |
| rs3948495 | T | -0.0089 | 0.0014 | 0.0074 | 0.0020 |
| rs399821 | A | -0.0078 | 0.0014 | 0.0019 | 0.0020 |
| rs39998 | A | 0.0090 | 0.0015 | 0.0025 | 0.0022 |
| rs401526 | T | -0.0104 | 0.0014 | 0.0021 | 0.0020 |
| rs401966 | C | -0.0097 | 0.0014 | 0.0056 | 0.0021 |
| rs4127499 | A | 0.0087 | 0.0015 | -0.0027 | 0.0021 |
| rs41282553 | A | 0.0237 | 0.0040 | -0.0051 | 0.0057 |
| rs42210 | C | -0.0098 | 0.0016 | 0.0086 | 0.0022 |
| rs42302 | A | 0.0086 | 0.0015 | -0.0051 | 0.0021 |
| rs4255791 | A | -0.0084 | 0.0015 | 0.0049 | 0.0021 |
| rs4263475 | A | 0.0078 | 0.0014 | -0.0035 | 0.0020 |
| rs4283754 | A | -0.0084 | 0.0014 | 0.0003 | 0.0021 |
| rs429150 | T | 0.0085 | 0.0014 | -0.0037 | 0.0020 |
| rs4298514 | T | 0.0104 | 0.0015 | -0.0016 | 0.0022 |
| rs4320563 | A | 0.0109 | 0.0014 | -0.0037 | 0.0020 |
| rs4328757 | T | 0.0096 | 0.0014 | -0.0026 | 0.0020 |
| rs4352658 | T | -0.0190 | 0.0026 | 0.0168 | 0.0037 |
| rs4358081 | A | -0.0095 | 0.0014 | 0.0085 | 0.0020 |
| rs4358358 | A | 0.0109 | 0.0015 | -0.0009 | 0.0022 |
| rs4369924 | A | 0.0125 | 0.0019 | -0.0018 | 0.0027 |
| rs4384309 | A | 0.0102 | 0.0014 | -0.0061 | 0.0020 |
| rs4396896 | A | -0.0107 | 0.0014 | 0.0041 | 0.0020 |
| rs4423373 | A | -0.0087 | 0.0014 | 0.0015 | 0.0020 |
| rs4426420 | T | -0.0140 | 0.0019 | 0.0045 | 0.0027 |
| rs4434676 | A | -0.0089 | 0.0014 | -0.0006 | 0.0020 |
| rs4458044 | C | 0.0101 | 0.0016 | 0.0030 | 0.0023 |
| rs4467547 | T | 0.0121 | 0.0014 | -0.0071 | 0.0020 |
| rs4469771 | T | 0.0101 | 0.0015 | 0.0020 | 0.0021 |
| rs4480339 | A | -0.0097 | 0.0015 | 0.0048 | 0.0022 |
| rs4490539 | A | 0.0142 | 0.0015 | -0.0073 | 0.0022 |
| rs4497562 | A | 0.0112 | 0.0016 | -0.0036 | 0.0023 |
| rs4500930 | T | -0.0109 | 0.0015 | 0.0025 | 0.0021 |
| rs4500960 | T | -0.0131 | 0.0014 | 0.0102 | 0.0020 |
| rs4502401 | T | 0.0085 | 0.0015 | -0.0047 | 0.0021 |
| rs4652135 | A | 0.0103 | 0.0016 | -0.0052 | 0.0022 |
| rs4658019 | T | 0.0084 | 0.0014 | 0.0003 | 0.0020 |
| rs4663617 | A | 0.0104 | 0.0017 | -0.0010 | 0.0024 |
| rs4664983 | T | -0.0103 | 0.0018 | -0.0015 | 0.0025 |
| rs4673840 | T | -0.0138 | 0.0019 | -0.0001 | 0.0027 |
| rs4675248 | A | -0.0094 | 0.0014 | 0.0134 | 0.0020 |
| rs4685405 | T | -0.0111 | 0.0018 | 0.0099 | 0.0026 |
| rs4687735 | T | -0.0200 | 0.0033 | 0.0146 | 0.0046 |
| rs4691601 | A | -0.0115 | 0.0014 | 0.0025 | 0.0020 |
| rs4705763 | C | 0.0089 | 0.0014 | -0.0049 | 0.0020 |
| rs4712371 | A | -0.0124 | 0.0021 | 0.0069 | 0.0031 |
| rs4719460 | T | -0.0088 | 0.0015 | 0.0002 | 0.0021 |
| rs4719944 | T | -0.0121 | 0.0014 | 0.0074 | 0.0020 |
| rs4724083 | T | 0.0083 | 0.0015 | -0.0016 | 0.0022 |
| rs4726070 | A | 0.0122 | 0.0014 | -0.0020 | 0.0020 |
| rs4730020 | T | 0.0090 | 0.0016 | -0.0030 | 0.0023 |
| rs4731413 | A | 0.0111 | 0.0017 | -0.0020 | 0.0025 |
| rs4735297 | A | -0.0086 | 0.0015 | 0.0114 | 0.0021 |
| rs4739235 | A | 0.0109 | 0.0019 | -0.0087 | 0.0027 |
| rs4741571 | A | -0.0102 | 0.0015 | 0.0037 | 0.0022 |
| rs4757957 | C | 0.0133 | 0.0015 | -0.0038 | 0.0021 |
| rs4766424 | C | 0.0142 | 0.0021 | -0.0064 | 0.0030 |
| rs4778058 | T | -0.0092 | 0.0014 | 0.0018 | 0.0020 |
| rs4785187 | A | -0.0100 | 0.0017 | 0.0120 | 0.0024 |
| rs4787028 | T | 0.0086 | 0.0015 | 0.0006 | 0.0021 |
| rs4787457 | A | 0.0162 | 0.0015 | -0.0035 | 0.0021 |
| rs4788115 | A | 0.0111 | 0.0019 | -0.0038 | 0.0027 |
| rs479018 | A | 0.0109 | 0.0015 | -0.0052 | 0.0022 |
| rs4793090 | A | 0.0084 | 0.0015 | -0.0069 | 0.0021 |
| rs4810894 | A | -0.0085 | 0.0015 | -0.0009 | 0.0021 |
| rs4812697 | T | 0.0171 | 0.0027 | -0.0034 | 0.0038 |
| rs481940 | T | 0.0100 | 0.0016 | -0.0023 | 0.0023 |
| rs482787 | T | 0.0100 | 0.0015 | -0.0038 | 0.0021 |
| rs4839155 | T | 0.0115 | 0.0017 | -0.0064 | 0.0024 |
| rs4846010 | A | 0.0099 | 0.0018 | 0.0006 | 0.0025 |
| rs4846724 | A | 0.0098 | 0.0014 | -0.0025 | 0.0020 |
| rs4848924 | A | -0.0117 | 0.0015 | 0.0034 | 0.0022 |
| rs4850954 | T | 0.0077 | 0.0014 | -0.0003 | 0.0020 |
| rs4851263 | A | -0.0151 | 0.0023 | 0.0031 | 0.0033 |
| rs4877516 | A | -0.0101 | 0.0014 | 0.0123 | 0.0020 |
| rs4881269 | A | 0.0100 | 0.0014 | -0.0006 | 0.0021 |
| rs488476 | C | -0.0113 | 0.0015 | 0.0074 | 0.0020 |
| rs4894658 | C | 0.0091 | 0.0016 | -0.0078 | 0.0023 |
| rs4895650 | T | 0.0093 | 0.0014 | -0.0069 | 0.0020 |
| rs4899012 | C | -0.0103 | 0.0014 | 0.0042 | 0.0020 |
| rs4904523 | A | -0.0082 | 0.0014 | 0.0029 | 0.0020 |
| rs4915735 | A | 0.0116 | 0.0020 | -0.0003 | 0.0029 |
| rs4919624 | A | 0.0193 | 0.0018 | -0.0102 | 0.0025 |
| rs4925065 | T | -0.0078 | 0.0014 | 0.0002 | 0.0020 |
| rs4925109 | A | -0.0096 | 0.0015 | 0.0030 | 0.0022 |
| rs4938815 | T | 0.0085 | 0.0015 | -0.0023 | 0.0022 |
| rs4941735 | T | 0.0092 | 0.0014 | -0.0045 | 0.0020 |
| rs4972748 | T | 0.0105 | 0.0018 | -0.0075 | 0.0026 |
| rs4977885 | A | -0.0084 | 0.0014 | 0.0062 | 0.0020 |
| rs4981245 | A | 0.0080 | 0.0014 | 0.0000 | 0.0020 |
| rs4984541 | A | -0.0138 | 0.0017 | 0.0065 | 0.0024 |
| rs4984613 | T | 0.0174 | 0.0028 | 0.0023 | 0.0041 |
| rs4984682 | C | -0.0138 | 0.0017 | 0.0122 | 0.0024 |
| rs548897 | A | 0.0081 | 0.0014 | -0.0048 | 0.0020 |
| rs55675587 | T | -0.0103 | 0.0018 | 0.0055 | 0.0026 |
| rs55736314 | C | -0.0154 | 0.0014 | 0.0075 | 0.0020 |
| rs55826493 | T | -0.0127 | 0.0021 | 0.0042 | 0.0030 |
| rs55986781 | T | 0.0187 | 0.0026 | -0.0120 | 0.0035 |
| rs56085180 | A | -0.0269 | 0.0037 | 0.0010 | 0.0052 |
| rs56099375 | T | 0.0120 | 0.0017 | -0.0095 | 0.0023 |
| rs56171318 | T | -0.0137 | 0.0020 | 0.0033 | 0.0028 |
| rs56174996 | A | -0.0143 | 0.0020 | 0.0080 | 0.0029 |
| rs56306882 | A | 0.0086 | 0.0016 | -0.0082 | 0.0023 |
| rs56319902 | T | -0.0206 | 0.0017 | -0.0084 | 0.0024 |
| rs56391344 | A | 0.0148 | 0.0016 | -0.0105 | 0.0023 |
| rs563954 | A | -0.0081 | 0.0014 | 0.0079 | 0.0020 |
| rs56405138 | A | 0.0153 | 0.0022 | -0.0118 | 0.0031 |
| rs56408528 | T | -0.0082 | 0.0014 | -0.0029 | 0.0020 |
| rs567003 | A | 0.0094 | 0.0015 | -0.0017 | 0.0021 |
| rs56794817 | A | 0.0122 | 0.0019 | -0.0014 | 0.0028 |
| rs57016874 | T | 0.0209 | 0.0037 | -0.0054 | 0.0051 |
| rs57148205 | A | 0.0095 | 0.0016 | -0.0032 | 0.0024 |
| rs57204268 | A | 0.0114 | 0.0020 | -0.0041 | 0.0028 |
| rs57349798 | A | 0.0093 | 0.0014 | -0.0109 | 0.0020 |
| rs57352738 | A | -0.0152 | 0.0017 | 0.0110 | 0.0025 |
| rs57437407 | A | -0.0145 | 0.0023 | 0.0076 | 0.0032 |
| rs575113 | A | 0.0133 | 0.0015 | -0.0023 | 0.0022 |
| rs5754581 | T | -0.0082 | 0.0014 | -0.0021 | 0.0020 |
| rs5754753 | T | -0.0126 | 0.0016 | 0.0006 | 0.0022 |
| rs5754762 | A | 0.0197 | 0.0030 | -0.0159 | 0.0044 |
| rs57661533 | T | -0.0114 | 0.0021 | 0.0036 | 0.0030 |
| rs580652 | T | -0.0147 | 0.0024 | 0.0018 | 0.0034 |
| rs585557 | A | 0.0128 | 0.0018 | -0.0053 | 0.0025 |
| rs58779949 | A | 0.0110 | 0.0019 | -0.0086 | 0.0026 |
| rs58859557 | T | 0.0187 | 0.0028 | -0.0136 | 0.0039 |
| rs58921703 | T | 0.0109 | 0.0015 | -0.0042 | 0.0021 |
| rs58950082 | T | 0.0095 | 0.0017 | -0.0037 | 0.0025 |
| rs58996896 | A | -0.0094 | 0.0016 | 0.0103 | 0.0023 |
| rs590013 | T | 0.0100 | 0.0015 | -0.0060 | 0.0021 |
| rs59300999 | T | -0.0164 | 0.0028 | 0.0096 | 0.0040 |
| rs59480703 | C | -0.0123 | 0.0018 | 0.0082 | 0.0025 |
| rs59813324 | T | 0.0097 | 0.0018 | -0.0027 | 0.0025 |
| rs59967356 | T | -0.0127 | 0.0021 | 0.0027 | 0.0030 |
| rs60096640 | A | 0.0159 | 0.0023 | -0.0091 | 0.0033 |
| rs6020560 | T | -0.0083 | 0.0014 | 0.0062 | 0.0020 |
| rs6043521 | T | -0.0083 | 0.0014 | -0.0002 | 0.0021 |
| rs60589532 | A | 0.0168 | 0.0028 | -0.0196 | 0.0042 |
| rs6060308 | A | 0.0095 | 0.0016 | -0.0089 | 0.0023 |
| rs6065080 | T | -0.0130 | 0.0015 | -0.0013 | 0.0021 |
| rs6065784 | C | 0.0110 | 0.0015 | -0.0047 | 0.0022 |
| rs60717745 | C | -0.0118 | 0.0019 | 0.0010 | 0.0028 |
| rs60726488 | T | -0.0090 | 0.0016 | -0.0026 | 0.0023 |
| rs6091570 | A | 0.0081 | 0.0015 | -0.0004 | 0.0021 |
| rs61104616 | A | -0.0172 | 0.0014 | 0.0036 | 0.0020 |
| rs6123924 | A | 0.0129 | 0.0019 | -0.0010 | 0.0028 |
| rs613872 | T | -0.0163 | 0.0019 | 0.0074 | 0.0026 |
| rs61387839 | A | 0.0088 | 0.0015 | 0.0025 | 0.0022 |
| rs61527214 | A | 0.0110 | 0.0014 | -0.0030 | 0.0020 |
| rs61739710 | A | -0.0099 | 0.0016 | 0.0010 | 0.0022 |
| rs61748951 | A | -0.0273 | 0.0048 | 0.0176 | 0.0067 |
| rs61755388 | T | 0.0125 | 0.0017 | -0.0003 | 0.0024 |
| rs61757207 | A | 0.0359 | 0.0063 | -0.0157 | 0.0082 |
| rs61798586 | A | -0.0112 | 0.0020 | -0.0016 | 0.0028 |
| rs61853335 | T | -0.0109 | 0.0018 | 0.0118 | 0.0026 |
| rs61958175 | A | -0.0198 | 0.0034 | 0.0227 | 0.0047 |
| rs61997667 | T | -0.0144 | 0.0020 | 0.0078 | 0.0028 |
| rs62007304 | A | 0.0264 | 0.0047 | -0.0148 | 0.0061 |
| rs62018216 | T | 0.0113 | 0.0019 | -0.0059 | 0.0028 |
| rs62051146 | A | -0.0134 | 0.0022 | 0.0051 | 0.0031 |
| rs62090515 | A | -0.0090 | 0.0015 | 0.0007 | 0.0021 |
| rs62092949 | T | -0.0087 | 0.0014 | -0.0022 | 0.0020 |
| rs62103198 | T | -0.0147 | 0.0025 | 0.0113 | 0.0035 |
| rs62109862 | A | 0.0136 | 0.0020 | -0.0021 | 0.0029 |
| rs62142891 | A | 0.0108 | 0.0016 | -0.0054 | 0.0023 |
| rs62155350 | A | 0.0338 | 0.0053 | -0.0024 | 0.0072 |
| rs62155770 | A | -0.0196 | 0.0032 | 0.0083 | 0.0043 |
| rs62155873 | T | -0.0151 | 0.0021 | 0.0244 | 0.0030 |
| rs62174974 | A | -0.0105 | 0.0018 | 0.0031 | 0.0025 |
| rs62177359 | A | -0.0275 | 0.0038 | 0.0290 | 0.0055 |
| rs62179650 | A | 0.0112 | 0.0016 | -0.0115 | 0.0022 |
| rs62182994 | T | 0.0137 | 0.0015 | -0.0082 | 0.0021 |
| rs62190914 | T | 0.0089 | 0.0015 | 0.0039 | 0.0021 |
| rs62194170 | A | 0.0111 | 0.0017 | -0.0054 | 0.0025 |
| rs622169 | T | 0.0086 | 0.0014 | 0.0011 | 0.0021 |
| rs62247449 | C | 0.0114 | 0.0014 | -0.0017 | 0.0020 |
| rs62252819 | A | -0.0094 | 0.0017 | 0.0045 | 0.0023 |
| rs62256284 | T | 0.0111 | 0.0017 | 0.0020 | 0.0024 |
| rs62260764 | C | -0.0132 | 0.0016 | 0.0082 | 0.0023 |
| rs62262671 | A | 0.0213 | 0.0020 | -0.0149 | 0.0029 |
| rs62340636 | T | 0.0103 | 0.0014 | -0.0033 | 0.0021 |
| rs62370510 | A | -0.0109 | 0.0019 | -0.0006 | 0.0026 |
| rs62379838 | T | 0.0121 | 0.0015 | 0.0005 | 0.0022 |
| rs62409395 | T | 0.0101 | 0.0018 | -0.0039 | 0.0025 |
| rs62420387 | T | -0.0184 | 0.0027 | 0.0053 | 0.0038 |
| rs62506074 | T | 0.0091 | 0.0015 | -0.0019 | 0.0021 |
| rs62506104 | A | -0.0098 | 0.0018 | 0.0015 | 0.0026 |
| rs628993 | A | 0.0128 | 0.0023 | -0.0042 | 0.0034 |
| rs631287 | A | 0.0084 | 0.0014 | -0.0128 | 0.0020 |
| rs633279 | A | -0.0089 | 0.0015 | -0.0031 | 0.0022 |
| rs635754 | A | 0.0119 | 0.0014 | -0.0009 | 0.0020 |
| rs6428587 | T | 0.0084 | 0.0015 | -0.0047 | 0.0021 |
| rs6435326 | A | -0.0091 | 0.0014 | 0.0032 | 0.0020 |
| rs6442126 | A | -0.0121 | 0.0015 | 0.0097 | 0.0021 |
| rs6445633 | A | 0.0088 | 0.0015 | 0.0013 | 0.0022 |
| rs6449503 | A | 0.0185 | 0.0014 | -0.0088 | 0.0020 |
| rs6450476 | A | -0.0116 | 0.0015 | -0.0039 | 0.0022 |
| rs6452793 | T | 0.0175 | 0.0017 | -0.0065 | 0.0024 |
| rs6461536 | A | 0.0088 | 0.0015 | -0.0015 | 0.0022 |
| rs6469654 | C | -0.0103 | 0.0017 | 0.0059 | 0.0024 |
| rs6472208 | T | -0.0091 | 0.0014 | -0.0050 | 0.0020 |
| rs6480234 | T | -0.0084 | 0.0014 | 0.0044 | 0.0020 |
| rs648044 | A | 0.0083 | 0.0015 | -0.0067 | 0.0021 |
| rs6490618 | T | 0.0088 | 0.0015 | -0.0017 | 0.0021 |
| rs6493265 | T | -0.0122 | 0.0014 | 0.0132 | 0.0020 |
| rs6493275 | A | 0.0131 | 0.0017 | -0.0234 | 0.0024 |
| rs6503409 | T | 0.0087 | 0.0015 | -0.0002 | 0.0021 |
| rs6534338 | T | 0.0103 | 0.0015 | -0.0101 | 0.0022 |
| rs6543810 | T | 0.0082 | 0.0015 | 0.0019 | 0.0021 |
| rs6557171 | T | -0.0155 | 0.0015 | 0.0081 | 0.0021 |
| rs6567288 | A | -0.0079 | 0.0014 | 0.0039 | 0.0020 |
| rs6569077 | T | 0.0118 | 0.0014 | -0.0044 | 0.0021 |
| rs6573552 | T | -0.0090 | 0.0014 | 0.0009 | 0.0020 |
| rs6573559 | T | 0.0105 | 0.0015 | 0.0032 | 0.0022 |
| rs660001 | A | -0.0164 | 0.0017 | 0.0019 | 0.0024 |
| rs663251 | T | 0.0081 | 0.0015 | -0.0047 | 0.0021 |
| rs66482320 | C | 0.0124 | 0.0022 | -0.0024 | 0.0032 |
| rs66568921 | T | -0.0181 | 0.0015 | 0.0106 | 0.0021 |
| rs66671632 | T | -0.0133 | 0.0021 | 0.0057 | 0.0030 |
| rs66721975 | A | -0.0087 | 0.0015 | 0.0011 | 0.0022 |
| rs6672986 | C | 0.0094 | 0.0014 | -0.0059 | 0.0020 |
| rs6689641 | A | 0.0078 | 0.0014 | -0.0052 | 0.0020 |
| rs6690195 | T | -0.0113 | 0.0014 | 0.0103 | 0.0020 |
| rs6695132 | T | 0.0097 | 0.0017 | -0.0029 | 0.0024 |
| rs6696068 | T | -0.0088 | 0.0014 | -0.0013 | 0.0020 |
| rs6697584 | T | -0.0126 | 0.0017 | 0.0034 | 0.0024 |
| rs6704768 | A | -0.0116 | 0.0014 | -0.0028 | 0.0020 |
| rs6706275 | T | 0.0083 | 0.0015 | 0.0021 | 0.0021 |
| rs6711399 | T | -0.0107 | 0.0019 | 0.0007 | 0.0026 |
| rs6715321 | T | -0.0097 | 0.0014 | 0.0043 | 0.0020 |
| rs6715849 | A | -0.0122 | 0.0014 | 0.0031 | 0.0020 |
| rs6720515 | A | 0.0103 | 0.0017 | -0.0018 | 0.0024 |
| rs67224963 | A | 0.0113 | 0.0017 | -0.0011 | 0.0025 |
| rs6729612 | T | 0.0092 | 0.0015 | 0.0029 | 0.0021 |
| rs6731373 | A | -0.0115 | 0.0015 | -0.0067 | 0.0021 |
| rs6736025 | T | 0.0079 | 0.0014 | -0.0039 | 0.0020 |
| rs6738860 | A | -0.0088 | 0.0014 | 0.0111 | 0.0020 |
| rs6743032 | A | -0.0165 | 0.0024 | 0.0042 | 0.0034 |
| rs6744040 | A | -0.0080 | 0.0014 | 0.0057 | 0.0020 |
| rs67456868 | A | 0.0119 | 0.0018 | -0.0010 | 0.0026 |
| rs6752228 | T | -0.0080 | 0.0014 | 0.0062 | 0.0020 |
| rs6757087 | T | -0.0085 | 0.0014 | 0.0056 | 0.0020 |
| rs67661275 | A | 0.0134 | 0.0025 | 0.0011 | 0.0035 |
| rs6774533 | T | 0.0090 | 0.0015 | -0.0016 | 0.0022 |
| rs67790232 | A | 0.0101 | 0.0017 | -0.0062 | 0.0025 |
| rs6780414 | T | 0.0105 | 0.0018 | 0.0008 | 0.0026 |
| rs6792805 | A | -0.0090 | 0.0016 | 0.0056 | 0.0022 |
| rs6812533 | T | 0.0106 | 0.0016 | -0.0072 | 0.0023 |
| rs68145588 | T | -0.0127 | 0.0021 | 0.0037 | 0.0028 |
| rs6824567 | T | 0.0130 | 0.0016 | -0.0130 | 0.0022 |
| rs6853599 | T | 0.0108 | 0.0019 | -0.0066 | 0.0026 |
| rs6861925 | C | -0.0090 | 0.0014 | 0.0064 | 0.0020 |
| rs6871635 | A | -0.0087 | 0.0014 | 0.0112 | 0.0020 |
| rs6881581 | A | -0.0096 | 0.0015 | 0.0018 | 0.0021 |
| rs6881733 | T | -0.0086 | 0.0016 | -0.0010 | 0.0022 |
| rs6917154 | T | 0.0127 | 0.0021 | -0.0072 | 0.0030 |
| rs6917204 | T | 0.0119 | 0.0017 | -0.0045 | 0.0025 |
| rs6918506 | C | 0.0113 | 0.0014 | -0.0084 | 0.0020 |
| rs6924023 | A | 0.0107 | 0.0016 | -0.0062 | 0.0023 |
| rs6932108 | C | -0.0210 | 0.0027 | 0.0088 | 0.0036 |
| rs6946136 | A | 0.0081 | 0.0014 | -0.0016 | 0.0020 |
| rs6946362 | T | 0.0091 | 0.0015 | -0.0039 | 0.0022 |
| rs6951996 | A | -0.0182 | 0.0032 | 0.0064 | 0.0046 |
| rs6956283 | T | 0.0098 | 0.0016 | -0.0053 | 0.0022 |
| rs6969783 | A | -0.0098 | 0.0014 | 0.0146 | 0.0020 |
| rs6977237 | T | 0.0082 | 0.0014 | -0.0023 | 0.0020 |
| rs6989141 | A | 0.0100 | 0.0017 | -0.0047 | 0.0025 |
| rs6994287 | A | 0.0102 | 0.0014 | -0.0013 | 0.0020 |
| rs7012546 | T | 0.0092 | 0.0014 | -0.0001 | 0.0020 |
| rs7016302 | C | -0.0125 | 0.0019 | -0.0008 | 0.0027 |
| rs702606 | T | 0.0118 | 0.0021 | -0.0031 | 0.0029 |
| rs7029718 | A | 0.0244 | 0.0014 | -0.0050 | 0.0020 |
| rs7030373 | A | -0.0098 | 0.0017 | 0.0031 | 0.0025 |
| rs7035315 | A | -0.0090 | 0.0015 | 0.0018 | 0.0021 |
| rs7040995 | C | 0.0097 | 0.0014 | -0.0033 | 0.0020 |
| rs7041702 | A | -0.0111 | 0.0016 | 0.0046 | 0.0023 |
| rs7084508 | T | -0.0085 | 0.0015 | 0.0046 | 0.0021 |
| rs710629 | A | 0.0082 | 0.0015 | 0.0051 | 0.0021 |
| rs7108020 | A | 0.0111 | 0.0015 | -0.0047 | 0.0021 |
| rs711793 | T | 0.0103 | 0.0015 | 0.0012 | 0.0021 |
| rs7127580 | C | -0.0150 | 0.0023 | 0.0039 | 0.0032 |
| rs713584 | A | -0.0080 | 0.0014 | 0.0020 | 0.0020 |
| rs71413877 | A | 0.0351 | 0.0036 | -0.0076 | 0.0051 |
| rs7146625 | A | 0.0091 | 0.0016 | -0.0015 | 0.0023 |
| rs7147473 | A | -0.0103 | 0.0015 | 0.0052 | 0.0021 |
| rs7158218 | A | -0.0090 | 0.0015 | 0.0018 | 0.0022 |
| rs71646142 | T | 0.0104 | 0.0018 | -0.0023 | 0.0025 |
| rs716513 | A | 0.0084 | 0.0014 | -0.0073 | 0.0020 |
| rs7167688 | T | -0.0094 | 0.0014 | 0.0003 | 0.0020 |
| rs7171405 | A | -0.0101 | 0.0016 | 0.0029 | 0.0023 |
| rs717996 | T | -0.0122 | 0.0014 | 0.0091 | 0.0020 |
| rs7182216 | C | -0.0096 | 0.0017 | -0.0022 | 0.0024 |
| rs7190 | A | -0.0095 | 0.0015 | 0.0061 | 0.0021 |
| rs721579 | T | -0.0110 | 0.0016 | -0.0024 | 0.0023 |
| rs7218235 | A | -0.0097 | 0.0017 | 0.0091 | 0.0025 |
| rs7223311 | C | -0.0079 | 0.0014 | 0.0002 | 0.0020 |
| rs7226824 | T | -0.0078 | 0.0014 | 0.0021 | 0.0020 |
| rs72482130 | T | 0.0141 | 0.0019 | -0.0118 | 0.0028 |
| rs7254263 | T | -0.0119 | 0.0016 | -0.0001 | 0.0022 |
| rs7255223 | A | 0.0097 | 0.0016 | -0.0035 | 0.0023 |
| rs72622559 | T | -0.0120 | 0.0017 | 0.0055 | 0.0026 |
| rs72624911 | T | 0.0282 | 0.0034 | -0.0076 | 0.0048 |
| rs72636697 | T | -0.0140 | 0.0020 | 0.0102 | 0.0029 |
| rs72667460 | T | 0.0197 | 0.0031 | -0.0094 | 0.0044 |
| rs72671456 | A | -0.0174 | 0.0020 | 0.0036 | 0.0029 |
| rs72672052 | A | -0.0101 | 0.0018 | 0.0038 | 0.0026 |
| rs72673097 | A | 0.0257 | 0.0042 | 0.0038 | 0.0055 |
| rs72677177 | A | 0.0100 | 0.0014 | -0.0065 | 0.0020 |
| rs72694479 | T | 0.0127 | 0.0020 | 0.0010 | 0.0028 |
| rs72709560 | A | 0.0085 | 0.0015 | -0.0060 | 0.0021 |
| rs72771860 | T | -0.0211 | 0.0029 | 0.0047 | 0.0042 |
| rs72792395 | T | -0.0145 | 0.0022 | 0.0023 | 0.0032 |
| rs728054 | A | -0.0128 | 0.0015 | -0.0004 | 0.0021 |
| rs72819118 | T | -0.0231 | 0.0024 | 0.0009 | 0.0034 |
| rs72824753 | T | 0.0133 | 0.0024 | -0.0042 | 0.0033 |
| rs72828517 | T | -0.0168 | 0.0019 | -0.0005 | 0.0026 |
| rs72829857 | A | -0.0149 | 0.0016 | 0.0063 | 0.0024 |
| rs72834698 | A | 0.0117 | 0.0020 | -0.0092 | 0.0029 |
| rs72881110 | A | 0.0146 | 0.0027 | -0.0033 | 0.0038 |
| rs72883760 | A | 0.0101 | 0.0018 | -0.0083 | 0.0025 |
| rs72896637 | T | 0.0162 | 0.0028 | -0.0056 | 0.0038 |
| rs72902523 | T | -0.0103 | 0.0016 | -0.0025 | 0.0023 |
| rs72906124 | T | 0.0185 | 0.0030 | -0.0063 | 0.0043 |
| rs72917504 | T | -0.0178 | 0.0030 | 0.0040 | 0.0042 |
| rs72919450 | T | -0.0132 | 0.0023 | -0.0002 | 0.0033 |
| rs72944064 | T | -0.0098 | 0.0016 | 0.0015 | 0.0023 |
| rs72962169 | T | -0.0174 | 0.0019 | 0.0108 | 0.0027 |
| rs72972965 | A | 0.0083 | 0.0015 | -0.0033 | 0.0021 |
| rs72993796 | T | 0.0127 | 0.0022 | -0.0022 | 0.0031 |
| rs73034295 | A | 0.0099 | 0.0018 | -0.0056 | 0.0026 |
| rs730384 | A | 0.0101 | 0.0014 | -0.0025 | 0.0020 |
| rs73039077 | C | 0.0092 | 0.0016 | -0.0088 | 0.0023 |
| rs73055556 | A | 0.0111 | 0.0020 | -0.0081 | 0.0029 |
| rs73082325 | C | -0.0252 | 0.0044 | 0.0106 | 0.0059 |
| rs73106136 | T | -0.0147 | 0.0026 | 0.0012 | 0.0037 |
| rs73154546 | A | 0.0197 | 0.0036 | 0.0047 | 0.0050 |
| rs7317761 | C | 0.0109 | 0.0016 | -0.0116 | 0.0023 |
| rs73191311 | A | -0.0094 | 0.0015 | -0.0026 | 0.0021 |
| rs7321274 | A | 0.0114 | 0.0018 | -0.0051 | 0.0025 |
| rs73219806 | A | 0.0106 | 0.0019 | -0.0105 | 0.0027 |
| rs7323027 | A | 0.0130 | 0.0015 | -0.0074 | 0.0021 |
| rs7326331 | A | -0.0128 | 0.0016 | 0.0024 | 0.0022 |
| rs73344830 | A | 0.0170 | 0.0014 | -0.0071 | 0.0020 |
| rs73405293 | A | 0.0113 | 0.0020 | -0.0088 | 0.0028 |
| rs73457936 | A | 0.0144 | 0.0025 | -0.0066 | 0.0035 |
| rs7356536 | T | -0.0129 | 0.0021 | 0.0006 | 0.0029 |
| rs73581580 | A | -0.0181 | 0.0022 | 0.0112 | 0.0030 |
| rs736281 | T | 0.0086 | 0.0014 | 0.0006 | 0.0020 |
| rs73643713 | T | -0.0165 | 0.0027 | 0.0015 | 0.0038 |
| rs736471 | T | 0.0083 | 0.0014 | 0.0008 | 0.0020 |
| rs73648455 | T | -0.0183 | 0.0026 | 0.0086 | 0.0037 |
| rs737945 | C | -0.0115 | 0.0014 | -0.0012 | 0.0020 |
| rs73874335 | T | -0.0178 | 0.0029 | 0.0048 | 0.0044 |
| rs73961845 | A | -0.0114 | 0.0014 | 0.0008 | 0.0021 |
| rs74091672 | A | -0.0153 | 0.0022 | 0.0067 | 0.0031 |
| rs7430651 | T | -0.0111 | 0.0016 | 0.0020 | 0.0022 |
| rs743316 | T | 0.0112 | 0.0017 | 0.0012 | 0.0025 |
| rs74415461 | T | 0.0176 | 0.0026 | -0.0148 | 0.0036 |
| rs74453875 | A | 0.0214 | 0.0037 | -0.0035 | 0.0054 |
| rs74462621 | T | 0.0125 | 0.0020 | -0.0107 | 0.0029 |
| rs7449561 | A | 0.0102 | 0.0017 | -0.0068 | 0.0024 |
| rs7451726 | A | 0.0109 | 0.0017 | -0.0004 | 0.0025 |
| rs74545339 | A | -0.0158 | 0.0022 | 0.0020 | 0.0032 |
| rs7460106 | T | -0.0123 | 0.0017 | -0.0019 | 0.0024 |
| rs74615093 | A | 0.0145 | 0.0026 | -0.0008 | 0.0037 |
| rs746839 | C | 0.0135 | 0.0015 | -0.0021 | 0.0021 |
| rs74747621 | T | 0.0153 | 0.0022 | -0.0027 | 0.0032 |
| rs74787922 | A | 0.0182 | 0.0028 | -0.0087 | 0.0039 |
| rs74944275 | T | 0.0231 | 0.0035 | -0.0074 | 0.0048 |
| rs74944857 | T | -0.0089 | 0.0015 | -0.0023 | 0.0021 |
| rs7495033 | A | 0.0095 | 0.0015 | -0.0066 | 0.0022 |
| rs75033012 | C | -0.0258 | 0.0038 | -0.0029 | 0.0051 |
| rs75177132 | T | 0.0298 | 0.0036 | -0.0222 | 0.0047 |
| rs75203411 | T | 0.0135 | 0.0019 | -0.0025 | 0.0027 |
| rs7526112 | T | 0.0124 | 0.0015 | 0.0013 | 0.0021 |
| rs75308819 | A | 0.0287 | 0.0047 | 0.0034 | 0.0067 |
| rs75434274 | A | 0.0151 | 0.0026 | -0.0063 | 0.0036 |
| rs75500877 | T | 0.0207 | 0.0033 | 0.0122 | 0.0047 |
| rs7552964 | A | 0.0148 | 0.0023 | -0.0066 | 0.0034 |
| rs7560871 | A | -0.0193 | 0.0027 | 0.0050 | 0.0039 |
| rs7561705 | A | 0.0078 | 0.0014 | 0.0042 | 0.0020 |
| rs7575637 | A | 0.0107 | 0.0014 | -0.0054 | 0.0020 |
| rs75756843 | A | 0.0288 | 0.0049 | -0.0143 | 0.0074 |
| rs7575938 | A | 0.0090 | 0.0015 | -0.0038 | 0.0021 |
| rs7590368 | T | -0.0146 | 0.0016 | -0.0002 | 0.0023 |
| rs7597126 | T | -0.0094 | 0.0014 | -0.0044 | 0.0020 |
| rs7597412 | C | 0.0088 | 0.0015 | -0.0022 | 0.0021 |
| rs7603132 | A | 0.0137 | 0.0018 | -0.0058 | 0.0025 |
| rs76076331 | T | 0.0206 | 0.0021 | -0.0070 | 0.0030 |
| rs76077165 | A | 0.0165 | 0.0025 | -0.0111 | 0.0036 |
| rs76167224 | T | -0.0187 | 0.0033 | 0.0047 | 0.0045 |
| rs7617204 | A | -0.0097 | 0.0014 | -0.0023 | 0.0020 |
| rs76235882 | A | 0.0241 | 0.0039 | -0.0136 | 0.0059 |
| rs76241605 | A | 0.0139 | 0.0022 | -0.0029 | 0.0031 |
| rs76246107 | A | -0.0162 | 0.0026 | -0.0099 | 0.0038 |
| rs7625428 | T | 0.0098 | 0.0014 | -0.0038 | 0.0020 |
| rs76267866 | A | 0.0104 | 0.0017 | 0.0009 | 0.0025 |
| rs7630133 | A | 0.0082 | 0.0015 | -0.0015 | 0.0021 |
| rs763553 | A | 0.0087 | 0.0014 | 0.0015 | 0.0020 |
| rs7640424 | T | 0.0113 | 0.0015 | -0.0123 | 0.0022 |
| rs7650602 | T | -0.0089 | 0.0014 | 0.0017 | 0.0020 |
| rs76552497 | T | -0.0120 | 0.0019 | 0.0085 | 0.0027 |
| rs76577427 | C | 0.0203 | 0.0024 | -0.0037 | 0.0032 |
| rs7672622 | A | -0.0088 | 0.0016 | 0.0017 | 0.0023 |
| rs767943 | A | -0.0122 | 0.0016 | 0.0067 | 0.0023 |
| rs7683416 | T | 0.0139 | 0.0014 | 0.0050 | 0.0020 |
| rs76876592 | A | 0.0126 | 0.0020 | -0.0007 | 0.0029 |
| rs76878669 | C | 0.0141 | 0.0017 | -0.0002 | 0.0024 |
| rs7692359 | T | -0.0109 | 0.0017 | 0.0016 | 0.0024 |
| rs76957677 | T | -0.0297 | 0.0054 | 0.0171 | 0.0072 |
| rs77025239 | A | -0.0140 | 0.0019 | 0.0026 | 0.0028 |
| rs7714719 | C | 0.0116 | 0.0020 | -0.0080 | 0.0029 |
| rs7716161 | C | 0.0105 | 0.0019 | -0.0023 | 0.0026 |
| rs77201694 | A | 0.0156 | 0.0022 | -0.0077 | 0.0031 |
| rs77370942 | A | -0.0147 | 0.0027 | 0.0083 | 0.0037 |
| rs77554090 | T | -0.0186 | 0.0027 | 0.0083 | 0.0038 |
| rs77609760 | A | 0.0236 | 0.0028 | -0.0039 | 0.0040 |
| rs7766240 | T | -0.0134 | 0.0024 | 0.0073 | 0.0034 |
| rs77702622 | A | -0.0213 | 0.0029 | 0.0055 | 0.0040 |
| rs77702819 | T | 0.0187 | 0.0025 | -0.0048 | 0.0035 |
| rs77719387 | A | -0.0403 | 0.0058 | 0.0098 | 0.0080 |
| rs7772172 | A | 0.0091 | 0.0014 | 0.0000 | 0.0020 |
| rs7775100 | T | -0.0087 | 0.0014 | 0.0035 | 0.0020 |
| rs77826402 | T | 0.0151 | 0.0027 | -0.0059 | 0.0039 |
| rs7796103 | C | -0.0077 | 0.0014 | 0.0028 | 0.0020 |
| rs7799141 | A | 0.0099 | 0.0015 | -0.0120 | 0.0021 |
| rs77999825 | A | 0.0294 | 0.0039 | -0.0010 | 0.0054 |
| rs7803932 | A | 0.0127 | 0.0019 | -0.0004 | 0.0027 |
| rs780569 | A | -0.0085 | 0.0016 | -0.0005 | 0.0022 |
| rs78116078 | C | 0.0086 | 0.0016 | -0.0066 | 0.0022 |
| rs7816777 | T | -0.0088 | 0.0015 | 0.0004 | 0.0021 |
| rs78193153 | A | 0.0191 | 0.0029 | -0.0092 | 0.0041 |
| rs7823700 | T | 0.0132 | 0.0024 | -0.0028 | 0.0034 |
| rs78365243 | T | 0.0181 | 0.0033 | -0.0098 | 0.0046 |
| rs78440611 | A | -0.0147 | 0.0024 | 0.0001 | 0.0034 |
| rs7849480 | A | -0.0084 | 0.0014 | 0.0057 | 0.0020 |
| rs7849487 | T | -0.0167 | 0.0015 | 0.0037 | 0.0021 |
| rs7855503 | C | 0.0088 | 0.0015 | 0.0016 | 0.0021 |
| rs78648104 | T | -0.0141 | 0.0025 | 0.0118 | 0.0036 |
| rs78702390 | C | 0.0167 | 0.0030 | -0.0051 | 0.0043 |
| rs78714229 | T | 0.0172 | 0.0030 | -0.0104 | 0.0043 |
| rs7875078 | A | -0.0078 | 0.0014 | 0.0072 | 0.0020 |
| rs78918150 | T | -0.0080 | 0.0015 | -0.0002 | 0.0021 |
| rs7894722 | T | 0.0085 | 0.0014 | -0.0080 | 0.0021 |
| rs7899270 | A | -0.0084 | 0.0015 | -0.0003 | 0.0021 |
| rs7905192 | T | 0.0084 | 0.0014 | -0.0064 | 0.0020 |
| rs790647 | A | -0.0145 | 0.0017 | 0.0077 | 0.0024 |
| rs7910403 | T | 0.0122 | 0.0018 | -0.0068 | 0.0025 |
| rs7914674 | T | -0.0139 | 0.0022 | 0.0108 | 0.0025 |
| rs7920624 | A | 0.0120 | 0.0014 | -0.0021 | 0.0020 |
| rs7924036 | T | 0.0133 | 0.0014 | -0.0102 | 0.0020 |
| rs79265434 | A | -0.0197 | 0.0022 | 0.0008 | 0.0031 |
| rs7928017 | A | 0.0096 | 0.0014 | -0.0074 | 0.0020 |
| rs7931563 | T | -0.0109 | 0.0014 | 0.0052 | 0.0021 |
| rs795230 | T | 0.0081 | 0.0014 | 0.0027 | 0.0020 |
| rs7958371 | A | -0.0085 | 0.0015 | 0.0041 | 0.0021 |
| rs79585412 | T | -0.0178 | 0.0031 | 0.0079 | 0.0043 |
| rs7965154 | A | -0.0085 | 0.0015 | 0.0038 | 0.0022 |
| rs7967550 | A | -0.0086 | 0.0014 | -0.0001 | 0.0020 |
| rs7972246 | T | 0.0110 | 0.0015 | -0.0059 | 0.0021 |
| rs79728014 | A | -0.0134 | 0.0020 | 0.0100 | 0.0028 |
| rs7974852 | A | 0.0117 | 0.0014 | -0.0040 | 0.0020 |
| rs7977614 | A | -0.0134 | 0.0016 | 0.0084 | 0.0022 |
| rs79798166 | A | 0.0172 | 0.0025 | 0.0004 | 0.0036 |
| rs79855925 | T | -0.0194 | 0.0033 | 0.0064 | 0.0048 |
| rs79994730 | T | 0.0240 | 0.0035 | -0.0221 | 0.0046 |
| rs79997166 | A | 0.0172 | 0.0030 | -0.0053 | 0.0042 |
| rs80037907 | T | -0.0107 | 0.0019 | 0.0032 | 0.0027 |
| rs8008382 | T | -0.0106 | 0.0015 | -0.0003 | 0.0022 |
| rs8009933 | A | 0.0088 | 0.0015 | -0.0047 | 0.0021 |
| rs8016504 | A | -0.0087 | 0.0014 | 0.0093 | 0.0020 |
| rs80223410 | T | -0.0118 | 0.0020 | 0.0016 | 0.0029 |
| rs8024 | A | -0.0115 | 0.0015 | 0.0069 | 0.0021 |
| rs80257979 | T | -0.0252 | 0.0040 | 0.0081 | 0.0059 |
| rs8030487 | A | 0.0085 | 0.0015 | -0.0044 | 0.0022 |
| rs803619 | T | -0.0173 | 0.0023 | 0.0061 | 0.0033 |
| rs8046072 | A | 0.0101 | 0.0018 | -0.0047 | 0.0025 |
| rs8052523 | T | -0.0084 | 0.0014 | 0.0018 | 0.0020 |
| rs8066044 | A | 0.0090 | 0.0016 | -0.0041 | 0.0023 |
| rs806816 | A | 0.0101 | 0.0016 | -0.0049 | 0.0023 |
| rs8097125 | T | 0.0086 | 0.0014 | -0.0042 | 0.0021 |
| rs8103741 | A | -0.0108 | 0.0019 | 0.0021 | 0.0026 |
| rs818415 | T | -0.0110 | 0.0018 | 0.0026 | 0.0025 |
| rs8192465 | A | -0.0272 | 0.0049 | -0.0041 | 0.0067 |
| rs852771 | T | 0.0092 | 0.0015 | -0.0005 | 0.0022 |
| rs853286 | T | 0.0124 | 0.0023 | -0.0050 | 0.0033 |
| rs854796 | A | 0.0090 | 0.0015 | 0.0000 | 0.0022 |
| rs868456 | A | 0.0089 | 0.0015 | -0.0011 | 0.0022 |
| rs870589 | A | -0.0079 | 0.0014 | 0.0063 | 0.0020 |
| rs884108 | A | -0.0094 | 0.0017 | 0.0007 | 0.0024 |
| rs891793 | C | -0.0099 | 0.0014 | -0.0015 | 0.0020 |
| rs893522 | A | 0.0149 | 0.0025 | -0.0015 | 0.0034 |
| rs902820 | A | -0.0096 | 0.0014 | 0.0035 | 0.0020 |
| rs911149 | T | 0.0091 | 0.0017 | 0.0031 | 0.0024 |
| rs912883 | T | 0.0087 | 0.0015 | -0.0074 | 0.0021 |
| rs913509 | A | 0.0096 | 0.0015 | -0.0073 | 0.0021 |
| rs925161 | C | -0.0078 | 0.0014 | 0.0052 | 0.0020 |
| rs9267658 | T | 0.0138 | 0.0020 | -0.0023 | 0.0028 |
| rs9267677 | T | 0.0180 | 0.0024 | -0.0123 | 0.0034 |
| rs9289300 | T | -0.0164 | 0.0019 | -0.0017 | 0.0027 |
| rs9291437 | C | -0.0081 | 0.0014 | 0.0059 | 0.0021 |
| rs9294770 | T | 0.0083 | 0.0014 | -0.0021 | 0.0021 |
| rs929511 | T | -0.0160 | 0.0021 | 0.0048 | 0.0030 |
| rs9300612 | T | 0.0084 | 0.0015 | -0.0028 | 0.0022 |
| rs933738 | A | -0.0110 | 0.0018 | 0.0130 | 0.0026 |
| rs9349956 | A | -0.0159 | 0.0018 | 0.0054 | 0.0026 |
| rs9359939 | A | -0.0105 | 0.0016 | 0.0056 | 0.0024 |
| rs936496 | A | -0.0079 | 0.0014 | 0.0048 | 0.0021 |
| rs9371883 | C | -0.0087 | 0.0015 | -0.0001 | 0.0021 |
| rs9373363 | A | -0.0111 | 0.0016 | 0.0060 | 0.0023 |
| rs9375188 | T | 0.0212 | 0.0014 | -0.0007 | 0.0020 |
| rs9375403 | C | 0.0126 | 0.0016 | 0.0059 | 0.0024 |
| rs9386110 | T | -0.0163 | 0.0023 | 0.0036 | 0.0032 |
| rs9388490 | T | 0.0091 | 0.0014 | -0.0066 | 0.0020 |
| rs939400 | T | -0.0095 | 0.0015 | 0.0065 | 0.0021 |
| rs9411331 | A | 0.0145 | 0.0015 | -0.0107 | 0.0021 |
| rs9435340 | A | 0.0086 | 0.0015 | -0.0121 | 0.0021 |
| rs9442750 | A | -0.0089 | 0.0016 | 0.0055 | 0.0023 |
| rs9446060 | A | 0.0077 | 0.0014 | -0.0032 | 0.0020 |
| rs9465509 | A | -0.0084 | 0.0014 | 0.0023 | 0.0020 |
| rs9490512 | A | -0.0136 | 0.0014 | 0.0050 | 0.0020 |
| rs9492774 | C | 0.0080 | 0.0015 | 0.0035 | 0.0021 |
| rs9513416 | A | -0.0116 | 0.0019 | -0.0002 | 0.0027 |
| rs9513754 | T | 0.0096 | 0.0016 | -0.0079 | 0.0022 |
| rs9527662 | A | 0.0094 | 0.0014 | -0.0074 | 0.0021 |
| rs9527905 | A | -0.0094 | 0.0014 | 0.0008 | 0.0020 |
| rs9529146 | T | 0.0127 | 0.0017 | -0.0016 | 0.0024 |
| rs9536462 | A | 0.0133 | 0.0020 | -0.0062 | 0.0029 |
| rs9540718 | A | -0.0089 | 0.0014 | 0.0069 | 0.0020 |
| rs9545395 | T | 0.0138 | 0.0021 | -0.0014 | 0.0030 |
| rs9556958 | T | -0.0110 | 0.0014 | 0.0016 | 0.0020 |
| rs9563168 | A | 0.0101 | 0.0017 | -0.0040 | 0.0025 |
| rs9568798 | T | -0.0093 | 0.0016 | 0.0007 | 0.0022 |
| rs9597907 | T | 0.0152 | 0.0027 | -0.0050 | 0.0039 |
| rs9611597 | A | -0.0107 | 0.0019 | -0.0041 | 0.0027 |
| rs9616906 | A | 0.0122 | 0.0014 | 0.0024 | 0.0020 |
| rs9616947 | T | -0.0134 | 0.0015 | 0.0029 | 0.0021 |
| rs9633970 | T | 0.0091 | 0.0016 | -0.0013 | 0.0024 |
| rs9649 | T | 0.0108 | 0.0019 | -0.0081 | 0.0027 |
| rs9655780 | A | -0.0153 | 0.0019 | 0.0038 | 0.0027 |
| rs9666728 | T | -0.0082 | 0.0014 | 0.0035 | 0.0020 |
| rs9683585 | C | 0.0084 | 0.0014 | -0.0026 | 0.0020 |
| rs969512 | A | -0.0106 | 0.0015 | 0.0139 | 0.0021 |
| rs9771228 | T | 0.0096 | 0.0015 | -0.0159 | 0.0021 |
| rs977143 | A | 0.0085 | 0.0015 | 0.0030 | 0.0021 |
| rs978807 | A | 0.0139 | 0.0018 | -0.0013 | 0.0025 |
| rs981230 | T | 0.0100 | 0.0014 | -0.0025 | 0.0020 |
| rs981883 | A | 0.0099 | 0.0014 | -0.0020 | 0.0020 |
| rs9820604 | T | 0.0115 | 0.0018 | -0.0001 | 0.0026 |
| rs9821664 | A | 0.0094 | 0.0017 | -0.0024 | 0.0025 |
| rs9830359 | T | -0.0147 | 0.0022 | 0.0115 | 0.0031 |
| rs9844755 | A | 0.0083 | 0.0015 | -0.0058 | 0.0021 |
| rs9849884 | A | -0.0090 | 0.0014 | 0.0037 | 0.0020 |
| rs9853928 | T | -0.0127 | 0.0017 | 0.0033 | 0.0025 |
| rs9858921 | A | -0.0087 | 0.0014 | 0.0019 | 0.0020 |
| rs9859556 | T | 0.0290 | 0.0015 | -0.0094 | 0.0022 |
| rs9866123 | A | 0.0086 | 0.0014 | -0.0045 | 0.0020 |
| rs9870317 | A | 0.0086 | 0.0015 | -0.0006 | 0.0021 |
| rs9877225 | T | 0.0090 | 0.0016 | -0.0060 | 0.0023 |
| rs9882532 | T | 0.0124 | 0.0015 | -0.0117 | 0.0021 |
| rs9886703 | A | -0.0136 | 0.0019 | -0.0013 | 0.0027 |
| rs9888796 | T | 0.0113 | 0.0016 | -0.0006 | 0.0023 |
| rs9916901 | T | 0.0092 | 0.0016 | -0.0100 | 0.0023 |
| rs9924031 | C | 0.0093 | 0.0015 | -0.0061 | 0.0021 |
| rs9926649 | T | 0.0177 | 0.0031 | 0.0004 | 0.0046 |
| rs9927049 | A | 0.0099 | 0.0016 | 0.0033 | 0.0023 |
| rs9927137 | A | 0.0082 | 0.0014 | -0.0062 | 0.0020 |
| rs9927842 | T | -0.0130 | 0.0020 | 0.0055 | 0.0027 |
| rs9929556 | T | -0.0099 | 0.0014 | -0.0005 | 0.0020 |
| rs9929762 | A | 0.0096 | 0.0014 | -0.0025 | 0.0020 |
| rs9929993 | T | -0.0102 | 0.0015 | 0.0008 | 0.0021 |
| rs9933256 | A | 0.0119 | 0.0014 | -0.0076 | 0.0020 |
| rs995698 | A | -0.0099 | 0.0014 | 0.0042 | 0.0020 |
| rs9974899 | T | -0.0093 | 0.0016 | 0.0010 | 0.0023 |
| rs9977825 | T | -0.0092 | 0.0015 | 0.0092 | 0.0021 |

#### Supplementary table 15: Genetic association estimates for the IVW ratio-method MR analysis estimating the effect of education on stroke. ea=effect allele; gx=education; gy=stroke; se=standard error

| SNP | ea | gx | gx_se | gy | gy_se |
| --- | --- | --- | --- | --- | --- |
| rs10006235 | T | -0.0093 | 0.0016 | 0.0003 | 0.0086 |
| rs10009513 | A | 0.0088 | 0.0014 | -0.0064 | 0.0080 |
| rs10021733 | T | -0.0164 | 0.0027 | -0.0238 | 0.0131 |
| rs10032941 | T | -0.0080 | 0.0015 | 0.0135 | 0.0081 |
| rs10041403 | T | 0.0151 | 0.0019 | -0.0167 | 0.0122 |
| rs10042828 | A | -0.0145 | 0.0022 | 0.0025 | 0.0146 |
| rs10046069 | A | 0.0127 | 0.0022 | -0.0157 | 0.0117 |
| rs10057590 | A | 0.0106 | 0.0014 | -0.0169 | 0.0080 |
| rs10060023 | T | 0.0110 | 0.0015 | -0.0144 | 0.0085 |
| rs10061420 | A | 0.0082 | 0.0014 | 0.0026 | 0.0080 |
| rs1006749 | A | 0.0079 | 0.0014 | 0.0137 | 0.0084 |
| rs10074178 | A | 0.0079 | 0.0014 | 0.0001 | 0.0078 |
| rs1007731 | A | -0.0143 | 0.0022 | -0.0170 | 0.0156 |
| rs10080647 | A | 0.0124 | 0.0020 | -0.0049 | 0.0107 |
| rs1008078 | T | -0.0182 | 0.0014 | 0.0103 | 0.0083 |
| rs1009470 | T | 0.0086 | 0.0014 | 0.0120 | 0.0079 |
| rs10098073 | A | -0.0091 | 0.0014 | -0.0059 | 0.0082 |
| rs10122669 | A | -0.0079 | 0.0014 | 0.0110 | 0.0080 |
| rs10128888 | A | -0.0083 | 0.0015 | 0.0105 | 0.0087 |
| rs10145520 | T | -0.0122 | 0.0018 | 0.0095 | 0.0098 |
| rs10166286 | T | 0.0151 | 0.0019 | 0.0069 | 0.0098 |
| rs10169002 | A | -0.0080 | 0.0014 | 0.0027 | 0.0081 |
| rs10181071 | A | 0.0102 | 0.0019 | -0.0154 | 0.0110 |
| rs10189857 | A | 0.0158 | 0.0014 | -0.0086 | 0.0084 |
| rs10192369 | A | -0.0078 | 0.0014 | 0.0048 | 0.0079 |
| rs10192834 | T | 0.0083 | 0.0015 | 0.0053 | 0.0084 |
| rs10193498 | A | 0.0097 | 0.0016 | -0.0114 | 0.0086 |
| rs10204051 | T | -0.0101 | 0.0014 | 0.0029 | 0.0085 |
| rs10205057 | T | 0.0090 | 0.0014 | -0.0148 | 0.0078 |
| rs10208 | T | 0.0091 | 0.0015 | 0.0056 | 0.0087 |
| rs10215082 | A | -0.0149 | 0.0014 | 0.0074 | 0.0086 |
| rs1024268 | T | -0.0084 | 0.0014 | 0.0014 | 0.0081 |
| rs10251438 | A | 0.0078 | 0.0014 | -0.0104 | 0.0079 |
| rs10264573 | A | 0.0175 | 0.0032 | 0.0219 | 0.0191 |
| rs1035578 | A | -0.0094 | 0.0014 | 0.0023 | 0.0080 |
| rs10402747 | T | 0.0077 | 0.0014 | 0.0010 | 0.0086 |
| rs10411759 | A | 0.0116 | 0.0020 | -0.0025 | 0.0103 |
| rs10444280 | T | 0.0100 | 0.0018 | -0.0097 | 0.0095 |
| rs10460095 | A | -0.0124 | 0.0014 | 0.0066 | 0.0082 |
| rs10480450 | A | 0.0158 | 0.0028 | -0.0070 | 0.0137 |
| rs10496091 | A | -0.0131 | 0.0016 | -0.0134 | 0.0091 |
| rs10499535 | A | 0.0080 | 0.0014 | -0.0096 | 0.0080 |
| rs1050847 | T | 0.0092 | 0.0014 | -0.0083 | 0.0091 |
| rs10509251 | T | -0.0093 | 0.0017 | 0.0103 | 0.0092 |
| rs10515007 | T | 0.0143 | 0.0019 | 0.0094 | 0.0103 |
| rs1051860 | A | -0.0079 | 0.0014 | -0.0032 | 0.0080 |
| rs10519504 | T | 0.0119 | 0.0019 | 0.0096 | 0.0107 |
| rs1054442 | A | -0.0136 | 0.0015 | 0.0114 | 0.0081 |
| rs1061801 | A | -0.0110 | 0.0018 | -0.0025 | 0.0119 |
| rs1066769 | A | -0.0230 | 0.0041 | -0.0104 | 0.0195 |
| rs10742591 | A | -0.0084 | 0.0014 | -0.0013 | 0.0082 |
| rs10748841 | A | -0.0111 | 0.0020 | -0.0042 | 0.0088 |
| rs10750539 | A | -0.0095 | 0.0015 | -0.0028 | 0.0088 |
| rs10752262 | T | 0.0102 | 0.0014 | -0.0008 | 0.0082 |
| rs1075228 | A | -0.0081 | 0.0014 | 0.0134 | 0.0092 |
| rs10761202 | T | 0.0088 | 0.0014 | -0.0028 | 0.0078 |
| rs10761251 | A | 0.0113 | 0.0015 | 0.0065 | 0.0081 |
| rs10772644 | C | 0.0142 | 0.0022 | 0.0064 | 0.0129 |
| rs10773002 | A | 0.0185 | 0.0016 | -0.0150 | 0.0088 |
| rs10773208 | T | -0.0110 | 0.0016 | 0.0016 | 0.0093 |
| rs10782651 | T | -0.0082 | 0.0014 | 0.0058 | 0.0078 |
| rs10783243 | A | -0.0092 | 0.0014 | -0.0078 | 0.0078 |
| rs10789285 | T | 0.0091 | 0.0016 | 0.0026 | 0.0088 |
| rs10793903 | A | 0.0126 | 0.0022 | -0.0072 | 0.0120 |
| rs10795831 | T | -0.0099 | 0.0017 | -0.0051 | 0.0090 |
| rs10798888 | T | -0.0140 | 0.0019 | -0.0107 | 0.0101 |
| rs10799615 | A | -0.0089 | 0.0016 | -0.0016 | 0.0086 |
| rs10805383 | A | -0.0103 | 0.0014 | 0.0172 | 0.0081 |
| rs10810099 | A | -0.0146 | 0.0016 | 0.0094 | 0.0087 |
| rs10810145 | A | 0.0078 | 0.0014 | -0.0119 | 0.0085 |
| rs10830858 | T | 0.0092 | 0.0014 | -0.0008 | 0.0081 |
| rs10831656 | T | 0.0090 | 0.0015 | -0.0073 | 0.0084 |
| rs10844179 | A | 0.0104 | 0.0017 | -0.0021 | 0.0092 |
| rs10845051 | A | 0.0083 | 0.0014 | 0.0022 | 0.0086 |
| rs10875121 | C | 0.0174 | 0.0019 | -0.0007 | 0.0111 |
| rs10877283 | T | 0.0085 | 0.0014 | -0.0001 | 0.0080 |
| rs10879676 | T | -0.0083 | 0.0014 | -0.0070 | 0.0082 |
| rs10886012 | C | 0.0078 | 0.0014 | 0.0018 | 0.0079 |
| rs10892807 | T | 0.0109 | 0.0014 | -0.0068 | 0.0083 |
| rs10899282 | A | 0.0140 | 0.0017 | -0.0168 | 0.0096 |
| rs10927053 | A | -0.0125 | 0.0022 | -0.0083 | 0.0120 |
| rs10928190 | T | -0.0087 | 0.0014 | 0.0084 | 0.0081 |
| rs10931821 | A | -0.0141 | 0.0014 | 0.0063 | 0.0079 |
| rs10937240 | T | -0.0123 | 0.0018 | -0.0047 | 0.0092 |
| rs10940540 | A | 0.0078 | 0.0014 | 0.0005 | 0.0082 |
| rs10940921 | T | 0.0111 | 0.0014 | 0.0111 | 0.0081 |
| rs10943588 | A | 0.0084 | 0.0015 | -0.0027 | 0.0082 |
| rs10947439 | T | 0.0090 | 0.0015 | -0.0059 | 0.0085 |
| rs10949263 | A | -0.0091 | 0.0016 | 0.0079 | 0.0090 |
| rs10951590 | T | -0.0109 | 0.0015 | 0.0079 | 0.0085 |
| rs10974256 | A | -0.0093 | 0.0015 | 0.0001 | 0.0084 |
| rs1097784 | T | -0.0083 | 0.0014 | 0.0011 | 0.0082 |
| rs10979613 | T | -0.0117 | 0.0015 | -0.0043 | 0.0080 |
| rs10983324 | A | 0.0084 | 0.0015 | -0.0055 | 0.0084 |
| rs10984445 | A | 0.0115 | 0.0014 | -0.0075 | 0.0080 |
| rs10985402 | T | 0.0089 | 0.0016 | 0.0033 | 0.0092 |
| rs10995639 | T | 0.0078 | 0.0014 | 0.0035 | 0.0080 |
| rs10996167 | C | 0.0113 | 0.0020 | -0.0024 | 0.0090 |
| rs11003463 | T | 0.0091 | 0.0015 | -0.0181 | 0.0099 |
| rs11021432 | A | 0.0103 | 0.0015 | 0.0066 | 0.0086 |
| rs11023764 | A | 0.0091 | 0.0015 | -0.0072 | 0.0091 |
| rs11030084 | T | 0.0106 | 0.0018 | -0.0120 | 0.0099 |
| rs11030102 | C | 0.0101 | 0.0016 | -0.0094 | 0.0102 |
| rs1106090 | A | 0.0094 | 0.0014 | -0.0035 | 0.0081 |
| rs11076962 | T | 0.0086 | 0.0016 | -0.0099 | 0.0087 |
| rs1107871 | A | -0.0083 | 0.0014 | -0.0060 | 0.0080 |
| rs11081529 | T | 0.0126 | 0.0015 | 0.0003 | 0.0088 |
| rs11082011 | T | 0.0194 | 0.0015 | 0.0049 | 0.0085 |
| rs11100237 | A | -0.0101 | 0.0014 | 0.0038 | 0.0079 |
| rs11115056 | A | 0.0187 | 0.0023 | 0.0076 | 0.0152 |
| rs11121177 | A | 0.0151 | 0.0018 | -0.0293 | 0.0105 |
| rs111226181 | T | -0.0201 | 0.0018 | 0.0319 | 0.0112 |
| rs111235962 | T | 0.0133 | 0.0024 | 0.0015 | 0.0140 |
| rs11130380 | T | 0.0090 | 0.0014 | 0.0124 | 0.0086 |
| rs111370527 | T | 0.0151 | 0.0027 | -0.0061 | 0.0185 |
| rs11138947 | T | 0.0105 | 0.0016 | -0.0050 | 0.0084 |
| rs111530150 | T | 0.0269 | 0.0048 | -0.0007 | 0.0318 |
| rs11155813 | T | -0.0129 | 0.0023 | 0.0076 | 0.0109 |
| rs11157931 | A | -0.0139 | 0.0014 | -0.0005 | 0.0084 |
| rs11158800 | A | -0.0091 | 0.0014 | -0.0005 | 0.0079 |
| rs1117152 | T | 0.0106 | 0.0015 | -0.0096 | 0.0085 |
| rs11174399 | A | 0.0111 | 0.0018 | -0.0083 | 0.0099 |
| rs111821073 | T | 0.0124 | 0.0019 | -0.0041 | 0.0114 |
| rs111852224 | T | 0.0156 | 0.0022 | -0.0132 | 0.0144 |
| rs11190955 | T | -0.0083 | 0.0015 | -0.0069 | 0.0086 |
| rs11191237 | T | 0.0132 | 0.0024 | 0.0073 | 0.0119 |
| rs1120924 | T | 0.0108 | 0.0015 | -0.0043 | 0.0087 |
| rs11209894 | A | 0.0119 | 0.0021 | 0.0406 | 0.0115 |
| rs11210228 | T | -0.0105 | 0.0019 | -0.0016 | 0.0087 |
| rs11210400 | A | 0.0120 | 0.0014 | -0.0045 | 0.0083 |
| rs11211123 | A | 0.0104 | 0.0017 | -0.0168 | 0.0091 |
| rs11213482 | A | -0.0106 | 0.0019 | -0.0159 | 0.0132 |
| rs112210983 | A | 0.0105 | 0.0017 | 0.0089 | 0.0104 |
| rs1123285 | C | -0.0094 | 0.0015 | 0.0124 | 0.0085 |
| rs112375785 | A | 0.0184 | 0.0021 | -0.0046 | 0.0148 |
| rs112379405 | C | -0.0082 | 0.0014 | -0.0151 | 0.0084 |
| rs112512729 | T | -0.0166 | 0.0027 | 0.0079 | 0.0174 |
| rs112603734 | A | -0.0131 | 0.0022 | 0.0131 | 0.0106 |
| rs11265191 | T | 0.0083 | 0.0015 | -0.0014 | 0.0082 |
| rs112806496 | C | -0.0160 | 0.0025 | 0.0036 | 0.0167 |
| rs1128956 | T | -0.0140 | 0.0019 | -0.0084 | 0.0109 |
| rs113205706 | A | -0.0135 | 0.0024 | 0.0182 | 0.0156 |
| rs113520408 | A | 0.0155 | 0.0016 | -0.0111 | 0.0100 |
| rs113588399 | T | -0.0104 | 0.0017 | -0.0075 | 0.0106 |
| rs113615161 | T | -0.0123 | 0.0021 | 0.0088 | 0.0115 |
| rs113720505 | A | 0.0164 | 0.0029 | 0.0140 | 0.0214 |
| rs114192810 | T | -0.0279 | 0.0051 | 0.0445 | 0.0340 |
| rs114468556 | A | 0.0293 | 0.0036 | -0.0170 | 0.0263 |
| rs114593137 | A | -0.0122 | 0.0017 | 0.0083 | 0.0105 |
| rs1146079 | C | -0.0087 | 0.0016 | -0.0032 | 0.0092 |
| rs114810763 | A | -0.0121 | 0.0020 | -0.0156 | 0.0147 |
| rs114952970 | T | 0.0265 | 0.0040 | 0.0243 | 0.0274 |
| rs114958262 | T | -0.0220 | 0.0039 | -0.0242 | 0.0285 |
| rs115000530 | A | -0.0250 | 0.0031 | -0.0002 | 0.0205 |
| rs115017135 | A | -0.0206 | 0.0038 | 0.0255 | 0.0271 |
| rs11542663 | A | 0.0112 | 0.0015 | 0.0085 | 0.0084 |
| rs115438240 | T | -0.0169 | 0.0030 | 0.0250 | 0.0193 |
| rs115693355 | A | 0.0309 | 0.0053 | -0.0045 | 0.0364 |
| rs11588857 | A | 0.0199 | 0.0017 | 0.0141 | 0.0096 |
| rs11591870 | T | 0.0098 | 0.0017 | -0.0044 | 0.0115 |
| rs11592299 | A | 0.0097 | 0.0018 | -0.0063 | 0.0115 |
| rs11596387 | T | 0.0258 | 0.0046 | 0.0450 | 0.0345 |
| rs11598765 | A | -0.0106 | 0.0018 | 0.0062 | 0.0099 |
| rs11599236 | T | -0.0106 | 0.0014 | -0.0113 | 0.0081 |
| rs11613431 | A | -0.0094 | 0.0016 | -0.0030 | 0.0099 |
| rs11620355 | A | 0.0182 | 0.0025 | -0.0127 | 0.0160 |
| rs11623285 | T | -0.0135 | 0.0021 | 0.0114 | 0.0129 |
| rs116386746 | C | 0.0249 | 0.0045 | -0.0321 | 0.0316 |
| rs11640569 | A | 0.0112 | 0.0020 | -0.0053 | 0.0128 |
| rs11644446 | A | 0.0117 | 0.0019 | -0.0329 | 0.0120 |
| rs11647188 | A | 0.0079 | 0.0014 | -0.0029 | 0.0083 |
| rs11652522 | A | -0.0154 | 0.0023 | 0.0092 | 0.0136 |
| rs11657342 | A | 0.0140 | 0.0019 | -0.0005 | 0.0105 |
| rs11657979 | A | -0.0091 | 0.0017 | 0.0037 | 0.0096 |
| rs11663602 | A | -0.0130 | 0.0016 | 0.0105 | 0.0094 |
| rs11663678 | T | -0.0121 | 0.0022 | 0.0273 | 0.0132 |
| rs116656374 | T | -0.0242 | 0.0038 | -0.0157 | 0.0289 |
| rs11675476 | T | -0.0127 | 0.0014 | 0.0130 | 0.0079 |
| rs1167827 | A | 0.0104 | 0.0014 | -0.0328 | 0.0093 |
| rs11678980 | A | -0.0166 | 0.0014 | 0.0024 | 0.0082 |
| rs11687736 | A | -0.0105 | 0.0015 | 0.0127 | 0.0086 |
| rs11693885 | A | -0.0091 | 0.0014 | 0.0046 | 0.0081 |
| rs116967397 | A | 0.0315 | 0.0056 | 0.0075 | 0.0397 |
| rs117005905 | T | 0.0150 | 0.0022 | -0.0219 | 0.0126 |
| rs11703948 | A | -0.0168 | 0.0024 | -0.0041 | 0.0134 |
| rs1171040 | A | -0.0113 | 0.0017 | 0.0079 | 0.0094 |
| rs11716398 | A | 0.0159 | 0.0021 | -0.0173 | 0.0103 |
| rs11720093 | T | 0.0150 | 0.0026 | -0.0132 | 0.0138 |
| rs11720985 | T | 0.0085 | 0.0015 | 0.0103 | 0.0094 |
| rs11724690 | T | 0.0093 | 0.0015 | 0.0018 | 0.0095 |
| rs117273411 | T | -0.0211 | 0.0034 | 0.0329 | 0.0249 |
| rs11732160 | A | 0.0105 | 0.0016 | -0.0054 | 0.0091 |
| rs11732657 | A | -0.0126 | 0.0016 | -0.0081 | 0.0091 |
| rs11733439 | A | -0.0096 | 0.0017 | 0.0167 | 0.0096 |
| rs11736863 | A | 0.0142 | 0.0018 | -0.0064 | 0.0107 |
| rs117398064 | C | -0.0141 | 0.0025 | -0.0122 | 0.0174 |
| rs117468730 | A | -0.0310 | 0.0050 | -0.0494 | 0.0338 |
| rs117623407 | A | -0.0118 | 0.0020 | 0.0011 | 0.0136 |
| rs11765387 | T | 0.0086 | 0.0015 | -0.0074 | 0.0083 |
| rs117668569 | T | 0.0234 | 0.0042 | -0.0348 | 0.0277 |
| rs117677995 | A | -0.0180 | 0.0032 | 0.0191 | 0.0253 |
| rs11772232 | T | 0.0177 | 0.0019 | -0.0217 | 0.0114 |
| rs11774212 | T | 0.0128 | 0.0014 | 0.0007 | 0.0085 |
| rs117799466 | C | 0.0106 | 0.0016 | -0.0043 | 0.0101 |
| rs11780023 | T | 0.0085 | 0.0014 | 0.0144 | 0.0084 |
| rs11789013 | T | 0.0106 | 0.0016 | -0.0009 | 0.0095 |
| rs118040169 | A | -0.0248 | 0.0038 | 0.0166 | 0.0258 |
| rs118093058 | T | 0.0146 | 0.0021 | 0.0284 | 0.0123 |
| rs118134876 | T | -0.0223 | 0.0030 | 0.0270 | 0.0207 |
| rs11845781 | T | -0.0079 | 0.0014 | 0.0099 | 0.0080 |
| rs11871429 | A | 0.0128 | 0.0017 | -0.0178 | 0.0091 |
| rs11876620 | T | 0.0143 | 0.0024 | -0.0043 | 0.0149 |
| rs11894424 | A | 0.0209 | 0.0033 | 0.0036 | 0.0137 |
| rs11917701 | A | -0.0087 | 0.0014 | 0.0085 | 0.0081 |
| rs11919835 | C | 0.0089 | 0.0015 | -0.0079 | 0.0085 |
| rs1196760 | C | 0.0139 | 0.0025 | -0.0084 | 0.0161 |
| rs12005151 | A | 0.0087 | 0.0015 | -0.0125 | 0.0085 |
| rs12028010 | T | 0.0164 | 0.0017 | 0.0028 | 0.0093 |
| rs12028229 | T | 0.0099 | 0.0016 | -0.0069 | 0.0117 |
| rs12054166 | C | 0.0096 | 0.0016 | 0.0153 | 0.0095 |
| rs12076635 | C | 0.0207 | 0.0017 | -0.0041 | 0.0091 |
| rs12113634 | T | 0.0088 | 0.0015 | 0.0064 | 0.0083 |
| rs12123293 | T | 0.0086 | 0.0015 | -0.0015 | 0.0083 |
| rs12126231 | A | 0.0095 | 0.0014 | -0.0183 | 0.0082 |
| rs12127928 | T | 0.0108 | 0.0018 | 0.0002 | 0.0109 |
| rs12145078 | A | -0.0127 | 0.0017 | 0.0168 | 0.0101 |
| rs12151248 | T | -0.0149 | 0.0023 | -0.0235 | 0.0150 |
| rs12170452 | A | 0.0105 | 0.0014 | -0.0138 | 0.0085 |
| rs1220779 | A | -0.0079 | 0.0014 | -0.0015 | 0.0083 |
| rs12234369 | A | 0.0087 | 0.0015 | -0.0088 | 0.0089 |
| rs12238011 | T | -0.0141 | 0.0026 | 0.0104 | 0.0136 |
| rs12273435 | A | -0.0132 | 0.0017 | 0.0081 | 0.0114 |
| rs12285074 | A | 0.0161 | 0.0023 | 0.0194 | 0.0107 |
| rs12304188 | A | 0.0111 | 0.0019 | -0.0199 | 0.0100 |
| rs12325727 | A | 0.0088 | 0.0014 | -0.0152 | 0.0081 |
| rs12332731 | A | 0.0135 | 0.0018 | 0.0020 | 0.0106 |
| rs12342546 | A | -0.0128 | 0.0017 | -0.0047 | 0.0095 |
| rs12359372 | T | -0.0092 | 0.0015 | 0.0149 | 0.0086 |
| rs12375949 | T | -0.0138 | 0.0014 | 0.0023 | 0.0081 |
| rs12405889 | T | -0.0098 | 0.0014 | -0.0075 | 0.0078 |
| rs12431682 | T | -0.0093 | 0.0015 | 0.0017 | 0.0082 |
| rs12438177 | A | 0.0093 | 0.0015 | 0.0028 | 0.0081 |
| rs12453682 | T | 0.0102 | 0.0015 | 0.0062 | 0.0083 |
| rs1245829 | A | -0.0099 | 0.0014 | 0.0032 | 0.0088 |
| rs12467175 | T | -0.0081 | 0.0014 | -0.0070 | 0.0080 |
| rs12468040 | T | 0.0137 | 0.0014 | 0.0015 | 0.0082 |
| rs12473986 | T | -0.0085 | 0.0015 | 0.0057 | 0.0085 |
| rs12477385 | T | 0.0109 | 0.0017 | -0.0275 | 0.0096 |
| rs12478156 | T | 0.0108 | 0.0015 | -0.0042 | 0.0083 |
| rs12503522 | T | -0.0113 | 0.0016 | 0.0100 | 0.0086 |
| rs12506221 | T | -0.0130 | 0.0014 | -0.0026 | 0.0079 |
| rs12516990 | C | -0.0099 | 0.0017 | -0.0012 | 0.0105 |
| rs12519073 | T | -0.0107 | 0.0017 | 0.0149 | 0.0098 |
| rs12524795 | T | 0.0102 | 0.0014 | -0.0071 | 0.0081 |
| rs12568153 | T | -0.0091 | 0.0015 | 0.0030 | 0.0088 |
| rs12571549 | A | 0.0151 | 0.0020 | -0.0006 | 0.0102 |
| rs12574281 | A | -0.0080 | 0.0015 | 0.0029 | 0.0090 |
| rs12591647 | T | -0.0152 | 0.0018 | 0.0149 | 0.0096 |
| rs12601380 | A | -0.0082 | 0.0014 | -0.0102 | 0.0082 |
| rs12602286 | T | 0.0160 | 0.0021 | -0.0160 | 0.0110 |
| rs12613500 | C | 0.0098 | 0.0014 | -0.0062 | 0.0081 |
| rs12614263 | A | 0.0083 | 0.0014 | -0.0026 | 0.0080 |
| rs12620796 | A | 0.0121 | 0.0019 | -0.0075 | 0.0086 |
| rs12638072 | A | -0.0107 | 0.0015 | 0.0040 | 0.0092 |
| rs12643771 | T | 0.0130 | 0.0015 | -0.0131 | 0.0090 |
| rs12646216 | T | 0.0084 | 0.0014 | -0.0161 | 0.0083 |
| rs12646523 | T | -0.0132 | 0.0016 | 0.0024 | 0.0090 |
| rs12659776 | T | 0.0085 | 0.0014 | -0.0079 | 0.0079 |
| rs12670376 | A | 0.0092 | 0.0014 | -0.0152 | 0.0079 |
| rs1267062 | C | 0.0113 | 0.0018 | -0.0029 | 0.0111 |
| rs12682775 | T | -0.0119 | 0.0017 | 0.0230 | 0.0094 |
| rs12694681 | T | 0.0102 | 0.0015 | -0.0092 | 0.0085 |
| rs12709186 | A | -0.0091 | 0.0015 | -0.0037 | 0.0090 |
| rs12712269 | T | -0.0107 | 0.0014 | 0.0062 | 0.0081 |
| rs12746551 | C | 0.0248 | 0.0041 | 0.0161 | 0.0227 |
| rs12750688 | C | 0.0116 | 0.0016 | -0.0077 | 0.0100 |
| rs12761729 | T | -0.0098 | 0.0016 | 0.0168 | 0.0105 |
| rs12761761 | T | 0.0153 | 0.0017 | 0.0033 | 0.0103 |
| rs12764593 | C | 0.0202 | 0.0029 | -0.0426 | 0.0198 |
| rs12765185 | A | -0.0088 | 0.0016 | -0.0102 | 0.0096 |
| rs12774577 | T | 0.0142 | 0.0020 | -0.0556 | 0.0119 |
| rs12789313 | T | 0.0092 | 0.0014 | -0.0097 | 0.0082 |
| rs12790196 | T | -0.0101 | 0.0015 | 0.0129 | 0.0089 |
| rs12810587 | T | 0.0094 | 0.0015 | -0.0012 | 0.0087 |
| rs12875339 | A | -0.0108 | 0.0015 | 0.0003 | 0.0086 |
| rs12888615 | T | 0.0120 | 0.0018 | -0.0170 | 0.0113 |
| rs12891042 | T | 0.0092 | 0.0014 | 0.0027 | 0.0080 |
| rs12891191 | C | 0.0080 | 0.0014 | -0.0059 | 0.0084 |
| rs12912465 | T | -0.0104 | 0.0016 | 0.0027 | 0.0106 |
| rs12952191 | T | -0.0081 | 0.0014 | 0.0102 | 0.0081 |
| rs12953422 | A | -0.0080 | 0.0014 | 0.0102 | 0.0086 |
| rs12957463 | A | 0.0146 | 0.0018 | -0.0073 | 0.0095 |
| rs12962845 | C | -0.0123 | 0.0021 | 0.0130 | 0.0115 |
| rs12967010 | T | 0.0114 | 0.0017 | -0.0041 | 0.0101 |
| rs12970264 | A | -0.0085 | 0.0014 | -0.0254 | 0.0092 |
| rs12981405 | T | -0.0112 | 0.0019 | -0.0055 | 0.0102 |
| rs13009915 | T | 0.0212 | 0.0037 | -0.0085 | 0.0259 |
| rs13010288 | T | 0.0198 | 0.0021 | -0.0297 | 0.0129 |
| rs13010566 | A | -0.0091 | 0.0014 | -0.0069 | 0.0083 |
| rs13015496 | C | 0.0119 | 0.0018 | -0.0007 | 0.0096 |
| rs13016316 | A | -0.0138 | 0.0024 | -0.0099 | 0.0124 |
| rs1301838 | T | -0.0096 | 0.0015 | -0.0003 | 0.0089 |
| rs13018640 | T | -0.0215 | 0.0014 | 0.0054 | 0.0081 |
| rs13026611 | A | -0.0084 | 0.0015 | 0.0066 | 0.0085 |
| rs13029602 | T | -0.0092 | 0.0014 | -0.0020 | 0.0083 |
| rs13034349 | T | -0.0082 | 0.0014 | 0.0177 | 0.0084 |
| rs13050131 | A | -0.0081 | 0.0015 | 0.0009 | 0.0088 |
| rs13085461 | C | 0.0103 | 0.0014 | -0.0058 | 0.0079 |
| rs13099165 | T | -0.0118 | 0.0017 | 0.0142 | 0.0105 |
| rs13107325 | T | -0.0239 | 0.0027 | -0.0005 | 0.0193 |
| rs13117856 | A | 0.0084 | 0.0015 | -0.0017 | 0.0091 |
| rs13130765 | C | -0.0091 | 0.0014 | 0.0155 | 0.0088 |
| rs13133213 | A | 0.0096 | 0.0014 | -0.0201 | 0.0079 |
| rs13145650 | T | -0.0173 | 0.0025 | 0.0013 | 0.0134 |
| rs13154429 | C | -0.0136 | 0.0023 | 0.0123 | 0.0142 |
| rs13163062 | T | 0.0115 | 0.0014 | -0.0079 | 0.0080 |
| rs13163845 | T | -0.0149 | 0.0020 | -0.0104 | 0.0112 |
| rs13168136 | A | -0.0092 | 0.0017 | 0.0104 | 0.0089 |
| rs13169187 | A | -0.0079 | 0.0014 | 0.0024 | 0.0080 |
| rs13177031 | A | -0.0085 | 0.0015 | 0.0012 | 0.0084 |
| rs13190235 | T | -0.0138 | 0.0024 | 0.0048 | 0.0119 |
| rs13197257 | T | 0.0109 | 0.0016 | -0.0109 | 0.0093 |
| rs1320139 | C | -0.0149 | 0.0014 | 0.0082 | 0.0080 |
| rs13212041 | T | 0.0101 | 0.0018 | 0.0049 | 0.0094 |
| rs13240401 | T | 0.0176 | 0.0017 | -0.0142 | 0.0101 |
| rs13246220 | T | -0.0093 | 0.0014 | -0.0079 | 0.0082 |
| rs13261773 | C | 0.0122 | 0.0019 | 0.0058 | 0.0108 |
| rs13266287 | A | -0.0085 | 0.0015 | 0.0159 | 0.0084 |
| rs13281564 | A | 0.0088 | 0.0014 | 0.0015 | 0.0083 |
| rs13284516 | C | -0.0082 | 0.0014 | -0.0054 | 0.0087 |
| rs1329044 | T | 0.0134 | 0.0019 | -0.0026 | 0.0105 |
| rs1329125 | T | -0.0097 | 0.0015 | -0.0042 | 0.0092 |
| rs13296345 | T | 0.0090 | 0.0015 | -0.0202 | 0.0081 |
| rs13327482 | A | -0.0121 | 0.0018 | 0.0148 | 0.0115 |
| rs1334297 | A | 0.0257 | 0.0016 | -0.0066 | 0.0092 |
| rs1335482 | T | 0.0096 | 0.0014 | -0.0221 | 0.0089 |
| rs13361043 | T | -0.0158 | 0.0028 | 0.0062 | 0.0127 |
| rs13381557 | A | -0.0087 | 0.0014 | -0.0010 | 0.0080 |
| rs13388333 | A | 0.0085 | 0.0015 | 0.0040 | 0.0087 |
| rs13397529 | C | -0.0097 | 0.0017 | 0.0085 | 0.0102 |
| rs13398860 | A | 0.0086 | 0.0015 | 0.0028 | 0.0085 |
| rs13402497 | A | -0.0080 | 0.0014 | 0.0075 | 0.0077 |
| rs13422673 | T | -0.0110 | 0.0014 | -0.0042 | 0.0083 |
| rs13425585 | C | -0.0089 | 0.0014 | 0.0134 | 0.0080 |
| rs1368250 | T | 0.0250 | 0.0045 | 0.0055 | 0.0367 |
| rs137079 | T | 0.0128 | 0.0020 | 0.0039 | 0.0119 |
| rs137858393 | A | 0.0193 | 0.0029 | -0.0163 | 0.0194 |
| rs138096147 | A | 0.0234 | 0.0042 | 0.0273 | 0.0320 |
| rs1391513 | T | -0.0095 | 0.0015 | 0.0127 | 0.0087 |
| rs139244147 | A | -0.0193 | 0.0030 | 0.0006 | 0.0166 |
| rs1392816 | T | 0.0082 | 0.0014 | -0.0117 | 0.0085 |
| rs139980871 | T | 0.0180 | 0.0029 | -0.0069 | 0.0192 |
| rs1404549 | A | -0.0089 | 0.0015 | 0.0047 | 0.0089 |
| rs1405876 | T | 0.0109 | 0.0015 | 0.0059 | 0.0087 |
| rs140711597 | C | 0.0366 | 0.0055 | -0.0578 | 0.0418 |
| rs1408284 | C | -0.0127 | 0.0020 | 0.0334 | 0.0131 |
| rs1408430 | C | 0.0096 | 0.0016 | 0.0089 | 0.0088 |
| rs141586924 | A | -0.0090 | 0.0015 | -0.0016 | 0.0098 |
| rs141729694 | T | 0.0195 | 0.0026 | 0.0160 | 0.0151 |
| rs142014757 | A | 0.0102 | 0.0018 | -0.0143 | 0.0131 |
| rs1426619 | T | 0.0096 | 0.0014 | 0.0041 | 0.0079 |
| rs1427298 | T | 0.0086 | 0.0014 | -0.0096 | 0.0083 |
| rs142747148 | A | 0.0329 | 0.0052 | -0.0206 | 0.0371 |
| rs1427829 | A | -0.0083 | 0.0014 | 0.0170 | 0.0080 |
| rs143148393 | T | -0.0211 | 0.0039 | -0.0213 | 0.0271 |
| rs1434630 | T | -0.0137 | 0.0020 | -0.0147 | 0.0108 |
| rs143743568 | A | 0.0118 | 0.0020 | -0.0001 | 0.0116 |
| rs143812851 | A | -0.0113 | 0.0019 | 0.0061 | 0.0118 |
| rs1440930 | C | 0.0081 | 0.0014 | -0.0024 | 0.0084 |
| rs144336753 | A | 0.0388 | 0.0053 | 0.0087 | 0.0357 |
| rs1445591 | A | 0.0085 | 0.0015 | -0.0039 | 0.0098 |
| rs145018899 | A | 0.0095 | 0.0015 | -0.0006 | 0.0092 |
| rs1464297 | T | -0.0109 | 0.0015 | 0.0013 | 0.0084 |
| rs1467737 | T | -0.0084 | 0.0014 | 0.0022 | 0.0082 |
| rs1485300 | C | 0.0098 | 0.0015 | -0.0103 | 0.0086 |
| rs1490612 | T | 0.0193 | 0.0026 | -0.0228 | 0.0169 |
| rs150252215 | A | 0.0204 | 0.0035 | -0.0201 | 0.0224 |
| rs150537577 | A | -0.0151 | 0.0026 | -0.0209 | 0.0140 |
| rs1505676 | C | 0.0084 | 0.0015 | -0.0041 | 0.0082 |
| rs151381 | T | -0.0087 | 0.0014 | 0.0057 | 0.0080 |
| rs1518890 | T | -0.0090 | 0.0016 | -0.0080 | 0.0090 |
| rs1527878 | A | -0.0105 | 0.0016 | 0.0020 | 0.0098 |
| rs1529597 | A | 0.0223 | 0.0038 | -0.0330 | 0.0236 |
| rs1538389 | T | -0.0109 | 0.0018 | 0.0053 | 0.0103 |
| rs1542354 | A | -0.0084 | 0.0014 | 0.0018 | 0.0082 |
| rs1544 | A | -0.0096 | 0.0016 | 0.0090 | 0.0090 |
| rs1554798 | A | -0.0080 | 0.0014 | 0.0134 | 0.0081 |
| rs1564347 | T | 0.0088 | 0.0015 | 0.0024 | 0.0084 |
| rs1569092 | A | 0.0160 | 0.0020 | -0.0013 | 0.0111 |
| rs1593022 | T | -0.0099 | 0.0017 | 0.0041 | 0.0099 |
| rs1603460 | T | 0.0090 | 0.0014 | -0.0081 | 0.0081 |
| rs162445 | A | 0.0152 | 0.0026 | 0.0236 | 0.0121 |
| rs163229 | C | -0.0260 | 0.0047 | 0.0090 | 0.0311 |
| rs1637770 | T | 0.0191 | 0.0028 | 0.0306 | 0.0174 |
| rs164938 | T | -0.0088 | 0.0014 | 0.0107 | 0.0081 |
| rs165633 | A | -0.0116 | 0.0016 | 0.0074 | 0.0101 |
| rs1656614 | C | 0.0122 | 0.0015 | -0.0105 | 0.0094 |
| rs1671269 | T | -0.0100 | 0.0016 | -0.0110 | 0.0088 |
| rs1671770 | A | 0.0125 | 0.0018 | -0.0051 | 0.0104 |
| rs16851779 | T | -0.0153 | 0.0027 | 0.0001 | 0.0146 |
| rs16871807 | T | 0.0090 | 0.0015 | 0.0052 | 0.0083 |
| rs1689510 | C | 0.0194 | 0.0015 | -0.0187 | 0.0088 |
| rs16901689 | T | 0.0111 | 0.0019 | -0.0045 | 0.0107 |
| rs1693584 | T | -0.0087 | 0.0015 | 0.0180 | 0.0088 |
| rs16958559 | T | 0.0096 | 0.0017 | 0.0092 | 0.0106 |
| rs16966271 | T | 0.0092 | 0.0016 | 0.0034 | 0.0087 |
| rs16975275 | A | -0.0102 | 0.0017 | 0.0049 | 0.0095 |
| rs17048801 | A | -0.0106 | 0.0014 | 0.0099 | 0.0085 |
| rs17048855 | A | 0.0108 | 0.0015 | -0.0128 | 0.0083 |
| rs17069646 | T | 0.0085 | 0.0015 | 0.0032 | 0.0082 |
| rs17110109 | T | -0.0093 | 0.0014 | 0.0262 | 0.0081 |
| rs17113730 | A | -0.0170 | 0.0027 | 0.0041 | 0.0165 |
| rs17131123 | A | -0.0243 | 0.0039 | 0.0484 | 0.0289 |
| rs17133297 | A | -0.0146 | 0.0023 | 0.0075 | 0.0117 |
| rs17144467 | A | 0.0098 | 0.0015 | -0.0060 | 0.0084 |
| rs17148998 | A | 0.0101 | 0.0017 | -0.0060 | 0.0108 |
| rs171697 | C | 0.0133 | 0.0015 | -0.0200 | 0.0093 |
| rs17170519 | C | 0.0132 | 0.0023 | -0.0074 | 0.0114 |
| rs1717204 | A | -0.0117 | 0.0018 | 0.0093 | 0.0112 |
| rs1718188 | A | 0.0096 | 0.0014 | 0.0011 | 0.0079 |
| rs17186106 | T | 0.0108 | 0.0018 | -0.0281 | 0.0109 |
| rs17190418 | T | -0.0213 | 0.0031 | 0.0199 | 0.0210 |
| rs17205908 | T | -0.0092 | 0.0015 | -0.0072 | 0.0097 |
| rs17248751 | A | -0.0136 | 0.0017 | 0.0152 | 0.0111 |
| rs1728118 | A | 0.0089 | 0.0016 | -0.0013 | 0.0085 |
| rs1729412 | T | -0.0107 | 0.0014 | -0.0010 | 0.0080 |
| rs173003 | A | -0.0082 | 0.0014 | -0.0022 | 0.0083 |
| rs1734370 | A | 0.0099 | 0.0016 | -0.0036 | 0.0088 |
| rs1738050 | C | -0.0093 | 0.0014 | 0.0066 | 0.0084 |
| rs17411339 | A | 0.0140 | 0.0014 | -0.0101 | 0.0088 |
| rs17428076 | C | 0.0137 | 0.0016 | -0.0089 | 0.0099 |
| rs17440885 | A | 0.0160 | 0.0029 | 0.0177 | 0.0187 |
| rs1747714 | T | 0.0106 | 0.0014 | -0.0143 | 0.0084 |
| rs1747817 | T | -0.0092 | 0.0016 | 0.0171 | 0.0100 |
| rs17489649 | A | 0.0127 | 0.0015 | -0.0146 | 0.0093 |
| rs17502934 | T | -0.0167 | 0.0020 | 0.0061 | 0.0120 |
| rs175325 | A | -0.0100 | 0.0014 | 0.0071 | 0.0084 |
| rs17536059 | C | -0.0108 | 0.0019 | 0.0029 | 0.0114 |
| rs17551064 | A | 0.0141 | 0.0019 | 0.0025 | 0.0111 |
| rs17563464 | A | -0.0147 | 0.0017 | 0.0295 | 0.0119 |
| rs17565975 | A | -0.0106 | 0.0014 | -0.0042 | 0.0080 |
| rs17568389 | A | 0.0127 | 0.0014 | -0.0004 | 0.0081 |
| rs17574007 | A | -0.0129 | 0.0021 | 0.0114 | 0.0137 |
| rs1758747 | A | 0.0092 | 0.0015 | -0.0072 | 0.0085 |
| rs17598675 | T | -0.0116 | 0.0014 | 0.0048 | 0.0085 |
| rs17604349 | A | 0.0127 | 0.0018 | -0.0036 | 0.0111 |
| rs17609255 | T | -0.0087 | 0.0014 | 0.0125 | 0.0081 |
| rs176218 | T | 0.0200 | 0.0018 | -0.0010 | 0.0094 |
| rs17622379 | T | -0.0113 | 0.0018 | 0.0094 | 0.0105 |
| rs17650634 | T | -0.0104 | 0.0019 | 0.0088 | 0.0117 |
| rs17667540 | A | -0.0087 | 0.0015 | -0.0025 | 0.0081 |
| rs17669337 | T | -0.0110 | 0.0014 | -0.0085 | 0.0080 |
| rs17680712 | T | -0.0092 | 0.0014 | 0.0050 | 0.0078 |
| rs17686649 | T | -0.0098 | 0.0017 | 0.0164 | 0.0102 |
| rs17732878 | T | -0.0095 | 0.0017 | -0.0069 | 0.0103 |
| rs17742342 | A | -0.0119 | 0.0018 | 0.0006 | 0.0105 |
| rs17747544 | A | -0.0090 | 0.0014 | -0.0008 | 0.0082 |
| rs1779549 | A | 0.0081 | 0.0014 | -0.0148 | 0.0080 |
| rs178183 | T | 0.0145 | 0.0016 | -0.0018 | 0.0091 |
| rs17882802 | A | 0.0084 | 0.0014 | -0.0153 | 0.0086 |
| rs17883331 | A | -0.0108 | 0.0018 | 0.0074 | 0.0112 |
| rs1792602 | A | -0.0097 | 0.0014 | 0.0082 | 0.0082 |
| rs181214 | T | -0.0104 | 0.0017 | 0.0087 | 0.0113 |
| rs182902112 | A | -0.0300 | 0.0053 | -0.0179 | 0.0357 |
| rs1835339 | T | 0.0084 | 0.0014 | 0.0105 | 0.0082 |
| rs183869217 | C | 0.0174 | 0.0030 | 0.0077 | 0.0189 |
| rs1841023 | A | -0.0092 | 0.0015 | -0.0013 | 0.0092 |
| rs1842713 | A | -0.0117 | 0.0017 | -0.0313 | 0.0093 |
| rs1843815 | A | 0.0077 | 0.0014 | -0.0146 | 0.0080 |
| rs1861786 | A | -0.0081 | 0.0014 | 0.0046 | 0.0081 |
| rs1865955 | T | 0.0122 | 0.0019 | -0.0076 | 0.0096 |
| rs1866823 | A | 0.0107 | 0.0014 | 0.0009 | 0.0079 |
| rs187580 | T | -0.0104 | 0.0017 | -0.0028 | 0.0106 |
| rs187951956 | A | -0.0315 | 0.0058 | 0.0283 | 0.0418 |
| rs1880088 | A | -0.0097 | 0.0016 | 0.0184 | 0.0089 |
| rs1880692 | A | 0.0086 | 0.0014 | 0.0039 | 0.0081 |
| rs1890132 | T | -0.0089 | 0.0015 | 0.0114 | 0.0098 |
| rs1898111 | A | 0.0101 | 0.0018 | -0.0051 | 0.0098 |
| rs190102446 | T | -0.0210 | 0.0037 | 0.0167 | 0.0241 |
| rs1905616 | A | 0.0083 | 0.0015 | -0.0035 | 0.0086 |
| rs1910005 | T | -0.0090 | 0.0015 | -0.0046 | 0.0090 |
| rs1918394 | T | 0.0119 | 0.0019 | -0.0080 | 0.0103 |
| rs192436652 | T | -0.0319 | 0.0045 | 0.0266 | 0.0318 |
| rs1933264 | T | 0.0100 | 0.0016 | -0.0090 | 0.0090 |
| rs1952183 | A | -0.0089 | 0.0014 | -0.0017 | 0.0082 |
| rs1955250 | A | -0.0140 | 0.0025 | 0.0105 | 0.0133 |
| rs1963381 | A | 0.0099 | 0.0016 | 0.0147 | 0.0090 |
| rs1980129 | A | 0.0085 | 0.0014 | 0.0016 | 0.0079 |
| rs198262 | T | -0.0223 | 0.0035 | 0.0039 | 0.0227 |
| rs1991585 | T | -0.0101 | 0.0015 | 0.0065 | 0.0087 |
| rs1995181 | A | -0.0078 | 0.0014 | -0.0036 | 0.0077 |
| rs2007655 | T | 0.0086 | 0.0014 | -0.0031 | 0.0078 |
| rs2011603 | A | -0.0091 | 0.0016 | 0.0262 | 0.0089 |
| rs2014830 | T | 0.0110 | 0.0015 | -0.0106 | 0.0088 |
| rs201495 | T | -0.0112 | 0.0019 | 0.0003 | 0.0106 |
| rs2023016 | C | 0.0107 | 0.0017 | -0.0169 | 0.0093 |
| rs2024568 | T | -0.0097 | 0.0016 | 0.0280 | 0.0089 |
| rs2029401 | A | -0.0091 | 0.0014 | 0.0089 | 0.0082 |
| rs2034631 | T | 0.0089 | 0.0015 | -0.0152 | 0.0082 |
| rs2039204 | A | 0.0082 | 0.0014 | -0.0088 | 0.0079 |
| rs2043187 | A | 0.0107 | 0.0015 | -0.0108 | 0.0090 |
| rs2052285 | A | 0.0112 | 0.0014 | 0.0017 | 0.0084 |
| rs2055940 | A | 0.0082 | 0.0015 | -0.0161 | 0.0089 |
| rs2077235 | T | 0.0097 | 0.0017 | 0.0013 | 0.0090 |
| rs2081652 | A | 0.0123 | 0.0015 | -0.0041 | 0.0087 |
| rs2082317 | T | -0.0078 | 0.0014 | 0.0087 | 0.0081 |
| rs2088913 | A | 0.0081 | 0.0014 | -0.0044 | 0.0079 |
| rs2092248 | T | -0.0083 | 0.0015 | 0.0010 | 0.0087 |
| rs2098526 | A | -0.0241 | 0.0043 | 0.0110 | 0.0203 |
| rs2100249 | T | -0.0089 | 0.0015 | 0.0034 | 0.0082 |
| rs2126069 | T | 0.0080 | 0.0014 | 0.0061 | 0.0082 |
| rs2131167 | A | 0.0080 | 0.0014 | -0.0025 | 0.0081 |
| rs214626 | A | 0.0101 | 0.0018 | 0.0028 | 0.0096 |
| rs2160514 | A | -0.0094 | 0.0014 | 0.0027 | 0.0081 |
| rs2179152 | T | -0.0131 | 0.0015 | 0.0039 | 0.0087 |
| rs2183271 | T | 0.0084 | 0.0015 | 0.0144 | 0.0088 |
| rs2199409 | T | 0.0103 | 0.0017 | 0.0129 | 0.0100 |
| rs2212430 | T | -0.0100 | 0.0016 | 0.0003 | 0.0087 |
| rs2216144 | T | 0.0098 | 0.0014 | -0.0103 | 0.0078 |
| rs2220926 | T | -0.0095 | 0.0014 | 0.0062 | 0.0078 |
| rs2250660 | C | 0.0080 | 0.0014 | -0.0123 | 0.0081 |
| rs2252098 | T | -0.0077 | 0.0014 | 0.0003 | 0.0086 |
| rs2254681 | A | 0.0101 | 0.0016 | 0.0054 | 0.0094 |
| rs2256965 | A | 0.0106 | 0.0014 | -0.0060 | 0.0084 |
| rs2276209 | A | 0.0085 | 0.0015 | 0.0159 | 0.0083 |
| rs2283076 | A | 0.0115 | 0.0017 | 0.0079 | 0.0103 |
| rs2287838 | A | -0.0104 | 0.0014 | -0.0024 | 0.0081 |
| rs2290601 | T | -0.0149 | 0.0017 | 0.0103 | 0.0088 |
| rs2297293 | C | 0.0099 | 0.0015 | -0.0006 | 0.0088 |
| rs2299156 | T | 0.0099 | 0.0018 | 0.0041 | 0.0098 |
| rs2314338 | T | 0.0093 | 0.0016 | -0.0063 | 0.0093 |
| rs2321157 | A | -0.0081 | 0.0014 | 0.0087 | 0.0079 |
| rs232496 | T | 0.0090 | 0.0015 | -0.0015 | 0.0082 |
| rs2332179 | A | 0.0121 | 0.0020 | 0.0105 | 0.0117 |
| rs2336721 | T | 0.0095 | 0.0015 | -0.0040 | 0.0088 |
| rs2358628 | A | -0.0091 | 0.0015 | 0.0135 | 0.0085 |
| rs236318 | C | -0.0102 | 0.0017 | -0.0082 | 0.0092 |
| rs2364544 | A | -0.0105 | 0.0014 | -0.0040 | 0.0083 |
| rs2365376 | A | 0.0092 | 0.0015 | 0.0065 | 0.0084 |
| rs2368831 | T | -0.0092 | 0.0014 | -0.0026 | 0.0096 |
| rs2373124 | A | -0.0096 | 0.0016 | 0.0138 | 0.0096 |
| rs2406253 | A | 0.0135 | 0.0018 | -0.0115 | 0.0108 |
| rs2414072 | A | -0.0083 | 0.0014 | 0.0164 | 0.0078 |
| rs2416214 | A | 0.0079 | 0.0014 | 0.0030 | 0.0082 |
| rs2416759 | A | -0.0093 | 0.0015 | -0.0134 | 0.0085 |
| rs2416845 | T | 0.0263 | 0.0048 | -0.0177 | 0.0247 |
| rs2431023 | A | 0.0093 | 0.0014 | -0.0121 | 0.0080 |
| rs2434672 | A | 0.0091 | 0.0014 | 0.0040 | 0.0082 |
| rs2436760 | T | 0.0232 | 0.0042 | -0.0124 | 0.0186 |
| rs2447097 | T | 0.0096 | 0.0014 | -0.0011 | 0.0080 |
| rs2458370 | T | -0.0086 | 0.0015 | 0.0136 | 0.0080 |
| rs2469226 | A | 0.0096 | 0.0017 | -0.0061 | 0.0100 |
| rs2470966 | C | -0.0103 | 0.0015 | 0.0157 | 0.0098 |
| rs2478208 | C | -0.0103 | 0.0014 | -0.0018 | 0.0079 |
| rs2496482 | T | 0.0121 | 0.0015 | -0.0137 | 0.0092 |
| rs2517086 | C | -0.0088 | 0.0014 | 0.0011 | 0.0085 |
| rs2521602 | A | -0.0266 | 0.0049 | -0.0114 | 0.0218 |
| rs2522545 | T | -0.0083 | 0.0014 | -0.0002 | 0.0081 |
| rs2526398 | C | -0.0222 | 0.0015 | 0.0172 | 0.0090 |
| rs2529069 | T | -0.0106 | 0.0016 | 0.0111 | 0.0087 |
| rs252991 | A | 0.0095 | 0.0015 | -0.0068 | 0.0081 |
| rs2542673 | A | 0.0088 | 0.0015 | 0.0083 | 0.0083 |
| rs2545795 | A | 0.0125 | 0.0014 | 0.0004 | 0.0082 |
| rs255053 | A | -0.0111 | 0.0018 | -0.0069 | 0.0098 |
| rs2554835 | A | 0.0088 | 0.0014 | -0.0080 | 0.0085 |
| rs2568955 | T | -0.0165 | 0.0016 | 0.0067 | 0.0098 |
| rs2569041 | T | 0.0077 | 0.0014 | -0.0009 | 0.0085 |
| rs2570497 | T | -0.0121 | 0.0015 | 0.0102 | 0.0081 |
| rs2588959 | T | -0.0080 | 0.0014 | 0.0066 | 0.0078 |
| rs2657283 | T | 0.0085 | 0.0014 | -0.0067 | 0.0085 |
| rs2665668 | A | -0.0094 | 0.0015 | -0.0046 | 0.0087 |
| rs2702576 | A | -0.0090 | 0.0014 | 0.0172 | 0.0086 |
| rs2718277 | T | 0.0142 | 0.0026 | -0.0030 | 0.0147 |
| rs2718791 | T | -0.0081 | 0.0015 | -0.0097 | 0.0087 |
| rs27220 | A | -0.0114 | 0.0015 | -0.0080 | 0.0082 |
| rs2725370 | T | -0.0141 | 0.0015 | -0.0069 | 0.0092 |
| rs2736752 | T | 0.0095 | 0.0017 | -0.0125 | 0.0110 |
| rs2740795 | A | 0.0098 | 0.0016 | 0.0183 | 0.0089 |
| rs2761438 | A | 0.0098 | 0.0015 | -0.0082 | 0.0083 |
| rs2764684 | T | 0.0143 | 0.0019 | 0.0105 | 0.0118 |
| rs2805064 | C | 0.0093 | 0.0016 | -0.0084 | 0.0095 |
| rs2838006 | T | 0.0132 | 0.0015 | -0.0087 | 0.0084 |
| rs28482086 | A | 0.0086 | 0.0015 | -0.0004 | 0.0086 |
| rs28505285 | C | 0.0106 | 0.0018 | 0.0071 | 0.0115 |
| rs28512462 | C | 0.0112 | 0.0014 | 0.0023 | 0.0082 |
| rs28513882 | A | -0.0110 | 0.0018 | -0.0004 | 0.0116 |
| rs2852349 | T | -0.0093 | 0.0014 | -0.0009 | 0.0081 |
| rs28587776 | T | 0.0088 | 0.0015 | -0.0097 | 0.0086 |
| rs28669886 | A | -0.0084 | 0.0015 | 0.0132 | 0.0086 |
| rs2870281 | A | -0.0083 | 0.0014 | 0.0064 | 0.0080 |
| rs28735993 | A | 0.0162 | 0.0022 | 0.0107 | 0.0127 |
| rs2885198 | A | 0.0088 | 0.0014 | -0.0138 | 0.0082 |
| rs2898191 | A | 0.0093 | 0.0015 | -0.0002 | 0.0088 |
| rs2910823 | T | 0.0115 | 0.0014 | 0.0121 | 0.0085 |
| rs2916490 | A | -0.0092 | 0.0015 | 0.0087 | 0.0090 |
| rs2923424 | A | 0.0120 | 0.0014 | -0.0013 | 0.0080 |
| rs2926702 | T | -0.0124 | 0.0021 | 0.0148 | 0.0108 |
| rs2929032 | A | -0.0081 | 0.0014 | -0.0042 | 0.0078 |
| rs2929860 | T | 0.0110 | 0.0018 | 0.0001 | 0.0115 |
| rs2942884 | A | -0.0090 | 0.0014 | -0.0020 | 0.0079 |
| rs2954114 | A | -0.0081 | 0.0014 | 0.0004 | 0.0081 |
| rs2958182 | A | 0.0082 | 0.0015 | 0.0016 | 0.0087 |
| rs2962378 | A | 0.0101 | 0.0017 | -0.0014 | 0.0092 |
| rs2964199 | T | -0.0099 | 0.0015 | 0.0061 | 0.0085 |
| rs2964255 | A | 0.0088 | 0.0015 | -0.0161 | 0.0085 |
| rs2976397 | T | 0.0081 | 0.0014 | 0.0109 | 0.0084 |
| rs2977464 | T | 0.0102 | 0.0018 | 0.0016 | 0.0095 |
| rs29792 | A | -0.0093 | 0.0015 | -0.0257 | 0.0090 |
| rs2980813 | A | 0.0088 | 0.0014 | -0.0104 | 0.0081 |
| rs2989476 | C | 0.0078 | 0.0014 | -0.0188 | 0.0080 |
| rs2989751 | A | -0.0092 | 0.0017 | 0.0014 | 0.0089 |
| rs2992037 | A | 0.0149 | 0.0015 | -0.0038 | 0.0091 |
| rs2994326 | T | -0.0100 | 0.0018 | 0.0013 | 0.0108 |
| rs2998299 | T | -0.0136 | 0.0017 | -0.0022 | 0.0096 |
| rs3026996 | A | 0.0138 | 0.0016 | 0.0031 | 0.0106 |
| rs303752 | A | -0.0110 | 0.0014 | 0.0291 | 0.0089 |
| rs3111251 | T | -0.0090 | 0.0014 | -0.0021 | 0.0080 |
| rs312927 | A | -0.0147 | 0.0022 | 0.0194 | 0.0136 |
| rs312945 | A | -0.0091 | 0.0015 | -0.0081 | 0.0087 |
| rs317050 | T | 0.0078 | 0.0014 | -0.0164 | 0.0079 |
| rs322614 | A | -0.0087 | 0.0014 | 0.0066 | 0.0080 |
| rs322744 | T | -0.0090 | 0.0016 | -0.0001 | 0.0087 |
| rs324885 | A | 0.0142 | 0.0014 | -0.0079 | 0.0081 |
| rs32940 | T | -0.0096 | 0.0015 | -0.0022 | 0.0088 |
| rs339057 | A | -0.0081 | 0.0014 | 0.0111 | 0.0081 |
| rs34067381 | T | -0.0104 | 0.0015 | 0.0310 | 0.0090 |
| rs34098770 | A | -0.0160 | 0.0020 | 0.0024 | 0.0106 |
| rs34106693 | C | 0.0159 | 0.0019 | -0.0124 | 0.0122 |
| rs34122915 | C | 0.0084 | 0.0014 | -0.0034 | 0.0093 |
| rs341504 | A | 0.0099 | 0.0015 | -0.0116 | 0.0082 |
| rs34155847 | A | 0.0135 | 0.0016 | -0.0091 | 0.0094 |
| rs34262657 | T | -0.0240 | 0.0038 | -0.0276 | 0.0255 |
| rs34286836 | T | -0.0089 | 0.0014 | -0.0136 | 0.0080 |
| rs34298584 | A | 0.0111 | 0.0019 | -0.0003 | 0.0115 |
| rs34305371 | A | 0.0314 | 0.0024 | -0.0241 | 0.0164 |
| rs34309 | A | 0.0080 | 0.0014 | -0.0022 | 0.0083 |
| rs34316274 | A | 0.0097 | 0.0017 | -0.0052 | 0.0091 |
| rs34363861 | A | 0.0087 | 0.0014 | -0.0031 | 0.0083 |
| rs34394051 | A | -0.0117 | 0.0020 | 0.0113 | 0.0103 |
| rs34410 | C | -0.0083 | 0.0014 | 0.0077 | 0.0080 |
| rs34624793 | T | 0.0116 | 0.0021 | -0.0145 | 0.0144 |
| rs34720381 | T | -0.0180 | 0.0024 | 0.0377 | 0.0141 |
| rs34748029 | A | 0.0185 | 0.0025 | 0.0138 | 0.0176 |
| rs347661 | T | -0.0081 | 0.0014 | 0.0017 | 0.0084 |
| rs34780702 | A | 0.0111 | 0.0017 | -0.0002 | 0.0092 |
| rs34807077 | A | 0.0128 | 0.0019 | 0.0018 | 0.0099 |
| rs34811474 | A | 0.0104 | 0.0017 | -0.0161 | 0.0128 |
| rs34967082 | A | -0.0080 | 0.0014 | -0.0064 | 0.0084 |
| rs35016816 | T | 0.0176 | 0.0023 | 0.0041 | 0.0128 |
| rs350281 | T | -0.0117 | 0.0015 | -0.0102 | 0.0112 |
| rs35104491 | A | 0.0124 | 0.0018 | 0.0105 | 0.0096 |
| rs35111506 | C | -0.0175 | 0.0030 | -0.0021 | 0.0192 |
| rs35192107 | T | -0.0091 | 0.0017 | 0.0225 | 0.0098 |
| rs35319653 | T | 0.0119 | 0.0015 | 0.0028 | 0.0086 |
| rs35417702 | T | -0.0152 | 0.0014 | 0.0030 | 0.0078 |
| rs35493937 | C | 0.0124 | 0.0017 | -0.0016 | 0.0094 |
| rs35606437 | A | 0.0105 | 0.0016 | -0.0011 | 0.0090 |
| rs356999 | A | -0.0101 | 0.0014 | -0.0049 | 0.0085 |
| rs35745455 | A | 0.0080 | 0.0014 | -0.0031 | 0.0084 |
| rs35751693 | T | -0.0246 | 0.0040 | -0.0001 | 0.0308 |
| rs35754740 | T | 0.0083 | 0.0014 | 0.0053 | 0.0080 |
| rs35811586 | T | 0.0217 | 0.0031 | -0.0050 | 0.0225 |
| rs35929923 | A | -0.0114 | 0.0016 | 0.0110 | 0.0097 |
| rs36085856 | C | 0.0201 | 0.0036 | -0.0112 | 0.0183 |
| rs36120534 | T | 0.0100 | 0.0018 | 0.0000 | 0.0114 |
| rs362307 | T | -0.0226 | 0.0027 | 0.0607 | 0.0176 |
| rs363096 | T | -0.0143 | 0.0014 | 0.0140 | 0.0079 |
| rs3735478 | T | 0.0108 | 0.0016 | 0.0053 | 0.0096 |
| rs3751331 | A | -0.0097 | 0.0014 | -0.0109 | 0.0084 |
| rs3766979 | A | 0.0085 | 0.0015 | -0.0051 | 0.0084 |
| rs3768480 | C | 0.0101 | 0.0014 | -0.0093 | 0.0085 |
| rs3781339 | T | -0.0104 | 0.0018 | 0.0254 | 0.0099 |
| rs3790609 | T | 0.0107 | 0.0019 | -0.0051 | 0.0105 |
| rs3809169 | T | 0.0140 | 0.0025 | -0.0153 | 0.0145 |
| rs3809634 | A | -0.0109 | 0.0015 | 0.0030 | 0.0091 |
| rs3812281 | T | 0.0113 | 0.0014 | -0.0035 | 0.0081 |
| rs3817923 | A | -0.0121 | 0.0022 | 0.0149 | 0.0131 |
| rs382196 | T | -0.0082 | 0.0014 | -0.0053 | 0.0083 |
| rs3847228 | T | 0.0083 | 0.0014 | 0.0089 | 0.0086 |
| rs3848715 | A | -0.0080 | 0.0014 | 0.0054 | 0.0084 |
| rs3859523 | T | 0.0124 | 0.0018 | 0.0031 | 0.0108 |
| rs387027 | A | -0.0083 | 0.0015 | 0.0114 | 0.0085 |
| rs3895736 | A | 0.0144 | 0.0019 | -0.0115 | 0.0115 |
| rs3897821 | A | 0.0154 | 0.0015 | 0.0000 | 0.0085 |
| rs3948495 | T | -0.0089 | 0.0014 | 0.0062 | 0.0083 |
| rs399821 | A | -0.0078 | 0.0014 | -0.0046 | 0.0078 |
| rs39998 | A | 0.0090 | 0.0015 | -0.0033 | 0.0094 |
| rs401526 | T | -0.0104 | 0.0014 | 0.0060 | 0.0081 |
| rs401966 | C | -0.0097 | 0.0014 | 0.0076 | 0.0088 |
| rs4127499 | A | 0.0087 | 0.0015 | -0.0037 | 0.0091 |
| rs41282553 | A | 0.0237 | 0.0040 | -0.0155 | 0.0256 |
| rs42210 | C | -0.0098 | 0.0016 | -0.0039 | 0.0088 |
| rs42302 | A | 0.0086 | 0.0015 | -0.0085 | 0.0084 |
| rs4255791 | A | -0.0084 | 0.0015 | -0.0014 | 0.0092 |
| rs4263475 | A | 0.0078 | 0.0014 | -0.0089 | 0.0086 |
| rs4283754 | A | -0.0084 | 0.0014 | -0.0006 | 0.0098 |
| rs429150 | T | 0.0085 | 0.0014 | -0.0005 | 0.0085 |
| rs4298514 | T | 0.0104 | 0.0015 | -0.0108 | 0.0084 |
| rs4320563 | A | 0.0109 | 0.0014 | -0.0158 | 0.0078 |
| rs4328757 | T | 0.0096 | 0.0014 | 0.0037 | 0.0082 |
| rs4352658 | T | -0.0190 | 0.0026 | -0.0091 | 0.0141 |
| rs4358081 | A | -0.0095 | 0.0014 | -0.0047 | 0.0082 |
| rs4358358 | A | 0.0109 | 0.0015 | -0.0048 | 0.0090 |
| rs4369924 | A | 0.0125 | 0.0019 | -0.0239 | 0.0102 |
| rs4384309 | A | 0.0102 | 0.0014 | -0.0084 | 0.0083 |
| rs4396896 | A | -0.0107 | 0.0014 | 0.0019 | 0.0079 |
| rs4423373 | A | -0.0087 | 0.0014 | 0.0037 | 0.0086 |
| rs4426420 | T | -0.0140 | 0.0019 | 0.0005 | 0.0111 |
| rs4434676 | A | -0.0089 | 0.0014 | 0.0246 | 0.0085 |
| rs4458044 | C | 0.0101 | 0.0016 | -0.0178 | 0.0110 |
| rs4467547 | T | 0.0121 | 0.0014 | -0.0187 | 0.0091 |
| rs4469771 | T | 0.0101 | 0.0015 | -0.0082 | 0.0083 |
| rs4480339 | A | -0.0097 | 0.0015 | 0.0017 | 0.0094 |
| rs4490539 | A | 0.0142 | 0.0015 | -0.0021 | 0.0092 |
| rs4497562 | A | 0.0112 | 0.0016 | -0.0125 | 0.0088 |
| rs4500930 | T | -0.0109 | 0.0015 | 0.0114 | 0.0084 |
| rs4500960 | T | -0.0131 | 0.0014 | -0.0180 | 0.0079 |
| rs4502401 | T | 0.0085 | 0.0015 | -0.0027 | 0.0086 |
| rs4652135 | A | 0.0103 | 0.0016 | -0.0046 | 0.0087 |
| rs4658019 | T | 0.0084 | 0.0014 | 0.0021 | 0.0081 |
| rs4663617 | A | 0.0104 | 0.0017 | 0.0024 | 0.0098 |
| rs4664983 | T | -0.0103 | 0.0018 | 0.0037 | 0.0093 |
| rs4673840 | T | -0.0138 | 0.0019 | 0.0008 | 0.0105 |
| rs4675248 | A | -0.0094 | 0.0014 | -0.0031 | 0.0082 |
| rs4685405 | T | -0.0111 | 0.0018 | 0.0010 | 0.0098 |
| rs4687735 | T | -0.0200 | 0.0033 | 0.0532 | 0.0226 |
| rs4691601 | A | -0.0115 | 0.0014 | 0.0000 | 0.0080 |
| rs4705763 | C | 0.0089 | 0.0014 | 0.0011 | 0.0079 |
| rs4712371 | A | -0.0124 | 0.0021 | 0.0158 | 0.0116 |
| rs4719460 | T | -0.0088 | 0.0015 | 0.0167 | 0.0083 |
| rs4719944 | T | -0.0121 | 0.0014 | -0.0011 | 0.0079 |
| rs4724083 | T | 0.0083 | 0.0015 | 0.0015 | 0.0087 |
| rs4726070 | A | 0.0122 | 0.0014 | 0.0011 | 0.0081 |
| rs4730020 | T | 0.0090 | 0.0016 | -0.0127 | 0.0101 |
| rs4731413 | A | 0.0111 | 0.0017 | -0.0090 | 0.0111 |
| rs4735297 | A | -0.0086 | 0.0015 | -0.0011 | 0.0082 |
| rs4739235 | A | 0.0109 | 0.0019 | -0.0206 | 0.0101 |
| rs4741571 | A | -0.0102 | 0.0015 | 0.0016 | 0.0093 |
| rs4757957 | C | 0.0133 | 0.0015 | 0.0013 | 0.0087 |
| rs4766424 | C | 0.0142 | 0.0021 | -0.0109 | 0.0135 |
| rs4778058 | T | -0.0092 | 0.0014 | 0.0051 | 0.0082 |
| rs4785187 | A | -0.0100 | 0.0017 | 0.0033 | 0.0092 |
| rs4787028 | T | 0.0086 | 0.0015 | -0.0031 | 0.0083 |
| rs4787457 | A | 0.0162 | 0.0015 | 0.0053 | 0.0085 |
| rs4788115 | A | 0.0111 | 0.0019 | -0.0144 | 0.0114 |
| rs479018 | A | 0.0109 | 0.0015 | 0.0024 | 0.0099 |
| rs4793090 | A | 0.0084 | 0.0015 | 0.0039 | 0.0081 |
| rs4810894 | A | -0.0085 | 0.0015 | -0.0065 | 0.0082 |
| rs4812697 | T | 0.0171 | 0.0027 | -0.0353 | 0.0179 |
| rs481940 | T | 0.0100 | 0.0016 | 0.0112 | 0.0094 |
| rs482787 | T | 0.0100 | 0.0015 | 0.0108 | 0.0083 |
| rs4839155 | T | 0.0115 | 0.0017 | 0.0133 | 0.0096 |
| rs4846010 | A | 0.0099 | 0.0018 | -0.0147 | 0.0121 |
| rs4846724 | A | 0.0098 | 0.0014 | -0.0013 | 0.0079 |
| rs4848924 | A | -0.0117 | 0.0015 | 0.0084 | 0.0085 |
| rs4850954 | T | 0.0077 | 0.0014 | -0.0005 | 0.0082 |
| rs4851263 | A | -0.0151 | 0.0023 | 0.0077 | 0.0119 |
| rs4877516 | A | -0.0101 | 0.0014 | 0.0102 | 0.0079 |
| rs4881269 | A | 0.0100 | 0.0014 | -0.0063 | 0.0081 |
| rs488476 | C | -0.0113 | 0.0015 | 0.0065 | 0.0085 |
| rs4894658 | C | 0.0091 | 0.0016 | 0.0032 | 0.0088 |
| rs4895650 | T | 0.0093 | 0.0014 | 0.0083 | 0.0088 |
| rs4899012 | C | -0.0103 | 0.0014 | 0.0087 | 0.0084 |
| rs4904523 | A | -0.0082 | 0.0014 | 0.0082 | 0.0079 |
| rs4915735 | A | 0.0116 | 0.0020 | -0.0047 | 0.0120 |
| rs4919624 | A | 0.0193 | 0.0018 | -0.0043 | 0.0100 |
| rs4925065 | T | -0.0078 | 0.0014 | 0.0014 | 0.0081 |
| rs4925109 | A | -0.0096 | 0.0015 | 0.0213 | 0.0086 |
| rs4938815 | T | 0.0085 | 0.0015 | 0.0061 | 0.0088 |
| rs4941735 | T | 0.0092 | 0.0014 | 0.0052 | 0.0083 |
| rs4972748 | T | 0.0105 | 0.0018 | -0.0115 | 0.0096 |
| rs4977885 | A | -0.0084 | 0.0014 | 0.0134 | 0.0085 |
| rs4981245 | A | 0.0080 | 0.0014 | -0.0045 | 0.0085 |
| rs4984541 | A | -0.0138 | 0.0017 | -0.0005 | 0.0093 |
| rs4984613 | T | 0.0174 | 0.0028 | -0.0208 | 0.0147 |
| rs4984682 | C | -0.0138 | 0.0017 | -0.0034 | 0.0091 |
| rs548897 | A | 0.0081 | 0.0014 | -0.0100 | 0.0083 |
| rs55675587 | T | -0.0103 | 0.0018 | 0.0071 | 0.0098 |
| rs55736314 | C | -0.0154 | 0.0014 | 0.0183 | 0.0090 |
| rs55826493 | T | -0.0127 | 0.0021 | -0.0179 | 0.0135 |
| rs55986781 | T | 0.0187 | 0.0026 | -0.0291 | 0.0166 |
| rs56085180 | A | -0.0269 | 0.0037 | -0.0020 | 0.0236 |
| rs56099375 | T | 0.0120 | 0.0017 | 0.0042 | 0.0108 |
| rs56171318 | T | -0.0137 | 0.0020 | -0.0105 | 0.0121 |
| rs56174996 | A | -0.0143 | 0.0020 | 0.0001 | 0.0131 |
| rs56306882 | A | 0.0086 | 0.0016 | 0.0065 | 0.0097 |
| rs56319902 | T | -0.0206 | 0.0017 | 0.0211 | 0.0125 |
| rs56391344 | A | 0.0148 | 0.0016 | -0.0135 | 0.0103 |
| rs563954 | A | -0.0081 | 0.0014 | 0.0097 | 0.0086 |
| rs56405138 | A | 0.0153 | 0.0022 | -0.0070 | 0.0127 |
| rs56408528 | T | -0.0082 | 0.0014 | 0.0095 | 0.0080 |
| rs567003 | A | 0.0094 | 0.0015 | -0.0048 | 0.0090 |
| rs56794817 | A | 0.0122 | 0.0019 | -0.0089 | 0.0101 |
| rs57016874 | T | 0.0209 | 0.0037 | -0.0085 | 0.0211 |
| rs57148205 | A | 0.0095 | 0.0016 | -0.0160 | 0.0097 |
| rs57204268 | A | 0.0114 | 0.0020 | -0.0010 | 0.0125 |
| rs57349798 | A | 0.0093 | 0.0014 | -0.0171 | 0.0087 |
| rs57352738 | A | -0.0152 | 0.0017 | -0.0072 | 0.0103 |
| rs57437407 | A | -0.0145 | 0.0023 | -0.0229 | 0.0137 |
| rs575113 | A | 0.0133 | 0.0015 | -0.0151 | 0.0087 |
| rs5754581 | T | -0.0082 | 0.0014 | -0.0041 | 0.0080 |
| rs5754753 | T | -0.0126 | 0.0016 | -0.0033 | 0.0099 |
| rs5754762 | A | 0.0197 | 0.0030 | -0.0291 | 0.0139 |
| rs57661533 | T | -0.0114 | 0.0021 | -0.0002 | 0.0103 |
| rs580652 | T | -0.0147 | 0.0024 | 0.0080 | 0.0111 |
| rs585557 | A | 0.0128 | 0.0018 | -0.0294 | 0.0095 |
| rs58779949 | A | 0.0110 | 0.0019 | -0.0208 | 0.0112 |
| rs58859557 | T | 0.0187 | 0.0028 | -0.0561 | 0.0150 |
| rs58921703 | T | 0.0109 | 0.0015 | -0.0155 | 0.0083 |
| rs58950082 | T | 0.0095 | 0.0017 | -0.0093 | 0.0094 |
| rs58996896 | A | -0.0094 | 0.0016 | 0.0059 | 0.0094 |
| rs590013 | T | 0.0100 | 0.0015 | -0.0085 | 0.0086 |
| rs59300999 | T | -0.0164 | 0.0028 | 0.0065 | 0.0121 |
| rs59480703 | C | -0.0123 | 0.0018 | -0.0160 | 0.0096 |
| rs59813324 | T | 0.0097 | 0.0018 | -0.0060 | 0.0095 |
| rs59967356 | T | -0.0127 | 0.0021 | -0.0050 | 0.0144 |
| rs60096640 | A | 0.0159 | 0.0023 | -0.0006 | 0.0108 |
| rs6020560 | T | -0.0083 | 0.0014 | -0.0078 | 0.0080 |
| rs6043521 | T | -0.0083 | 0.0014 | -0.0057 | 0.0084 |
| rs60589532 | A | 0.0168 | 0.0028 | 0.0086 | 0.0152 |
| rs6060308 | A | 0.0095 | 0.0016 | -0.0208 | 0.0092 |
| rs6065080 | T | -0.0130 | 0.0015 | 0.0001 | 0.0082 |
| rs6065784 | C | 0.0110 | 0.0015 | -0.0042 | 0.0092 |
| rs60717745 | C | -0.0118 | 0.0019 | -0.0092 | 0.0105 |
| rs60726488 | T | -0.0090 | 0.0016 | 0.0021 | 0.0093 |
| rs6091570 | A | 0.0081 | 0.0015 | -0.0106 | 0.0082 |
| rs61104616 | A | -0.0172 | 0.0014 | 0.0047 | 0.0080 |
| rs6123924 | A | 0.0129 | 0.0019 | -0.0163 | 0.0102 |
| rs613872 | T | -0.0163 | 0.0019 | 0.0168 | 0.0121 |
| rs61387839 | A | 0.0088 | 0.0015 | -0.0028 | 0.0095 |
| rs61527214 | A | 0.0110 | 0.0014 | 0.0000 | 0.0085 |
| rs61739710 | A | -0.0099 | 0.0016 | 0.0027 | 0.0091 |
| rs61748951 | A | -0.0273 | 0.0048 | 0.0547 | 0.0362 |
| rs61755388 | T | 0.0125 | 0.0017 | 0.0202 | 0.0107 |
| rs61798586 | A | -0.0112 | 0.0020 | -0.0017 | 0.0115 |
| rs61853335 | T | -0.0109 | 0.0018 | 0.0190 | 0.0100 |
| rs61958175 | A | -0.0198 | 0.0034 | 0.0378 | 0.0251 |
| rs61997667 | T | -0.0144 | 0.0020 | 0.0071 | 0.0118 |
| rs62007304 | A | 0.0264 | 0.0047 | -0.0032 | 0.0368 |
| rs62018216 | T | 0.0113 | 0.0019 | -0.0066 | 0.0121 |
| rs62051146 | A | -0.0134 | 0.0022 | 0.0071 | 0.0122 |
| rs62090515 | A | -0.0090 | 0.0015 | 0.0231 | 0.0085 |
| rs62092949 | T | -0.0087 | 0.0014 | -0.0033 | 0.0081 |
| rs62103198 | T | -0.0147 | 0.0025 | -0.0212 | 0.0186 |
| rs62109862 | A | 0.0136 | 0.0020 | -0.0236 | 0.0110 |
| rs62142891 | A | 0.0108 | 0.0016 | -0.0138 | 0.0100 |
| rs62155350 | A | 0.0338 | 0.0053 | 0.0803 | 0.0407 |
| rs62155770 | A | -0.0196 | 0.0032 | -0.0111 | 0.0213 |
| rs62155873 | T | -0.0151 | 0.0021 | -0.0017 | 0.0136 |
| rs62174974 | A | -0.0105 | 0.0018 | 0.0237 | 0.0095 |
| rs62177359 | A | -0.0275 | 0.0038 | 0.0216 | 0.0291 |
| rs62179650 | A | 0.0112 | 0.0016 | 0.0109 | 0.0094 |
| rs62182994 | T | 0.0137 | 0.0015 | -0.0092 | 0.0097 |
| rs62190914 | T | 0.0089 | 0.0015 | 0.0091 | 0.0085 |
| rs62194170 | A | 0.0111 | 0.0017 | 0.0000 | 0.0095 |
| rs622169 | T | 0.0086 | 0.0014 | -0.0158 | 0.0097 |
| rs62247449 | C | 0.0114 | 0.0014 | 0.0045 | 0.0078 |
| rs62252819 | A | -0.0094 | 0.0017 | -0.0043 | 0.0103 |
| rs62256284 | T | 0.0111 | 0.0017 | 0.0001 | 0.0098 |
| rs62260764 | C | -0.0132 | 0.0016 | -0.0054 | 0.0102 |
| rs62262671 | A | 0.0213 | 0.0020 | 0.0020 | 0.0138 |
| rs62340636 | T | 0.0103 | 0.0014 | -0.0105 | 0.0080 |
| rs62370510 | A | -0.0109 | 0.0019 | 0.0277 | 0.0115 |
| rs62379838 | T | 0.0121 | 0.0015 | -0.0005 | 0.0095 |
| rs62409395 | T | 0.0101 | 0.0018 | -0.0072 | 0.0137 |
| rs62420387 | T | -0.0184 | 0.0027 | 0.0002 | 0.0174 |
| rs62506074 | T | 0.0091 | 0.0015 | -0.0024 | 0.0091 |
| rs62506104 | A | -0.0098 | 0.0018 | 0.0008 | 0.0105 |
| rs628993 | A | 0.0128 | 0.0023 | 0.0276 | 0.0122 |
| rs631287 | A | 0.0084 | 0.0014 | 0.0045 | 0.0078 |
| rs633279 | A | -0.0089 | 0.0015 | 0.0134 | 0.0086 |
| rs635754 | A | 0.0119 | 0.0014 | -0.0044 | 0.0078 |
| rs6428587 | T | 0.0084 | 0.0015 | -0.0027 | 0.0081 |
| rs6435326 | A | -0.0091 | 0.0014 | -0.0170 | 0.0081 |
| rs6442126 | A | -0.0121 | 0.0015 | 0.0090 | 0.0093 |
| rs6445633 | A | 0.0088 | 0.0015 | -0.0021 | 0.0096 |
| rs6449503 | A | 0.0185 | 0.0014 | -0.0038 | 0.0088 |
| rs6450476 | A | -0.0116 | 0.0015 | 0.0162 | 0.0089 |
| rs6452793 | T | 0.0175 | 0.0017 | -0.0147 | 0.0100 |
| rs6461536 | A | 0.0088 | 0.0015 | -0.0169 | 0.0087 |
| rs6469654 | C | -0.0103 | 0.0017 | 0.0051 | 0.0091 |
| rs6472208 | T | -0.0091 | 0.0014 | 0.0164 | 0.0087 |
| rs6480234 | T | -0.0084 | 0.0014 | 0.0182 | 0.0081 |
| rs648044 | A | 0.0083 | 0.0015 | 0.0043 | 0.0089 |
| rs6490618 | T | 0.0088 | 0.0015 | -0.0096 | 0.0083 |
| rs6493265 | T | -0.0122 | 0.0014 | 0.0179 | 0.0085 |
| rs6493275 | A | 0.0131 | 0.0017 | 0.0070 | 0.0102 |
| rs6503409 | T | 0.0087 | 0.0015 | -0.0035 | 0.0089 |
| rs6534338 | T | 0.0103 | 0.0015 | -0.0051 | 0.0086 |
| rs6543810 | T | 0.0082 | 0.0015 | -0.0012 | 0.0085 |
| rs6557171 | T | -0.0155 | 0.0015 | 0.0040 | 0.0083 |
| rs6567288 | A | -0.0079 | 0.0014 | 0.0277 | 0.0081 |
| rs6569077 | T | 0.0118 | 0.0014 | -0.0049 | 0.0082 |
| rs6573552 | T | -0.0090 | 0.0014 | 0.0079 | 0.0080 |
| rs6573559 | T | 0.0105 | 0.0015 | 0.0055 | 0.0083 |
| rs660001 | A | -0.0164 | 0.0017 | 0.0200 | 0.0096 |
| rs663251 | T | 0.0081 | 0.0015 | -0.0070 | 0.0080 |
| rs66482320 | C | 0.0124 | 0.0022 | -0.0113 | 0.0121 |
| rs66568921 | T | -0.0181 | 0.0015 | 0.0109 | 0.0082 |
| rs66671632 | T | -0.0133 | 0.0021 | -0.0137 | 0.0116 |
| rs66721975 | A | -0.0087 | 0.0015 | 0.0036 | 0.0091 |
| rs6672986 | C | 0.0094 | 0.0014 | -0.0081 | 0.0080 |
| rs6689641 | A | 0.0078 | 0.0014 | -0.0034 | 0.0081 |
| rs6690195 | T | -0.0113 | 0.0014 | -0.0080 | 0.0087 |
| rs6695132 | T | 0.0097 | 0.0017 | -0.0019 | 0.0092 |
| rs6696068 | T | -0.0088 | 0.0014 | 0.0047 | 0.0081 |
| rs6697584 | T | -0.0126 | 0.0017 | -0.0001 | 0.0100 |
| rs6704768 | A | -0.0116 | 0.0014 | 0.0099 | 0.0077 |
| rs6706275 | T | 0.0083 | 0.0015 | -0.0058 | 0.0094 |
| rs6711399 | T | -0.0107 | 0.0019 | -0.0083 | 0.0095 |
| rs6715321 | T | -0.0097 | 0.0014 | 0.0156 | 0.0082 |
| rs6715849 | A | -0.0122 | 0.0014 | 0.0114 | 0.0079 |
| rs6720515 | A | 0.0103 | 0.0017 | -0.0004 | 0.0090 |
| rs67224963 | A | 0.0113 | 0.0017 | -0.0214 | 0.0095 |
| rs6729612 | T | 0.0092 | 0.0015 | 0.0149 | 0.0086 |
| rs6731373 | A | -0.0115 | 0.0015 | -0.0018 | 0.0091 |
| rs6736025 | T | 0.0079 | 0.0014 | 0.0037 | 0.0079 |
| rs6738860 | A | -0.0088 | 0.0014 | -0.0145 | 0.0085 |
| rs6743032 | A | -0.0165 | 0.0024 | -0.0119 | 0.0144 |
| rs6744040 | A | -0.0080 | 0.0014 | -0.0155 | 0.0079 |
| rs67456868 | A | 0.0119 | 0.0018 | -0.0184 | 0.0112 |
| rs6752228 | T | -0.0080 | 0.0014 | -0.0037 | 0.0079 |
| rs6757087 | T | -0.0085 | 0.0014 | -0.0008 | 0.0084 |
| rs6760887 | T | 0.0155 | 0.0022 | 0.0005 | 0.0102 |
| rs67661275 | A | 0.0134 | 0.0025 | -0.0320 | 0.0173 |
| rs6774533 | T | 0.0090 | 0.0015 | -0.0018 | 0.0086 |
| rs67790232 | A | 0.0101 | 0.0017 | -0.0088 | 0.0101 |
| rs6780414 | T | 0.0105 | 0.0018 | -0.0043 | 0.0102 |
| rs6792805 | A | -0.0090 | 0.0016 | 0.0043 | 0.0088 |
| rs6812533 | T | 0.0106 | 0.0016 | 0.0112 | 0.0090 |
| rs68145588 | T | -0.0127 | 0.0021 | -0.0146 | 0.0120 |
| rs6824567 | T | 0.0130 | 0.0016 | -0.0063 | 0.0095 |
| rs6853599 | T | 0.0108 | 0.0019 | -0.0346 | 0.0120 |
| rs6861925 | C | -0.0090 | 0.0014 | 0.0043 | 0.0079 |
| rs6871635 | A | -0.0087 | 0.0014 | 0.0073 | 0.0081 |
| rs6881581 | A | -0.0096 | 0.0015 | 0.0094 | 0.0086 |
| rs6881733 | T | -0.0086 | 0.0016 | 0.0073 | 0.0084 |
| rs6917154 | T | 0.0127 | 0.0021 | -0.0031 | 0.0140 |
| rs6917204 | T | 0.0119 | 0.0017 | -0.0125 | 0.0092 |
| rs6918506 | C | 0.0113 | 0.0014 | 0.0057 | 0.0080 |
| rs6924023 | A | 0.0107 | 0.0016 | -0.0018 | 0.0093 |
| rs6932108 | C | -0.0210 | 0.0027 | 0.0220 | 0.0184 |
| rs6946136 | A | 0.0081 | 0.0014 | -0.0047 | 0.0084 |
| rs6946362 | T | 0.0091 | 0.0015 | -0.0004 | 0.0090 |
| rs6951996 | A | -0.0182 | 0.0032 | -0.0013 | 0.0177 |
| rs6956283 | T | 0.0098 | 0.0016 | 0.0090 | 0.0092 |
| rs6969783 | A | -0.0098 | 0.0014 | 0.0093 | 0.0079 |
| rs6977237 | T | 0.0082 | 0.0014 | 0.0139 | 0.0079 |
| rs6989141 | A | 0.0100 | 0.0017 | -0.0375 | 0.0109 |
| rs6994287 | A | 0.0102 | 0.0014 | 0.0002 | 0.0080 |
| rs7012546 | T | 0.0092 | 0.0014 | -0.0114 | 0.0082 |
| rs7016302 | C | -0.0125 | 0.0019 | -0.0178 | 0.0113 |
| rs702606 | T | 0.0118 | 0.0021 | 0.0122 | 0.0118 |
| rs7029718 | A | 0.0244 | 0.0014 | -0.0136 | 0.0081 |
| rs7030373 | A | -0.0098 | 0.0017 | 0.0089 | 0.0089 |
| rs7035315 | A | -0.0090 | 0.0015 | -0.0032 | 0.0081 |
| rs7040995 | C | 0.0097 | 0.0014 | 0.0117 | 0.0079 |
| rs7041702 | A | -0.0111 | 0.0016 | 0.0028 | 0.0092 |
| rs7084508 | T | -0.0085 | 0.0015 | 0.0005 | 0.0088 |
| rs710629 | A | 0.0082 | 0.0015 | 0.0053 | 0.0084 |
| rs7108020 | A | 0.0111 | 0.0015 | 0.0001 | 0.0090 |
| rs711793 | T | 0.0103 | 0.0015 | 0.0001 | 0.0096 |
| rs7127580 | C | -0.0150 | 0.0023 | 0.0025 | 0.0135 |
| rs713584 | A | -0.0080 | 0.0014 | 0.0006 | 0.0083 |
| rs71413877 | A | 0.0351 | 0.0036 | -0.0397 | 0.0254 |
| rs7146625 | A | 0.0091 | 0.0016 | -0.0046 | 0.0090 |
| rs7147473 | A | -0.0103 | 0.0015 | 0.0047 | 0.0093 |
| rs7158218 | A | -0.0090 | 0.0015 | 0.0207 | 0.0092 |
| rs71646142 | T | 0.0104 | 0.0018 | 0.0087 | 0.0113 |
| rs716513 | A | 0.0084 | 0.0014 | 0.0100 | 0.0082 |
| rs7167688 | T | -0.0094 | 0.0014 | 0.0005 | 0.0081 |
| rs7171405 | A | -0.0101 | 0.0016 | 0.0002 | 0.0092 |
| rs717996 | T | -0.0122 | 0.0014 | -0.0012 | 0.0084 |
| rs7182216 | C | -0.0096 | 0.0017 | -0.0032 | 0.0105 |
| rs7190 | A | -0.0095 | 0.0015 | 0.0132 | 0.0093 |
| rs721579 | T | -0.0110 | 0.0016 | -0.0094 | 0.0089 |
| rs7218235 | A | -0.0097 | 0.0017 | 0.0078 | 0.0105 |
| rs7223311 | C | -0.0079 | 0.0014 | 0.0153 | 0.0080 |
| rs7226824 | T | -0.0078 | 0.0014 | 0.0009 | 0.0079 |
| rs72482130 | T | 0.0141 | 0.0019 | -0.0165 | 0.0113 |
| rs72486027 | T | -0.0104 | 0.0016 | 0.0213 | 0.0096 |
| rs7254263 | T | -0.0119 | 0.0016 | 0.0095 | 0.0089 |
| rs7255223 | A | 0.0097 | 0.0016 | -0.0112 | 0.0089 |
| rs72622559 | T | -0.0120 | 0.0017 | 0.0133 | 0.0093 |
| rs72624911 | T | 0.0282 | 0.0034 | -0.0212 | 0.0259 |
| rs72636697 | T | -0.0140 | 0.0020 | 0.0063 | 0.0112 |
| rs72667460 | T | 0.0197 | 0.0031 | -0.0343 | 0.0208 |
| rs72671456 | A | -0.0174 | 0.0020 | 0.0336 | 0.0127 |
| rs72672052 | A | -0.0101 | 0.0018 | -0.0008 | 0.0102 |
| rs72673097 | A | 0.0257 | 0.0042 | -0.0738 | 0.0324 |
| rs72677177 | A | 0.0100 | 0.0014 | 0.0047 | 0.0086 |
| rs72694479 | T | 0.0127 | 0.0020 | -0.0090 | 0.0127 |
| rs72709560 | A | 0.0085 | 0.0015 | -0.0123 | 0.0092 |
| rs72771860 | T | -0.0211 | 0.0029 | -0.0064 | 0.0218 |
| rs72792395 | T | -0.0145 | 0.0022 | 0.0041 | 0.0135 |
| rs728054 | A | -0.0128 | 0.0015 | 0.0241 | 0.0087 |
| rs72819118 | T | -0.0231 | 0.0024 | 0.0255 | 0.0159 |
| rs72824753 | T | 0.0133 | 0.0024 | -0.0227 | 0.0145 |
| rs72828517 | T | -0.0168 | 0.0019 | 0.0071 | 0.0108 |
| rs72829857 | A | -0.0149 | 0.0016 | -0.0050 | 0.0096 |
| rs72834698 | A | 0.0117 | 0.0020 | -0.0127 | 0.0133 |
| rs72881110 | A | 0.0146 | 0.0027 | -0.0276 | 0.0157 |
| rs72883760 | A | 0.0101 | 0.0018 | 0.0069 | 0.0105 |
| rs72896637 | T | 0.0162 | 0.0028 | -0.0073 | 0.0191 |
| rs72902523 | T | -0.0103 | 0.0016 | -0.0076 | 0.0089 |
| rs72906124 | T | 0.0185 | 0.0030 | -0.0342 | 0.0190 |
| rs72917504 | T | -0.0178 | 0.0030 | 0.0162 | 0.0192 |
| rs72919450 | T | -0.0132 | 0.0023 | 0.0232 | 0.0125 |
| rs72944064 | T | -0.0098 | 0.0016 | 0.0085 | 0.0098 |
| rs72962169 | T | -0.0174 | 0.0019 | 0.0079 | 0.0116 |
| rs72972965 | A | 0.0083 | 0.0015 | 0.0118 | 0.0087 |
| rs72993796 | T | 0.0127 | 0.0022 | -0.0091 | 0.0148 |
| rs73034295 | A | 0.0099 | 0.0018 | -0.0166 | 0.0115 |
| rs730384 | A | 0.0101 | 0.0014 | -0.0032 | 0.0082 |
| rs73039077 | C | 0.0092 | 0.0016 | -0.0033 | 0.0101 |
| rs73055556 | A | 0.0111 | 0.0020 | -0.0400 | 0.0108 |
| rs73082325 | C | -0.0252 | 0.0044 | 0.0017 | 0.0331 |
| rs73106136 | T | -0.0147 | 0.0026 | -0.0138 | 0.0166 |
| rs73154546 | A | 0.0197 | 0.0036 | -0.0312 | 0.0248 |
| rs7317761 | C | 0.0109 | 0.0016 | -0.0012 | 0.0088 |
| rs73191311 | A | -0.0094 | 0.0015 | -0.0017 | 0.0086 |
| rs7321274 | A | 0.0114 | 0.0018 | 0.0024 | 0.0100 |
| rs73219806 | A | 0.0106 | 0.0019 | -0.0222 | 0.0101 |
| rs7323027 | A | 0.0130 | 0.0015 | 0.0027 | 0.0083 |
| rs7326331 | A | -0.0128 | 0.0016 | 0.0228 | 0.0090 |
| rs73344830 | A | 0.0170 | 0.0014 | 0.0025 | 0.0082 |
| rs73405293 | A | 0.0113 | 0.0020 | 0.0180 | 0.0112 |
| rs73457936 | A | 0.0144 | 0.0025 | 0.0085 | 0.0160 |
| rs73518807 | C | -0.0128 | 0.0021 | -0.0068 | 0.0091 |
| rs7356536 | T | -0.0129 | 0.0021 | 0.0142 | 0.0130 |
| rs73581580 | A | -0.0181 | 0.0022 | 0.0271 | 0.0169 |
| rs736281 | T | 0.0086 | 0.0014 | 0.0013 | 0.0084 |
| rs73643713 | T | -0.0165 | 0.0027 | -0.0224 | 0.0144 |
| rs736471 | T | 0.0083 | 0.0014 | 0.0066 | 0.0083 |
| rs73648455 | T | -0.0183 | 0.0026 | 0.0060 | 0.0140 |
| rs737945 | C | -0.0115 | 0.0014 | 0.0137 | 0.0081 |
| rs73874335 | T | -0.0178 | 0.0029 | -0.0042 | 0.0146 |
| rs73961845 | A | -0.0114 | 0.0014 | 0.0045 | 0.0085 |
| rs74091672 | A | -0.0153 | 0.0022 | 0.0014 | 0.0147 |
| rs7430651 | T | -0.0111 | 0.0016 | 0.0159 | 0.0090 |
| rs743316 | T | 0.0112 | 0.0017 | -0.0005 | 0.0109 |
| rs74415461 | T | 0.0176 | 0.0026 | 0.0009 | 0.0168 |
| rs74453875 | A | 0.0214 | 0.0037 | 0.0329 | 0.0197 |
| rs74462621 | T | 0.0125 | 0.0020 | 0.0132 | 0.0127 |
| rs7449561 | A | 0.0102 | 0.0017 | 0.0081 | 0.0093 |
| rs7451726 | A | 0.0109 | 0.0017 | 0.0030 | 0.0093 |
| rs74545339 | A | -0.0158 | 0.0022 | -0.0127 | 0.0132 |
| rs7460106 | T | -0.0123 | 0.0017 | -0.0218 | 0.0098 |
| rs74615093 | A | 0.0145 | 0.0026 | -0.0316 | 0.0151 |
| rs746839 | C | 0.0135 | 0.0015 | -0.0076 | 0.0091 |
| rs74747621 | T | 0.0153 | 0.0022 | -0.0283 | 0.0137 |
| rs74787922 | A | 0.0182 | 0.0028 | 0.0055 | 0.0176 |
| rs74944275 | T | 0.0231 | 0.0035 | 0.0307 | 0.0233 |
| rs74944857 | T | -0.0089 | 0.0015 | -0.0092 | 0.0096 |
| rs7495033 | A | 0.0095 | 0.0015 | -0.0084 | 0.0087 |
| rs75033012 | C | -0.0258 | 0.0038 | -0.0025 | 0.0292 |
| rs75177132 | T | 0.0298 | 0.0036 | -0.0217 | 0.0288 |
| rs75203411 | T | 0.0135 | 0.0019 | -0.0008 | 0.0107 |
| rs7526112 | T | 0.0124 | 0.0015 | 0.0012 | 0.0084 |
| rs75308819 | A | 0.0287 | 0.0047 | 0.0221 | 0.0353 |
| rs75434274 | A | 0.0151 | 0.0026 | -0.0014 | 0.0184 |
| rs75500877 | T | 0.0207 | 0.0033 | -0.0043 | 0.0205 |
| rs7552964 | A | 0.0148 | 0.0023 | -0.0095 | 0.0149 |
| rs7560871 | A | -0.0193 | 0.0027 | 0.0261 | 0.0167 |
| rs7561705 | A | 0.0078 | 0.0014 | 0.0120 | 0.0080 |
| rs7575637 | A | 0.0107 | 0.0014 | -0.0073 | 0.0078 |
| rs75756843 | A | 0.0288 | 0.0049 | 0.0884 | 0.0336 |
| rs7575938 | A | 0.0090 | 0.0015 | -0.0072 | 0.0085 |
| rs7590368 | T | -0.0146 | 0.0016 | 0.0057 | 0.0094 |
| rs7597126 | T | -0.0094 | 0.0014 | 0.0001 | 0.0086 |
| rs7597412 | C | 0.0088 | 0.0015 | -0.0180 | 0.0089 |
| rs7603132 | A | 0.0137 | 0.0018 | 0.0039 | 0.0099 |
| rs76076331 | T | 0.0206 | 0.0021 | -0.0229 | 0.0128 |
| rs76077165 | A | 0.0165 | 0.0025 | -0.0073 | 0.0163 |
| rs76167224 | T | -0.0187 | 0.0033 | 0.0690 | 0.0262 |
| rs7617204 | A | -0.0097 | 0.0014 | 0.0015 | 0.0081 |
| rs76235882 | A | 0.0241 | 0.0039 | -0.0104 | 0.0253 |
| rs76241605 | A | 0.0139 | 0.0022 | 0.0064 | 0.0151 |
| rs76246107 | A | -0.0162 | 0.0026 | 0.0060 | 0.0197 |
| rs7625428 | T | 0.0098 | 0.0014 | 0.0095 | 0.0084 |
| rs76267866 | A | 0.0104 | 0.0017 | 0.0027 | 0.0106 |
| rs7630133 | A | 0.0082 | 0.0015 | -0.0029 | 0.0088 |
| rs763553 | A | 0.0087 | 0.0014 | 0.0154 | 0.0083 |
| rs7640424 | T | 0.0113 | 0.0015 | -0.0127 | 0.0097 |
| rs7650602 | T | -0.0089 | 0.0014 | -0.0071 | 0.0082 |
| rs76552497 | T | -0.0120 | 0.0019 | 0.0157 | 0.0107 |
| rs76577427 | C | 0.0203 | 0.0024 | -0.0182 | 0.0160 |
| rs7672622 | A | -0.0088 | 0.0016 | 0.0015 | 0.0088 |
| rs767943 | A | -0.0122 | 0.0016 | 0.0052 | 0.0092 |
| rs7683416 | T | 0.0139 | 0.0014 | 0.0031 | 0.0080 |
| rs76876592 | A | 0.0126 | 0.0020 | -0.0075 | 0.0123 |
| rs76878669 | C | 0.0141 | 0.0017 | -0.0022 | 0.0092 |
| rs7692359 | T | -0.0109 | 0.0017 | 0.0044 | 0.0105 |
| rs76957677 | T | -0.0297 | 0.0054 | 0.0526 | 0.0442 |
| rs77025239 | A | -0.0140 | 0.0019 | -0.0070 | 0.0102 |
| rs7714719 | C | 0.0116 | 0.0020 | -0.0025 | 0.0114 |
| rs7716161 | C | 0.0105 | 0.0019 | 0.0155 | 0.0112 |
| rs77201694 | A | 0.0156 | 0.0022 | -0.0156 | 0.0154 |
| rs77370942 | A | -0.0147 | 0.0027 | 0.0090 | 0.0176 |
| rs77554090 | T | -0.0186 | 0.0027 | -0.0061 | 0.0173 |
| rs77609760 | A | 0.0236 | 0.0028 | -0.0027 | 0.0185 |
| rs7766240 | T | -0.0134 | 0.0024 | 0.0007 | 0.0135 |
| rs77702622 | A | -0.0213 | 0.0029 | 0.0030 | 0.0198 |
| rs77702819 | T | 0.0187 | 0.0025 | -0.0077 | 0.0158 |
| rs7772172 | A | 0.0091 | 0.0014 | -0.0011 | 0.0090 |
| rs7775100 | T | -0.0087 | 0.0014 | 0.0133 | 0.0079 |
| rs77826402 | T | 0.0151 | 0.0027 | 0.0275 | 0.0154 |
| rs7796103 | C | -0.0077 | 0.0014 | 0.0192 | 0.0084 |
| rs7799141 | A | 0.0099 | 0.0015 | 0.0066 | 0.0085 |
| rs77999825 | A | 0.0294 | 0.0039 | -0.0102 | 0.0286 |
| rs7803932 | A | 0.0127 | 0.0019 | -0.0169 | 0.0117 |
| rs780569 | A | -0.0085 | 0.0016 | 0.0085 | 0.0087 |
| rs78116078 | C | 0.0086 | 0.0016 | 0.0059 | 0.0099 |
| rs7816777 | T | -0.0088 | 0.0015 | 0.0017 | 0.0087 |
| rs78193153 | A | 0.0191 | 0.0029 | 0.0037 | 0.0129 |
| rs7823700 | T | 0.0132 | 0.0024 | -0.0069 | 0.0169 |
| rs78365243 | T | 0.0181 | 0.0033 | -0.0199 | 0.0212 |
| rs78440611 | A | -0.0147 | 0.0024 | 0.0039 | 0.0153 |
| rs7849480 | A | -0.0084 | 0.0014 | 0.0203 | 0.0080 |
| rs7849487 | T | -0.0167 | 0.0015 | -0.0203 | 0.0081 |
| rs7855503 | C | 0.0088 | 0.0015 | 0.0117 | 0.0083 |
| rs78648104 | T | -0.0141 | 0.0025 | 0.0584 | 0.0142 |
| rs78702390 | C | 0.0167 | 0.0030 | -0.0433 | 0.0183 |
| rs78714229 | T | 0.0172 | 0.0030 | -0.0076 | 0.0192 |
| rs7875078 | A | -0.0078 | 0.0014 | 0.0204 | 0.0081 |
| rs78918150 | T | -0.0080 | 0.0015 | 0.0098 | 0.0088 |
| rs7894722 | T | 0.0085 | 0.0014 | -0.0062 | 0.0084 |
| rs7899270 | A | -0.0084 | 0.0015 | -0.0014 | 0.0088 |
| rs7905192 | T | 0.0084 | 0.0014 | 0.0011 | 0.0083 |
| rs790647 | A | -0.0145 | 0.0017 | 0.0117 | 0.0091 |
| rs7910403 | T | 0.0122 | 0.0018 | -0.0015 | 0.0112 |
| rs7914674 | T | -0.0139 | 0.0022 | -0.0185 | 0.0099 |
| rs7920624 | A | 0.0120 | 0.0014 | -0.0232 | 0.0080 |
| rs7924036 | T | 0.0133 | 0.0014 | -0.0064 | 0.0080 |
| rs79265434 | A | -0.0197 | 0.0022 | 0.0390 | 0.0132 |
| rs7928017 | A | 0.0096 | 0.0014 | -0.0160 | 0.0081 |
| rs7931563 | T | -0.0109 | 0.0014 | 0.0018 | 0.0081 |
| rs795230 | T | 0.0081 | 0.0014 | 0.0031 | 0.0081 |
| rs7958371 | A | -0.0085 | 0.0015 | -0.0047 | 0.0084 |
| rs79585412 | T | -0.0178 | 0.0031 | 0.0036 | 0.0175 |
| rs7965154 | A | -0.0085 | 0.0015 | 0.0028 | 0.0085 |
| rs7967550 | A | -0.0086 | 0.0014 | 0.0068 | 0.0085 |
| rs7972246 | T | 0.0110 | 0.0015 | -0.0020 | 0.0085 |
| rs79728014 | A | -0.0134 | 0.0020 | 0.0051 | 0.0126 |
| rs7974852 | A | 0.0117 | 0.0014 | 0.0100 | 0.0079 |
| rs7977614 | A | -0.0134 | 0.0016 | -0.0133 | 0.0096 |
| rs79798166 | A | 0.0172 | 0.0025 | 0.0082 | 0.0165 |
| rs79855925 | T | -0.0194 | 0.0033 | -0.0114 | 0.0226 |
| rs79994730 | T | 0.0240 | 0.0035 | 0.0617 | 0.0244 |
| rs79997166 | A | 0.0172 | 0.0030 | 0.0364 | 0.0208 |
| rs80037907 | T | -0.0107 | 0.0019 | 0.0186 | 0.0118 |
| rs8008382 | T | -0.0106 | 0.0015 | 0.0069 | 0.0085 |
| rs8009933 | A | 0.0088 | 0.0015 | -0.0068 | 0.0083 |
| rs8016504 | A | -0.0087 | 0.0014 | -0.0032 | 0.0077 |
| rs80223410 | T | -0.0118 | 0.0020 | 0.0273 | 0.0143 |
| rs8024 | A | -0.0115 | 0.0015 | 0.0254 | 0.0089 |
| rs80257979 | T | -0.0252 | 0.0040 | -0.0009 | 0.0260 |
| rs8030487 | A | 0.0085 | 0.0015 | -0.0218 | 0.0095 |
| rs803619 | T | -0.0173 | 0.0023 | 0.0057 | 0.0132 |
| rs8046072 | A | 0.0101 | 0.0018 | 0.0006 | 0.0104 |
| rs8052523 | T | -0.0084 | 0.0014 | 0.0068 | 0.0085 |
| rs8066044 | A | 0.0090 | 0.0016 | -0.0233 | 0.0090 |
| rs806816 | A | 0.0101 | 0.0016 | 0.0173 | 0.0100 |
| rs8097125 | T | 0.0086 | 0.0014 | 0.0140 | 0.0086 |
| rs8103741 | A | -0.0108 | 0.0019 | 0.0019 | 0.0105 |
| rs818415 | T | -0.0110 | 0.0018 | 0.0178 | 0.0097 |
| rs8192465 | A | -0.0272 | 0.0049 | 0.0545 | 0.0373 |
| rs852771 | T | 0.0092 | 0.0015 | -0.0039 | 0.0091 |
| rs853286 | T | 0.0124 | 0.0023 | -0.0111 | 0.0114 |
| rs854796 | A | 0.0090 | 0.0015 | -0.0046 | 0.0086 |
| rs868456 | A | 0.0089 | 0.0015 | 0.0056 | 0.0089 |
| rs870589 | A | -0.0079 | 0.0014 | 0.0002 | 0.0080 |
| rs884108 | A | -0.0094 | 0.0017 | -0.0140 | 0.0097 |
| rs891793 | C | -0.0099 | 0.0014 | 0.0082 | 0.0080 |
| rs893522 | A | 0.0149 | 0.0025 | -0.0153 | 0.0151 |
| rs902820 | A | -0.0096 | 0.0014 | -0.0013 | 0.0080 |
| rs911149 | T | 0.0091 | 0.0017 | 0.0000 | 0.0099 |
| rs912883 | T | 0.0087 | 0.0015 | 0.0136 | 0.0083 |
| rs913509 | A | 0.0096 | 0.0015 | 0.0027 | 0.0085 |
| rs925161 | C | -0.0078 | 0.0014 | 0.0081 | 0.0081 |
| rs9267658 | T | 0.0138 | 0.0020 | -0.0209 | 0.0136 |
| rs9267677 | T | 0.0180 | 0.0024 | 0.0240 | 0.0147 |
| rs9289300 | T | -0.0164 | 0.0019 | -0.0026 | 0.0115 |
| rs9291437 | C | -0.0081 | 0.0014 | -0.0035 | 0.0080 |
| rs9294770 | T | 0.0083 | 0.0014 | 0.0035 | 0.0083 |
| rs929511 | T | -0.0160 | 0.0021 | 0.0139 | 0.0121 |
| rs9300612 | T | 0.0084 | 0.0015 | 0.0042 | 0.0083 |
| rs933738 | A | -0.0110 | 0.0018 | -0.0045 | 0.0104 |
| rs9349956 | A | -0.0159 | 0.0018 | 0.0158 | 0.0116 |
| rs9359939 | A | -0.0105 | 0.0016 | -0.0101 | 0.0091 |
| rs936496 | A | -0.0079 | 0.0014 | 0.0134 | 0.0085 |
| rs9371883 | C | -0.0087 | 0.0015 | 0.0066 | 0.0086 |
| rs9373363 | A | -0.0111 | 0.0016 | -0.0104 | 0.0087 |
| rs9375188 | T | 0.0212 | 0.0014 | 0.0015 | 0.0079 |
| rs9375403 | C | 0.0126 | 0.0016 | -0.0049 | 0.0099 |
| rs9386110 | T | -0.0163 | 0.0023 | 0.0059 | 0.0124 |
| rs9388490 | T | 0.0091 | 0.0014 | 0.0163 | 0.0086 |
| rs939400 | T | -0.0095 | 0.0015 | 0.0114 | 0.0083 |
| rs9411331 | A | 0.0145 | 0.0015 | -0.0240 | 0.0083 |
| rs9435340 | A | 0.0086 | 0.0015 | -0.0002 | 0.0082 |
| rs9442750 | A | -0.0089 | 0.0016 | -0.0079 | 0.0097 |
| rs9446060 | A | 0.0077 | 0.0014 | -0.0099 | 0.0082 |
| rs9465509 | A | -0.0084 | 0.0014 | -0.0009 | 0.0080 |
| rs9490512 | A | -0.0136 | 0.0014 | 0.0000 | 0.0079 |
| rs9492774 | C | 0.0080 | 0.0015 | -0.0028 | 0.0079 |
| rs9513416 | A | -0.0116 | 0.0019 | -0.0203 | 0.0122 |
| rs9513754 | T | 0.0096 | 0.0016 | 0.0079 | 0.0091 |
| rs9527662 | A | 0.0094 | 0.0014 | -0.0017 | 0.0091 |
| rs9527905 | A | -0.0094 | 0.0014 | 0.0134 | 0.0080 |
| rs9529146 | T | 0.0127 | 0.0017 | 0.0024 | 0.0090 |
| rs9536462 | A | 0.0133 | 0.0020 | 0.0088 | 0.0106 |
| rs9540718 | A | -0.0089 | 0.0014 | -0.0105 | 0.0080 |
| rs9545395 | T | 0.0138 | 0.0021 | -0.0204 | 0.0107 |
| rs9556958 | T | -0.0110 | 0.0014 | 0.0109 | 0.0075 |
| rs9563168 | A | 0.0101 | 0.0017 | -0.0208 | 0.0100 |
| rs9568798 | T | -0.0093 | 0.0016 | -0.0068 | 0.0091 |
| rs9597907 | T | 0.0152 | 0.0027 | -0.0127 | 0.0158 |
| rs9611597 | A | -0.0107 | 0.0019 | 0.0011 | 0.0128 |
| rs9616906 | A | 0.0122 | 0.0014 | -0.0177 | 0.0086 |
| rs9616947 | T | -0.0134 | 0.0015 | 0.0116 | 0.0087 |
| rs9633970 | T | 0.0091 | 0.0016 | -0.0146 | 0.0093 |
| rs9649 | T | 0.0108 | 0.0019 | -0.0175 | 0.0122 |
| rs9655780 | A | -0.0153 | 0.0019 | -0.0130 | 0.0119 |
| rs9666728 | T | -0.0082 | 0.0014 | 0.0162 | 0.0079 |
| rs9683585 | C | 0.0084 | 0.0014 | 0.0073 | 0.0078 |
| rs969512 | A | -0.0106 | 0.0015 | 0.0051 | 0.0085 |
| rs9771228 | T | 0.0096 | 0.0015 | -0.0075 | 0.0086 |
| rs977143 | A | 0.0085 | 0.0015 | -0.0067 | 0.0088 |
| rs978807 | A | 0.0139 | 0.0018 | -0.0070 | 0.0095 |
| rs981230 | T | 0.0100 | 0.0014 | -0.0087 | 0.0084 |
| rs981883 | A | 0.0099 | 0.0014 | -0.0232 | 0.0080 |
| rs9820604 | T | 0.0115 | 0.0018 | -0.0056 | 0.0098 |
| rs9821664 | A | 0.0094 | 0.0017 | -0.0073 | 0.0092 |
| rs9830359 | T | -0.0147 | 0.0022 | 0.0128 | 0.0122 |
| rs9844755 | A | 0.0083 | 0.0015 | -0.0032 | 0.0085 |
| rs9849884 | A | -0.0090 | 0.0014 | -0.0011 | 0.0084 |
| rs9853928 | T | -0.0127 | 0.0017 | 0.0062 | 0.0089 |
| rs9858921 | A | -0.0087 | 0.0014 | 0.0075 | 0.0078 |
| rs9859556 | T | 0.0290 | 0.0015 | -0.0071 | 0.0094 |
| rs9866123 | A | 0.0086 | 0.0014 | -0.0034 | 0.0081 |
| rs9870317 | A | 0.0086 | 0.0015 | 0.0093 | 0.0082 |
| rs9877225 | T | 0.0090 | 0.0016 | -0.0128 | 0.0093 |
| rs9882532 | T | 0.0124 | 0.0015 | 0.0005 | 0.0087 |
| rs9886703 | A | -0.0136 | 0.0019 | -0.0060 | 0.0099 |
| rs9888796 | T | 0.0113 | 0.0016 | 0.0008 | 0.0095 |
| rs9916901 | T | 0.0092 | 0.0016 | -0.0071 | 0.0085 |
| rs9924031 | C | 0.0093 | 0.0015 | -0.0112 | 0.0091 |
| rs9926649 | T | 0.0177 | 0.0031 | -0.0197 | 0.0171 |
| rs9927049 | A | 0.0099 | 0.0016 | 0.0165 | 0.0099 |
| rs9927137 | A | 0.0082 | 0.0014 | 0.0033 | 0.0087 |
| rs9927842 | T | -0.0130 | 0.0020 | 0.0065 | 0.0107 |
| rs9929556 | T | -0.0099 | 0.0014 | 0.0020 | 0.0081 |
| rs9929762 | A | 0.0096 | 0.0014 | -0.0053 | 0.0079 |
| rs9929993 | T | -0.0102 | 0.0015 | 0.0093 | 0.0081 |
| rs9933256 | A | 0.0119 | 0.0014 | -0.0138 | 0.0092 |
| rs995698 | A | -0.0099 | 0.0014 | -0.0107 | 0.0086 |
| rs9974899 | T | -0.0093 | 0.0016 | -0.0068 | 0.0098 |
| rs9977825 | T | -0.0092 | 0.0015 | 0.0178 | 0.0090 |

#### Supplementary table 16: Genetic association estimates for the two-sample regression-based MVMR analysis estimating the effect of SBP on CAD, adjusted for education. ea=effect allele; gx=SBP; gy=CAD; gz=education; se=standard error

| SNP | ea | gx | gx_se | gy | gy_se | gz | gz_se |
| --- | --- | --- | --- | --- | --- | --- | --- |
| rs10024506 | G | 0.3036 | 0.0521 | 0.0131 | 0.0102 | 0.0004 | 0.0020 |
| rs10048404 | C | 0.2683 | 0.0458 | 0.0192 | 0.0100 | -0.0055 | 0.0018 |
| rs1010064 | A | 0.4007 | 0.0571 | 0.0401 | 0.0114 | -0.0015 | 0.0022 |
| rs10195405 | A | 0.2918 | 0.0486 | 0.0166 | 0.0104 | -0.0035 | 0.0019 |
| rs10212058 | A | 0.3278 | 0.0558 | 0.0355 | 0.0122 | -0.0003 | 0.0022 |
| rs1051447 | C | 0.2715 | 0.0489 | 0.0065 | 0.0099 | 0.0003 | 0.0019 |
| rs10775143 | C | 0.2586 | 0.0453 | -0.0066 | 0.0096 | 0.0075 | 0.0017 |
| rs1077795 | A | 0.2939 | 0.0503 | 0.0193 | 0.0111 | 0.0000 | 0.0019 |
| rs10817007 | T | -0.4373 | 0.0661 | -0.0231 | 0.0131 | 0.0045 | 0.0026 |
| rs10883543 | G | -0.6091 | 0.0705 | 0.0147 | 0.0168 | 0.0000 | 0.0027 |
| rs10977670 | A | 0.3011 | 0.0463 | -0.0118 | 0.0097 | 0.0012 | 0.0018 |
| rs10995311 | C | 0.3440 | 0.0446 | -0.0113 | 0.0096 | 0.0059 | 0.0017 |
| rs11070245 | T | -0.2798 | 0.0444 | 0.0050 | 0.0093 | -0.0052 | 0.0017 |
| rs11145807 | A | 0.2734 | 0.0453 | 0.0081 | 0.0107 | 0.0017 | 0.0018 |
| rs11146456 | A | -0.4230 | 0.0695 | -0.0022 | 0.0190 | 0.0026 | 0.0027 |
| rs11187838 | G | 0.4660 | 0.0446 | -0.0242 | 0.0093 | 0.0028 | 0.0017 |
| rs11191580 | T | 0.9647 | 0.0829 | 0.0732 | 0.0142 | 0.0022 | 0.0031 |
| rs113044050 | C | 0.3637 | 0.0641 | 0.0232 | 0.0130 | -0.0046 | 0.0024 |
| rs113523959 | G | 0.3852 | 0.0627 | -0.0031 | 0.0162 | -0.0046 | 0.0025 |
| rs113605702 | G | 0.4460 | 0.0798 | 0.0097 | 0.0171 | -0.0134 | 0.0031 |
| rs115262049 | A | 0.4875 | 0.0780 | 0.0321 | 0.0200 | -0.0032 | 0.0030 |
| rs11605215 | G | -0.3191 | 0.0559 | -0.0157 | 0.0117 | -0.0014 | 0.0021 |
| rs11616710 | C | -0.5056 | 0.0778 | -0.0179 | 0.0196 | 0.0028 | 0.0029 |
| rs11641308 | T | -0.3272 | 0.0470 | -0.0311 | 0.0104 | 0.0008 | 0.0018 |
| rs11667829 | G | -0.2636 | 0.0472 | -0.0359 | 0.0108 | 0.0046 | 0.0018 |
| rs116734066 | C | 0.4365 | 0.0764 | -0.0227 | 0.0164 | 0.0024 | 0.0030 |
| rs11697820 | C | 0.2499 | 0.0451 | 0.0068 | 0.0096 | -0.0020 | 0.0017 |
| rs11719057 | G | 0.2714 | 0.0467 | 0.0253 | 0.0100 | -0.0014 | 0.0018 |
| rs11721038 | T | 0.4978 | 0.0841 | 0.0464 | 0.0154 | -0.0024 | 0.0032 |
| rs1173771 | A | -0.6369 | 0.0450 | -0.0119 | 0.0093 | 0.0000 | 0.0017 |
| rs11774829 | T | 0.4089 | 0.0739 | 0.0209 | 0.0175 | 0.0029 | 0.0028 |
| rs11795365 | G | -0.3642 | 0.0642 | 0.0185 | 0.0129 | -0.0015 | 0.0025 |
| rs11821781 | A | -0.3392 | 0.0507 | -0.0038 | 0.0109 | 0.0026 | 0.0020 |
| rs11874 | G | -0.6251 | 0.0642 | -0.0497 | 0.0156 | 0.0068 | 0.0025 |
| rs12035750 | T | -0.3499 | 0.0454 | -0.0138 | 0.0101 | 0.0085 | 0.0018 |
| rs12055028 | G | -0.3149 | 0.0479 | 0.0012 | 0.0098 | 0.0000 | 0.0018 |
| rs12147852 | G | -0.3294 | 0.0516 | -0.0149 | 0.0110 | -0.0012 | 0.0020 |
| rs12258967 | C | 0.6150 | 0.0484 | 0.0257 | 0.0112 | 0.0011 | 0.0019 |
| rs1229984 | T | -1.0787 | 0.1483 | -0.0333 | 0.0186 | 0.0229 | 0.0057 |
| rs1250258 | C | 0.3174 | 0.0501 | 0.0328 | 0.0108 | 0.0021 | 0.0019 |
| rs12509892 | T | -0.3709 | 0.0443 | -0.0397 | 0.0094 | 0.0037 | 0.0017 |
| rs12543884 | A | -0.3423 | 0.0579 | -0.0105 | 0.0111 | -0.0027 | 0.0022 |
| rs12544711 | G | 0.2944 | 0.0449 | 0.0011 | 0.0093 | -0.0002 | 0.0017 |
| rs12567136 | C | 0.7688 | 0.0595 | 0.0232 | 0.0128 | 0.0015 | 0.0023 |
| rs12608469 | T | -0.5593 | 0.0935 | 0.0495 | 0.0212 | -0.0057 | 0.0036 |
| rs1262894 | A | 0.4733 | 0.0836 | -0.0130 | 0.0191 | -0.0053 | 0.0034 |
| rs1263146 | G | 0.2574 | 0.0444 | -0.0026 | 0.0095 | -0.0045 | 0.0017 |
| rs12643599 | A | 0.3161 | 0.0459 | 0.0512 | 0.0095 | -0.0011 | 0.0018 |
| rs12699561 | G | -0.2481 | 0.0456 | -0.0246 | 0.0098 | 0.0055 | 0.0018 |
| rs12705390 | G | -0.7711 | 0.0555 | -0.0174 | 0.0110 | 0.0051 | 0.0021 |
| rs12822344 | C | -0.2893 | 0.0448 | -0.0109 | 0.0095 | 0.0005 | 0.0017 |
| rs12906962 | T | -0.2770 | 0.0477 | -0.0166 | 0.0102 | -0.0041 | 0.0018 |
| rs1290784 | C | -0.3710 | 0.0446 | -0.0114 | 0.0094 | -0.0011 | 0.0017 |
| rs13107325 | C | 0.6251 | 0.0841 | 0.0067 | 0.0223 | 0.0188 | 0.0034 |
| rs13129779 | C | 0.2546 | 0.0446 | 0.0251 | 0.0094 | -0.0004 | 0.0017 |
| rs13219548 | C | -0.4362 | 0.0445 | -0.0066 | 0.0092 | 0.0009 | 0.0017 |
| rs13436194 | A | 0.3628 | 0.0446 | 0.0206 | 0.0093 | 0.0039 | 0.0017 |
| rs137923903 | C | 1.2912 | 0.1937 | 0.0361 | 0.0454 | 0.0046 | 0.0078 |
| rs1397985 | G | 0.3098 | 0.0473 | 0.0075 | 0.0099 | 0.0015 | 0.0018 |
| rs1422279 | C | -0.3138 | 0.0456 | -0.0060 | 0.0096 | -0.0045 | 0.0018 |
| rs1436138 | A | 0.3017 | 0.0463 | 0.0081 | 0.0098 | 0.0029 | 0.0018 |
| rs145339349 | G | -1.1270 | 0.1676 | -0.0356 | 0.0429 | -0.0010 | 0.0065 |
| rs1458038 | C | -0.7179 | 0.0487 | -0.0461 | 0.0102 | -0.0028 | 0.0019 |
| rs149932962 | G | -0.3024 | 0.0511 | 0.0241 | 0.0123 | 0.0014 | 0.0020 |
| rs150857355 | G | -0.8776 | 0.1531 | -0.0783 | 0.0444 | -0.0029 | 0.0060 |
| rs151157965 | G | -0.4568 | 0.0751 | 0.0079 | 0.0162 | 0.0059 | 0.0029 |
| rs1530558 | T | 0.3654 | 0.0643 | -0.0072 | 0.0152 | -0.0035 | 0.0027 |
| rs1542977 | G | -0.2805 | 0.0443 | -0.0295 | 0.0093 | 0.0065 | 0.0017 |
| rs1543270 | C | 0.3593 | 0.0445 | -0.0226 | 0.0094 | -0.0028 | 0.0017 |
| rs1543927 | T | 0.4427 | 0.0502 | 0.0205 | 0.0101 | -0.0087 | 0.0019 |
| rs167479 | G | 0.5274 | 0.0442 | 0.0146 | 0.0135 | 0.0009 | 0.0020 |
| rs17011002 | C | -0.5018 | 0.0638 | -0.0103 | 0.0130 | 0.0041 | 0.0025 |
| rs17173238 | A | -0.2918 | 0.0489 | -0.0154 | 0.0113 | -0.0018 | 0.0019 |
| rs1757915 | G | -0.3347 | 0.0461 | -0.0185 | 0.0096 | -0.0007 | 0.0018 |
| rs17677603 | A | -0.3326 | 0.0454 | -0.0283 | 0.0096 | -0.0027 | 0.0018 |
| rs1779240 | G | 0.3653 | 0.0521 | 0.0183 | 0.0107 | -0.0032 | 0.0020 |
| rs1887320 | G | -0.3471 | 0.0444 | -0.0226 | 0.0091 | 0.0020 | 0.0017 |
| rs2017199 | G | 0.3274 | 0.0485 | -0.0030 | 0.0106 | -0.0007 | 0.0019 |
| rs2052692 | A | 0.2673 | 0.0475 | -0.0084 | 0.0108 | 0.0064 | 0.0018 |
| rs2062316 | A | -0.3519 | 0.0445 | -0.0107 | 0.0100 | 0.0030 | 0.0017 |
| rs2111834 | G | 0.4380 | 0.0684 | -0.0223 | 0.0163 | -0.0107 | 0.0027 |
| rs2129561 | A | -0.3354 | 0.0451 | -0.0166 | 0.0094 | 0.0004 | 0.0017 |
| rs2177843 | C | -0.4184 | 0.0628 | 0.0015 | 0.0125 | 0.0024 | 0.0024 |
| rs2237306 | C | 0.3706 | 0.0642 | 0.0023 | 0.0126 | -0.0205 | 0.0024 |
| rs2249105 | A | 0.3532 | 0.0458 | -0.0033 | 0.0097 | -0.0039 | 0.0018 |
| rs2285815 | C | 0.2952 | 0.0500 | 0.0184 | 0.0101 | -0.0039 | 0.0019 |
| rs2288276 | G | -0.4101 | 0.0729 | -0.0071 | 0.0136 | -0.0017 | 0.0028 |
| rs2301597 | T | 0.4335 | 0.0448 | -0.0002 | 0.0094 | -0.0052 | 0.0017 |
| rs2303083 | G | 0.4692 | 0.0559 | 0.0158 | 0.0125 | -0.0105 | 0.0022 |
| rs2306363 | G | 0.4059 | 0.0546 | 0.0499 | 0.0117 | -0.0052 | 0.0022 |
| rs231708 | G | 0.2696 | 0.0476 | 0.0035 | 0.0100 | 0.0010 | 0.0018 |
| rs2423514 | A | 0.3053 | 0.0443 | 0.0181 | 0.0095 | 0.0005 | 0.0017 |
| rs2469133 | G | -0.2652 | 0.0466 | -0.0220 | 0.0098 | -0.0036 | 0.0018 |
| rs2469997 | G | 0.3808 | 0.0570 | -0.0012 | 0.0127 | 0.0011 | 0.0022 |
| rs2493296 | C | -0.4144 | 0.0642 | -0.0560 | 0.0147 | -0.0003 | 0.0025 |
| rs2499801 | G | 0.3601 | 0.0569 | -0.0518 | 0.0123 | -0.0012 | 0.0022 |
| rs2606738 | T | -0.3227 | 0.0470 | -0.0130 | 0.0095 | 0.0065 | 0.0018 |
| rs2610989 | T | -0.3378 | 0.0503 | -0.0239 | 0.0112 | 0.0077 | 0.0019 |
| rs2613203 | A | -0.3618 | 0.0569 | -0.0335 | 0.0118 | -0.0040 | 0.0022 |
| rs263017 | A | 0.2394 | 0.0444 | 0.0017 | 0.0092 | -0.0032 | 0.0017 |
| rs2643826 | C | -0.3654 | 0.0445 | -0.0001 | 0.0096 | 0.0014 | 0.0017 |
| rs2681492 | T | 0.7161 | 0.0589 | -0.0735 | 0.0113 | 0.0021 | 0.0023 |
| rs268263 | T | -0.5684 | 0.0517 | -0.0418 | 0.0109 | -0.0050 | 0.0020 |
| rs2764834 | T | -0.3148 | 0.0539 | -0.0042 | 0.0109 | 0.0011 | 0.0021 |
| rs2782652 | T | -0.2821 | 0.0443 | -0.0223 | 0.0097 | -0.0121 | 0.0017 |
| rs2782980 | T | -0.3734 | 0.0492 | -0.0149 | 0.0106 | -0.0027 | 0.0019 |
| rs2867114 | C | 0.4161 | 0.0759 | 0.0416 | 0.0126 | -0.0053 | 0.0025 |
| rs2897515 | T | 0.2629 | 0.0445 | 0.0118 | 0.0093 | 0.0031 | 0.0017 |
| rs2978456 | T | -0.2517 | 0.0443 | 0.0026 | 0.0101 | 0.0066 | 0.0018 |
| rs3101442 | T | -0.3048 | 0.0443 | -0.0138 | 0.0094 | 0.0017 | 0.0017 |
| rs3211995 | G | 0.4331 | 0.0607 | 0.0062 | 0.0141 | -0.0090 | 0.0024 |
| rs34991912 | T | 0.2671 | 0.0449 | 0.0465 | 0.0094 | -0.0003 | 0.0017 |
| rs35021474 | C | 0.4695 | 0.0455 | -0.0006 | 0.0104 | -0.0067 | 0.0018 |
| rs35443 | G | 0.3896 | 0.0454 | -0.0015 | 0.0096 | 0.0009 | 0.0018 |
| rs35726503 | A | 0.3459 | 0.0447 | -0.0004 | 0.0098 | -0.0041 | 0.0017 |
| rs35807464 | G | -0.5299 | 0.0889 | -0.0220 | 0.0184 | -0.0087 | 0.0032 |
| rs3786516 | T | -0.3211 | 0.0538 | 0.0149 | 0.0110 | -0.0023 | 0.0020 |
| rs3790604 | C | -0.7446 | 0.0846 | -0.0138 | 0.0161 | 0.0033 | 0.0032 |
| rs3826537 | A | -0.2872 | 0.0447 | 0.0037 | 0.0100 | 0.0018 | 0.0017 |
| rs3827750 | C | -0.5341 | 0.0705 | -0.0395 | 0.0131 | -0.0052 | 0.0027 |
| rs3887266 | C | -0.3884 | 0.0694 | -0.0179 | 0.0143 | -0.0007 | 0.0027 |
| rs4141663 | C | 0.2757 | 0.0449 | 0.0094 | 0.0093 | 0.0018 | 0.0017 |
| rs42038 | C | 0.3773 | 0.0484 | 0.0313 | 0.0109 | 0.0011 | 0.0019 |
| rs4246616 | C | -0.2718 | 0.0445 | -0.0007 | 0.0098 | -0.0041 | 0.0017 |
| rs434578 | C | 0.3810 | 0.0634 | 0.0369 | 0.0141 | -0.0012 | 0.0025 |
| rs4480845 | T | 0.4017 | 0.0463 | -0.0173 | 0.0098 | -0.0006 | 0.0018 |
| rs4648815 | G | 0.2905 | 0.0447 | 0.0050 | 0.0112 | -0.0074 | 0.0018 |
| rs4666493 | G | -0.3577 | 0.0448 | -0.0130 | 0.0094 | -0.0007 | 0.0017 |
| rs4690775 | A | 0.2629 | 0.0451 | 0.0203 | 0.0095 | 0.0031 | 0.0017 |
| rs4937515 | G | 0.3455 | 0.0453 | 0.0075 | 0.0103 | -0.0046 | 0.0017 |
| rs55924432 | C | -0.2770 | 0.0451 | -0.0213 | 0.0100 | -0.0023 | 0.0018 |
| rs56352102 | C | -0.5273 | 0.0570 | -0.0295 | 0.0120 | -0.0004 | 0.0022 |
| rs57139556 | A | 0.5824 | 0.0857 | 0.0457 | 0.0157 | -0.0026 | 0.0033 |
| rs57301765 | G | -0.5131 | 0.0608 | -0.0697 | 0.0113 | 0.0052 | 0.0023 |
| rs597808 | A | 0.4025 | 0.0443 | 0.0612 | 0.0106 | -0.0024 | 0.0017 |
| rs604723 | T | -0.6293 | 0.0497 | -0.0327 | 0.0101 | 0.0049 | 0.0019 |
| rs6062625 | T | -0.3285 | 0.0512 | -0.0201 | 0.0118 | 0.0035 | 0.0020 |
| rs61148001 | C | 0.3262 | 0.0546 | 0.0086 | 0.0108 | 0.0000 | 0.0021 |
| rs62162674 | G | 0.2691 | 0.0445 | 0.0115 | 0.0096 | 0.0006 | 0.0017 |
| rs62229372 | C | -0.3966 | 0.0672 | -0.0043 | 0.0168 | 0.0037 | 0.0026 |
| rs6453400 | G | 0.2515 | 0.0445 | 0.0053 | 0.0096 | -0.0035 | 0.0017 |
| rs6461992 | A | -0.7171 | 0.0853 | 0.0089 | 0.0172 | 0.0014 | 0.0032 |
| rs6704991 | A | -0.2749 | 0.0461 | -0.0128 | 0.0099 | 0.0019 | 0.0018 |
| rs6708660 | T | 0.2771 | 0.0452 | -0.0143 | 0.0097 | -0.0002 | 0.0017 |
| rs6775384 | G | -0.2988 | 0.0441 | -0.0294 | 0.0094 | 0.0249 | 0.0017 |
| rs6787069 | C | -0.3038 | 0.0496 | 0.0119 | 0.0104 | 0.0085 | 0.0019 |
| rs6815273 | G | 0.2850 | 0.0450 | 0.0124 | 0.0101 | 0.0002 | 0.0017 |
| rs6963105 | A | -0.2766 | 0.0446 | -0.0167 | 0.0111 | 0.0092 | 0.0018 |
| rs7019061 | C | 0.2794 | 0.0489 | 0.0130 | 0.0100 | -0.0031 | 0.0019 |
| rs7070797 | G | 0.5992 | 0.0635 | 0.0129 | 0.0154 | -0.0046 | 0.0025 |
| rs7107356 | A | -0.4367 | 0.0441 | -0.0063 | 0.0093 | 0.0055 | 0.0017 |
| rs7123754 | T | 0.3299 | 0.0472 | 0.0164 | 0.0098 | -0.0032 | 0.0018 |
| rs7125196 | T | 0.4486 | 0.0693 | 0.0172 | 0.0127 | 0.0106 | 0.0026 |
| rs71313932 | G | -0.2825 | 0.0491 | -0.0359 | 0.0105 | -0.0010 | 0.0019 |
| rs7200432 | G | 0.3351 | 0.0484 | 0.0109 | 0.0101 | 0.0025 | 0.0019 |
| rs72683923 | T | 1.0062 | 0.1588 | 0.0390 | 0.0368 | -0.0110 | 0.0062 |
| rs72792829 | C | 0.3598 | 0.0534 | 0.0131 | 0.0104 | -0.0010 | 0.0021 |
| rs72812818 | G | 0.2708 | 0.0482 | 0.0109 | 0.0108 | -0.0009 | 0.0018 |
| rs72843959 | C | 0.2788 | 0.0471 | -0.0239 | 0.0271 | 0.0031 | 0.0018 |
| rs7302981 | A | 0.3398 | 0.0457 | 0.0003 | 0.0100 | -0.0007 | 0.0018 |
| rs73046792 | G | 0.4241 | 0.0595 | 0.0163 | 0.0163 | 0.0001 | 0.0024 |
| rs73050466 | G | 0.8133 | 0.1316 | -0.0128 | 0.0340 | -0.0064 | 0.0048 |
| rs7306710 | T | -0.2759 | 0.0445 | -0.0212 | 0.0098 | 0.0082 | 0.0017 |
| rs73075659 | A | 0.3378 | 0.0467 | 0.0062 | 0.0106 | 0.0026 | 0.0018 |
| rs7315688 | G | 0.2693 | 0.0484 | 0.0191 | 0.0102 | -0.0024 | 0.0019 |
| rs73563812 | G | 0.3946 | 0.0521 | 0.0202 | 0.0107 | 0.0065 | 0.0020 |
| rs7439366 | T | -0.2487 | 0.0443 | 0.0001 | 0.0102 | 0.0005 | 0.0017 |
| rs7459106 | A | 0.3574 | 0.0562 | 0.0327 | 0.0115 | -0.0045 | 0.0022 |
| rs75271047 | G | 0.4691 | 0.0800 | 0.0607 | 0.0169 | -0.0057 | 0.0031 |
| rs7532726 | A | 0.2910 | 0.0500 | 0.0208 | 0.0107 | -0.0002 | 0.0019 |
| rs7553325 | C | -0.2516 | 0.0443 | 0.0201 | 0.0097 | 0.0000 | 0.0017 |
| rs75777337 | T | -0.6027 | 0.0724 | -0.0420 | 0.0159 | 0.0026 | 0.0027 |
| rs75989961 | T | -0.4855 | 0.0807 | -0.0312 | 0.0216 | 0.0027 | 0.0032 |
| rs77870048 | C | -0.9363 | 0.0989 | -0.0144 | 0.0304 | -0.0063 | 0.0040 |
| rs79349366 | C | 0.7946 | 0.1255 | -0.0266 | 0.0351 | 0.0157 | 0.0048 |
| rs7938342 | T | -0.5398 | 0.0449 | -0.0097 | 0.0104 | 0.0015 | 0.0018 |
| rs7963801 | T | -0.2501 | 0.0448 | -0.0218 | 0.0102 | 0.0107 | 0.0017 |
| rs8027450 | C | -0.5420 | 0.0474 | -0.0573 | 0.0109 | 0.0012 | 0.0018 |
| rs8076588 | C | 0.2841 | 0.0443 | 0.0088 | 0.0095 | 0.0050 | 0.0017 |
| rs861585 | G | -0.2584 | 0.0445 | -0.0075 | 0.0094 | 0.0067 | 0.0017 |
| rs891511 | G | 0.3979 | 0.0479 | 0.0214 | 0.0109 | -0.0042 | 0.0019 |
| rs9294987 | T | -0.2568 | 0.0445 | 0.0239 | 0.0101 | -0.0012 | 0.0017 |
| rs9373523 | T | 0.2556 | 0.0450 | 0.0009 | 0.0093 | -0.0022 | 0.0017 |
| rs940743 | T | -0.2682 | 0.0444 | -0.0333 | 0.0098 | -0.0002 | 0.0017 |
| rs9472040 | A | -0.2797 | 0.0473 | -0.0200 | 0.0096 | -0.0006 | 0.0018 |
| rs9773022 | T | -0.2937 | 0.0443 | 0.0078 | 0.0110 | -0.0008 | 0.0018 |
| rs9907379 | T | -0.4026 | 0.0542 | -0.0358 | 0.0111 | -0.0009 | 0.0021 |
| rs9993149 | T | 0.2797 | 0.0442 | 0.0043 | 0.0092 | -0.0053 | 0.0017 |

#### Supplementary table 17: Genetic association estimates for the two-sample regression-based MVMR analysis estimating the effect of SBP on MI, adjusted for education. ea=effect allele; gx=SBP; gy=MI; gz=education; se=standard error

| SNP | ea | gx | gx_se | gy | gy_se | gz | gz_se |
| --- | --- | --- | --- | --- | --- | --- | --- |
| rs10024506 | G | 0.3036 | 0.0521 | 0.0213 | 0.0114 | 0.0004 | 0.0020 |
| rs10048404 | C | 0.2683 | 0.0458 | 0.0068 | 0.0111 | -0.0055 | 0.0018 |
| rs1010064 | A | 0.4007 | 0.0571 | 0.0406 | 0.0127 | -0.0015 | 0.0022 |
| rs10195405 | A | 0.2918 | 0.0486 | 0.0086 | 0.0111 | -0.0035 | 0.0019 |
| rs10212058 | A | 0.3278 | 0.0558 | 0.0274 | 0.0134 | -0.0003 | 0.0022 |
| rs1051447 | C | 0.2715 | 0.0489 | 0.0131 | 0.0109 | 0.0003 | 0.0019 |
| rs10775143 | C | 0.2586 | 0.0453 | -0.0001 | 0.0107 | 0.0075 | 0.0017 |
| rs1077795 | A | 0.2939 | 0.0503 | 0.0086 | 0.0121 | 0.0000 | 0.0019 |
| rs10817007 | T | -0.4373 | 0.0661 | -0.0317 | 0.0146 | 0.0045 | 0.0026 |
| rs10883543 | G | -0.6091 | 0.0705 | 0.0193 | 0.0183 | 0.0000 | 0.0027 |
| rs10977670 | A | 0.3011 | 0.0463 | -0.0025 | 0.0107 | 0.0012 | 0.0018 |
| rs10995311 | C | 0.3440 | 0.0446 | -0.0162 | 0.0105 | 0.0059 | 0.0017 |
| rs11070245 | T | -0.2798 | 0.0444 | 0.0075 | 0.0103 | -0.0052 | 0.0017 |
| rs11145807 | A | 0.2734 | 0.0453 | 0.0155 | 0.0120 | 0.0017 | 0.0018 |
| rs11146456 | A | -0.4230 | 0.0695 | -0.0195 | 0.0212 | 0.0026 | 0.0027 |
| rs11187838 | G | 0.4660 | 0.0446 | -0.0190 | 0.0104 | 0.0028 | 0.0017 |
| rs11191580 | T | 0.9647 | 0.0829 | 0.0729 | 0.0158 | 0.0022 | 0.0031 |
| rs113044050 | C | 0.3637 | 0.0641 | 0.0130 | 0.0143 | -0.0046 | 0.0024 |
| rs113523959 | G | 0.3852 | 0.0627 | 0.0004 | 0.0180 | -0.0046 | 0.0025 |
| rs113605702 | G | 0.4460 | 0.0798 | 0.0049 | 0.0192 | -0.0134 | 0.0031 |
| rs115262049 | A | 0.4875 | 0.0780 | 0.0264 | 0.0218 | -0.0032 | 0.0030 |
| rs11605215 | G | -0.3191 | 0.0559 | 0.0029 | 0.0131 | -0.0014 | 0.0021 |
| rs11616710 | C | -0.5056 | 0.0778 | -0.0098 | 0.0215 | 0.0028 | 0.0029 |
| rs11641308 | T | -0.3272 | 0.0470 | -0.0149 | 0.0116 | 0.0008 | 0.0018 |
| rs11667829 | G | -0.2636 | 0.0472 | -0.0314 | 0.0118 | 0.0046 | 0.0018 |
| rs116734066 | C | 0.4365 | 0.0764 | -0.0453 | 0.0178 | 0.0024 | 0.0030 |
| rs11697820 | C | 0.2499 | 0.0451 | 0.0104 | 0.0106 | -0.0020 | 0.0017 |
| rs11719057 | G | 0.2714 | 0.0467 | 0.0101 | 0.0111 | -0.0014 | 0.0018 |
| rs11721038 | T | 0.4978 | 0.0841 | 0.0381 | 0.0171 | -0.0024 | 0.0032 |
| rs1173771 | A | -0.6369 | 0.0450 | -0.0048 | 0.0105 | 0.0000 | 0.0017 |
| rs11774829 | T | 0.4089 | 0.0739 | 0.0116 | 0.0196 | 0.0029 | 0.0028 |
| rs11795365 | G | -0.3642 | 0.0642 | 0.0282 | 0.0145 | -0.0015 | 0.0025 |
| rs11821781 | A | -0.3392 | 0.0507 | -0.0104 | 0.0120 | 0.0026 | 0.0020 |
| rs11874 | G | -0.6251 | 0.0642 | -0.0494 | 0.0171 | 0.0068 | 0.0025 |
| rs12035750 | T | -0.3499 | 0.0454 | -0.0035 | 0.0112 | 0.0085 | 0.0018 |
| rs12055028 | G | -0.3149 | 0.0479 | 0.0049 | 0.0108 | 0.0000 | 0.0018 |
| rs12147852 | G | -0.3294 | 0.0516 | -0.0116 | 0.0123 | -0.0012 | 0.0020 |
| rs12258967 | C | 0.6150 | 0.0484 | 0.0294 | 0.0122 | 0.0011 | 0.0019 |
| rs1229984 | T | -1.0787 | 0.1483 | -0.0308 | 0.0208 | 0.0229 | 0.0057 |
| rs1250258 | C | 0.3174 | 0.0501 | 0.0306 | 0.0116 | 0.0021 | 0.0019 |
| rs12509892 | T | -0.3709 | 0.0443 | -0.0320 | 0.0103 | 0.0037 | 0.0017 |
| rs12543884 | A | -0.3423 | 0.0579 | -0.0153 | 0.0124 | -0.0027 | 0.0022 |
| rs12544711 | G | 0.2944 | 0.0449 | -0.0002 | 0.0104 | -0.0002 | 0.0017 |
| rs12567136 | C | 0.7688 | 0.0595 | 0.0182 | 0.0141 | 0.0015 | 0.0023 |
| rs12608469 | T | -0.5593 | 0.0935 | 0.0490 | 0.0239 | -0.0057 | 0.0036 |
| rs1262894 | A | 0.4733 | 0.0836 | -0.0187 | 0.0214 | -0.0053 | 0.0034 |
| rs1263146 | G | 0.2574 | 0.0444 | -0.0056 | 0.0102 | -0.0045 | 0.0017 |
| rs12643599 | A | 0.3161 | 0.0459 | 0.0531 | 0.0105 | -0.0011 | 0.0018 |
| rs12699561 | G | -0.2481 | 0.0456 | -0.0253 | 0.0110 | 0.0055 | 0.0018 |
| rs12705390 | G | -0.7711 | 0.0555 | 0.0028 | 0.0123 | 0.0051 | 0.0021 |
| rs12822344 | C | -0.2893 | 0.0448 | -0.0098 | 0.0106 | 0.0005 | 0.0017 |
| rs12906962 | T | -0.2770 | 0.0477 | -0.0240 | 0.0115 | -0.0041 | 0.0018 |
| rs1290784 | C | -0.3710 | 0.0446 | -0.0073 | 0.0104 | -0.0011 | 0.0017 |
| rs13107325 | C | 0.6251 | 0.0841 | 0.0015 | 0.0244 | 0.0188 | 0.0034 |
| rs13129779 | C | 0.2546 | 0.0446 | 0.0168 | 0.0104 | -0.0004 | 0.0017 |
| rs13219548 | C | -0.4362 | 0.0445 | -0.0044 | 0.0102 | 0.0009 | 0.0017 |
| rs13436194 | A | 0.3628 | 0.0446 | 0.0287 | 0.0102 | 0.0039 | 0.0017 |
| rs137923903 | C | 1.2912 | 0.1937 | 0.0681 | 0.0487 | 0.0046 | 0.0078 |
| rs1397985 | G | 0.3098 | 0.0473 | 0.0008 | 0.0110 | 0.0015 | 0.0018 |
| rs1422279 | C | -0.3138 | 0.0456 | -0.0150 | 0.0106 | -0.0045 | 0.0018 |
| rs1436138 | A | 0.3017 | 0.0463 | -0.0086 | 0.0110 | 0.0029 | 0.0018 |
| rs145339349 | G | -1.1270 | 0.1676 | -0.0589 | 0.0480 | -0.0010 | 0.0065 |
| rs1458038 | C | -0.7179 | 0.0487 | -0.0467 | 0.0113 | -0.0028 | 0.0019 |
| rs149932962 | G | -0.3024 | 0.0511 | 0.0287 | 0.0136 | 0.0014 | 0.0020 |
| rs150857355 | G | -0.8776 | 0.1531 | -0.0799 | 0.0504 | -0.0029 | 0.0060 |
| rs151157965 | G | -0.4568 | 0.0751 | -0.0109 | 0.0181 | 0.0059 | 0.0029 |
| rs1530558 | T | 0.3654 | 0.0643 | -0.0206 | 0.0161 | -0.0035 | 0.0027 |
| rs1542977 | G | -0.2805 | 0.0443 | -0.0279 | 0.0104 | 0.0065 | 0.0017 |
| rs1543270 | C | 0.3593 | 0.0445 | -0.0139 | 0.0104 | -0.0028 | 0.0017 |
| rs1543927 | T | 0.4427 | 0.0502 | 0.0348 | 0.0113 | -0.0087 | 0.0019 |
| rs167479 | G | 0.5274 | 0.0442 | 0.0128 | 0.0158 | 0.0009 | 0.0020 |
| rs17011002 | C | -0.5018 | 0.0638 | 0.0023 | 0.0146 | 0.0041 | 0.0025 |
| rs17173238 | A | -0.2918 | 0.0489 | -0.0091 | 0.0125 | -0.0018 | 0.0019 |
| rs1757915 | G | -0.3347 | 0.0461 | -0.0107 | 0.0106 | -0.0007 | 0.0018 |
| rs17677603 | A | -0.3326 | 0.0454 | -0.0240 | 0.0106 | -0.0027 | 0.0018 |
| rs1779240 | G | 0.3653 | 0.0521 | 0.0217 | 0.0118 | -0.0032 | 0.0020 |
| rs1887320 | G | -0.3471 | 0.0444 | -0.0115 | 0.0102 | 0.0020 | 0.0017 |
| rs2017199 | G | 0.3274 | 0.0485 | -0.0043 | 0.0117 | -0.0007 | 0.0019 |
| rs2052692 | A | 0.2673 | 0.0475 | -0.0123 | 0.0121 | 0.0064 | 0.0018 |
| rs2062316 | A | -0.3519 | 0.0445 | -0.0120 | 0.0110 | 0.0030 | 0.0017 |
| rs2111834 | G | 0.4380 | 0.0684 | -0.0009 | 0.0180 | -0.0107 | 0.0027 |
| rs2129561 | A | -0.3354 | 0.0451 | -0.0146 | 0.0107 | 0.0004 | 0.0017 |
| rs2177843 | C | -0.4184 | 0.0628 | -0.0073 | 0.0140 | 0.0024 | 0.0024 |
| rs2237306 | C | 0.3706 | 0.0642 | 0.0096 | 0.0143 | -0.0205 | 0.0024 |
| rs2249105 | A | 0.3532 | 0.0458 | -0.0006 | 0.0105 | -0.0039 | 0.0018 |
| rs2285815 | C | 0.2952 | 0.0500 | 0.0230 | 0.0112 | -0.0039 | 0.0019 |
| rs2288276 | G | -0.4101 | 0.0729 | -0.0116 | 0.0151 | -0.0017 | 0.0028 |
| rs2301597 | T | 0.4335 | 0.0448 | 0.0010 | 0.0103 | -0.0052 | 0.0017 |
| rs2303083 | G | 0.4692 | 0.0559 | 0.0139 | 0.0138 | -0.0105 | 0.0022 |
| rs2306363 | G | 0.4059 | 0.0546 | 0.0421 | 0.0129 | -0.0052 | 0.0022 |
| rs231708 | G | 0.2696 | 0.0476 | 0.0109 | 0.0111 | 0.0010 | 0.0018 |
| rs2423514 | A | 0.3053 | 0.0443 | 0.0251 | 0.0104 | 0.0005 | 0.0017 |
| rs2469133 | G | -0.2652 | 0.0466 | -0.0373 | 0.0109 | -0.0036 | 0.0018 |
| rs2469997 | G | 0.3808 | 0.0570 | -0.0035 | 0.0140 | 0.0011 | 0.0022 |
| rs2493296 | C | -0.4144 | 0.0642 | -0.0596 | 0.0162 | -0.0003 | 0.0025 |
| rs2499801 | G | 0.3601 | 0.0569 | -0.0526 | 0.0134 | -0.0012 | 0.0022 |
| rs2606738 | T | -0.3227 | 0.0470 | -0.0061 | 0.0105 | 0.0065 | 0.0018 |
| rs2610989 | T | -0.3378 | 0.0503 | -0.0374 | 0.0125 | 0.0077 | 0.0019 |
| rs2613203 | A | -0.3618 | 0.0569 | -0.0191 | 0.0131 | -0.0040 | 0.0022 |
| rs263017 | A | 0.2394 | 0.0444 | 0.0085 | 0.0102 | -0.0032 | 0.0017 |
| rs2643826 | C | -0.3654 | 0.0445 | -0.0006 | 0.0106 | 0.0014 | 0.0017 |
| rs2681492 | T | 0.7161 | 0.0589 | -0.0715 | 0.0125 | 0.0021 | 0.0023 |
| rs268263 | T | -0.5684 | 0.0517 | -0.0408 | 0.0117 | -0.0050 | 0.0020 |
| rs2764834 | T | -0.3148 | 0.0539 | -0.0020 | 0.0120 | 0.0011 | 0.0021 |
| rs2782652 | T | -0.2821 | 0.0443 | -0.0184 | 0.0108 | -0.0121 | 0.0017 |
| rs2782980 | T | -0.3734 | 0.0492 | -0.0218 | 0.0118 | -0.0027 | 0.0019 |
| rs2867114 | C | 0.4161 | 0.0759 | 0.0213 | 0.0139 | -0.0053 | 0.0025 |
| rs2897515 | T | 0.2629 | 0.0445 | 0.0124 | 0.0103 | 0.0031 | 0.0017 |
| rs2978456 | T | -0.2517 | 0.0443 | 0.0047 | 0.0113 | 0.0066 | 0.0018 |
| rs3101442 | T | -0.3048 | 0.0443 | -0.0181 | 0.0105 | 0.0017 | 0.0017 |
| rs3211995 | G | 0.4331 | 0.0607 | -0.0074 | 0.0158 | -0.0090 | 0.0024 |
| rs34991912 | T | 0.2671 | 0.0449 | 0.0466 | 0.0104 | -0.0003 | 0.0017 |
| rs35021474 | C | 0.4695 | 0.0455 | -0.0134 | 0.0115 | -0.0067 | 0.0018 |
| rs35443 | G | 0.3896 | 0.0454 | 0.0008 | 0.0107 | 0.0009 | 0.0018 |
| rs35726503 | A | 0.3459 | 0.0447 | -0.0022 | 0.0109 | -0.0041 | 0.0017 |
| rs35807464 | G | -0.5299 | 0.0889 | -0.0309 | 0.0208 | -0.0087 | 0.0032 |
| rs3786516 | T | -0.3211 | 0.0538 | 0.0184 | 0.0121 | -0.0023 | 0.0020 |
| rs3790604 | C | -0.7446 | 0.0846 | -0.0184 | 0.0182 | 0.0033 | 0.0032 |
| rs3826537 | A | -0.2872 | 0.0447 | 0.0144 | 0.0111 | 0.0018 | 0.0017 |
| rs3827750 | C | -0.5341 | 0.0705 | -0.0408 | 0.0145 | -0.0052 | 0.0027 |
| rs3887266 | C | -0.3884 | 0.0694 | -0.0282 | 0.0156 | -0.0007 | 0.0027 |
| rs4141663 | C | 0.2757 | 0.0449 | 0.0069 | 0.0104 | 0.0018 | 0.0017 |
| rs42038 | C | 0.3773 | 0.0484 | 0.0234 | 0.0122 | 0.0011 | 0.0019 |
| rs4246616 | C | -0.2718 | 0.0445 | -0.0056 | 0.0105 | -0.0041 | 0.0017 |
| rs434578 | C | 0.3810 | 0.0634 | 0.0292 | 0.0153 | -0.0012 | 0.0025 |
| rs4480845 | T | 0.4017 | 0.0463 | -0.0164 | 0.0108 | -0.0006 | 0.0018 |
| rs4648815 | G | 0.2905 | 0.0447 | 0.0030 | 0.0123 | -0.0074 | 0.0018 |
| rs4666493 | G | -0.3577 | 0.0448 | -0.0107 | 0.0104 | -0.0007 | 0.0017 |
| rs4690775 | A | 0.2629 | 0.0451 | 0.0233 | 0.0105 | 0.0031 | 0.0017 |
| rs4937515 | G | 0.3455 | 0.0453 | -0.0018 | 0.0113 | -0.0046 | 0.0017 |
| rs55924432 | C | -0.2770 | 0.0451 | -0.0055 | 0.0112 | -0.0023 | 0.0018 |
| rs56352102 | C | -0.5273 | 0.0570 | -0.0187 | 0.0132 | -0.0004 | 0.0022 |
| rs57139556 | A | 0.5824 | 0.0857 | 0.0526 | 0.0174 | -0.0026 | 0.0033 |
| rs57301765 | G | -0.5131 | 0.0608 | -0.0412 | 0.0126 | 0.0052 | 0.0023 |
| rs597808 | A | 0.4025 | 0.0443 | 0.0699 | 0.0118 | -0.0024 | 0.0017 |
| rs604723 | T | -0.6293 | 0.0497 | -0.0288 | 0.0112 | 0.0049 | 0.0019 |
| rs6062625 | T | -0.3285 | 0.0512 | -0.0355 | 0.0132 | 0.0035 | 0.0020 |
| rs61148001 | C | 0.3262 | 0.0546 | 0.0144 | 0.0120 | 0.0000 | 0.0021 |
| rs62162674 | G | 0.2691 | 0.0445 | 0.0014 | 0.0103 | 0.0006 | 0.0017 |
| rs62229372 | C | -0.3966 | 0.0672 | 0.0046 | 0.0192 | 0.0037 | 0.0026 |
| rs6453400 | G | 0.2515 | 0.0445 | 0.0149 | 0.0107 | -0.0035 | 0.0017 |
| rs6461992 | A | -0.7171 | 0.0853 | 0.0095 | 0.0197 | 0.0014 | 0.0032 |
| rs6704991 | A | -0.2749 | 0.0461 | 0.0017 | 0.0107 | 0.0019 | 0.0018 |
| rs6708660 | T | 0.2771 | 0.0452 | -0.0090 | 0.0106 | -0.0002 | 0.0017 |
| rs6775384 | G | -0.2988 | 0.0441 | -0.0281 | 0.0104 | 0.0249 | 0.0017 |
| rs6787069 | C | -0.3038 | 0.0496 | 0.0187 | 0.0114 | 0.0085 | 0.0019 |
| rs6815273 | G | 0.2850 | 0.0450 | 0.0001 | 0.0112 | 0.0002 | 0.0017 |
| rs6963105 | A | -0.2766 | 0.0446 | -0.0123 | 0.0121 | 0.0092 | 0.0018 |
| rs7019061 | C | 0.2794 | 0.0489 | 0.0069 | 0.0111 | -0.0031 | 0.0019 |
| rs7070797 | G | 0.5992 | 0.0635 | 0.0233 | 0.0169 | -0.0046 | 0.0025 |
| rs7107356 | A | -0.4367 | 0.0441 | -0.0109 | 0.0104 | 0.0055 | 0.0017 |
| rs7123754 | T | 0.3299 | 0.0472 | 0.0157 | 0.0108 | -0.0032 | 0.0018 |
| rs7125196 | T | 0.4486 | 0.0693 | 0.0287 | 0.0143 | 0.0106 | 0.0026 |
| rs71313932 | G | -0.2825 | 0.0491 | -0.0308 | 0.0117 | -0.0010 | 0.0019 |
| rs7200432 | G | 0.3351 | 0.0484 | 0.0168 | 0.0112 | 0.0025 | 0.0019 |
| rs72683923 | T | 1.0062 | 0.1588 | 0.0812 | 0.0397 | -0.0110 | 0.0062 |
| rs72792829 | C | 0.3598 | 0.0534 | 0.0253 | 0.0116 | -0.0010 | 0.0021 |
| rs72812818 | G | 0.2708 | 0.0482 | 0.0082 | 0.0121 | -0.0009 | 0.0018 |
| rs72843959 | C | 0.2788 | 0.0471 | -0.0234 | 0.0315 | 0.0031 | 0.0018 |
| rs7302981 | A | 0.3398 | 0.0457 | 0.0036 | 0.0111 | -0.0007 | 0.0018 |
| rs73046792 | G | 0.4241 | 0.0595 | 0.0067 | 0.0180 | 0.0001 | 0.0024 |
| rs73050466 | G | 0.8133 | 0.1316 | 0.0107 | 0.0394 | -0.0064 | 0.0048 |
| rs7306710 | T | -0.2759 | 0.0445 | -0.0223 | 0.0108 | 0.0082 | 0.0017 |
| rs73075659 | A | 0.3378 | 0.0467 | 0.0130 | 0.0116 | 0.0026 | 0.0018 |
| rs7315688 | G | 0.2693 | 0.0484 | 0.0103 | 0.0115 | -0.0024 | 0.0019 |
| rs73563812 | G | 0.3946 | 0.0521 | 0.0115 | 0.0119 | 0.0065 | 0.0020 |
| rs7439366 | T | -0.2487 | 0.0443 | -0.0004 | 0.0113 | 0.0005 | 0.0017 |
| rs7459106 | A | 0.3574 | 0.0562 | 0.0345 | 0.0130 | -0.0045 | 0.0022 |
| rs75271047 | G | 0.4691 | 0.0800 | 0.0350 | 0.0185 | -0.0057 | 0.0031 |
| rs7532726 | A | 0.2910 | 0.0500 | 0.0214 | 0.0117 | -0.0002 | 0.0019 |
| rs7553325 | C | -0.2516 | 0.0443 | 0.0179 | 0.0107 | 0.0000 | 0.0017 |
| rs75777337 | T | -0.6027 | 0.0724 | -0.0433 | 0.0174 | 0.0026 | 0.0027 |
| rs75989961 | T | -0.4855 | 0.0807 | -0.0405 | 0.0236 | 0.0027 | 0.0032 |
| rs77870048 | C | -0.9363 | 0.0989 | -0.0007 | 0.0339 | -0.0063 | 0.0040 |
| rs79349366 | C | 0.7946 | 0.1255 | -0.0311 | 0.0388 | 0.0157 | 0.0048 |
| rs7938342 | T | -0.5398 | 0.0449 | 0.0004 | 0.0116 | 0.0015 | 0.0018 |
| rs7963801 | T | -0.2501 | 0.0448 | -0.0309 | 0.0113 | 0.0107 | 0.0017 |
| rs8027450 | C | -0.5420 | 0.0474 | -0.0602 | 0.0122 | 0.0012 | 0.0018 |
| rs8076588 | C | 0.2841 | 0.0443 | 0.0129 | 0.0105 | 0.0050 | 0.0017 |
| rs861585 | G | -0.2584 | 0.0445 | -0.0046 | 0.0104 | 0.0067 | 0.0017 |
| rs891511 | G | 0.3979 | 0.0479 | 0.0175 | 0.0123 | -0.0042 | 0.0019 |
| rs9294987 | T | -0.2568 | 0.0445 | 0.0146 | 0.0111 | -0.0012 | 0.0017 |
| rs9373523 | T | 0.2556 | 0.0450 | -0.0131 | 0.0104 | -0.0022 | 0.0017 |
| rs940743 | T | -0.2682 | 0.0444 | -0.0340 | 0.0108 | -0.0002 | 0.0017 |
| rs9472040 | A | -0.2797 | 0.0473 | -0.0164 | 0.0106 | -0.0006 | 0.0018 |
| rs9773022 | T | -0.2937 | 0.0443 | 0.0044 | 0.0120 | -0.0008 | 0.0018 |
| rs9907379 | T | -0.4026 | 0.0542 | -0.0229 | 0.0123 | -0.0009 | 0.0021 |
| rs9993149 | T | 0.2797 | 0.0442 | -0.0017 | 0.0102 | -0.0053 | 0.0017 |

#### Supplementary table 18: Genetic association estimates for the two-sample regression-based MVMR analysis estimating the effect of SBP on stroke, adjusted for education. ea=effect allele; gx=SBP; gy=stroke; gz=education; se=standard error

| SNP | ea | gx | gx_se | gy | gy_se | gz | gz_se |
| --- | --- | --- | --- | --- | --- | --- | --- |
| rs10024506 | G | 0.3036 | 0.0521 | 0.0048 | 0.0090 | 0.0004 | 0.0020 |
| rs10048404 | C | 0.2683 | 0.0458 | 0.0130 | 0.0088 | -0.0055 | 0.0018 |
| rs1010064 | A | 0.4007 | 0.0571 | -0.0100 | 0.0096 | -0.0015 | 0.0022 |
| rs10195405 | A | 0.2918 | 0.0486 | 0.0053 | 0.0087 | -0.0035 | 0.0019 |
| rs10212058 | A | 0.3278 | 0.0558 | 0.0045 | 0.0103 | -0.0003 | 0.0022 |
| rs1051447 | C | 0.2715 | 0.0489 | 0.0267 | 0.0088 | 0.0003 | 0.0019 |
| rs10775143 | C | 0.2586 | 0.0453 | -0.0040 | 0.0079 | 0.0075 | 0.0017 |
| rs1077795 | A | 0.2939 | 0.0503 | 0.0310 | 0.0102 | 0.0000 | 0.0019 |
| rs10817007 | T | -0.4373 | 0.0661 | 0.0088 | 0.0108 | 0.0045 | 0.0026 |
| rs10883543 | G | -0.6091 | 0.0705 | 0.0279 | 0.0144 | 0.0000 | 0.0027 |
| rs10977670 | A | 0.3011 | 0.0463 | -0.0016 | 0.0082 | 0.0012 | 0.0018 |
| rs10995311 | C | 0.3440 | 0.0446 | -0.0038 | 0.0086 | 0.0059 | 0.0017 |
| rs11070245 | T | -0.2798 | 0.0444 | 0.0061 | 0.0079 | -0.0052 | 0.0017 |
| rs11145807 | A | 0.2734 | 0.0453 | 0.0168 | 0.0093 | 0.0017 | 0.0018 |
| rs11146456 | A | -0.4230 | 0.0695 | -0.0037 | 0.0155 | 0.0026 | 0.0027 |
| rs11187838 | G | 0.4660 | 0.0446 | 0.0166 | 0.0078 | 0.0028 | 0.0017 |
| rs11191580 | T | 0.9647 | 0.0829 | 0.0023 | 0.0127 | 0.0022 | 0.0031 |
| rs113044050 | C | 0.3637 | 0.0641 | -0.0004 | 0.0118 | -0.0046 | 0.0024 |
| rs113523959 | G | 0.3852 | 0.0627 | 0.0234 | 0.0142 | -0.0046 | 0.0025 |
| rs113605702 | G | 0.4460 | 0.0798 | 0.0096 | 0.0142 | -0.0134 | 0.0031 |
| rs115262049 | A | 0.4875 | 0.0780 | 0.0043 | 0.0172 | -0.0032 | 0.0030 |
| rs11605215 | G | -0.3191 | 0.0559 | 0.0003 | 0.0096 | -0.0014 | 0.0021 |
| rs11616710 | C | -0.5056 | 0.0778 | -0.0360 | 0.0165 | 0.0028 | 0.0029 |
| rs11641308 | T | -0.3272 | 0.0470 | -0.0083 | 0.0085 | 0.0008 | 0.0018 |
| rs11667829 | G | -0.2636 | 0.0472 | -0.0048 | 0.0098 | 0.0046 | 0.0018 |
| rs116734066 | C | 0.4365 | 0.0764 | -0.0196 | 0.0157 | 0.0024 | 0.0030 |
| rs11697820 | C | 0.2499 | 0.0451 | 0.0053 | 0.0082 | -0.0020 | 0.0017 |
| rs11719057 | G | 0.2714 | 0.0467 | 0.0168 | 0.0089 | -0.0014 | 0.0018 |
| rs11721038 | T | 0.4978 | 0.0841 | 0.0224 | 0.0131 | -0.0024 | 0.0032 |
| rs1173771 | A | -0.6369 | 0.0450 | -0.0291 | 0.0081 | 0.0000 | 0.0017 |
| rs11774829 | T | 0.4089 | 0.0739 | 0.0289 | 0.0144 | 0.0029 | 0.0028 |
| rs11795365 | G | -0.3642 | 0.0642 | -0.0323 | 0.0108 | -0.0015 | 0.0025 |
| rs11821781 | A | -0.3392 | 0.0507 | 0.0101 | 0.0093 | 0.0026 | 0.0020 |
| rs11874 | G | -0.6251 | 0.0642 | -0.0502 | 0.0137 | 0.0068 | 0.0025 |
| rs12035750 | T | -0.3499 | 0.0454 | -0.0154 | 0.0084 | 0.0085 | 0.0018 |
| rs12055028 | G | -0.3149 | 0.0479 | 0.0010 | 0.0085 | 0.0000 | 0.0018 |
| rs12147852 | G | -0.3294 | 0.0516 | 0.0128 | 0.0094 | -0.0012 | 0.0020 |
| rs12258967 | C | 0.6150 | 0.0484 | 0.0256 | 0.0098 | 0.0011 | 0.0019 |
| rs1229984 | T | -1.0787 | 0.1483 | -0.0429 | 0.0174 | 0.0229 | 0.0057 |
| rs1250258 | C | 0.3174 | 0.0501 | 0.0251 | 0.0094 | 0.0021 | 0.0019 |
| rs12509892 | T | -0.3709 | 0.0443 | -0.0074 | 0.0082 | 0.0037 | 0.0017 |
| rs12543884 | A | -0.3423 | 0.0579 | 0.0012 | 0.0097 | -0.0027 | 0.0022 |
| rs12544711 | G | 0.2944 | 0.0449 | -0.0157 | 0.0081 | -0.0002 | 0.0017 |
| rs12567136 | C | 0.7688 | 0.0595 | 0.0203 | 0.0109 | 0.0015 | 0.0023 |
| rs12608469 | T | -0.5593 | 0.0935 | -0.0088 | 0.0167 | -0.0057 | 0.0036 |
| rs1262894 | A | 0.4733 | 0.0836 | 0.0165 | 0.0153 | -0.0053 | 0.0034 |
| rs1263146 | G | 0.2574 | 0.0444 | 0.0045 | 0.0079 | -0.0045 | 0.0017 |
| rs12643599 | A | 0.3161 | 0.0459 | 0.0185 | 0.0081 | -0.0011 | 0.0018 |
| rs12699561 | G | -0.2481 | 0.0456 | -0.0197 | 0.0081 | 0.0055 | 0.0018 |
| rs12705390 | G | -0.7711 | 0.0555 | -0.0323 | 0.0107 | 0.0051 | 0.0021 |
| rs12822344 | C | -0.2893 | 0.0448 | -0.0021 | 0.0081 | 0.0005 | 0.0017 |
| rs12906962 | T | -0.2770 | 0.0477 | -0.0017 | 0.0086 | -0.0041 | 0.0018 |
| rs1290784 | C | -0.3710 | 0.0446 | -0.0161 | 0.0083 | -0.0011 | 0.0017 |
| rs13107325 | C | 0.6251 | 0.0841 | 0.0005 | 0.0193 | 0.0188 | 0.0034 |
| rs13129779 | C | 0.2546 | 0.0446 | -0.0006 | 0.0082 | -0.0004 | 0.0017 |
| rs13219548 | C | -0.4362 | 0.0445 | -0.0266 | 0.0079 | 0.0009 | 0.0017 |
| rs13436194 | A | 0.3628 | 0.0446 | 0.0066 | 0.0081 | 0.0039 | 0.0017 |
| rs1397985 | G | 0.3098 | 0.0473 | 0.0060 | 0.0084 | 0.0015 | 0.0018 |
| rs1422279 | C | -0.3138 | 0.0456 | -0.0181 | 0.0080 | -0.0045 | 0.0018 |
| rs1436138 | A | 0.3017 | 0.0463 | 0.0036 | 0.0083 | 0.0029 | 0.0018 |
| rs145339349 | G | -1.1270 | 0.1676 | -0.0159 | 0.0393 | -0.0010 | 0.0065 |
| rs1458038 | C | -0.7179 | 0.0487 | -0.0261 | 0.0087 | -0.0028 | 0.0019 |
| rs149932962 | G | -0.3024 | 0.0511 | 0.0031 | 0.0104 | 0.0014 | 0.0020 |
| rs150857355 | G | -0.8776 | 0.1531 | -0.0029 | 0.0384 | -0.0029 | 0.0060 |
| rs151157965 | G | -0.4568 | 0.0751 | 0.0157 | 0.0133 | 0.0059 | 0.0029 |
| rs1530558 | T | 0.3654 | 0.0643 | -0.0020 | 0.0131 | -0.0035 | 0.0027 |
| rs1542977 | G | -0.2805 | 0.0443 | -0.0061 | 0.0079 | 0.0065 | 0.0017 |
| rs1543270 | C | 0.3593 | 0.0445 | 0.0189 | 0.0077 | -0.0028 | 0.0017 |
| rs1543927 | T | 0.4427 | 0.0502 | 0.0160 | 0.0086 | -0.0087 | 0.0019 |
| rs167479 | G | 0.5274 | 0.0442 | 0.0214 | 0.0116 | 0.0009 | 0.0020 |
| rs17011002 | C | -0.5018 | 0.0638 | -0.0068 | 0.0110 | 0.0041 | 0.0025 |
| rs17173238 | A | -0.2918 | 0.0489 | -0.0132 | 0.0095 | -0.0018 | 0.0019 |
| rs1757915 | G | -0.3347 | 0.0461 | -0.0183 | 0.0084 | -0.0007 | 0.0018 |
| rs17677603 | A | -0.3326 | 0.0454 | -0.0275 | 0.0081 | -0.0027 | 0.0018 |
| rs1779240 | G | 0.3653 | 0.0521 | 0.0109 | 0.0094 | -0.0032 | 0.0020 |
| rs1887320 | G | -0.3471 | 0.0444 | -0.0269 | 0.0080 | 0.0020 | 0.0017 |
| rs2017199 | G | 0.3274 | 0.0485 | 0.0005 | 0.0095 | -0.0007 | 0.0019 |
| rs2052692 | A | 0.2673 | 0.0475 | -0.0298 | 0.0091 | 0.0064 | 0.0018 |
| rs2062316 | A | -0.3519 | 0.0445 | -0.0299 | 0.0086 | 0.0030 | 0.0017 |
| rs2111834 | G | 0.4380 | 0.0684 | 0.0198 | 0.0140 | -0.0107 | 0.0027 |
| rs2129561 | A | -0.3354 | 0.0451 | -0.0221 | 0.0082 | 0.0004 | 0.0017 |
| rs2177843 | C | -0.4184 | 0.0628 | 0.0042 | 0.0105 | 0.0024 | 0.0024 |
| rs2237306 | C | 0.3706 | 0.0642 | 0.0106 | 0.0100 | -0.0205 | 0.0024 |
| rs2249105 | A | 0.3532 | 0.0458 | 0.0201 | 0.0081 | -0.0039 | 0.0018 |
| rs2285815 | C | 0.2952 | 0.0500 | -0.0088 | 0.0087 | -0.0039 | 0.0019 |
| rs2288276 | G | -0.4101 | 0.0729 | 0.0073 | 0.0115 | -0.0017 | 0.0028 |
| rs2301597 | T | 0.4335 | 0.0448 | -0.0086 | 0.0081 | -0.0052 | 0.0017 |
| rs2303083 | G | 0.4692 | 0.0559 | 0.0021 | 0.0108 | -0.0105 | 0.0022 |
| rs2306363 | G | 0.4059 | 0.0546 | 0.0274 | 0.0098 | -0.0052 | 0.0022 |
| rs231708 | G | 0.2696 | 0.0476 | 0.0246 | 0.0086 | 0.0010 | 0.0018 |
| rs2423514 | A | 0.3053 | 0.0443 | 0.0032 | 0.0084 | 0.0005 | 0.0017 |
| rs2469133 | G | -0.2652 | 0.0466 | -0.0173 | 0.0081 | -0.0036 | 0.0018 |
| rs2469997 | G | 0.3808 | 0.0570 | -0.0060 | 0.0110 | 0.0011 | 0.0022 |
| rs2493296 | C | -0.4144 | 0.0642 | -0.0425 | 0.0124 | -0.0003 | 0.0025 |
| rs2499801 | G | 0.3601 | 0.0569 | 0.0074 | 0.0103 | -0.0012 | 0.0022 |
| rs2606738 | T | -0.3227 | 0.0470 | -0.0081 | 0.0081 | 0.0065 | 0.0018 |
| rs2610989 | T | -0.3378 | 0.0503 | -0.0315 | 0.0091 | 0.0077 | 0.0019 |
| rs2613203 | A | -0.3618 | 0.0569 | 0.0139 | 0.0102 | -0.0040 | 0.0022 |
| rs263017 | A | 0.2394 | 0.0444 | 0.0037 | 0.0079 | -0.0032 | 0.0017 |
| rs2643826 | C | -0.3654 | 0.0445 | -0.0250 | 0.0083 | 0.0014 | 0.0017 |
| rs2681492 | T | 0.7161 | 0.0589 | 0.0361 | 0.0099 | 0.0021 | 0.0023 |
| rs268263 | T | -0.5684 | 0.0517 | -0.0027 | 0.0090 | -0.0050 | 0.0020 |
| rs2764834 | T | -0.3148 | 0.0539 | -0.0141 | 0.0092 | 0.0011 | 0.0021 |
| rs2782652 | T | -0.2821 | 0.0443 | -0.0014 | 0.0082 | -0.0121 | 0.0017 |
| rs2782980 | T | -0.3734 | 0.0492 | 0.0016 | 0.0089 | -0.0027 | 0.0019 |
| rs2867114 | C | 0.4161 | 0.0759 | 0.0020 | 0.0115 | -0.0053 | 0.0025 |
| rs2897515 | T | 0.2629 | 0.0445 | 0.0109 | 0.0079 | 0.0031 | 0.0017 |
| rs2978456 | T | -0.2517 | 0.0443 | 0.0036 | 0.0084 | 0.0066 | 0.0018 |
| rs3101442 | T | -0.3048 | 0.0443 | -0.0065 | 0.0080 | 0.0017 | 0.0017 |
| rs3211995 | G | 0.4331 | 0.0607 | 0.0082 | 0.0119 | -0.0090 | 0.0024 |
| rs34991912 | T | 0.2671 | 0.0449 | 0.0153 | 0.0081 | -0.0003 | 0.0017 |
| rs35443 | G | 0.3896 | 0.0454 | 0.0455 | 0.0083 | 0.0009 | 0.0018 |
| rs35726503 | A | 0.3459 | 0.0447 | -0.0003 | 0.0084 | -0.0041 | 0.0017 |
| rs35807464 | G | -0.5299 | 0.0889 | -0.0050 | 0.0151 | -0.0087 | 0.0032 |
| rs3786516 | T | -0.3211 | 0.0538 | 0.0094 | 0.0098 | -0.0023 | 0.0020 |
| rs3790604 | C | -0.7446 | 0.0846 | -0.0671 | 0.0129 | 0.0033 | 0.0032 |
| rs3826537 | A | -0.2872 | 0.0447 | -0.0034 | 0.0080 | 0.0018 | 0.0017 |
| rs3827750 | C | -0.5341 | 0.0705 | -0.0319 | 0.0115 | -0.0052 | 0.0027 |
| rs3887266 | C | -0.3884 | 0.0694 | -0.0384 | 0.0133 | -0.0007 | 0.0027 |
| rs4141663 | C | 0.2757 | 0.0449 | 0.0173 | 0.0079 | 0.0018 | 0.0017 |
| rs42038 | C | 0.3773 | 0.0484 | 0.0508 | 0.0095 | 0.0011 | 0.0019 |
| rs4246616 | C | -0.2718 | 0.0445 | -0.0140 | 0.0083 | -0.0041 | 0.0017 |
| rs434578 | C | 0.3810 | 0.0634 | 0.0176 | 0.0127 | -0.0012 | 0.0025 |
| rs4480845 | T | 0.4017 | 0.0463 | 0.0002 | 0.0084 | -0.0006 | 0.0018 |
| rs4648815 | G | 0.2905 | 0.0447 | 0.0050 | 0.0090 | -0.0074 | 0.0018 |
| rs4666493 | G | -0.3577 | 0.0448 | 0.0038 | 0.0082 | -0.0007 | 0.0017 |
| rs4690775 | A | 0.2629 | 0.0451 | 0.0257 | 0.0083 | 0.0031 | 0.0017 |
| rs4937515 | G | 0.3455 | 0.0453 | -0.0095 | 0.0089 | -0.0046 | 0.0017 |
| rs55924432 | C | -0.2770 | 0.0451 | -0.0339 | 0.0084 | -0.0023 | 0.0018 |
| rs56352102 | C | -0.5273 | 0.0570 | -0.0135 | 0.0105 | -0.0004 | 0.0022 |
| rs57139556 | A | 0.5824 | 0.0857 | 0.0172 | 0.0140 | -0.0026 | 0.0033 |
| rs57301765 | G | -0.5131 | 0.0608 | -0.0669 | 0.0096 | 0.0052 | 0.0023 |
| rs597808 | A | 0.4025 | 0.0443 | 0.0576 | 0.0093 | -0.0024 | 0.0017 |
| rs604723 | T | -0.6293 | 0.0497 | -0.0158 | 0.0083 | 0.0049 | 0.0019 |
| rs6062625 | T | -0.3285 | 0.0512 | -0.0117 | 0.0101 | 0.0035 | 0.0020 |
| rs61148001 | C | 0.3262 | 0.0546 | 0.0024 | 0.0090 | 0.0000 | 0.0021 |
| rs62162674 | G | 0.2691 | 0.0445 | 0.0249 | 0.0082 | 0.0006 | 0.0017 |
| rs62229372 | C | -0.3966 | 0.0672 | -0.0182 | 0.0135 | 0.0037 | 0.0026 |
| rs6453400 | G | 0.2515 | 0.0445 | 0.0079 | 0.0080 | -0.0035 | 0.0017 |
| rs6461992 | A | -0.7171 | 0.0853 | -0.0315 | 0.0125 | 0.0014 | 0.0032 |
| rs6704991 | A | -0.2749 | 0.0461 | -0.0090 | 0.0083 | 0.0019 | 0.0018 |
| rs6708660 | T | 0.2771 | 0.0452 | 0.0265 | 0.0091 | -0.0002 | 0.0017 |
| rs6775384 | G | -0.2988 | 0.0441 | -0.0093 | 0.0087 | 0.0249 | 0.0017 |
| rs6787069 | C | -0.3038 | 0.0496 | -0.0033 | 0.0092 | 0.0085 | 0.0019 |
| rs6815273 | G | 0.2850 | 0.0450 | 0.0125 | 0.0086 | 0.0002 | 0.0017 |
| rs6963105 | A | -0.2766 | 0.0446 | -0.0369 | 0.0096 | 0.0092 | 0.0018 |
| rs7019061 | C | 0.2794 | 0.0489 | -0.0003 | 0.0086 | -0.0031 | 0.0019 |
| rs7070797 | G | 0.5992 | 0.0635 | 0.0315 | 0.0132 | -0.0046 | 0.0025 |
| rs7107356 | A | -0.4367 | 0.0441 | 0.0154 | 0.0079 | 0.0055 | 0.0017 |
| rs7123754 | T | 0.3299 | 0.0472 | 0.0156 | 0.0081 | -0.0032 | 0.0018 |
| rs7125196 | T | 0.4486 | 0.0693 | 0.0240 | 0.0108 | 0.0106 | 0.0026 |
| rs71313932 | G | -0.2825 | 0.0491 | -0.0139 | 0.0090 | -0.0010 | 0.0019 |
| rs7200432 | G | 0.3351 | 0.0484 | 0.0150 | 0.0089 | 0.0025 | 0.0019 |
| rs72683923 | T | 1.0062 | 0.1588 | 0.0357 | 0.0380 | -0.0110 | 0.0062 |
| rs72792829 | C | 0.3598 | 0.0534 | -0.0132 | 0.0091 | -0.0010 | 0.0021 |
| rs72812818 | G | 0.2708 | 0.0482 | 0.0145 | 0.0091 | -0.0009 | 0.0018 |
| rs72843959 | C | 0.2788 | 0.0471 | 0.0223 | 0.0097 | 0.0031 | 0.0018 |
| rs7302981 | A | 0.3398 | 0.0457 | -0.0011 | 0.0089 | -0.0007 | 0.0018 |
| rs73046792 | G | 0.4241 | 0.0595 | 0.0098 | 0.0131 | 0.0001 | 0.0024 |
| rs73050466 | G | 0.8133 | 0.1316 | -0.0124 | 0.0287 | -0.0064 | 0.0048 |
| rs7306710 | T | -0.2759 | 0.0445 | -0.0109 | 0.0084 | 0.0082 | 0.0017 |
| rs73075659 | A | 0.3378 | 0.0467 | 0.0144 | 0.0093 | 0.0026 | 0.0018 |
| rs7315688 | G | 0.2693 | 0.0484 | 0.0338 | 0.0086 | -0.0024 | 0.0019 |
| rs73563812 | G | 0.3946 | 0.0521 | 0.0069 | 0.0089 | 0.0065 | 0.0020 |
| rs7439366 | T | -0.2487 | 0.0443 | -0.0134 | 0.0090 | 0.0005 | 0.0017 |
| rs7459106 | A | 0.3574 | 0.0562 | 0.0107 | 0.0095 | -0.0045 | 0.0022 |
| rs75271047 | G | 0.4691 | 0.0800 | 0.0234 | 0.0151 | -0.0057 | 0.0031 |
| rs7532726 | A | 0.2910 | 0.0500 | 0.0035 | 0.0098 | -0.0002 | 0.0019 |
| rs7553325 | C | -0.2516 | 0.0443 | -0.0008 | 0.0086 | 0.0000 | 0.0017 |
| rs75777337 | T | -0.6027 | 0.0724 | -0.0126 | 0.0136 | 0.0026 | 0.0027 |
| rs75989961 | T | -0.4855 | 0.0807 | -0.0035 | 0.0178 | 0.0027 | 0.0032 |
| rs79349366 | C | 0.7946 | 0.1255 | 0.0165 | 0.0276 | 0.0157 | 0.0048 |
| rs7938342 | T | -0.5398 | 0.0449 | -0.0329 | 0.0084 | 0.0015 | 0.0018 |
| rs7963801 | T | -0.2501 | 0.0448 | -0.0044 | 0.0088 | 0.0107 | 0.0017 |
| rs8027450 | C | -0.5420 | 0.0474 | -0.0303 | 0.0092 | 0.0012 | 0.0018 |
| rs8076588 | C | 0.2841 | 0.0443 | 0.0144 | 0.0079 | 0.0050 | 0.0017 |
| rs861585 | G | -0.2584 | 0.0445 | -0.0150 | 0.0081 | 0.0067 | 0.0017 |
| rs891511 | G | 0.3979 | 0.0479 | 0.0233 | 0.0089 | -0.0042 | 0.0019 |
| rs9294987 | T | -0.2568 | 0.0445 | 0.0030 | 0.0086 | -0.0012 | 0.0017 |
| rs9373523 | T | 0.2556 | 0.0450 | -0.0203 | 0.0081 | -0.0022 | 0.0017 |
| rs940743 | T | -0.2682 | 0.0444 | -0.0004 | 0.0092 | -0.0002 | 0.0017 |
| rs9472040 | A | -0.2797 | 0.0473 | -0.0449 | 0.0081 | -0.0006 | 0.0018 |
| rs9773022 | T | -0.2937 | 0.0443 | -0.0174 | 0.0088 | -0.0008 | 0.0018 |
| rs9907379 | T | -0.4026 | 0.0542 | -0.0257 | 0.0093 | -0.0009 | 0.0021 |
| rs9993149 | T | 0.2797 | 0.0442 | 0.0133 | 0.0078 | -0.0053 | 0.0017 |

#### Supplementary table 19: Genetic association estimates for the two-sample regression-based MVMR analysis estimating the effect of smoking on CAD, adjusted for education. ea=effect allele; gx=smoking; gy=CAD; gz=education; se=standard error

| SNP | ea | gx | gx_se | gy | gy_se | gz | gz_se |
| --- | --- | --- | --- | --- | --- | --- | --- |
| rs10282292 | C | 0.0128 | 0.0021 | 0.0227 | 0.0096 | -0.0010 | 0.0018 |
| rs1050847 | C | 0.0114 | 0.0020 | 0.0262 | 0.0103 | -0.0108 | 0.0017 |
| rs10516044 | G | -0.0199 | 0.0031 | -0.0038 | 0.0147 | 0.0055 | 0.0027 |
| rs10879871 | T | -0.0137 | 0.0021 | -0.0021 | 0.0095 | 0.0084 | 0.0018 |
| rs10918701 | G | 0.0116 | 0.0021 | 0.0236 | 0.0096 | -0.0010 | 0.0018 |
| rs10922907 | A | 0.0145 | 0.0020 | 0.0058 | 0.0094 | -0.0161 | 0.0017 |
| rs111243290 | T | -0.0290 | 0.0053 | 0.0007 | 0.0282 | 0.0190 | 0.0046 |
| rs1118814 | G | 0.0121 | 0.0021 | -0.0032 | 0.0098 | -0.0069 | 0.0018 |
| rs11210229 | A | 0.0167 | 0.0020 | 0.0001 | 0.0094 | -0.0043 | 0.0017 |
| rs112282219 | G | -0.0333 | 0.0050 | 0.0020 | 0.0273 | 0.0017 | 0.0044 |
| rs11255908 | T | -0.0145 | 0.0023 | 0.0102 | 0.0102 | 0.0050 | 0.0020 |
| rs113382419 | C | -0.0409 | 0.0032 | -0.0207 | 0.0190 | -0.0053 | 0.0028 |
| rs1149331 | G | -0.0110 | 0.0020 | -0.0077 | 0.0095 | 0.0075 | 0.0017 |
| rs11768481 | C | 0.0129 | 0.0021 | 0.0141 | 0.0104 | -0.0010 | 0.0018 |
| rs11783093 | C | 0.0227 | 0.0027 | 0.0350 | 0.0138 | -0.0024 | 0.0023 |
| rs11861214 | G | 0.0135 | 0.0024 | -0.0022 | 0.0112 | -0.0114 | 0.0020 |
| rs11948770 | T | -0.0148 | 0.0024 | 0.0058 | 0.0113 | -0.0028 | 0.0020 |
| rs1221148 | C | 0.0133 | 0.0020 | -0.0008 | 0.0095 | 0.0001 | 0.0017 |
| rs12244388 | G | -0.0192 | 0.0021 | 0.0363 | 0.0094 | 0.0065 | 0.0018 |
| rs1246265 | T | -0.0127 | 0.0022 | -0.0091 | 0.0101 | 0.0048 | 0.0019 |
| rs12481282 | G | -0.0129 | 0.0022 | 0.0266 | 0.0104 | 0.0027 | 0.0019 |
| rs12623702 | A | -0.0141 | 0.0021 | 0.0086 | 0.0099 | 0.0099 | 0.0017 |
| rs12708665 | A | -0.0131 | 0.0022 | -0.0107 | 0.0107 | 0.0089 | 0.0019 |
| rs12831617 | C | -0.0131 | 0.0023 | -0.0238 | 0.0115 | 0.0048 | 0.0020 |
| rs12967855 | A | 0.0117 | 0.0021 | -0.0096 | 0.0097 | -0.0194 | 0.0018 |
| rs13009008 | A | 0.0123 | 0.0021 | 0.0033 | 0.0101 | -0.0095 | 0.0018 |
| rs13016665 | C | -0.0123 | 0.0020 | -0.0051 | 0.0098 | 0.0012 | 0.0017 |
| rs13296519 | G | -0.0138 | 0.0020 | -0.0195 | 0.0099 | 0.0078 | 0.0017 |
| rs136233 | A | -0.0141 | 0.0025 | -0.0307 | 0.0119 | 0.0058 | 0.0022 |
| rs147412694 | G | -0.0168 | 0.0028 | -0.0145 | 0.0141 | -0.0008 | 0.0024 |
| rs17309874 | G | -0.0164 | 0.0023 | -0.0246 | 0.0118 | 0.0083 | 0.0019 |
| rs17553262 | A | -0.0182 | 0.0031 | 0.0046 | 0.0165 | 0.0112 | 0.0027 |
| rs17576594 | G | 0.0157 | 0.0022 | -0.0013 | 0.0103 | -0.0113 | 0.0019 |
| rs1827535 | T | -0.0124 | 0.0022 | -0.0131 | 0.0106 | 0.0019 | 0.0018 |
| rs1922018 | C | 0.0145 | 0.0021 | -0.0064 | 0.0100 | -0.0057 | 0.0018 |
| rs1931263 | G | -0.0109 | 0.0020 | -0.0138 | 0.0098 | 0.0120 | 0.0017 |
| rs1933270 | T | 0.0132 | 0.0021 | 0.0057 | 0.0094 | -0.0016 | 0.0018 |
| rs202645 | A | -0.0145 | 0.0025 | -0.0072 | 0.0113 | -0.0001 | 0.0021 |
| rs2062882 | G | -0.0117 | 0.0020 | -0.0006 | 0.0095 | 0.0080 | 0.0017 |
| rs2080870 | A | 0.0125 | 0.0023 | -0.0146 | 0.0112 | -0.0158 | 0.0020 |
| rs215600 | G | 0.0165 | 0.0021 | 0.0150 | 0.0095 | -0.0116 | 0.0018 |
| rs2160896 | C | -0.0120 | 0.0021 | -0.0144 | 0.0098 | 0.0106 | 0.0018 |
| rs2254710 | C | 0.0130 | 0.0023 | 0.0194 | 0.0105 | -0.0031 | 0.0020 |
| rs2401924 | G | 0.0151 | 0.0020 | 0.0066 | 0.0092 | 0.0044 | 0.0017 |
| rs245774 | A | -0.0129 | 0.0022 | -0.0113 | 0.0112 | 0.0070 | 0.0019 |
| rs2675638 | G | 0.0122 | 0.0020 | 0.0061 | 0.0093 | -0.0068 | 0.0017 |
| rs2678670 | A | 0.0126 | 0.0020 | 0.0097 | 0.0102 | -0.0099 | 0.0017 |
| rs2838834 | C | -0.0133 | 0.0022 | -0.0217 | 0.0098 | -0.0009 | 0.0019 |
| rs28485305 | C | 0.0115 | 0.0021 | 0.0112 | 0.0094 | -0.0008 | 0.0018 |
| rs2867112 | T | 0.0211 | 0.0027 | 0.0414 | 0.0125 | -0.0048 | 0.0023 |
| rs2890772 | G | -0.0196 | 0.0020 | 0.0039 | 0.0097 | -0.0036 | 0.0017 |
| rs2894808 | T | -0.0219 | 0.0037 | -0.0045 | 0.0167 | 0.0076 | 0.0031 |
| rs317021 | T | -0.0168 | 0.0026 | -0.0008 | 0.0127 | 0.0057 | 0.0022 |
| rs326341 | G | 0.0137 | 0.0020 | -0.0037 | 0.0094 | -0.0063 | 0.0017 |
| rs329120 | C | 0.0139 | 0.0020 | -0.0201 | 0.0093 | -0.0083 | 0.0017 |
| rs348809 | A | -0.0118 | 0.0021 | -0.0069 | 0.0097 | 0.0050 | 0.0018 |
| rs35120974 | A | -0.0121 | 0.0020 | 0.0018 | 0.0094 | -0.0049 | 0.0017 |
| rs35169606 | T | 0.0126 | 0.0021 | 0.0126 | 0.0112 | -0.0025 | 0.0018 |
| rs35175834 | G | -0.0236 | 0.0024 | 0.0112 | 0.0120 | 0.0129 | 0.0021 |
| rs35343344 | C | 0.0131 | 0.0023 | 0.0151 | 0.0112 | -0.0059 | 0.0020 |
| rs359243 | T | -0.0126 | 0.0021 | -0.0028 | 0.0098 | 0.0064 | 0.0017 |
| rs369230 | G | -0.0130 | 0.0022 | -0.0265 | 0.0124 | 0.0052 | 0.0019 |
| rs3742365 | T | -0.0155 | 0.0020 | -0.0162 | 0.0095 | -0.0059 | 0.0017 |
| rs3769949 | T | -0.0118 | 0.0020 | -0.0145 | 0.0095 | 0.0050 | 0.0017 |
| rs3811038 | T | -0.0137 | 0.0022 | 0.0045 | 0.0110 | 0.0066 | 0.0019 |
| rs3896224 | A | 0.0139 | 0.0020 | 0.0120 | 0.0093 | -0.0088 | 0.0017 |
| rs421983 | T | 0.0125 | 0.0020 | 0.0006 | 0.0092 | -0.0033 | 0.0017 |
| rs4391802 | A | 0.0149 | 0.0022 | 0.0048 | 0.0102 | -0.0072 | 0.0019 |
| rs4473348 | A | -0.0149 | 0.0023 | -0.0202 | 0.0111 | -0.0026 | 0.0020 |
| rs4543592 | T | -0.0124 | 0.0020 | -0.0067 | 0.0096 | 0.0067 | 0.0017 |
| rs4571506 | C | 0.0113 | 0.0020 | 0.0157 | 0.0094 | -0.0115 | 0.0017 |
| rs4671357 | T | -0.0136 | 0.0020 | -0.0083 | 0.0095 | 0.0091 | 0.0017 |
| rs4814873 | C | 0.0140 | 0.0024 | 0.0211 | 0.0109 | -0.0064 | 0.0020 |
| rs4949465 | T | -0.0168 | 0.0030 | -0.0229 | 0.0122 | 0.0129 | 0.0025 |
| rs4957528 | A | -0.0145 | 0.0025 | -0.0146 | 0.0127 | 0.0051 | 0.0021 |
| rs549845 | G | 0.0161 | 0.0022 | 0.0082 | 0.0098 | -0.0177 | 0.0019 |
| rs6011779 | C | 0.0276 | 0.0025 | 0.0187 | 0.0135 | -0.0042 | 0.0022 |
| rs60952428 | T | 0.0192 | 0.0035 | 0.0144 | 0.0139 | -0.0079 | 0.0031 |
| rs6119897 | G | -0.0185 | 0.0023 | -0.0202 | 0.0105 | 0.0069 | 0.0020 |
| rs62098013 | G | -0.0122 | 0.0021 | -0.0081 | 0.0107 | 0.0126 | 0.0018 |
| rs62135536 | C | 0.0352 | 0.0057 | 0.0038 | 0.0298 | -0.0092 | 0.0049 |
| rs62155874 | A | -0.0244 | 0.0030 | 0.0173 | 0.0154 | 0.0142 | 0.0026 |
| rs62175972 | T | 0.0313 | 0.0055 | 0.0303 | 0.0304 | -0.0300 | 0.0048 |
| rs624833 | T | 0.0132 | 0.0022 | 0.0104 | 0.0107 | -0.0132 | 0.0018 |
| rs6598539 | T | -0.0118 | 0.0020 | -0.0150 | 0.0092 | 0.0065 | 0.0017 |
| rs6741228 | T | 0.0114 | 0.0020 | 0.0202 | 0.0093 | -0.0075 | 0.0017 |
| rs67596067 | G | -0.0128 | 0.0021 | -0.0171 | 0.0100 | 0.0036 | 0.0018 |
| rs6778080 | T | 0.0158 | 0.0023 | 0.0106 | 0.0114 | -0.0139 | 0.0019 |
| rs6779302 | G | -0.0125 | 0.0021 | 0.0026 | 0.0098 | 0.0113 | 0.0018 |
| rs6935954 | A | 0.0137 | 0.0020 | -0.0109 | 0.0099 | -0.0138 | 0.0017 |
| rs6957896 | C | -0.0109 | 0.0020 | -0.0073 | 0.0096 | 0.0016 | 0.0017 |
| rs6962772 | A | 0.0159 | 0.0028 | 0.0087 | 0.0114 | -0.0040 | 0.0023 |
| rs7039819 | G | 0.0126 | 0.0020 | 0.0144 | 0.0095 | -0.0106 | 0.0017 |
| rs7077678 | C | 0.0123 | 0.0021 | 0.0116 | 0.0093 | 0.0016 | 0.0017 |
| rs71367545 | G | -0.0148 | 0.0025 | 0.0039 | 0.0122 | 0.0125 | 0.0021 |
| rs7144406 | A | -0.0135 | 0.0025 | -0.0036 | 0.0130 | 0.0051 | 0.0021 |
| rs7155595 | A | -0.0126 | 0.0021 | -0.0086 | 0.0103 | 0.0073 | 0.0018 |
| rs72678864 | G | 0.0178 | 0.0026 | 0.0103 | 0.0140 | -0.0123 | 0.0023 |
| rs7297175 | T | -0.0115 | 0.0020 | 0.0041 | 0.0098 | 0.0156 | 0.0017 |
| rs732083 | G | 0.0120 | 0.0021 | 0.0207 | 0.0098 | -0.0081 | 0.0018 |
| rs73220544 | A | -0.0156 | 0.0028 | -0.0082 | 0.0133 | 0.0069 | 0.0024 |
| rs7333559 | G | 0.0152 | 0.0025 | 0.0052 | 0.0120 | -0.0060 | 0.0021 |
| rs74086911 | G | 0.0213 | 0.0038 | 0.0019 | 0.0190 | -0.0184 | 0.0034 |
| rs7528604 | G | 0.0138 | 0.0020 | 0.0100 | 0.0093 | -0.0059 | 0.0017 |
| rs7553348 | G | 0.0138 | 0.0020 | -0.0033 | 0.0094 | -0.0004 | 0.0017 |
| rs7569203 | A | -0.0156 | 0.0022 | 0.0009 | 0.0098 | 0.0026 | 0.0018 |
| rs75742406 | G | 0.0139 | 0.0023 | 0.0080 | 0.0106 | -0.0010 | 0.0019 |
| rs76608582 | C | 0.0311 | 0.0049 | -0.0330 | 0.0379 | -0.0280 | 0.0045 |
| rs775758 | A | 0.0115 | 0.0020 | 0.0036 | 0.0093 | -0.0030 | 0.0017 |
| rs7766610 | C | 0.0181 | 0.0026 | 0.0122 | 0.0110 | 0.0008 | 0.0022 |
| rs7775552 | A | -0.0119 | 0.0020 | -0.0025 | 0.0093 | 0.0020 | 0.0017 |
| rs7807019 | A | -0.0150 | 0.0020 | -0.0189 | 0.0092 | 0.0106 | 0.0017 |
| rs7897434 | G | 0.0116 | 0.0021 | 0.0090 | 0.0103 | -0.0038 | 0.0018 |
| rs8042134 | T | -0.0141 | 0.0020 | 0.0139 | 0.0097 | 0.0079 | 0.0017 |
| rs8042849 | C | 0.0276 | 0.0021 | -0.0039 | 0.0101 | 0.0002 | 0.0018 |
| rs860326 | C | 0.0119 | 0.0020 | 0.0078 | 0.0096 | -0.0088 | 0.0017 |
| rs8614 | C | -0.0165 | 0.0026 | -0.0213 | 0.0117 | 0.0041 | 0.0022 |
| rs889398 | C | 0.0134 | 0.0020 | 0.0268 | 0.0097 | -0.0018 | 0.0017 |
| rs912780 | T | 0.0121 | 0.0021 | 0.0042 | 0.0095 | -0.0042 | 0.0018 |
| rs9435340 | T | 0.0121 | 0.0021 | 0.0111 | 0.0097 | -0.0063 | 0.0018 |
| rs9842947 | C | -0.0125 | 0.0021 | -0.0009 | 0.0099 | 0.0009 | 0.0018 |
| rs986391 | G | 0.0159 | 0.0021 | -0.0010 | 0.0095 | 0.0016 | 0.0018 |
| rs9904288 | T | 0.0122 | 0.0022 | 0.0455 | 0.0103 | -0.0106 | 0.0019 |
| rs9919670 | G | -0.0217 | 0.0020 | 0.0073 | 0.0095 | 0.0074 | 0.0017 |

#### Supplementary table 20: Genetic association estimates for the two-sample regression-based MVMR analysis estimating the effect of smoking on MI, adjusted for education. ea=effect allele; gx=smoking; gy=MI; gz=education; se=standard error

| SNP | ea | gx | gx_se | gy | gy_se | gz | gz_se |
| --- | --- | --- | --- | --- | --- | --- | --- |
| rs10282292 | C | 0.0128 | 0.0021 | 0.0157 | 0.0107 | -0.0010 | 0.0018 |
| rs1050847 | C | 0.0114 | 0.0020 | 0.0304 | 0.0116 | -0.0108 | 0.0017 |
| rs10516044 | G | -0.0199 | 0.0031 | 0.0084 | 0.0162 | 0.0055 | 0.0027 |
| rs10879871 | T | -0.0137 | 0.0021 | 0.0033 | 0.0105 | 0.0084 | 0.0018 |
| rs10918701 | G | 0.0116 | 0.0021 | 0.0226 | 0.0106 | -0.0010 | 0.0018 |
| rs10922907 | A | 0.0145 | 0.0020 | 0.0102 | 0.0105 | -0.0161 | 0.0017 |
| rs111243290 | T | -0.0290 | 0.0053 | -0.0222 | 0.0308 | 0.0190 | 0.0046 |
| rs1118814 | G | 0.0121 | 0.0021 | -0.0029 | 0.0109 | -0.0069 | 0.0018 |
| rs11210229 | A | 0.0167 | 0.0020 | 0.0074 | 0.0104 | -0.0043 | 0.0017 |
| rs112282219 | G | -0.0333 | 0.0050 | -0.0076 | 0.0299 | 0.0017 | 0.0044 |
| rs11255908 | T | -0.0145 | 0.0023 | 0.0125 | 0.0113 | 0.0050 | 0.0020 |
| rs113382419 | C | -0.0409 | 0.0032 | -0.0165 | 0.0208 | -0.0053 | 0.0028 |
| rs1149331 | G | -0.0110 | 0.0020 | -0.0042 | 0.0106 | 0.0075 | 0.0017 |
| rs11768481 | C | 0.0129 | 0.0021 | 0.0055 | 0.0117 | -0.0010 | 0.0018 |
| rs11783093 | C | 0.0227 | 0.0027 | 0.0260 | 0.0151 | -0.0024 | 0.0023 |
| rs11861214 | G | 0.0135 | 0.0024 | -0.0053 | 0.0125 | -0.0114 | 0.0020 |
| rs11948770 | T | -0.0148 | 0.0024 | 0.0013 | 0.0127 | -0.0028 | 0.0020 |
| rs1221148 | C | 0.0133 | 0.0020 | 0.0017 | 0.0106 | 0.0001 | 0.0017 |
| rs12244388 | G | -0.0192 | 0.0021 | 0.0297 | 0.0104 | 0.0065 | 0.0018 |
| rs1246265 | T | -0.0127 | 0.0022 | -0.0203 | 0.0113 | 0.0048 | 0.0019 |
| rs12481282 | G | -0.0129 | 0.0022 | 0.0229 | 0.0115 | 0.0027 | 0.0019 |
| rs12623702 | A | -0.0141 | 0.0021 | 0.0190 | 0.0107 | 0.0099 | 0.0017 |
| rs12708665 | A | -0.0131 | 0.0022 | -0.0186 | 0.0119 | 0.0089 | 0.0019 |
| rs12831617 | C | -0.0131 | 0.0023 | -0.0142 | 0.0127 | 0.0048 | 0.0020 |
| rs12967855 | A | 0.0117 | 0.0021 | -0.0135 | 0.0107 | -0.0194 | 0.0018 |
| rs13009008 | A | 0.0123 | 0.0021 | -0.0051 | 0.0109 | -0.0095 | 0.0018 |
| rs13016665 | C | -0.0123 | 0.0020 | -0.0029 | 0.0105 | 0.0012 | 0.0017 |
| rs13296519 | G | -0.0138 | 0.0020 | -0.0237 | 0.0111 | 0.0078 | 0.0017 |
| rs136233 | A | -0.0141 | 0.0025 | -0.0278 | 0.0134 | 0.0058 | 0.0022 |
| rs147412694 | G | -0.0168 | 0.0028 | -0.0065 | 0.0155 | -0.0008 | 0.0024 |
| rs17309874 | G | -0.0164 | 0.0023 | -0.0193 | 0.0130 | 0.0083 | 0.0019 |
| rs17553262 | A | -0.0182 | 0.0031 | 0.0043 | 0.0186 | 0.0112 | 0.0027 |
| rs17576594 | G | 0.0157 | 0.0022 | 0.0010 | 0.0116 | -0.0113 | 0.0019 |
| rs1827535 | T | -0.0124 | 0.0022 | -0.0141 | 0.0116 | 0.0019 | 0.0018 |
| rs1922018 | C | 0.0145 | 0.0021 | -0.0012 | 0.0112 | -0.0057 | 0.0018 |
| rs1931263 | G | -0.0109 | 0.0020 | -0.0118 | 0.0107 | 0.0120 | 0.0017 |
| rs1933270 | T | 0.0132 | 0.0021 | 0.0018 | 0.0104 | -0.0016 | 0.0018 |
| rs202645 | A | -0.0145 | 0.0025 | 0.0073 | 0.0123 | -0.0001 | 0.0021 |
| rs2062882 | G | -0.0117 | 0.0020 | -0.0053 | 0.0105 | 0.0080 | 0.0017 |
| rs2080870 | A | 0.0125 | 0.0023 | -0.0151 | 0.0123 | -0.0158 | 0.0020 |
| rs215600 | G | 0.0165 | 0.0021 | 0.0190 | 0.0106 | -0.0116 | 0.0018 |
| rs2160896 | C | -0.0120 | 0.0021 | -0.0086 | 0.0110 | 0.0106 | 0.0018 |
| rs2254710 | C | 0.0130 | 0.0023 | 0.0174 | 0.0116 | -0.0031 | 0.0020 |
| rs2401924 | G | 0.0151 | 0.0020 | 0.0078 | 0.0102 | 0.0044 | 0.0017 |
| rs245774 | A | -0.0129 | 0.0022 | -0.0183 | 0.0126 | 0.0070 | 0.0019 |
| rs2675638 | G | 0.0122 | 0.0020 | -0.0136 | 0.0104 | -0.0068 | 0.0017 |
| rs2678670 | A | 0.0126 | 0.0020 | 0.0177 | 0.0109 | -0.0099 | 0.0017 |
| rs2838834 | C | -0.0133 | 0.0022 | -0.0192 | 0.0110 | -0.0009 | 0.0019 |
| rs28485305 | C | 0.0115 | 0.0021 | 0.0102 | 0.0105 | -0.0008 | 0.0018 |
| rs2867112 | T | 0.0211 | 0.0027 | 0.0210 | 0.0138 | -0.0048 | 0.0023 |
| rs2890772 | G | -0.0196 | 0.0020 | -0.0001 | 0.0104 | -0.0036 | 0.0017 |
| rs2894808 | T | -0.0219 | 0.0037 | 0.0116 | 0.0184 | 0.0076 | 0.0031 |
| rs317021 | T | -0.0168 | 0.0026 | -0.0003 | 0.0139 | 0.0057 | 0.0022 |
| rs326341 | G | 0.0137 | 0.0020 | -0.0013 | 0.0104 | -0.0063 | 0.0017 |
| rs329120 | C | 0.0139 | 0.0020 | -0.0101 | 0.0105 | -0.0083 | 0.0017 |
| rs348809 | A | -0.0118 | 0.0021 | 0.0011 | 0.0108 | 0.0050 | 0.0018 |
| rs35120974 | A | -0.0121 | 0.0020 | -0.0005 | 0.0104 | -0.0049 | 0.0017 |
| rs35169606 | T | 0.0126 | 0.0021 | 0.0104 | 0.0123 | -0.0025 | 0.0018 |
| rs35175834 | G | -0.0236 | 0.0024 | 0.0079 | 0.0133 | 0.0129 | 0.0021 |
| rs35343344 | C | 0.0131 | 0.0023 | 0.0118 | 0.0125 | -0.0059 | 0.0020 |
| rs359243 | T | -0.0126 | 0.0021 | -0.0008 | 0.0105 | 0.0064 | 0.0017 |
| rs369230 | G | -0.0130 | 0.0022 | -0.0316 | 0.0140 | 0.0052 | 0.0019 |
| rs3742365 | T | -0.0155 | 0.0020 | -0.0122 | 0.0106 | -0.0059 | 0.0017 |
| rs3769949 | T | -0.0118 | 0.0020 | -0.0083 | 0.0102 | 0.0050 | 0.0017 |
| rs3811038 | T | -0.0137 | 0.0022 | -0.0004 | 0.0118 | 0.0066 | 0.0019 |
| rs3896224 | A | 0.0139 | 0.0020 | 0.0176 | 0.0103 | -0.0088 | 0.0017 |
| rs421983 | T | 0.0125 | 0.0020 | -0.0070 | 0.0102 | -0.0033 | 0.0017 |
| rs4391802 | A | 0.0149 | 0.0022 | 0.0120 | 0.0113 | -0.0072 | 0.0019 |
| rs4473348 | A | -0.0149 | 0.0023 | -0.0262 | 0.0119 | -0.0026 | 0.0020 |
| rs4543592 | T | -0.0124 | 0.0020 | -0.0039 | 0.0107 | 0.0067 | 0.0017 |
| rs4571506 | C | 0.0113 | 0.0020 | -0.0005 | 0.0103 | -0.0115 | 0.0017 |
| rs4671357 | T | -0.0136 | 0.0020 | -0.0132 | 0.0102 | 0.0091 | 0.0017 |
| rs4814873 | C | 0.0140 | 0.0024 | 0.0240 | 0.0121 | -0.0064 | 0.0020 |
| rs4949465 | T | -0.0168 | 0.0030 | -0.0272 | 0.0135 | 0.0129 | 0.0025 |
| rs4957528 | A | -0.0145 | 0.0025 | -0.0081 | 0.0140 | 0.0051 | 0.0021 |
| rs549845 | G | 0.0161 | 0.0022 | 0.0202 | 0.0109 | -0.0177 | 0.0019 |
| rs6011779 | C | 0.0276 | 0.0025 | 0.0175 | 0.0153 | -0.0042 | 0.0022 |
| rs60952428 | T | 0.0192 | 0.0035 | 0.0162 | 0.0155 | -0.0079 | 0.0031 |
| rs6119897 | G | -0.0185 | 0.0023 | -0.0160 | 0.0118 | 0.0069 | 0.0020 |
| rs62098013 | G | -0.0122 | 0.0021 | -0.0069 | 0.0117 | 0.0126 | 0.0018 |
| rs62135536 | C | 0.0352 | 0.0057 | -0.0006 | 0.0328 | -0.0092 | 0.0049 |
| rs62155874 | A | -0.0244 | 0.0030 | 0.0166 | 0.0166 | 0.0142 | 0.0026 |
| rs62175972 | T | 0.0313 | 0.0055 | 0.0103 | 0.0318 | -0.0300 | 0.0048 |
| rs624833 | T | 0.0132 | 0.0022 | 0.0199 | 0.0118 | -0.0132 | 0.0018 |
| rs6598539 | T | -0.0118 | 0.0020 | -0.0190 | 0.0102 | 0.0065 | 0.0017 |
| rs6741228 | T | 0.0114 | 0.0020 | 0.0172 | 0.0104 | -0.0075 | 0.0017 |
| rs67596067 | G | -0.0128 | 0.0021 | -0.0078 | 0.0110 | 0.0036 | 0.0018 |
| rs6778080 | T | 0.0158 | 0.0023 | 0.0095 | 0.0128 | -0.0139 | 0.0019 |
| rs6779302 | G | -0.0125 | 0.0021 | 0.0024 | 0.0108 | 0.0113 | 0.0018 |
| rs6935954 | A | 0.0137 | 0.0020 | -0.0173 | 0.0108 | -0.0138 | 0.0017 |
| rs6957896 | C | -0.0109 | 0.0020 | -0.0125 | 0.0108 | 0.0016 | 0.0017 |
| rs6962772 | A | 0.0159 | 0.0028 | 0.0085 | 0.0130 | -0.0040 | 0.0023 |
| rs7039819 | G | 0.0126 | 0.0020 | 0.0175 | 0.0105 | -0.0106 | 0.0017 |
| rs7077678 | C | 0.0123 | 0.0021 | 0.0197 | 0.0103 | 0.0016 | 0.0017 |
| rs71367545 | G | -0.0148 | 0.0025 | -0.0138 | 0.0134 | 0.0125 | 0.0021 |
| rs7144406 | A | -0.0135 | 0.0025 | -0.0056 | 0.0143 | 0.0051 | 0.0021 |
| rs7155595 | A | -0.0126 | 0.0021 | -0.0131 | 0.0113 | 0.0073 | 0.0018 |
| rs72678864 | G | 0.0178 | 0.0026 | 0.0079 | 0.0153 | -0.0123 | 0.0023 |
| rs7297175 | T | -0.0115 | 0.0020 | -0.0040 | 0.0110 | 0.0156 | 0.0017 |
| rs732083 | G | 0.0120 | 0.0021 | 0.0187 | 0.0109 | -0.0081 | 0.0018 |
| rs73220544 | A | -0.0156 | 0.0028 | -0.0305 | 0.0149 | 0.0069 | 0.0024 |
| rs7333559 | G | 0.0152 | 0.0025 | 0.0103 | 0.0133 | -0.0060 | 0.0021 |
| rs74086911 | G | 0.0213 | 0.0038 | -0.0037 | 0.0211 | -0.0184 | 0.0034 |
| rs7528604 | G | 0.0138 | 0.0020 | 0.0069 | 0.0103 | -0.0059 | 0.0017 |
| rs7553348 | G | 0.0138 | 0.0020 | -0.0023 | 0.0105 | -0.0004 | 0.0017 |
| rs7569203 | A | -0.0156 | 0.0022 | -0.0012 | 0.0109 | 0.0026 | 0.0018 |
| rs75742406 | G | 0.0139 | 0.0023 | 0.0096 | 0.0117 | -0.0010 | 0.0019 |
| rs76608582 | C | 0.0311 | 0.0049 | -0.0376 | 0.0421 | -0.0280 | 0.0045 |
| rs775758 | A | 0.0115 | 0.0020 | 0.0138 | 0.0104 | -0.0030 | 0.0017 |
| rs7766610 | C | 0.0181 | 0.0026 | 0.0147 | 0.0124 | 0.0008 | 0.0022 |
| rs7775552 | A | -0.0119 | 0.0020 | 0.0032 | 0.0102 | 0.0020 | 0.0017 |
| rs7807019 | A | -0.0150 | 0.0020 | -0.0210 | 0.0102 | 0.0106 | 0.0017 |
| rs7897434 | G | 0.0116 | 0.0021 | -0.0050 | 0.0114 | -0.0038 | 0.0018 |
| rs8042134 | T | -0.0141 | 0.0020 | 0.0169 | 0.0109 | 0.0079 | 0.0017 |
| rs8042849 | C | 0.0276 | 0.0021 | 0.0013 | 0.0112 | 0.0002 | 0.0018 |
| rs860326 | C | 0.0119 | 0.0020 | -0.0004 | 0.0107 | -0.0088 | 0.0017 |
| rs8614 | C | -0.0165 | 0.0026 | -0.0182 | 0.0130 | 0.0041 | 0.0022 |
| rs889398 | C | 0.0134 | 0.0020 | 0.0183 | 0.0109 | -0.0018 | 0.0017 |
| rs912780 | T | 0.0121 | 0.0021 | -0.0038 | 0.0106 | -0.0042 | 0.0018 |
| rs9435340 | T | 0.0121 | 0.0021 | 0.0137 | 0.0107 | -0.0063 | 0.0018 |
| rs9842947 | C | -0.0125 | 0.0021 | 0.0011 | 0.0110 | 0.0009 | 0.0018 |
| rs986391 | G | 0.0159 | 0.0021 | 0.0036 | 0.0105 | 0.0016 | 0.0018 |
| rs9904288 | T | 0.0122 | 0.0022 | 0.0441 | 0.0114 | -0.0106 | 0.0019 |
| rs9919670 | G | -0.0217 | 0.0020 | 0.0066 | 0.0104 | 0.0074 | 0.0017 |

#### Supplementary table 21: Genetic association estimates for the two-sample regression-based MVMR analysis estimating the effect of smoking on stroke, adjusted for education. ea=effect allele; gx=smoking; gy=stroke; gz=education; se=standard error

| SNP | ea | gx | gx_se | gy | gy_se | gz | gz_se |
| --- | --- | --- | --- | --- | --- | --- | --- |
| rs10282292 | C | 0.0128 | 0.0021 | 0.0066 | 0.0080 | -0.0010 | 0.0018 |
| rs1050847 | C | 0.0114 | 0.0020 | 0.0083 | 0.0091 | -0.0108 | 0.0017 |
| rs10516044 | G | -0.0199 | 0.0031 | -0.0148 | 0.0133 | 0.0055 | 0.0027 |
| rs10879871 | T | -0.0137 | 0.0021 | -0.0047 | 0.0082 | 0.0084 | 0.0018 |
| rs10918701 | G | 0.0116 | 0.0021 | 0.0012 | 0.0082 | -0.0010 | 0.0018 |
| rs10922907 | A | 0.0145 | 0.0020 | 0.0057 | 0.0078 | -0.0161 | 0.0017 |
| rs111243290 | T | -0.0290 | 0.0053 | -0.0367 | 0.0222 | 0.0190 | 0.0046 |
| rs1118814 | G | 0.0121 | 0.0021 | 0.0120 | 0.0083 | -0.0069 | 0.0018 |
| rs11210229 | A | 0.0167 | 0.0020 | 0.0040 | 0.0080 | -0.0043 | 0.0017 |
| rs112282219 | G | -0.0333 | 0.0050 | -0.0288 | 0.0236 | 0.0017 | 0.0044 |
| rs11255908 | T | -0.0145 | 0.0023 | 0.0025 | 0.0090 | 0.0050 | 0.0020 |
| rs113382419 | C | -0.0409 | 0.0032 | -0.0028 | 0.0157 | -0.0053 | 0.0028 |
| rs1149331 | G | -0.0110 | 0.0020 | -0.0050 | 0.0084 | 0.0075 | 0.0017 |
| rs11768481 | C | 0.0129 | 0.0021 | 0.0008 | 0.0099 | -0.0010 | 0.0018 |
| rs11783093 | C | 0.0227 | 0.0027 | 0.0017 | 0.0123 | -0.0024 | 0.0023 |
| rs11861214 | G | 0.0135 | 0.0024 | -0.0093 | 0.0093 | -0.0114 | 0.0020 |
| rs11948770 | T | -0.0148 | 0.0024 | 0.0004 | 0.0095 | -0.0028 | 0.0020 |
| rs1221148 | C | 0.0133 | 0.0020 | 0.0010 | 0.0078 | 0.0001 | 0.0017 |
| rs12244388 | G | -0.0192 | 0.0021 | -0.0151 | 0.0079 | 0.0065 | 0.0018 |
| rs1246265 | T | -0.0127 | 0.0022 | -0.0227 | 0.0087 | 0.0048 | 0.0019 |
| rs12481282 | G | -0.0129 | 0.0022 | 0.0061 | 0.0087 | 0.0027 | 0.0019 |
| rs12623702 | A | -0.0141 | 0.0021 | 0.0059 | 0.0082 | 0.0099 | 0.0017 |
| rs12708665 | A | -0.0131 | 0.0022 | -0.0058 | 0.0093 | 0.0089 | 0.0019 |
| rs12831617 | C | -0.0131 | 0.0023 | -0.0094 | 0.0108 | 0.0048 | 0.0020 |
| rs12967855 | A | 0.0117 | 0.0021 | -0.0056 | 0.0085 | -0.0194 | 0.0018 |
| rs13009008 | A | 0.0123 | 0.0021 | 0.0215 | 0.0083 | -0.0095 | 0.0018 |
| rs13016665 | C | -0.0123 | 0.0020 | -0.0108 | 0.0080 | 0.0012 | 0.0017 |
| rs13296519 | G | -0.0138 | 0.0020 | 0.0084 | 0.0086 | 0.0078 | 0.0017 |
| rs136233 | A | -0.0141 | 0.0025 | -0.0009 | 0.0104 | 0.0058 | 0.0022 |
| rs147412694 | G | -0.0168 | 0.0028 | -0.0203 | 0.0127 | -0.0008 | 0.0024 |
| rs17309874 | G | -0.0164 | 0.0023 | -0.0090 | 0.0102 | 0.0083 | 0.0019 |
| rs17553262 | A | -0.0182 | 0.0031 | 0.0083 | 0.0132 | 0.0112 | 0.0027 |
| rs17576594 | G | 0.0157 | 0.0022 | 0.0043 | 0.0087 | -0.0113 | 0.0019 |
| rs1827535 | T | -0.0124 | 0.0022 | -0.0154 | 0.0092 | 0.0019 | 0.0018 |
| rs1922018 | C | 0.0145 | 0.0021 | -0.0109 | 0.0084 | -0.0057 | 0.0018 |
| rs1931263 | G | -0.0109 | 0.0020 | 0.0091 | 0.0085 | 0.0120 | 0.0017 |
| rs1933270 | T | 0.0132 | 0.0021 | 0.0001 | 0.0081 | -0.0016 | 0.0018 |
| rs202645 | A | -0.0145 | 0.0025 | 0.0008 | 0.0108 | -0.0001 | 0.0021 |
| rs2062882 | G | -0.0117 | 0.0020 | 0.0130 | 0.0087 | 0.0080 | 0.0017 |
| rs2080870 | A | 0.0125 | 0.0023 | 0.0061 | 0.0099 | -0.0158 | 0.0020 |
| rs215600 | G | 0.0165 | 0.0021 | 0.0095 | 0.0082 | -0.0116 | 0.0018 |
| rs2160896 | C | -0.0120 | 0.0021 | 0.0061 | 0.0084 | 0.0106 | 0.0018 |
| rs2254710 | C | 0.0130 | 0.0023 | 0.0150 | 0.0091 | -0.0031 | 0.0020 |
| rs2401924 | G | 0.0151 | 0.0020 | 0.0077 | 0.0079 | 0.0044 | 0.0017 |
| rs245774 | A | -0.0129 | 0.0022 | -0.0169 | 0.0093 | 0.0070 | 0.0019 |
| rs2675638 | G | 0.0122 | 0.0020 | 0.0182 | 0.0080 | -0.0068 | 0.0017 |
| rs2678670 | A | 0.0126 | 0.0020 | 0.0100 | 0.0088 | -0.0099 | 0.0017 |
| rs2838834 | C | -0.0133 | 0.0022 | 0.0050 | 0.0086 | -0.0009 | 0.0019 |
| rs28485305 | C | 0.0115 | 0.0021 | 0.0155 | 0.0080 | -0.0008 | 0.0018 |
| rs2867112 | T | 0.0211 | 0.0027 | 0.0031 | 0.0108 | -0.0048 | 0.0023 |
| rs2890772 | G | -0.0196 | 0.0020 | -0.0040 | 0.0081 | -0.0036 | 0.0017 |
| rs2894808 | T | -0.0219 | 0.0037 | 0.0112 | 0.0148 | 0.0076 | 0.0031 |
| rs317021 | T | -0.0168 | 0.0026 | 0.0075 | 0.0121 | 0.0057 | 0.0022 |
| rs326341 | G | 0.0137 | 0.0020 | 0.0076 | 0.0078 | -0.0063 | 0.0017 |
| rs329120 | C | 0.0139 | 0.0020 | 0.0054 | 0.0078 | -0.0083 | 0.0017 |
| rs348809 | A | -0.0118 | 0.0021 | 0.0062 | 0.0084 | 0.0050 | 0.0018 |
| rs35120974 | A | -0.0121 | 0.0020 | -0.0146 | 0.0080 | -0.0049 | 0.0017 |
| rs35169606 | T | 0.0126 | 0.0021 | 0.0102 | 0.0096 | -0.0025 | 0.0018 |
| rs35175834 | G | -0.0236 | 0.0024 | 0.0060 | 0.0103 | 0.0129 | 0.0021 |
| rs35343344 | C | 0.0131 | 0.0023 | 0.0219 | 0.0095 | -0.0059 | 0.0020 |
| rs359243 | T | -0.0126 | 0.0021 | 0.0015 | 0.0087 | 0.0064 | 0.0017 |
| rs369230 | G | -0.0130 | 0.0022 | -0.0073 | 0.0093 | 0.0052 | 0.0019 |
| rs3742365 | T | -0.0155 | 0.0020 | 0.0004 | 0.0080 | -0.0059 | 0.0017 |
| rs3769949 | T | -0.0118 | 0.0020 | -0.0131 | 0.0080 | 0.0050 | 0.0017 |
| rs3811038 | T | -0.0137 | 0.0022 | 0.0065 | 0.0091 | 0.0066 | 0.0019 |
| rs3896224 | A | 0.0139 | 0.0020 | -0.0038 | 0.0080 | -0.0088 | 0.0017 |
| rs421983 | T | 0.0125 | 0.0020 | 0.0063 | 0.0079 | -0.0033 | 0.0017 |
| rs4391802 | A | 0.0149 | 0.0022 | 0.0073 | 0.0087 | -0.0072 | 0.0019 |
| rs4473348 | A | -0.0149 | 0.0023 | -0.0096 | 0.0091 | -0.0026 | 0.0020 |
| rs4543592 | T | -0.0124 | 0.0020 | -0.0003 | 0.0082 | 0.0067 | 0.0017 |
| rs4571506 | C | 0.0113 | 0.0020 | 0.0165 | 0.0079 | -0.0115 | 0.0017 |
| rs4671357 | T | -0.0136 | 0.0020 | -0.0041 | 0.0079 | 0.0091 | 0.0017 |
| rs4814873 | C | 0.0140 | 0.0024 | 0.0105 | 0.0093 | -0.0064 | 0.0020 |
| rs4949465 | T | -0.0168 | 0.0030 | 0.0086 | 0.0109 | 0.0129 | 0.0025 |
| rs4957528 | A | -0.0145 | 0.0025 | -0.0022 | 0.0104 | 0.0051 | 0.0021 |
| rs549845 | G | 0.0161 | 0.0022 | 0.0124 | 0.0084 | -0.0177 | 0.0019 |
| rs6011779 | C | 0.0276 | 0.0025 | 0.0000 | 0.0111 | -0.0042 | 0.0022 |
| rs60952428 | T | 0.0192 | 0.0035 | -0.0149 | 0.0120 | -0.0079 | 0.0031 |
| rs6119897 | G | -0.0185 | 0.0023 | 0.0103 | 0.0092 | 0.0069 | 0.0020 |
| rs62098013 | G | -0.0122 | 0.0021 | -0.0089 | 0.0092 | 0.0126 | 0.0018 |
| rs62135536 | C | 0.0352 | 0.0057 | -0.0021 | 0.0283 | -0.0092 | 0.0049 |
| rs62155874 | A | -0.0244 | 0.0030 | 0.0020 | 0.0130 | 0.0142 | 0.0026 |
| rs62175972 | T | 0.0313 | 0.0055 | 0.0276 | 0.0280 | -0.0300 | 0.0048 |
| rs624833 | T | 0.0132 | 0.0022 | 0.0029 | 0.0093 | -0.0132 | 0.0018 |
| rs6598539 | T | -0.0118 | 0.0020 | -0.0046 | 0.0080 | 0.0065 | 0.0017 |
| rs6741228 | T | 0.0114 | 0.0020 | 0.0106 | 0.0080 | -0.0075 | 0.0017 |
| rs67596067 | G | -0.0128 | 0.0021 | -0.0035 | 0.0086 | 0.0036 | 0.0018 |
| rs6778080 | T | 0.0158 | 0.0023 | -0.0119 | 0.0102 | -0.0139 | 0.0019 |
| rs6779302 | G | -0.0125 | 0.0021 | 0.0035 | 0.0088 | 0.0113 | 0.0018 |
| rs6935954 | A | 0.0137 | 0.0020 | -0.0043 | 0.0088 | -0.0138 | 0.0017 |
| rs6957896 | C | -0.0109 | 0.0020 | -0.0075 | 0.0084 | 0.0016 | 0.0017 |
| rs6962772 | A | 0.0159 | 0.0028 | 0.0117 | 0.0104 | -0.0040 | 0.0023 |
| rs7039819 | G | 0.0126 | 0.0020 | 0.0091 | 0.0086 | -0.0106 | 0.0017 |
| rs7077678 | C | 0.0123 | 0.0021 | -0.0048 | 0.0080 | 0.0016 | 0.0017 |
| rs71367545 | G | -0.0148 | 0.0025 | -0.0090 | 0.0113 | 0.0125 | 0.0021 |
| rs7144406 | A | -0.0135 | 0.0025 | -0.0022 | 0.0112 | 0.0051 | 0.0021 |
| rs7155595 | A | -0.0126 | 0.0021 | -0.0117 | 0.0093 | 0.0073 | 0.0018 |
| rs72678864 | G | 0.0178 | 0.0026 | -0.0055 | 0.0127 | -0.0123 | 0.0023 |
| rs7297175 | T | -0.0115 | 0.0020 | -0.0186 | 0.0083 | 0.0156 | 0.0017 |
| rs732083 | G | 0.0120 | 0.0021 | 0.0131 | 0.0083 | -0.0081 | 0.0018 |
| rs73220544 | A | -0.0156 | 0.0028 | -0.0090 | 0.0113 | 0.0069 | 0.0024 |
| rs7333559 | G | 0.0152 | 0.0025 | 0.0014 | 0.0097 | -0.0060 | 0.0021 |
| rs74086911 | G | 0.0213 | 0.0038 | 0.0027 | 0.0168 | -0.0184 | 0.0034 |
| rs7528604 | G | 0.0138 | 0.0020 | 0.0202 | 0.0079 | -0.0059 | 0.0017 |
| rs7553348 | G | 0.0138 | 0.0020 | -0.0042 | 0.0081 | -0.0004 | 0.0017 |
| rs7569203 | A | -0.0156 | 0.0022 | 0.0174 | 0.0085 | 0.0026 | 0.0018 |
| rs75742406 | G | 0.0139 | 0.0023 | -0.0236 | 0.0091 | -0.0010 | 0.0019 |
| rs775758 | A | 0.0115 | 0.0020 | -0.0066 | 0.0080 | -0.0030 | 0.0017 |
| rs7766610 | C | 0.0181 | 0.0026 | -0.0033 | 0.0093 | 0.0008 | 0.0022 |
| rs7775552 | A | -0.0119 | 0.0020 | 0.0055 | 0.0087 | 0.0020 | 0.0017 |
| rs7807019 | A | -0.0150 | 0.0020 | -0.0091 | 0.0078 | 0.0106 | 0.0017 |
| rs7897434 | G | 0.0116 | 0.0021 | -0.0048 | 0.0086 | -0.0038 | 0.0018 |
| rs8042134 | T | -0.0141 | 0.0020 | 0.0056 | 0.0082 | 0.0079 | 0.0017 |
| rs8042849 | C | 0.0276 | 0.0021 | 0.0086 | 0.0086 | 0.0002 | 0.0018 |
| rs860326 | C | 0.0119 | 0.0020 | 0.0008 | 0.0081 | -0.0088 | 0.0017 |
| rs8614 | C | -0.0165 | 0.0026 | -0.0133 | 0.0098 | 0.0041 | 0.0022 |
| rs889398 | C | 0.0134 | 0.0020 | 0.0115 | 0.0083 | -0.0018 | 0.0017 |
| rs912780 | T | 0.0121 | 0.0021 | 0.0063 | 0.0081 | -0.0042 | 0.0018 |
| rs9435340 | T | 0.0121 | 0.0021 | 0.0002 | 0.0082 | -0.0063 | 0.0018 |
| rs9842947 | C | -0.0125 | 0.0021 | 0.0015 | 0.0088 | 0.0009 | 0.0018 |
| rs986391 | G | 0.0159 | 0.0021 | 0.0026 | 0.0085 | 0.0016 | 0.0018 |
| rs9904288 | T | 0.0122 | 0.0022 | 0.0147 | 0.0088 | -0.0106 | 0.0019 |
| rs9919670 | G | -0.0217 | 0.0020 | -0.0151 | 0.0085 | 0.0074 | 0.0017 |

#### Supplementary table 22: One-sample MR sensitivity analyses. The risk difference scale was used for the weighted median, and the log odds ratio scale was used for all other measures.

| **Education-CVD** | **Estimate** | **95% LCI** | **95% UCI** | **P** |
| --- | --- | --- | --- | --- |
| IVW | -0.594 | -0.873 | -0.316 | 0.000 |
| MR-Egger | -0.647 | -0.946 | -0.349 | 0.000 |
| MR-Egger intercept |  |  |  | 0.330 |
| Weighted median | -0.027 | -0.049 | -0.006 | 0.011 |
| **Education-CHD** |  |  |  |  |
| IVW | -0.722 | -1.143 | -0.302 | 0.001 |
| MR-Egger | -0.799 | -1.249 | -0.349 | 0.001 |
| MR-Egger intercept |  |  |  | 0.346 |
| Weighted median | -0.020 | -0.032 | -0.009 | 0.000 |
| **Education-MI** |  |  |  |  |
| IVW | -0.802 | -1.249 | -0.354 | 0.000 |
| MR-Egger | -0.858 | -1.339 | -0.376 | 0.000 |
| MR-Egger intercept |  |  |  | 0.527 |
| Weighted median | -0.008 | -0.014 | -0.002 | 0.014 |
| **Education-Stroke** |  |  |  |  |
| IVW | -1.119 | -1.529 | -0.709 | 0.000 |
| MR-Egger | -1.246 | -1.685 | -0.807 | 0.000 |
| MR-Egger intercept |  |  |  | 0.132 |
| Weighted median | -0.006 | -0.013 | 0.000 | 0.056 |
| **Education_BMI** |  |  |  |  |
| IVW | -0.080 | -0.143 | -0.016 | 0.014 |
| MR-Egger | -0.068 | -0.136 | 0.000 | 0.053 |
| MR-Egger intercept |  |  |  | 0.351 |
| Weighted median | -0.455 | -0.554 | -0.357 | 0.000 |
| **Education-SBP** |  |  |  |  |
| IVW | -0.472 | -0.665 | -0.278 | 0.000 |
| MR-Egger | -0.516 | -0.723 | -0.308 | 0.000 |
| MR-Egger intercept |  |  |  | 0.268 |
| Weighted median | -0.158 | -0.237 | -0.080 | 0.000 |
| **Education-Smoking** |  |  |  |  |
| IVW | -0.021 | -0.075 | 0.009 | 0.442 |
| MR-Egger | -0.022 | -0.081 | 0.010 | 0.463 |
| MR-Egger intercept |  |  |  | 0.950 |
| Weighted median | -0.319 | -0.401 | -0.238 | 0.000 |

#### Supplementary table 23: Two-sample MR sensitivity analyses. CHD, MI and stroke are in log OR units, while BMI, SBP and smoking are in SD units.

| Education-CHD | Estimate | 95% lower CI | 95% upper CI | P value |
| --- | --- | --- | --- | --- |
| IVW | -0.460 | -0.516 | -0.404 | 0.000 |
| MR-Egger | -0.383 | -0.609 | -0.157 | 0.001 |
| Intercept |  |  |  | 0.370 |
| Weighted median | -0.481 | -0.563 | -0.399 | 0.000 |
| Education-MI |  |  |  |  |
| IVW | -0.489 | -0.551 | -0.427 | 0.000 |
| MR-Egger | -0.407 | -0.654 | -0.160 | 0.001 |
| Intercept |  |  |  | 0.384 |
| Weighted median | -0.522 | -0.616 | -0.429 | 0.000 |
| Education-Stroke |  |  |  |  |
| IVW | -0.337 | -0.385 | -0.289 | 0.000 |
| MR-Egger | -0.325 | -0.514 | -0.135 | 0.001 |
| Intercept |  |  |  | 0.757 |
| Weighted median | -0.343 | -0.413 | -0.272 | 0.000 |
| Education-BMI |  |  |  |  |
| IVW | -0.221 | -0.239 | -0.202 | 0.000 |
| MR-Egger | -0.282 | -0.493 | -0.072 | 0.009 |
| Intercept |  |  |  | 0.989 |
| Weighted median | -0.267 | -0.301 | -0.232 | 0.000 |
| Education-SBP |  |  |  |  |
| IVW | -2.861 | -3.124 | -2.599 | 0.000 |
| MR-Egger | -2.406 | -3.927 | -0.886 | 0.002 |
| Intercept |  |  |  | 0.325 |
| Weighted median | -3.386 | -3.822 | -2.951 | 0.000 |
| Education-Smoking |  |  |  |  |
| IVW | -0.320 | -0.332 | -0.307 | 0.000 |
| MR-Egger | -0.290 | -0.359 | -0.220 | 0.000 |
| Intercept |  |  |  | 0.057 |
| Weighted median | -0.348 | -0.369 | -0.327 | 0.000 |

#### Supplementary table 24: Unadjusted estimates for the proportion mediated in individual level UK Biobank analyses estimated from the log OR scale

| **Outcome** | **Method** | **Mediator** | **Proportion Mediated (%) (95% CI)** |
| --- | --- | --- | --- |
| **CVD** | Multivariable observational | BMI | 15.84 (14.96, 16.72) |
|  |  | SBP | 19.52 (16.70, 22.34) |
|  |  | Smoking | 16.19 (14.76, 17.63) |
|  | One-sample MR | BMI | 10.01 (-54.48, 74.68) |
|  |  | SBP | 9.79 (-4.54, 24.12) |
|  |  | Smoking | 22.12 (-47.78, 92.02) |
| **Stroke** | Multivariable observational | BMI | 8.73 (4.23, 13.22) |
|  |  | SBP | 22.23 (16.28, 28.19) |
|  |  | Smoking | 13.39 (11.32, 15.45) |
|  | One-sample MR | BMI | 12.85 (-274.10, 299.80) |
|  |  | SBP | 12.44 (-1057.89, 1082.77) |
|  |  | Smoking | 26.41 (5.39, 47.44) |
| **MI** | Multivariable observational | BMI | 12.02 (7.08, 16.96) |
|  |  | SBP | 22.77 (18.54, 26.99) |
|  |  | Smoking | 21.84 (15.85, 27.83) |
|  | One-sample MR | BMI | -5.63 (-30.70, 19.43) |
|  |  | SBP | 8.37 (1.72, 15.01) |
|  |  | Smoking | 13.88 (-4.38, 32.15) |
| **CHD** | Multivariable observational | BMI | 14.40 (11.88, 16.93) |
|  |  | SBP | 21.97 (17.04, 26.89) |
|  |  | Smoking | 16.64 (13.85, 19.43) |
|  | One-sample MR | BMI | -5.49 (-12.57, 1.59) |
|  |  | SBP | 11.40 (3.81, 18.99) |
|  |  | Smoking | -0.68 (-56.92, 55.55) |

#### Supplementary table 25: Age and sex adjusted estimates for mediation by BMI, SBP and smoking on the association of education to CVD in individual level UK Biobank analyses

| **Outcome** | **Method** | **Mediator** | **Proportion Mediated (%)** |
| --- | --- | --- | --- |
| **CVD** | Multivariable observational | BMI | 20.00 (15.16, 24.84) |
|  |  | SBP | 8.55 (6.68, 10.41) |
|  |  | Smoking | 20.15 (18.04, 22.26) |
|  | One-sample MR | BMI | 9.52 (-10.51, 29.56) |
|  |  | SBP | 11.80 (0.48, 23.11) |
|  |  | Smoking | 23.72 (-30.28, 77.72) |
| **Stroke** | Multivariable observational | BMI | 12.56 (2.48, 22.64) |
|  |  | SBP | 15.92 (6.24, 23.80) |
|  |  | Smoking | 20.16 (13.66, 26.66) |
|  | One-sample MR | BMI | 11.65 (-28.16, 51.47) |
|  |  | SBP | 14.47 (-3.88, 32.81) |
|  |  | Smoking | 25.95 (-6.52, 58.42) |
| **MI** | Multivariable observational | BMI | 11.80 (8.36, 15.24) |
|  |  | SBP | 10.97 (7.28, 14.66) |
|  |  | Smoking | 21.00 (14.75, 27.26) |
|  | One-sample MR | BMI | -5.34 (-13.31, 2.64) |
|  |  | SBP | 10.77 (9.85, 11.68) |
|  |  | Smoking | 16.24 (-1.87, 34.35) |
| **CHD** | Multivariable observational | BMI | 17.60 (12.35, 22.84) |
|  |  | SBP | 10.69 (9.51, 11.88) |
|  |  | Smoking | 18.61 (15.21, 22.01) |
|  | One-sample MR | BMI | -4.78 (-16.26, 6.70) |
|  |  | SBP | 14.22 (0.71, 27.73) |
|  |  | Smoking | 4.24 (-28.15, 36.63) |
